# A simple mass-action model predicts genome-wide protein timecourses from mRNA trajectories during a dynamic response in two strains of *Saccharomyces cerevisiae*

**DOI:** 10.1101/805846

**Authors:** Sidney Kuo, Jarrett D. Egertson, Gennifer E. Merrihew, Michael J. MacCoss, Daniel A. Pollard, Scott A. Rifkin

## Abstract

Although mRNA is a necessary precursor to protein, several studies have argued that the relationship between mRNA and protein levels is often weak. This claim undermines the functional relevance of conclusions based on quantitative analyses of mRNA levels, which are ubiquitous in modern biology from the single gene to the whole genome scale. Furthermore, if post-translational processes vary between strains and species, then comparative studies based on mRNA alone would miss an important driver of diversity. However, gene expression is dynamic, and most studies examining relationship between mRNA and protein levels at the genome scale have analyzed single timepoints. We measure yeast gene expression after pheromone exposure and show that, for most genes, protein timecourses can be predicted from mRNA timecourses through a simple, gene-specific, generative model. By comparing model parameters and predictions between strains, we find that while mRNA variation often leads to protein differences, evolution also manipulates protein-specific processes to amplify or buffer transcriptional regulation.

## Introduction

Nearly two decades of genome-scale analyses of gene expression variation have resulted in a broad characterization of the location and activities of genetic variants affecting mRNA abundances. Expression quantitative trait locus (eQTL) mapping experiments have revealed that mRNA abundance variation results from a large number of gene-specific genetic variants located within the locus of the varying gene, as well as a smaller number of genetic variants that act on many genes from distant locations in the genome (Yvert et al. 2003; Doss et al. 2005; Gibson and Weir 2005; Andrie et al. 2014; Pai et al. 2015; Gilad et al. 2008). Allele-specific expression measurements in F1 hybrids confirmed that variation in mRNA abundances is the result of both *cis*-acting (allele-specific) and *trans*-acting (non-allele-specific) genetic variation (Wittkopp et al. 2004). Integrating causal studies of mRNA variation into analyses of higher phenotypic variation promised to be a way to mechanistically explain how polymorphisms lead to phenotypic differences (Schadt et al. 2005).

However, while the genetics of mRNA abundance variation has advanced, its relevance to higher-level phenotypic variation is still unclear. Quantitative measurements of protein levels have shown that steady state mRNA abundances are not necessarily good predictors of relative protein expression across genes (Taniguchi et al. 2010; Lackner et al. 2012; Vogel and Marcotte 2012; Edfors et al. 2016; Fortelny et al. 2017), highlighting the role that gene-specific translation rates and protein stabilities can play in protein expression. This disconnect has been found not just across genes, but across individuals and strains as well. Although the reliability and sensitivity of quantitative proteomic technologies still lag behind their mRNA counterparts, initial proteomic surveys of natural variation in steady state protein levels have found that mRNA variation is not well correlated with protein variation (Minagawa et al. 2008; Khan et al. 2013; Chick et al. 2016; Consoli et al. 2002; Cenik et al. 2015; Battle et al. 2015). A mass-spectroscopy study of genetic variation in yeast underlying steady state protein levels for highly expressed genes found that a minority of variants were linked to mRNA variation (Foss et al. 2011). Other studies of the same yeast strains using fluorescent protein levels instead of mass-spectroscopy found that around half of genetic variation affecting protein levels also affects mRNA levels (Parts et al. 2014; Albert et al. 2014a), suggesting that this disconnect between mRNA and protein variation is not simply the result of technical limitations of mass-spectrometry. These studies imply that a specific class of genetic variants acts directly on protein levels to impact phenotypic variation. If variation in mRNA levels is only weakly associated with variation in protein levels, then mRNA differences would have limited relevance to downstream phenotypes (Yang et al. 2017), despite the fact that most genome-scale, comparative gene expression studies have been performed at the mRNA level (Brem et al. 2002; Rockman and Kruglyak 2006).

Genetic variants could affect protein expression through translation (initiation, elongation, and termination), protein folding, stability, and degradation. These effects could act in *cis* through variants within the locus of the gene or could act in *trans* through auto-regulation or the action of other genes. For translation rates, *trans*-acting polymorphisms have been found to be more common than *cis*-acting ones in yeast (Torabi and Kruglyak 2011; McManus et al. 2014). Genetic variants could also act at both the mRNA and protein levels, reinforcing or compensating for each other (Khan et al. 2013; McManus et al. 2014; Artieri and Fraser 2014; Albert et al. 2014b).

Many of the studies that document a disconnect between mRNA and protein levels across genes or across strains/individuals are based on snapshot measurements of cells (Zheng et al. 2010; Roberts et al. 2000) or organisms *at steady state*. As a consequence, their conclusions about the relationship between mRNA and protein may not be generalizable to common, dynamic situations such as development, disease progression, or response to environmental fluctuations. Moreover, timecourse data can reveal regulatory connections within cells that are not readily apparent even from analyses of large knockout panels (Gonçalves et al. 2017).

## Results

To determine (a) whether mRNA levels are a good proxy for protein levels during the dynamic response of an organism to an environmental stimulus (Liu et al. 2016) and (b) whether differences in mRNA levels between strains translate into differences in protein levels, we profiled the transcriptomes and proteomes of two strains of *Saccharomyces cerevisiae* during their response to pheromone. mRNA levels in strains S288c and YJM145 have previously been shown to differ at 30 minutes after exposure to alpha factor pheromone (Zheng et al. 2010) and ribosome loading and transcriptional differences have been compared between these strains at steady state (Sun et al. 2016). We collected mRNA samples at 0, 0. 5, 1, 1.6, 2.4, 3.2, 4.8, 6.4, and 8 hours after exposure and protein samples at 0, 1.6, 3.2, 4.8, 6.4, and 8 hours for each strain in three biological replicates with mRNA and protein collected from the same flasks. We measured genome-wide levels of mRNA by RNA-seq and protein levels by quantitative mass spectrometry (Chapman et al. 2014).

Even during this response to a specific environmental signal, gene expression was pervasive genome-wide throughout the timecourse. We detected transcripts for 6,231 genes at one or more time points in at least one strain and 6,078 genes with transcripts at one or more timepoints in both strains of which a shared set of 5,595 genes could be analyzed through our DESeq2-based pipeline (Love et al. 2014). This comprises approximately 85% of yeast genes (Supplemental Methods). 2,345 different proteins were detected, and all but two were present at every timepoint in both strains (Supplemental Methods). This is approximately 35% of the genome. Proteins that are present in low abundances may not be detected via mass spectrometry (Bantscheff et al. 2007; Peshkin et al. 2015), which may partially account for fewer proteins being present than as detected by RNA-seq.

Although large numbers of genes had detectable levels of mRNA or protein, not all were affected by pheromone induction. We tested for significant induction or repression by checking whether at least one timepoint was significantly different from time 0 (FDR < 0.05) (Spies et al. 2019). With some variation between strains, around 25% of analyzed mRNA and 34% of analyzed proteins were significantly induced or repressed, with the majority shared between strains (Table 1; Figure 1). The classification of genes as significantly induced or repressed and/or different between strains at 1 hour post-induction were not, in general, representative of analyses over the whole timecourse (Figure 2).

**Table 1:** Number of genes induced or repressed via RNA-seq and protein-MS

|  | mRNA (Total 5,595<br>genes detected) | Protein (Total 2345<br>genes detected) |
| --- | --- | --- |
| S288c | 1,274 (22.8%) | 793 (33.8%) |
| YJM145 | 1,676 (30.0%) | 796 (33.9%) |
| Both | 900 (16.1%) | 497 (21.2%) |

**Figure 1:**
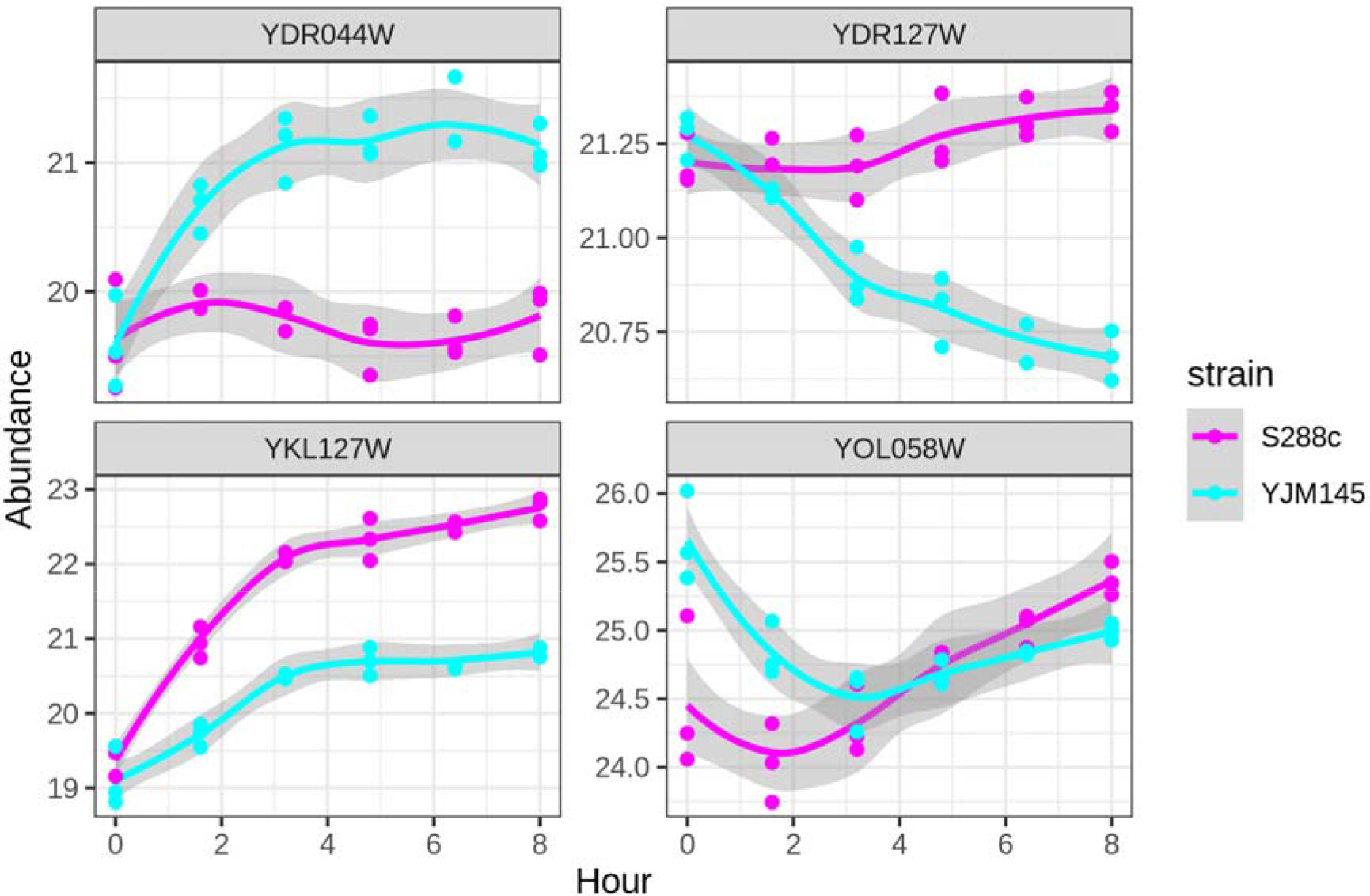
Example protein trajectories. Protein abundances were plotted over time, with the color denoting the strain. Dots represent actual experimentally measured time points, whereas the curves are the loess-smoothed fits of the linear model with shaded 95% confidence intervals, which represent our best estimate of average abundance at that time point. The trajectories for these four genes were identified to be significantly different between strains by a likelihood ratio test (see below). Top-left: YDR044W (HEM13) is involved in the heme biosynthetic pathway. Top-right: YDR127W (ARO1) is involved in chromisate biosynthesis, which is the precursor to aromatic amino acids. Bottom-left: YKL127W (PGM1) codes for phosphoglucomutase, which converts glucose-1-phosphate to glucose-6-phosphate, a process necessary for hexose metabolism. Bottom-right: YOL058W (ARG1) is an enzyme in the arginine biosynthesis pathway.

**Figure 2:**
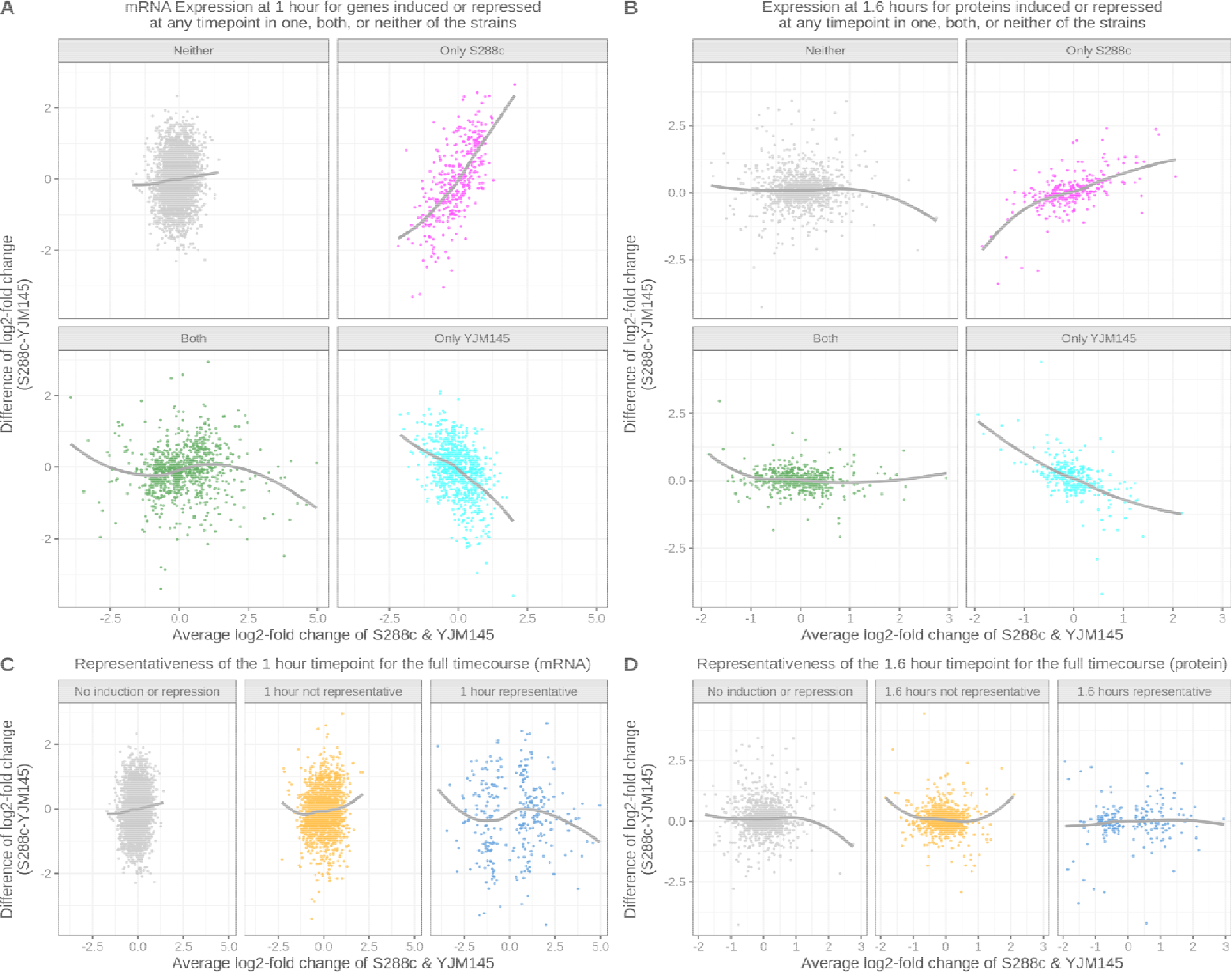
Studies comparing one timepoint to a baseline misclassify differential expression during a dynamic response. **(A)** MA plots of mRNA expression for the two strains at 1 hour after pheromone exposure separated into whether the mRNA is significantly induced or repressed during the timecourse in one, both, or neither strain. Grey lines are loess fits through the data as guides to the eye. **(B)** Like (A) except for protein expression at 1.6 hours after induction. **(C)** Like (A) except separated by whether the induction at 1 hour was representative of the whole timecourse. **(D)** Like (C except for protein at 1.6 hours.

Across genes, the correlation *between strains* for mRNA is 0.49 at 1 hour post-induction and for protein is 0.48 at 1.6 hours post-induction. However, despite seemingly low correlations, if we test for differences between strains over entire mRNA or protein trajectories by evaluating the significance of strain × timepoint interactions in a likelihood ratio test, we detect 3.9% (217 genes) of mRNA trajectories being significantly different between strains (FDR<0.05), and 4.6% (108 genes) of protein trajectories being significantly different (FDR<0.05). For mRNA, expression trajectories of genes involved in translation (including ribosomal genes) tended to be more divergent than expected while protein divergences were dominated by genes involve in amino-acid synthesis (Supplemental Data S1). These tend to be the categories of genes that are differentially expressed at the two levels. There were only 14 genes which were different between strains for both mRNA and protein. These numbers may be conservative since the false negative rate for mRNA and protein differences is not necessarily the same, but if we rank the p-values of the likelihood ratio tests for between strain differences, there is not a noticeable relationship between those for mRNA and protein (Supplemental Figure S1). This suggests that post-transcriptional regulation plays an important role, both in maintaining protein levels while mRNA differs, and in affecting protein levels without significant changes in mRNA trajectories. Large differences in mRNA do not necessarily lead to large differences in protein between strains.

If we restrict our focus to only those genes that significantly change during the response in both RNA and protein in either strain, the mean correlation between mRNA and protein trajectories is 0.19, although significantly changing genes are enriched for higher correlations (Supplemental Figure S2). The low correlation is in line with the literature (Christiano et al. 2014; Maier et al. 2009) and corroborates the idea that taking only the magnitude and direction of mRNA differential expression is a poor predictor of the behavior of its corresponding protein product, presumably because it does not take into account properties like protein degradation rates. Indeed, even qualitatively, mRNA and protein are not necessarily congruent. When the mRNA changes significantly during the timecourse, only around 2/3 of proteins follow the direction of this change. (Figure 3). Even among the significantly changing protein targets of STE12 – a transcriptional activator that is known to respond to pheromone – mRNA and protein change in the same direction only for 79 of 130 targets in s288c. This suggests that (a) even qualitative conclusions drawn from dynamic changes in mRNA abundance are not always representative of the behavior in protein levels and that (b) generalizations about protein behavior based on mRNA measurements at single timepoints can be strikingly mistaken.

**Figure 3.**
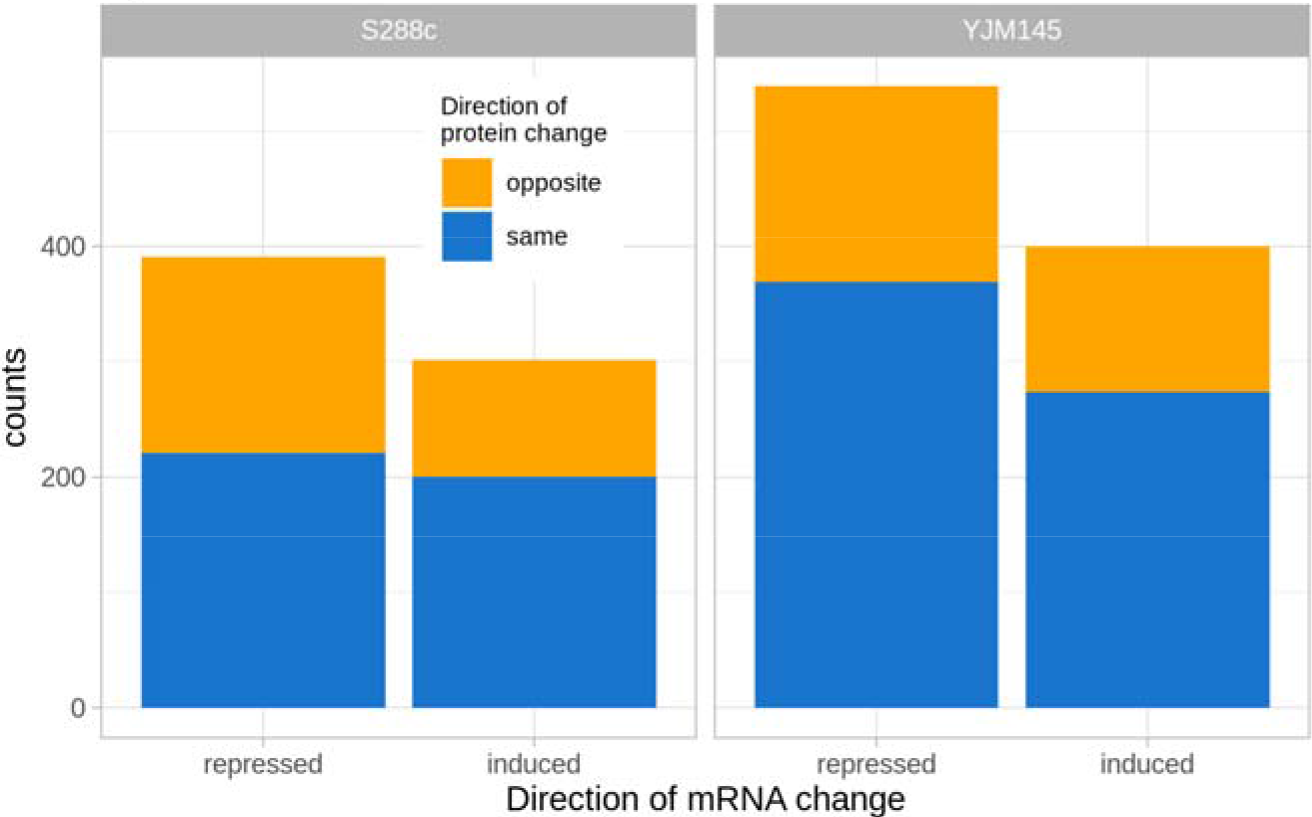
Concordance of induction or repression between mRNA and protein.

In order to bridge the disconnect between mRNA and protein levels and to study variation in the protein production process, we fit a mass action model to our protein data (Lee et al. 2011) of the form *dp/dt* = *k*_*s*_*R*(*t-τ*) – *k*_*D*_*P*(*t*), where *k*_*s*_ is the protein synthesis rate after pheromone exposure, *R*(*t−τ*) is the measured amount of mRNA at some amount of time *τ* before time *t*, *k*_*D*_ is the protein degradation rate after pheromone exposure, and *P*(*t*) is the amount of protein at time *t*. Because individual parameter estimates are often poorly constrained in systems biology models (Gutenkunst et al. 2007; Erguler and Stumpf 2011), we estimated their joint posterior distribution for each strain separately and for a model where both strains shared the same synthesis and degradation values (Supplemental Methods). This Bayesian framework allowed us to compare these rates between strains along with the structure of their covariation while accounting for the uncertainties in the parameter estimates (Gutenkunst et al. 2007; McElreath 2018).

The model fit the data well. For each gene and strain we simulated trajectories from the posterior (Figure 4). The mean normalized root mean squared error was less than 2% for approximately 60% of the genes and less than 10% for 98% of them. At the maximum *a posteriori* estimates, protein synthesis rates during the pheromone response range are log-normally distributed with the middle 50% ranging from 1.9 to 22.5 per minute. The middle 50% of protein half lives range from 1.9 to 10.3 hours with 13.1% (S288c) and 10.7% (YJM145) of proteins with half-lives less than one hour. The predicted protein half-lives were also approximately log-normally distributed (Belle et al. 2006), although we found an overabundance of short lived proteins and an underrepresentation of long-lived ones (Christiano et al. 2014) (Figure 5; Supplemental Figure S3). While this could be the result of MS bias, which tends to only pick up more abundant proteins, pheromone response induces substantial physiological changes such as cell cycle arrest, which may lead to active protein degradation for certain proteins compared with steady-state growth conditions (Bardwell 2005).

**Figure 4:**
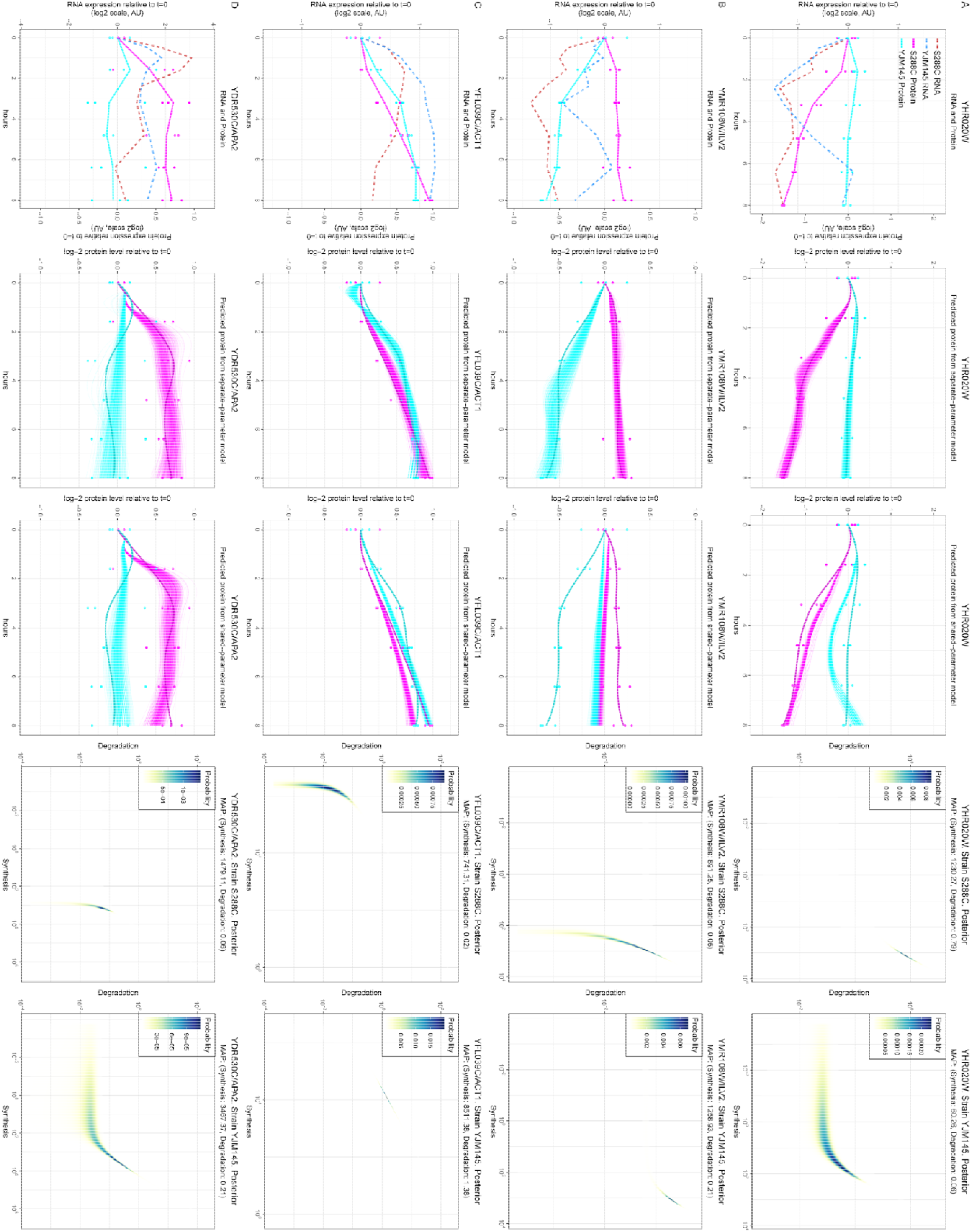
RNA, protein, and parameter variation can take several different forms. The figure is described rotated into landscape orientation. From left to right, the 5 panels in each row are: (1) the mean mRNA and protein trajectories with protein data plotted as filled circles. Dotted lines are RNA (red: S288C; blue: YJM145) and solid lines are protein (magenta: S288C; cyan: YJM145). The values are on a log scale relative to the value at t=0. (2) 500 trajectories simulated from the posterior of the separate-parameter model. Spline fits to the actual protein data are shaded in a slightly darker color than the simulations. Protein data is plotted as filled circles. As in (1), the values are on a log scale relative to the value at t=0. (3) Same as (2) except simulated trajectories are generated from the shared-parameter model. (4) Heat map of the posterior density estimate for S288C. (5) Heat map of the posterior density estimate for YJM145. (**A**): YHR020W has significantly different mRNA, protein, and parameters; (**B**) YMR108W (ILV2) has significantly different protein and parameters but not mRNA trajectories; (**C**) YFL039C (ACT1) has significantly different mRNA trajectories and parameters but not protein trajectories; (**D**) YDR530C (APA2) has shared parameter values between strains but significantly different mRNA and protein trajectories.

**Figure 5:**
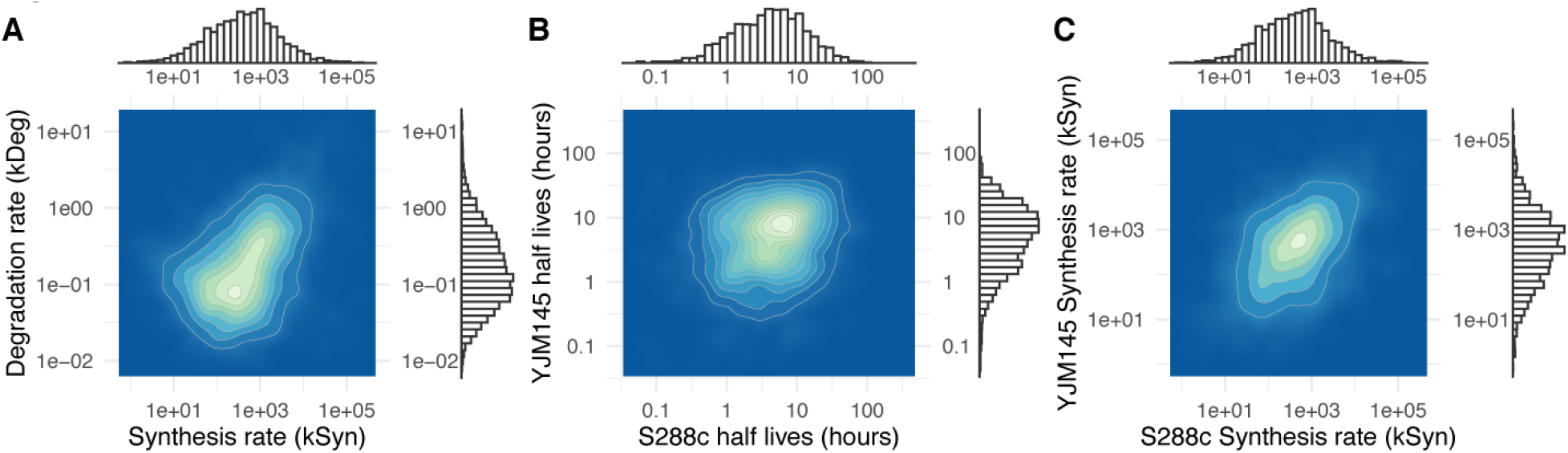
Maximum *a posteriori* parameter estimates. (**A**) Density plot and histograms of synthesis rate and degradation rate pairs for both strains together. (**B**) Density plot and histograms of protein half-lives. (**C**) Density plot and histograms of estimated protein synthesis rates during pheromone response.

In order to lead to differences in protein trajectories, mutations must alter either RNA trajectories or biochemical parameters such as the translation and degradation rates we modeled here. We found that strain-specific protein synthesis and degradation rates gave demonstrably better protein trajectory predictions than a shared parameter model based on WAIC (Watanabe 2010; McElreath 2018) for 300 of the 2,316 genes that had good data for both RNA and protein (FDR<0.05; Figure 5). Genes involved in small molecule metabolic processes were most likely to have divergent parameters between strains (FDR < 1E-13; Supplemental Data S1), particularly those active in the mitochondria (Eden et al. 2009). This apparent divergence in the parameters controlling mitochondrial protein dynamics is consistent with other results showing that YJM145 is divergent from S288c-like lab strains in mitochondrial expression (Sun et al. 2016).

Of these 2,316 genes, 406 significantly diverged between strains (FDR<0.05) in either RNA, protein, or parameters or a combination of factors (Figures 4,5). Over half of these only had detectable differences in parameter values. For 71 genes, dominated by genes involved in translation, the divergence was in RNA alone, only 1 gene showed divergence in RNA and protein trajectories without a concomitant change in parameters, and 60 genes, including those involved in biosynthesis, had significant divergence in protein trajectories and parameters. These tests have different statistical power, but we should be better able to detect RNA differences than protein differences and so we likely overestimate the number of genes diverging in RNA alone and underestimate those diverging in protein or protein and parameters. In either case, these numbers make it clear both that studies focusing on the mRNA level alone will miss a large fraction of potentially functional divergence and that parameters of these dynamic biological processes can vary between strains without leading to substantial differences in protein expression. These phenotypically neutral parameter shifts in our study place cells from the two strains in different regions of the parameter space and, we hypothesize, mean that the strains have different potentials to respond to new physiological or evolutionary pressures.

## Discussion

After an initial explosion of microarray and then RNA-seq studies exploring genome-wide variation in mRNA levels (Rifkin et al. 2003; Landry et al. 2007; Gilad et al. 2005; Ranz et al. 2003; Meiklejohn et al. 2003; Brem et al. 2002), an increasing number of studies are uncovering genetic variation affecting post-transcriptional processes (Schaefke et al. 2018; Pai et al. 2015; Pai and Gilad 2014; Battle et al. 2015). These include RNA splicing (Bradley et al. 2012; Barbosa-Morais et al. 2012; Manning and Cooper 2017), poly-adenylation (Xiao et al. 2016), RNA decay (which also affects RNA trajectories) (Andrie et al. 2014), and protein-specific processes like those investigated here (McManus et al. 2014; Wu et al. 2013; Foss et al. 2007; Horvatovich et al. 2014; Stark et al. 2014; Sun et al. 2016; Albert et al. 2014b). This variation complicates the link between DNA variation and phenotypic variation, and it is not yet clear whether there are general, systematic patterns that distinguish the kinds of biological processes that are likely to be affected by transcriptional variation versus post-transcriptional variation.

Our results show that, for any given gene, mRNA levels are not necessarily reliable predictors of protein levels even during an active response to a stimulus and that differences in mRNA abundances between strains do not necessarily translate into differences at the protein level. Much of the documented mRNA expression variation between strains and species is probably screened off from selection by processes acting downstream. This allows transcriptional variation to build up and form a reservoir that can fuel future evolutionary change.

However, although the relationship between mRNA and protein levels at a given timepoint can be tenuous, protein trajectories can be predicted from mRNA by a simple, generative, mass-action model that uses physiologically plausible parameter values to represent the post-transcriptional processes of protein synthesis and degradation. These parameter values can differ between strains, and they certainly depend upon cellular context (Bardwell 2005). Modeling the data in this way highlights the fact that protein production itself is a buffering process that can smooth over transcriptional bursts (Raj et al. 2006) and temper differences in mRNA trajectories. When protein degradation rates are slow compared to those of mRNA, then protein production will act as if it is integrating mRNA levels over time, and tweaks to protein synthesis and degradation parameters could keep two strains' protein levels similar to each other despite mRNA divergence. If, as often postulated for enzyme kinetics, the cellular effect of a protein is a saturating function of protein level (Kacser and Burns 1981; Savageau 1976), then this could allow synthesis and degradation rates to drift and build up evolutionary potential as long as they jointly maintain at least a minimum protein level. On the flip side, reconfiguring cellular state to face a new environmental challenge could require gene-specific shifts in the balance of synthesis and degradation so that the cell's complement of proteins can be rapidly turned over without having to wait for the effects of changes in mRNA numbers to percolate to the protein level. We find cases where the mRNA levels do not appreciably change after pheromone exposure while the protein increases or declines.

Our estimated posterior densities (Figure 5) show that, in general, coordinated changes to protein synthesis and degradation rates can maintain consistent protein trajectory predictions from an mRNA timecourse. However, most of the genes for which we found protein differences did not differ in their mRNA trajectories, and so buffering is not necessarily the main task of parameter variation. Instead, modifications to protein synthesis and degradation rates can directly generate differences at the protein level from the same mRNA substrate. These three knobs – mRNA levels, synthesis rates, and degradation – can be changed dynamically and either in a coordinated manner or independently of each other and give evolution the ability to finely tune protein trajectories to meet specific cellular exigencies. Over evolutionary time, they can drift, loosely bounded by stabilizing selection, and build up a reservoir of untested variation in a population that could fuel future evolutionary change.

## Methods

### Strains and experimental conditions

The two *S. cerevisiae* strains used in this study were S288c and YJM145. Both are mating type **a** with the following genes knocked out via the loxP system (Gueldener et al. 2002): *URA3*, *HO*, *AMN1*, and *BAR1*. In both strains, *FIG1* had been tagged with yECitrine using the Ca.URA3 marker for other studies. Strains were grown in 50 mL YPD overnight as a starter culture, with the experimental culture being in 1L YPD in a 3L Fernbach flask to provide proper aeration. Experimental cultures were diluted to an OD of ~0.1 and incubated for 2 hours before inducing with pheromone. Pheromone was added to achieve a final concentration of 50 nM in the media. Samples taken for the mRNA were at hours 0, 0.5, 1, 1.6, 2.4, 3.2, 4.8, 6.4, 8 where hour 0 is taken immediately before induction. Samples for the protein MS were taken from hours 0, 1.6, 3.2, 4.8, 6.4, 8. Fifty mL of each sample was taken from the flask at each time point, spun down, and flash frozen in liquid nitrogen. Experiments were performed on three separate days with the RNA and protein handled together afterwards.

Total RNA extraction of all the samples was done using the Ambion RiboPure kit. Multiplexed RNA-Seq libraries were generated from the 3' end of transcripts according to a published protocol (Kadoki et al. 2017) The protocol produces mRNA reads proportional to the number of transcripts independent of transcript length. Sequencing was done on an Illumina HiSeq4000 using single reads 50 base pairs in length. Quantitative MS was performed by the MacCoss laboratory including processing of raw MS data (see Supplemental Methods).

### Processing and statistical methods

Processing of RNA-Seq data was done on Galaxy, using BWA to align reads to the *S. cerevisiae* reference genome (R64-2-1, 2015-01-13) and htseq-count to extract read counts for each transcript(Li and Durbin 2009; Anders et al. 2015; Afgan et al. 2016). Estimation of gene counts was performed using DESeq2 (Love et al. 2014) with a generalized linear model parameterized as *count* ~ *timepoint* + *strain* + *timepoint:strain*, with *timepoint* being a factor variable(Love et al. 2014). Likelihood ratio tests between the strains use the form *count* ~ *timepoint* + *strain* as the reduced model, which would not include differences in the pre-induction time point between the strains.

Pre-processing of MS intensity values was done using Skyline, followed by sample normalization and log transformation via MSStats (Choi et al. 2014; Egertson et al. 2015) (Supplemental Methods). A general linear model using log intensities for each protein that follows the same form as the RNA-seq model was fit to the data, and the likelihood ratio test was performed the same way.

All analysis and modeling was performed in R (R Core Team 2019). Although each of our RNA replicates corresponded to a protein replicate from the same flask, including this pairing in the modeling did not improve the fit in general. We estimated posterior densities for each gene for a model where the synthesis and degradation parameters were shared between strains and one where they were allowed to differ using a weakly-informative prior on the parameters (Supplemental Methods). All fits were on a log scale as errors are thought to be symmetrical for protein levels on a log scale.

## Data access

All raw RNA-seq data generated in this study have been submitted to the NCBI Gene Expression Omnibus (GEO; https://www.ncbi.nlm.nih.gov/geo/) under accession number GSE131211. All mass spectrometry data generated in this study have been submitted to Panorama public at the URL https://panoramaweb.org/KuoRifkin.url and has the ProteomeXchange ID PXD015745. The complete processed data, R code, and outputs from the analyses will be available from the UCSD library digital collection archive at https://library.ucsd.edu/dc/

## Acknowledgements

This project was supported by National Science Foundation grant NSF MCB-1517482 to SAR and NSF MCB-1518314 to DP. SK was partially supported on NIH training grant T32GM007240-39A1.

## Author Contributions

SK, SAR, and DP designed the study. SK performed the experiments. JE, GM, MM performed the MS measurements. SK and SAR analyzed the data and wrote the paper.

## Disclosure Declaration

The authors report no conflicts of interest.

## Supplemental material

<u>Supplemental Methods</u>

<u>Supplemental Figures</u> (S1-S3)

<u>Supplemental Data S1.</u> Zip archive with the results of the GOrilla (Eden et al. 2009) GO analyses for between-strain differences of mRNA, protein, and parameters for Process, Function, and Component ontologies.

