## Supplemental Methods for "A simple mass-action model predicts genome-wide protein timecourses from mRNA trajectories during a dynamic response in two strains of *Saccharomyces cerevisiae*"

Contents

1. Methods for Data Independent Acquisition LC-MS/MS
    1.1 Sample preparation for timecourse analysis
        1.1.1 Sample Key
        1.1.2 Transfer from Filter Paper into Solution
        1.1.3 Cell Lysis
        1.1.4 BCA Assay
        1.1.5 Sample Digestion
        1.1.6 MCX Cleanup
    1.2 Data Acquistion for timecourse analysis
        1.2.1 LC-MS/MS Equipment
        1.2.2 DIA Acquisition Method
        1.2.3 Run Order
    1.3 Data Analysis
        1.3.1 Demultiplexing
        1.3.2 Data Query
        1.3.3 Post Processing
            1.3.3.1 Outlier Removal
            1.3.3.2 Imputation
            1.3.3.3 Transition Selection
            1.3.3.4 Peptide Filtering
        1.3.4 Peptide Quantification
2. Analysis of transcript counts
3. Analysis of protein counts
4. Comparisons between transcripts and proteins
5. Modeling
    5.1 Least-squares fit
    5.2 Estimation of standard deviation and time-delay parameters
    5.3 Posterior density estimation and analysis

### 1. Methods for Data Independent Acquisition LC-MS/MS

#### 1.1 Sample preparation for timecourse analysis

Samples were arranged in block randomized fashion prior to processing to avoid batch effects while maintaining statistical power. The time course replicates are run in three separate blocks. Within each of these blocks, the order of time points is randomized. For each time point within a block, the order of strains analyzed for that point is random. The batch used for any particular strain x time combination is ordered completely randomly.

##### 1.1.1 Sample Key

| Sample Number | Sample Key |
|---|---|
| **1** | 2-115-D |
| **2** | 3-104-D |
| **3** | 1-104-A |

| | |
|---|---|
| **4** | 2-115-A |
| **5** | 1-115-B |
| **6** | 2-104-B |
| **7** | 3-115-F |
| **8** | 1-104-F |
| **9** | 1-104-E |
| **10** | 2-115-E |
| **11** | 2-104-C |
| **12** | 3-115-C |
| **13** | 2-115-F |
| **14** | 2-104-F |
| **15** | 1-115-D |
| **16** | 1-104-D |
| **17** | 2-104-A |
| **18** | 3-115-A |
| **19** | 2-104-E |
| **20** | 1-115-E |
| **21** | 1-104-B |
| **22** | 2-115-B |
| **23** | 1-115-C |
| **24** | 1-104-C |
| **25** | 3-104-A |
| **26** | 1-115-A |
| **27** | 2-104-D |
| **28** | 3-115-D |
| **29** | 3-115-E |
| **30** | 3-104-E |
| **31** | 1-115-F |
| **32** | 3-104-F |
| **33** | 3-115-B |
| **34** | 3-104-B |
| **35** | 3-104-C |
| **36** | 2-115-C |

#### 1.1.2 Transfer from Filter Paper into Solution

Yeast cultures were provided on filter paper. To keep cold, samples were processed two at a time. Each sample was submerged in 3 mL 50 mM ammonium bicarbonate at Ph 7.8 to completely cover the filter paper. Samples were vortexed vigorously. If filter paper was very folded up, an attempt was made to unfold it using tweezers and resubmerge. Tubes were spun at 3000 rpm, 4°C for 2 minutes. Filter paper was removed, and tubes spun again at 3000 rpm, 4°C for 2 minutes. 1.5 mL of supernatant above the cell pellet was removed and the remaining supernatant and cells transferred into a new 1.5 mL microcentrifuge tube. The sample was spun at 4°C for 5 minutes, supernatant completely removed and cell pellet frozen at -80°C.

#### 1.1.3 Cell Lysis

Samples were lysed in batches:

| Lysis batch | Samples |
|---|---|
| **1** | 1-8 |
| **2** | 9-18 |
| **3** | 19-27 |
| **4** | 28-36 |

Cell were resuspended in 250 µL of 0.1% PPS in 50 mM ammonium bicarbonate (pH 7.8). (NOTE: A minor error was made during this step – samples 1-4 were resuspended in 750 µL instead of 250 µL). Each sample was combined with about 100 µL of glass beads in a 1.5 mL bead tube. The samples were beat with beads for 3 rounds of 1 minute beating followed by 1 minute cooling on ice. The samples were transferred upside down into a freezer and allowed to cool. A hole was poked into the bottom of each sample tube with a heated push pin, and the sample tube was placed over a new tube and returned to the freezer until all tubes were ready to be spun. Tubes are centrifuged at 4000 RPM, 4°C for 5 minutes to displace liquid into the new tube (10 minutes for 750 µL volume).

#### 1.1.4 BCA Assay

BCA standards were made with 0.1% RapiGest in 50 mM ammonium bicarbonate with BSA as standards. BCA working reagent was prepared in two batches of 4900 µL BCA Reagent A + 100 µL Reagent B.

| BSA Standard Concentration (µg/µL) | OD600 |
|---|---|
| **0.025** | 0.236 |
| **0.125** | 0.532 |
| **0.25** | 0.877 |
| **0.5** | 1.497 |
| **0.75** | 2.016 |
| **1.0** | 2.645 |

BSA Standard Curve Measurements

Sample concentration is assessed at 1/8 dilution (3.1 /ul/ sample + 21.9 /ul/ of 50 mM ammonium bicarbonate).

| # | Sample code | OD600 | Conc (µg/µL) | Volume (µL) | Protein (µg) |
|---|---|---|---|---|---|
| **1** | 2-115-D | 0.734 | 1.67 | 750 | 1252.5 |
| **2** | 3-104-D | 1.432 | 3.83 | 750 | 2872.5 |
| **3** | 1-104-A | 0.248 | 0.21 | 750 | 157.5 |
| **4** | 2-115-A | 0.296 | 0.25 | 750 | 187.5 |
| **5** | 1-115-B | 0.77 | 1.76 | 250 | 440 |
| **6** | 2-104-B | 0.437 | 0.82 | 250 | 205 |
| **7** | 3-115-F | 2.223 | 6.62 | 250 | 1655 |
| **8** | 1-104-F | 2.195 | 6.53 | 250 | 1632.5 |
| **9** | 1-104-E | 1.328 | 3.55 | 250 | 887.5 |
| **10** | 2-115-E | 1.113 | 2.97 | 250 | 742.5 |
| **11** | 2-104-C | 0.916 | 2.09 | 250 | 522.5 |
| **12** | 3-115-C | 2.159 | 6.43 | 250 | 1607.5 |
| **13** | 2-115-F | 2.131 | 6.34 | 250 | 1585 |
| **14** | 2-104-F | 1.222 | 3.27 | 250 | 817.5 |
| **15** | 1-115-D | 1.247 | 3.33 | 250 | 832.5 |
| **16** | 1-104-D | 1.154 | 3.08 | 250 | 770 |
| **17** | 2-104-A | 0.271 | 0.23 | 250 | 57.5 |
| **18** | 3-115-A | 0.579 | 1.09 | 250 | 272.5 |
| **19** | 2-104-E | 1.852 | 5.51 | 250 | 1377.5 |
| **20** | 1-115-E | 1.181 | 2.69 | 250 | 672.5 |
| **21** | 1-104-B | 0.636 | 1.2 | 250 | 300 |
| **22** | 2-115-B | 0.582 | 1.09 | 250 | 272.5 |
| **23** | 1-115-C | 0.853 | 1.95 | 250 | 487.5 |
| **24** | 1-104-C | 0.978 | 2.23 | 250 | 557.5 |
| **25** | 3-104-A | 0.78 | 1.78 | 250 | 445 |
| **26** | 1-115-A | 0.283 | 0.24 | 250 | 60 |
| **27** | 2-104-D | 1.462 | 3.91 | 250 | 977.5 |
| **28** | 3-115-D | 2.232 | 6.64 | 250 | 1660 |
| **29** | 3-115-E | 2.181 (half) | 12.98 | 250 | 3245 |
| **30** | 3-104-E | 1.993 (half) | 11.86 | 250 | 2965 |

| | | | | | |
|---|---|---|---|---|---|
| **31** | 1-115-F | 1.280 (half) | 6.84 | 250 | 1710 |
| **32** | 3-104-F | 2.283 (half) | 13.59 | 250 | 3397.5 |
| **33** | 3-115-B | 1.706 | 4.56 | 250 | 1140 |
| **34** | 3-104-B | 1.038 | 2.37 | 250 | 592.5 |
| **35** | 3-104-C | 1.657 | 4.43 | 250 | 1107.5 |
| **36** | 2-115-C | 1.461 | 3.9 | 250 | 975 |

Sample concentrations. (half) means sample was too concentrated and had to be havled in the spectrophotometer (50 µL water + 50 µL BCA sample

#### 1.1.5 Sample Digestion

Samples were digested in two batches (samples 1-18, and samples 19-36). For each sample, 50 µg of protein was digested. Prior to digestion, 188 ng of a human ApoA1 protein standard was added to assess digestion. Samples were reduced with the addition of 500 mM dithiothreitol (DTT) to a concentration of 5 mM and incubation at 60°C for 30 minutes. Reduced samples are alkylated via the addition of 500 mM iodoacetamide (IAA) to a final concentration of 15 mM and incubation at room temperature (22-25°C) for 30 minutes in the dark. Alkylation reactions are quenched via the addition of an additional aliquot of 500 mM dithiothreitol to bring the final concentration to 10 mM. Each reduced and alkylated plasma dilution is digested with 1 µg (for 1:50 trypsin:protein) of sequencing grade modified porcine trypsin for 4 hours at 37°C/700 RPM. Digestion is quenched by the addition of 5M HCl to a final concentration of 200 mM. Acidifed digests are incubated for one hour at room temperature to facilitate hydrolysis of the PPS surfactant. Digested standards are centrifuged at 1400 rpm/4 °C for 5 minutes in a bench-top centrifuge to pellet.

#### 1.1.6 MCX Cleanup

Samples were cleaned using Waters Oasis MCX cartridges according to the manufacturer’s protocol. MCX cleanup was performed in batches of 9:

| Batch | Samples |
|---|---|
| **1** | 1-9 |
| **2** | 10-18 |
| **3** | 19-27 |
| **4** | 28-36 |

After MCX, samples were dried by speed vaccuum and resuspended in 199.5 µL of 0.1% formic acid. The Thermo Peptide Retention Time Calibration (PRTC) standard was added to each resuspended digest to a concentration of 50 fmol/µL and total volume of 200 µL as a loading control. 20 µL of each sample digest was transferred to polypropylene auto-sampler vials with snap-on lids and stored at -20 °C while queued for injection. The rest of the sample (~180 µL) is stored at -80 °C.

### 1.2 Data Acquistion

#### 1.2.1 LC-MS/MS Equipment

A Waters nanoAcquity UPLC system coupled in-line to a Thermo Orbitrap Fusion mass spectrom- eter is used for LC-MS/MS analysis. A trapping column is used for sample cleanup prior to LC separation. The trapping column is a 150 µM inner diameter fused-silica column with a Kasil frit that is packed with 3 µM C18 reversed phase packing material (Reprosil Pur 120 C-18-AQ from Dr. Maisch GmbH) to a bed length of ~4cm. The trapping column is coupled to an analytical column. The analytical column is a 75 µM inner diameter PicoFrit column ordered from New Objective and packed to 30 cm with the same stationary phase (C18) as the trapping column. In each mass spectrometry run, 4 µL of sample (diluted to 250 ng/µL in buffer A) is loaded onto the trapping column and washed with a mixture of 98% buffer A (0.1% formic acid in water) and 2% buffer B (0.1% formic acid in acetonitrile). After trapping, the sample is loaded onto the analytical column and separated over a 90 minute linear gradient from 2-35% buffer B. As the digest is separated, it is eluted from the column and ionized into the mass spectrometer via an electrospray ionization interface with a 2kV spray voltage.

#### 1.2.2 DIA Acquisition Method

The DIA acquisition method acquires comprehensive MS/MS data on all precursors between 500 and 900 m/z. The acquisition consists of a cycle of MS/MS scan acquisition with an MS scan acquired every 21 MS/MS scans.

MS1 Acquisition Parameters

| Acquisition Parameter | Value |
|---|---|
| **Isolation Mode** | Quadrupole |
| **Isolation Window** | 485 - 925 m/z |
| **Detector** | Orbitrap |
| **Resolution** | 60,000 |
| **Scan Range** | 350 - 1000 m/z |
| **Maximum Injection Time** | 20 ms |
| **AGC Target** | 2e5 |
| **S-Lens RF** | 60 |

MS2 Acquisition Parameters

| Acquisition Parameter | Value |
|---|---|
| **Isolation Mode** | Quadrupole |
| **Isolation Window Width** | 20 m/z |
| **Detector** | Orbitrap |
| **Resolution** | 30,000 |
| **Scan Range** | 100 - 2000 m/z |
| **Maximum Injection Time** | 60 ms |
| **AGC Target** | 1e5 |

| | |
|---|---|
| **S-Lens RF** | 60 |
| **Activation Type** | HCD |
| **Normalized Collision Energy** | 30 |
| **Default Charge State** | 2 |

Acquisition Cycle for MS/MS Scans

| Scan Event | Target m/z |
|---|---|
| **1** | 510.481923 |
| **2** | 530.491018 |
| **3** | 550.500112 |
| **4** | 570.509207 |
| **5** | 590.518302 |
| **6** | 610.527397 |
| **7** | 630.536492 |
| **8** | 650.545587 |
| **9** | 670.554682 |
| **10** | 690.563777 |
| **11** | 710.572872 |
| **12** | 730.581967 |
| **13** | 750.591062 |
| **14** | 770.600157 |
| **15** | 790.609252 |
| **16** | 810.618347 |
| **17** | 830.627442 |
| **18** | 850.636537 |
| **19** | 870.645632 |
| **20** | 890.654727 |
| **21** | 910.663822 |
| **22** | 500.477375 |
| **23** | 520.48647 |
| **24** | 540.495565 |
| **25** | 560.50466 |
| **26** | 580.513755 |
| **27** | 600.52285 |

| | |
|---|---|
| **28** | 620.531945 |
| **29** | 640.54104 |
| **30** | 660.550135 |
| **31** | 680.55923 |
| **32** | 700.568325 |
| **33** | 720.57742 |
| **34** | 740.586515 |
| **35** | 760.59561 |
| **36** | 780.604705 |
| **37** | 800.6138 |
| **38** | 820.622895 |
| **39** | 840.63199 |
| **40** | 860.641085 |
| **41** | 880.65018 |
| **42** | 900.659275 |

#### 1.2.3 Run Order

Sample order is randomized prior to mass spectrometry data acquisition.

| UW # | Filename | UCSD number | UCSD name | Date | Strain | Time |
|---|---|---|---|---|---|---|
| **QC** | 29July2016-FU-QC01 | - | - | - | - | - |
| **QC** | 29July2016-FU-QC02 | - | - | - | - | - |
| **QC** | 29July2016-FU-QC03 | - | - | - | - | - |
| **QC** | 29July2016-FU-QC04 | - | - | - | - | - |
| **10** | 29July2016-FU-UCSD-10-rep2-115-6 4hr-pre-01 | 2-115-E | Rep2-115-6.4hr | 22-Mar | 115 | 6.4 hr |
| **QC** | 29July2016-FU-QC05 | - | - | - | - | - |
| **10** | 29July2016-FU-UCSD-10-rep2-115-6 4hr-01 | 2-115-E | Rep2-115-6.4hr | 22-Mar | 115 | 6.4 hr |
| **25** | 29July2016-FU-UCSD-25-rep3-104-0hr-01 | 3-104-A | Rep3-104-0hr | 30-Mar | 104 | 0 hr |
| **28** | 29July2016-FU-UCSD-28-rep3-115-4 8hr-01 | 3-115-D | Rep3-115-4.8hr | 30-Mar | 115 | 4.8 hr |
| **23** | 29July2016-FU-UCSD-23-rep1-115-3 2hr-01 | 1-115-C | Rep1-115-3.2hr | 18-Mar | 115 | 3.2 hr |
| **34** | 29July2016-FU-UCSD-34-rep3-104-1 6hr-01 | 3-104-B | Rep3-104-1.6hr | 30-Mar | 104 | 1.6 hr |
| **26** | 29July2016-FU-UCSD-26-rep1-115-0hr-01 | 1-115-A | Rep1-115-0hr | 18-Mar | 115 | 0 hr |
| **QC** | 29July2016-FU-QC06 | - | - | - | - | - |
| **35** | 29July2016-FU-UCSD-35-rep3-104-3 2hr-01 | 3-104-C | Rep3-104-3.2hr | 30-Mar | 104 | 3.2 hr |
| **5** | 29July2016-FU-UCSD-5-rep1-115- | 1-115-B | Rep1-115-1.6hr | 18-Mar | 115 | 1.6 hr |

| | | | | | | |
|---|---|---|---|---|---|---|
| | 1 6hr-01 | | | | | |
| **7** | 29July2016-FU-UCSD-7-rep3-115-8hr-01 | 3-115-F | Rep3-115-8hr | 30-Mar | 115 | 8 hr |
| **9** | 29July2016-FU-UCSD-9-rep1-104-6 4hr-01 | 1-104-E | Rep1-104-6.4hr | 18-Mar | 104 | 6.4 hr |
| **24** | 29July2016-FU-UCSD-24-rep1-104-3 2hr-01 | 1-104-C | Rep1-104-3.2hr | 18-Mar | 104 | 3.2 hr |
| **27** | 29July2016-FU-UCSD-27-rep2-104-4 8hr-01 | 2-104-D | Rep2-104-4.8hr | 22-Mar | 104 | 4.8 hr |
| **QC** | 29July2016-FU-QC07 | - | - | - | - | - |
| **17** | 29July2016-FU-UCSD-17-rep2-104-0hr-01 | 2-104-A | Rep2-104-0hr | 22-Mar | 104 | 0 hr |
| **6** | 29July2016-FU-UCSD-6-rep2-104-1 6hr-01 | 2-104-B | Rep2-104-1.6hr | 22-Mar | 104 | 1.6 hr |
| **1** | 29July2016-FU-UCSD-1-rep2-115-4 8hr-01 | 2-115-D | Rep2-115-4.8hr | 22-Mar | 115 | 4.8 hr |
| **30** | 29July2016-FU-UCSD-30-rep3-104-6 4hr-01 | 3-104-E | Rep3-104-6.4hr | 30-Mar | 104 | 6.4 hr |
| **19** | 29July2016-FU-UCSD-19-rep2-104-6 4hr-01 | 2-104-E | Rep2-104-6.4hr | 22-Mar | 104 | 6.4 hr |
| **32** | 29July2016-FU-UCSD-32-rep3-104-8hr-01 | 3-104-F | Rep3-104-8hr | 30-Mar | 104 | 8 hr |
| **QC** | 29July2016-FU-QC08 | - | - | - | - | - |
| **4** | 29July2016-FU-UCSD-4-rep2-115-0hr-01 | 2-115-A | Rep2-115-0hr | 22-Mar | 115 | 0 hr |
| **22** | 29July2016-FU-UCSD-22-rep2-115-1 6hr-01 | 2-115-B | Rep2-115-1.6hr | 22-Mar | 115 | 1.6 hr |
| **13** | 29July2016-FU-UCSD-13-rep2-115-8hr-01 | 2-115-F | Rep2-115-8hr | 22-Mar | 115 | 8 hr |
| **20** | 29July2016-FU-UCSD-20-rep1-115-6 4hr-01 | 1-115-E | Rep1-115-6.4hr | 18-Mar | 115 | 6.4 hr |
| **14** | 29July2016-FU-UCSD-14-rep2-104-8hr-01 | 2-104-F | Rep2-104-8hr | 22-Mar | 104 | 8 hr |
| **8** | 29July2016-FU-UCSD-8-rep1-104-8hr-01 | 1-104-F | Rep1-104-8hr | 18-Mar | 104 | 8 hr |
| **QC** | 29July2016-FU-QC09 | - | - | - | - | - |
| **29** | 29July2016-FU-UCSD-28-rep3-115-6 4hr-01* | 3-115-E | Rep3-115-6.4hr | 30-Mar | 115 | 6.4 hr |
| **2** | 29July2016-FU-UCSD-2-rep3-104-4 8hr-01 | 3-104-D | Rep3-104-4.8hr | 30-Mar | 104 | 4.8 hr |
| **12** | 29July2016-FU-UCSD-12-rep3-115-3 2hr-01 | 3-115-C | Rep3-115-3.2hr | 30-Mar | 115 | 3.2 hr |
| **31** | 29July2016-FU-UCSD-31-rep1-115-8hr-01 | 1-115-F | Rep1-115-8hr | 18-Mar | 115 | 8 hr |
| **16** | 29July2016-FU-UCSD-16-rep1-104-4 8hr-01 | 1-104-D | Rep1-104-4.8hr | 18-Mar | 104 | 4.8 hr |
| **36** | 29July2016-FU-UCSD-36-rep2-115-3 2hr-01 | 2-115-C | Rep2-115-3.2hr | 22-Mar | 115 | 3.2 hr |
| **QC** | 29July2016-FU-QC10 | - | - | - | - | - |
| **21** | 29July2016-FU-UCSD-21-rep1-104-1 6hr-01 | 1-104-B | Rep1-104-1.6hr | 18-Mar | 104 | 1.6 hr |
| **33** | 29July2016-FU-UCSD-33-rep3- | 3-115-B | Rep3-115-1.6hr | 30-Mar | 115 | 1.6 hr |

| | | | | | | |
|---|---|---|---|---|---|---|
| | 115-1 6hr-01 | | | | | |
| **18** | 29July2016-FU-UCSD-18-rep3-115-0hr-01 | 3-115-A | Rep3-115-0hr | 30-Mar | 115 | 0 hr |
| **11** | 29July2016-FU-UCSD-11-rep2-104-3 2hr-01 | 2-104-C | Rep2-104-3.2hr | 22-Mar | 104 | 3.2 hr |
| **15** | 29July2016-FU-UCSD-15-rep1-115-6 4hr-01* | 1-115-D | Rep1-115-4.8hr | 18-Mar | 115 | 4.8 hr |
| **3** | 29July2016-FU-UCSD-3-rep1-104-0hr-01 | 1-104-A | Rep1-104-0hr | 18-Mar | 104 | 0 hr |
| **QC** | 29July2016-FU-QC11 | - | - | - | - | - |

Mass Spectrometry Data Acquisition Order Files annotated with an asterisk have an error in the filename with the erroneous portion in bold.

### 1.3 Data Analysis

### 1.3.1
#### Demultiplexing

The DIA data was acquired using an overlapping window isolation scheme (see Section 2.1.2: DIA Acquisition Method). The overlapping windows are demultiplexed to effectively double the pre- cursor selectivity (20 m/z + 10 m/z). First, the .raw files were converted to vendor-neutral .mzML files. During this step, the data were also centroided using Thermo's proprietary centroid- ing algorithm. This conversion was done using `msconvert` from ProteoWizard release 3.0.9810 (`http://proteowizard.sourceforge.net/`). The converted mzML files were demultiplexed using `PRISM` version 0.0.1 (`https://bitbucket.org/maccosslab/prism`).

`msconvert` Parameters

| Parameter Name | Value |
|---|---|
| **outdir** | .\mzml |
| **mzML** | true |
| **zlib** | true |
| **mz64** | true |
| **inten64** | true |
| **simAsSpectra** | true |
| **filter** | "peakPicking vendor msLevel=1-2" |

`PRISM` Parameters

| Parameter Name | Value |
|---|---|
| **optimization** | overlap only |

#### 1.3.2 Data Query

Demultiplexed files were queried using the MacCoss lab yeast library generated from Saccharomyces cerevisiae strain S288C. The data query was performed using `EncyclopeDIA` version 0.2.8. `EncyclopeDIA` returns peptide detections with an experiment-wide FDR threshold of 1.0%. `EncyclopeDIA` also determines peak integration boundaries for detected peptides and filters out transitions that contain interference.

EncyclopeDIA Parameters

| Parameter Name | Value |
|---|---|
| **Data Acquisition Type** | Non-Overlapping DIA |
| **Precursor Window Width** | 10 |
| **Enzyme** | Trypsin |
| **Fragmentation** | CID (B/Y) |
| **Proteome Type** | Standard Proteome |
| **Precursor (PPM)** | 10 |
| **Fragment (PPM)** | 10 |
| **Number of Quantitative Ions** | 5 |
| **Number of Cores** | 7 |
| **RT Align?** | yes |

#### 1.3.3 Post Processing

##### 1.3.3.1 Outlier Removal

At times, peptide false discoveries will result in a peptide being detected in one or more runs at a retention time that is inconsistent with the majority of other runs. A python script is used to detect and remove these outliers. To detect outliers, a non-parametric (support vector regressor) alignment is used to align detected peptide retention times from all runs to a single “reference” run (randomly chosen). After alignment, the distribution of aligned retention time differences for all shared peptides between any pair of files can be calculated. A normal distribution is fit to the inner 99th percentile of this distribution as an estimate of the distribution of retention times for non-outlier peptides. Outliers are determined on a peptide, by peptide basis. For each peptide, the biggest “cluster” of peak retention times is calculated. All peaks in a cluster are within 3-sigma of the RT error calculated for non-outlier peptides with its nearest-neighbor. If the largest cluster does not represent at least 50% of the peaks considered, the peptide is removed from all downstream analysis. If the largest cluster is the majority, all peaks outside of this cluster are marked as outliers.

##### 1.3.3.2 Imputation

Many peptides will not be detected in every LC-MS/MS run. Additionally, some peptides will have outlier detections removed (see 3.3.1 Outlier Removal), resulting in peptides with “missing values”in a subset of the acquired mass spectrometry runs. To fill in the blanks, retention time boundaries are imputed from MS runs that have a dection for the peptide using a pair-wise non-parametric (support vector regressor) retention time alignment.

##### 1.3.3.3 Transition Selection

EncyclopeDIA filters out transitions with interference (see 3.2 Data Query). However, to quantify peptides across multiple runs, the same transitions must be used in every run. The transitions used are the top 5 most intense transitions without interference from the LC-MS/MS run where the peptide had the highest intensity. Any peptides with less than three transitions selected after this step are removed from the analysis.

##### 1.3.3.4 Peptide Filtering

Usually, it is preferable to quantify only peptides that uniquely map to a single protein, based on existing sequence data (open reading frames). The quantitative analysis is performed with, and without this filter. The filtering for unique peptides was calculated using an S288C reference sequence database from the Saccharomyces Genome Database. The ORF sequences were down- loaded on 8/15/2016 from `http://downloads.yeastgenome.org/sequence/S288C_reference/orf_protein/orf_trans.fasta.gz`. “Dubious” ORFs and pseudogenes were not included in this analysis. If a peptide mapped to two related proteins (e.g. YPL257W-A and YPL257W-B), it was not excluded from the analysis unless it also mapped to another unrelated protein.

1.3.4 Peptide Quantification

`Skyline-daily` version 3.5.1.999 (`https://skyline.gs.washington.edu`) was used to extract chromatograms from the data for quantification. The chromatograms to extract are defined by the transitions selected for each peptide in the section Peptide Filtering. The integration boundaries are also already determined by `EncyclopeDIA` (Section 1.3.2: Data Query) or imputation (Section 1.3.3.2: Imputation). Skyline-daily integrates the chromatograms and performs background subtraction to improve quantitation. Similar to the `EncyclopeDIA` analysis, chromatograms are extracted from the centroided, demultiplexed data with a +/- 10 ppm tolerance. Data is exported from the Skyline document as a “report” file (comma-separated-value text) containing integrated peak areas for each peptide / run. After this report export, the report is modified to normalize each individual LC-MS/MS run by its integrated total ion current (TIC) extracted from MS scans. This corrects for differences in total material analyzed in each LC-MS/MS run, which may vary due to inaccu- racies in the protein concentration assessment (see Section 1.1.4: BCA Assay) or imprecision in the volume of sample loaded onto the LC-MS/MS column in each run by the autosampler (Section 1.2.1: LC-MS/MS Equipment**).**

2. Analysis of transcript counts

Processing of RNA-Seq data was done on Galaxy, using BWA to align reads to the *S. cerevisiae* reference genome (R64-2-1, 2015-01-13) and htseq-count to extract read counts for each transcript.(Li and Durbin 2009; Anders et al. 2015; Afgan et al. 2016) All subsequent analyses were performed in R (version 3.6.1).(R Core Team 2019; Wickham and Grolemund 2016)

*file: FullAnalysis.Rmd* [Analyses below are annotated with the chunk in the .Rmd file]
We filtered the count data to remove rows with all zeros [chunk 2] and performed an initial PCA analysis to identify and remove outlier experiments [chunks 3-4]. Using this cleaned dataset, we used DESeq2 (version 1.24.0)(Love et al. 2014) to estimate log-fold change in mRNA expression relative to the initial, pre-exposure timepoint [chunk 8]. Although ideally we would treat the timecourses as timecourses, for short time series classic pairwise-comparison tools like DESeq2 are still more robust and have fewer false positives than specialized RNA-seq timecourse methods.(Spies et al. 2019; Fischer et al. 2018; Nueda et al. 2014; Conesa et al. 2006; Conesa and Nueda 2013; Äijö et al. 2014; Liang and Kelemen 2018; Bacher et al. 2018; Topa and Honkela 2018) Genes with zero counts for most samples give rise to log-fold estimates of zero or NA. We removed these from the dataset [chunk 9]. To reduce the decrease in power due to multiple testing adjustment, before applying the Benjamini-Hochberg FDR procedure,(Benjamini and Hochberg 1995) DESeq2 initially filters out genes with mean normalized counts below a threshold since these are unlikely to be detected as differentially expressed and then bases its adjustment on the genes that remain.(Love et al. 2014) We modified this procedure, basing the filtering on the maximum instead of the mean, maximizing the number of genes identified as differentially expressed, and using a false discovery rate of 5% instead of the default 10%. The results are similar with either cutoff [chunk 9; file: *pvalueAdjustment_modified.R*].

Gene expression is dynamic, but many studies of differential expression, including a previous analysis of pheromone response in these strains,(Zheng et al. 2010) compare only two timepoints. We calculated Pearson correlation of mRNA levels between strains at the 1 hour timepoint and also determined whether the genes identified as significantly induced or repressed and as different between strains at one timepoint were representative of the whole timecourse [chunk 11].

In order to compare mRNA trajectories between strains, we reran DESeq2 using a likelihood ratio test, comparing a full model that included main effects for strain and timepoint and a strain x timepoint interaction term to a reduced model with no interaction. We controlled for multiple testing using FDR as above. [chunks 13-14].

### 3. Analysis of the protein counts

We removed duplicate peptides from the Skyline output and used MSstats(Choi et al. 2014) to transform the raw data into experiment-normalized data for analysis [chunks 18-19]. We analyzed the protein data to mimic our mRNA analysis, analyzing induction or repression after pheromone exposure within each strain [chunks 20-22, 24-25] and using likelihood ratio tests as described above to compare the strains to each other [chunks 23-24, 28].

As for the mRNA, we evaluated results based solely on differential expression at the 1.6 hour timepoint with results based on the whole timecourse [chunk 27].

### 4. Comparisons between mRNA and protein

For analyses involving both RNA and protein, we restricted the dataset to genes with good data for both strains for both molecule types [chunks 30-33]. We calculated Pearson correlations between mRNA and protein for each gene just using the 6 timepoints they shared [chunk 34].

### 5. Modeling

As described in the main text, we fit a simple mass-action model of the form $dp/dt = k_s R(t-\tau) - k_D P(t)$ to the data, predicting protein expression trajectories from mRNA trajectories. $k_s$ is the protein synthesis rate after pheromone exposure, $R(t-\tau)$ is the measured amount of mRNA at some amount of time $\tau$ before time $t$, $k_D$ is the protein degradation rate after pheromone exposure, and $P(t)$ is the amount of protein at time $t$. To simulate the protein timecourse, we fit an interpolating spline(Forsythe et al. 1977) to the mean RNA estimate from DESeq2 and used this to generate a quasi-continuous protein curve. We used non-log RNA values to generate non-log protein predictions, but evaluated the protein fits on a log-2 scale. We carried out this modeling in four stages.

#### 5.1 Least squares fit

In order to make reasonable priors for the posterior density estimation, we first fit the model using the L-BFGS-B method which allows for bounded optimization(Byrd et al. 1995), minimizing the root mean squared error between the prediction and the three replicates. We fixed timepoint zero at the exponentiated mean of the log protein values and so we did not include this timepoint in our error calculation. The parameter $\tau$ represents a delay in protein synthesis and protein folding, and we constrained it between 0.01 and 1 hours. All estimation was done separately for each gene x strain combination [chunk 37].

We used these point estimates for the parameters to generate bounded, weakly informative priors for Bayesian estimation of the posterior parameter densities. We assumed that errors on a log scale were normally distributed, and so the Bayesian model fit to each gene x strain combination involved four parameters: the three above and a standard deviation of the normal distribution for the likelihood estimates. [file: *BayesAnalysis_CompareSingleStrains.Rmd*; chunk b1.3] The priors we used were:

$$\log_{10}(kSyn) \sim Normal(2.2, 2.55);\ \ -3.8 \le \log_{10}(kSyn) \le 8.2$$

$$\log_{10}(kDeg) \sim Normal(0.7, 1.2);\ \ -3.7 \le \log_{10}(kDeg) \le 2.3$$

$$\log_{10}(SD) \sim Normal(-0.5, 0.96);\ \ --2.5 \le \log_{10}(SD) \le 1.5$$

$$\tau \sim Uniform(0.1, 1)$$

5.2 Estimation of standard deviation and time-delay parameters
For each gene x strain combination, we ran an MCMC algorithm using the DEzs differential evolution sampler in the R package BayesianTools with 3 chains, 11,000 iterations, and a burn-in of 1,000 iterations. [chunks b1.4, b1.9].(ter Braak and Vrugt 2008; Hartig et al. 2019) This gave a rough picture of the 4-dimensional posterior along with marginal posterior distributions for each parameter. Generally, the standard deviations had a tight posterior mode while the time delays were not very resolved. The standard deviation of the errors for the likelihood estimates and the time delay are nuisance parameters in that they are useful for fitting the model but do not carry the same biological interest as the synthesis and degradation rates. The time delay could be interesting, but we do not have the resolution early in the timecourse to truly narrow the estimate. In order to improve the posterior resolution for the synthesis and degradation rates and to facilitate later comparisons between the posteriors for the two strains and for a shared model, we fixed the standard deviation and time delay for each gene x strain combination at their values at the maximum *a posteriori* estimate of the 4D posterior from the MCMC [chunk b1.9]. We then carried out a grid approximation to the posterior on the 2D synthesis-degradation parameter plane.

5.3 Posterior density estimation and analysis
Separately for each gene x strain combination, we sampled the synthesis-degradation parameter plane with a resolution of 0.01 (on a log10 scale) starting by filling in the convex hull of the region identified by the MCMC run and proceeding outwards until the posterior estimate dropped below a threshold of 10,000 fold less than the maximum *a posteriori* estimate [chunks b1.8-b1.9]. This procedure resulted in a 2D posterior distribution of synthesis and degradation rates for each gene x strain.

For two genes (YKL127W and YBR208C) the posterior distributions estimated in (3) occupied disjoint regions in the parameter space; all of the rest had some overlap in their support, but the posterior support for one strain often did not include the high-probability regions for the other. In order to facilitate between-strain comparisons and to make it possible to compare a strain-specific parameter model to a model where the synthesis and degradation parameters were shared between strains, we expanded our estimates of the posterior for each strain over all parameter values in the support of either strain, also filling in any holes between the support regions for each strain [file: *BayesAnalysis_CompareSharedDiffDoubleStrains.Rmd;* chunks 6, 9-10].

We compared strain posteriors to each other by a variety of methods, including the Jensen-Shannon metric [chunk b2.17].(Endres and Schindelin 2003) By these metrics, the strain-specific posteriors were very different for the majority of genes. However, our key comparison was whether a model with strain-specific synthesis and degradation parameters made significantly better predictions than a model where the strains shared these parameters.

Strain - specific parameter model

$$\frac{dP_{S288c}}{dt} = kSyn_{S288c} * mRNA_{S288c}(t - \tau_{S288c}) - kDeg_{S288c} * P_{S288c}(t)$$

$$\frac{dP_{YJM145}}{dt} = kSyn_{YJM145} * mRNA_{YJM145}(t - \tau_{YJM145}) - kDeg_{YJM145} * P_{YJM145}(t)$$

Shared parameter model

$$\frac{dP_{S288c}}{dt} = kSyn_{shared} * mRNA_{S288c}(t - \tau_{S288c}) - kDeg_{shared} * P_{S288c}(t)$$

$$\frac{dP_{YJM145}}{dt} = kSyn_{shared} * mRNA_{YJM145}(t - \tau_{YJM145}) - kDeg_{shared} * P_{YJM145}(t)$$

The posterior for the strain-specific parameter model can be considered to be the Kronecker product of the two 2D densities, while the shared parameter model is a 2D slice through this 4D space. We used WAIC (Watanabe 2010; Gelman et al. 2014; McElreath 2018) to compare these two models, although our datapoints are not independent. Because WAIC is a sum of components calculated for each datapoint individually, model performance can be paired at each datapoint, making it possible to estimate a standard error for the difference in overall WAIC values(McElreath 2018) and therefore to calculate a standardized statistic for the WAIC difference [chunk b2.7-b2.8] that can be used to compare the predictive ability of the different models. We estimated a standardized statistic in this way and compared the standardized statistic to a t-distribution with 29 degrees of freedom.

The posterior densities form an oriented cloud in the parameter space. We estimated the sloppy and stiff directions of the parameters by sampling 100,000 parameter pairs within 0.25 log10 units from the MAP estimate, weighted by their posterior probabilities, and then estimating the eigenvectors of the synthesis-degradation covariance matrix. These eigenvectors capture the principal directions of variation at the MAP. The eigenvalues and their ratio are less meaningful because they depend upon the number of pairs and the radius around the MAP [chunks b2.13-b2.14].
