## Supplemental Figures for "A simple mass-action model predicts genome-wide protein timecourses from mRNA trajectories during a dynamic response in two strains of *Saccharomyces cerevisiae*"

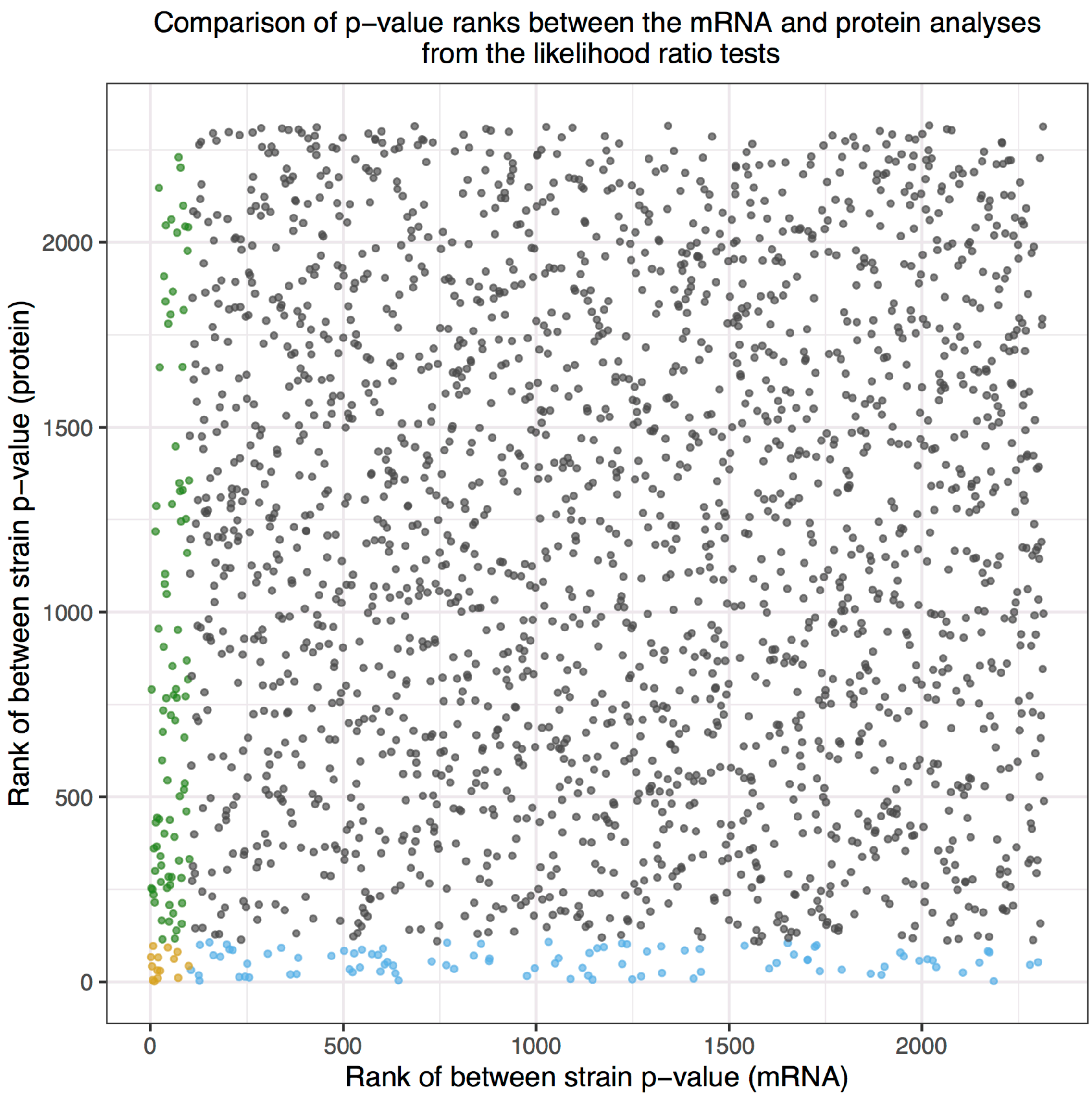


Supplemental Figure S1. Between strain differences in mRNA do not necessarily translate into between strain differences in protein. Scatter plot of the ranks of the likelihood ratio test statistic for between strain differences. Each dot represents a gene. Green indicates genes that are significantly different (FDR<0.05) between strains for mRNA but not protein. Blue: significantly different (FDR<0.05) for protein but not mRNA. Yellow: significantly different for both mRNA and protein (FDR<0.05).

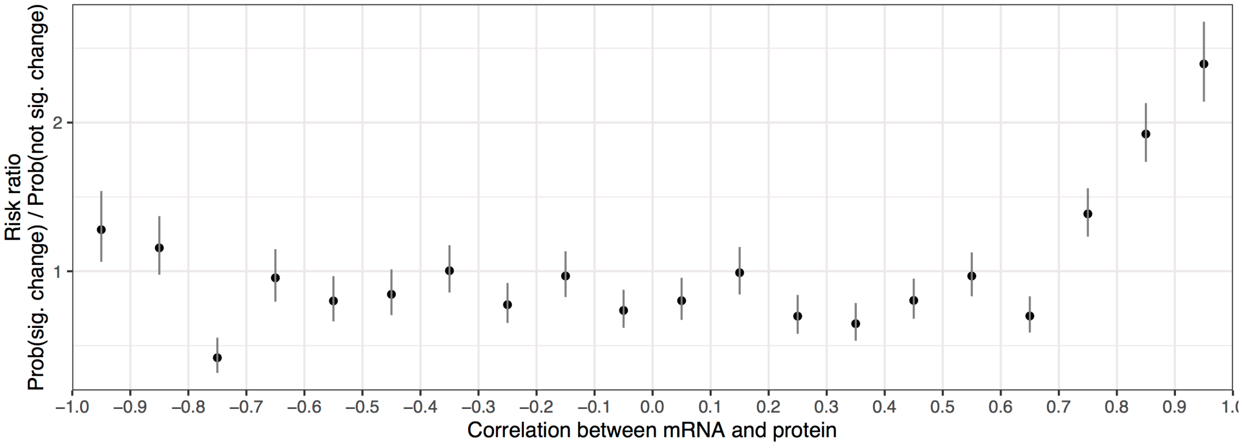


Supplemental Figure S2. Genes with both mRNA and protein significantly changing are enriched for higher correlations. The correlations for genes x strain combinations with both mRNA and protein significantly changing were distributed into bins of correlation ranging from -1 to 1. The same was done for gene x strain combinations where only one or neither of mRNA or protein was significantly changing. The fraction of significant gene x strain combinations in each bin was divided by the fraction of non-significant gene x strain combinations in each bin, making a risk ratio. The error bars mark standard errors of these risk ratios within each bin.

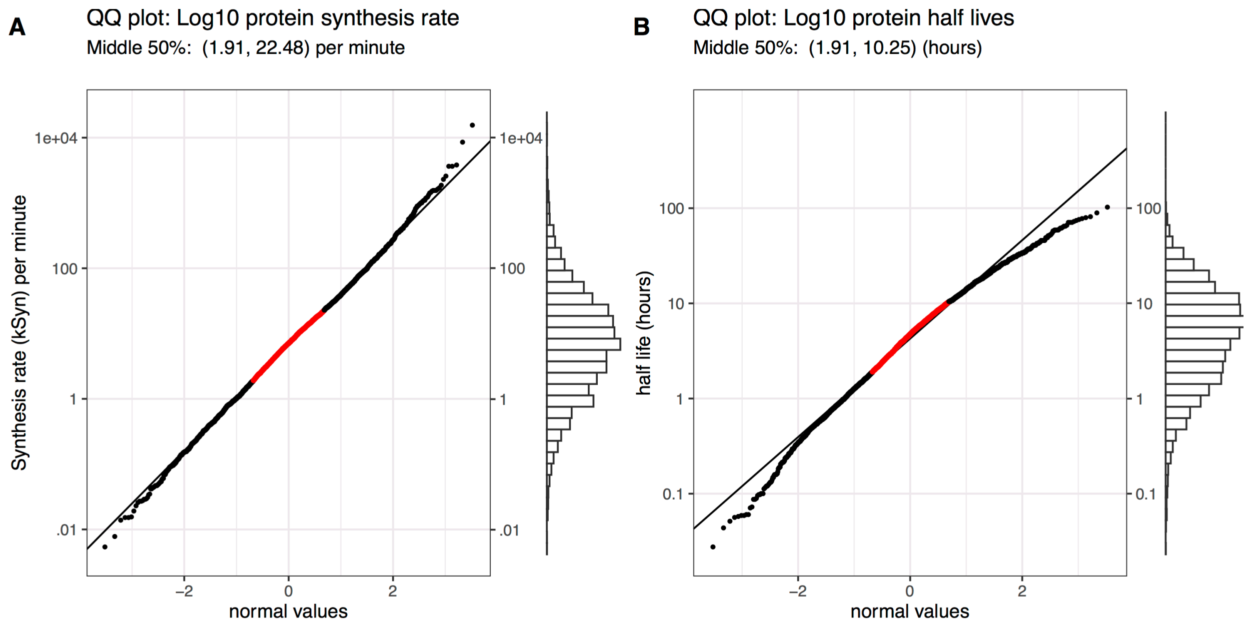


Supplemental Figure S3. QQ plots and histograms of the maximum *a posteriori* estimates of the $\log_{10}$ synthesis parameter **(A)** and the $\log_{10}$ half lives **(B)** against a normal distribution. Red stretches indicate the middle 50% of the values.
