## Supplemental Data S1 for "A simple mass-action model predicts genome-wide protein timecourses from mRNA trajectories during a dynamic response in two strains of *Saccharomyces cerevisiae*": GOCOMPONENT_par.html

Results


 **Parameters** 

*P-value color scale*

|  |  |  |  |  |
| --- | --- | --- | --- | --- |
| > 10^-3^ | 10^-3^ to 10^-5^ | 10^-5^ to 10^-7^ | 10^-7^ to 10^-9^ | < 10^-9^ |


|  |  |  |  |  |  |
| --- | --- | --- | --- | --- | --- |
| **GO term** | **Description** | **P-value** | **FDR q-value** | **Enrichment (N, B, n, b)** | **Genes** |
| GO:0005739 | mitochondrion | 1.48E-7 | 1.3E-4 | 1.54 (2298,517,358,124) | [+] Show genes  IDH1 - isocitrate dehydrogenase (nad(+)) idh1  RSP5 - nedd4 family e3 ubiquitin-protein ligase  MSC6 - msc6p  ENO2 - phosphopyruvate hydratase eno2  MOS1 - mos1p  MYO4 - myosin 4  GPD1 - glycerol-3-phosphate dehydrogenase (nad(+)) gpd1  YSA1 - ysa1p  ARP2 - actin-related protein 2  COA4 - coa4p  BAT2 - bat2p  RSM7 - rsm7p  RMD9 - rmd9p  FAA1 - long-chain fatty acid-coa ligase faa1  QCR2 - ubiquinol--cytochrome-c reductase subunit 2  CIT1 - citrate (si)-synthase cit1  ILV2 - acetolactate synthase catalytic subunit  CBR1 - cbr1p  SSA2 - hsp70 family chaperone ssa2  GLR1 - glutathione-disulfide reductase glr1  AIM18 - aim18p  RSM26 - rsm26p  MAS1 - mas1p  HFD1 - hfd1p  ECM33 - ecm33p  NFS1 - nfs1p  TCB1 - tcb1p  YTA12 - m-aaa protease subunit yta12  ATP2 - atp2p  HXK1 - hexokinase 1  ILV6 - acetolactate synthase regulatory subunit  VPS21 - vps21p  ATP4 - atp4p  RHO1 - rho1p  ARG5,6 - bifunctional acetylglutamate kinase/n-acetyl-gamma-glutamyl-phosphate reductase  CYM1 - cym1p  YDL119C - hypothetical protein  DCS1 - dcs1p  DIS3 - dis3p  KGD2 - alpha-ketoglutarate dehydrogenase kgd2  APE2 - ape2p  ATP7 - f1f0 atp synthase subunit d  ATP3 - atp3p  HSC82 - hsp90 family chaperone hsc82  TIM23 - tim23p  COX17 - cox17p  PEP4 - pep4p  MDH3 - malate dehydrogenase mdh3  YDL086W - carboxymethylenebutenolidase  YKT6 - ykt6p  ILV1 - threonine ammonia-lyase ilv1  NDI1 - nadh-ubiquinone reductase (h(+)-translocating) ndi1  MRPS12 - putative mitochondrial 37s ribosomal protein mrps12  SDH2 - succinate dehydrogenase iron-sulfur protein subunit sdh2  FCJ1 - fcj1p  YNL208W - hypothetical protein  NDE1 - nadh-ubiquinone reductase (h(+)-translocating) nde1  DNM1 - dnm1p  ALD5 - aldehyde dehydrogenase (nad(p)(+)) ald5  MRPL19 - mitochondrial 54s ribosomal protein yml19  TRX3 - trx3p  ODC2 - odc2p  ILV3 - ilv3p  ACH1 - ach1p  ERG6 - sterol 24-c-methyltransferase  KGD1 - alpha-ketoglutarate dehydrogenase kgd1  FMP52 - fmp52p  ILV5 - ketol-acid reductoisomerase  AYR1 - acylglycerone-phosphate reductase  ENO1 - phosphopyruvate hydratase eno1  GGC1 - ggc1p  GPX2 - glutathione peroxidase gpx2  YDR341C - arginine--trna ligase  COQ1 - trans-hexaprenyltranstransferase  MDH1 - malate dehydrogenase mdh1  ISC1 - inositol phosphosphingolipid phospholipase  FUM1 - fumarase fum1  AFG3 - aaa family atpase afg3  TUF1 - tuf1p  HTS1 - histidine--trna ligase  MAE1 - malate dehydrogenase (oxaloacetate-decarboxylating)  COR1 - ubiquinol--cytochrome-c reductase subunit cor1  GDH2 - glutamate dehydrogenase (nad(+))  ATP15 - f1f0 atp synthase subunit epsilon  UGA1 - 4-aminobutyrate transaminase  GUT2 - glycerol-3-phosphate dehydrogenase  PHB2 - phb2p  PET9 - pet9p  ARG7 - glutamate n-acetyltransferase  MAS2 - mas2p  PDR5 - atp-binding cassette multidrug transporter pdr5  DLD3 - dld3p  SLT2 - slt2p  MRH1 - mrh1p  ATP5 - atp5p  RAS2 - ras2p  LSC1 - succinate--coa ligase (gdp-forming) subunit alpha  HXK2 - hexokinase 2  RPO21 - rpo21p  ARO3 - 3-deoxy-7-phosphoheptulonate synthase aro3  EHD3 - ehd3p  IDH2 - isocitrate dehydrogenase (nad(+)) idh2  CCA1 - cca1p  CMC2 - cmc2p  MCR1 - mcr1p  NEW1 - new1p  EHT1 - eht1p  SDH1 - succinate dehydrogenase flavoprotein subunit sdh1  PRX1 - prx1p  ARC18 - arc18p  ALD4 - aldehyde dehydrogenase (nadp(+)) ald4  GLT1 - glutamate synthase (nadh)  DLD1 - dld1p  GCN1 - gcn1p  AIM17 - aim17p  RIP1 - ubiquinol--cytochrome-c reductase catalytic subunit rip1  DLD2 - dld2p  PIL1 - pil1p  OXR1 - oxr1p  ATP1 - f1f0 atp synthase subunit alpha  LSP1 - lsp1p  TPS2 - trehalose-phosphatase tps2  PTP1 - ptp1p  ZEO1 - zeo1p |
| GO:0044429 | mitochondrial part | 1.59E-5 | 7E-3 | 1.64 (2298,297,358,76) | [+] Show genes  IDH1 - isocitrate dehydrogenase (nad(+)) idh1  MRPL19 - mitochondrial 54s ribosomal protein yml19  MSC6 - msc6p  ODC2 - odc2p  ERG6 - sterol 24-c-methyltransferase  MOS1 - mos1p  KGD1 - alpha-ketoglutarate dehydrogenase kgd1  FMP52 - fmp52p  ILV5 - ketol-acid reductoisomerase  AYR1 - acylglycerone-phosphate reductase  GGC1 - ggc1p  GPX2 - glutathione peroxidase gpx2  COA4 - coa4p  COQ1 - trans-hexaprenyltranstransferase  MDH1 - malate dehydrogenase mdh1  ISC1 - inositol phosphosphingolipid phospholipase  RSM7 - rsm7p  FUM1 - fumarase fum1  PPX1 - ppx1p  AFG3 - aaa family atpase afg3  RMD9 - rmd9p  FAA1 - long-chain fatty acid-coa ligase faa1  MAE1 - malate dehydrogenase (oxaloacetate-decarboxylating)  QCR2 - ubiquinol--cytochrome-c reductase subunit 2  CIT1 - citrate (si)-synthase cit1  CBR1 - cbr1p  COR1 - ubiquinol--cytochrome-c reductase subunit cor1  RSM26 - rsm26p  MAS1 - mas1p  ATP15 - f1f0 atp synthase subunit epsilon  HFD1 - hfd1p  RIB3 - 3,4-dihydroxy-2-butanone-4-phosphate synthase rib3  YTA12 - m-aaa protease subunit yta12  GUT2 - glycerol-3-phosphate dehydrogenase  PHB2 - phb2p  ATP2 - atp2p  PET9 - pet9p  ILV6 - acetolactate synthase regulatory subunit  VPS21 - vps21p  RHO1 - rho1p  ATP4 - atp4p  ARG5,6 - bifunctional acetylglutamate kinase/n-acetyl-gamma-glutamyl-phosphate reductase  ARG7 - glutamate n-acetyltransferase  MAS2 - mas2p  CYM1 - cym1p  YDL119C - hypothetical protein  ATP5 - atp5p  KGD2 - alpha-ketoglutarate dehydrogenase kgd2  LSC1 - succinate--coa ligase (gdp-forming) subunit alpha  ATP7 - f1f0 atp synthase subunit d  EHD3 - ehd3p  IDH2 - isocitrate dehydrogenase (nad(+)) idh2  CMC2 - cmc2p  CCA1 - cca1p  ARC15 - arc15p  MCR1 - mcr1p  ATP3 - atp3p  TIM23 - tim23p  EHT1 - eht1p  COX17 - cox17p  SDH1 - succinate dehydrogenase flavoprotein subunit sdh1  ALD4 - aldehyde dehydrogenase (nadp(+)) ald4  DLD1 - dld1p  NDI1 - nadh-ubiquinone reductase (h(+)-translocating) ndi1  MRPS12 - putative mitochondrial 37s ribosomal protein mrps12  SDH2 - succinate dehydrogenase iron-sulfur protein subunit sdh2  RIP1 - ubiquinol--cytochrome-c reductase catalytic subunit rip1  FCJ1 - fcj1p  DLD2 - dld2p  PIL1 - pil1p  NDE1 - nadh-ubiquinone reductase (h(+)-translocating) nde1  ATP1 - f1f0 atp synthase subunit alpha  LSP1 - lsp1p  DNM1 - dnm1p  ZEO1 - zeo1p  ALD5 - aldehyde dehydrogenase (nad(p)(+)) ald5 |
| GO:0005743 | mitochondrial inner membrane | 3.72E-5 | 1.09E-2 | 2.26 (2298,91,358,32) | [+] Show genes  MAS2 - mas2p  ODC2 - odc2p  MOS1 - mos1p  YDL119C - hypothetical protein  ATP5 - atp5p  ATP7 - f1f0 atp synthase subunit d  GPX2 - glutathione peroxidase gpx2  GGC1 - ggc1p  CMC2 - cmc2p  COA4 - coa4p  COQ1 - trans-hexaprenyltranstransferase  TIM23 - tim23p  ATP3 - atp3p  SDH1 - succinate dehydrogenase flavoprotein subunit sdh1  RMD9 - rmd9p  AFG3 - aaa family atpase afg3  QCR2 - ubiquinol--cytochrome-c reductase subunit 2  DLD1 - dld1p  COR1 - ubiquinol--cytochrome-c reductase subunit cor1  MAS1 - mas1p  ATP15 - f1f0 atp synthase subunit epsilon  NDI1 - nadh-ubiquinone reductase (h(+)-translocating) ndi1  SDH2 - succinate dehydrogenase iron-sulfur protein subunit sdh2  FCJ1 - fcj1p  RIP1 - ubiquinol--cytochrome-c reductase catalytic subunit rip1  YTA12 - m-aaa protease subunit yta12  GUT2 - glycerol-3-phosphate dehydrogenase  ATP1 - f1f0 atp synthase subunit alpha  PHB2 - phb2p  ATP2 - atp2p  PET9 - pet9p  ATP4 - atp4p |
| GO:0019866 | organelle inner membrane | 4.68E-5 | 1.03E-2 | 2.21 (2298,96,358,33) | [+] Show genes  MAS2 - mas2p  ODC2 - odc2p  MOS1 - mos1p  YDL119C - hypothetical protein  ATP5 - atp5p  ATP7 - f1f0 atp synthase subunit d  GGC1 - ggc1p  GPX2 - glutathione peroxidase gpx2  CMC2 - cmc2p  COQ1 - trans-hexaprenyltranstransferase  COA4 - coa4p  TIM23 - tim23p  ATP3 - atp3p  SDH1 - succinate dehydrogenase flavoprotein subunit sdh1  RMD9 - rmd9p  AFG3 - aaa family atpase afg3  QCR2 - ubiquinol--cytochrome-c reductase subunit 2  DLD1 - dld1p  COR1 - ubiquinol--cytochrome-c reductase subunit cor1  MAS1 - mas1p  ATP15 - f1f0 atp synthase subunit epsilon  NDI1 - nadh-ubiquinone reductase (h(+)-translocating) ndi1  SDH2 - succinate dehydrogenase iron-sulfur protein subunit sdh2  RIP1 - ubiquinol--cytochrome-c reductase catalytic subunit rip1  FCJ1 - fcj1p  SSM4 - e3 ubiquitin-protein ligase ssm4  YTA12 - m-aaa protease subunit yta12  GUT2 - glycerol-3-phosphate dehydrogenase  ATP1 - f1f0 atp synthase subunit alpha  PHB2 - phb2p  ATP2 - atp2p  PET9 - pet9p  ATP4 - atp4p |
| GO:0005829 | cytosol | 1.1E-4 | 1.93E-2 | 1.31 (2298,417,684,162) | [+] Show genes  TPS1 - alpha,alpha-trehalose-phosphate synthase (udp-forming) tps1  NMD3 - nmd3p  FUN12 - fun12p  ENO2 - phosphopyruvate hydratase eno2  TSL1 - tsl1p  ARO2 - bifunctional chorismate synthase/riboflavin reductase [nad(p)h] aro2  ALD6 - aldehyde dehydrogenase (nadp(+)) ald6  URM1 - ubiquitin-related modifier urm1  BET3 - bet3p  GPD1 - glycerol-3-phosphate dehydrogenase (nad(+)) gpd1  ASC1 - asc1p  ASN2 - asparagine synthase (glutamine-hydrolyzing) 2  YSA1 - ysa1p  CKA2 - cka2p  GCD6 - gcd6p  GLK1 - glucokinase  FES1 - fes1p  CPR6 - peptidylprolyl isomerase cpr6  EMI2 - putative glucokinase  GND1 - phosphogluconate dehydrogenase (decarboxylating) gnd1  PPX1 - ppx1p  SIS1 - sis1p  BRO1 - bro1p  RFC3 - replication factor c subunit 3  FRA1 - fra1p  PGM2 - phosphoglucomutase pgm2  URA8 - ura8p  SSA2 - hsp70 family chaperone ssa2  GLR1 - glutathione-disulfide reductase glr1  SNF4 - snf4p  RIB3 - 3,4-dihydroxy-2-butanone-4-phosphate synthase rib3  NFS1 - nfs1p  LEU2 - 3-isopropylmalate dehydrogenase  BCY1 - bcy1p  SEC18 - sec18p  PTI1 - pti1p  MUK1 - muk1p  RET2 - ret2p  FBP26 - fbp26p  RPN12 - proteasome regulatory particle lid subunit rpn12  CAR2 - ornithine-oxo-acid transaminase  HXK1 - hexokinase 1  THI4 - thi4p  PBI2 - pbi2p  VPS35 - vps35p  HSP150 - hsp150p  VPS21 - vps21p  NUP133 - nup133p  PRP19 - e3 ubiquitin-protein ligase prp19  RPS1B - ribosomal 40s subunit protein s1b  DSL1 - dsl1p  BUD17 - putative pyridoxal kinase bud17  DAK1 - dak1p  YDL124W - aldo-keto reductase superfamily protein  GRE3 - trifunctional aldehyde reductase/xylose reductase/glucose 1-dehydrogenase (nadp(+))  OSH6 - oxysterol-binding protein osh6  RVS161 - rvs161p  RPN9 - proteasome regulatory particle lid subunit rpn9  KAP120 - kap120p  ASN1 - asparagine synthase (glutamine-hydrolyzing) 1  SSE1 - sse1p  YRB2 - yrb2p  GUK1 - guanylate kinase  DPS1 - aspartate--trna ligase dps1  GET3 - guanine nucleotide exchange factor get3  HSC82 - hsp90 family chaperone hsc82  PDR16 - pdr16p  COX17 - cox17p  COG8 - cog8p  RHO3 - rho3p  HEM2 - porphobilinogen synthase hem2  SLY1 - sly1p  GCD1 - gcd1p  YKT6 - ykt6p  SGT2 - sgt2p  YJL068C - hypothetical protein  CAB5 - putative dephospho-coa kinase  ARC35 - arc35p  SPP1 - spp1p  TRX2 - trx2p  RTS1 - rts1p  HMO1 - hmo1p  KES1 - kes1p  DNM1 - dnm1p  BRE5 - bre5p  HSP82 - hsp90 family chaperone hsp82  YPT7 - ypt7p  MVP1 - mvp1p  CHS5 - chs5p  SRP54 - srp54p  ACH1 - ach1p  NPT1 - nicotinate phosphoribosyltransferase  PDC1 - indolepyruvate decarboxylase 1  TRM732 - trm732p  TRP4 - anthranilate phosphoribosyltransferase  TRX1 - trx1p  GRX3 - grx3p  GAD1 - glutamate decarboxylase gad1  ENO1 - phosphopyruvate hydratase eno1  HSM3 - hsm3p  LEU1 - 3-isopropylmalate dehydratase leu1  GPX2 - glutathione peroxidase gpx2  YLR345W - bifunctional fructose-2,6-bisphosphate 2-phosphatase/6-phosphofructo-2-kinase  YDR341C - arginine--trna ligase  CPR7 - cpr7p  ADE16 - bifunctional phosphoribosylaminoimidazolecarboxamide formyltransferase/imp cyclohydrolase ade16  FUM1 - fumarase fum1  ACO1 - aconitate hydratase aco1  YAP1802 - yap1802p  PEP5 - pep5p  TRS85 - trs85p  HTS1 - histidine--trna ligase  ARG1 - argininosuccinate synthase  PRE7 - proteasome core particle subunit beta 6  UGA1 - 4-aminobutyrate transaminase  AMD1 - amp deaminase  GLN4 - glutamine--trna ligase  CSE1 - cse1p  MEF1 - mef1p  IMH1 - imh1p  VPS74 - vps74p  CDC25 - cdc25p  NAS6 - nas6p  ADE5,7 - bifunctional aminoimidazole ribotide synthase/glycinamide ribotide synthase  ERG12 - mevalonate kinase  SDS24 - sds24p  NCP1 - ncp1p  PGM1 - phosphoglucomutase pgm1  ACB1 - long-chain fatty acid transporter acb1  GRX4 - grx4p  YPK1 - ypk1p  CDC11 - cdc11p  APL2 - apl2p  SLT2 - slt2p  RDI1 - rdi1p  CHC1 - chc1p  LSC1 - succinate--coa ligase (gdp-forming) subunit alpha  HSP12 - hsp12p  HXK2 - hexokinase 2  SSE2 - sse2p  SSA3 - hsp70 family atpase ssa3  YMR074C - hypothetical protein  KRS1 - lysine--trna ligase krs1  GPM1 - phosphoglycerate mutase gpm1  DED81 - asparagine--trna ligase ded81  HEM13 - coproporphyrinogen oxidase  ACO2 - aco2p  CAB1 - pantothenate kinase  MET12 - methylenetetrahydrofolate reductase (nad(p)h) met12  TRS33 - trs33p  BLM10 - blm10p  GCN1 - gcn1p  YDJ1 - ydj1p  GEA2 - gea2p  PDC5 - indolepyruvate decarboxylase 5  VAC8 - vac8p  ARA1 - d-arabinose 1-dehydrogenase (nad(p)(+)) ara1  ARA2 - d-arabinose 1-dehydrogenase (nad(p)(+)) ara2  SSB1 - hsp70 family atpase ssb1  TPS2 - trehalose-phosphatase tps2  CCS1 - ccs1p  CDC55 - cdc55p |
| GO:0098800 | inner mitochondrial membrane protein complex | 1.22E-4 | 1.79E-2 | 2.95 (2298,37,358,17) | [+] Show genes  AFG3 - aaa family atpase afg3  QCR2 - ubiquinol--cytochrome-c reductase subunit 2  COR1 - ubiquinol--cytochrome-c reductase subunit cor1  MOS1 - mos1p  ATP5 - atp5p  ATP15 - f1f0 atp synthase subunit epsilon  SDH2 - succinate dehydrogenase iron-sulfur protein subunit sdh2  RIP1 - ubiquinol--cytochrome-c reductase catalytic subunit rip1  FCJ1 - fcj1p  YTA12 - m-aaa protease subunit yta12  ATP7 - f1f0 atp synthase subunit d  ATP2 - atp2p  ATP1 - f1f0 atp synthase subunit alpha  TIM23 - tim23p  ATP3 - atp3p  ATP4 - atp4p  SDH1 - succinate dehydrogenase flavoprotein subunit sdh1 |
| GO:0031966 | mitochondrial membrane | 2.55E-4 | 3.2E-2 | 1.81 (2298,163,358,46) | [+] Show genes  MAS2 - mas2p  ODC2 - odc2p  ERG6 - sterol 24-c-methyltransferase  MOS1 - mos1p  YDL119C - hypothetical protein  ATP5 - atp5p  FMP52 - fmp52p  AYR1 - acylglycerone-phosphate reductase  ATP7 - f1f0 atp synthase subunit d  GGC1 - ggc1p  GPX2 - glutathione peroxidase gpx2  CMC2 - cmc2p  COQ1 - trans-hexaprenyltranstransferase  COA4 - coa4p  MCR1 - mcr1p  ATP3 - atp3p  TIM23 - tim23p  EHT1 - eht1p  SDH1 - succinate dehydrogenase flavoprotein subunit sdh1  AFG3 - aaa family atpase afg3  RMD9 - rmd9p  FAA1 - long-chain fatty acid-coa ligase faa1  QCR2 - ubiquinol--cytochrome-c reductase subunit 2  CBR1 - cbr1p  DLD1 - dld1p  COR1 - ubiquinol--cytochrome-c reductase subunit cor1  ATP15 - f1f0 atp synthase subunit epsilon  MAS1 - mas1p  HFD1 - hfd1p  NDI1 - nadh-ubiquinone reductase (h(+)-translocating) ndi1  SDH2 - succinate dehydrogenase iron-sulfur protein subunit sdh2  RIP1 - ubiquinol--cytochrome-c reductase catalytic subunit rip1  YTA12 - m-aaa protease subunit yta12  FCJ1 - fcj1p  PIL1 - pil1p  GUT2 - glycerol-3-phosphate dehydrogenase  ATP1 - f1f0 atp synthase subunit alpha  PHB2 - phb2p  ATP2 - atp2p  PET9 - pet9p  LSP1 - lsp1p  DNM1 - dnm1p  VPS21 - vps21p  ZEO1 - zeo1p  RHO1 - rho1p  ATP4 - atp4p |
| GO:0048471 | perinuclear region of cytoplasm | 3.42E-4 | 3.76E-2 | 11.40 (2298,14,72,5) | [+] Show genes  HSP82 - hsp90 family chaperone hsp82  YDJ1 - ydj1p  DCS2 - dcs2p  DCS1 - dcs1p  HSC82 - hsp90 family chaperone hsc82 |
| GO:0045239 | tricarboxylic acid cycle enzyme complex | 4.54E-4 | 4.44E-2 | 3.25 (2298,8,707,8) | [+] Show genes  KGD1 - alpha-ketoglutarate dehydrogenase kgd1  IDH1 - isocitrate dehydrogenase (nad(+)) idh1  FUM1 - fumarase fum1  KGD2 - alpha-ketoglutarate dehydrogenase kgd2  YMR31 - mitochondrial 37s ribosomal protein ymr31  LPD1 - dihydrolipoyl dehydrogenase  IDH2 - isocitrate dehydrogenase (nad(+)) idh2  LSC2 - succinate--coa ligase (gdp-forming) subunit beta |
| GO:0044444 | cytoplasmic part | 5.55E-4 | 4.88E-2 | 1.09 (2298,1502,735,523) | [+] Show genes  YKR070W - hypothetical protein  ARO2 - bifunctional chorismate synthase/riboflavin reductase [nad(p)h] aro2  GOS1 - gos1p  RPL9A - ribosomal 60s subunit protein l9a  MOS1 - mos1p  BET3 - bet3p  GPD1 - glycerol-3-phosphate dehydrogenase (nad(+)) gpd1  ASC1 - asc1p  FMP46 - fmp46p  EMC1 - emc1p  GLK1 - glucokinase  ARP2 - actin-related protein 2  SRO9 - sro9p  MXR2 - mxr2p  ATP16 - f1f0 atp synthase subunit delta  RMD9 - rmd9p  SEH1 - seh1p  PGM2 - phosphoglucomutase pgm2  PTR2 - ptr2p  QCR2 - ubiquinol--cytochrome-c reductase subunit 2  ILV2 - acetolactate synthase catalytic subunit  SNF4 - snf4p  MYO5 - myosin 5  HFD1 - hfd1p  NFS1 - nfs1p  SEC23 - sec23p  LEU2 - 3-isopropylmalate dehydrogenase  RPS2 - ribosomal 40s subunit protein s2  YTA12 - m-aaa protease subunit yta12  ILV6 - acetolactate synthase regulatory subunit  NUP133 - nup133p  RHO1 - rho1p  CAP1 - cap1p  KRE6 - kre6p  CDC10 - septin cdc10  YCP4 - ycp4p  YDL124W - aldo-keto reductase superfamily protein  OSH6 - oxysterol-binding protein osh6  RVS161 - rvs161p  YDL119C - hypothetical protein  ASN1 - asparagine synthase (glutamine-hydrolyzing) 1  GUP1 - gup1p  CWH43 - cwh43p  PIN3 - pin3p  ATP7 - f1f0 atp synthase subunit d  ARC19 - arc19p  UFD4 - putative ubiquitin-protein ligase ufd4  SSO2 - sso2p  HSC82 - hsp90 family chaperone hsc82  GET3 - guanine nucleotide exchange factor get3  RRP43 - rrp43p  OCH1 - och1p  PNC1 - nicotinamidase  RPL31A - ribosomal 60s subunit protein l31a  MDH3 - malate dehydrogenase mdh3  HEM2 - porphobilinogen synthase hem2  PAI3 - pai3p  YDL086W - carboxymethylenebutenolidase  FCJ1 - fcj1p  OLE1 - stearoyl-coa 9-desaturase  NDE1 - nadh-ubiquinone reductase (h(+)-translocating) nde1  YET3 - yet3p  YPR114W - hypothetical protein  RSN1 - rsn1p  TRX3 - trx3p  YDR089W - hypothetical protein  TIF11 - tif11p  TRM732 - trm732p  FAA4 - long-chain fatty acid-coa ligase faa4  GRX3 - grx3p  GAD1 - glutamate decarboxylase gad1  ERG7 - lanosterol synthase erg7  HSM3 - hsm3p  YHM2 - yhm2p  MDH1 - malate dehydrogenase mdh1  PEP5 - pep5p  TRS85 - trs85p  COX6 - cytochrome c oxidase subunit vi  MAE1 - malate dehydrogenase (oxaloacetate-decarboxylating)  TRM1 - trm1p  UGP1 - utp glucose-1-phosphate uridylyltransferase  PTM1 - ptm1p  SAC6 - sac6p  EMP47 - emp47p  GAS3 - gas3p  MSC7 - msc7p  CSE1 - cse1p  ADE5,7 - bifunctional aminoimidazole ribotide synthase/glycinamide ribotide synthase  YCF1 - atp-binding cassette glutathione s-conjugate transporter ycf1  ERG12 - mevalonate kinase  SCJ1 - scj1p  NCP1 - ncp1p  GYP8 - gyp8p  MYO3 - myosin 3  PGM1 - phosphoglucomutase pgm1  GCV1 - glycine decarboxylase subunit t  MAS2 - mas2p  YPK1 - ypk1p  MYO1 - myosin 1  RPL22B - ribosomal 60s subunit protein l22b  SLT2 - slt2p  APL2 - apl2p  CHC1 - chc1p  ACT1 - actin  OCT1 - oct1p  MRH1 - mrh1p  SOD2 - superoxide dismutase sod2  MDJ1 - mdj1p  HSP12 - hsp12p  HXK2 - hexokinase 2  ARO3 - 3-deoxy-7-phosphoheptulonate synthase aro3  EHD3 - ehd3p  LPD1 - dihydrolipoyl dehydrogenase  KRS1 - lysine--trna ligase krs1  GNA1 - glucosamine 6-phosphate n-acetyltransferase  GPM1 - phosphoglycerate mutase gpm1  YME2 - yme2p  MCR1 - mcr1p  DED81 - asparagine--trna ligase ded81  SDH1 - succinate dehydrogenase flavoprotein subunit sdh1  HEM13 - coproporphyrinogen oxidase  APE1 - ape1p  BLM10 - blm10p  TIM10 - tim10p  GCN1 - gcn1p  LCB2 - serine c-palmitoyltransferase lcb2  AIM7 - aim7p  AIM17 - aim17p  RPS20 - ribosomal 40s subunit protein s20  RPS13 - ribosomal 40s subunit protein s13  GCN20 - putative aaa family atpase gcn20  TPS2 - trehalose-phosphatase tps2  CDC55 - cdc55p  RPL29 - ribosomal 60s subunit protein l29  TSL1 - tsl1p  IVY1 - ivy1p  ALD6 - aldehyde dehydrogenase (nadp(+)) ald6  ASN2 - asparagine synthase (glutamine-hydrolyzing) 2  SPT6 - spt6p  CKA2 - cka2p  GCD6 - gcd6p  TUB1 - tub1p  CPR3 - peptidylprolyl isomerase cpr3  SEC16 - sec16p  BRO1 - bro1p  FRA1 - fra1p  RPL3 - ribosomal 60s subunit protein l3  HSP104 - chaperone atpase hsp104  SSA2 - hsp70 family chaperone ssa2  GLR1 - glutathione-disulfide reductase glr1  PHB1 - phb1p  PNG1 - png1p  OST3 - ost3p  TCB1 - tcb1p  CYS4 - cystathionine beta-synthase cys4  YMR31 - mitochondrial 37s ribosomal protein ymr31  PTI1 - pti1p  MUK1 - muk1p  RET2 - ret2p  RPN12 - proteasome regulatory particle lid subunit rpn12  LYS1 - saccharopine dehydrogenase (nad+, l-lysine-forming)  HXK1 - hexokinase 1  THI4 - thi4p  VPS13 - vps13p  RPL24B - ribosomal 60s subunit protein l24b  VPS21 - vps21p  PRP19 - e3 ubiquitin-protein ligase prp19  YGR149W - hypothetical protein  ATP4 - atp4p  PTC5 - ptc5p  RPS1B - ribosomal 40s subunit protein s1b  MRPS35 - mitochondrial 37s ribosomal protein mrps35  GAS5 - gas5p  DAK1 - dak1p  KAP120 - kap120p  TIM13 - tim13p  SSE1 - sse1p  DIS3 - dis3p  KGD2 - alpha-ketoglutarate dehydrogenase kgd2  RNR4 - ribonucleotide-diphosphate reductase subunit rnr4  DPS1 - aspartate--trna ligase dps1  COX17 - cox17p  RPL6A - ribosomal 60s subunit protein l6a  COG8 - cog8p  PEP4 - pep4p  HMG1 - hydroxymethylglutaryl-coa reductase (nadph) hmg1  SLY1 - sly1p  SGT2 - sgt2p  CAB5 - putative dephospho-coa kinase  NDI1 - nadh-ubiquinone reductase (h(+)-translocating) ndi1  SPP1 - spp1p  SDH2 - succinate dehydrogenase iron-sulfur protein subunit sdh2  TRX2 - trx2p  YBT1 - bile acid-transporting atpase ybt1  RTS1 - rts1p  RPL5 - ribosomal 60s subunit protein l5  HMO1 - hmo1p  KES1 - kes1p  DNM1 - dnm1p  YPT7 - ypt7p  RPL24A - ribosomal 60s subunit protein l24a  RPL30 - ribosomal 60s subunit protein l30  MVP1 - mvp1p  CWH41 - cwh41p  MRP1 - mitochondrial 37s ribosomal protein mrp1  SRP54 - srp54p  ERG6 - sterol 24-c-methyltransferase  PDC1 - indolepyruvate decarboxylase 1  TRX1 - trx1p  TRP4 - anthranilate phosphoribosyltransferase  ERG4 - delta(24(24(1)))-sterol reductase  LEU1 - 3-isopropylmalate dehydratase leu1  YDR341C - arginine--trna ligase  VMA9 - vma9p  RPL15A - ribosomal 60s subunit protein l15a  ERG26 - sterol-4-alpha-carboxylate 3-dehydrogenase (decarboxylating)  ADE16 - bifunctional phosphoribosylaminoimidazolecarboxamide formyltransferase/imp cyclohydrolase ade16  HXT2 - hxt2p  HTS1 - histidine--trna ligase  ARG1 - argininosuccinate synthase  LSM6 - lsm6p  BUD20 - bud20p  RPL10 - ribosomal 60s subunit protein l10  UGA1 - 4-aminobutyrate transaminase  AMD1 - amp deaminase  YMC1 - ymc1p  RPS15 - ribosomal 40s subunit protein s15  RPP2A - ribosomal protein p2a  MEF1 - mef1p  VPS74 - vps74p  ERG3 - c-5 sterol desaturase  REX2 - rex2p  RPL22A - ribosomal 60s subunit protein l22a  AIM45 - aim45p  ARG7 - glutamate n-acetyltransferase  ACB1 - long-chain fatty acid transporter acb1  RPL26B - ribosomal 60s subunit protein l26b  MHR1 - mhr1p  DPL1 - sphinganine-1-phosphate aldolase dpl1  PRO1 - glutamate 5-kinase  ATP5 - atp5p  YGR054W - hypothetical protein  YME1 - yme1p  YMR074C - hypothetical protein  RRP45 - rrp45p  NYV1 - nyv1p  DPP1 - bifunctional diacylglycerol diphosphate phospatase/phosphatidate phosphatase  XDJ1 - xdj1p  ERG10 - acetyl-coa c-acetyltransferase  STM1 - stm1p  MET12 - methylenetetrahydrofolate reductase (nad(p)h) met12  TWF1 - twf1p  VTC3 - vtc3p  CPR5 - peptidylprolyl isomerase family protein cpr5  RRP12 - rrp12p  TAE2 - tae2p  PIL1 - pil1p  PDC5 - indolepyruvate decarboxylase 5  RRP46 - rrp46p  ARA2 - d-arabinose 1-dehydrogenase (nad(p)(+)) ara2  LSP1 - lsp1p  TFB1 - tfb1p  CCS1 - ccs1p  ZEO1 - zeo1p  IDH1 - isocitrate dehydrogenase (nad(+)) idh1  NMT1 - nmt1p  PYK2 - pyruvate kinase pyk2  PHS1 - phs1p  HCR1 - hcr1p  RSP5 - nedd4 family e3 ubiquitin-protein ligase  MSC6 - msc6p  CIR2 - cir2p  MSS51 - mss51p  COA4 - coa4p  CPR6 - peptidylprolyl isomerase cpr6  EMI2 - putative glucokinase  MTC1 - mtc1p  PBS2 - pbs2p  SIS1 - sis1p  GCD14 - gcd14p  FAA1 - long-chain fatty acid-coa ligase faa1  YDR476C - hypothetical protein  CIT1 - citrate (si)-synthase cit1  MRPL28 - mitochondrial 54s ribosomal protein yml28  RPS31 - ubiquitin-ribosomal 40s subunit protein s31 fusion protein  MYO2 - myosin 2  MAS1 - mas1p  RIB3 - 3,4-dihydroxy-2-butanone-4-phosphate synthase rib3  TFS1 - tfs1p  CPA1 - carbamoyl-phosphate synthase (glutamine-hydrolyzing) cpa1  COX15 - cox15p  GLC7 - glc7p  FBP26 - fbp26p  VPS35 - vps35p  PBI2 - pbi2p  HSP150 - hsp150p  ARG5,6 - bifunctional acetylglutamate kinase/n-acetyl-gamma-glutamyl-phosphate reductase  MPD1 - protein disulfide isomerase mpd1  KAR2 - hsp70 family atpase kar2  BUD17 - putative pyridoxal kinase bud17  GPI17 - gpi17p  CYM1 - cym1p  RPN9 - proteasome regulatory particle lid subunit rpn9  DCS1 - dcs1p  FCY2 - fcy2p  GUK1 - guanylate kinase  LRO1 - phospholipid:diacylglycerol acyltransferase  TIM23 - tim23p  RAD51 - recombinase rad51  TUM1 - tum1p  RPB7 - rpb7p  RPT4 - proteasome regulatory particle base subunit rpt4  CDC42 - cdc42p  RPT3 - proteasome regulatory particle base subunit rpt3  GCD1 - gcd1p  TRP2 - anthranilate synthase trp2  RPN8 - proteasome regulatory particle lid subunit rpn8  ILV1 - threonine ammonia-lyase ilv1  YJL068C - hypothetical protein  MRPS12 - putative mitochondrial 37s ribosomal protein mrps12  ARC35 - arc35p  RPL33B - ribosomal 60s subunit protein l33b  SCP160 - scp160p  MET7 - tetrahydrofolate synthase  TRM112 - trm112p  SSP120 - ssp120p  ABP140 - abp140p  BRE5 - bre5p  ALD5 - aldehyde dehydrogenase (nad(p)(+)) ald5  HSP82 - hsp90 family chaperone hsp82  ODC2 - odc2p  CHS5 - chs5p  ACH1 - ach1p  NPT1 - nicotinate phosphoribosyltransferase  BGL2 - bgl2p  RPL38 - ribosomal 60s subunit protein l38  SCW4 - scw4p  ILV5 - ketol-acid reductoisomerase  ENO1 - phosphopyruvate hydratase eno1  YLR345W - bifunctional fructose-2,6-bisphosphate 2-phosphatase/6-phosphofructo-2-kinase  RPL26A - ribosomal 60s subunit protein l26a  COQ1 - trans-hexaprenyltranstransferase  FKS1 - fks1p  RPP0 - ribosomal protein p0  FUM1 - fumarase fum1  ACO1 - aconitate hydratase aco1  TUF1 - tuf1p  YAP1802 - yap1802p  EDE1 - ede1p  COR1 - ubiquinol--cytochrome-c reductase subunit cor1  LSC2 - succinate--coa ligase (gdp-forming) subunit beta  ATP15 - f1f0 atp synthase subunit epsilon  PRE7 - proteasome core particle subunit beta 6  GLN4 - glutamine--trna ligase  BUD6 - bud6p  DCS2 - dcs2p  CDC3 - septin cdc3  CRM1 - crm1p  HEK2 - hek2p  PHB2 - phb2p  IMH1 - imh1p  PET9 - pet9p  CDC25 - cdc25p  NAS6 - nas6p  MRP21 - mitochondrial 37s ribosomal protein mrp21  RPL17B - rpl17bp  PGC1 - pgc1p  GRX4 - grx4p  RPL7B - ribosomal 60s subunit protein l7b  RPL32 - ribosomal 60s subunit protein l32  PDR5 - atp-binding cassette multidrug transporter pdr5  RPB2 - rpb2p  LSC1 - succinate--coa ligase (gdp-forming) subunit alpha  AST1 - ast1p  CBP3 - cbp3p  SSA3 - hsp70 family atpase ssa3  RPS22A - rps22ap  IDH2 - isocitrate dehydrogenase (nad(+)) idh2  CMC2 - cmc2p  CCA1 - cca1p  ATG27 - atg27p  YLR413W - hypothetical protein  NEW1 - new1p  PRX1 - prx1p  DAP1 - dap1p  RPS22B - ribosomal 40s subunit protein s22b  ACO2 - aco2p  ARC18 - arc18p  SSQ1 - hsp70 family atpase ssq1  CAB1 - pantothenate kinase  TRS33 - trs33p  POS5 - pos5p  INP53 - phosphatidylinositol-3-/phosphoinositide 5-phosphatase inp53  YCR075W-A - hypothetical protein  MRN1 - mrn1p  MIC23 - mic23p  ATP1 - f1f0 atp synthase subunit alpha  KTR1 - ktr1p  OXR1 - oxr1p  BNA4 - kynurenine 3-monooxygenase  RPS7A - ribosomal 40s subunit protein s7a  TPS1 - alpha,alpha-trehalose-phosphate synthase (udp-forming) tps1  MGM101 - mgm101p  NMD3 - nmd3p  FUN12 - fun12p  ENO2 - phosphopyruvate hydratase eno2  ATP11 - atp11p  MET5 - met5p  GRS1 - glycine--trna ligase  MYO4 - myosin 4  URM1 - ubiquitin-related modifier urm1  CMD1 - cmd1p  GPI16 - gpi16p  YSA1 - ysa1p  FES1 - fes1p  BAT2 - bat2p  YMC2 - ymc2p  GND1 - phosphogluconate dehydrogenase (decarboxylating) gnd1  RSM7 - rsm7p  PPX1 - ppx1p  RFC3 - replication factor c subunit 3  CPA2 - cpa2p  PBY1 - pby1p  SEC39 - sec39p  CBR1 - cbr1p  URA8 - ura8p  AIM18 - aim18p  RSM26 - rsm26p  IST2 - ist2p  RPL6B - ribosomal 60s subunit protein l6b  ECM33 - ecm33p  CAP2 - cap2p  CRN1 - crn1p  SSM4 - e3 ubiquitin-protein ligase ssm4  SEC18 - sec18p  BCY1 - bcy1p  ERG24 - delta(14)-sterol reductase  TAT1 - tat1p  IST1 - ist1p  ATP2 - atp2p  CAR2 - ornithine-oxo-acid transaminase  EMC5 - emc5p  TIM44 - tim44p  RPL17A - ribosomal 60s subunit protein l17a  DSL1 - dsl1p  CDC12 - septin cdc12  TPO5 - tpo5p  GRE3 - trifunctional aldehyde reductase/xylose reductase/glucose 1-dehydrogenase (nadp(+))  YRB2 - yrb2p  SUI1 - sui1p  MAM33 - mam33p  APE2 - ape2p  ARC15 - arc15p  ATP3 - atp3p  MCD4 - mcd4p  PDR16 - pdr16p  SDS22 - sds22p  RHO3 - rho3p  ETR1 - etr1p  YKT6 - ykt6p  GRX7 - glutathione-disulfide reductase grx7  YNL208W - hypothetical protein  RHO5 - rho5p  RPS3 - ribosomal 40s subunit protein s3  MRPL19 - mitochondrial 54s ribosomal protein yml19  ILV3 - ilv3p  SPC1 - spc1p  CHS1 - chitin synthase chs1  KGD1 - alpha-ketoglutarate dehydrogenase kgd1  FMP52 - fmp52p  AYR1 - acylglycerone-phosphate reductase  GPX2 - glutathione peroxidase gpx2  GGC1 - ggc1p  CPR7 - cpr7p  ISC1 - inositol phosphosphingolipid phospholipase  AFG3 - aaa family atpase afg3  PGA2 - pga2p  PCS60 - pcs60p  MCX1 - mcx1p  GDH2 - glutamate dehydrogenase (nad(+))  NRK1 - ribosylnicotinamide kinase  GUT2 - glycerol-3-phosphate dehydrogenase  SRV2 - srv2p  ERG28 - erg28p  SUC2 - beta-fructofuranosidase suc2  SDS24 - sds24p  LYS21 - homocitrate synthase lys21  ARP3 - arp3p  RPS9B - ribosomal 40s subunit protein s9b  DBP2 - dbp2p  DLD3 - dld3p  PRB1 - prb1p  CDC11 - cdc11p  RDI1 - rdi1p  PGI1 - glucose-6-phosphate isomerase  COP1 - cop1p  RAS2 - ras2p  SSE2 - sse2p  RPO21 - rpo21p  LEU4 - 2-isopropylmalate synthase leu4  POL1 - pol1p  EHT1 - eht1p  ALD4 - aldehyde dehydrogenase (nadp(+)) ald4  GLT1 - glutamate synthase (nadh)  RPS12 - ribosomal 40s subunit protein s12  SPF1 - spf1p  DLD1 - dld1p  SCP1 - scp1p  CDC28 - cdc28p  MKT1 - mkt1p  YDJ1 - ydj1p  GEA2 - gea2p  RIP1 - ubiquinol--cytochrome-c reductase catalytic subunit rip1  DLD2 - dld2p  GCD10 - gcd10p  VAC8 - vac8p  LAT1 - dihydrolipoyllysine-residue acetyltransferase  ARA1 - d-arabinose 1-dehydrogenase (nad(p)(+)) ara1  SSB1 - hsp70 family atpase ssb1  RPL9B - ribosomal 60s subunit protein l9b  CCT5 - cct5p  PTP1 - ptp1p  RPL16B - ribosomal 60s subunit protein l16b |
| GO:0045261 | proton-transporting ATP synthase complex, catalytic core F(1) | 8.13E-4 | 6.5E-2 | 9.48 (2298,5,194,4) | [+] Show genes  ATP1 - f1f0 atp synthase subunit alpha  ATP2 - atp2p  ATP15 - f1f0 atp synthase subunit epsilon  ATP3 - atp3p |
| GO:0043233 | organelle lumen | 8.22E-4 | 6.02E-2 | 1.67 (2298,137,493,49) | [+] Show genes  IDH1 - isocitrate dehydrogenase (nad(+)) idh1  MSC6 - msc6p  TRX1 - trx1p  KGD1 - alpha-ketoglutarate dehydrogenase kgd1  COA4 - coa4p  CPR3 - peptidylprolyl isomerase cpr3  MDH1 - malate dehydrogenase mdh1  PPX1 - ppx1p  FUM1 - fumarase fum1  ACO1 - aconitate hydratase aco1  MAE1 - malate dehydrogenase (oxaloacetate-decarboxylating)  QCR2 - ubiquinol--cytochrome-c reductase subunit 2  PCS60 - pcs60p  CIT1 - citrate (si)-synthase cit1  SNF4 - snf4p  MAS1 - mas1p  RIB3 - 3,4-dihydroxy-2-butanone-4-phosphate synthase rib3  TFS1 - tfs1p  GUT2 - glycerol-3-phosphate dehydrogenase  ERG3 - c-5 sterol desaturase  SCJ1 - scj1p  TIM44 - tim44p  PTC5 - ptc5p  ARG5,6 - bifunctional acetylglutamate kinase/n-acetyl-gamma-glutamyl-phosphate reductase  AIM45 - aim45p  ARG7 - glutamate n-acetyltransferase  MAS2 - mas2p  CYM1 - cym1p  PRB1 - prb1p  OCT1 - oct1p  MAM33 - mam33p  MDJ1 - mdj1p  IDH2 - isocitrate dehydrogenase (nad(+)) idh2  CCA1 - cca1p  GPM1 - phosphoglycerate mutase gpm1  CMC2 - cmc2p  MCR1 - mcr1p  COX17 - cox17p  MDH3 - malate dehydrogenase mdh3  ALD4 - aldehyde dehydrogenase (nadp(+)) ald4  TIM10 - tim10p  NDI1 - nadh-ubiquinone reductase (h(+)-translocating) ndi1  POS5 - pos5p  CPR5 - peptidylprolyl isomerase family protein cpr5  GRX7 - glutathione-disulfide reductase grx7  MIC23 - mic23p  DLD2 - dld2p  NDE1 - nadh-ubiquinone reductase (h(+)-translocating) nde1  ALD5 - aldehyde dehydrogenase (nad(p)(+)) ald5 |
| GO:0070013 | intracellular organelle lumen | 8.22E-4 | 5.56E-2 | 1.67 (2298,137,493,49) | [+] Show genes  IDH1 - isocitrate dehydrogenase (nad(+)) idh1  MSC6 - msc6p  TRX1 - trx1p  KGD1 - alpha-ketoglutarate dehydrogenase kgd1  COA4 - coa4p  CPR3 - peptidylprolyl isomerase cpr3  MDH1 - malate dehydrogenase mdh1  PPX1 - ppx1p  FUM1 - fumarase fum1  ACO1 - aconitate hydratase aco1  MAE1 - malate dehydrogenase (oxaloacetate-decarboxylating)  QCR2 - ubiquinol--cytochrome-c reductase subunit 2  PCS60 - pcs60p  CIT1 - citrate (si)-synthase cit1  SNF4 - snf4p  MAS1 - mas1p  RIB3 - 3,4-dihydroxy-2-butanone-4-phosphate synthase rib3  TFS1 - tfs1p  GUT2 - glycerol-3-phosphate dehydrogenase  ERG3 - c-5 sterol desaturase  SCJ1 - scj1p  PTC5 - ptc5p  TIM44 - tim44p  ARG5,6 - bifunctional acetylglutamate kinase/n-acetyl-gamma-glutamyl-phosphate reductase  AIM45 - aim45p  ARG7 - glutamate n-acetyltransferase  MAS2 - mas2p  CYM1 - cym1p  PRB1 - prb1p  OCT1 - oct1p  MAM33 - mam33p  MDJ1 - mdj1p  IDH2 - isocitrate dehydrogenase (nad(+)) idh2  CMC2 - cmc2p  GPM1 - phosphoglycerate mutase gpm1  CCA1 - cca1p  MCR1 - mcr1p  COX17 - cox17p  MDH3 - malate dehydrogenase mdh3  ALD4 - aldehyde dehydrogenase (nadp(+)) ald4  TIM10 - tim10p  NDI1 - nadh-ubiquinone reductase (h(+)-translocating) ndi1  POS5 - pos5p  CPR5 - peptidylprolyl isomerase family protein cpr5  GRX7 - glutathione-disulfide reductase grx7  DLD2 - dld2p  MIC23 - mic23p  NDE1 - nadh-ubiquinone reductase (h(+)-translocating) nde1  ALD5 - aldehyde dehydrogenase (nad(p)(+)) ald5 |
| GO:0031974 | membrane-enclosed lumen | 8.22E-4 | 5.16E-2 | 1.67 (2298,137,493,49) | [+] Show genes  IDH1 - isocitrate dehydrogenase (nad(+)) idh1  MSC6 - msc6p  TRX1 - trx1p  KGD1 - alpha-ketoglutarate dehydrogenase kgd1  COA4 - coa4p  CPR3 - peptidylprolyl isomerase cpr3  MDH1 - malate dehydrogenase mdh1  PPX1 - ppx1p  FUM1 - fumarase fum1  ACO1 - aconitate hydratase aco1  MAE1 - malate dehydrogenase (oxaloacetate-decarboxylating)  QCR2 - ubiquinol--cytochrome-c reductase subunit 2  PCS60 - pcs60p  CIT1 - citrate (si)-synthase cit1  SNF4 - snf4p  MAS1 - mas1p  RIB3 - 3,4-dihydroxy-2-butanone-4-phosphate synthase rib3  TFS1 - tfs1p  GUT2 - glycerol-3-phosphate dehydrogenase  ERG3 - c-5 sterol desaturase  SCJ1 - scj1p  PTC5 - ptc5p  TIM44 - tim44p  ARG5,6 - bifunctional acetylglutamate kinase/n-acetyl-gamma-glutamyl-phosphate reductase  AIM45 - aim45p  ARG7 - glutamate n-acetyltransferase  MAS2 - mas2p  CYM1 - cym1p  PRB1 - prb1p  OCT1 - oct1p  MAM33 - mam33p  MDJ1 - mdj1p  IDH2 - isocitrate dehydrogenase (nad(+)) idh2  CMC2 - cmc2p  GPM1 - phosphoglycerate mutase gpm1  CCA1 - cca1p  MCR1 - mcr1p  COX17 - cox17p  MDH3 - malate dehydrogenase mdh3  ALD4 - aldehyde dehydrogenase (nadp(+)) ald4  TIM10 - tim10p  NDI1 - nadh-ubiquinone reductase (h(+)-translocating) ndi1  POS5 - pos5p  CPR5 - peptidylprolyl isomerase family protein cpr5  GRX7 - glutathione-disulfide reductase grx7  DLD2 - dld2p  MIC23 - mic23p  NDE1 - nadh-ubiquinone reductase (h(+)-translocating) nde1  ALD5 - aldehyde dehydrogenase (nad(p)(+)) ald5 |

Species used: Saccharomyces cerevisiae

The system has recognized 2316 genes out of 2316 gene terms entered by the user.  
 136 genes were recognized by gene symbol and 2180 genes by other gene IDs .  
Only 2298 of these genes are associated with a GO term.

  
 Output in Microsoft Excel format  

% List genereted using GOrilla
% http://cbl-gorilla.cs.technion.ac.il/
% GO term pValue
GO:0005739 1.48E-7
GO:0044429 1.59E-5
GO:0005743 3.72E-5
GO:0019866 4.68E-5
GO:0005829 1.1E-4
GO:0098800 1.22E-4
GO:0031966 2.55E-4
GO:0048471 3.42E-4
GO:0045239 4.54E-4
GO:0044444 5.55E-4
GO:0045261 8.13E-4
GO:0043233 8.22E-4
GO:0070013 8.22E-4
GO:0031974 8.22E-4
 Visualize output in REViGO   

The GOrilla database is periodically updated using the GO database and other sources.  
The GOrilla database was last updated on Sep 21, 2019

This results page will be available on this site for one month from now (until
Oct 23, 2019
). You can bookmark this page and come back to it later.

  
**'P-value'** is the enrichment
p-value computed according to the mHG or HG model. This p-value is not
corrected for multiple testing of 879 GO terms.  
  
**'FDR q-value'** is the correction of the above p-value for multiple testing using the Benjamini and Hochberg (1995) method.   
Namely, for the i^th^ term (ranked according to p-value) the FDR q-value is (p-value \* number of GO terms) / i.   
  
**Enrichment (N, B, n, b)** is defined as follows:  
N - is the total number of genes  
B - is the total number of genes associated with a specific GO term  
n - is the number of genes in the top of the user's input list or in the target set when appropriate  
b - is the number of genes in the intersection  
Enrichment = (b/n) / (B/N)  
  
**Genes:** For each GO term you can see the list of associated genes that appear in the optimal top of the list.  
Each gene name is specified by gene symbol followed by a short description of the gene   

Back to GOrilla analysis results
