## Supplemental Data S1 for "A simple mass-action model predicts genome-wide protein timecourses from mRNA trajectories during a dynamic response in two strains of *Saccharomyces cerevisiae*": GOCOMPONENT_prot.html

Results


 **protein** 

*P-value color scale*

|  |  |  |  |  |
| --- | --- | --- | --- | --- |
| > 10^-3^ | 10^-3^ to 10^-5^ | 10^-5^ to 10^-7^ | 10^-7^ to 10^-9^ | < 10^-9^ |


|  |  |  |  |  |  |
| --- | --- | --- | --- | --- | --- |
| **GO term** | **Description** | **P-value** | **FDR q-value** | **Enrichment (N, B, n, b)** | **Genes** |
| GO:0022625 | cytosolic large ribosomal subunit | 1.15E-5 | 1.01E-2 | 2.60 (2325,51,438,25) | [+] Show genes  RPL29 - ribosomal 60s subunit protein l29  RPL25 - ribosomal 60s subunit protein l25  RPL24A - ribosomal 60s subunit protein l24a  RPL30 - ribosomal 60s subunit protein l30  RPL17B - rpl17bp  RPL7B - ribosomal 60s subunit protein l7b  RPL32 - ribosomal 60s subunit protein l32  RPL26B - ribosomal 60s subunit protein l26b  RPL9A - ribosomal 60s subunit protein l9a  RPL8A - ribosomal 60s subunit protein l8a  REI1 - rei1p  RPL26A - ribosomal 60s subunit protein l26a  RPL15A - ribosomal 60s subunit protein l15a  RPP0 - ribosomal protein p0  RPL6A - ribosomal 60s subunit protein l6a  RPL3 - ribosomal 60s subunit protein l3  RPL14B - ribosomal 60s subunit protein l14b  RPL31A - ribosomal 60s subunit protein l31a  RPL10 - ribosomal 60s subunit protein l10  RPL6B - ribosomal 60s subunit protein l6b  RPL33B - ribosomal 60s subunit protein l33b  RPP2A - ribosomal protein p2a  RPL5 - ribosomal 60s subunit protein l5  RPL33A - ribosomal 60s subunit protein l33a  RPL16B - ribosomal 60s subunit protein l16b |
| GO:0044391 | ribosomal subunit | 1.79E-4 | 7.87E-2 | 1.87 (2325,116,451,42) | [+] Show genes  RPS1B - ribosomal 40s subunit protein s1b  RPL29 - ribosomal 60s subunit protein l29  RPL25 - ribosomal 60s subunit protein l25  RPS9B - ribosomal 40s subunit protein s9b  RPL24A - ribosomal 60s subunit protein l24a  RPS3 - ribosomal 40s subunit protein s3  RPL17B - rpl17bp  RPL30 - ribosomal 60s subunit protein l30  RPL7B - ribosomal 60s subunit protein l7b  MRPL19 - mitochondrial 54s ribosomal protein yml19  RPL32 - ribosomal 60s subunit protein l32  RPL26B - ribosomal 60s subunit protein l26b  RPL9A - ribosomal 60s subunit protein l9a  RPL8A - ribosomal 60s subunit protein l8a  REI1 - rei1p  YGR054W - hypothetical protein  ASC1 - asc1p  EHD3 - ehd3p  RSM18 - mitochondrial 37s ribosomal protein rsm18  RPL26A - ribosomal 60s subunit protein l26a  RPL15A - ribosomal 60s subunit protein l15a  RPP0 - ribosomal protein p0  RSM7 - rsm7p  RPL6A - ribosomal 60s subunit protein l6a  RPL3 - ribosomal 60s subunit protein l3  RPL14B - ribosomal 60s subunit protein l14b  RPL31A - ribosomal 60s subunit protein l31a  RPL10 - ribosomal 60s subunit protein l10  RSM26 - rsm26p  RPL6B - ribosomal 60s subunit protein l6b  MRPS12 - putative mitochondrial 37s ribosomal protein mrps12  RPS2 - ribosomal 40s subunit protein s2  RPL33B - ribosomal 60s subunit protein l33b  RPP2A - ribosomal protein p2a  RPS13 - ribosomal 40s subunit protein s13  RPS5 - rps5p  RPL5 - ribosomal 60s subunit protein l5  RSM25 - mitochondrial 37s ribosomal protein rsm25  RPS7A - ribosomal 40s subunit protein s7a  MRPL6 - mitochondrial 54s ribosomal protein yml16  RPL33A - ribosomal 60s subunit protein l33a  RPL16B - ribosomal 60s subunit protein l16b |
| GO:0045242 | isocitrate dehydrogenase complex (NAD+) | 5.2E-4 | 1.53E-1 | 61.18 (2325,2,38,2) | [+] Show genes  IDH1 - isocitrate dehydrogenase (nad(+)) idh1  IDH2 - isocitrate dehydrogenase (nad(+)) idh2 |
| GO:0005962 | mitochondrial isocitrate dehydrogenase complex (NAD+) | 5.2E-4 | 1.14E-1 | 61.18 (2325,2,38,2) | [+] Show genes  IDH1 - isocitrate dehydrogenase (nad(+)) idh1  IDH2 - isocitrate dehydrogenase (nad(+)) idh2 |

Species used: Saccharomyces cerevisiae

The system has recognized 2343 genes out of 2345 gene terms entered by the user.  
 141 genes were recognized by gene symbol and 2202 genes by other gene IDs .  
Only 2325 of these genes are associated with a GO term.

  
 Output in Microsoft Excel format  

% List genereted using GOrilla
% http://cbl-gorilla.cs.technion.ac.il/
% GO term pValue
GO:0022625 1.15E-5
GO:0044391 1.79E-4
GO:0045242 5.2E-4
GO:0005962 5.2E-4
 Visualize output in REViGO   

The GOrilla database is periodically updated using the GO database and other sources.  
The GOrilla database was last updated on Sep 21, 2019

This results page will be available on this site for one month from now (until
Oct 23, 2019
). You can bookmark this page and come back to it later.

  
**'P-value'** is the enrichment
p-value computed according to the mHG or HG model. This p-value is not
corrected for multiple testing of 880 GO terms.  
  
**'FDR q-value'** is the correction of the above p-value for multiple testing using the Benjamini and Hochberg (1995) method.   
Namely, for the i^th^ term (ranked according to p-value) the FDR q-value is (p-value \* number of GO terms) / i.   
  
**Enrichment (N, B, n, b)** is defined as follows:  
N - is the total number of genes  
B - is the total number of genes associated with a specific GO term  
n - is the number of genes in the top of the user's input list or in the target set when appropriate  
b - is the number of genes in the intersection  
Enrichment = (b/n) / (B/N)  
  
**Genes:** For each GO term you can see the list of associated genes that appear in the optimal top of the list.  
Each gene name is specified by gene symbol followed by a short description of the gene   

Back to GOrilla analysis results
