## Supplemental Data S1 for "A simple mass-action model predicts genome-wide protein timecourses from mRNA trajectories during a dynamic response in two strains of *Saccharomyces cerevisiae*": GOCOMPONENT_rna.html

Results


 **rna** 

*P-value color scale*

|  |  |  |  |  |
| --- | --- | --- | --- | --- |
| > 10^-3^ | 10^-3^ to 10^-5^ | 10^-5^ to 10^-7^ | 10^-7^ to 10^-9^ | < 10^-9^ |


|  |  |  |  |  |  |
| --- | --- | --- | --- | --- | --- |
| **GO term** | **Description** | **P-value** | **FDR q-value** | **Enrichment (N, B, n, b)** | **Genes** |
| GO:0044445 | cytosolic part | 1.54E-46 | 1.63E-43 | 10.07 (5247,207,156,62) | [+] Show genes  RPL29 - ribosomal 60s subunit protein l29  RPL2B - ribosomal 60s subunit protein l2b  RPS0B - ribosomal 40s subunit protein s0b  RPL24A - ribosomal 60s subunit protein l24a  RPL30 - ribosomal 60s subunit protein l30  RPL9A - ribosomal 60s subunit protein l9a  RPL40A - ubiquitin-ribosomal 60s subunit protein l40a fusion protein  RPL23B - ribosomal 60s subunit protein l23b  RPL38 - ribosomal 60s subunit protein l38  RPL13B - ribosomal 60s subunit protein l13b  RPS16A - ribosomal 40s subunit protein s16a  RPL8A - ribosomal 60s subunit protein l8a  ASC1 - asc1p  RPS8B - ribosomal 40s subunit protein s8b  RPL26A - ribosomal 60s subunit protein l26a  RPL15A - ribosomal 60s subunit protein l15a  RPL11A - ribosomal 60s subunit protein l11a  RPP0 - ribosomal protein p0  RPL27B - ribosomal 60s subunit protein l27b  RPL34B - ribosomal 60s subunit protein l34b  RPL3 - ribosomal 60s subunit protein l3  RPS18B - ribosomal 40s subunit protein s18b  RPL19A - ribosomal 60s subunit protein l19a  RPS21B - rps21bp  RPS10B - ribosomal 40s subunit protein s10b  RPP2B - ribosomal protein p2b  RLP7 - rlp7p  RPL6B - ribosomal 60s subunit protein l6b  RPS15 - ribosomal 40s subunit protein s15  RPP2A - ribosomal protein p2a  RPS5 - rps5p  RPS17A - ribosomal 40s subunit protein s17a  RPL37A - ribosomal 60s subunit protein l37a  RPL24B - ribosomal 60s subunit protein l24b  RPS28A - ribosomal 40s subunit protein s28a  RPS27B - ribosomal 40s subunit protein s27b  RPL17A - ribosomal 60s subunit protein l17a  RPS1B - ribosomal 40s subunit protein s1b  RPP1B - ribosomal protein p1b  RPL21A - ribosomal 60s subunit protein l21a  RPL14A - ribosomal 60s subunit protein l14a  RPL17B - rpl17bp  RPL32 - ribosomal 60s subunit protein l32  RPL26B - ribosomal 60s subunit protein l26b  RPL22B - ribosomal 60s subunit protein l22b  RPL35B - ribosomal 60s subunit protein l35b  RPS10A - ribosomal 40s subunit protein s10a  RPL34A - ribosomal 60s subunit protein l34a  RPL27A - ribosomal 60s subunit protein l27a  RNR4 - ribonucleotide-diphosphate reductase subunit rnr4  RPS22A - rps22ap  RPS17B - ribosomal 40s subunit protein s17b  RPL6A - ribosomal 60s subunit protein l6a  RPS23B - ribosomal 40s subunit protein s23b  RPL42B - ribosomal 60s subunit protein l42b  RPS14A - ribosomal 40s subunit protein s14a  RPL31A - ribosomal 60s subunit protein l31a  RPS13 - ribosomal 40s subunit protein s13  RPS26A - ribosomal 40s subunit protein s26a  RPS7A - ribosomal 40s subunit protein s7a  RPL16B - ribosomal 60s subunit protein l16b  RPS24A - ribosomal 40s subunit protein s24a |
| GO:0044391 | ribosomal subunit | 7.13E-46 | 3.78E-43 | 9.85 (5247,213,155,62) | [+] Show genes  RPL29 - ribosomal 60s subunit protein l29  RPL2B - ribosomal 60s subunit protein l2b  RPS0B - ribosomal 40s subunit protein s0b  RPL24A - ribosomal 60s subunit protein l24a  RPL30 - ribosomal 60s subunit protein l30  RPL9A - ribosomal 60s subunit protein l9a  RPL40A - ubiquitin-ribosomal 60s subunit protein l40a fusion protein  RPL23B - ribosomal 60s subunit protein l23b  RPL38 - ribosomal 60s subunit protein l38  RPL13B - ribosomal 60s subunit protein l13b  RPS16A - ribosomal 40s subunit protein s16a  RPL8A - ribosomal 60s subunit protein l8a  ASC1 - asc1p  RPS8B - ribosomal 40s subunit protein s8b  RPL26A - ribosomal 60s subunit protein l26a  RPL11A - ribosomal 60s subunit protein l11a  RPL15A - ribosomal 60s subunit protein l15a  RPP0 - ribosomal protein p0  RPL27B - ribosomal 60s subunit protein l27b  RPL34B - ribosomal 60s subunit protein l34b  RPL3 - ribosomal 60s subunit protein l3  RPS18B - ribosomal 40s subunit protein s18b  RPL19A - ribosomal 60s subunit protein l19a  RPS21B - rps21bp  RPS10B - ribosomal 40s subunit protein s10b  RPP2B - ribosomal protein p2b  RLP7 - rlp7p  RPL6B - ribosomal 60s subunit protein l6b  RPS15 - ribosomal 40s subunit protein s15  RPP2A - ribosomal protein p2a  RPS5 - rps5p  RPS17A - ribosomal 40s subunit protein s17a  RPL37A - ribosomal 60s subunit protein l37a  RPL24B - ribosomal 60s subunit protein l24b  RPS28A - ribosomal 40s subunit protein s28a  RPS27B - ribosomal 40s subunit protein s27b  RPL17A - ribosomal 60s subunit protein l17a  RPS1B - ribosomal 40s subunit protein s1b  RPP1B - ribosomal protein p1b  RPL21A - ribosomal 60s subunit protein l21a  RPL14A - ribosomal 60s subunit protein l14a  RPL17B - rpl17bp  MRPL24 - mitochondrial 54s ribosomal protein yml24/yml14  RPL32 - ribosomal 60s subunit protein l32  RPL26B - ribosomal 60s subunit protein l26b  RPL22B - ribosomal 60s subunit protein l22b  RPL35B - ribosomal 60s subunit protein l35b  RPS10A - ribosomal 40s subunit protein s10a  RPL34A - ribosomal 60s subunit protein l34a  RPL27A - ribosomal 60s subunit protein l27a  RPS22A - rps22ap  RPS17B - ribosomal 40s subunit protein s17b  RPL6A - ribosomal 60s subunit protein l6a  RPS23B - ribosomal 40s subunit protein s23b  RPL42B - ribosomal 60s subunit protein l42b  RPS14A - ribosomal 40s subunit protein s14a  RPL31A - ribosomal 60s subunit protein l31a  RPS13 - ribosomal 40s subunit protein s13  RPS26A - ribosomal 40s subunit protein s26a  RPS7A - ribosomal 40s subunit protein s7a  RPL16B - ribosomal 60s subunit protein l16b  RPS24A - ribosomal 40s subunit protein s24a |
| GO:0005840 | ribosome | 1.58E-45 | 5.6E-43 | 9.48 (5247,225,155,63) | [+] Show genes  RPL29 - ribosomal 60s subunit protein l29  RPL2B - ribosomal 60s subunit protein l2b  RPS0B - ribosomal 40s subunit protein s0b  RPL24A - ribosomal 60s subunit protein l24a  RPL30 - ribosomal 60s subunit protein l30  SED1 - sed1p  RPL9A - ribosomal 60s subunit protein l9a  RPL40A - ubiquitin-ribosomal 60s subunit protein l40a fusion protein  RPL23B - ribosomal 60s subunit protein l23b  RPL38 - ribosomal 60s subunit protein l38  RPL13B - ribosomal 60s subunit protein l13b  TEF1 - tef1p  RPS16A - ribosomal 40s subunit protein s16a  RPL8A - ribosomal 60s subunit protein l8a  ASC1 - asc1p  RPS8B - ribosomal 40s subunit protein s8b  RPL26A - ribosomal 60s subunit protein l26a  RPL11A - ribosomal 60s subunit protein l11a  RPL15A - ribosomal 60s subunit protein l15a  RPP0 - ribosomal protein p0  RPL27B - ribosomal 60s subunit protein l27b  RPL34B - ribosomal 60s subunit protein l34b  RPL3 - ribosomal 60s subunit protein l3  RPS18B - ribosomal 40s subunit protein s18b  RPL19A - ribosomal 60s subunit protein l19a  RPS21B - rps21bp  RPS10B - ribosomal 40s subunit protein s10b  RPP2B - ribosomal protein p2b  RPL6B - ribosomal 60s subunit protein l6b  RPS15 - ribosomal 40s subunit protein s15  RPP2A - ribosomal protein p2a  RPS5 - rps5p  RPS17A - ribosomal 40s subunit protein s17a  RPL37A - ribosomal 60s subunit protein l37a  RPL24B - ribosomal 60s subunit protein l24b  RPS28A - ribosomal 40s subunit protein s28a  RPS27B - ribosomal 40s subunit protein s27b  RPL17A - ribosomal 60s subunit protein l17a  RPS1B - ribosomal 40s subunit protein s1b  RPL21A - ribosomal 60s subunit protein l21a  RPP1B - ribosomal protein p1b  RPL14A - ribosomal 60s subunit protein l14a  RPL17B - rpl17bp  MRPL24 - mitochondrial 54s ribosomal protein yml24/yml14  RPL32 - ribosomal 60s subunit protein l32  RPL26B - ribosomal 60s subunit protein l26b  RPL22B - ribosomal 60s subunit protein l22b  RPL35B - ribosomal 60s subunit protein l35b  RPS10A - ribosomal 40s subunit protein s10a  RPL34A - ribosomal 60s subunit protein l34a  RPL27A - ribosomal 60s subunit protein l27a  RPS22A - rps22ap  RPS17B - ribosomal 40s subunit protein s17b  RPL6A - ribosomal 60s subunit protein l6a  RPS23B - ribosomal 40s subunit protein s23b  RPL42B - ribosomal 60s subunit protein l42b  RPS14A - ribosomal 40s subunit protein s14a  RPL31A - ribosomal 60s subunit protein l31a  RPS13 - ribosomal 40s subunit protein s13  RPS26A - ribosomal 40s subunit protein s26a  RPS7A - ribosomal 40s subunit protein s7a  RPL16B - ribosomal 60s subunit protein l16b  RPS24A - ribosomal 40s subunit protein s24a |
| GO:0022625 | cytosolic large ribosomal subunit | 8.74E-42 | 2.32E-39 | 14.75 (5247,76,206,44) | [+] Show genes  RPL17A - ribosomal 60s subunit protein l17a  RPL29 - ribosomal 60s subunit protein l29  RPL2B - ribosomal 60s subunit protein l2b  RPL14A - ribosomal 60s subunit protein l14a  RPL21A - ribosomal 60s subunit protein l21a  RPP1B - ribosomal protein p1b  RPL24A - ribosomal 60s subunit protein l24a  RPL17B - rpl17bp  RPL30 - ribosomal 60s subunit protein l30  RPL31B - ribosomal 60s subunit protein l31b  RPL32 - ribosomal 60s subunit protein l32  RPL26B - ribosomal 60s subunit protein l26b  RPL22B - ribosomal 60s subunit protein l22b  RPL35B - ribosomal 60s subunit protein l35b  RPL9A - ribosomal 60s subunit protein l9a  RPL40A - ubiquitin-ribosomal 60s subunit protein l40a fusion protein  RPL23B - ribosomal 60s subunit protein l23b  RPL41B - ribosomal 60s subunit protein l41b  RPL38 - ribosomal 60s subunit protein l38  RPL13B - ribosomal 60s subunit protein l13b  RPL8A - ribosomal 60s subunit protein l8a  RPL34A - ribosomal 60s subunit protein l34a  RPL12A - ribosomal 60s subunit protein l12a  RPL27A - ribosomal 60s subunit protein l27a  RPL16A - ribosomal 60s subunit protein l16a  RPL26A - ribosomal 60s subunit protein l26a  RPL11A - ribosomal 60s subunit protein l11a  RPL15A - ribosomal 60s subunit protein l15a  RPP0 - ribosomal protein p0  RPL6A - ribosomal 60s subunit protein l6a  RPL34B - ribosomal 60s subunit protein l34b  RPL27B - ribosomal 60s subunit protein l27b  RPL42B - ribosomal 60s subunit protein l42b  RPL3 - ribosomal 60s subunit protein l3  RPL31A - ribosomal 60s subunit protein l31a  RPL19A - ribosomal 60s subunit protein l19a  RPP2B - ribosomal protein p2b  RLP7 - rlp7p  RPL6B - ribosomal 60s subunit protein l6b  RPP2A - ribosomal protein p2a  RPL11B - ribosomal 60s subunit protein l11b  RPL37A - ribosomal 60s subunit protein l37a  RPL16B - ribosomal 60s subunit protein l16b  RPL24B - ribosomal 60s subunit protein l24b |
| GO:0015934 | large ribosomal subunit | 1.95E-31 | 4.15E-29 | 9.80 (5247,122,193,44) | [+] Show genes  RPL2B - ribosomal 60s subunit protein l2b  RPL29 - ribosomal 60s subunit protein l29  RPL17A - ribosomal 60s subunit protein l17a  RPL21A - ribosomal 60s subunit protein l21a  RPP1B - ribosomal protein p1b  RPL14A - ribosomal 60s subunit protein l14a  RPL24A - ribosomal 60s subunit protein l24a  RPL17B - rpl17bp  RPL30 - ribosomal 60s subunit protein l30  RPL31B - ribosomal 60s subunit protein l31b  RPL32 - ribosomal 60s subunit protein l32  MRPL24 - mitochondrial 54s ribosomal protein yml24/yml14  RPL26B - ribosomal 60s subunit protein l26b  RPL22B - ribosomal 60s subunit protein l22b  RPL35B - ribosomal 60s subunit protein l35b  RPL9A - ribosomal 60s subunit protein l9a  RPL23B - ribosomal 60s subunit protein l23b  RPL40A - ubiquitin-ribosomal 60s subunit protein l40a fusion protein  RPL41B - ribosomal 60s subunit protein l41b  RPL13B - ribosomal 60s subunit protein l13b  RPL38 - ribosomal 60s subunit protein l38  RPL8A - ribosomal 60s subunit protein l8a  RPL34A - ribosomal 60s subunit protein l34a  RPL27A - ribosomal 60s subunit protein l27a  RPL16A - ribosomal 60s subunit protein l16a  RPL26A - ribosomal 60s subunit protein l26a  RPL15A - ribosomal 60s subunit protein l15a  RPL11A - ribosomal 60s subunit protein l11a  RPP0 - ribosomal protein p0  RPL6A - ribosomal 60s subunit protein l6a  RPL27B - ribosomal 60s subunit protein l27b  RPL34B - ribosomal 60s subunit protein l34b  RPL42B - ribosomal 60s subunit protein l42b  RPL3 - ribosomal 60s subunit protein l3  RPL31A - ribosomal 60s subunit protein l31a  RPL19A - ribosomal 60s subunit protein l19a  RPP2B - ribosomal protein p2b  RPL6B - ribosomal 60s subunit protein l6b  RLP7 - rlp7p  RPP2A - ribosomal protein p2a  RPL11B - ribosomal 60s subunit protein l11b  RPL37A - ribosomal 60s subunit protein l37a  RPL16B - ribosomal 60s subunit protein l16b  RPL24B - ribosomal 60s subunit protein l24b |
| GO:1990904 | ribonucleoprotein complex | 4.12E-28 | 7.29E-26 | 4.29 (5247,616,143,72) | [+] Show genes  RPL2B - ribosomal 60s subunit protein l2b  RPL29 - ribosomal 60s subunit protein l29  ECM16 - ecm16p  RPS0B - ribosomal 40s subunit protein s0b  VMA2 - vma2p  RPL30 - ribosomal 60s subunit protein l30  SED1 - sed1p  EFG1 - efg1p  RPL40A - ubiquitin-ribosomal 60s subunit protein l40a fusion protein  RPL23B - ribosomal 60s subunit protein l23b  RPL38 - ribosomal 60s subunit protein l38  RPL13B - ribosomal 60s subunit protein l13b  RPS16A - ribosomal 40s subunit protein s16a  TEF1 - tef1p  RPL8A - ribosomal 60s subunit protein l8a  UTP14 - utp14p  ASC1 - asc1p  RPS8B - ribosomal 40s subunit protein s8b  RPL26A - ribosomal 60s subunit protein l26a  RPL11A - ribosomal 60s subunit protein l11a  RPL15A - ribosomal 60s subunit protein l15a  RPP0 - ribosomal protein p0  RPL34B - ribosomal 60s subunit protein l34b  RPL27B - ribosomal 60s subunit protein l27b  RPL3 - ribosomal 60s subunit protein l3  RPS18B - ribosomal 40s subunit protein s18b  RPL19A - ribosomal 60s subunit protein l19a  RPS21B - rps21bp  SMB1 - smb1p  RPS10B - ribosomal 40s subunit protein s10b  RPP2B - ribosomal protein p2b  RPL6B - ribosomal 60s subunit protein l6b  RLP7 - rlp7p  RPS15 - ribosomal 40s subunit protein s15  RPP2A - ribosomal protein p2a  RPS5 - rps5p  RPS17A - ribosomal 40s subunit protein s17a  RPL37A - ribosomal 60s subunit protein l37a  RPL24B - ribosomal 60s subunit protein l24b  RPS28A - ribosomal 40s subunit protein s28a  RPL17A - ribosomal 60s subunit protein l17a  RPS1B - ribosomal 40s subunit protein s1b  RPS27B - ribosomal 40s subunit protein s27b  RPL21A - ribosomal 60s subunit protein l21a  RPP1B - ribosomal protein p1b  RPL14A - ribosomal 60s subunit protein l14a  RPL17B - rpl17bp  HCA4 - hca4p  RPL32 - ribosomal 60s subunit protein l32  MRPL24 - mitochondrial 54s ribosomal protein yml24/yml14  RPL26B - ribosomal 60s subunit protein l26b  RPL22B - ribosomal 60s subunit protein l22b  RPS10A - ribosomal 40s subunit protein s10a  NSR1 - nsr1p  RPL34A - ribosomal 60s subunit protein l34a  SSE1 - sse1p  RPL27A - ribosomal 60s subunit protein l27a  RPS22A - rps22ap  RPS17B - ribosomal 40s subunit protein s17b  NEW1 - new1p  RPL6A - ribosomal 60s subunit protein l6a  RPS23B - ribosomal 40s subunit protein s23b  RPL42B - ribosomal 60s subunit protein l42b  RPS14A - ribosomal 40s subunit protein s14a  RPL31A - ribosomal 60s subunit protein l31a  STM1 - stm1p  RIO1 - rio1p  RPS13 - ribosomal 40s subunit protein s13  RPS26A - ribosomal 40s subunit protein s26a  RPS7A - ribosomal 40s subunit protein s7a  RPL16B - ribosomal 60s subunit protein l16b  RPS24A - ribosomal 40s subunit protein s24a |
| GO:0022627 | cytosolic small ribosomal subunit | 4.74E-19 | 7.19E-17 | 14.36 (5247,57,141,22) | [+] Show genes  RPS0B - ribosomal 40s subunit protein s0b  RPS27B - ribosomal 40s subunit protein s27b  RPS1B - ribosomal 40s subunit protein s1b  RPS23B - ribosomal 40s subunit protein s23b  RPS14A - ribosomal 40s subunit protein s14a  RPS18B - ribosomal 40s subunit protein s18b  RPS21B - rps21bp  RPS10A - ribosomal 40s subunit protein s10a  RPS10B - ribosomal 40s subunit protein s10b  RPS16A - ribosomal 40s subunit protein s16a  RPS15 - ribosomal 40s subunit protein s15  ASC1 - asc1p  RPS13 - ribosomal 40s subunit protein s13  RPS5 - rps5p  RPS22A - rps22ap  RPS8B - ribosomal 40s subunit protein s8b  RPS17A - ribosomal 40s subunit protein s17a  RPS26A - ribosomal 40s subunit protein s26a  RPS17B - ribosomal 40s subunit protein s17b  RPS7A - ribosomal 40s subunit protein s7a  RPS24A - ribosomal 40s subunit protein s24a  RPS28A - ribosomal 40s subunit protein s28a |
| GO:0043228 | non-membrane-bounded organelle | 3.33E-18 | 4.42E-16 | 2.79 (5247,972,155,80) | [+] Show genes  RPL2B - ribosomal 60s subunit protein l2b  RPS0B - ribosomal 40s subunit protein s0b  RPL29 - ribosomal 60s subunit protein l29  ECM16 - ecm16p  MGM101 - mgm101p  RPL24A - ribosomal 60s subunit protein l24a  VMA2 - vma2p  RPL30 - ribosomal 60s subunit protein l30  RRB1 - rrb1p  SED1 - sed1p  EFG1 - efg1p  RPL9A - ribosomal 60s subunit protein l9a  RPL40A - ubiquitin-ribosomal 60s subunit protein l40a fusion protein  RPL23B - ribosomal 60s subunit protein l23b  RPL13B - ribosomal 60s subunit protein l13b  RPL38 - ribosomal 60s subunit protein l38  TEF1 - tef1p  RPS16A - ribosomal 40s subunit protein s16a  RPL8A - ribosomal 60s subunit protein l8a  SLA1 - sla1p  UTP14 - utp14p  ILV5 - ketol-acid reductoisomerase  ASC1 - asc1p  RPS8B - ribosomal 40s subunit protein s8b  RPL26A - ribosomal 60s subunit protein l26a  RPL11A - ribosomal 60s subunit protein l11a  RPL15A - ribosomal 60s subunit protein l15a  RPP0 - ribosomal protein p0  RPL34B - ribosomal 60s subunit protein l34b  RPL27B - ribosomal 60s subunit protein l27b  RPL3 - ribosomal 60s subunit protein l3  RPS18B - ribosomal 40s subunit protein s18b  RPL19A - ribosomal 60s subunit protein l19a  RPS21B - rps21bp  RPS10B - ribosomal 40s subunit protein s10b  RPP2B - ribosomal protein p2b  FPR4 - peptidylprolyl isomerase fpr4  RLP7 - rlp7p  RPL6B - ribosomal 60s subunit protein l6b  RPS15 - ribosomal 40s subunit protein s15  RPP2A - ribosomal protein p2a  RPS5 - rps5p  RPS17A - ribosomal 40s subunit protein s17a  RPL37A - ribosomal 60s subunit protein l37a  RPL24B - ribosomal 60s subunit protein l24b  RPS28A - ribosomal 40s subunit protein s28a  RPS27B - ribosomal 40s subunit protein s27b  RPS1B - ribosomal 40s subunit protein s1b  RPL17A - ribosomal 60s subunit protein l17a  RPL21A - ribosomal 60s subunit protein l21a  RPP1B - ribosomal protein p1b  RPL14A - ribosomal 60s subunit protein l14a  RPL17B - rpl17bp  HCA4 - hca4p  RPL32 - ribosomal 60s subunit protein l32  MRPL24 - mitochondrial 54s ribosomal protein yml24/yml14  RPL26B - ribosomal 60s subunit protein l26b  RPL22B - ribosomal 60s subunit protein l22b  RPL35B - ribosomal 60s subunit protein l35b  RPS10A - ribosomal 40s subunit protein s10a  ACT1 - actin  NSR1 - nsr1p  RPL34A - ribosomal 60s subunit protein l34a  RPL27A - ribosomal 60s subunit protein l27a  RPS22A - rps22ap  RPS17B - ribosomal 40s subunit protein s17b  PAH1 - phosphatidate phosphatase pah1  RPL6A - ribosomal 60s subunit protein l6a  TPM1 - tpm1p  RPS23B - ribosomal 40s subunit protein s23b  RPL42B - ribosomal 60s subunit protein l42b  RPS14A - ribosomal 40s subunit protein s14a  RPL31A - ribosomal 60s subunit protein l31a  HSL1 - hsl1p  SUA5 - sua5p  RPS13 - ribosomal 40s subunit protein s13  RPS26A - ribosomal 40s subunit protein s26a  RPS7A - ribosomal 40s subunit protein s7a  RPS24A - ribosomal 40s subunit protein s24a  RPL16B - ribosomal 60s subunit protein l16b |
| GO:0043232 | intracellular non-membrane-bounded organelle | 3.33E-18 | 3.92E-16 | 2.79 (5247,972,155,80) | [+] Show genes  RPL2B - ribosomal 60s subunit protein l2b  RPS0B - ribosomal 40s subunit protein s0b  RPL29 - ribosomal 60s subunit protein l29  ECM16 - ecm16p  MGM101 - mgm101p  RPL24A - ribosomal 60s subunit protein l24a  VMA2 - vma2p  RPL30 - ribosomal 60s subunit protein l30  RRB1 - rrb1p  SED1 - sed1p  EFG1 - efg1p  RPL9A - ribosomal 60s subunit protein l9a  RPL40A - ubiquitin-ribosomal 60s subunit protein l40a fusion protein  RPL23B - ribosomal 60s subunit protein l23b  RPL13B - ribosomal 60s subunit protein l13b  RPL38 - ribosomal 60s subunit protein l38  TEF1 - tef1p  RPS16A - ribosomal 40s subunit protein s16a  RPL8A - ribosomal 60s subunit protein l8a  SLA1 - sla1p  UTP14 - utp14p  ILV5 - ketol-acid reductoisomerase  ASC1 - asc1p  RPS8B - ribosomal 40s subunit protein s8b  RPL26A - ribosomal 60s subunit protein l26a  RPL11A - ribosomal 60s subunit protein l11a  RPL15A - ribosomal 60s subunit protein l15a  RPP0 - ribosomal protein p0  RPL34B - ribosomal 60s subunit protein l34b  RPL27B - ribosomal 60s subunit protein l27b  RPL3 - ribosomal 60s subunit protein l3  RPS18B - ribosomal 40s subunit protein s18b  RPL19A - ribosomal 60s subunit protein l19a  RPS21B - rps21bp  RPS10B - ribosomal 40s subunit protein s10b  RPP2B - ribosomal protein p2b  FPR4 - peptidylprolyl isomerase fpr4  RLP7 - rlp7p  RPL6B - ribosomal 60s subunit protein l6b  RPS15 - ribosomal 40s subunit protein s15  RPP2A - ribosomal protein p2a  RPS5 - rps5p  RPS17A - ribosomal 40s subunit protein s17a  RPL37A - ribosomal 60s subunit protein l37a  RPL24B - ribosomal 60s subunit protein l24b  RPS28A - ribosomal 40s subunit protein s28a  RPS27B - ribosomal 40s subunit protein s27b  RPS1B - ribosomal 40s subunit protein s1b  RPL17A - ribosomal 60s subunit protein l17a  RPL21A - ribosomal 60s subunit protein l21a  RPP1B - ribosomal protein p1b  RPL14A - ribosomal 60s subunit protein l14a  RPL17B - rpl17bp  HCA4 - hca4p  RPL32 - ribosomal 60s subunit protein l32  MRPL24 - mitochondrial 54s ribosomal protein yml24/yml14  RPL26B - ribosomal 60s subunit protein l26b  RPL22B - ribosomal 60s subunit protein l22b  RPL35B - ribosomal 60s subunit protein l35b  RPS10A - ribosomal 40s subunit protein s10a  ACT1 - actin  NSR1 - nsr1p  RPL34A - ribosomal 60s subunit protein l34a  RPL27A - ribosomal 60s subunit protein l27a  RPS22A - rps22ap  RPS17B - ribosomal 40s subunit protein s17b  PAH1 - phosphatidate phosphatase pah1  RPL6A - ribosomal 60s subunit protein l6a  TPM1 - tpm1p  RPS23B - ribosomal 40s subunit protein s23b  RPL42B - ribosomal 60s subunit protein l42b  RPS14A - ribosomal 40s subunit protein s14a  RPL31A - ribosomal 60s subunit protein l31a  HSL1 - hsl1p  SUA5 - sua5p  RPS13 - ribosomal 40s subunit protein s13  RPS26A - ribosomal 40s subunit protein s26a  RPS7A - ribosomal 40s subunit protein s7a  RPS24A - ribosomal 40s subunit protein s24a  RPL16B - ribosomal 60s subunit protein l16b |
| GO:0015935 | small ribosomal subunit | 7.18E-14 | 7.62E-12 | 9.00 (5247,91,141,22) | [+] Show genes  RPS0B - ribosomal 40s subunit protein s0b  RPS1B - ribosomal 40s subunit protein s1b  RPS27B - ribosomal 40s subunit protein s27b  RPS23B - ribosomal 40s subunit protein s23b  RPS14A - ribosomal 40s subunit protein s14a  RPS18B - ribosomal 40s subunit protein s18b  RPS21B - rps21bp  RPS10A - ribosomal 40s subunit protein s10a  RPS10B - ribosomal 40s subunit protein s10b  RPS16A - ribosomal 40s subunit protein s16a  RPS15 - ribosomal 40s subunit protein s15  ASC1 - asc1p  RPS13 - ribosomal 40s subunit protein s13  RPS5 - rps5p  RPS22A - rps22ap  RPS8B - ribosomal 40s subunit protein s8b  RPS17A - ribosomal 40s subunit protein s17a  RPS26A - ribosomal 40s subunit protein s26a  RPS17B - ribosomal 40s subunit protein s17b  RPS7A - ribosomal 40s subunit protein s7a  RPS24A - ribosomal 40s subunit protein s24a  RPS28A - ribosomal 40s subunit protein s28a |
| GO:0030684 | preribosome | 2.98E-12 | 2.87E-10 | 2.18 (5247,154,1201,77) | [+] Show genes  NSA2 - nsa2p  ECM16 - ecm16p  LCP5 - lcp5p  RPS3 - ribosomal 40s subunit protein s3  EFG1 - efg1p  UTP30 - utp30p  RRP1 - rrp1p  KRR1 - krr1p  RPF1 - rpf1p  RPS16A - ribosomal 40s subunit protein s16a  NOP16 - nop16p  NOP7 - nop7p  UTP10 - utp10p  UTP14 - utp14p  REI1 - rei1p  BUD22 - bud22p  NUG1 - nug1p  SSF1 - ssf1p  NOC2 - noc2p  KRI1 - kri1p  RPS8B - ribosomal 40s subunit protein s8b  UTP20 - utp20p  RPP0 - ribosomal protein p0  RPL27B - ribosomal 60s subunit protein l27b  RPS1A - ribosomal 40s subunit protein s1a  CIC1 - cic1p  RPS6A - ribosomal 40s subunit protein s6a  BUD20 - bud20p  UTP8 - utp8p  UTP9 - utp9p  RLP7 - rlp7p  NSA1 - nsa1p  CBF5 - pseudouridine synthase cbf5  RPS2 - ribosomal 40s subunit protein s2  BUD21 - bud21p  RPS5 - rps5p  MPP10 - mpp10p  RPL37A - ribosomal 60s subunit protein l37a  ENP2 - enp2p  YBL028C - hypothetical protein  RPS9A - ribosomal 40s subunit protein s9a  NOP58 - nop58p  RPL17A - ribosomal 60s subunit protein l17a  RPS1B - ribosomal 40s subunit protein s1b  RPL25 - ribosomal 60s subunit protein l25  RPL17B - rpl17bp  HCA4 - hca4p  NIP7 - nip7p  RRS1 - rrs1p  RPL35B - ribosomal 60s subunit protein l35b  PNO1 - pno1p  RPL34A - ribosomal 60s subunit protein l34a  RPS11B - ribosomal 40s subunit protein s11b  RPS14B - rps14bp  BMS1 - bms1p  EBP2 - ebp2p  SAS10 - sas10p  DRS1 - drs1p  YTM1 - ytm1p  NOP14 - nop14p  NOG2 - nog2p  RPS14A - ribosomal 40s subunit protein s14a  UTP5 - utp5p  SLX9 - slx9p  ECM1 - ecm1p  RIO1 - rio1p  HAS1 - atp-dependent rna helicase has1  ROK1 - rna-dependent atpase rok1  UTP11 - utp11p  RPS0A - ribosomal 40s subunit protein s0a  MAK21 - mak21p  RRP12 - rrp12p  RPS13 - ribosomal 40s subunit protein s13  RIO2 - rio2p  RPS7A - ribosomal 40s subunit protein s7a  MRD1 - mrd1p  PWP2 - pwp2p |
| GO:0030686 | 90S preribosome | 1.11E-6 | 9.78E-5 | 2.43 (5247,76,993,35) | [+] Show genes  ECM16 - ecm16p  RPS1B - ribosomal 40s subunit protein s1b  RPS3 - ribosomal 40s subunit protein s3  UTP30 - utp30p  KRR1 - krr1p  PNO1 - pno1p  RPS16A - ribosomal 40s subunit protein s16a  UTP10 - utp10p  RPS11B - ribosomal 40s subunit protein s11b  BUD22 - bud22p  KRI1 - kri1p  BMS1 - bms1p  RPS8B - ribosomal 40s subunit protein s8b  UTP20 - utp20p  RPP0 - ribosomal protein p0  NOP14 - nop14p  RPS14A - ribosomal 40s subunit protein s14a  RPS1A - ribosomal 40s subunit protein s1a  CIC1 - cic1p  RPS6A - ribosomal 40s subunit protein s6a  SLX9 - slx9p  HAS1 - atp-dependent rna helicase has1  UTP8 - utp8p  UTP9 - utp9p  RPS0A - ribosomal 40s subunit protein s0a  CBF5 - pseudouridine synthase cbf5  RRP12 - rrp12p  BUD21 - bud21p  RPS13 - ribosomal 40s subunit protein s13  RPS5 - rps5p  MPP10 - mpp10p  RPS7A - ribosomal 40s subunit protein s7a  RPS9A - ribosomal 40s subunit protein s9a  MRD1 - mrd1p  PWP2 - pwp2p |
| GO:0005730 | nucleolus | 1.69E-6 | 1.38E-4 | 1.47 (5247,268,1635,123) | [+] Show genes  ECM16 - ecm16p  NSA2 - nsa2p  LCP5 - lcp5p  FAF1 - faf1p  SIR2 - sir2p  RRB1 - rrb1p  UTP30 - utp30p  LHP1 - lhp1p  KRR1 - krr1p  NOP7 - nop7p  UTP10 - utp10p  UTP14 - utp14p  KRI1 - kri1p  RPA190 - rpa190p  TRF5 - non-canonical poly(a) polymerase trf5  GRC3 - grc3p  RPS6A - ribosomal 40s subunit protein s6a  UTP8 - utp8p  FPR4 - peptidylprolyl isomerase fpr4  RLP7 - rlp7p  UTP9 - utp9p  NSA1 - nsa1p  RPF2 - rpf2p  CBF5 - pseudouridine synthase cbf5  YNL022C - hypothetical protein  RPS2 - ribosomal 40s subunit protein s2  BUD21 - bud21p  RPA34 - rpa34p  MAF1 - maf1p  JIP5 - jip5p  ENP2 - enp2p  RPS9A - ribosomal 40s subunit protein s9a  NOP58 - nop58p  RRP36 - rrp36p  LRS4 - lrs4p  HCA4 - hca4p  RPB9 - rpb9p  RRS1 - rrs1p  MUM2 - mum2p  NSR1 - nsr1p  MAK11 - mak11p  RPA14 - rpa14p  DBP9 - dbp9p  EBP2 - ebp2p  RRN11 - rrn11p  DRS1 - drs1p  YTM1 - ytm1p  RCL1 - rcl1p  IFH1 - ifh1p  RPC10 - rpc10p  NOG2 - nog2p  RPS14A - ribosomal 40s subunit protein s14a  UTP5 - utp5p  FPR3 - peptidylprolyl isomerase fpr3  ECM1 - ecm1p  TGS1 - tgs1p  YIL096C - hypothetical protein  SPO12 - spo12p  BUD23 - bud23p  RPC40 - rpc40p  NOP53 - nop53p  TRM112 - trm112p  DBP7 - dbp7p  MRD1 - mrd1p  PWP2 - pwp2p  NCL1 - ncl1p  EFG1 - efg1p  RRN9 - rrn9p  ZUO1 - zuo1p  RRP1 - rrp1p  UBP10 - ubp10p  RPF1 - rpf1p  PXR1 - pxr1p  RSA4 - rsa4p  RRP8 - rrp8p  NOP16 - nop16p  GAR1 - gar1p  BUD22 - bud22p  NUG1 - nug1p  SSF1 - ssf1p  DHR2 - dhr2p  BRX1 - brx1p  NOC2 - noc2p  ARX1 - arx1p  UTP20 - utp20p  LSM7 - lsm7p  CIC1 - cic1p  ULS1 - uls1p  RRP3 - rna-dependent atpase rrp3  SDA1 - sda1p  CTK2 - ctk2p  ESF1 - esf1p  NOP9 - nop9p  FYV7 - fyv7p  EDC2 - edc2p  MPP10 - mpp10p  YBL028C - hypothetical protein  HRR25 - hrr25p  RPC19 - rpc19p  RPL7B - ribosomal 60s subunit protein l7b  NOP15 - nop15p  NIP7 - nip7p  PNO1 - pno1p  POL5 - pol5p  RPS14B - rps14bp  BMS1 - bms1p  SAS10 - sas10p  NOP14 - nop14p  ACS2 - acetate--coa ligase acs2  SHU1 - shu1p  SLX9 - slx9p  AIR2 - air2p  HAS1 - atp-dependent rna helicase has1  ROK1 - rna-dependent atpase rok1  UTP11 - utp11p  YBR141C - hypothetical protein  MAK21 - mak21p  RRP12 - rrp12p  RPS13 - ribosomal 40s subunit protein s13  RRP46 - rrp46p  RPA12 - rpa12p  RAD27 - rad27p  RPS7A - ribosomal 40s subunit protein s7a |
| GO:0005737 | cytoplasm | 2.29E-6 | 1.74E-4 | 1.48 (5247,2232,160,101) | [+] Show genes  RPL2B - ribosomal 60s subunit protein l2b  RPL29 - ribosomal 60s subunit protein l29  VMA2 - vma2p  BOP3 - bop3p  TSL1 - tsl1p  RPL9A - ribosomal 60s subunit protein l9a  RPL23B - ribosomal 60s subunit protein l23b  RPL13B - ribosomal 60s subunit protein l13b  RPS16A - ribosomal 40s subunit protein s16a  RPL8A - ribosomal 60s subunit protein l8a  ASC1 - asc1p  POR1 - por1p  PPT1 - ppt1p  RPS8B - ribosomal 40s subunit protein s8b  RPL34B - ribosomal 60s subunit protein l34b  RPL27B - ribosomal 60s subunit protein l27b  RPL3 - ribosomal 60s subunit protein l3  YER156C - hypothetical protein  RPL19A - ribosomal 60s subunit protein l19a  RPS21B - rps21bp  RPL6B - ribosomal 60s subunit protein l6b  RPS5 - rps5p  RPL37A - ribosomal 60s subunit protein l37a  YRB1 - yrb1p  RPL24B - ribosomal 60s subunit protein l24b  RPS1B - ribosomal 40s subunit protein s1b  RPL17A - ribosomal 60s subunit protein l17a  RPL14A - ribosomal 60s subunit protein l14a  BUD14 - bud14p  YDR161W - hypothetical protein  RPS10A - ribosomal 40s subunit protein s10a  RTS3 - rts3p  PGK1 - phosphoglycerate kinase  ABZ1 - 4-amino-4-deoxychorismate synthase  RPL34A - ribosomal 60s subunit protein l34a  SSE1 - sse1p  SRD1 - srd1p  RNR4 - ribonucleotide-diphosphate reductase subunit rnr4  RPS17B - ribosomal 40s subunit protein s17b  PAH1 - phosphatidate phosphatase pah1  RPL6A - ribosomal 60s subunit protein l6a  RPS23B - ribosomal 40s subunit protein s23b  RPL42B - ribosomal 60s subunit protein l42b  RPS14A - ribosomal 40s subunit protein s14a  RPL31A - ribosomal 60s subunit protein l31a  ECM22 - ecm22p  BNA5 - kynureninase  CHS3 - chitin synthase chs3  THR4 - threonine synthase thr4  RPS24A - ribosomal 40s subunit protein s24a  RPS0B - ribosomal 40s subunit protein s0b  RPL24A - ribosomal 60s subunit protein l24a  RPL30 - ribosomal 60s subunit protein l30  RPL40A - ubiquitin-ribosomal 60s subunit protein l40a fusion protein  TMA10 - tma10p  RPL38 - ribosomal 60s subunit protein l38  TEF1 - tef1p  SLA1 - sla1p  RPL26A - ribosomal 60s subunit protein l26a  RPL15A - ribosomal 60s subunit protein l15a  RPL11A - ribosomal 60s subunit protein l11a  RPP0 - ribosomal protein p0  FUM1 - fumarase fum1  RPS18B - ribosomal 40s subunit protein s18b  RPS10B - ribosomal 40s subunit protein s10b  SMB1 - smb1p  RPP2B - ribosomal protein p2b  RPS15 - ribosomal 40s subunit protein s15  RPP2A - ribosomal protein p2a  DUR1,2 - bifunctional urea carboxylase/allophanate hydrolase  RPS17A - ribosomal 40s subunit protein s17a  RPS28A - ribosomal 40s subunit protein s28a  RPS27B - ribosomal 40s subunit protein s27b  PMU1 - pmu1p  RPP1B - ribosomal protein p1b  RPL21A - ribosomal 60s subunit protein l21a  RPL17B - rpl17bp  RPL32 - ribosomal 60s subunit protein l32  RPL26B - ribosomal 60s subunit protein l26b  RPL22B - ribosomal 60s subunit protein l22b  ARF2 - arf2p  SLT2 - slt2p  RPL35B - ribosomal 60s subunit protein l35b  ACT1 - actin  NCS2 - ncs2p  RPL27A - ribosomal 60s subunit protein l27a  RPS22A - rps22ap  FRA2 - fra2p  NEW1 - new1p  YHR020W - proline--trna ligase  TPM1 - tpm1p  APA2 - apa2p  STM1 - stm1p  SUA5 - sua5p  HSL1 - hsl1p  RIO1 - rio1p  RPS13 - ribosomal 40s subunit protein s13  MIC23 - mic23p  RPS26A - ribosomal 40s subunit protein s26a  RPS7A - ribosomal 40s subunit protein s7a  RPL16B - ribosomal 60s subunit protein l16b |
| GO:0044444 | cytoplasmic part | 2.58E-6 | 1.82E-4 | 1.35 (5247,3045,145,114) | [+] Show genes  RPL2B - ribosomal 60s subunit protein l2b  YLR194C - hypothetical protein  ECM16 - ecm16p  RPL29 - ribosomal 60s subunit protein l29  MGM101 - mgm101p  VMA2 - vma2p  PFA3 - pfa3p  KTR2 - ktr2p  TSL1 - tsl1p  MSS51 - mss51p  RPL23B - ribosomal 60s subunit protein l23b  RPL13B - ribosomal 60s subunit protein l13b  RPS16A - ribosomal 40s subunit protein s16a  RPL8A - ribosomal 60s subunit protein l8a  ASC1 - asc1p  PPT1 - ppt1p  RPS8B - ribosomal 40s subunit protein s8b  ATP18 - atp18p  RPL34B - ribosomal 60s subunit protein l34b  RPL27B - ribosomal 60s subunit protein l27b  RPL3 - ribosomal 60s subunit protein l3  CYT1 - ubiquinol--cytochrome-c reductase catalytic subunit cyt1  RPL19A - ribosomal 60s subunit protein l19a  MFA1 - mfa1p  RPS21B - rps21bp  FLO11 - flo11p  RPL6B - ribosomal 60s subunit protein l6b  RLP7 - rlp7p  TCB1 - tcb1p  RPS5 - rps5p  RPL37A - ribosomal 60s subunit protein l37a  HSP150 - hsp150p  YKE4 - zn(2+) transporter yke4  RPL24B - ribosomal 60s subunit protein l24b  CIS3 - cis3p  RPS1B - ribosomal 40s subunit protein s1b  RPL17A - ribosomal 60s subunit protein l17a  RPL14A - ribosomal 60s subunit protein l14a  MRPL24 - mitochondrial 54s ribosomal protein yml24/yml14  RPS10A - ribosomal 40s subunit protein s10a  NSR1 - nsr1p  PGK1 - phosphoglycerate kinase  RPL34A - ribosomal 60s subunit protein l34a  SSE1 - sse1p  RPS17B - ribosomal 40s subunit protein s17b  PAH1 - phosphatidate phosphatase pah1  RPL6A - ribosomal 60s subunit protein l6a  PRM4 - prm4p  AGA2 - aga2p  RPS23B - ribosomal 40s subunit protein s23b  RPL42B - ribosomal 60s subunit protein l42b  RPS14A - ribosomal 40s subunit protein s14a  RPL31A - ribosomal 60s subunit protein l31a  PRM5 - prm5p  ECM22 - ecm22p  CHS3 - chitin synthase chs3  PRY2 - pry2p  RPS24A - ribosomal 40s subunit protein s24a  RPS0B - ribosomal 40s subunit protein s0b  RPL30 - ribosomal 60s subunit protein l30  SED1 - sed1p  RPL40A - ubiquitin-ribosomal 60s subunit protein l40a fusion protein  CHS1 - chitin synthase chs1  RPL38 - ribosomal 60s subunit protein l38  TEF1 - tef1p  SCW4 - scw4p  SLA1 - sla1p  ILV5 - ketol-acid reductoisomerase  HOR7 - hor7p  RPL26A - ribosomal 60s subunit protein l26a  RPL15A - ribosomal 60s subunit protein l15a  RPL11A - ribosomal 60s subunit protein l11a  RPP0 - ribosomal protein p0  FUM1 - fumarase fum1  DFG5 - dfg5p  RPS18B - ribosomal 40s subunit protein s18b  RPS10B - ribosomal 40s subunit protein s10b  RPP2B - ribosomal protein p2b  RPS15 - ribosomal 40s subunit protein s15  RPP2A - ribosomal protein p2a  RPS17A - ribosomal 40s subunit protein s17a  RPS28A - ribosomal 40s subunit protein s28a  RPS27B - ribosomal 40s subunit protein s27b  STF1 - stf1p  RPP1B - ribosomal protein p1b  RPL21A - ribosomal 60s subunit protein l21a  RPL17B - rpl17bp  RPL32 - ribosomal 60s subunit protein l32  RPL26B - ribosomal 60s subunit protein l26b  RPL22B - ribosomal 60s subunit protein l22b  ARF2 - arf2p  SLT2 - slt2p  PRB1 - prb1p  QCR8 - qcr8p  FMP33 - fmp33p  ACT1 - actin  RPL27A - ribosomal 60s subunit protein l27a  RPS22A - rps22ap  FRA2 - fra2p  NEW1 - new1p  TPM1 - tpm1p  STM1 - stm1p  HSL1 - hsl1p  RIO1 - rio1p  PST1 - pst1p  DLD1 - dld1p  CWP2 - cwp2p  UIP4 - uip4p  RPS13 - ribosomal 40s subunit protein s13  MIC23 - mic23p  RPS26A - ribosomal 40s subunit protein s26a  RPS7A - ribosomal 40s subunit protein s7a  RPL16B - ribosomal 60s subunit protein l16b  YET2 - yet2p |
| GO:0005576 | extracellular region | 3.41E-6 | 2.26E-4 | 1.76 (5247,99,1742,58) | [+] Show genes  SED1 - sed1p  PPA2 - ppa2p  EGT2 - egt2p  FYV5 - fyv5p  TIR3 - tir3p  BGL2 - bgl2p  SCW4 - scw4p  YLR042C - hypothetical protein  YNL190W - hypothetical protein  SIM1 - sim1p  YGP1 - ygp1p  DAN4 - dan4p  TIR1 - tir1p  FIG2 - fig2p  PLB1 - plb1p  TOS6 - tos6p  DSE4 - dse4p  YIL169C - hypothetical protein  MFA1 - mfa1p  SSA2 - hsp70 family chaperone ssa2  MSB2 - msb2p  FLO11 - flo11p  TIP1 - tip1p  FLO1 - flo1p  HSP150 - hsp150p  SUC2 - beta-fructofuranosidase suc2  CIS3 - cis3p  YPS1 - yps1p  ACB1 - long-chain fatty acid transporter acb1  GAS5 - gas5p  YBR056W - 17-beta-hydroxysteroid dehydrogenase-like protein  RNY1 - rny1p  PST2 - pst2p  PIR1 - pir1p  ZPS1 - zps1p  HPF1 - hpf1p  APE2 - ape2p  SCW10 - scw10p  YOL159C - hypothetical protein  PLB3 - plb3p  CTS1 - cts1p  DSE2 - dse2p  UTR2 - utr2p  FLO9 - flo9p  CRH1 - crh1p  PST1 - pst1p  CWP1 - cwp1p  CWP2 - cwp2p  TOS1 - tos1p  TIR4 - tir4p  TIR2 - tir2p  SUN4 - sun4p  PRY2 - pry2p  IPP1 - ipp1p  YOR389W - hypothetical protein  AGA1 - aga1p  YPS3 - yps3p  FIT3 - fit3p |
| GO:0030687 | preribosome, large subunit precursor | 5.39E-6 | 3.36E-4 | 2.43 (5247,59,1097,30) | [+] Show genes  NSA2 - nsa2p  RPL17A - ribosomal 60s subunit protein l17a  RPL25 - ribosomal 60s subunit protein l25  RPL17B - rpl17bp  RRS1 - rrs1p  NIP7 - nip7p  RRP1 - rrp1p  RPL35B - ribosomal 60s subunit protein l35b  RPF1 - rpf1p  NOP16 - nop16p  NOP7 - nop7p  REI1 - rei1p  RPL34A - ribosomal 60s subunit protein l34a  NUG1 - nug1p  SSF1 - ssf1p  NOC2 - noc2p  EBP2 - ebp2p  DRS1 - drs1p  YTM1 - ytm1p  NOG2 - nog2p  RPL27B - ribosomal 60s subunit protein l27b  CIC1 - cic1p  ECM1 - ecm1p  HAS1 - atp-dependent rna helicase has1  BUD20 - bud20p  RLP7 - rlp7p  NSA1 - nsa1p  MAK21 - mak21p  RPL37A - ribosomal 60s subunit protein l37a  YBL028C - hypothetical protein |
| GO:0030312 | external encapsulating structure | 1.07E-5 | 6.29E-4 | 7.64 (5247,104,66,10) | [+] Show genes  YLR194C - hypothetical protein  AGA2 - aga2p  SED1 - sed1p  PAU7 - pau7p  FIG2 - fig2p  FLO11 - flo11p  HSP150 - hsp150p  CWP2 - cwp2p  SCW4 - scw4p  CIS3 - cis3p |
| GO:0005618 | cell wall | 1.07E-5 | 5.96E-4 | 7.64 (5247,104,66,10) | [+] Show genes  YLR194C - hypothetical protein  AGA2 - aga2p  SED1 - sed1p  PAU7 - pau7p  FIG2 - fig2p  FLO11 - flo11p  HSP150 - hsp150p  CWP2 - cwp2p  SCW4 - scw4p  CIS3 - cis3p |
| GO:0032991 | protein-containing complex | 1.94E-5 | 1.03E-3 | 1.48 (5247,2077,159,93) | [+] Show genes  RPL2B - ribosomal 60s subunit protein l2b  RPS0B - ribosomal 40s subunit protein s0b  ECM16 - ecm16p  RPL29 - ribosomal 60s subunit protein l29  RPL24A - ribosomal 60s subunit protein l24a  RPL30 - ribosomal 60s subunit protein l30  VMA2 - vma2p  SED1 - sed1p  EFG1 - efg1p  TSL1 - tsl1p  RPL9A - ribosomal 60s subunit protein l9a  RPL40A - ubiquitin-ribosomal 60s subunit protein l40a fusion protein  RPL23B - ribosomal 60s subunit protein l23b  RPL38 - ribosomal 60s subunit protein l38  RPL13B - ribosomal 60s subunit protein l13b  RPS16A - ribosomal 40s subunit protein s16a  TEF1 - tef1p  RPL8A - ribosomal 60s subunit protein l8a  SLA1 - sla1p  UTP14 - utp14p  ASC1 - asc1p  POR1 - por1p  RPS8B - ribosomal 40s subunit protein s8b  ATP18 - atp18p  DPB3 - dpb3p  RPL26A - ribosomal 60s subunit protein l26a  IRE1 - ire1p  RPL15A - ribosomal 60s subunit protein l15a  RPL11A - ribosomal 60s subunit protein l11a  RPP0 - ribosomal protein p0  FUM1 - fumarase fum1  RPL34B - ribosomal 60s subunit protein l34b  RPL27B - ribosomal 60s subunit protein l27b  RPL3 - ribosomal 60s subunit protein l3  CYT1 - ubiquinol--cytochrome-c reductase catalytic subunit cyt1  RPS18B - ribosomal 40s subunit protein s18b  RPL19A - ribosomal 60s subunit protein l19a  RPS21B - rps21bp  SLX1 - slx1p  SMB1 - smb1p  RPS10B - ribosomal 40s subunit protein s10b  RPP2B - ribosomal protein p2b  RLP7 - rlp7p  RPL6B - ribosomal 60s subunit protein l6b  RPS15 - ribosomal 40s subunit protein s15  RPP2A - ribosomal protein p2a  RPS5 - rps5p  RPL37A - ribosomal 60s subunit protein l37a  RPS17A - ribosomal 40s subunit protein s17a  YRB1 - yrb1p  GSC2 - gsc2p  RPL24B - ribosomal 60s subunit protein l24b  RPS28A - ribosomal 40s subunit protein s28a  RPL17A - ribosomal 60s subunit protein l17a  RPS1B - ribosomal 40s subunit protein s1b  RPS27B - ribosomal 40s subunit protein s27b  RPL14A - ribosomal 60s subunit protein l14a  RPL21A - ribosomal 60s subunit protein l21a  NRG2 - nrg2p  RPP1B - ribosomal protein p1b  RPL17B - rpl17bp  BUD14 - bud14p  HCA4 - hca4p  RPL32 - ribosomal 60s subunit protein l32  MRPL24 - mitochondrial 54s ribosomal protein yml24/yml14  RPL26B - ribosomal 60s subunit protein l26b  RPL22B - ribosomal 60s subunit protein l22b  QCR8 - qcr8p  RPL35B - ribosomal 60s subunit protein l35b  RPS10A - ribosomal 40s subunit protein s10a  ACT1 - actin  NSR1 - nsr1p  RPL34A - ribosomal 60s subunit protein l34a  SSE1 - sse1p  RPL27A - ribosomal 60s subunit protein l27a  RNR4 - ribonucleotide-diphosphate reductase subunit rnr4  RPS22A - rps22ap  RPS17B - ribosomal 40s subunit protein s17b  NEW1 - new1p  YHR020W - proline--trna ligase  RPL6A - ribosomal 60s subunit protein l6a  RPC37 - rpc37p  RPS23B - ribosomal 40s subunit protein s23b  RPL42B - ribosomal 60s subunit protein l42b  RPS14A - ribosomal 40s subunit protein s14a  RPL31A - ribosomal 60s subunit protein l31a  STM1 - stm1p  RIO1 - rio1p  RPS13 - ribosomal 40s subunit protein s13  RPS26A - ribosomal 40s subunit protein s26a  RPS7A - ribosomal 40s subunit protein s7a  RPS24A - ribosomal 40s subunit protein s24a  RPL16B - ribosomal 60s subunit protein l16b |
| GO:0009277 | fungal-type cell wall | 3.41E-5 | 1.72E-3 | 35.33 (5247,99,6,4) | [+] Show genes  YLR194C - hypothetical protein  SED1 - sed1p  SCW4 - scw4p  CIS3 - cis3p |
| GO:0045009 | chitosome | 8.65E-5 | 4.17E-3 | 149.91 (5247,2,35,2) | [+] Show genes  CHS3 - chitin synthase chs3  CHS1 - chitin synthase chs1 |
| GO:0045275 | respiratory chain complex III | 4.28E-4 | 1.98E-2 | 5.17 (5247,10,710,7) | [+] Show genes  RIP1 - ubiquinol--cytochrome-c reductase catalytic subunit rip1  QCR7 - ubiquinol--cytochrome-c reductase subunit 7  CYT1 - ubiquinol--cytochrome-c reductase catalytic subunit cyt1  QCR2 - ubiquinol--cytochrome-c reductase subunit 2  QCR6 - ubiquinol--cytochrome-c reductase subunit 6  QCR8 - qcr8p  QCR10 - ubiquinol--cytochrome-c reductase subunit 10 |
| GO:0005750 | mitochondrial respiratory chain complex III | 4.28E-4 | 1.89E-2 | 5.17 (5247,10,710,7) | [+] Show genes  RIP1 - ubiquinol--cytochrome-c reductase catalytic subunit rip1  QCR7 - ubiquinol--cytochrome-c reductase subunit 7  CYT1 - ubiquinol--cytochrome-c reductase catalytic subunit cyt1  QCR6 - ubiquinol--cytochrome-c reductase subunit 6  QCR2 - ubiquinol--cytochrome-c reductase subunit 2  QCR8 - qcr8p  QCR10 - ubiquinol--cytochrome-c reductase subunit 10 |
| GO:0031225 | anchored component of membrane | 4.6E-4 | 1.95E-2 | 2.69 (5247,60,618,19) | [+] Show genes  YLR194C - hypothetical protein  SED1 - sed1p  DFG5 - dfg5p  GAS5 - gas5p  EGT2 - egt2p  PST1 - pst1p  TIR3 - tir3p  FLO11 - flo11p  CWP2 - cwp2p  TIR4 - tir4p  HPF1 - hpf1p  YDR524C-B - hypothetical protein  AGA1 - aga1p  PLB1 - plb1p  FIG2 - fig2p  TIP1 - tip1p  FLO1 - flo1p  YPS3 - yps3p  FIT3 - fit3p |

Species used: Saccharomyces cerevisiae

The system has recognized 5321 genes out of 5595 gene terms entered by the user.  
 628 genes were recognized by gene symbol and 4693 genes by other gene IDs .  
Only 5247 of these genes are associated with a GO term.

  
 Output in Microsoft Excel format  

% List genereted using GOrilla
% http://cbl-gorilla.cs.technion.ac.il/
% GO term pValue
GO:0044445 1.54E-46
GO:0044391 7.13E-46
GO:0005840 1.58E-45
GO:0022625 8.74E-42
GO:0015934 1.95E-31
GO:1990904 4.12E-28
GO:0022627 4.74E-19
GO:0043228 3.33E-18
GO:0043232 3.33E-18
GO:0015935 7.18E-14
GO:0030684 2.98E-12
GO:0030686 1.11E-6
GO:0005730 1.69E-6
GO:0005737 2.29E-6
GO:0044444 2.58E-6
GO:0005576 3.41E-6
GO:0030687 5.39E-6
GO:0030312 1.07E-5
GO:0005618 1.07E-5
GO:0032991 1.94E-5
GO:0009277 3.41E-5
GO:0045009 8.65E-5
GO:0045275 4.28E-4
GO:0005750 4.28E-4
GO:0031225 4.6E-4
 Visualize output in REViGO   

The GOrilla database is periodically updated using the GO database and other sources.  
The GOrilla database was last updated on Sep 21, 2019

This results page will be available on this site for one month from now (until
Oct 23, 2019
). You can bookmark this page and come back to it later.

  
**'P-value'** is the enrichment
p-value computed according to the mHG or HG model. This p-value is not
corrected for multiple testing of 1061 GO terms.  
  
**'FDR q-value'** is the correction of the above p-value for multiple testing using the Benjamini and Hochberg (1995) method.   
Namely, for the i^th^ term (ranked according to p-value) the FDR q-value is (p-value \* number of GO terms) / i.   
  
**Enrichment (N, B, n, b)** is defined as follows:  
N - is the total number of genes  
B - is the total number of genes associated with a specific GO term  
n - is the number of genes in the top of the user's input list or in the target set when appropriate  
b - is the number of genes in the intersection  
Enrichment = (b/n) / (B/N)  
  
**Genes:** For each GO term you can see the list of associated genes that appear in the optimal top of the list.  
Each gene name is specified by gene symbol followed by a short description of the gene   

Back to GOrilla analysis results
