## Supplemental Data S1 for "A simple mass-action model predicts genome-wide protein timecourses from mRNA trajectories during a dynamic response in two strains of *Saccharomyces cerevisiae*": GOFUNCTION_par.html

Results


 **Parameters** 

*P-value color scale*

|  |  |  |  |  |
| --- | --- | --- | --- | --- |
| > 10^-3^ | 10^-3^ to 10^-5^ | 10^-5^ to 10^-7^ | 10^-7^ to 10^-9^ | < 10^-9^ |


|  |  |  |  |  |  |
| --- | --- | --- | --- | --- | --- |
| **GO term** | **Description** | **P-value** | **FDR q-value** | **Enrichment (N, B, n, b)** | **Genes** |
| GO:0003824 | catalytic activity | 1.36E-9 | 2.55E-6 | 1.43 (2298,1204,190,142) | [+] Show genes  IDH1 - isocitrate dehydrogenase (nad(+)) idh1  PHS1 - phs1p  RSP5 - nedd4 family e3 ubiquitin-protein ligase  SIT4 - sit4p  ENO2 - phosphopyruvate hydratase eno2  TSL1 - tsl1p  ARO2 - bifunctional chorismate synthase/riboflavin reductase [nad(p)h] aro2  MYO4 - myosin 4  GPD1 - glycerol-3-phosphate dehydrogenase (nad(+)) gpd1  APA1 - apa1p  ASN2 - asparagine synthase (glutamine-hydrolyzing) 2  YSA1 - ysa1p  GCD6 - gcd6p  GLK1 - glucokinase  ARP2 - actin-related protein 2  CPR6 - peptidylprolyl isomerase cpr6  EMI2 - putative glucokinase  FRA1 - fra1p  PGM2 - phosphoglucomutase pgm2  HIS4 - trifunctional histidinol dehydrogenase/phosphoribosyl-amp cyclohydrolase/phosphoribosyl-atp diphosphatase  QCR2 - ubiquinol--cytochrome-c reductase subunit 2  CIT1 - citrate (si)-synthase cit1  HSP104 - chaperone atpase hsp104  ILV2 - acetolactate synthase catalytic subunit  GLR1 - glutathione-disulfide reductase glr1  SSA2 - hsp70 family chaperone ssa2  RSM26 - rsm26p  MAS1 - mas1p  HFD1 - hfd1p  NFS1 - nfs1p  GLC7 - glc7p  ATP2 - atp2p  CAR2 - ornithine-oxo-acid transaminase  THI4 - thi4p  ILV6 - acetolactate synthase regulatory subunit  HSP150 - hsp150p  NTH1 - alpha,alpha-trehalase nth1  VPS21 - vps21p  RHO1 - rho1p  ARG5,6 - bifunctional acetylglutamate kinase/n-acetyl-gamma-glutamyl-phosphate reductase  CAB2 - phosphopantothenate--cysteine ligase cab2  GPH1 - gph1p  GSY2 - glycogen (starch) synthase gsy2  HOR2 - glycerol-1-phosphatase hor2  GRE3 - trifunctional aldehyde reductase/xylose reductase/glucose 1-dehydrogenase (nadp(+))  DCS1 - dcs1p  AIM2 - aim2p  ASN1 - asparagine synthase (glutamine-hydrolyzing) 1  GUK1 - guanylate kinase  HOM3 - aspartate kinase  APE2 - ape2p  ZWF1 - glucose-6-phosphate dehydrogenase  ATP3 - atp3p  GDE1 - gde1p  HSC82 - hsp90 family chaperone hsc82  GET3 - guanine nucleotide exchange factor get3  PEP4 - pep4p  PNC1 - nicotinamidase  MDH3 - malate dehydrogenase mdh3  YGL039W - carbonyl reductase (nadph-dependent)  OLA1 - ola1p  HEM2 - porphobilinogen synthase hem2  YDL086W - carboxymethylenebutenolidase  TRP2 - anthranilate synthase trp2  ERG13 - hydroxymethylglutaryl-coa synthase  BDH1 - (r,r)-butanediol dehydrogenase  BNA5 - kynureninase  YIL108W - putative metalloendopeptidase  CDC60 - leucine--trna ligase cdc60  ALD2 - aldehyde dehydrogenase (nad(+)) ald2  SDH2 - succinate dehydrogenase iron-sulfur protein subunit sdh2  OLE1 - stearoyl-coa 9-desaturase  NDE1 - nadh-ubiquinone reductase (h(+)-translocating) nde1  DNM1 - dnm1p  ALD5 - aldehyde dehydrogenase (nad(p)(+)) ald5  HSP82 - hsp90 family chaperone hsp82  TRX3 - trx3p  CWH41 - cwh41p  ACH1 - ach1p  ILV3 - ilv3p  BGL2 - bgl2p  PDC1 - indolepyruvate decarboxylase 1  GLO1 - lactoylglutathione lyase glo1  SCW4 - scw4p  ILV5 - ketol-acid reductoisomerase  GRX3 - grx3p  ERG7 - lanosterol synthase erg7  ENO1 - phosphopyruvate hydratase eno1  UBC13 - e2 ubiquitin-conjugating protein ubc13  ADE16 - bifunctional phosphoribosylaminoimidazolecarboxamide formyltransferase/imp cyclohydrolase ade16  MDH1 - malate dehydrogenase mdh1  ARO4 - 3-deoxy-7-phosphoheptulonate synthase aro4  ISC1 - inositol phosphosphingolipid phospholipase  FUM1 - fumarase fum1  AFG3 - aaa family atpase afg3  ARG1 - argininosuccinate synthase  GLN1 - glutamate--ammonia ligase  PCS60 - pcs60p  COR1 - ubiquinol--cytochrome-c reductase subunit cor1  UGP1 - utp glucose-1-phosphate uridylyltransferase  AMD1 - amp deaminase  GAS3 - gas3p  DCS2 - dcs2p  ARO1 - pentafunctional protein aro1p  DUR1,2 - bifunctional urea carboxylase/allophanate hydrolase  ERG3 - c-5 sterol desaturase  ERG12 - mevalonate kinase  SHM2 - glycine hydroxymethyltransferase shm2  PGM1 - phosphoglucomutase pgm1  DLD3 - dld3p  PRB1 - prb1p  ARO8 - bifunctional 2-aminoadipate transaminase/aromatic-amino-acid:2-oxoglutarate transaminase  RAS2 - ras2p  LSC1 - succinate--coa ligase (gdp-forming) subunit alpha  HXK2 - hexokinase 2  RPO21 - rpo21p  ARO3 - 3-deoxy-7-phosphoheptulonate synthase aro3  IDH2 - isocitrate dehydrogenase (nad(+)) idh2  KRS1 - lysine--trna ligase krs1  ADE6 - phosphoribosylformylglycinamidine synthase  YKL151C - nadhx dehydratase  MCR1 - mcr1p  NEW1 - new1p  EHT1 - eht1p  PRX1 - prx1p  YHR020W - proline--trna ligase  ADE2 - phosphoribosylaminoimidazole carboxylase ade2  HEM13 - coproporphyrinogen oxidase  ALD4 - aldehyde dehydrogenase (nadp(+)) ald4  GLT1 - glutamate synthase (nadh)  APE1 - ape1p  DLD1 - dld1p  GFA1 - glutamine--fructose-6-phosphate transaminase (isomerizing) gfa1  CDC28 - cdc28p  MKT1 - mkt1p  LCB2 - serine c-palmitoyltransferase lcb2  AIM17 - aim17p  ATP1 - f1f0 atp synthase subunit alpha  ARA2 - d-arabinose 1-dehydrogenase (nad(p)(+)) ara2  GCN20 - putative aaa family atpase gcn20  TPS2 - trehalose-phosphatase tps2  PTP1 - ptp1p |
| GO:0050662 | coenzyme binding | 3.61E-9 | 3.39E-6 | 3.30 (2298,92,242,32) | [+] Show genes  IDH1 - isocitrate dehydrogenase (nad(+)) idh1  ARG5,6 - bifunctional acetylglutamate kinase/n-acetyl-gamma-glutamyl-phosphate reductase  ACB1 - long-chain fatty acid transporter acb1  GPH1 - gph1p  DLD3 - dld3p  ARO2 - bifunctional chorismate synthase/riboflavin reductase [nad(p)h] aro2  ARO8 - bifunctional 2-aminoadipate transaminase/aromatic-amino-acid:2-oxoglutarate transaminase  MET5 - met5p  PDC1 - indolepyruvate decarboxylase 1  KGD1 - alpha-ketoglutarate dehydrogenase kgd1  GPD1 - glycerol-3-phosphate dehydrogenase (nad(+)) gpd1  GAD1 - glutamate decarboxylase gad1  HOM2 - aspartate-semialdehyde dehydrogenase  ZWF1 - glucose-6-phosphate dehydrogenase  IDH2 - isocitrate dehydrogenase (nad(+)) idh2  MAE1 - malate dehydrogenase (oxaloacetate-decarboxylating)  YGL039W - carbonyl reductase (nadph-dependent)  HIS4 - trifunctional histidinol dehydrogenase/phosphoribosyl-amp cyclohydrolase/phosphoribosyl-atp diphosphatase  GLT1 - glutamate synthase (nadh)  PRO2 - glutamate-5-semialdehyde dehydrogenase  ILV2 - acetolactate synthase catalytic subunit  GLR1 - glutathione-disulfide reductase glr1  BNA5 - kynureninase  DLD1 - dld1p  ILV1 - threonine ammonia-lyase ilv1  YGR017W - hypothetical protein  LCB2 - serine c-palmitoyltransferase lcb2  NFS1 - nfs1p  YBL036C - hypothetical protein  DLD2 - dld2p  CAR2 - ornithine-oxo-acid transaminase  SHM2 - glycine hydroxymethyltransferase shm2 |
| GO:0048037 | cofactor binding | 5.14E-8 | 3.22E-5 | 2.74 (2298,128,242,37) | [+] Show genes  ARG5,6 - bifunctional acetylglutamate kinase/n-acetyl-gamma-glutamyl-phosphate reductase  IDH1 - isocitrate dehydrogenase (nad(+)) idh1  ACB1 - long-chain fatty acid transporter acb1  GPH1 - gph1p  DLD3 - dld3p  ARO2 - bifunctional chorismate synthase/riboflavin reductase [nad(p)h] aro2  ARO8 - bifunctional 2-aminoadipate transaminase/aromatic-amino-acid:2-oxoglutarate transaminase  ILV3 - ilv3p  MET5 - met5p  PDC1 - indolepyruvate decarboxylase 1  GPD1 - glycerol-3-phosphate dehydrogenase (nad(+)) gpd1  KGD1 - alpha-ketoglutarate dehydrogenase kgd1  LSC1 - succinate--coa ligase (gdp-forming) subunit alpha  GRX3 - grx3p  GAD1 - glutamate decarboxylase gad1  HOM2 - aspartate-semialdehyde dehydrogenase  ZWF1 - glucose-6-phosphate dehydrogenase  IDH2 - isocitrate dehydrogenase (nad(+)) idh2  MAE1 - malate dehydrogenase (oxaloacetate-decarboxylating)  HIS4 - trifunctional histidinol dehydrogenase/phosphoribosyl-amp cyclohydrolase/phosphoribosyl-atp diphosphatase  YGL039W - carbonyl reductase (nadph-dependent)  GLT1 - glutamate synthase (nadh)  ILV2 - acetolactate synthase catalytic subunit  PRO2 - glutamate-5-semialdehyde dehydrogenase  GLR1 - glutathione-disulfide reductase glr1  DLD1 - dld1p  BNA5 - kynureninase  YGR017W - hypothetical protein  ILV1 - threonine ammonia-lyase ilv1  LCB2 - serine c-palmitoyltransferase lcb2  NFS1 - nfs1p  YBL036C - hypothetical protein  SDH2 - succinate dehydrogenase iron-sulfur protein subunit sdh2  DLD2 - dld2p  OLE1 - stearoyl-coa 9-desaturase  CAR2 - ornithine-oxo-acid transaminase  SHM2 - glycine hydroxymethyltransferase shm2 |
| GO:0043167 | ion binding | 3.22E-7 | 1.51E-4 | 1.58 (2298,772,185,98) | [+] Show genes  IDH1 - isocitrate dehydrogenase (nad(+)) idh1  RSP5 - nedd4 family e3 ubiquitin-protein ligase  SIT4 - sit4p  ENO2 - phosphopyruvate hydratase eno2  ARO2 - bifunctional chorismate synthase/riboflavin reductase [nad(p)h] aro2  MYO4 - myosin 4  APA1 - apa1p  ASN2 - asparagine synthase (glutamine-hydrolyzing) 2  YSA1 - ysa1p  GLK1 - glucokinase  ARP2 - actin-related protein 2  EMI2 - putative glucokinase  FRA1 - fra1p  PGM2 - phosphoglucomutase pgm2  HIS4 - trifunctional histidinol dehydrogenase/phosphoribosyl-amp cyclohydrolase/phosphoribosyl-atp diphosphatase  QCR2 - ubiquinol--cytochrome-c reductase subunit 2  ILV2 - acetolactate synthase catalytic subunit  HSP104 - chaperone atpase hsp104  SSA2 - hsp70 family chaperone ssa2  GLR1 - glutathione-disulfide reductase glr1  RSM26 - rsm26p  MAS1 - mas1p  NFS1 - nfs1p  TCB1 - tcb1p  ATP2 - atp2p  GLC7 - glc7p  CAR2 - ornithine-oxo-acid transaminase  THI4 - thi4p  NTH1 - alpha,alpha-trehalase nth1  VPS21 - vps21p  RHO1 - rho1p  ARG5,6 - bifunctional acetylglutamate kinase/n-acetyl-gamma-glutamyl-phosphate reductase  GPH1 - gph1p  HOR2 - glycerol-1-phosphatase hor2  ASN1 - asparagine synthase (glutamine-hydrolyzing) 1  SSE1 - sse1p  GUK1 - guanylate kinase  APE2 - ape2p  HOM3 - aspartate kinase  HSC82 - hsp90 family chaperone hsc82  GET3 - guanine nucleotide exchange factor get3  PNC1 - nicotinamidase  HEM2 - porphobilinogen synthase hem2  BDH1 - (r,r)-butanediol dehydrogenase  TRP2 - anthranilate synthase trp2  BNA5 - kynureninase  CDC60 - leucine--trna ligase cdc60  YIL108W - putative metalloendopeptidase  SDH2 - succinate dehydrogenase iron-sulfur protein subunit sdh2  OLE1 - stearoyl-coa 9-desaturase  KES1 - kes1p  DNM1 - dnm1p  HSP82 - hsp90 family chaperone hsp82  ILV3 - ilv3p  PDC1 - indolepyruvate decarboxylase 1  GLO1 - lactoylglutathione lyase glo1  ILV5 - ketol-acid reductoisomerase  GRX3 - grx3p  ENO1 - phosphopyruvate hydratase eno1  UBC13 - e2 ubiquitin-conjugating protein ubc13  ISC1 - inositol phosphosphingolipid phospholipase  AFG3 - aaa family atpase afg3  GLN1 - glutamate--ammonia ligase  ARG1 - argininosuccinate synthase  COR1 - ubiquinol--cytochrome-c reductase subunit cor1  UGP1 - utp glucose-1-phosphate uridylyltransferase  YGR017W - hypothetical protein  AMD1 - amp deaminase  ARO1 - pentafunctional protein aro1p  DUR1,2 - bifunctional urea carboxylase/allophanate hydrolase  ERG3 - c-5 sterol desaturase  ERG12 - mevalonate kinase  SHM2 - glycine hydroxymethyltransferase shm2  SCJ1 - scj1p  ARP3 - arp3p  PGM1 - phosphoglucomutase pgm1  DLD3 - dld3p  ARO8 - bifunctional 2-aminoadipate transaminase/aromatic-amino-acid:2-oxoglutarate transaminase  RAS2 - ras2p  HXK2 - hexokinase 2  SSE2 - sse2p  RPO21 - rpo21p  KRS1 - lysine--trna ligase krs1  IDH2 - isocitrate dehydrogenase (nad(+)) idh2  ADE6 - phosphoribosylformylglycinamidine synthase  YKL151C - nadhx dehydratase  NEW1 - new1p  YHR020W - proline--trna ligase  ADE2 - phosphoribosylaminoimidazole carboxylase ade2  GLT1 - glutamate synthase (nadh)  APE1 - ape1p  DLD1 - dld1p  CDC28 - cdc28p  LCB2 - serine c-palmitoyltransferase lcb2  YDJ1 - ydj1p  AIM17 - aim17p  ATP1 - f1f0 atp synthase subunit alpha  GCN20 - putative aaa family atpase gcn20 |
| GO:0016614 | oxidoreductase activity, acting on CH-OH group of donors | 5.99E-6 | 2.25E-3 | 4.45 (2298,49,158,15) | [+] Show genes  IDH1 - isocitrate dehydrogenase (nad(+)) idh1  MDH3 - malate dehydrogenase mdh3  HIS4 - trifunctional histidinol dehydrogenase/phosphoribosyl-amp cyclohydrolase/phosphoribosyl-atp diphosphatase  DLD3 - dld3p  YGL039W - carbonyl reductase (nadph-dependent)  GRE3 - trifunctional aldehyde reductase/xylose reductase/glucose 1-dehydrogenase (nadp(+))  BDH1 - (r,r)-butanediol dehydrogenase  DLD1 - dld1p  GPD1 - glycerol-3-phosphate dehydrogenase (nad(+)) gpd1  ILV5 - ketol-acid reductoisomerase  ARO1 - pentafunctional protein aro1p  ZWF1 - glucose-6-phosphate dehydrogenase  ARA2 - d-arabinose 1-dehydrogenase (nad(p)(+)) ara2  IDH2 - isocitrate dehydrogenase (nad(+)) idh2  MDH1 - malate dehydrogenase mdh1 |
| GO:0046872 | metal ion binding | 1.33E-5 | 4.18E-3 | 2.01 (2298,353,146,45) | [+] Show genes  IDH1 - isocitrate dehydrogenase (nad(+)) idh1  PGM1 - phosphoglucomutase pgm1  SIT4 - sit4p  ENO2 - phosphopyruvate hydratase eno2  HOR2 - glycerol-1-phosphatase hor2  ILV3 - ilv3p  PDC1 - indolepyruvate decarboxylase 1  GLO1 - lactoylglutathione lyase glo1  ILV5 - ketol-acid reductoisomerase  ENO1 - phosphopyruvate hydratase eno1  APE2 - ape2p  RPO21 - rpo21p  ADE6 - phosphoribosylformylglycinamidine synthase  GET3 - guanine nucleotide exchange factor get3  ISC1 - inositol phosphosphingolipid phospholipase  ADE2 - phosphoribosylaminoimidazole carboxylase ade2  FRA1 - fra1p  PNC1 - nicotinamidase  PGM2 - phosphoglucomutase pgm2  QCR2 - ubiquinol--cytochrome-c reductase subunit 2  HIS4 - trifunctional histidinol dehydrogenase/phosphoribosyl-amp cyclohydrolase/phosphoribosyl-atp diphosphatase  GLT1 - glutamate synthase (nadh)  ILV2 - acetolactate synthase catalytic subunit  HEM2 - porphobilinogen synthase hem2  BDH1 - (r,r)-butanediol dehydrogenase  TRP2 - anthranilate synthase trp2  APE1 - ape1p  COR1 - ubiquinol--cytochrome-c reductase subunit cor1  RSM26 - rsm26p  UGP1 - utp glucose-1-phosphate uridylyltransferase  YIL108W - putative metalloendopeptidase  MAS1 - mas1p  AMD1 - amp deaminase  NFS1 - nfs1p  YDJ1 - ydj1p  AIM17 - aim17p  TCB1 - tcb1p  ARO1 - pentafunctional protein aro1p  DUR1,2 - bifunctional urea carboxylase/allophanate hydrolase  GLC7 - glc7p  THI4 - thi4p  ERG3 - c-5 sterol desaturase  ERG12 - mevalonate kinase  NTH1 - alpha,alpha-trehalase nth1  SCJ1 - scj1p |
| GO:0016616 | oxidoreductase activity, acting on the CH-OH group of donors, NAD or NADP as acceptor | 1.53E-5 | 4.11E-3 | 4.43 (2298,46,158,14) | [+] Show genes  IDH1 - isocitrate dehydrogenase (nad(+)) idh1  MDH3 - malate dehydrogenase mdh3  YGL039W - carbonyl reductase (nadph-dependent)  HIS4 - trifunctional histidinol dehydrogenase/phosphoribosyl-amp cyclohydrolase/phosphoribosyl-atp diphosphatase  GRE3 - trifunctional aldehyde reductase/xylose reductase/glucose 1-dehydrogenase (nadp(+))  BDH1 - (r,r)-butanediol dehydrogenase  DLD1 - dld1p  GPD1 - glycerol-3-phosphate dehydrogenase (nad(+)) gpd1  ILV5 - ketol-acid reductoisomerase  ARO1 - pentafunctional protein aro1p  ZWF1 - glucose-6-phosphate dehydrogenase  ARA2 - d-arabinose 1-dehydrogenase (nad(p)(+)) ara2  IDH2 - isocitrate dehydrogenase (nad(+)) idh2  MDH1 - malate dehydrogenase mdh1 |
| GO:0043169 | cation binding | 1.76E-5 | 4.13E-3 | 1.78 (2298,359,216,60) | [+] Show genes  IDH1 - isocitrate dehydrogenase (nad(+)) idh1  SIT4 - sit4p  ENO2 - phosphopyruvate hydratase eno2  ILV3 - ilv3p  PDC1 - indolepyruvate decarboxylase 1  GLO1 - lactoylglutathione lyase glo1  KGD1 - alpha-ketoglutarate dehydrogenase kgd1  ILV5 - ketol-acid reductoisomerase  GRX3 - grx3p  ENO1 - phosphopyruvate hydratase eno1  YSA1 - ysa1p  COQ1 - trans-hexaprenyltranstransferase  ISC1 - inositol phosphosphingolipid phospholipase  AFG3 - aaa family atpase afg3  FRA1 - fra1p  PGM2 - phosphoglucomutase pgm2  MAE1 - malate dehydrogenase (oxaloacetate-decarboxylating)  HIS4 - trifunctional histidinol dehydrogenase/phosphoribosyl-amp cyclohydrolase/phosphoribosyl-atp diphosphatase  QCR2 - ubiquinol--cytochrome-c reductase subunit 2  ILV2 - acetolactate synthase catalytic subunit  COR1 - ubiquinol--cytochrome-c reductase subunit cor1  UGP1 - utp glucose-1-phosphate uridylyltransferase  RSM26 - rsm26p  MAS1 - mas1p  AMD1 - amp deaminase  NFS1 - nfs1p  ARO1 - pentafunctional protein aro1p  TCB1 - tcb1p  DUR1,2 - bifunctional urea carboxylase/allophanate hydrolase  GLC7 - glc7p  THI4 - thi4p  ERG3 - c-5 sterol desaturase  ERG12 - mevalonate kinase  NTH1 - alpha,alpha-trehalase nth1  FCY1 - cytosine deaminase  SHM2 - glycine hydroxymethyltransferase shm2  SCJ1 - scj1p  ARG5,6 - bifunctional acetylglutamate kinase/n-acetyl-gamma-glutamyl-phosphate reductase  PGM1 - phosphoglucomutase pgm1  CYM1 - cym1p  HOR2 - glycerol-1-phosphatase hor2  APE2 - ape2p  RPO21 - rpo21p  IDH2 - isocitrate dehydrogenase (nad(+)) idh2  ADE6 - phosphoribosylformylglycinamidine synthase  GET3 - guanine nucleotide exchange factor get3  ADE2 - phosphoribosylaminoimidazole carboxylase ade2  PNC1 - nicotinamidase  GLT1 - glutamate synthase (nadh)  HEM2 - porphobilinogen synthase hem2  OLA1 - ola1p  YMR027W - hypothetical protein  TRP2 - anthranilate synthase trp2  APE1 - ape1p  BDH1 - (r,r)-butanediol dehydrogenase  YIL108W - putative metalloendopeptidase  YDJ1 - ydj1p  AIM17 - aim17p  SDH2 - succinate dehydrogenase iron-sulfur protein subunit sdh2  OLE1 - stearoyl-coa 9-desaturase |
| GO:0016491 | oxidoreductase activity | 1.9E-5 | 3.98E-3 | 2.05 (2298,203,243,44) | [+] Show genes  ARG5,6 - bifunctional acetylglutamate kinase/n-acetyl-gamma-glutamyl-phosphate reductase  IDH1 - isocitrate dehydrogenase (nad(+)) idh1  TRX3 - trx3p  DLD3 - dld3p  ARO2 - bifunctional chorismate synthase/riboflavin reductase [nad(p)h] aro2  GRE3 - trifunctional aldehyde reductase/xylose reductase/glucose 1-dehydrogenase (nadp(+))  MET5 - met5p  KGD1 - alpha-ketoglutarate dehydrogenase kgd1  GPD1 - glycerol-3-phosphate dehydrogenase (nad(+)) gpd1  ILV5 - ketol-acid reductoisomerase  GRX3 - grx3p  YMR315W - hypothetical protein  HOM2 - aspartate-semialdehyde dehydrogenase  ZWF1 - glucose-6-phosphate dehydrogenase  IDH2 - isocitrate dehydrogenase (nad(+)) idh2  MCR1 - mcr1p  MDH1 - malate dehydrogenase mdh1  PRX1 - prx1p  HEM13 - coproporphyrinogen oxidase  MDH3 - malate dehydrogenase mdh3  MAE1 - malate dehydrogenase (oxaloacetate-decarboxylating)  ALD4 - aldehyde dehydrogenase (nadp(+)) ald4  YGL039W - carbonyl reductase (nadph-dependent)  HIS4 - trifunctional histidinol dehydrogenase/phosphoribosyl-amp cyclohydrolase/phosphoribosyl-atp diphosphatase  QCR2 - ubiquinol--cytochrome-c reductase subunit 2  GLT1 - glutamate synthase (nadh)  PRO2 - glutamate-5-semialdehyde dehydrogenase  BDH1 - (r,r)-butanediol dehydrogenase  GLR1 - glutathione-disulfide reductase glr1  DLD1 - dld1p  COR1 - ubiquinol--cytochrome-c reductase subunit cor1  RSM26 - rsm26p  ALD2 - aldehyde dehydrogenase (nad(+)) ald2  HFD1 - hfd1p  AIM17 - aim17p  SDH2 - succinate dehydrogenase iron-sulfur protein subunit sdh2  ARO1 - pentafunctional protein aro1p  DLD2 - dld2p  OLE1 - stearoyl-coa 9-desaturase  ERG24 - delta(14)-sterol reductase  NDE1 - nadh-ubiquinone reductase (h(+)-translocating) nde1  ARA2 - d-arabinose 1-dehydrogenase (nad(p)(+)) ara2  ERG3 - c-5 sterol desaturase  ALD5 - aldehyde dehydrogenase (nad(p)(+)) ald5 |
| GO:0004029 | aldehyde dehydrogenase (NAD) activity | 2.78E-5 | 5.22E-3 | 4.53 (2298,8,507,8) | [+] Show genes  MSC7 - msc7p  ALD4 - aldehyde dehydrogenase (nadp(+)) ald4  YGL039W - carbonyl reductase (nadph-dependent)  YNL134C - hypothetical protein  ALD6 - aldehyde dehydrogenase (nadp(+)) ald6  ALD2 - aldehyde dehydrogenase (nad(+)) ald2  HFD1 - hfd1p  ALD5 - aldehyde dehydrogenase (nad(p)(+)) ald5 |
| GO:0016835 | carbon-oxygen lyase activity | 3.42E-5 | 5.84E-3 | 4.28 (2298,23,280,12) | [+] Show genes  HIS3 - imidazoleglycerol-phosphate dehydratase his3  FUM1 - fumarase fum1  PHS1 - phs1p  ARO1 - pentafunctional protein aro1p  ENO1 - phosphopyruvate hydratase eno1  TRP5 - tryptophan synthase trp5  ENO2 - phosphopyruvate hydratase eno2  HEM4 - uroporphyrinogen-iii synthase hem4  HEM2 - porphobilinogen synthase hem2  ARO2 - bifunctional chorismate synthase/riboflavin reductase [nad(p)h] aro2  ILV3 - ilv3p  YKL151C - nadhx dehydratase |
| GO:0016829 | lyase activity | 3.71E-5 | 5.81E-3 | 2.98 (2298,55,280,20) | [+] Show genes  FUM1 - fumarase fum1  ADE2 - phosphoribosylaminoimidazole carboxylase ade2  PHS1 - phs1p  TRP5 - tryptophan synthase trp5  MAE1 - malate dehydrogenase (oxaloacetate-decarboxylating)  ENO2 - phosphopyruvate hydratase eno2  HEM2 - porphobilinogen synthase hem2  ARO2 - bifunctional chorismate synthase/riboflavin reductase [nad(p)h] aro2  TRP2 - anthranilate synthase trp2  ILV3 - ilv3p  PDC1 - indolepyruvate decarboxylase 1  ILV1 - threonine ammonia-lyase ilv1  GLO1 - lactoylglutathione lyase glo1  HIS3 - imidazoleglycerol-phosphate dehydratase his3  RIB3 - 3,4-dihydroxy-2-butanone-4-phosphate synthase rib3  ARO1 - pentafunctional protein aro1p  GAD1 - glutamate decarboxylase gad1  ENO1 - phosphopyruvate hydratase eno1  HEM4 - uroporphyrinogen-iii synthase hem4  YKL151C - nadhx dehydratase |
| GO:0016836 | hydro-lyase activity | 4.34E-5 | 6.28E-3 | 4.51 (2298,20,280,11) | [+] Show genes  HIS3 - imidazoleglycerol-phosphate dehydratase his3  FUM1 - fumarase fum1  PHS1 - phs1p  ARO1 - pentafunctional protein aro1p  ENO1 - phosphopyruvate hydratase eno1  TRP5 - tryptophan synthase trp5  ENO2 - phosphopyruvate hydratase eno2  HEM4 - uroporphyrinogen-iii synthase hem4  HEM2 - porphobilinogen synthase hem2  ILV3 - ilv3p  YKL151C - nadhx dehydratase |
| GO:0016879 | ligase activity, forming carbon-nitrogen bonds | 5.13E-5 | 6.89E-3 | 18.53 (2298,31,20,5) | [+] Show genes  ASN1 - asparagine synthase (glutamine-hydrolyzing) 1  CAB2 - phosphopantothenate--cysteine ligase cab2  ASN2 - asparagine synthase (glutamine-hydrolyzing) 2  DUR1,2 - bifunctional urea carboxylase/allophanate hydrolase  ARG1 - argininosuccinate synthase |
| GO:0004066 | asparagine synthase (glutamine-hydrolyzing) activity | 5.15E-5 | 6.46E-3 | 135.18 (2298,2,17,2) | [+] Show genes  ASN1 - asparagine synthase (glutamine-hydrolyzing) 1  ASN2 - asparagine synthase (glutamine-hydrolyzing) 2 |
| GO:0036094 | small molecule binding | 6.73E-5 | 7.91E-3 | 1.52 (2298,505,264,88) | [+] Show genes  HSP82 - hsp90 family chaperone hsp82  IDH1 - isocitrate dehydrogenase (nad(+)) idh1  ARO2 - bifunctional chorismate synthase/riboflavin reductase [nad(p)h] aro2  MET5 - met5p  PDC1 - indolepyruvate decarboxylase 1  MYO4 - myosin 4  KGD1 - alpha-ketoglutarate dehydrogenase kgd1  GPD1 - glycerol-3-phosphate dehydrogenase (nad(+)) gpd1  APA1 - apa1p  GAD1 - glutamate decarboxylase gad1  ASN2 - asparagine synthase (glutamine-hydrolyzing) 2  UBC13 - e2 ubiquitin-conjugating protein ubc13  ARP2 - actin-related protein 2  GLK1 - glucokinase  EMI2 - putative glucokinase  AFG3 - aaa family atpase afg3  TUF1 - tuf1p  CPA2 - cpa2p  FAA1 - long-chain fatty acid-coa ligase faa1  HTS1 - histidine--trna ligase  MAE1 - malate dehydrogenase (oxaloacetate-decarboxylating)  ARG1 - argininosuccinate synthase  GLN1 - glutamate--ammonia ligase  HIS4 - trifunctional histidinol dehydrogenase/phosphoribosyl-amp cyclohydrolase/phosphoribosyl-atp diphosphatase  PRO2 - glutamate-5-semialdehyde dehydrogenase  ILV2 - acetolactate synthase catalytic subunit  HSP104 - chaperone atpase hsp104  GLR1 - glutathione-disulfide reductase glr1  SSA2 - hsp70 family chaperone ssa2  MYO2 - myosin 2  YGR017W - hypothetical protein  NFS1 - nfs1p  YBL036C - hypothetical protein  ARO1 - pentafunctional protein aro1p  DUR1,2 - bifunctional urea carboxylase/allophanate hydrolase  ATP2 - atp2p  CAR2 - ornithine-oxo-acid transaminase  ADE5,7 - bifunctional aminoimidazole ribotide synthase/glycinamide ribotide synthase  HXK1 - hexokinase 1  ERG12 - mevalonate kinase  SHM2 - glycine hydroxymethyltransferase shm2  VPS21 - vps21p  RVB2 - ruvb family atp-dependent dna helicase reptin  RHO1 - rho1p  ARP3 - arp3p  ARG5,6 - bifunctional acetylglutamate kinase/n-acetyl-gamma-glutamyl-phosphate reductase  ACB1 - long-chain fatty acid transporter acb1  ORC5 - origin recognition complex subunit 5  GPH1 - gph1p  PDR5 - atp-binding cassette multidrug transporter pdr5  DLD3 - dld3p  PAP1 - pap1p  SLT2 - slt2p  ARO8 - bifunctional 2-aminoadipate transaminase/aromatic-amino-acid:2-oxoglutarate transaminase  ACT1 - actin  RAS2 - ras2p  ASN1 - asparagine synthase (glutamine-hydrolyzing) 1  LSC1 - succinate--coa ligase (gdp-forming) subunit alpha  HIS1 - atp phosphoribosyltransferase  SSE1 - sse1p  GUK1 - guanylate kinase  HXK2 - hexokinase 2  HOM2 - aspartate-semialdehyde dehydrogenase  SSE2 - sse2p  ZWF1 - glucose-6-phosphate dehydrogenase  HOM3 - aspartate kinase  IDH2 - isocitrate dehydrogenase (nad(+)) idh2  KRS1 - lysine--trna ligase krs1  ADE6 - phosphoribosylformylglycinamidine synthase  YKL151C - nadhx dehydratase  NEW1 - new1p  HSC82 - hsp90 family chaperone hsc82  GET3 - guanine nucleotide exchange factor get3  YHR020W - proline--trna ligase  ADE2 - phosphoribosylaminoimidazole carboxylase ade2  GLT1 - glutamate synthase (nadh)  OLA1 - ola1p  DLD1 - dld1p  BNA5 - kynureninase  ILV1 - threonine ammonia-lyase ilv1  CDC60 - leucine--trna ligase cdc60  CDC28 - cdc28p  LCB2 - serine c-palmitoyltransferase lcb2  YDJ1 - ydj1p  DLD2 - dld2p  ATP1 - f1f0 atp synthase subunit alpha  GCN20 - putative aaa family atpase gcn20  DNM1 - dnm1p |
| GO:0016903 | oxidoreductase activity, acting on the aldehyde or oxo group of donors | 9.23E-5 | 1.02E-2 | 2.92 (2298,22,537,15) | [+] Show genes  ARG5,6 - bifunctional acetylglutamate kinase/n-acetyl-gamma-glutamyl-phosphate reductase  YGL039W - carbonyl reductase (nadph-dependent)  ALD4 - aldehyde dehydrogenase (nadp(+)) ald4  YDL124W - aldo-keto reductase superfamily protein  PRO2 - glutamate-5-semialdehyde dehydrogenase  LYS2 - l-aminoadipate-semialdehyde dehydrogenase  ALD6 - aldehyde dehydrogenase (nadp(+)) ald6  ALD2 - aldehyde dehydrogenase (nad(+)) ald2  HFD1 - hfd1p  KGD1 - alpha-ketoglutarate dehydrogenase kgd1  MSC7 - msc7p  YMR31 - mitochondrial 37s ribosomal protein ymr31  HOM2 - aspartate-semialdehyde dehydrogenase  YNL134C - hypothetical protein  ALD5 - aldehyde dehydrogenase (nad(p)(+)) ald5 |
| GO:0016620 | oxidoreductase activity, acting on the aldehyde or oxo group of donors, NAD or NADP as acceptor | 1.42E-4 | 1.48E-2 | 3.21 (2298,16,537,12) | [+] Show genes  ARG5,6 - bifunctional acetylglutamate kinase/n-acetyl-gamma-glutamyl-phosphate reductase  MSC7 - msc7p  HOM2 - aspartate-semialdehyde dehydrogenase  ALD4 - aldehyde dehydrogenase (nadp(+)) ald4  YGL039W - carbonyl reductase (nadph-dependent)  YNL134C - hypothetical protein  LYS2 - l-aminoadipate-semialdehyde dehydrogenase  PRO2 - glutamate-5-semialdehyde dehydrogenase  ALD2 - aldehyde dehydrogenase (nad(+)) ald2  ALD6 - aldehyde dehydrogenase (nadp(+)) ald6  ALD5 - aldehyde dehydrogenase (nad(p)(+)) ald5  HFD1 - hfd1p |
| GO:0019842 | vitamin binding | 2.03E-4 | 2.01E-2 | 3.03 (2298,39,311,16) | [+] Show genes  MET17 - bifunctional cysteine synthase/o-acetylhomoserine aminocarboxypropyltransferase met17  GPH1 - gph1p  ARO9 - aromatic-amino-acid:2-oxoglutarate transaminase  ILV2 - acetolactate synthase catalytic subunit  ARO8 - bifunctional 2-aminoadipate transaminase/aromatic-amino-acid:2-oxoglutarate transaminase  BNA5 - kynureninase  PDC1 - indolepyruvate decarboxylase 1  ILV1 - threonine ammonia-lyase ilv1  UGA1 - 4-aminobutyrate transaminase  KGD1 - alpha-ketoglutarate dehydrogenase kgd1  NFS1 - nfs1p  LCB2 - serine c-palmitoyltransferase lcb2  YBL036C - hypothetical protein  GAD1 - glutamate decarboxylase gad1  CAR2 - ornithine-oxo-acid transaminase  SHM2 - glycine hydroxymethyltransferase shm2 |
| GO:0016874 | ligase activity | 2.73E-4 | 2.56E-2 | 7.76 (2298,74,28,7) | [+] Show genes  ASN1 - asparagine synthase (glutamine-hydrolyzing) 1  CAB2 - phosphopantothenate--cysteine ligase cab2  LSC1 - succinate--coa ligase (gdp-forming) subunit alpha  ASN2 - asparagine synthase (glutamine-hydrolyzing) 2  DUR1,2 - bifunctional urea carboxylase/allophanate hydrolase  ARG1 - argininosuccinate synthase  PCS60 - pcs60p |
| GO:0004396 | hexokinase activity | 3.71E-4 | 3.32E-2 | 9.23 (2298,4,249,4) | [+] Show genes  HXK2 - hexokinase 2  HXK1 - hexokinase 1  GLK1 - glucokinase  EMI2 - putative glucokinase |
| GO:0004340 | glucokinase activity | 3.71E-4 | 3.17E-2 | 9.23 (2298,4,249,4) | [+] Show genes  HXK2 - hexokinase 2  HXK1 - hexokinase 1  GLK1 - glucokinase  EMI2 - putative glucokinase |
| GO:0008865 | fructokinase activity | 3.71E-4 | 3.03E-2 | 9.23 (2298,4,249,4) | [+] Show genes  HXK2 - hexokinase 2  HXK1 - hexokinase 1  GLK1 - glucokinase  EMI2 - putative glucokinase |
| GO:0005536 | glucose binding | 3.71E-4 | 2.91E-2 | 9.23 (2298,4,249,4) | [+] Show genes  HXK2 - hexokinase 2  HXK1 - hexokinase 1  GLK1 - glucokinase  EMI2 - putative glucokinase |
| GO:0019158 | mannokinase activity | 3.71E-4 | 2.79E-2 | 9.23 (2298,4,249,4) | [+] Show genes  HXK2 - hexokinase 2  HXK1 - hexokinase 1  GLK1 - glucokinase  EMI2 - putative glucokinase |
| GO:0030170 | pyridoxal phosphate binding | 3.73E-4 | 2.7E-2 | 3.31 (2298,29,311,13) | [+] Show genes  MET17 - bifunctional cysteine synthase/o-acetylhomoserine aminocarboxypropyltransferase met17  GPH1 - gph1p  ARO9 - aromatic-amino-acid:2-oxoglutarate transaminase  ARO8 - bifunctional 2-aminoadipate transaminase/aromatic-amino-acid:2-oxoglutarate transaminase  BNA5 - kynureninase  ILV1 - threonine ammonia-lyase ilv1  UGA1 - 4-aminobutyrate transaminase  LCB2 - serine c-palmitoyltransferase lcb2  NFS1 - nfs1p  GAD1 - glutamate decarboxylase gad1  YBL036C - hypothetical protein  CAR2 - ornithine-oxo-acid transaminase  SHM2 - glycine hydroxymethyltransferase shm2 |
| GO:0070279 | vitamin B6 binding | 3.73E-4 | 2.6E-2 | 3.31 (2298,29,311,13) | [+] Show genes  MET17 - bifunctional cysteine synthase/o-acetylhomoserine aminocarboxypropyltransferase met17  GPH1 - gph1p  ARO9 - aromatic-amino-acid:2-oxoglutarate transaminase  ARO8 - bifunctional 2-aminoadipate transaminase/aromatic-amino-acid:2-oxoglutarate transaminase  BNA5 - kynureninase  ILV1 - threonine ammonia-lyase ilv1  UGA1 - 4-aminobutyrate transaminase  LCB2 - serine c-palmitoyltransferase lcb2  NFS1 - nfs1p  GAD1 - glutamate decarboxylase gad1  YBL036C - hypothetical protein  CAR2 - ornithine-oxo-acid transaminase  SHM2 - glycine hydroxymethyltransferase shm2 |
| GO:0046933 | proton-transporting ATP synthase activity, rotational mechanism | 5.61E-4 | 3.77E-2 | 4.90 (2298,10,328,7) | [+] Show genes  ATP7 - f1f0 atp synthase subunit d  ATP1 - f1f0 atp synthase subunit alpha  ATP2 - atp2p  ATP5 - atp5p  ATP3 - atp3p  ATP15 - f1f0 atp synthase subunit epsilon  ATP4 - atp4p |
| GO:0004039 | allophanate hydrolase activity | 8.7E-4 | 5.65E-2 | 1,149.00 (2298,1,2,1) | [+] Show genes  DUR1,2 - bifunctional urea carboxylase/allophanate hydrolase |
| GO:0004847 | urea carboxylase activity | 8.7E-4 | 5.46E-2 | 1,149.00 (2298,1,2,1) | [+] Show genes  DUR1,2 - bifunctional urea carboxylase/allophanate hydrolase |

Species used: Saccharomyces cerevisiae

The system has recognized 2316 genes out of 2316 gene terms entered by the user.  
 136 genes were recognized by gene symbol and 2180 genes by other gene IDs .  
Only 2298 of these genes are associated with a GO term.

  
 Output in Microsoft Excel format  

% List genereted using GOrilla
% http://cbl-gorilla.cs.technion.ac.il/
% GO term pValue
GO:0003824 1.36E-9
GO:0050662 3.61E-9
GO:0048037 5.14E-8
GO:0043167 3.22E-7
GO:0016614 5.99E-6
GO:0046872 1.33E-5
GO:0016616 1.53E-5
GO:0043169 1.76E-5
GO:0016491 1.9E-5
GO:0004029 2.78E-5
GO:0016835 3.42E-5
GO:0016829 3.71E-5
GO:0016836 4.34E-5
GO:0016879 5.13E-5
GO:0004066 5.15E-5
GO:0036094 6.73E-5
GO:0016903 9.23E-5
GO:0016620 1.42E-4
GO:0019842 2.03E-4
GO:0016874 2.73E-4
GO:0004396 3.71E-4
GO:0004340 3.71E-4
GO:0008865 3.71E-4
GO:0005536 3.71E-4
GO:0019158 3.71E-4
GO:0030170 3.73E-4
GO:0070279 3.73E-4
GO:0046933 5.61E-4
GO:0004039 8.7E-4
GO:0004847 8.7E-4
 Visualize output in REViGO   

The GOrilla database is periodically updated using the GO database and other sources.  
The GOrilla database was last updated on Sep 21, 2019

This results page will be available on this site for one month from now (until
Oct 23, 2019
). You can bookmark this page and come back to it later.

  
**'P-value'** is the enrichment
p-value computed according to the mHG or HG model. This p-value is not
corrected for multiple testing of 1881 GO terms.  
  
**'FDR q-value'** is the correction of the above p-value for multiple testing using the Benjamini and Hochberg (1995) method.   
Namely, for the i^th^ term (ranked according to p-value) the FDR q-value is (p-value \* number of GO terms) / i.   
  
**Enrichment (N, B, n, b)** is defined as follows:  
N - is the total number of genes  
B - is the total number of genes associated with a specific GO term  
n - is the number of genes in the top of the user's input list or in the target set when appropriate  
b - is the number of genes in the intersection  
Enrichment = (b/n) / (B/N)  
  
**Genes:** For each GO term you can see the list of associated genes that appear in the optimal top of the list.  
Each gene name is specified by gene symbol followed by a short description of the gene   

Back to GOrilla analysis results
