## Supplemental Data S1 for "A simple mass-action model predicts genome-wide protein timecourses from mRNA trajectories during a dynamic response in two strains of *Saccharomyces cerevisiae*": GOFUNCTION_prot.html

Results


 **protein** 

*P-value color scale*

|  |  |  |  |  |
| --- | --- | --- | --- | --- |
| > 10^-3^ | 10^-3^ to 10^-5^ | 10^-5^ to 10^-7^ | 10^-7^ to 10^-9^ | < 10^-9^ |


|  |  |  |  |  |  |
| --- | --- | --- | --- | --- | --- |
| **GO term** | **Description** | **P-value** | **FDR q-value** | **Enrichment (N, B, n, b)** | **Genes** |
| GO:0050662 | coenzyme binding | 6.72E-10 | 1.27E-6 | 5.53 (2325,92,96,21) | [+] Show genes  IDH1 - isocitrate dehydrogenase (nad(+)) idh1  ARG5,6 - bifunctional acetylglutamate kinase/n-acetyl-gamma-glutamyl-phosphate reductase  GPH1 - gph1p  HIS4 - trifunctional histidinol dehydrogenase/phosphoribosyl-amp cyclohydrolase/phosphoribosyl-atp diphosphatase  DLD3 - dld3p  GLT1 - glutamate synthase (nadh)  ARO2 - bifunctional chorismate synthase/riboflavin reductase [nad(p)h] aro2  ILV2 - acetolactate synthase catalytic subunit  ARO8 - bifunctional 2-aminoadipate transaminase/aromatic-amino-acid:2-oxoglutarate transaminase  GLR1 - glutathione-disulfide reductase glr1  DLD1 - dld1p  BNA5 - kynureninase  PDC1 - indolepyruvate decarboxylase 1  ILV1 - threonine ammonia-lyase ilv1  GAD1 - glutamate decarboxylase gad1  HOM2 - aspartate-semialdehyde dehydrogenase  CYS4 - cystathionine beta-synthase cys4  ZWF1 - glucose-6-phosphate dehydrogenase  IDH2 - isocitrate dehydrogenase (nad(+)) idh2  YHB1 - yhb1p  MET10 - sulfite reductase subunit alpha |
| GO:0048037 | cofactor binding | 1.57E-8 | 1.49E-5 | 4.31 (2325,128,97,23) | [+] Show genes  IDH1 - isocitrate dehydrogenase (nad(+)) idh1  ARG5,6 - bifunctional acetylglutamate kinase/n-acetyl-gamma-glutamyl-phosphate reductase  GPH1 - gph1p  DLD3 - dld3p  HIS4 - trifunctional histidinol dehydrogenase/phosphoribosyl-amp cyclohydrolase/phosphoribosyl-atp diphosphatase  GLT1 - glutamate synthase (nadh)  ARO2 - bifunctional chorismate synthase/riboflavin reductase [nad(p)h] aro2  ILV2 - acetolactate synthase catalytic subunit  ARO8 - bifunctional 2-aminoadipate transaminase/aromatic-amino-acid:2-oxoglutarate transaminase  ILV3 - ilv3p  GLR1 - glutathione-disulfide reductase glr1  DLD1 - dld1p  BNA5 - kynureninase  PDC1 - indolepyruvate decarboxylase 1  ILV1 - threonine ammonia-lyase ilv1  GAD1 - glutamate decarboxylase gad1  CYS4 - cystathionine beta-synthase cys4  HOM2 - aspartate-semialdehyde dehydrogenase  OLE1 - stearoyl-coa 9-desaturase  ZWF1 - glucose-6-phosphate dehydrogenase  IDH2 - isocitrate dehydrogenase (nad(+)) idh2  YHB1 - yhb1p  MET10 - sulfite reductase subunit alpha |
| GO:0003824 | catalytic activity | 4.78E-8 | 3.02E-5 | 1.51 (2325,1216,114,90) | [+] Show genes  IDH1 - isocitrate dehydrogenase (nad(+)) idh1  GLC3 - 1,4-alpha-glucan branching enzyme  PPA2 - ppa2p  CWH41 - cwh41p  ARO2 - bifunctional chorismate synthase/riboflavin reductase [nad(p)h] aro2  ILV3 - ilv3p  PDC1 - indolepyruvate decarboxylase 1  MYO4 - myosin 4  SCW4 - scw4p  HIS3 - imidazoleglycerol-phosphate dehydratase his3  TAL1 - sedoheptulose-7-phosphate:d-glyceraldehyde-3-phosphate transaldolase tal1  FAA4 - long-chain fatty acid-coa ligase faa4  ILV5 - ketol-acid reductoisomerase  GAD1 - glutamate decarboxylase gad1  YSA1 - ysa1p  HIS7 - imidazoleglycerol-phosphate synthase  MET10 - sulfite reductase subunit alpha  ADE16 - bifunctional phosphoribosylaminoimidazolecarboxamide formyltransferase/imp cyclohydrolase ade16  CPR7 - cpr7p  ARO4 - 3-deoxy-7-phosphoheptulonate synthase aro4  ISC1 - inositol phosphosphingolipid phospholipase  FUM1 - fumarase fum1  TUF1 - tuf1p  CPA2 - cpa2p  PGM2 - phosphoglucomutase pgm2  ARG1 - argininosuccinate synthase  GLN1 - glutamate--ammonia ligase  PCS60 - pcs60p  HIS4 - trifunctional histidinol dehydrogenase/phosphoribosyl-amp cyclohydrolase/phosphoribosyl-atp diphosphatase  EXG1 - exg1p  ILV2 - acetolactate synthase catalytic subunit  GLR1 - glutathione-disulfide reductase glr1  SSA2 - hsp70 family chaperone ssa2  YCK2 - yck2p  ARO1 - pentafunctional protein aro1p  DUR1,2 - bifunctional urea carboxylase/allophanate hydrolase  CYS4 - cystathionine beta-synthase cys4  GLC7 - glc7p  ILV6 - acetolactate synthase regulatory subunit  YHB1 - yhb1p  HSP150 - hsp150p  ARG5,6 - bifunctional acetylglutamate kinase/n-acetyl-gamma-glutamyl-phosphate reductase  CAB2 - phosphopantothenate--cysteine ligase cab2  YMR196W - hypothetical protein  KRE6 - kre6p  PGM1 - phosphoglucomutase pgm1  ARG7 - glutamate n-acetyltransferase  GPH1 - gph1p  CDC10 - septin cdc10  DLD3 - dld3p  YDL124W - aldo-keto reductase superfamily protein  VPS1 - vps1p  ARO8 - bifunctional 2-aminoadipate transaminase/aromatic-amino-acid:2-oxoglutarate transaminase  TRP3 - bifunctional anthranilate synthase/indole-3-glycerol-phosphate synthase  ASN1 - asparagine synthase (glutamine-hydrolyzing) 1  RAS2 - ras2p  GUK1 - guanylate kinase  BPL1 - biotin--[acetyl-coa-carboxylase] ligase bpl1  HOM2 - aspartate-semialdehyde dehydrogenase  HOM3 - aspartate kinase  ZWF1 - glucose-6-phosphate dehydrogenase  ARO3 - 3-deoxy-7-phosphoheptulonate synthase aro3  RPO21 - rpo21p  EHD3 - ehd3p  LAP3 - lap3p  IDH2 - isocitrate dehydrogenase (nad(+)) idh2  LEU4 - 2-isopropylmalate synthase leu4  ARG4 - argininosuccinate lyase arg4  NEW1 - new1p  YHR020W - proline--trna ligase  HEM13 - coproporphyrinogen oxidase  APA2 - apa2p  ALD4 - aldehyde dehydrogenase (nadp(+)) ald4  OLA1 - ola1p  GLT1 - glutamate synthase (nadh)  TRP2 - anthranilate synthase trp2  BDH1 - (r,r)-butanediol dehydrogenase  BNA5 - kynureninase  DLD1 - dld1p  MCM3 - mcm3p  ILV1 - threonine ammonia-lyase ilv1  YPR127W - pyridoxine 4-dehydrogenase  MKT1 - mkt1p  LCB2 - serine c-palmitoyltransferase lcb2  INP53 - phosphatidylinositol-3-/phosphoinositide 5-phosphatase inp53  CPR5 - peptidylprolyl isomerase family protein cpr5  OLE1 - stearoyl-coa 9-desaturase  PGM3 - phosphoglucomutase pgm3  TPD3 - tpd3p  ALD5 - aldehyde dehydrogenase (nad(p)(+)) ald5 |
| GO:0016829 | lyase activity | 4.59E-6 | 2.17E-3 | 5.45 (2325,56,99,13) | [+] Show genes  FUM1 - fumarase fum1  ARO2 - bifunctional chorismate synthase/riboflavin reductase [nad(p)h] aro2  ILV3 - ilv3p  TRP2 - anthranilate synthase trp2  PDC1 - indolepyruvate decarboxylase 1  TRP3 - bifunctional anthranilate synthase/indole-3-glycerol-phosphate synthase  ILV1 - threonine ammonia-lyase ilv1  HIS3 - imidazoleglycerol-phosphate dehydratase his3  GAD1 - glutamate decarboxylase gad1  ARO1 - pentafunctional protein aro1p  CYS4 - cystathionine beta-synthase cys4  HIS7 - imidazoleglycerol-phosphate synthase  ARG4 - argininosuccinate lyase arg4 |
| GO:0036094 | small molecule binding | 1.28E-5 | 4.87E-3 | 1.82 (2325,509,138,55) | [+] Show genes  IDH1 - isocitrate dehydrogenase (nad(+)) idh1  ARO2 - bifunctional chorismate synthase/riboflavin reductase [nad(p)h] aro2  PDC1 - indolepyruvate decarboxylase 1  MYO4 - myosin 4  FAA4 - long-chain fatty acid-coa ligase faa4  GAD1 - glutamate decarboxylase gad1  MET10 - sulfite reductase subunit alpha  TUF1 - tuf1p  CPA2 - cpa2p  ARG1 - argininosuccinate synthase  GLN1 - glutamate--ammonia ligase  HIS4 - trifunctional histidinol dehydrogenase/phosphoribosyl-amp cyclohydrolase/phosphoribosyl-atp diphosphatase  ILV2 - acetolactate synthase catalytic subunit  GLR1 - glutathione-disulfide reductase glr1  SSA2 - hsp70 family chaperone ssa2  YCK2 - yck2p  ARO1 - pentafunctional protein aro1p  CYS4 - cystathionine beta-synthase cys4  DUR1,2 - bifunctional urea carboxylase/allophanate hydrolase  KIN1 - kin1p  CAR2 - ornithine-oxo-acid transaminase  YCF1 - atp-binding cassette glutathione s-conjugate transporter ycf1  YHB1 - yhb1p  RVB2 - ruvb family atp-dependent dna helicase reptin  ARG5,6 - bifunctional acetylglutamate kinase/n-acetyl-gamma-glutamyl-phosphate reductase  GPH1 - gph1p  CDC10 - septin cdc10  DLD3 - dld3p  VPS1 - vps1p  ARO8 - bifunctional 2-aminoadipate transaminase/aromatic-amino-acid:2-oxoglutarate transaminase  RAS2 - ras2p  ASN1 - asparagine synthase (glutamine-hydrolyzing) 1  SSE1 - sse1p  HIS1 - atp phosphoribosyltransferase  BPL1 - biotin--[acetyl-coa-carboxylase] ligase bpl1  GUK1 - guanylate kinase  HOM2 - aspartate-semialdehyde dehydrogenase  SSE2 - sse2p  HOM3 - aspartate kinase  ZWF1 - glucose-6-phosphate dehydrogenase  IDH2 - isocitrate dehydrogenase (nad(+)) idh2  HSC82 - hsp90 family chaperone hsc82  NEW1 - new1p  YHR020W - proline--trna ligase  APA2 - apa2p  GLT1 - glutamate synthase (nadh)  OLA1 - ola1p  DLD1 - dld1p  BNA5 - kynureninase  MCM3 - mcm3p  ILV1 - threonine ammonia-lyase ilv1  LCB2 - serine c-palmitoyltransferase lcb2  POS5 - pos5p  MVD1 - diphosphomevalonate decarboxylase mvd1  DNM1 - dnm1p |
| GO:0016614 | oxidoreductase activity, acting on CH-OH group of donors | 3.51E-5 | 1.11E-2 | 8.52 (2325,52,42,8) | [+] Show genes  IDH1 - isocitrate dehydrogenase (nad(+)) idh1  ILV5 - ketol-acid reductoisomerase  ARO1 - pentafunctional protein aro1p  DLD3 - dld3p  HIS4 - trifunctional histidinol dehydrogenase/phosphoribosyl-amp cyclohydrolase/phosphoribosyl-atp diphosphatase  IDH2 - isocitrate dehydrogenase (nad(+)) idh2  BDH1 - (r,r)-butanediol dehydrogenase  DLD1 - dld1p |
| GO:0043167 | ion binding | 4E-5 | 1.08E-2 | 1.24 (2325,779,586,243) | [+] Show genes  IDH1 - isocitrate dehydrogenase (nad(+)) idh1  RSP5 - nedd4 family e3 ubiquitin-protein ligase  SIT4 - sit4p  GSH1 - gsh1p  ARO2 - bifunctional chorismate synthase/riboflavin reductase [nad(p)h] aro2  SAM2 - methionine adenosyltransferase sam2  FOL3 - dihydrofolate synthase  APA1 - apa1p  GLK1 - glucokinase  ARP2 - actin-related protein 2  PRS2 - ribose phosphate diphosphokinase subunit prs2  EMI2 - putative glucokinase  FRE1 - fre1p  FAA1 - long-chain fatty acid-coa ligase faa1  PGM2 - phosphoglucomutase pgm2  HIS4 - trifunctional histidinol dehydrogenase/phosphoribosyl-amp cyclohydrolase/phosphoribosyl-atp diphosphatase  SNF1 - snf1p  ILV2 - acetolactate synthase catalytic subunit  RPS31 - ubiquitin-ribosomal 40s subunit protein s31 fusion protein  SNF4 - snf4p  MAS1 - mas1p  SEC23 - sec23p  TFS1 - tfs1p  SNQ2 - atp-binding cassette transporter snq2  CPA1 - carbamoyl-phosphate synthase (glutamine-hydrolyzing) cpa1  GLC7 - glc7p  HBS1 - hbs1p  ARG5,6 - bifunctional acetylglutamate kinase/n-acetyl-gamma-glutamyl-phosphate reductase  ATG20 - atg20p  GPH1 - gph1p  MOT2 - ccr4-not core ubiquitin-protein ligase subunit mot2  CDC10 - septin cdc10  RPB9 - rpb9p  VPS1 - vps1p  YPK9 - ypk9p  ASN1 - asparagine synthase (glutamine-hydrolyzing) 1  HIS1 - atp phosphoribosyltransferase  YSH1 - ysh1p  GUK1 - guanylate kinase  HOM3 - aspartate kinase  MCM5 - mcm5p  HSC82 - hsp90 family chaperone hsc82  NOG2 - nog2p  PNC1 - nicotinamidase  CDC42 - cdc42p  TRP2 - anthranilate synthase trp2  BNA5 - kynureninase  SUB2 - sub2p  ILV1 - threonine ammonia-lyase ilv1  CDC31 - cdc31p  MVD1 - diphosphomevalonate decarboxylase mvd1  OLE1 - stearoyl-coa 9-desaturase  PDR15 - atp-binding cassette multidrug transporter pdr15  HSP82 - hsp90 family chaperone hsp82  TMA19 - tma19p  RFC1 - replication factor c subunit 1  PPA2 - ppa2p  TUP1 - tup1p  FUS3 - fus3p  YOR1 - atp-binding cassette transporter yor1  HIS3 - imidazoleglycerol-phosphate dehydratase his3  FAA4 - long-chain fatty acid-coa ligase faa4  REI1 - rei1p  LHS1 - hsp70 family chaperone lhs1  ILV5 - ketol-acid reductoisomerase  RKR1 - ubiquitin-protein ligase rkr1  GRX3 - grx3p  GAD1 - glutamate decarboxylase gad1  ENO1 - phosphopyruvate hydratase eno1  DNF2 - aminophospholipid-translocating p4-type atpase dnf2  FOL2 - gtp cyclohydrolase i  YCR087C-A - hypothetical protein  COQ1 - trans-hexaprenyltranstransferase  URA6 - bifunctional uridylate/adenylate kinase  TUF1 - tuf1p  YBL055C - 3'-5'-exodeoxyribonuclease  YAP1802 - yap1802p  PEP5 - pep5p  RRP3 - rna-dependent atpase rrp3  YKL033W-A - hypothetical protein  SAM4 - sam4p  GLN4 - glutamine--trna ligase  YBL036C - hypothetical protein  ARO1 - pentafunctional protein aro1p  CDC3 - septin cdc3  KIN1 - kin1p  YCF1 - atp-binding cassette glutathione s-conjugate transporter ycf1  FAS2 - trifunctional fatty acid synthase subunit fas2  YHB1 - yhb1p  ERG12 - mevalonate kinase  SCJ1 - scj1p  NCP1 - ncp1p  RVB2 - ruvb family atp-dependent dna helicase reptin  MYO3 - myosin 3  PGM1 - phosphoglucomutase pgm1  ISN1 - imp 5'-nucleotidase  MAS2 - mas2p  PDR5 - atp-binding cassette multidrug transporter pdr5  ARO8 - bifunctional 2-aminoadipate transaminase/aromatic-amino-acid:2-oxoglutarate transaminase  AFG2 - aaa family atpase afg2  ACT1 - actin  THI80 - thiamine diphosphokinase  ARP8 - arp8p  HXK2 - hexokinase 2  VIP1 - inositol polyphosphate kinase vip1  IDH2 - isocitrate dehydrogenase (nad(+)) idh2  KRS1 - lysine--trna ligase krs1  CCA1 - cca1p  NEW1 - new1p  YHR020W - proline--trna ligase  SAD1 - sad1p  APA2 - apa2p  LCB1 - serine c-palmitoyltransferase lcb1  GTS1 - gts1p  CAB1 - pantothenate kinase  APE1 - ape1p  SEC4 - sec4p  TIM10 - tim10p  POS5 - pos5p  LCB2 - serine c-palmitoyltransferase lcb2  PGM3 - phosphoglucomutase pgm3  RAS1 - ras1p  ARL3 - arl3p  ENO2 - phosphopyruvate hydratase eno2  LYS2 - l-aminoadipate-semialdehyde dehydrogenase  GRX5 - grx5p  MYO4 - myosin 4  FAA3 - long-chain fatty acid-coa ligase faa3  UTP14 - utp14p  ASN2 - asparagine synthase (glutamine-hydrolyzing) 2  YSA1 - ysa1p  LSB3 - lsb3p  TCP1 - tcp1p  CLA4 - cla4p  TUB1 - tub1p  PAN1 - pan1p  MET10 - sulfite reductase subunit alpha  NOB1 - nob1p  RFC3 - replication factor c subunit 3  CPA2 - cpa2p  FRA1 - fra1p  SSA2 - hsp70 family chaperone ssa2  GLR1 - glutathione-disulfide reductase glr1  RSM26 - rsm26p  YNL284C-B - gag-pol fusion protein  TCB1 - tcb1p  SSM4 - e3 ubiquitin-protein ligase ssm4  CYS4 - cystathionine beta-synthase cys4  YDR248C - gluconokinase  SEC18 - sec18p  BCY1 - bcy1p  HXK1 - hexokinase 1  CAR2 - ornithine-oxo-acid transaminase  THI4 - thi4p  ARF3 - arf3p  CNA1 - cna1p  VPS21 - vps21p  PTC5 - ptc5p  SNU114 - snu114p  CDC12 - septin cdc12  DAK1 - dak1p  OXP1 - oxp1p  SSE1 - sse1p  FUR1 - uracil phosphoribosyltransferase  BPT1 - atp-binding cassette bilirubin transporter bpt1  APE2 - ape2p  DPS1 - aspartate--trna ligase dps1  ERG1 - squalene monooxygenase  NUP60 - nup60p  COX17 - cox17p  RPC10 - rpc10p  TCB3 - tcb3p  PDX3 - pyridoxamine-phosphate oxidase pdx3  ARO9 - aromatic-amino-acid:2-oxoglutarate transaminase  RHO3 - rho3p  RVB1 - ruvb family atp-dependent dna helicase pontin  YCK1 - yck1p  OLA1 - ola1p  BDH1 - (r,r)-butanediol dehydrogenase  CDC60 - leucine--trna ligase cdc60  SPP1 - spp1p  GRX7 - glutathione-disulfide reductase grx7  YBT1 - bile acid-transporting atpase ybt1  CCT6 - cct6p  SER2 - phosphoserine phosphatase  KES1 - kes1p  DNM1 - dnm1p  PRS1 - ribose phosphate diphosphokinase subunit prs1  LYS12 - homoisocitrate dehydrogenase  CCT2 - cct2p  GLC3 - 1,4-alpha-glucan branching enzyme  MVP1 - mvp1p  SRP54 - srp54p  ILV3 - ilv3p  PDC1 - indolepyruvate decarboxylase 1  KGD1 - alpha-ketoglutarate dehydrogenase kgd1  STH1 - sth1p  LEU1 - 3-isopropylmalate dehydratase leu1  ASG1 - asg1p  ISC1 - inositol phosphosphingolipid phospholipase  HTS1 - histidine--trna ligase  GLN1 - glutamate--ammonia ligase  ARG1 - argininosuccinate synthase  YCK2 - yck2p  PRS5 - ribose phosphate diphosphokinase subunit prs5  YGR017W - hypothetical protein  AMD1 - amp deaminase  NRK1 - ribosylnicotinamide kinase  DUR1,2 - bifunctional urea carboxylase/allophanate hydrolase  MEF1 - mef1p  VTC4 - vtc4p  VPS74 - vps74p  ERG3 - c-5 sterol desaturase  FCY1 - cytosine deaminase  ARP3 - arp3p  AIM45 - aim45p  ACB1 - long-chain fatty acid transporter acb1  FET3 - fet3p  CDC48 - aaa family atpase cdc48  DLD3 - dld3p  CDC11 - cdc11p  PRO1 - glutamate 5-kinase  RAS2 - ras2p  BPL1 - biotin--[acetyl-coa-carboxylase] ligase bpl1  TYR1 - pprephenate dehydrogenase (nadp(+))  SSE2 - sse2p  RCO1 - rco1p  RPO21 - rpo21p  XDJ1 - xdj1p  POL1 - pol1p  UTR4 - putative acireductone synthase utr4  PEP3 - pep3p  MET12 - methylenetetrahydrofolate reductase (nad(p)h) met12  GLT1 - glutamate synthase (nadh)  SPF1 - spf1p  YMR027W - hypothetical protein  DLD1 - dld1p  UBC7 - e2 ubiquitin-conjugating protein ubc7  MCM3 - mcm3p  DLD2 - dld2p  PDC5 - indolepyruvate decarboxylase 5  SSB1 - hsp70 family atpase ssb1  CCT5 - cct5p |
| GO:0043168 | anion binding | 8.52E-5 | 2.02E-2 | 1.30 (2325,517,586,170) | [+] Show genes  ARL3 - arl3p  RSP5 - nedd4 family e3 ubiquitin-protein ligase  GSH1 - gsh1p  LYS2 - l-aminoadipate-semialdehyde dehydrogenase  ARO2 - bifunctional chorismate synthase/riboflavin reductase [nad(p)h] aro2  MYO4 - myosin 4  SAM2 - methionine adenosyltransferase sam2  FAA3 - long-chain fatty acid-coa ligase faa3  UTP14 - utp14p  FOL3 - dihydrofolate synthase  APA1 - apa1p  ASN2 - asparagine synthase (glutamine-hydrolyzing) 2  LSB3 - lsb3p  TCP1 - tcp1p  CLA4 - cla4p  TUB1 - tub1p  GLK1 - glucokinase  ARP2 - actin-related protein 2  PRS2 - ribose phosphate diphosphokinase subunit prs2  MET10 - sulfite reductase subunit alpha  EMI2 - putative glucokinase  CPA2 - cpa2p  RFC3 - replication factor c subunit 3  FAA1 - long-chain fatty acid-coa ligase faa1  HIS4 - trifunctional histidinol dehydrogenase/phosphoribosyl-amp cyclohydrolase/phosphoribosyl-atp diphosphatase  SNF1 - snf1p  ILV2 - acetolactate synthase catalytic subunit  GLR1 - glutathione-disulfide reductase glr1  SSA2 - hsp70 family chaperone ssa2  SNF4 - snf4p  YNL284C-B - gag-pol fusion protein  TFS1 - tfs1p  TCB1 - tcb1p  SNQ2 - atp-binding cassette transporter snq2  YDR248C - gluconokinase  CYS4 - cystathionine beta-synthase cys4  SEC18 - sec18p  BCY1 - bcy1p  CPA1 - carbamoyl-phosphate synthase (glutamine-hydrolyzing) cpa1  CAR2 - ornithine-oxo-acid transaminase  HXK1 - hexokinase 1  ARF3 - arf3p  HBS1 - hbs1p  VPS21 - vps21p  ARG5,6 - bifunctional acetylglutamate kinase/n-acetyl-gamma-glutamyl-phosphate reductase  ATG20 - atg20p  GPH1 - gph1p  CDC10 - septin cdc10  SNU114 - snu114p  CDC12 - septin cdc12  DAK1 - dak1p  VPS1 - vps1p  OXP1 - oxp1p  YPK9 - ypk9p  ASN1 - asparagine synthase (glutamine-hydrolyzing) 1  SSE1 - sse1p  HIS1 - atp phosphoribosyltransferase  FUR1 - uracil phosphoribosyltransferase  GUK1 - guanylate kinase  BPT1 - atp-binding cassette bilirubin transporter bpt1  HOM3 - aspartate kinase  MCM5 - mcm5p  DPS1 - aspartate--trna ligase dps1  ERG1 - squalene monooxygenase  NUP60 - nup60p  HSC82 - hsp90 family chaperone hsc82  TCB3 - tcb3p  PDX3 - pyridoxamine-phosphate oxidase pdx3  NOG2 - nog2p  ARO9 - aromatic-amino-acid:2-oxoglutarate transaminase  RHO3 - rho3p  RVB1 - ruvb family atp-dependent dna helicase pontin  YCK1 - yck1p  OLA1 - ola1p  CDC42 - cdc42p  BNA5 - kynureninase  SUB2 - sub2p  ILV1 - threonine ammonia-lyase ilv1  CDC60 - leucine--trna ligase cdc60  MVD1 - diphosphomevalonate decarboxylase mvd1  YBT1 - bile acid-transporting atpase ybt1  CCT6 - cct6p  KES1 - kes1p  DNM1 - dnm1p  PRS1 - ribose phosphate diphosphokinase subunit prs1  PDR15 - atp-binding cassette multidrug transporter pdr15  HSP82 - hsp90 family chaperone hsp82  CCT2 - cct2p  RFC1 - replication factor c subunit 1  MVP1 - mvp1p  TUP1 - tup1p  SRP54 - srp54p  FUS3 - fus3p  YOR1 - atp-binding cassette transporter yor1  PDC1 - indolepyruvate decarboxylase 1  KGD1 - alpha-ketoglutarate dehydrogenase kgd1  FAA4 - long-chain fatty acid-coa ligase faa4  LHS1 - hsp70 family chaperone lhs1  STH1 - sth1p  GAD1 - glutamate decarboxylase gad1  DNF2 - aminophospholipid-translocating p4-type atpase dnf2  FOL2 - gtp cyclohydrolase i  URA6 - bifunctional uridylate/adenylate kinase  TUF1 - tuf1p  YAP1802 - yap1802p  PEP5 - pep5p  HTS1 - histidine--trna ligase  GLN1 - glutamate--ammonia ligase  ARG1 - argininosuccinate synthase  RRP3 - rna-dependent atpase rrp3  YCK2 - yck2p  PRS5 - ribose phosphate diphosphokinase subunit prs5  YGR017W - hypothetical protein  GLN4 - glutamine--trna ligase  NRK1 - ribosylnicotinamide kinase  YBL036C - hypothetical protein  ARO1 - pentafunctional protein aro1p  DUR1,2 - bifunctional urea carboxylase/allophanate hydrolase  CDC3 - septin cdc3  KIN1 - kin1p  VTC4 - vtc4p  MEF1 - mef1p  VPS74 - vps74p  YCF1 - atp-binding cassette glutathione s-conjugate transporter ycf1  YHB1 - yhb1p  ERG12 - mevalonate kinase  NCP1 - ncp1p  RVB2 - ruvb family atp-dependent dna helicase reptin  ARP3 - arp3p  AIM45 - aim45p  MYO3 - myosin 3  ACB1 - long-chain fatty acid transporter acb1  CDC48 - aaa family atpase cdc48  PDR5 - atp-binding cassette multidrug transporter pdr5  DLD3 - dld3p  CDC11 - cdc11p  ARO8 - bifunctional 2-aminoadipate transaminase/aromatic-amino-acid:2-oxoglutarate transaminase  AFG2 - aaa family atpase afg2  PRO1 - glutamate 5-kinase  ACT1 - actin  THI80 - thiamine diphosphokinase  RAS2 - ras2p  BPL1 - biotin--[acetyl-coa-carboxylase] ligase bpl1  TYR1 - pprephenate dehydrogenase (nadp(+))  ARP8 - arp8p  HXK2 - hexokinase 2  SSE2 - sse2p  VIP1 - inositol polyphosphate kinase vip1  KRS1 - lysine--trna ligase krs1  CCA1 - cca1p  NEW1 - new1p  YHR020W - proline--trna ligase  APA2 - apa2p  LCB1 - serine c-palmitoyltransferase lcb1  PEP3 - pep3p  CAB1 - pantothenate kinase  MET12 - methylenetetrahydrofolate reductase (nad(p)h) met12  GLT1 - glutamate synthase (nadh)  SPF1 - spf1p  DLD1 - dld1p  UBC7 - e2 ubiquitin-conjugating protein ubc7  MCM3 - mcm3p  SEC4 - sec4p  POS5 - pos5p  LCB2 - serine c-palmitoyltransferase lcb2  DLD2 - dld2p  PDC5 - indolepyruvate decarboxylase 5  RAS1 - ras1p  SSB1 - hsp70 family atpase ssb1  CCT5 - cct5p |
| GO:0050660 | flavin adenine dinucleotide binding | 1.14E-4 | 2.4E-2 | 8.07 (2325,21,96,7) | [+] Show genes  DLD3 - dld3p  GLT1 - glutamate synthase (nadh)  ILV2 - acetolactate synthase catalytic subunit  GLR1 - glutathione-disulfide reductase glr1  YHB1 - yhb1p  DLD1 - dld1p  MET10 - sulfite reductase subunit alpha |
| GO:0005524 | ATP binding | 1.17E-4 | 2.21E-2 | 7.00 (2325,332,6,6) | [+] Show genes  ASN1 - asparagine synthase (glutamine-hydrolyzing) 1  ARO1 - pentafunctional protein aro1p  GUK1 - guanylate kinase  DUR1,2 - bifunctional urea carboxylase/allophanate hydrolase  HOM3 - aspartate kinase  YHR020W - proline--trna ligase |
| GO:0032559 | adenyl ribonucleotide binding | 1.28E-4 | 2.2E-2 | 6.94 (2325,335,6,6) | [+] Show genes  ASN1 - asparagine synthase (glutamine-hydrolyzing) 1  ARO1 - pentafunctional protein aro1p  GUK1 - guanylate kinase  DUR1,2 - bifunctional urea carboxylase/allophanate hydrolase  HOM3 - aspartate kinase  YHR020W - proline--trna ligase |
| GO:0030554 | adenyl nucleotide binding | 1.34E-4 | 2.12E-2 | 6.86 (2325,339,6,6) | [+] Show genes  ASN1 - asparagine synthase (glutamine-hydrolyzing) 1  ARO1 - pentafunctional protein aro1p  GUK1 - guanylate kinase  DUR1,2 - bifunctional urea carboxylase/allophanate hydrolase  HOM3 - aspartate kinase  YHR020W - proline--trna ligase |
| GO:0003735 | structural constituent of ribosome | 1.38E-4 | 2.01E-2 | 1.93 (2325,104,451,39) | [+] Show genes  RPS1B - ribosomal 40s subunit protein s1b  RPL29 - ribosomal 60s subunit protein l29  RPL25 - ribosomal 60s subunit protein l25  RPS9B - ribosomal 40s subunit protein s9b  RPL24A - ribosomal 60s subunit protein l24a  RPS3 - ribosomal 40s subunit protein s3  RPL17B - rpl17bp  RPL30 - ribosomal 60s subunit protein l30  RPL7B - ribosomal 60s subunit protein l7b  MRPL19 - mitochondrial 54s ribosomal protein yml19  RPL32 - ribosomal 60s subunit protein l32  RPL26B - ribosomal 60s subunit protein l26b  RPL9A - ribosomal 60s subunit protein l9a  RPL8A - ribosomal 60s subunit protein l8a  EHD3 - ehd3p  RSM18 - mitochondrial 37s ribosomal protein rsm18  RPL26A - ribosomal 60s subunit protein l26a  RPL15A - ribosomal 60s subunit protein l15a  RPP0 - ribosomal protein p0  RSM7 - rsm7p  RPL6A - ribosomal 60s subunit protein l6a  RPL3 - ribosomal 60s subunit protein l3  RPL14B - ribosomal 60s subunit protein l14b  RPL31A - ribosomal 60s subunit protein l31a  RPL10 - ribosomal 60s subunit protein l10  RSM26 - rsm26p  RPL6B - ribosomal 60s subunit protein l6b  MRPS12 - putative mitochondrial 37s ribosomal protein mrps12  RPS2 - ribosomal 40s subunit protein s2  RPL33B - ribosomal 60s subunit protein l33b  RPP2A - ribosomal protein p2a  RPS13 - ribosomal 40s subunit protein s13  RPS5 - rps5p  RPL5 - ribosomal 60s subunit protein l5  RSM25 - mitochondrial 37s ribosomal protein rsm25  RPS7A - ribosomal 40s subunit protein s7a  MRPL6 - mitochondrial 54s ribosomal protein yml16  RPL33A - ribosomal 60s subunit protein l33a  RPL16B - ribosomal 60s subunit protein l16b |
| GO:0016616 | oxidoreductase activity, acting on the CH-OH group of donors, NAD or NADP as acceptor | 2.38E-4 | 3.23E-2 | 7.91 (2325,49,42,7) | [+] Show genes  IDH1 - isocitrate dehydrogenase (nad(+)) idh1  ILV5 - ketol-acid reductoisomerase  ARO1 - pentafunctional protein aro1p  HIS4 - trifunctional histidinol dehydrogenase/phosphoribosyl-amp cyclohydrolase/phosphoribosyl-atp diphosphatase  IDH2 - isocitrate dehydrogenase (nad(+)) idh2  BDH1 - (r,r)-butanediol dehydrogenase  DLD1 - dld1p |
| GO:0008144 | drug binding | 2.52E-4 | 3.19E-2 | 6.13 (2325,379,6,6) | [+] Show genes  ASN1 - asparagine synthase (glutamine-hydrolyzing) 1  ARO1 - pentafunctional protein aro1p  GUK1 - guanylate kinase  DUR1,2 - bifunctional urea carboxylase/allophanate hydrolase  HOM3 - aspartate kinase  YHR020W - proline--trna ligase |
| GO:0003849 | 3-deoxy-7-phosphoheptulonate synthase activity | 2.8E-4 | 3.31E-2 | 83.04 (2325,2,28,2) | [+] Show genes  ARO3 - 3-deoxy-7-phosphoheptulonate synthase aro3  ARO4 - 3-deoxy-7-phosphoheptulonate synthase aro4 |
| GO:0016491 | oxidoreductase activity | 3.02E-4 | 3.37E-2 | 2.94 (2325,206,69,18) | [+] Show genes  IDH1 - isocitrate dehydrogenase (nad(+)) idh1  ARG5,6 - bifunctional acetylglutamate kinase/n-acetyl-gamma-glutamyl-phosphate reductase  HEM13 - coproporphyrinogen oxidase  ALD4 - aldehyde dehydrogenase (nadp(+)) ald4  DLD3 - dld3p  HIS4 - trifunctional histidinol dehydrogenase/phosphoribosyl-amp cyclohydrolase/phosphoribosyl-atp diphosphatase  YDL124W - aldo-keto reductase superfamily protein  GLT1 - glutamate synthase (nadh)  ARO2 - bifunctional chorismate synthase/riboflavin reductase [nad(p)h] aro2  BDH1 - (r,r)-butanediol dehydrogenase  GLR1 - glutathione-disulfide reductase glr1  DLD1 - dld1p  ILV5 - ketol-acid reductoisomerase  ARO1 - pentafunctional protein aro1p  HOM2 - aspartate-semialdehyde dehydrogenase  IDH2 - isocitrate dehydrogenase (nad(+)) idh2  YHB1 - yhb1p  ALD5 - aldehyde dehydrogenase (nad(p)(+)) ald5 |
| GO:0019842 | vitamin binding | 3.39E-4 | 3.57E-2 | 4.66 (2325,39,128,10) | [+] Show genes  LCB2 - serine c-palmitoyltransferase lcb2  GAD1 - glutamate decarboxylase gad1  GPH1 - gph1p  CYS4 - cystathionine beta-synthase cys4  CAR2 - ornithine-oxo-acid transaminase  ILV2 - acetolactate synthase catalytic subunit  ARO8 - bifunctional 2-aminoadipate transaminase/aromatic-amino-acid:2-oxoglutarate transaminase  BNA5 - kynureninase  PDC1 - indolepyruvate decarboxylase 1  ILV1 - threonine ammonia-lyase ilv1 |
| GO:0035639 | purine ribonucleoside triphosphate binding | 3.46E-4 | 3.45E-2 | 5.89 (2325,395,6,6) | [+] Show genes  ASN1 - asparagine synthase (glutamine-hydrolyzing) 1  ARO1 - pentafunctional protein aro1p  GUK1 - guanylate kinase  DUR1,2 - bifunctional urea carboxylase/allophanate hydrolase  HOM3 - aspartate kinase  YHR020W - proline--trna ligase |
| GO:0032555 | purine ribonucleotide binding | 3.46E-4 | 3.28E-2 | 5.84 (2325,398,6,6) | [+] Show genes  ASN1 - asparagine synthase (glutamine-hydrolyzing) 1  ARO1 - pentafunctional protein aro1p  GUK1 - guanylate kinase  DUR1,2 - bifunctional urea carboxylase/allophanate hydrolase  HOM3 - aspartate kinase  YHR020W - proline--trna ligase |
| GO:0004042 | acetyl-CoA:L-glutamate N-acetyltransferase activity | 3.67E-4 | 3.31E-2 | 72.66 (2325,2,32,2) | [+] Show genes  ARG5,6 - bifunctional acetylglutamate kinase/n-acetyl-gamma-glutamyl-phosphate reductase  ARG7 - glutamate n-acetyltransferase |
| GO:0017076 | purine nucleotide binding | 3.95E-4 | 3.4E-2 | 5.77 (2325,403,6,6) | [+] Show genes  ASN1 - asparagine synthase (glutamine-hydrolyzing) 1  ARO1 - pentafunctional protein aro1p  GUK1 - guanylate kinase  DUR1,2 - bifunctional urea carboxylase/allophanate hydrolase  HOM3 - aspartate kinase  YHR020W - proline--trna ligase |
| GO:0004039 | allophanate hydrolase activity | 4.3E-4 | 3.54E-2 | 2,325.00 (2325,1,1,1) | [+] Show genes  DUR1,2 - bifunctional urea carboxylase/allophanate hydrolase |
| GO:0004847 | urea carboxylase activity | 4.3E-4 | 3.4E-2 | 2,325.00 (2325,1,1,1) | [+] Show genes  DUR1,2 - bifunctional urea carboxylase/allophanate hydrolase |
| GO:0032553 | ribonucleotide binding | 4.55E-4 | 3.45E-2 | 5.66 (2325,411,6,6) | [+] Show genes  ASN1 - asparagine synthase (glutamine-hydrolyzing) 1  ARO1 - pentafunctional protein aro1p  GUK1 - guanylate kinase  DUR1,2 - bifunctional urea carboxylase/allophanate hydrolase  HOM3 - aspartate kinase  YHR020W - proline--trna ligase |
| GO:0097367 | carbohydrate derivative binding | 4.93E-4 | 3.59E-2 | 5.54 (2325,420,6,6) | [+] Show genes  ASN1 - asparagine synthase (glutamine-hydrolyzing) 1  ARO1 - pentafunctional protein aro1p  GUK1 - guanylate kinase  DUR1,2 - bifunctional urea carboxylase/allophanate hydrolase  HOM3 - aspartate kinase  YHR020W - proline--trna ligase |
| GO:0004449 | isocitrate dehydrogenase (NAD+) activity | 5.2E-4 | 3.65E-2 | 61.18 (2325,2,38,2) | [+] Show genes  IDH1 - isocitrate dehydrogenase (nad(+)) idh1  IDH2 - isocitrate dehydrogenase (nad(+)) idh2 |
| GO:0051287 | NAD binding | 5.23E-4 | 3.54E-2 | 10.68 (2325,16,68,5) | [+] Show genes  IDH1 - isocitrate dehydrogenase (nad(+)) idh1  ARG5,6 - bifunctional acetylglutamate kinase/n-acetyl-gamma-glutamyl-phosphate reductase  HOM2 - aspartate-semialdehyde dehydrogenase  HIS4 - trifunctional histidinol dehydrogenase/phosphoribosyl-amp cyclohydrolase/phosphoribosyl-atp diphosphatase  IDH2 - isocitrate dehydrogenase (nad(+)) idh2 |
| GO:1901265 | nucleoside phosphate binding | 5.27E-4 | 3.44E-2 | 2.25 (2325,465,51,23) | [+] Show genes  IDH1 - isocitrate dehydrogenase (nad(+)) idh1  ARG5,6 - bifunctional acetylglutamate kinase/n-acetyl-gamma-glutamyl-phosphate reductase  CPA2 - cpa2p  ARG1 - argininosuccinate synthase  DLD3 - dld3p  HIS4 - trifunctional histidinol dehydrogenase/phosphoribosyl-amp cyclohydrolase/phosphoribosyl-atp diphosphatase  GLT1 - glutamate synthase (nadh)  ARO2 - bifunctional chorismate synthase/riboflavin reductase [nad(p)h] aro2  ILV2 - acetolactate synthase catalytic subunit  YCK2 - yck2p  DLD1 - dld1p  MCM3 - mcm3p  FAA4 - long-chain fatty acid-coa ligase faa4  RAS2 - ras2p  ASN1 - asparagine synthase (glutamine-hydrolyzing) 1  SSE1 - sse1p  ARO1 - pentafunctional protein aro1p  GUK1 - guanylate kinase  DUR1,2 - bifunctional urea carboxylase/allophanate hydrolase  HOM3 - aspartate kinase  IDH2 - isocitrate dehydrogenase (nad(+)) idh2  YHB1 - yhb1p  YHR020W - proline--trna ligase |
| GO:0000166 | nucleotide binding | 5.27E-4 | 3.33E-2 | 2.25 (2325,465,51,23) | [+] Show genes  IDH1 - isocitrate dehydrogenase (nad(+)) idh1  ARG5,6 - bifunctional acetylglutamate kinase/n-acetyl-gamma-glutamyl-phosphate reductase  CPA2 - cpa2p  ARG1 - argininosuccinate synthase  DLD3 - dld3p  HIS4 - trifunctional histidinol dehydrogenase/phosphoribosyl-amp cyclohydrolase/phosphoribosyl-atp diphosphatase  GLT1 - glutamate synthase (nadh)  ARO2 - bifunctional chorismate synthase/riboflavin reductase [nad(p)h] aro2  ILV2 - acetolactate synthase catalytic subunit  YCK2 - yck2p  DLD1 - dld1p  MCM3 - mcm3p  FAA4 - long-chain fatty acid-coa ligase faa4  RAS2 - ras2p  ASN1 - asparagine synthase (glutamine-hydrolyzing) 1  SSE1 - sse1p  ARO1 - pentafunctional protein aro1p  GUK1 - guanylate kinase  DUR1,2 - bifunctional urea carboxylase/allophanate hydrolase  HOM3 - aspartate kinase  IDH2 - isocitrate dehydrogenase (nad(+)) idh2  YHB1 - yhb1p  YHR020W - proline--trna ligase |
| GO:0004553 | hydrolase activity, hydrolyzing O-glycosyl compounds | 5.43E-4 | 3.32E-2 | 4.62 (2325,25,181,9) | [+] Show genes  KRE6 - kre6p  YMR196W - hypothetical protein  GLC3 - 1,4-alpha-glucan branching enzyme  EXG1 - exg1p  CWH41 - cwh41p  SCW10 - scw10p  CRH1 - crh1p  DSE4 - dse4p  SCW4 - scw4p |
| GO:1901363 | heterocyclic compound binding | 6.19E-4 | 3.67E-2 | 1.18 (2325,922,586,274) | [+] Show genes  IDH1 - isocitrate dehydrogenase (nad(+)) idh1  TFA2 - tfa2p  MSC6 - msc6p  GSH1 - gsh1p  UTP30 - utp30p  ARO2 - bifunctional chorismate synthase/riboflavin reductase [nad(p)h] aro2  LHP1 - lhp1p  RPL9A - ribosomal 60s subunit protein l9a  SAM2 - methionine adenosyltransferase sam2  RPL8A - ribosomal 60s subunit protein l8a  FOL3 - dihydrofolate synthase  APA1 - apa1p  GLE2 - gle2p  FPR2 - peptidylprolyl isomerase family protein fpr2  NAP1 - nap1p  LSM4 - lsm4p  GLK1 - glucokinase  ARP2 - actin-related protein 2  PRS2 - ribose phosphate diphosphokinase subunit prs2  STO1 - sto1p  EMI2 - putative glucokinase  SRO9 - sro9p  MLP1 - mlp1p  FAA1 - long-chain fatty acid-coa ligase faa1  SNF1 - snf1p  HIS4 - trifunctional histidinol dehydrogenase/phosphoribosyl-amp cyclohydrolase/phosphoribosyl-atp diphosphatase  ILV2 - acetolactate synthase catalytic subunit  LSM5 - lsm5p  SNF4 - snf4p  HRB1 - hrb1p  SPT15 - spt15p  RPS2 - ribosomal 40s subunit protein s2  SNQ2 - atp-binding cassette transporter snq2  CPA1 - carbamoyl-phosphate synthase (glutamine-hydrolyzing) cpa1  HBS1 - hbs1p  ARG5,6 - bifunctional acetylglutamate kinase/n-acetyl-gamma-glutamyl-phosphate reductase  MOT2 - ccr4-not core ubiquitin-protein ligase subunit mot2  GPH1 - gph1p  CDC10 - septin cdc10  RPB9 - rpb9p  VPS1 - vps1p  YPK9 - ypk9p  DCS1 - dcs1p  ASN1 - asparagine synthase (glutamine-hydrolyzing) 1  HIS1 - atp phosphoribosyltransferase  GUK1 - guanylate kinase  YSH1 - ysh1p  SRP14 - srp14p  HOM3 - aspartate kinase  MCM5 - mcm5p  RSM18 - mitochondrial 37s ribosomal protein rsm18  TDH1 - tdh1p  HSC82 - hsp90 family chaperone hsc82  RRP43 - rrp43p  NOG2 - nog2p  RPB7 - rpb7p  CDC42 - cdc42p  BNA5 - kynureninase  SUB2 - sub2p  SCD6 - scd6p  ILV1 - threonine ammonia-lyase ilv1  TIF4632 - tif4632p  MVD1 - diphosphomevalonate decarboxylase mvd1  OLE1 - stearoyl-coa 9-desaturase  PDR15 - atp-binding cassette multidrug transporter pdr15  HSP82 - hsp90 family chaperone hsp82  RFC1 - replication factor c subunit 1  LRP1 - lrp1p  FUS3 - fus3p  YOR1 - atp-binding cassette transporter yor1  TIF11 - tif11p  SCW4 - scw4p  REI1 - rei1p  FAA4 - long-chain fatty acid-coa ligase faa4  LHS1 - hsp70 family chaperone lhs1  ILV5 - ketol-acid reductoisomerase  BFR1 - bfr1p  GAD1 - glutamate decarboxylase gad1  DNF2 - aminophospholipid-translocating p4-type atpase dnf2  FOL2 - gtp cyclohydrolase i  YCR087C-A - hypothetical protein  RPL26A - ribosomal 60s subunit protein l26a  YHM2 - yhm2p  RPP0 - ribosomal protein p0  MDH1 - malate dehydrogenase mdh1  URA6 - bifunctional uridylate/adenylate kinase  TUF1 - tuf1p  RRP3 - rna-dependent atpase rrp3  DPB4 - dpb4p  TRM1 - trm1p  GLN4 - glutamine--trna ligase  YBL036C - hypothetical protein  ARO1 - pentafunctional protein aro1p  CDC3 - septin cdc3  KIN1 - kin1p  HEK2 - hek2p  YCF1 - atp-binding cassette glutathione s-conjugate transporter ycf1  YHB1 - yhb1p  ERG12 - mevalonate kinase  SMI1 - smi1p  NCP1 - ncp1p  RVB2 - ruvb family atp-dependent dna helicase reptin  DEF1 - def1p  MYO3 - myosin 3  PDR5 - atp-binding cassette multidrug transporter pdr5  ARO8 - bifunctional 2-aminoadipate transaminase/aromatic-amino-acid:2-oxoglutarate transaminase  AFG2 - aaa family atpase afg2  ACT1 - actin  SHE2 - she2p  THI80 - thiamine diphosphokinase  LSC1 - succinate--coa ligase (gdp-forming) subunit alpha  ARP8 - arp8p  HXK2 - hexokinase 2  VIP1 - inositol polyphosphate kinase vip1  GON7 - gon7p  IDH2 - isocitrate dehydrogenase (nad(+)) idh2  KRS1 - lysine--trna ligase krs1  CCA1 - cca1p  NEW1 - new1p  YHR020W - proline--trna ligase  APA2 - apa2p  LCB1 - serine c-palmitoyltransferase lcb1  RPL14B - ribosomal 60s subunit protein l14b  CAB1 - pantothenate kinase  LEO1 - leo1p  SEC4 - sec4p  POS5 - pos5p  LCB2 - serine c-palmitoyltransferase lcb2  TFC7 - tfc7p  RPS13 - ribosomal 40s subunit protein s13  RAS1 - ras1p  ARL3 - arl3p  STI1 - sti1p  PCF11 - pcf11p  MYO4 - myosin 4  FAA3 - long-chain fatty acid-coa ligase faa3  UTP14 - utp14p  ASN2 - asparagine synthase (glutamine-hydrolyzing) 2  TCP1 - tcp1p  CLA4 - cla4p  SPT6 - spt6p  GCD6 - gcd6p  TUB1 - tub1p  FES1 - fes1p  YDR210C-C - gag protein  NOP4 - nop4p  MET10 - sulfite reductase subunit alpha  NOB1 - nob1p  RSM7 - rsm7p  NHP6B - nhp6bp  POL30 - pol30p  CPA2 - cpa2p  RFC3 - replication factor c subunit 3  SSA2 - hsp70 family chaperone ssa2  GLR1 - glutathione-disulfide reductase glr1  FPR4 - peptidylprolyl isomerase fpr4  UTP9 - utp9p  RPL6B - ribosomal 60s subunit protein l6b  YNL284C-B - gag-pol fusion protein  BUD21 - bud21p  CYS4 - cystathionine beta-synthase cys4  YDR248C - gluconokinase  SEC18 - sec18p  BCY1 - bcy1p  PTI1 - pti1p  RPS5 - rps5p  HXK1 - hexokinase 1  CAR2 - ornithine-oxo-acid transaminase  ARF3 - arf3p  RPL24B - ribosomal 60s subunit protein l24b  VPS21 - vps21p  HSP26 - hsp26p  SNU114 - snu114p  CDC12 - septin cdc12  DAK1 - dak1p  YRA2 - yra2p  SER33 - phosphoglycerate dehydrogenase ser33  OXP1 - oxp1p  GRE3 - trifunctional aldehyde reductase/xylose reductase/glucose 1-dehydrogenase (nadp(+))  RPA49 - rpa49p  SSE1 - sse1p  FUR1 - uracil phosphoribosyltransferase  SUI1 - sui1p  MAM33 - mam33p  BPT1 - atp-binding cassette bilirubin transporter bpt1  HOM2 - aspartate-semialdehyde dehydrogenase  ZWF1 - glucose-6-phosphate dehydrogenase  ELP4 - elongator subunit elp4  DPS1 - aspartate--trna ligase dps1  CPR1 - peptidylprolyl isomerase cpr1  ERG1 - squalene monooxygenase  LAP3 - lap3p  RPC10 - rpc10p  TOP1 - dna topoisomerase 1  PDX3 - pyridoxamine-phosphate oxidase pdx3  RPL6A - ribosomal 60s subunit protein l6a  ARO9 - aromatic-amino-acid:2-oxoglutarate transaminase  RHO3 - rho3p  SKI6 - ski6p  RVB1 - ruvb family atp-dependent dna helicase pontin  YCK1 - yck1p  OLA1 - ola1p  CDC60 - leucine--trna ligase cdc60  RAP1 - rap1p  YBT1 - bile acid-transporting atpase ybt1  CCT6 - cct6p  RPL5 - ribosomal 60s subunit protein l5  HMO1 - hmo1p  DNM1 - dnm1p  MRPL6 - mitochondrial 54s ribosomal protein yml16  PRS1 - ribose phosphate diphosphokinase subunit prs1  LYS12 - homoisocitrate dehydrogenase  CCT2 - cct2p  RPL24A - ribosomal 60s subunit protein l24a  RPS3 - ribosomal 40s subunit protein s3  RPL30 - ribosomal 60s subunit protein l30  MRPL19 - mitochondrial 54s ribosomal protein yml19  POL31 - pol31p  SRP54 - srp54p  PDC1 - indolepyruvate decarboxylase 1  KGD1 - alpha-ketoglutarate dehydrogenase kgd1  ABD1 - abd1p  STH1 - sth1p  NOP13 - nop13p  ASG1 - asg1p  RPL15A - ribosomal 60s subunit protein l15a  HTS1 - histidine--trna ligase  GLN1 - glutamate--ammonia ligase  ARG1 - argininosuccinate synthase  PCS60 - pcs60p  YCK2 - yck2p  PRS5 - ribose phosphate diphosphokinase subunit prs5  YGR017W - hypothetical protein  NOP12 - nop12p  NRK1 - ribosylnicotinamide kinase  NOP9 - nop9p  DUR1,2 - bifunctional urea carboxylase/allophanate hydrolase  NHP6A - nhp6ap  MEF1 - mef1p  POP6 - pop6p  REX2 - rex2p  AIM45 - aim45p  ARP3 - arp3p  RPS9B - ribosomal 40s subunit protein s9b  RPL25 - ribosomal 60s subunit protein l25  ACB1 - long-chain fatty acid transporter acb1  CDC48 - aaa family atpase cdc48  DLD3 - dld3p  RPL26B - ribosomal 60s subunit protein l26b  CDC11 - cdc11p  PRO1 - glutamate 5-kinase  RAS2 - ras2p  YGR054W - hypothetical protein  BPL1 - biotin--[acetyl-coa-carboxylase] ligase bpl1  TYR1 - pprephenate dehydrogenase (nadp(+))  SSE2 - sse2p  YMR074C - hypothetical protein  RPO21 - rpo21p  RRP45 - rrp45p  POL1 - pol1p  STM1 - stm1p  MET12 - methylenetetrahydrofolate reductase (nad(p)h) met12  GLT1 - glutamate synthase (nadh)  SPF1 - spf1p  DLD1 - dld1p  UBC7 - e2 ubiquitin-conjugating protein ubc7  MCM3 - mcm3p  DLD2 - dld2p  PDC5 - indolepyruvate decarboxylase 5  RRP46 - rrp46p  CCT5 - cct5p  SSB1 - hsp70 family atpase ssb1  RPL16B - ribosomal 60s subunit protein l16b  SUB1 - sub1p |
| GO:0004030 | aldehyde dehydrogenase [NAD(P)+] activity | 6.37E-4 | 3.66E-2 | 62.00 (2325,3,25,2) | [+] Show genes  ALD4 - aldehyde dehydrogenase (nadp(+)) ald4  ALD5 - aldehyde dehydrogenase (nad(p)(+)) ald5 |
| GO:0000287 | magnesium ion binding | 7.04E-4 | 3.92E-2 | 2.10 (2325,42,605,23) | [+] Show genes  IDH1 - isocitrate dehydrogenase (nad(+)) idh1  PGM1 - phosphoglucomutase pgm1  UTR4 - putative acireductone synthase utr4  ISN1 - imp 5'-nucleotidase  PGM2 - phosphoglucomutase pgm2  PPA2 - ppa2p  ENO2 - phosphopyruvate hydratase eno2  ILV2 - acetolactate synthase catalytic subunit  YKL033W-A - hypothetical protein  PRS5 - ribose phosphate diphosphokinase subunit prs5  PDC1 - indolepyruvate decarboxylase 1  HIS1 - atp phosphoribosyltransferase  ENO1 - phosphopyruvate hydratase eno1  PGM3 - phosphoglucomutase pgm3  PDC5 - indolepyruvate decarboxylase 5  DNF2 - aminophospholipid-translocating p4-type atpase dnf2  IDH2 - isocitrate dehydrogenase (nad(+)) idh2  SER2 - phosphoserine phosphatase  FAS2 - trifunctional fatty acid synthase subunit fas2  PRS2 - ribose phosphate diphosphokinase subunit prs2  PRS1 - ribose phosphate diphosphokinase subunit prs1  CDC123 - cdc123p  LYS12 - homoisocitrate dehydrogenase |
| GO:0097159 | organic cyclic compound binding | 7.69E-4 | 4.16E-2 | 1.18 (2325,928,586,275) | [+] Show genes  IDH1 - isocitrate dehydrogenase (nad(+)) idh1  TFA2 - tfa2p  MSC6 - msc6p  GSH1 - gsh1p  UTP30 - utp30p  ARO2 - bifunctional chorismate synthase/riboflavin reductase [nad(p)h] aro2  LHP1 - lhp1p  RPL9A - ribosomal 60s subunit protein l9a  SAM2 - methionine adenosyltransferase sam2  RPL8A - ribosomal 60s subunit protein l8a  FOL3 - dihydrofolate synthase  APA1 - apa1p  GLE2 - gle2p  FPR2 - peptidylprolyl isomerase family protein fpr2  NAP1 - nap1p  LSM4 - lsm4p  GLK1 - glucokinase  ARP2 - actin-related protein 2  PRS2 - ribose phosphate diphosphokinase subunit prs2  STO1 - sto1p  EMI2 - putative glucokinase  SRO9 - sro9p  MLP1 - mlp1p  FAA1 - long-chain fatty acid-coa ligase faa1  SNF1 - snf1p  HIS4 - trifunctional histidinol dehydrogenase/phosphoribosyl-amp cyclohydrolase/phosphoribosyl-atp diphosphatase  ILV2 - acetolactate synthase catalytic subunit  LSM5 - lsm5p  SNF4 - snf4p  HRB1 - hrb1p  SPT15 - spt15p  RPS2 - ribosomal 40s subunit protein s2  SNQ2 - atp-binding cassette transporter snq2  CPA1 - carbamoyl-phosphate synthase (glutamine-hydrolyzing) cpa1  HBS1 - hbs1p  ARG5,6 - bifunctional acetylglutamate kinase/n-acetyl-gamma-glutamyl-phosphate reductase  MOT2 - ccr4-not core ubiquitin-protein ligase subunit mot2  GPH1 - gph1p  CDC10 - septin cdc10  RPB9 - rpb9p  VPS1 - vps1p  YPK9 - ypk9p  DCS1 - dcs1p  ASN1 - asparagine synthase (glutamine-hydrolyzing) 1  HIS1 - atp phosphoribosyltransferase  GUK1 - guanylate kinase  YSH1 - ysh1p  SRP14 - srp14p  HOM3 - aspartate kinase  MCM5 - mcm5p  RSM18 - mitochondrial 37s ribosomal protein rsm18  TDH1 - tdh1p  HSC82 - hsp90 family chaperone hsc82  RRP43 - rrp43p  NOG2 - nog2p  RPB7 - rpb7p  CDC42 - cdc42p  BNA5 - kynureninase  SUB2 - sub2p  SCD6 - scd6p  ILV1 - threonine ammonia-lyase ilv1  TIF4632 - tif4632p  MVD1 - diphosphomevalonate decarboxylase mvd1  OLE1 - stearoyl-coa 9-desaturase  PDR15 - atp-binding cassette multidrug transporter pdr15  HSP82 - hsp90 family chaperone hsp82  RFC1 - replication factor c subunit 1  LRP1 - lrp1p  FUS3 - fus3p  YOR1 - atp-binding cassette transporter yor1  TIF11 - tif11p  SCW4 - scw4p  REI1 - rei1p  FAA4 - long-chain fatty acid-coa ligase faa4  LHS1 - hsp70 family chaperone lhs1  ILV5 - ketol-acid reductoisomerase  BFR1 - bfr1p  GAD1 - glutamate decarboxylase gad1  DNF2 - aminophospholipid-translocating p4-type atpase dnf2  FOL2 - gtp cyclohydrolase i  YCR087C-A - hypothetical protein  RPL26A - ribosomal 60s subunit protein l26a  YHM2 - yhm2p  RPP0 - ribosomal protein p0  MDH1 - malate dehydrogenase mdh1  URA6 - bifunctional uridylate/adenylate kinase  TUF1 - tuf1p  RRP3 - rna-dependent atpase rrp3  DPB4 - dpb4p  TRM1 - trm1p  GLN4 - glutamine--trna ligase  YBL036C - hypothetical protein  ARO1 - pentafunctional protein aro1p  CDC3 - septin cdc3  KIN1 - kin1p  HEK2 - hek2p  YCF1 - atp-binding cassette glutathione s-conjugate transporter ycf1  YHB1 - yhb1p  ERG12 - mevalonate kinase  SMI1 - smi1p  NCP1 - ncp1p  RVB2 - ruvb family atp-dependent dna helicase reptin  DEF1 - def1p  MYO3 - myosin 3  PDR5 - atp-binding cassette multidrug transporter pdr5  ARO8 - bifunctional 2-aminoadipate transaminase/aromatic-amino-acid:2-oxoglutarate transaminase  AFG2 - aaa family atpase afg2  ACT1 - actin  SHE2 - she2p  THI80 - thiamine diphosphokinase  LSC1 - succinate--coa ligase (gdp-forming) subunit alpha  ARP8 - arp8p  HXK2 - hexokinase 2  VIP1 - inositol polyphosphate kinase vip1  GON7 - gon7p  IDH2 - isocitrate dehydrogenase (nad(+)) idh2  KRS1 - lysine--trna ligase krs1  CCA1 - cca1p  NEW1 - new1p  YHR020W - proline--trna ligase  APA2 - apa2p  LCB1 - serine c-palmitoyltransferase lcb1  RPL14B - ribosomal 60s subunit protein l14b  CAB1 - pantothenate kinase  LEO1 - leo1p  SEC4 - sec4p  POS5 - pos5p  LCB2 - serine c-palmitoyltransferase lcb2  TFC7 - tfc7p  RPS13 - ribosomal 40s subunit protein s13  RAS1 - ras1p  ARL3 - arl3p  STI1 - sti1p  PCF11 - pcf11p  MYO4 - myosin 4  FAA3 - long-chain fatty acid-coa ligase faa3  UTP14 - utp14p  ASN2 - asparagine synthase (glutamine-hydrolyzing) 2  TCP1 - tcp1p  CLA4 - cla4p  SPT6 - spt6p  GCD6 - gcd6p  TUB1 - tub1p  FES1 - fes1p  YDR210C-C - gag protein  NOP4 - nop4p  MET10 - sulfite reductase subunit alpha  NOB1 - nob1p  RSM7 - rsm7p  NHP6B - nhp6bp  POL30 - pol30p  RFC3 - replication factor c subunit 3  CPA2 - cpa2p  SSA2 - hsp70 family chaperone ssa2  GLR1 - glutathione-disulfide reductase glr1  FPR4 - peptidylprolyl isomerase fpr4  UTP9 - utp9p  RPL6B - ribosomal 60s subunit protein l6b  YNL284C-B - gag-pol fusion protein  BUD21 - bud21p  CYS4 - cystathionine beta-synthase cys4  YDR248C - gluconokinase  BCY1 - bcy1p  SEC18 - sec18p  PTI1 - pti1p  RPS5 - rps5p  HXK1 - hexokinase 1  CAR2 - ornithine-oxo-acid transaminase  ARF3 - arf3p  RPL24B - ribosomal 60s subunit protein l24b  VPS21 - vps21p  HSP26 - hsp26p  SNU114 - snu114p  CDC12 - septin cdc12  DAK1 - dak1p  YRA2 - yra2p  SER33 - phosphoglycerate dehydrogenase ser33  OXP1 - oxp1p  GRE3 - trifunctional aldehyde reductase/xylose reductase/glucose 1-dehydrogenase (nadp(+))  RPA49 - rpa49p  SSE1 - sse1p  FUR1 - uracil phosphoribosyltransferase  SUI1 - sui1p  MAM33 - mam33p  BPT1 - atp-binding cassette bilirubin transporter bpt1  HOM2 - aspartate-semialdehyde dehydrogenase  ZWF1 - glucose-6-phosphate dehydrogenase  ELP4 - elongator subunit elp4  DPS1 - aspartate--trna ligase dps1  CPR1 - peptidylprolyl isomerase cpr1  ERG1 - squalene monooxygenase  LAP3 - lap3p  RPC10 - rpc10p  TOP1 - dna topoisomerase 1  PDX3 - pyridoxamine-phosphate oxidase pdx3  RPL6A - ribosomal 60s subunit protein l6a  ARO9 - aromatic-amino-acid:2-oxoglutarate transaminase  RHO3 - rho3p  SKI6 - ski6p  RVB1 - ruvb family atp-dependent dna helicase pontin  OLA1 - ola1p  YCK1 - yck1p  CDC60 - leucine--trna ligase cdc60  RAP1 - rap1p  YBT1 - bile acid-transporting atpase ybt1  CCT6 - cct6p  RPL5 - ribosomal 60s subunit protein l5  HMO1 - hmo1p  KES1 - kes1p  DNM1 - dnm1p  MRPL6 - mitochondrial 54s ribosomal protein yml16  PRS1 - ribose phosphate diphosphokinase subunit prs1  LYS12 - homoisocitrate dehydrogenase  CCT2 - cct2p  RPL24A - ribosomal 60s subunit protein l24a  RPS3 - ribosomal 40s subunit protein s3  RPL30 - ribosomal 60s subunit protein l30  MRPL19 - mitochondrial 54s ribosomal protein yml19  POL31 - pol31p  SRP54 - srp54p  PDC1 - indolepyruvate decarboxylase 1  KGD1 - alpha-ketoglutarate dehydrogenase kgd1  ABD1 - abd1p  STH1 - sth1p  NOP13 - nop13p  ASG1 - asg1p  RPL15A - ribosomal 60s subunit protein l15a  HTS1 - histidine--trna ligase  ARG1 - argininosuccinate synthase  GLN1 - glutamate--ammonia ligase  PCS60 - pcs60p  YCK2 - yck2p  PRS5 - ribose phosphate diphosphokinase subunit prs5  YGR017W - hypothetical protein  NOP12 - nop12p  NRK1 - ribosylnicotinamide kinase  NOP9 - nop9p  DUR1,2 - bifunctional urea carboxylase/allophanate hydrolase  NHP6A - nhp6ap  MEF1 - mef1p  POP6 - pop6p  REX2 - rex2p  AIM45 - aim45p  ARP3 - arp3p  RPS9B - ribosomal 40s subunit protein s9b  RPL25 - ribosomal 60s subunit protein l25  ACB1 - long-chain fatty acid transporter acb1  CDC48 - aaa family atpase cdc48  DLD3 - dld3p  RPL26B - ribosomal 60s subunit protein l26b  CDC11 - cdc11p  PRO1 - glutamate 5-kinase  RAS2 - ras2p  YGR054W - hypothetical protein  BPL1 - biotin--[acetyl-coa-carboxylase] ligase bpl1  TYR1 - pprephenate dehydrogenase (nadp(+))  SSE2 - sse2p  YMR074C - hypothetical protein  RPO21 - rpo21p  RRP45 - rrp45p  POL1 - pol1p  STM1 - stm1p  MET12 - methylenetetrahydrofolate reductase (nad(p)h) met12  GLT1 - glutamate synthase (nadh)  SPF1 - spf1p  DLD1 - dld1p  UBC7 - e2 ubiquitin-conjugating protein ubc7  MCM3 - mcm3p  DLD2 - dld2p  PDC5 - indolepyruvate decarboxylase 5  RRP46 - rrp46p  CCT5 - cct5p  SSB1 - hsp70 family atpase ssb1  RPL16B - ribosomal 60s subunit protein l16b  SUB1 - sub1p |
| GO:0015926 | glucosidase activity | 9.94E-4 | 5.23E-2 | 9.24 (2325,17,74,5) | [+] Show genes  YMR196W - hypothetical protein  KRE6 - kre6p  EXG1 - exg1p  CWH41 - cwh41p  SCW4 - scw4p |

Species used: Saccharomyces cerevisiae

The system has recognized 2343 genes out of 2345 gene terms entered by the user.  
 141 genes were recognized by gene symbol and 2202 genes by other gene IDs .  
Only 2325 of these genes are associated with a GO term.

  
 Output in Microsoft Excel format  

% List genereted using GOrilla
% http://cbl-gorilla.cs.technion.ac.il/
% GO term pValue
GO:0050662 6.72E-10
GO:0048037 1.57E-8
GO:0003824 4.78E-8
GO:0016829 4.59E-6
GO:0036094 1.28E-5
GO:0016614 3.51E-5
GO:0043167 4E-5
GO:0043168 8.52E-5
GO:0050660 1.14E-4
GO:0005524 1.17E-4
GO:0032559 1.28E-4
GO:0030554 1.34E-4
GO:0003735 1.38E-4
GO:0016616 2.38E-4
GO:0008144 2.52E-4
GO:0003849 2.8E-4
GO:0016491 3.02E-4
GO:0019842 3.39E-4
GO:0035639 3.46E-4
GO:0032555 3.46E-4
GO:0004042 3.67E-4
GO:0017076 3.95E-4
GO:0004039 4.3E-4
GO:0004847 4.3E-4
GO:0032553 4.55E-4
GO:0097367 4.93E-4
GO:0004449 5.2E-4
GO:0051287 5.23E-4
GO:1901265 5.27E-4
GO:0000166 5.27E-4
GO:0004553 5.43E-4
GO:1901363 6.19E-4
GO:0004030 6.37E-4
GO:0000287 7.04E-4
GO:0097159 7.69E-4
GO:0015926 9.94E-4
 Visualize output in REViGO   

The GOrilla database is periodically updated using the GO database and other sources.  
The GOrilla database was last updated on Sep 21, 2019

This results page will be available on this site for one month from now (until
Oct 23, 2019
). You can bookmark this page and come back to it later.

  
**'P-value'** is the enrichment
p-value computed according to the mHG or HG model. This p-value is not
corrected for multiple testing of 1895 GO terms.  
  
**'FDR q-value'** is the correction of the above p-value for multiple testing using the Benjamini and Hochberg (1995) method.   
Namely, for the i^th^ term (ranked according to p-value) the FDR q-value is (p-value \* number of GO terms) / i.   
  
**Enrichment (N, B, n, b)** is defined as follows:  
N - is the total number of genes  
B - is the total number of genes associated with a specific GO term  
n - is the number of genes in the top of the user's input list or in the target set when appropriate  
b - is the number of genes in the intersection  
Enrichment = (b/n) / (B/N)  
  
**Genes:** For each GO term you can see the list of associated genes that appear in the optimal top of the list.  
Each gene name is specified by gene symbol followed by a short description of the gene   

Back to GOrilla analysis results
