## Supplemental Data S1 for "A simple mass-action model predicts genome-wide protein timecourses from mRNA trajectories during a dynamic response in two strains of *Saccharomyces cerevisiae*": GOFUNCTION_rna.html

Results


 **rna** 

*P-value color scale*

|  |  |  |  |  |
| --- | --- | --- | --- | --- |
| > 10^-3^ | 10^-3^ to 10^-5^ | 10^-5^ to 10^-7^ | 10^-7^ to 10^-9^ | < 10^-9^ |


|  |  |  |  |  |  |
| --- | --- | --- | --- | --- | --- |
| **GO term** | **Description** | **P-value** | **FDR q-value** | **Enrichment (N, B, n, b)** | **Genes** |
| GO:0003735 | structural constituent of ribosome | 1.37E-46 | 3.4E-43 | 10.38 (5247,199,155,61) | [+] Show genes  RPL29 - ribosomal 60s subunit protein l29  RPL2B - ribosomal 60s subunit protein l2b  RPS0B - ribosomal 40s subunit protein s0b  RPL24A - ribosomal 60s subunit protein l24a  RPL30 - ribosomal 60s subunit protein l30  RPL9A - ribosomal 60s subunit protein l9a  RPL23B - ribosomal 60s subunit protein l23b  RPL40A - ubiquitin-ribosomal 60s subunit protein l40a fusion protein  RPL38 - ribosomal 60s subunit protein l38  RPL13B - ribosomal 60s subunit protein l13b  RPS16A - ribosomal 40s subunit protein s16a  RPL8A - ribosomal 60s subunit protein l8a  RPS8B - ribosomal 40s subunit protein s8b  RPL26A - ribosomal 60s subunit protein l26a  RPL11A - ribosomal 60s subunit protein l11a  RPL15A - ribosomal 60s subunit protein l15a  RPP0 - ribosomal protein p0  RPL27B - ribosomal 60s subunit protein l27b  RPL34B - ribosomal 60s subunit protein l34b  RPL3 - ribosomal 60s subunit protein l3  RPS18B - ribosomal 40s subunit protein s18b  RPL19A - ribosomal 60s subunit protein l19a  RPS21B - rps21bp  RPS10B - ribosomal 40s subunit protein s10b  RPP2B - ribosomal protein p2b  RLP7 - rlp7p  RPL6B - ribosomal 60s subunit protein l6b  RPS15 - ribosomal 40s subunit protein s15  RPP2A - ribosomal protein p2a  RPS5 - rps5p  RPS17A - ribosomal 40s subunit protein s17a  RPL37A - ribosomal 60s subunit protein l37a  RPL24B - ribosomal 60s subunit protein l24b  RPS28A - ribosomal 40s subunit protein s28a  RPS27B - ribosomal 40s subunit protein s27b  RPL17A - ribosomal 60s subunit protein l17a  RPS1B - ribosomal 40s subunit protein s1b  RPP1B - ribosomal protein p1b  RPL21A - ribosomal 60s subunit protein l21a  RPL14A - ribosomal 60s subunit protein l14a  RPL17B - rpl17bp  MRPL24 - mitochondrial 54s ribosomal protein yml24/yml14  RPL32 - ribosomal 60s subunit protein l32  RPL26B - ribosomal 60s subunit protein l26b  RPL22B - ribosomal 60s subunit protein l22b  RPL35B - ribosomal 60s subunit protein l35b  RPS10A - ribosomal 40s subunit protein s10a  RPL34A - ribosomal 60s subunit protein l34a  RPL27A - ribosomal 60s subunit protein l27a  RPS22A - rps22ap  RPS17B - ribosomal 40s subunit protein s17b  RPL6A - ribosomal 60s subunit protein l6a  RPS23B - ribosomal 40s subunit protein s23b  RPL42B - ribosomal 60s subunit protein l42b  RPS14A - ribosomal 40s subunit protein s14a  RPL31A - ribosomal 60s subunit protein l31a  RPS13 - ribosomal 40s subunit protein s13  RPS26A - ribosomal 40s subunit protein s26a  RPS7A - ribosomal 40s subunit protein s7a  RPL16B - ribosomal 60s subunit protein l16b  RPS24A - ribosomal 40s subunit protein s24a |
| GO:0005198 | structural molecule activity | 2.84E-41 | 3.51E-38 | 7.28 (5247,316,155,68) | [+] Show genes  RPS0B - ribosomal 40s subunit protein s0b  YLR194C - hypothetical protein  RPL29 - ribosomal 60s subunit protein l29  RPL2B - ribosomal 60s subunit protein l2b  RPL24A - ribosomal 60s subunit protein l24a  RPL30 - ribosomal 60s subunit protein l30  SED1 - sed1p  RPL9A - ribosomal 60s subunit protein l9a  RPL40A - ubiquitin-ribosomal 60s subunit protein l40a fusion protein  RPL23B - ribosomal 60s subunit protein l23b  RPL38 - ribosomal 60s subunit protein l38  RPL13B - ribosomal 60s subunit protein l13b  RPS16A - ribosomal 40s subunit protein s16a  RPL8A - ribosomal 60s subunit protein l8a  RPS8B - ribosomal 40s subunit protein s8b  RPL26A - ribosomal 60s subunit protein l26a  RPL15A - ribosomal 60s subunit protein l15a  RPL11A - ribosomal 60s subunit protein l11a  RPP0 - ribosomal protein p0  RPL27B - ribosomal 60s subunit protein l27b  RPL34B - ribosomal 60s subunit protein l34b  RPL3 - ribosomal 60s subunit protein l3  RPS18B - ribosomal 40s subunit protein s18b  RPL19A - ribosomal 60s subunit protein l19a  RPS21B - rps21bp  RPS10B - ribosomal 40s subunit protein s10b  RPP2B - ribosomal protein p2b  RLP7 - rlp7p  RPL6B - ribosomal 60s subunit protein l6b  RPS15 - ribosomal 40s subunit protein s15  RPP2A - ribosomal protein p2a  RPS5 - rps5p  RPS17A - ribosomal 40s subunit protein s17a  RPL37A - ribosomal 60s subunit protein l37a  HSP150 - hsp150p  RPL24B - ribosomal 60s subunit protein l24b  CIS3 - cis3p  RPS28A - ribosomal 40s subunit protein s28a  RPS27B - ribosomal 40s subunit protein s27b  RPL17A - ribosomal 60s subunit protein l17a  RPS1B - ribosomal 40s subunit protein s1b  RPL21A - ribosomal 60s subunit protein l21a  RPP1B - ribosomal protein p1b  RPL14A - ribosomal 60s subunit protein l14a  RPL17B - rpl17bp  MRPL24 - mitochondrial 54s ribosomal protein yml24/yml14  RPL32 - ribosomal 60s subunit protein l32  RPL26B - ribosomal 60s subunit protein l26b  PAU7 - pau7p  RPL22B - ribosomal 60s subunit protein l22b  RPL35B - ribosomal 60s subunit protein l35b  RPS10A - ribosomal 40s subunit protein s10a  ACT1 - actin  RPL34A - ribosomal 60s subunit protein l34a  RPL27A - ribosomal 60s subunit protein l27a  RPS22A - rps22ap  RPS17B - ribosomal 40s subunit protein s17b  RPL6A - ribosomal 60s subunit protein l6a  RPS23B - ribosomal 40s subunit protein s23b  RPL42B - ribosomal 60s subunit protein l42b  RPS14A - ribosomal 40s subunit protein s14a  RPL31A - ribosomal 60s subunit protein l31a  CWP2 - cwp2p  RPS13 - ribosomal 40s subunit protein s13  RPS26A - ribosomal 40s subunit protein s26a  RPS7A - ribosomal 40s subunit protein s7a  RPL16B - ribosomal 60s subunit protein l16b  RPS24A - ribosomal 40s subunit protein s24a |
| GO:0005199 | structural constituent of cell wall | 2E-6 | 1.65E-3 | 73.38 (5247,22,13,4) | [+] Show genes  YLR194C - hypothetical protein  SED1 - sed1p  CWP2 - cwp2p  CIS3 - cis3p |
| GO:0019843 | rRNA binding | 3.03E-6 | 1.87E-3 | 3.39 (5247,70,487,22) | [+] Show genes  RPL2B - ribosomal 60s subunit protein l2b  RPL25 - ribosomal 60s subunit protein l25  RPS14A - ribosomal 40s subunit protein s14a  RPS18B - ribosomal 40s subunit protein s18b  RPL9A - ribosomal 60s subunit protein l9a  RPL23B - ribosomal 60s subunit protein l23b  RPF1 - rpf1p  RLP7 - rlp7p  RPS11B - ribosomal 40s subunit protein s11b  RPL12A - ribosomal 60s subunit protein l12a  RPS2 - ribosomal 40s subunit protein s2  RPL11B - ribosomal 60s subunit protein l11b  SSF1 - ssf1p  RPS13 - ribosomal 40s subunit protein s13  RPS14B - rps14bp  RPS5 - rps5p  RPL37A - ribosomal 60s subunit protein l37a  YEF3 - yef3p  RPL11A - ribosomal 60s subunit protein l11a  RPS9A - ribosomal 40s subunit protein s9a  MRD1 - mrd1p  RPP0 - ribosomal protein p0 |
| GO:0003723 | RNA binding | 1.78E-5 | 8.81E-3 | 2.04 (5247,563,210,46) | [+] Show genes  RPL2B - ribosomal 60s subunit protein l2b  ECM16 - ecm16p  RPL14A - ribosomal 60s subunit protein l14a  RPL24A - ribosomal 60s subunit protein l24a  RPL30 - ribosomal 60s subunit protein l30  DBP2 - dbp2p  HCA4 - hca4p  UTP30 - utp30p  RPL26B - ribosomal 60s subunit protein l26b  RPL22B - ribosomal 60s subunit protein l22b  RPL35B - ribosomal 60s subunit protein l35b  RPL9A - ribosomal 60s subunit protein l9a  RPL23B - ribosomal 60s subunit protein l23b  NCS2 - ncs2p  SCW4 - scw4p  RPS16A - ribosomal 40s subunit protein s16a  NSR1 - nsr1p  TEF1 - tef1p  YHR033W - putative glutamate 5-kinase  RPL8A - ribosomal 60s subunit protein l8a  RPL12A - ribosomal 60s subunit protein l12a  JSN1 - jsn1p  RPL16A - ribosomal 60s subunit protein l16a  RPL26A - ribosomal 60s subunit protein l26a  RPL15A - ribosomal 60s subunit protein l15a  RPL11A - ribosomal 60s subunit protein l11a  NEW1 - new1p  RPP0 - ribosomal protein p0  RPL6A - ribosomal 60s subunit protein l6a  RPS14A - ribosomal 40s subunit protein s14a  RPS18B - ribosomal 40s subunit protein s18b  RPL19A - ribosomal 60s subunit protein l19a  SUA5 - sua5p  SMB1 - smb1p  RPL6B - ribosomal 60s subunit protein l6b  RLP7 - rlp7p  RPS15 - ribosomal 40s subunit protein s15  TRM9 - trm9p  RPL11B - ribosomal 60s subunit protein l11b  RPS5 - rps5p  RPS13 - ribosomal 40s subunit protein s13  RPL37A - ribosomal 60s subunit protein l37a  RPS26A - ribosomal 40s subunit protein s26a  RPL16B - ribosomal 60s subunit protein l16b  RPL24B - ribosomal 60s subunit protein l24b  PWP2 - pwp2p |
| GO:0004100 | chitin synthase activity | 2.45E-4 | 1.01E-1 | 99.94 (5247,3,35,2) | [+] Show genes  CHS3 - chitin synthase chs3  CHS1 - chitin synthase chs1 |
| GO:0043531 | ADP binding | 2.73E-4 | 9.62E-2 | 19.08 (5247,3,275,3) | [+] Show genes  ATP1 - f1f0 atp synthase subunit alpha  HSP104 - chaperone atpase hsp104  PGK1 - phosphoglycerate kinase |

Species used: Saccharomyces cerevisiae

The system has recognized 5321 genes out of 5595 gene terms entered by the user.  
 628 genes were recognized by gene symbol and 4693 genes by other gene IDs .  
Only 5247 of these genes are associated with a GO term.

  
 Output in Microsoft Excel format  

% List genereted using GOrilla
% http://cbl-gorilla.cs.technion.ac.il/
% GO term pValue
GO:0003735 1.37E-46
GO:0005198 2.84E-41
GO:0005199 2E-6
GO:0019843 3.03E-6
GO:0003723 1.78E-5
GO:0004100 2.45E-4
GO:0043531 2.73E-4
 Visualize output in REViGO   

The GOrilla database is periodically updated using the GO database and other sources.  
The GOrilla database was last updated on Sep 21, 2019

This results page will be available on this site for one month from now (until
Oct 23, 2019
). You can bookmark this page and come back to it later.

  
**'P-value'** is the enrichment
p-value computed according to the mHG or HG model. This p-value is not
corrected for multiple testing of 2471 GO terms.  
  
**'FDR q-value'** is the correction of the above p-value for multiple testing using the Benjamini and Hochberg (1995) method.   
Namely, for the i^th^ term (ranked according to p-value) the FDR q-value is (p-value \* number of GO terms) / i.   
  
**Enrichment (N, B, n, b)** is defined as follows:  
N - is the total number of genes  
B - is the total number of genes associated with a specific GO term  
n - is the number of genes in the top of the user's input list or in the target set when appropriate  
b - is the number of genes in the intersection  
Enrichment = (b/n) / (B/N)  
  
**Genes:** For each GO term you can see the list of associated genes that appear in the optimal top of the list.  
Each gene name is specified by gene symbol followed by a short description of the gene   

Back to GOrilla analysis results
