## Supplemental Data S1 for "A simple mass-action model predicts genome-wide protein timecourses from mRNA trajectories during a dynamic response in two strains of *Saccharomyces cerevisiae*": GOPROCESS_par.html

Results


 **Parameters** 

*P-value color scale*

|  |  |  |  |  |
| --- | --- | --- | --- | --- |
| > 10^-3^ | 10^-3^ to 10^-5^ | 10^-5^ to 10^-7^ | 10^-7^ to 10^-9^ | < 10^-9^ |


|  |  |  |  |  |  |
| --- | --- | --- | --- | --- | --- |
| **GO term** | **Description** | **P-value** | **FDR q-value** | **Enrichment (N, B, n, b)** | **Genes** |
| GO:0044281 | small molecule metabolic process | 1.27E-18 | 5.22E-15 | 2.06 (2298,436,338,132) | [+] Show genes  IDH1 - isocitrate dehydrogenase (nad(+)) idh1  PHS1 - phs1p  ENO2 - phosphopyruvate hydratase eno2  ARO2 - bifunctional chorismate synthase/riboflavin reductase [nad(p)h] aro2  MET5 - met5p  GPD1 - glycerol-3-phosphate dehydrogenase (nad(+)) gpd1  APA1 - apa1p  ASN2 - asparagine synthase (glutamine-hydrolyzing) 2  YSA1 - ysa1p  GLK1 - glucokinase  BAT2 - bat2p  EMI2 - putative glucokinase  PPX1 - ppx1p  CPA2 - cpa2p  FAA1 - long-chain fatty acid-coa ligase faa1  PGM2 - phosphoglucomutase pgm2  HIS4 - trifunctional histidinol dehydrogenase/phosphoribosyl-amp cyclohydrolase/phosphoribosyl-atp diphosphatase  CIT1 - citrate (si)-synthase cit1  ILV2 - acetolactate synthase catalytic subunit  PRO2 - glutamate-5-semialdehyde dehydrogenase  HFD1 - hfd1p  RIB3 - 3,4-dihydroxy-2-butanone-4-phosphate synthase rib3  CPA1 - carbamoyl-phosphate synthase (glutamine-hydrolyzing) cpa1  ERG24 - delta(14)-sterol reductase  ATP2 - atp2p  CAR2 - ornithine-oxo-acid transaminase  HXK1 - hexokinase 1  THI4 - thi4p  ILV6 - acetolactate synthase regulatory subunit  ATP4 - atp4p  ARG5,6 - bifunctional acetylglutamate kinase/n-acetyl-gamma-glutamyl-phosphate reductase  CAB2 - phosphopantothenate--cysteine ligase cab2  YDL124W - aldo-keto reductase superfamily protein  HOR2 - glycerol-1-phosphatase hor2  GRE3 - trifunctional aldehyde reductase/xylose reductase/glucose 1-dehydrogenase (nadp(+))  ASN1 - asparagine synthase (glutamine-hydrolyzing) 1  KGD2 - alpha-ketoglutarate dehydrogenase kgd2  HIS1 - atp phosphoribosyltransferase  GUK1 - guanylate kinase  HOM2 - aspartate-semialdehyde dehydrogenase  ZWF1 - glucose-6-phosphate dehydrogenase  ATP7 - f1f0 atp synthase subunit d  HOM3 - aspartate kinase  HEM4 - uroporphyrinogen-iii synthase hem4  ATP3 - atp3p  URE2 - ure2p  PNC1 - nicotinamidase  MDH3 - malate dehydrogenase mdh3  ARO9 - aromatic-amino-acid:2-oxoglutarate transaminase  YGL039W - carbonyl reductase (nadph-dependent)  BDH1 - (r,r)-butanediol dehydrogenase  ERG13 - hydroxymethylglutaryl-coa synthase  TRP2 - anthranilate synthase trp2  BNA5 - kynureninase  CDC60 - leucine--trna ligase cdc60  ALD2 - aldehyde dehydrogenase (nad(+)) ald2  ILV1 - threonine ammonia-lyase ilv1  SDH2 - succinate dehydrogenase iron-sulfur protein subunit sdh2  OLE1 - stearoyl-coa 9-desaturase  NDE1 - nadh-ubiquinone reductase (h(+)-translocating) nde1  ALD5 - aldehyde dehydrogenase (nad(p)(+)) ald5  TRX3 - trx3p  TRP5 - tryptophan synthase trp5  RIB5 - riboflavin synthase  ILV3 - ilv3p  ACH1 - ach1p  ERG6 - sterol 24-c-methyltransferase  PDC1 - indolepyruvate decarboxylase 1  GLO1 - lactoylglutathione lyase glo1  KGD1 - alpha-ketoglutarate dehydrogenase kgd1  HIS3 - imidazoleglycerol-phosphate dehydratase his3  TAL1 - sedoheptulose-7-phosphate:d-glyceraldehyde-3-phosphate transaldolase tal1  ILV5 - ketol-acid reductoisomerase  ERG7 - lanosterol synthase erg7  GAD1 - glutamate decarboxylase gad1  ENO1 - phosphopyruvate hydratase eno1  FOL2 - gtp cyclohydrolase i  YDR341C - arginine--trna ligase  COQ1 - trans-hexaprenyltranstransferase  MDH1 - malate dehydrogenase mdh1  ADE16 - bifunctional phosphoribosylaminoimidazolecarboxamide formyltransferase/imp cyclohydrolase ade16  ARO4 - 3-deoxy-7-phosphoheptulonate synthase aro4  MET17 - bifunctional cysteine synthase/o-acetylhomoserine aminocarboxypropyltransferase met17  FUM1 - fumarase fum1  HTS1 - histidine--trna ligase  MAE1 - malate dehydrogenase (oxaloacetate-decarboxylating)  ARG1 - argininosuccinate synthase  GLN1 - glutamate--ammonia ligase  PCS60 - pcs60p  UGP1 - utp glucose-1-phosphate uridylyltransferase  GDH2 - glutamate dehydrogenase (nad(+))  ATP15 - f1f0 atp synthase subunit epsilon  UGA1 - 4-aminobutyrate transaminase  AMD1 - amp deaminase  ARO1 - pentafunctional protein aro1p  DUR1,2 - bifunctional urea carboxylase/allophanate hydrolase  GUT2 - glycerol-3-phosphate dehydrogenase  ADE5,7 - bifunctional aminoimidazole ribotide synthase/glycinamide ribotide synthase  ERG12 - mevalonate kinase  ERG3 - c-5 sterol desaturase  FCY1 - cytosine deaminase  SHM2 - glycine hydroxymethyltransferase shm2  PGM1 - phosphoglucomutase pgm1  ARG7 - glutamate n-acetyltransferase  ACB1 - long-chain fatty acid transporter acb1  ISN1 - imp 5'-nucleotidase  DLD3 - dld3p  ARO8 - bifunctional 2-aminoadipate transaminase/aromatic-amino-acid:2-oxoglutarate transaminase  ATP5 - atp5p  LSC1 - succinate--coa ligase (gdp-forming) subunit alpha  YMR315W - hypothetical protein  HXK2 - hexokinase 2  ARO3 - 3-deoxy-7-phosphoheptulonate synthase aro3  EHD3 - ehd3p  IDH2 - isocitrate dehydrogenase (nad(+)) idh2  KRS1 - lysine--trna ligase krs1  GNA1 - glucosamine 6-phosphate n-acetyltransferase  ADE6 - phosphoribosylformylglycinamidine synthase  YKL151C - nadhx dehydratase  MCR1 - mcr1p  EHT1 - eht1p  YHR020W - proline--trna ligase  SDH1 - succinate dehydrogenase flavoprotein subunit sdh1  ADE4 - amidophosphoribosyltransferase  ADE2 - phosphoribosylaminoimidazole carboxylase ade2  ALD4 - aldehyde dehydrogenase (nadp(+)) ald4  GLT1 - glutamate synthase (nadh)  DLD1 - dld1p  GFA1 - glutamine--fructose-6-phosphate transaminase (isomerizing) gfa1  INP53 - phosphatidylinositol-3-/phosphoinositide 5-phosphatase inp53  DLD2 - dld2p  ATP1 - f1f0 atp synthase subunit alpha |
| GO:0019752 | carboxylic acid metabolic process | 4.6E-14 | 9.49E-11 | 2.46 (2298,252,285,77) | [+] Show genes  IDH1 - isocitrate dehydrogenase (nad(+)) idh1  PHS1 - phs1p  TRP5 - tryptophan synthase trp5  ENO2 - phosphopyruvate hydratase eno2  ARO2 - bifunctional chorismate synthase/riboflavin reductase [nad(p)h] aro2  ILV3 - ilv3p  ACH1 - ach1p  MET5 - met5p  PDC1 - indolepyruvate decarboxylase 1  GLO1 - lactoylglutathione lyase glo1  KGD1 - alpha-ketoglutarate dehydrogenase kgd1  HIS3 - imidazoleglycerol-phosphate dehydratase his3  ILV5 - ketol-acid reductoisomerase  GAD1 - glutamate decarboxylase gad1  ENO1 - phosphopyruvate hydratase eno1  ASN2 - asparagine synthase (glutamine-hydrolyzing) 2  GLK1 - glucokinase  YDR341C - arginine--trna ligase  BAT2 - bat2p  EMI2 - putative glucokinase  MDH1 - malate dehydrogenase mdh1  ARO4 - 3-deoxy-7-phosphoheptulonate synthase aro4  FUM1 - fumarase fum1  HTS1 - histidine--trna ligase  FAA1 - long-chain fatty acid-coa ligase faa1  CPA2 - cpa2p  GLN1 - glutamate--ammonia ligase  ARG1 - argininosuccinate synthase  MAE1 - malate dehydrogenase (oxaloacetate-decarboxylating)  PCS60 - pcs60p  HIS4 - trifunctional histidinol dehydrogenase/phosphoribosyl-amp cyclohydrolase/phosphoribosyl-atp diphosphatase  CIT1 - citrate (si)-synthase cit1  PRO2 - glutamate-5-semialdehyde dehydrogenase  ILV2 - acetolactate synthase catalytic subunit  HFD1 - hfd1p  ARO1 - pentafunctional protein aro1p  DUR1,2 - bifunctional urea carboxylase/allophanate hydrolase  CAR2 - ornithine-oxo-acid transaminase  HXK1 - hexokinase 1  ILV6 - acetolactate synthase regulatory subunit  SHM2 - glycine hydroxymethyltransferase shm2  ARG5,6 - bifunctional acetylglutamate kinase/n-acetyl-gamma-glutamyl-phosphate reductase  ARG7 - glutamate n-acetyltransferase  ACB1 - long-chain fatty acid transporter acb1  DLD3 - dld3p  ARO8 - bifunctional 2-aminoadipate transaminase/aromatic-amino-acid:2-oxoglutarate transaminase  ASN1 - asparagine synthase (glutamine-hydrolyzing) 1  KGD2 - alpha-ketoglutarate dehydrogenase kgd2  HIS1 - atp phosphoribosyltransferase  LSC1 - succinate--coa ligase (gdp-forming) subunit alpha  HOM2 - aspartate-semialdehyde dehydrogenase  HXK2 - hexokinase 2  HOM3 - aspartate kinase  ARO3 - 3-deoxy-7-phosphoheptulonate synthase aro3  EHD3 - ehd3p  HEM4 - uroporphyrinogen-iii synthase hem4  IDH2 - isocitrate dehydrogenase (nad(+)) idh2  KRS1 - lysine--trna ligase krs1  ADE6 - phosphoribosylformylglycinamidine synthase  EHT1 - eht1p  YHR020W - proline--trna ligase  MDH3 - malate dehydrogenase mdh3  ALD4 - aldehyde dehydrogenase (nadp(+)) ald4  ARO9 - aromatic-amino-acid:2-oxoglutarate transaminase  GLT1 - glutamate synthase (nadh)  TRP2 - anthranilate synthase trp2  DLD1 - dld1p  BNA5 - kynureninase  GFA1 - glutamine--fructose-6-phosphate transaminase (isomerizing) gfa1  CDC60 - leucine--trna ligase cdc60  ILV1 - threonine ammonia-lyase ilv1  ALD2 - aldehyde dehydrogenase (nad(+)) ald2  SDH2 - succinate dehydrogenase iron-sulfur protein subunit sdh2  DLD2 - dld2p  OLE1 - stearoyl-coa 9-desaturase  NDE1 - nadh-ubiquinone reductase (h(+)-translocating) nde1  ALD5 - aldehyde dehydrogenase (nad(p)(+)) ald5 |
| GO:0043436 | oxoacid metabolic process | 5.41E-14 | 7.44E-11 | 2.21 (2298,264,338,86) | [+] Show genes  IDH1 - isocitrate dehydrogenase (nad(+)) idh1  PHS1 - phs1p  TRP5 - tryptophan synthase trp5  ENO2 - phosphopyruvate hydratase eno2  ARO2 - bifunctional chorismate synthase/riboflavin reductase [nad(p)h] aro2  ILV3 - ilv3p  ACH1 - ach1p  MET5 - met5p  PDC1 - indolepyruvate decarboxylase 1  GLO1 - lactoylglutathione lyase glo1  KGD1 - alpha-ketoglutarate dehydrogenase kgd1  HIS3 - imidazoleglycerol-phosphate dehydratase his3  ILV5 - ketol-acid reductoisomerase  GAD1 - glutamate decarboxylase gad1  ASN2 - asparagine synthase (glutamine-hydrolyzing) 2  ENO1 - phosphopyruvate hydratase eno1  FOL2 - gtp cyclohydrolase i  GLK1 - glucokinase  YDR341C - arginine--trna ligase  BAT2 - bat2p  EMI2 - putative glucokinase  MDH1 - malate dehydrogenase mdh1  ARO4 - 3-deoxy-7-phosphoheptulonate synthase aro4  MET17 - bifunctional cysteine synthase/o-acetylhomoserine aminocarboxypropyltransferase met17  PPX1 - ppx1p  FUM1 - fumarase fum1  HTS1 - histidine--trna ligase  CPA2 - cpa2p  FAA1 - long-chain fatty acid-coa ligase faa1  ARG1 - argininosuccinate synthase  GLN1 - glutamate--ammonia ligase  MAE1 - malate dehydrogenase (oxaloacetate-decarboxylating)  PCS60 - pcs60p  HIS4 - trifunctional histidinol dehydrogenase/phosphoribosyl-amp cyclohydrolase/phosphoribosyl-atp diphosphatase  CIT1 - citrate (si)-synthase cit1  ILV2 - acetolactate synthase catalytic subunit  PRO2 - glutamate-5-semialdehyde dehydrogenase  GDH2 - glutamate dehydrogenase (nad(+))  UGA1 - 4-aminobutyrate transaminase  HFD1 - hfd1p  ARO1 - pentafunctional protein aro1p  DUR1,2 - bifunctional urea carboxylase/allophanate hydrolase  CPA1 - carbamoyl-phosphate synthase (glutamine-hydrolyzing) cpa1  CAR2 - ornithine-oxo-acid transaminase  HXK1 - hexokinase 1  ILV6 - acetolactate synthase regulatory subunit  SHM2 - glycine hydroxymethyltransferase shm2  ARG5,6 - bifunctional acetylglutamate kinase/n-acetyl-gamma-glutamyl-phosphate reductase  ACB1 - long-chain fatty acid transporter acb1  ARG7 - glutamate n-acetyltransferase  DLD3 - dld3p  ARO8 - bifunctional 2-aminoadipate transaminase/aromatic-amino-acid:2-oxoglutarate transaminase  ASN1 - asparagine synthase (glutamine-hydrolyzing) 1  KGD2 - alpha-ketoglutarate dehydrogenase kgd2  HIS1 - atp phosphoribosyltransferase  LSC1 - succinate--coa ligase (gdp-forming) subunit alpha  HOM2 - aspartate-semialdehyde dehydrogenase  HXK2 - hexokinase 2  HOM3 - aspartate kinase  ARO3 - 3-deoxy-7-phosphoheptulonate synthase aro3  EHD3 - ehd3p  HEM4 - uroporphyrinogen-iii synthase hem4  IDH2 - isocitrate dehydrogenase (nad(+)) idh2  KRS1 - lysine--trna ligase krs1  ADE6 - phosphoribosylformylglycinamidine synthase  EHT1 - eht1p  SDH1 - succinate dehydrogenase flavoprotein subunit sdh1  YHR020W - proline--trna ligase  ADE4 - amidophosphoribosyltransferase  URE2 - ure2p  MDH3 - malate dehydrogenase mdh3  ALD4 - aldehyde dehydrogenase (nadp(+)) ald4  ARO9 - aromatic-amino-acid:2-oxoglutarate transaminase  GLT1 - glutamate synthase (nadh)  TRP2 - anthranilate synthase trp2  DLD1 - dld1p  BNA5 - kynureninase  GFA1 - glutamine--fructose-6-phosphate transaminase (isomerizing) gfa1  CDC60 - leucine--trna ligase cdc60  ALD2 - aldehyde dehydrogenase (nad(+)) ald2  ILV1 - threonine ammonia-lyase ilv1  SDH2 - succinate dehydrogenase iron-sulfur protein subunit sdh2  DLD2 - dld2p  OLE1 - stearoyl-coa 9-desaturase  NDE1 - nadh-ubiquinone reductase (h(+)-translocating) nde1  ALD5 - aldehyde dehydrogenase (nad(p)(+)) ald5 |
| GO:0006082 | organic acid metabolic process | 6.92E-14 | 7.13E-11 | 2.21 (2298,265,338,86) | [+] Show genes  IDH1 - isocitrate dehydrogenase (nad(+)) idh1  PHS1 - phs1p  TRP5 - tryptophan synthase trp5  ENO2 - phosphopyruvate hydratase eno2  ARO2 - bifunctional chorismate synthase/riboflavin reductase [nad(p)h] aro2  ILV3 - ilv3p  ACH1 - ach1p  MET5 - met5p  PDC1 - indolepyruvate decarboxylase 1  GLO1 - lactoylglutathione lyase glo1  KGD1 - alpha-ketoglutarate dehydrogenase kgd1  HIS3 - imidazoleglycerol-phosphate dehydratase his3  ILV5 - ketol-acid reductoisomerase  GAD1 - glutamate decarboxylase gad1  ASN2 - asparagine synthase (glutamine-hydrolyzing) 2  ENO1 - phosphopyruvate hydratase eno1  FOL2 - gtp cyclohydrolase i  GLK1 - glucokinase  YDR341C - arginine--trna ligase  BAT2 - bat2p  EMI2 - putative glucokinase  MDH1 - malate dehydrogenase mdh1  ARO4 - 3-deoxy-7-phosphoheptulonate synthase aro4  MET17 - bifunctional cysteine synthase/o-acetylhomoserine aminocarboxypropyltransferase met17  PPX1 - ppx1p  FUM1 - fumarase fum1  HTS1 - histidine--trna ligase  CPA2 - cpa2p  FAA1 - long-chain fatty acid-coa ligase faa1  GLN1 - glutamate--ammonia ligase  ARG1 - argininosuccinate synthase  MAE1 - malate dehydrogenase (oxaloacetate-decarboxylating)  PCS60 - pcs60p  HIS4 - trifunctional histidinol dehydrogenase/phosphoribosyl-amp cyclohydrolase/phosphoribosyl-atp diphosphatase  CIT1 - citrate (si)-synthase cit1  ILV2 - acetolactate synthase catalytic subunit  PRO2 - glutamate-5-semialdehyde dehydrogenase  GDH2 - glutamate dehydrogenase (nad(+))  UGA1 - 4-aminobutyrate transaminase  HFD1 - hfd1p  ARO1 - pentafunctional protein aro1p  DUR1,2 - bifunctional urea carboxylase/allophanate hydrolase  CPA1 - carbamoyl-phosphate synthase (glutamine-hydrolyzing) cpa1  HXK1 - hexokinase 1  CAR2 - ornithine-oxo-acid transaminase  ILV6 - acetolactate synthase regulatory subunit  SHM2 - glycine hydroxymethyltransferase shm2  ARG5,6 - bifunctional acetylglutamate kinase/n-acetyl-gamma-glutamyl-phosphate reductase  ACB1 - long-chain fatty acid transporter acb1  ARG7 - glutamate n-acetyltransferase  DLD3 - dld3p  ARO8 - bifunctional 2-aminoadipate transaminase/aromatic-amino-acid:2-oxoglutarate transaminase  ASN1 - asparagine synthase (glutamine-hydrolyzing) 1  KGD2 - alpha-ketoglutarate dehydrogenase kgd2  HIS1 - atp phosphoribosyltransferase  LSC1 - succinate--coa ligase (gdp-forming) subunit alpha  HOM2 - aspartate-semialdehyde dehydrogenase  HXK2 - hexokinase 2  HOM3 - aspartate kinase  ARO3 - 3-deoxy-7-phosphoheptulonate synthase aro3  EHD3 - ehd3p  HEM4 - uroporphyrinogen-iii synthase hem4  KRS1 - lysine--trna ligase krs1  IDH2 - isocitrate dehydrogenase (nad(+)) idh2  ADE6 - phosphoribosylformylglycinamidine synthase  EHT1 - eht1p  YHR020W - proline--trna ligase  SDH1 - succinate dehydrogenase flavoprotein subunit sdh1  ADE4 - amidophosphoribosyltransferase  URE2 - ure2p  MDH3 - malate dehydrogenase mdh3  ALD4 - aldehyde dehydrogenase (nadp(+)) ald4  ARO9 - aromatic-amino-acid:2-oxoglutarate transaminase  GLT1 - glutamate synthase (nadh)  TRP2 - anthranilate synthase trp2  DLD1 - dld1p  BNA5 - kynureninase  GFA1 - glutamine--fructose-6-phosphate transaminase (isomerizing) gfa1  CDC60 - leucine--trna ligase cdc60  ALD2 - aldehyde dehydrogenase (nad(+)) ald2  ILV1 - threonine ammonia-lyase ilv1  SDH2 - succinate dehydrogenase iron-sulfur protein subunit sdh2  DLD2 - dld2p  OLE1 - stearoyl-coa 9-desaturase  NDE1 - nadh-ubiquinone reductase (h(+)-translocating) nde1  ALD5 - aldehyde dehydrogenase (nad(p)(+)) ald5 |
| GO:0044283 | small molecule biosynthetic process | 3.44E-11 | 2.84E-8 | 2.33 (2298,229,284,66) | [+] Show genes  IDH1 - isocitrate dehydrogenase (nad(+)) idh1  PHS1 - phs1p  TRP5 - tryptophan synthase trp5  RIB5 - riboflavin synthase  ENO2 - phosphopyruvate hydratase eno2  ARO2 - bifunctional chorismate synthase/riboflavin reductase [nad(p)h] aro2  ILV3 - ilv3p  MET5 - met5p  PDC1 - indolepyruvate decarboxylase 1  HIS3 - imidazoleglycerol-phosphate dehydratase his3  ILV5 - ketol-acid reductoisomerase  ERG7 - lanosterol synthase erg7  ENO1 - phosphopyruvate hydratase eno1  ASN2 - asparagine synthase (glutamine-hydrolyzing) 2  GLK1 - glucokinase  COQ1 - trans-hexaprenyltranstransferase  BAT2 - bat2p  EMI2 - putative glucokinase  ARO4 - 3-deoxy-7-phosphoheptulonate synthase aro4  CPA2 - cpa2p  GLN1 - glutamate--ammonia ligase  ARG1 - argininosuccinate synthase  HIS4 - trifunctional histidinol dehydrogenase/phosphoribosyl-amp cyclohydrolase/phosphoribosyl-atp diphosphatase  CIT1 - citrate (si)-synthase cit1  PRO2 - glutamate-5-semialdehyde dehydrogenase  ILV2 - acetolactate synthase catalytic subunit  HFD1 - hfd1p  RIB3 - 3,4-dihydroxy-2-butanone-4-phosphate synthase rib3  ARO1 - pentafunctional protein aro1p  ERG24 - delta(14)-sterol reductase  HXK1 - hexokinase 1  CAR2 - ornithine-oxo-acid transaminase  THI4 - thi4p  ERG3 - c-5 sterol desaturase  ILV6 - acetolactate synthase regulatory subunit  ERG12 - mevalonate kinase  SHM2 - glycine hydroxymethyltransferase shm2  FCY1 - cytosine deaminase  ARG5,6 - bifunctional acetylglutamate kinase/n-acetyl-gamma-glutamyl-phosphate reductase  ACB1 - long-chain fatty acid transporter acb1  ARG7 - glutamate n-acetyltransferase  DLD3 - dld3p  HOR2 - glycerol-1-phosphatase hor2  ARO8 - bifunctional 2-aminoadipate transaminase/aromatic-amino-acid:2-oxoglutarate transaminase  ASN1 - asparagine synthase (glutamine-hydrolyzing) 1  HIS1 - atp phosphoribosyltransferase  GUK1 - guanylate kinase  HXK2 - hexokinase 2  HOM2 - aspartate-semialdehyde dehydrogenase  HOM3 - aspartate kinase  ARO3 - 3-deoxy-7-phosphoheptulonate synthase aro3  HEM4 - uroporphyrinogen-iii synthase hem4  IDH2 - isocitrate dehydrogenase (nad(+)) idh2  MCR1 - mcr1p  EHT1 - eht1p  ALD4 - aldehyde dehydrogenase (nadp(+)) ald4  ARO9 - aromatic-amino-acid:2-oxoglutarate transaminase  GLT1 - glutamate synthase (nadh)  BDH1 - (r,r)-butanediol dehydrogenase  TRP2 - anthranilate synthase trp2  ERG13 - hydroxymethylglutaryl-coa synthase  BNA5 - kynureninase  ALD2 - aldehyde dehydrogenase (nad(+)) ald2  ILV1 - threonine ammonia-lyase ilv1  OLE1 - stearoyl-coa 9-desaturase  ALD5 - aldehyde dehydrogenase (nad(p)(+)) ald5 |
| GO:0016053 | organic acid biosynthetic process | 2.24E-10 | 1.54E-7 | 2.63 (2298,154,284,50) | [+] Show genes  IDH1 - isocitrate dehydrogenase (nad(+)) idh1  PHS1 - phs1p  TRP5 - tryptophan synthase trp5  ENO2 - phosphopyruvate hydratase eno2  ARO2 - bifunctional chorismate synthase/riboflavin reductase [nad(p)h] aro2  ILV3 - ilv3p  MET5 - met5p  HIS3 - imidazoleglycerol-phosphate dehydratase his3  ILV5 - ketol-acid reductoisomerase  ASN2 - asparagine synthase (glutamine-hydrolyzing) 2  ENO1 - phosphopyruvate hydratase eno1  GLK1 - glucokinase  BAT2 - bat2p  EMI2 - putative glucokinase  ARO4 - 3-deoxy-7-phosphoheptulonate synthase aro4  CPA2 - cpa2p  ARG1 - argininosuccinate synthase  GLN1 - glutamate--ammonia ligase  HIS4 - trifunctional histidinol dehydrogenase/phosphoribosyl-amp cyclohydrolase/phosphoribosyl-atp diphosphatase  CIT1 - citrate (si)-synthase cit1  PRO2 - glutamate-5-semialdehyde dehydrogenase  ILV2 - acetolactate synthase catalytic subunit  ARO1 - pentafunctional protein aro1p  HXK1 - hexokinase 1  CAR2 - ornithine-oxo-acid transaminase  ILV6 - acetolactate synthase regulatory subunit  SHM2 - glycine hydroxymethyltransferase shm2  ARG5,6 - bifunctional acetylglutamate kinase/n-acetyl-gamma-glutamyl-phosphate reductase  ARG7 - glutamate n-acetyltransferase  ACB1 - long-chain fatty acid transporter acb1  DLD3 - dld3p  ARO8 - bifunctional 2-aminoadipate transaminase/aromatic-amino-acid:2-oxoglutarate transaminase  ASN1 - asparagine synthase (glutamine-hydrolyzing) 1  HIS1 - atp phosphoribosyltransferase  HOM2 - aspartate-semialdehyde dehydrogenase  HXK2 - hexokinase 2  HOM3 - aspartate kinase  ARO3 - 3-deoxy-7-phosphoheptulonate synthase aro3  HEM4 - uroporphyrinogen-iii synthase hem4  IDH2 - isocitrate dehydrogenase (nad(+)) idh2  EHT1 - eht1p  ARO9 - aromatic-amino-acid:2-oxoglutarate transaminase  ALD4 - aldehyde dehydrogenase (nadp(+)) ald4  GLT1 - glutamate synthase (nadh)  TRP2 - anthranilate synthase trp2  BNA5 - kynureninase  ALD2 - aldehyde dehydrogenase (nad(+)) ald2  ILV1 - threonine ammonia-lyase ilv1  OLE1 - stearoyl-coa 9-desaturase  ALD5 - aldehyde dehydrogenase (nad(p)(+)) ald5 |
| GO:0046394 | carboxylic acid biosynthetic process | 2.24E-10 | 1.32E-7 | 2.63 (2298,154,284,50) | [+] Show genes  IDH1 - isocitrate dehydrogenase (nad(+)) idh1  PHS1 - phs1p  TRP5 - tryptophan synthase trp5  ENO2 - phosphopyruvate hydratase eno2  ARO2 - bifunctional chorismate synthase/riboflavin reductase [nad(p)h] aro2  ILV3 - ilv3p  MET5 - met5p  HIS3 - imidazoleglycerol-phosphate dehydratase his3  ILV5 - ketol-acid reductoisomerase  ASN2 - asparagine synthase (glutamine-hydrolyzing) 2  ENO1 - phosphopyruvate hydratase eno1  GLK1 - glucokinase  BAT2 - bat2p  EMI2 - putative glucokinase  ARO4 - 3-deoxy-7-phosphoheptulonate synthase aro4  CPA2 - cpa2p  GLN1 - glutamate--ammonia ligase  ARG1 - argininosuccinate synthase  HIS4 - trifunctional histidinol dehydrogenase/phosphoribosyl-amp cyclohydrolase/phosphoribosyl-atp diphosphatase  CIT1 - citrate (si)-synthase cit1  ILV2 - acetolactate synthase catalytic subunit  PRO2 - glutamate-5-semialdehyde dehydrogenase  ARO1 - pentafunctional protein aro1p  CAR2 - ornithine-oxo-acid transaminase  HXK1 - hexokinase 1  ILV6 - acetolactate synthase regulatory subunit  SHM2 - glycine hydroxymethyltransferase shm2  ARG5,6 - bifunctional acetylglutamate kinase/n-acetyl-gamma-glutamyl-phosphate reductase  ACB1 - long-chain fatty acid transporter acb1  ARG7 - glutamate n-acetyltransferase  DLD3 - dld3p  ARO8 - bifunctional 2-aminoadipate transaminase/aromatic-amino-acid:2-oxoglutarate transaminase  ASN1 - asparagine synthase (glutamine-hydrolyzing) 1  HIS1 - atp phosphoribosyltransferase  HOM2 - aspartate-semialdehyde dehydrogenase  HXK2 - hexokinase 2  HOM3 - aspartate kinase  ARO3 - 3-deoxy-7-phosphoheptulonate synthase aro3  HEM4 - uroporphyrinogen-iii synthase hem4  IDH2 - isocitrate dehydrogenase (nad(+)) idh2  EHT1 - eht1p  ARO9 - aromatic-amino-acid:2-oxoglutarate transaminase  ALD4 - aldehyde dehydrogenase (nadp(+)) ald4  GLT1 - glutamate synthase (nadh)  TRP2 - anthranilate synthase trp2  BNA5 - kynureninase  ALD2 - aldehyde dehydrogenase (nad(+)) ald2  ILV1 - threonine ammonia-lyase ilv1  OLE1 - stearoyl-coa 9-desaturase  ALD5 - aldehyde dehydrogenase (nad(p)(+)) ald5 |
| GO:0008652 | cellular amino acid biosynthetic process | 1.04E-9 | 5.37E-7 | 1.98 (2298,94,766,62) | [+] Show genes  IDH1 - isocitrate dehydrogenase (nad(+)) idh1  TRP5 - tryptophan synthase trp5  ARO2 - bifunctional chorismate synthase/riboflavin reductase [nad(p)h] aro2  LYS2 - l-aminoadipate-semialdehyde dehydrogenase  ILV3 - ilv3p  MET5 - met5p  TRP4 - anthranilate phosphoribosyltransferase  HIS3 - imidazoleglycerol-phosphate dehydratase his3  ILV5 - ketol-acid reductoisomerase  ASN2 - asparagine synthase (glutamine-hydrolyzing) 2  LEU1 - 3-isopropylmalate dehydratase leu1  HIS2 - histidinol-phosphatase  HIS7 - imidazoleglycerol-phosphate synthase  BAT2 - bat2p  ARO4 - 3-deoxy-7-phosphoheptulonate synthase aro4  MET17 - bifunctional cysteine synthase/o-acetylhomoserine aminocarboxypropyltransferase met17  CPA2 - cpa2p  PRO3 - pyrroline-5-carboxylate reductase  GLN1 - glutamate--ammonia ligase  ARG1 - argininosuccinate synthase  HIS4 - trifunctional histidinol dehydrogenase/phosphoribosyl-amp cyclohydrolase/phosphoribosyl-atp diphosphatase  MET22 - met22p  CIT1 - citrate (si)-synthase cit1  PRO2 - glutamate-5-semialdehyde dehydrogenase  ILV2 - acetolactate synthase catalytic subunit  SAM4 - sam4p  SER1 - o-phospho-l-serine:2-oxoglutarate transaminase  LEU2 - 3-isopropylmalate dehydrogenase  ARO1 - pentafunctional protein aro1p  CYS4 - cystathionine beta-synthase cys4  CPA1 - carbamoyl-phosphate synthase (glutamine-hydrolyzing) cpa1  LYS1 - saccharopine dehydrogenase (nad+, l-lysine-forming)  CAR2 - ornithine-oxo-acid transaminase  ILV6 - acetolactate synthase regulatory subunit  SHM2 - glycine hydroxymethyltransferase shm2  LYS21 - homocitrate synthase lys21  ARG5,6 - bifunctional acetylglutamate kinase/n-acetyl-gamma-glutamyl-phosphate reductase  ARG7 - glutamate n-acetyltransferase  MHT1 - mht1p  ARO8 - bifunctional 2-aminoadipate transaminase/aromatic-amino-acid:2-oxoglutarate transaminase  PRO1 - glutamate 5-kinase  ABZ1 - 4-amino-4-deoxychorismate synthase  ASN1 - asparagine synthase (glutamine-hydrolyzing) 1  HIS1 - atp phosphoribosyltransferase  TYR1 - pprephenate dehydrogenase (nadp(+))  HOM2 - aspartate-semialdehyde dehydrogenase  HOM3 - aspartate kinase  ARO3 - 3-deoxy-7-phosphoheptulonate synthase aro3  IDH2 - isocitrate dehydrogenase (nad(+)) idh2  LPD1 - dihydrolipoyl dehydrogenase  LEU4 - 2-isopropylmalate synthase leu4  ACO2 - aco2p  HIS5 - histidinol-phosphate transaminase  ARO9 - aromatic-amino-acid:2-oxoglutarate transaminase  MET12 - methylenetetrahydrofolate reductase (nad(p)h) met12  GLT1 - glutamate synthase (nadh)  TRP2 - anthranilate synthase trp2  MET6 - 5-methyltetrahydropteroyltriglutamate-homocysteine s-methyltransferase  ILV1 - threonine ammonia-lyase ilv1  ALD2 - aldehyde dehydrogenase (nad(+)) ald2  MRI1 - s-methyl-5-thioribose-1-phosphate isomerase mri1  LYS9 - saccharopine dehydrogenase (nadp+, l-glutamate-forming) |
| GO:0006520 | cellular amino acid metabolic process | 2.31E-9 | 1.06E-6 | 2.36 (2298,164,320,54) | [+] Show genes  IDH1 - isocitrate dehydrogenase (nad(+)) idh1  TRP5 - tryptophan synthase trp5  ARO2 - bifunctional chorismate synthase/riboflavin reductase [nad(p)h] aro2  ILV3 - ilv3p  MET5 - met5p  PDC1 - indolepyruvate decarboxylase 1  HIS3 - imidazoleglycerol-phosphate dehydratase his3  ILV5 - ketol-acid reductoisomerase  GAD1 - glutamate decarboxylase gad1  ASN2 - asparagine synthase (glutamine-hydrolyzing) 2  YDR341C - arginine--trna ligase  BAT2 - bat2p  ARO4 - 3-deoxy-7-phosphoheptulonate synthase aro4  MET17 - bifunctional cysteine synthase/o-acetylhomoserine aminocarboxypropyltransferase met17  CPA2 - cpa2p  HTS1 - histidine--trna ligase  MAE1 - malate dehydrogenase (oxaloacetate-decarboxylating)  ARG1 - argininosuccinate synthase  GLN1 - glutamate--ammonia ligase  HIS4 - trifunctional histidinol dehydrogenase/phosphoribosyl-amp cyclohydrolase/phosphoribosyl-atp diphosphatase  CIT1 - citrate (si)-synthase cit1  ILV2 - acetolactate synthase catalytic subunit  PRO2 - glutamate-5-semialdehyde dehydrogenase  HFD1 - hfd1p  UGA1 - 4-aminobutyrate transaminase  ARO1 - pentafunctional protein aro1p  DUR1,2 - bifunctional urea carboxylase/allophanate hydrolase  CPA1 - carbamoyl-phosphate synthase (glutamine-hydrolyzing) cpa1  CAR2 - ornithine-oxo-acid transaminase  ILV6 - acetolactate synthase regulatory subunit  SHM2 - glycine hydroxymethyltransferase shm2  ARG5,6 - bifunctional acetylglutamate kinase/n-acetyl-gamma-glutamyl-phosphate reductase  ARG7 - glutamate n-acetyltransferase  ARO8 - bifunctional 2-aminoadipate transaminase/aromatic-amino-acid:2-oxoglutarate transaminase  ASN1 - asparagine synthase (glutamine-hydrolyzing) 1  HIS1 - atp phosphoribosyltransferase  KGD2 - alpha-ketoglutarate dehydrogenase kgd2  HOM2 - aspartate-semialdehyde dehydrogenase  HOM3 - aspartate kinase  ARO3 - 3-deoxy-7-phosphoheptulonate synthase aro3  EHD3 - ehd3p  IDH2 - isocitrate dehydrogenase (nad(+)) idh2  KRS1 - lysine--trna ligase krs1  ADE6 - phosphoribosylformylglycinamidine synthase  YHR020W - proline--trna ligase  ADE4 - amidophosphoribosyltransferase  ARO9 - aromatic-amino-acid:2-oxoglutarate transaminase  GLT1 - glutamate synthase (nadh)  TRP2 - anthranilate synthase trp2  GFA1 - glutamine--fructose-6-phosphate transaminase (isomerizing) gfa1  BNA5 - kynureninase  ALD2 - aldehyde dehydrogenase (nad(+)) ald2  ILV1 - threonine ammonia-lyase ilv1  CDC60 - leucine--trna ligase cdc60 |
| GO:0043648 | dicarboxylic acid metabolic process | 2.44E-9 | 1.01E-6 | 4.46 (2298,45,252,22) | [+] Show genes  IDH1 - isocitrate dehydrogenase (nad(+)) idh1  FUM1 - fumarase fum1  MAE1 - malate dehydrogenase (oxaloacetate-decarboxylating)  MDH3 - malate dehydrogenase mdh3  PCS60 - pcs60p  CIT1 - citrate (si)-synthase cit1  GLT1 - glutamate synthase (nadh)  ARO2 - bifunctional chorismate synthase/riboflavin reductase [nad(p)h] aro2  BNA5 - kynureninase  KGD1 - alpha-ketoglutarate dehydrogenase kgd1  KGD2 - alpha-ketoglutarate dehydrogenase kgd2  LSC1 - succinate--coa ligase (gdp-forming) subunit alpha  ARO1 - pentafunctional protein aro1p  GAD1 - glutamate decarboxylase gad1  HOM2 - aspartate-semialdehyde dehydrogenase  HOM3 - aspartate kinase  ARO3 - 3-deoxy-7-phosphoheptulonate synthase aro3  CAR2 - ornithine-oxo-acid transaminase  IDH2 - isocitrate dehydrogenase (nad(+)) idh2  SHM2 - glycine hydroxymethyltransferase shm2  MDH1 - malate dehydrogenase mdh1  ARO4 - 3-deoxy-7-phosphoheptulonate synthase aro4 |
| GO:1901605 | alpha-amino acid metabolic process | 2.24E-8 | 8.41E-6 | 1.77 (2298,127,766,75) | [+] Show genes  IDH1 - isocitrate dehydrogenase (nad(+)) idh1  TRP5 - tryptophan synthase trp5  LYS2 - l-aminoadipate-semialdehyde dehydrogenase  ILV3 - ilv3p  MET5 - met5p  PDC1 - indolepyruvate decarboxylase 1  SAM2 - methionine adenosyltransferase sam2  TRP4 - anthranilate phosphoribosyltransferase  HIS3 - imidazoleglycerol-phosphate dehydratase his3  BNA1 - bna1p  ILV5 - ketol-acid reductoisomerase  GAD1 - glutamate decarboxylase gad1  ASN2 - asparagine synthase (glutamine-hydrolyzing) 2  LEU1 - 3-isopropylmalate dehydratase leu1  HIS2 - histidinol-phosphatase  HIS7 - imidazoleglycerol-phosphate synthase  BAT2 - bat2p  MET17 - bifunctional cysteine synthase/o-acetylhomoserine aminocarboxypropyltransferase met17  CPA2 - cpa2p  PRO3 - pyrroline-5-carboxylate reductase  ARG1 - argininosuccinate synthase  GLN1 - glutamate--ammonia ligase  HIS4 - trifunctional histidinol dehydrogenase/phosphoribosyl-amp cyclohydrolase/phosphoribosyl-atp diphosphatase  MET22 - met22p  CIT1 - citrate (si)-synthase cit1  PRO2 - glutamate-5-semialdehyde dehydrogenase  ILV2 - acetolactate synthase catalytic subunit  URA8 - ura8p  SAM4 - sam4p  GDH2 - glutamate dehydrogenase (nad(+))  SER1 - o-phospho-l-serine:2-oxoglutarate transaminase  HFD1 - hfd1p  LEU2 - 3-isopropylmalate dehydrogenase  DUR1,2 - bifunctional urea carboxylase/allophanate hydrolase  CYS4 - cystathionine beta-synthase cys4  CPA1 - carbamoyl-phosphate synthase (glutamine-hydrolyzing) cpa1  CAR2 - ornithine-oxo-acid transaminase  LYS1 - saccharopine dehydrogenase (nad+, l-lysine-forming)  ILV6 - acetolactate synthase regulatory subunit  SHM2 - glycine hydroxymethyltransferase shm2  LYS21 - homocitrate synthase lys21  ARG5,6 - bifunctional acetylglutamate kinase/n-acetyl-gamma-glutamyl-phosphate reductase  ARG7 - glutamate n-acetyltransferase  MHT1 - mht1p  YHR112C - putative cystathionine beta-lyase  GCV1 - glycine decarboxylase subunit t  ARO8 - bifunctional 2-aminoadipate transaminase/aromatic-amino-acid:2-oxoglutarate transaminase  PRO1 - glutamate 5-kinase  ABZ1 - 4-amino-4-deoxychorismate synthase  ASN1 - asparagine synthase (glutamine-hydrolyzing) 1  KGD2 - alpha-ketoglutarate dehydrogenase kgd2  HIS1 - atp phosphoribosyltransferase  TYR1 - pprephenate dehydrogenase (nadp(+))  HOM2 - aspartate-semialdehyde dehydrogenase  HOM3 - aspartate kinase  IDH2 - isocitrate dehydrogenase (nad(+)) idh2  LPD1 - dihydrolipoyl dehydrogenase  ADE6 - phosphoribosylformylglycinamidine synthase  LEU4 - 2-isopropylmalate synthase leu4  ADE4 - amidophosphoribosyltransferase  TUM1 - tum1p  ACO2 - aco2p  HIS5 - histidinol-phosphate transaminase  MET12 - methylenetetrahydrofolate reductase (nad(p)h) met12  ARO9 - aromatic-amino-acid:2-oxoglutarate transaminase  GLT1 - glutamate synthase (nadh)  TRP2 - anthranilate synthase trp2  GFA1 - glutamine--fructose-6-phosphate transaminase (isomerizing) gfa1  BNA5 - kynureninase  MET6 - 5-methyltetrahydropteroyltriglutamate-homocysteine s-methyltransferase  ILV1 - threonine ammonia-lyase ilv1  MRI1 - s-methyl-5-thioribose-1-phosphate isomerase mri1  PDC5 - indolepyruvate decarboxylase 5  BNA4 - kynurenine 3-monooxygenase  LYS9 - saccharopine dehydrogenase (nadp+, l-glutamate-forming) |
| GO:0017144 | drug metabolic process | 3.14E-8 | 1.08E-5 | 3.49 (2298,150,123,28) | [+] Show genes  IDH1 - isocitrate dehydrogenase (nad(+)) idh1  ENO2 - phosphopyruvate hydratase eno2  ACH1 - ach1p  ARO8 - bifunctional 2-aminoadipate transaminase/aromatic-amino-acid:2-oxoglutarate transaminase  PDC1 - indolepyruvate decarboxylase 1  ASN1 - asparagine synthase (glutamine-hydrolyzing) 1  LSC1 - succinate--coa ligase (gdp-forming) subunit alpha  ASN2 - asparagine synthase (glutamine-hydrolyzing) 2  ENO1 - phosphopyruvate hydratase eno1  HXK2 - hexokinase 2  GLK1 - glucokinase  ATP3 - atp3p  EMI2 - putative glucokinase  MDH1 - malate dehydrogenase mdh1  PRX1 - prx1p  MDH3 - malate dehydrogenase mdh3  ALD4 - aldehyde dehydrogenase (nadp(+)) ald4  CIT1 - citrate (si)-synthase cit1  GLT1 - glutamate synthase (nadh)  GFA1 - glutamine--fructose-6-phosphate transaminase (isomerizing) gfa1  AMD1 - amp deaminase  DUR1,2 - bifunctional urea carboxylase/allophanate hydrolase  NDE1 - nadh-ubiquinone reductase (h(+)-translocating) nde1  ATP1 - f1f0 atp synthase subunit alpha  ATP2 - atp2p  CAR2 - ornithine-oxo-acid transaminase  THI4 - thi4p  ALD5 - aldehyde dehydrogenase (nad(p)(+)) ald5 |
| GO:1901607 | alpha-amino acid biosynthetic process | 6.73E-8 | 2.13E-5 | 1.91 (2298,88,766,56) | [+] Show genes  IDH1 - isocitrate dehydrogenase (nad(+)) idh1  TRP5 - tryptophan synthase trp5  LYS2 - l-aminoadipate-semialdehyde dehydrogenase  ILV3 - ilv3p  MET5 - met5p  TRP4 - anthranilate phosphoribosyltransferase  HIS3 - imidazoleglycerol-phosphate dehydratase his3  ILV5 - ketol-acid reductoisomerase  ASN2 - asparagine synthase (glutamine-hydrolyzing) 2  LEU1 - 3-isopropylmalate dehydratase leu1  HIS2 - histidinol-phosphatase  HIS7 - imidazoleglycerol-phosphate synthase  BAT2 - bat2p  MET17 - bifunctional cysteine synthase/o-acetylhomoserine aminocarboxypropyltransferase met17  CPA2 - cpa2p  PRO3 - pyrroline-5-carboxylate reductase  GLN1 - glutamate--ammonia ligase  ARG1 - argininosuccinate synthase  HIS4 - trifunctional histidinol dehydrogenase/phosphoribosyl-amp cyclohydrolase/phosphoribosyl-atp diphosphatase  MET22 - met22p  CIT1 - citrate (si)-synthase cit1  ILV2 - acetolactate synthase catalytic subunit  PRO2 - glutamate-5-semialdehyde dehydrogenase  SAM4 - sam4p  SER1 - o-phospho-l-serine:2-oxoglutarate transaminase  LEU2 - 3-isopropylmalate dehydrogenase  CYS4 - cystathionine beta-synthase cys4  CPA1 - carbamoyl-phosphate synthase (glutamine-hydrolyzing) cpa1  CAR2 - ornithine-oxo-acid transaminase  LYS1 - saccharopine dehydrogenase (nad+, l-lysine-forming)  ILV6 - acetolactate synthase regulatory subunit  SHM2 - glycine hydroxymethyltransferase shm2  ARG5,6 - bifunctional acetylglutamate kinase/n-acetyl-gamma-glutamyl-phosphate reductase  LYS21 - homocitrate synthase lys21  ARG7 - glutamate n-acetyltransferase  MHT1 - mht1p  ARO8 - bifunctional 2-aminoadipate transaminase/aromatic-amino-acid:2-oxoglutarate transaminase  PRO1 - glutamate 5-kinase  ASN1 - asparagine synthase (glutamine-hydrolyzing) 1  HIS1 - atp phosphoribosyltransferase  TYR1 - pprephenate dehydrogenase (nadp(+))  HOM2 - aspartate-semialdehyde dehydrogenase  HOM3 - aspartate kinase  IDH2 - isocitrate dehydrogenase (nad(+)) idh2  LPD1 - dihydrolipoyl dehydrogenase  LEU4 - 2-isopropylmalate synthase leu4  ACO2 - aco2p  HIS5 - histidinol-phosphate transaminase  MET12 - methylenetetrahydrofolate reductase (nad(p)h) met12  ARO9 - aromatic-amino-acid:2-oxoglutarate transaminase  GLT1 - glutamate synthase (nadh)  TRP2 - anthranilate synthase trp2  MET6 - 5-methyltetrahydropteroyltriglutamate-homocysteine s-methyltransferase  ILV1 - threonine ammonia-lyase ilv1  MRI1 - s-methyl-5-thioribose-1-phosphate isomerase mri1  LYS9 - saccharopine dehydrogenase (nadp+, l-glutamate-forming) |
| GO:1901566 | organonitrogen compound biosynthetic process | 1.05E-7 | 3.11E-5 | 1.35 (2298,462,769,208) | [+] Show genes  IDH1 - isocitrate dehydrogenase (nad(+)) idh1  HCR1 - hcr1p  PHS1 - phs1p  PYK2 - pyruvate kinase pyk2  GSH1 - gsh1p  ARO2 - bifunctional chorismate synthase/riboflavin reductase [nad(p)h] aro2  RPL9A - ribosomal 60s subunit protein l9a  GLK1 - glucokinase  PRS2 - ribose phosphate diphosphokinase subunit prs2  EMI2 - putative glucokinase  ATP16 - f1f0 atp synthase subunit delta  HIS4 - trifunctional histidinol dehydrogenase/phosphoribosyl-amp cyclohydrolase/phosphoribosyl-atp diphosphatase  CIT1 - citrate (si)-synthase cit1  ILV2 - acetolactate synthase catalytic subunit  PRO2 - glutamate-5-semialdehyde dehydrogenase  MRPL28 - mitochondrial 54s ribosomal protein yml28  RPS31 - ubiquitin-ribosomal 40s subunit protein s31 fusion protein  RIB3 - 3,4-dihydroxy-2-butanone-4-phosphate synthase rib3  LEU2 - 3-isopropylmalate dehydrogenase  RPS2 - ribosomal 40s subunit protein s2  CPA1 - carbamoyl-phosphate synthase (glutamine-hydrolyzing) cpa1  COX15 - cox15p  ILV6 - acetolactate synthase regulatory subunit  ARG5,6 - bifunctional acetylglutamate kinase/n-acetyl-gamma-glutamyl-phosphate reductase  BUD17 - putative pyridoxal kinase bud17  YDL119C - hypothetical protein  ABZ1 - 4-amino-4-deoxychorismate synthase  ASN1 - asparagine synthase (glutamine-hydrolyzing) 1  HIS1 - atp phosphoribosyltransferase  GUK1 - guanylate kinase  ATP7 - f1f0 atp synthase subunit d  HOM3 - aspartate kinase  HEM4 - uroporphyrinogen-iii synthase hem4  RPL31A - ribosomal 60s subunit protein l31a  PNC1 - nicotinamidase  HPT1 - hypoxanthine phosphoribosyltransferase  HEM2 - porphobilinogen synthase hem2  TRP2 - anthranilate synthase trp2  GCD1 - gcd1p  BNA5 - kynureninase  MET6 - 5-methyltetrahydropteroyltriglutamate-homocysteine s-methyltransferase  ALD2 - aldehyde dehydrogenase (nad(+)) ald2  ILV1 - threonine ammonia-lyase ilv1  MRI1 - s-methyl-5-thioribose-1-phosphate isomerase mri1  MRPS12 - putative mitochondrial 37s ribosomal protein mrps12  RPL33B - ribosomal 60s subunit protein l33b  MET7 - tetrahydrofolate synthase  LYS9 - saccharopine dehydrogenase (nadp+, l-glutamate-forming)  CAB4 - putative pantetheine-phosphate adenylyltransferase  NMA1 - nicotinamide-nucleotide adenylyltransferase nma1  NPT1 - nicotinate phosphoribosyltransferase  TIF11 - tif11p  RPL38 - ribosomal 60s subunit protein l38  HIS3 - imidazoleglycerol-phosphate dehydratase his3  ILV5 - ketol-acid reductoisomerase  ENO1 - phosphopyruvate hydratase eno1  FOL2 - gtp cyclohydrolase i  RPL26A - ribosomal 60s subunit protein l26a  RPP0 - ribosomal protein p0  MET17 - bifunctional cysteine synthase/o-acetylhomoserine aminocarboxypropyltransferase met17  TUF1 - tuf1p  SAM4 - sam4p  SER1 - o-phospho-l-serine:2-oxoglutarate transaminase  ATP15 - f1f0 atp synthase subunit epsilon  GLN4 - glutamine--trna ligase  ARO1 - pentafunctional protein aro1p  ADE5,7 - bifunctional aminoimidazole ribotide synthase/glycinamide ribotide synthase  RPL17B - rpl17bp  MRP21 - mitochondrial 37s ribosomal protein mrp21  ISN1 - imp 5'-nucleotidase  RPL7B - ribosomal 60s subunit protein l7b  RPL32 - ribosomal 60s subunit protein l32  RPL22B - ribosomal 60s subunit protein l22b  ARO8 - bifunctional 2-aminoadipate transaminase/aromatic-amino-acid:2-oxoglutarate transaminase  HXK2 - hexokinase 2  ARO3 - 3-deoxy-7-phosphoheptulonate synthase aro3  RPS22A - rps22ap  EHD3 - ehd3p  GON7 - gon7p  LPD1 - dihydrolipoyl dehydrogenase  KRS1 - lysine--trna ligase krs1  IDH2 - isocitrate dehydrogenase (nad(+)) idh2  GPM1 - phosphoglycerate mutase gpm1  DED81 - asparagine--trna ligase ded81  YHR020W - proline--trna ligase  ADE4 - amidophosphoribosyltransferase  ADE2 - phosphoribosylaminoimidazole carboxylase ade2  HEM13 - coproporphyrinogen oxidase  RPS22B - ribosomal 40s subunit protein s22b  ACO2 - aco2p  RPL14B - ribosomal 60s subunit protein l14b  CAB1 - pantothenate kinase  GFA1 - glutamine--fructose-6-phosphate transaminase (isomerizing) gfa1  LCB2 - serine c-palmitoyltransferase lcb2  POS5 - pos5p  AIM17 - aim17p  RPS20 - ribosomal 40s subunit protein s20  PGM3 - phosphoglucomutase pgm3  RPS13 - ribosomal 40s subunit protein s13  KTR1 - ktr1p  ATP1 - f1f0 atp synthase subunit alpha  BNA4 - kynurenine 3-monooxygenase  RPS7A - ribosomal 40s subunit protein s7a  RPL29 - ribosomal 60s subunit protein l29  FUN12 - fun12p  URA5 - orotate phosphoribosyltransferase ura5  ENO2 - phosphopyruvate hydratase eno2  LYS2 - l-aminoadipate-semialdehyde dehydrogenase  MET5 - met5p  GRS1 - glycine--trna ligase  XPT1 - xpt1p  ASN2 - asparagine synthase (glutamine-hydrolyzing) 2  GCD6 - gcd6p  HIS2 - histidinol-phosphatase  BAT2 - bat2p  RSM7 - rsm7p  RPL3 - ribosomal 60s subunit protein l3  CPA2 - cpa2p  URA8 - ura8p  RSM26 - rsm26p  RPL6B - ribosomal 60s subunit protein l6b  RPG1 - rpg1p  CYS4 - cystathionine beta-synthase cys4  YMR31 - mitochondrial 37s ribosomal protein ymr31  ATP2 - atp2p  LYS1 - saccharopine dehydrogenase (nad+, l-lysine-forming)  HXK1 - hexokinase 1  CAR2 - ornithine-oxo-acid transaminase  THI4 - thi4p  RPL24B - ribosomal 60s subunit protein l24b  ATP4 - atp4p  YGR149W - hypothetical protein  RPS1B - ribosomal 40s subunit protein s1b  RPL17A - ribosomal 60s subunit protein l17a  CAB2 - phosphopantothenate--cysteine ligase cab2  MHT1 - mht1p  MRPS35 - mitochondrial 37s ribosomal protein mrps35  FUR1 - uracil phosphoribosyltransferase  SUI1 - sui1p  HOM2 - aspartate-semialdehyde dehydrogenase  DPS1 - aspartate--trna ligase dps1  ATP3 - atp3p  RPL6A - ribosomal 60s subunit protein l6a  HIS5 - histidinol-phosphate transaminase  ARO9 - aromatic-amino-acid:2-oxoglutarate transaminase  ADE12 - adenylosuccinate synthase  CDC60 - leucine--trna ligase cdc60  CAB5 - putative dephospho-coa kinase  RPL5 - ribosomal 60s subunit protein l5  RSM24 - mitochondrial 37s ribosomal protein rsm24  RPL24A - ribosomal 60s subunit protein l24a  RPS3 - ribosomal 40s subunit protein s3  RPL30 - ribosomal 60s subunit protein l30  TRP5 - tryptophan synthase trp5  MRPL19 - mitochondrial 54s ribosomal protein yml19  RIB5 - riboflavin synthase  MRP1 - mitochondrial 37s ribosomal protein mrp1  ILV3 - ilv3p  CHS1 - chitin synthase chs1  TRP4 - anthranilate phosphoribosyltransferase  BNA1 - bna1p  LEU1 - 3-isopropylmalate dehydratase leu1  YDR341C - arginine--trna ligase  VMA9 - vma9p  HIS7 - imidazoleglycerol-phosphate synthase  RPL15A - ribosomal 60s subunit protein l15a  ADE16 - bifunctional phosphoribosylaminoimidazolecarboxamide formyltransferase/imp cyclohydrolase ade16  ARO4 - 3-deoxy-7-phosphoheptulonate synthase aro4  ISC1 - inositol phosphosphingolipid phospholipase  AFG3 - aaa family atpase afg3  HTS1 - histidine--trna ligase  PRO3 - pyrroline-5-carboxylate reductase  ARG1 - argininosuccinate synthase  GLN1 - glutamate--ammonia ligase  MET22 - met22p  RPL10 - ribosomal 60s subunit protein l10  AMD1 - amp deaminase  RPS15 - ribosomal 40s subunit protein s15  YMC1 - ymc1p  NRK1 - ribosylnicotinamide kinase  RPP2A - ribosomal protein p2a  MEF1 - mef1p  FCY1 - cytosine deaminase  SHM2 - glycine hydroxymethyltransferase shm2  RPL22A - ribosomal 60s subunit protein l22a  LYS21 - homocitrate synthase lys21  RPS9B - ribosomal 40s subunit protein s9b  ARG7 - glutamate n-acetyltransferase  RPL26B - ribosomal 60s subunit protein l26b  MHR1 - mhr1p  PRO1 - glutamate 5-kinase  ATP5 - atp5p  PGI1 - glucose-6-phosphate isomerase  YGR054W - hypothetical protein  PCM1 - phosphoacetylglucosamine mutase pcm1  TYR1 - pprephenate dehydrogenase (nadp(+))  LEU4 - 2-isopropylmalate synthase leu4  ADE6 - phosphoribosylformylglycinamidine synthase  MET12 - methylenetetrahydrofolate reductase (nad(p)h) met12  GLT1 - glutamate synthase (nadh)  MRPL35 - mitochondrial 54s ribosomal protein yml35  RPS12 - ribosomal 40s subunit protein s12  SPE4 - spermine synthase  TAE2 - tae2p  LAT1 - dihydrolipoyllysine-residue acetyltransferase  SSB1 - hsp70 family atpase ssb1  RPL9B - ribosomal 60s subunit protein l9b  RPL16B - ribosomal 60s subunit protein l16b |
| GO:0005975 | carbohydrate metabolic process | 1.46E-7 | 4E-5 | 3.46 (2298,125,138,26) | [+] Show genes  PGM1 - phosphoglucomutase pgm1  GPH1 - gph1p  GSY2 - glycogen (starch) synthase gsy2  ENO2 - phosphopyruvate hydratase eno2  CWH41 - cwh41p  HOR2 - glycerol-1-phosphatase hor2  TSL1 - tsl1p  GRE3 - trifunctional aldehyde reductase/xylose reductase/glucose 1-dehydrogenase (nadp(+))  BGL2 - bgl2p  PDC1 - indolepyruvate decarboxylase 1  GPD1 - glycerol-3-phosphate dehydrogenase (nad(+)) gpd1  ENO1 - phosphopyruvate hydratase eno1  HXK2 - hexokinase 2  ZWF1 - glucose-6-phosphate dehydrogenase  GLK1 - glucokinase  EMI2 - putative glucokinase  MDH1 - malate dehydrogenase mdh1  PGM2 - phosphoglucomutase pgm2  MDH3 - malate dehydrogenase mdh3  CIT1 - citrate (si)-synthase cit1  HSP104 - chaperone atpase hsp104  GFA1 - glutamine--fructose-6-phosphate transaminase (isomerizing) gfa1  UGP1 - utp glucose-1-phosphate uridylyltransferase  NDE1 - nadh-ubiquinone reductase (h(+)-translocating) nde1  GLC7 - glc7p  NTH1 - alpha,alpha-trehalase nth1 |
| GO:0072350 | tricarboxylic acid metabolic process | 2.65E-7 | 6.84E-5 | 3.67 (2298,19,527,16) | [+] Show genes  IDH1 - isocitrate dehydrogenase (nad(+)) idh1  FUM1 - fumarase fum1  ACO1 - aconitate hydratase aco1  ACO2 - aco2p  ARG1 - argininosuccinate synthase  MDH3 - malate dehydrogenase mdh3  CIT1 - citrate (si)-synthase cit1  LSC2 - succinate--coa ligase (gdp-forming) subunit beta  KGD1 - alpha-ketoglutarate dehydrogenase kgd1  LSC1 - succinate--coa ligase (gdp-forming) subunit alpha  KGD2 - alpha-ketoglutarate dehydrogenase kgd2  SDH2 - succinate dehydrogenase iron-sulfur protein subunit sdh2  YMR31 - mitochondrial 37s ribosomal protein ymr31  IDH2 - isocitrate dehydrogenase (nad(+)) idh2  MDH1 - malate dehydrogenase mdh1  SDH1 - succinate dehydrogenase flavoprotein subunit sdh1 |
| GO:0055114 | oxidation-reduction process | 3.15E-7 | 7.65E-5 | 1.88 (2298,240,341,67) | [+] Show genes  IDH1 - isocitrate dehydrogenase (nad(+)) idh1  GLC3 - 1,4-alpha-glucan branching enzyme  TRX3 - trx3p  ARO2 - bifunctional chorismate synthase/riboflavin reductase [nad(p)h] aro2  MET5 - met5p  PDC1 - indolepyruvate decarboxylase 1  KGD1 - alpha-ketoglutarate dehydrogenase kgd1  GPD1 - glycerol-3-phosphate dehydrogenase (nad(+)) gpd1  ILV5 - ketol-acid reductoisomerase  GRX3 - grx3p  AYR1 - acylglycerone-phosphate reductase  MDH1 - malate dehydrogenase mdh1  RMD9 - rmd9p  PGM2 - phosphoglucomutase pgm2  MAE1 - malate dehydrogenase (oxaloacetate-decarboxylating)  HIS4 - trifunctional histidinol dehydrogenase/phosphoribosyl-amp cyclohydrolase/phosphoribosyl-atp diphosphatase  QCR2 - ubiquinol--cytochrome-c reductase subunit 2  PRO2 - glutamate-5-semialdehyde dehydrogenase  CBR1 - cbr1p  GLR1 - glutathione-disulfide reductase glr1  COR1 - ubiquinol--cytochrome-c reductase subunit cor1  UGP1 - utp glucose-1-phosphate uridylyltransferase  RSM26 - rsm26p  GDH2 - glutamate dehydrogenase (nad(+))  HFD1 - hfd1p  RIB3 - 3,4-dihydroxy-2-butanone-4-phosphate synthase rib3  OST3 - ost3p  ARO1 - pentafunctional protein aro1p  GUT2 - glycerol-3-phosphate dehydrogenase  ERG24 - delta(14)-sterol reductase  GLC7 - glc7p  PET9 - pet9p  ERG3 - c-5 sterol desaturase  AAP1 - aap1p  ARG5,6 - bifunctional acetylglutamate kinase/n-acetyl-gamma-glutamyl-phosphate reductase  PGM1 - phosphoglucomutase pgm1  GPH1 - gph1p  GSY2 - glycogen (starch) synthase gsy2  DLD3 - dld3p  YDL124W - aldo-keto reductase superfamily protein  GRE3 - trifunctional aldehyde reductase/xylose reductase/glucose 1-dehydrogenase (nadp(+))  YMR315W - hypothetical protein  HOM2 - aspartate-semialdehyde dehydrogenase  ZWF1 - glucose-6-phosphate dehydrogenase  EHD3 - ehd3p  IDH2 - isocitrate dehydrogenase (nad(+)) idh2  MCR1 - mcr1p  SDH1 - succinate dehydrogenase flavoprotein subunit sdh1  PRX1 - prx1p  URE2 - ure2p  HEM13 - coproporphyrinogen oxidase  MDH3 - malate dehydrogenase mdh3  ALD4 - aldehyde dehydrogenase (nadp(+)) ald4  YGL039W - carbonyl reductase (nadph-dependent)  GLT1 - glutamate synthase (nadh)  BDH1 - (r,r)-butanediol dehydrogenase  DLD1 - dld1p  ALD2 - aldehyde dehydrogenase (nad(+)) ald2  AIM17 - aim17p  SDH2 - succinate dehydrogenase iron-sulfur protein subunit sdh2  RIP1 - ubiquinol--cytochrome-c reductase catalytic subunit rip1  DLD2 - dld2p  OLE1 - stearoyl-coa 9-desaturase  NDE1 - nadh-ubiquinone reductase (h(+)-translocating) nde1  ARA1 - d-arabinose 1-dehydrogenase (nad(p)(+)) ara1  ARA2 - d-arabinose 1-dehydrogenase (nad(p)(+)) ara2  ALD5 - aldehyde dehydrogenase (nad(p)(+)) ald5 |
| GO:0005991 | trehalose metabolic process | 3.27E-7 | 7.5E-5 | 9.24 (2298,9,221,8) | [+] Show genes  TPS1 - alpha,alpha-trehalose-phosphate synthase (udp-forming) tps1  PGM1 - phosphoglucomutase pgm1  PGM2 - phosphoglucomutase pgm2  TSL1 - tsl1p  HSP104 - chaperone atpase hsp104  NTH1 - alpha,alpha-trehalase nth1  TPS2 - trehalose-phosphatase tps2  UGP1 - utp glucose-1-phosphate uridylyltransferase |
| GO:1901564 | organonitrogen compound metabolic process | 4.37E-7 | 9.5E-5 | 1.20 (2298,944,743,367) | [+] Show genes  IDH1 - isocitrate dehydrogenase (nad(+)) idh1  NMT1 - nmt1p  PYK2 - pyruvate kinase pyk2  PHS1 - phs1p  HCR1 - hcr1p  RSP5 - nedd4 family e3 ubiquitin-protein ligase  SIT4 - sit4p  GSH1 - gsh1p  ARO2 - bifunctional chorismate synthase/riboflavin reductase [nad(p)h] aro2  RPL9A - ribosomal 60s subunit protein l9a  GPD1 - glycerol-3-phosphate dehydrogenase (nad(+)) gpd1  APA1 - apa1p  EFM1 - efm1p  LAP2 - lap2p  GLK1 - glucokinase  PRS2 - ribose phosphate diphosphokinase subunit prs2  CPR6 - peptidylprolyl isomerase cpr6  EMI2 - putative glucokinase  MXR2 - mxr2p  PBS2 - pbs2p  SIS1 - sis1p  ATP16 - f1f0 atp synthase subunit delta  FAA1 - long-chain fatty acid-coa ligase faa1  HIS4 - trifunctional histidinol dehydrogenase/phosphoribosyl-amp cyclohydrolase/phosphoribosyl-atp diphosphatase  QCR2 - ubiquinol--cytochrome-c reductase subunit 2  CIT1 - citrate (si)-synthase cit1  PRO2 - glutamate-5-semialdehyde dehydrogenase  ILV2 - acetolactate synthase catalytic subunit  MRPL28 - mitochondrial 54s ribosomal protein yml28  RPS31 - ubiquitin-ribosomal 40s subunit protein s31 fusion protein  SNF4 - snf4p  MAS1 - mas1p  HFD1 - hfd1p  RIB3 - 3,4-dihydroxy-2-butanone-4-phosphate synthase rib3  LEU2 - 3-isopropylmalate dehydrogenase  RPS2 - ribosomal 40s subunit protein s2  YTA12 - m-aaa protease subunit yta12  CPA1 - carbamoyl-phosphate synthase (glutamine-hydrolyzing) cpa1  COX15 - cox15p  GLC7 - glc7p  ILV6 - acetolactate synthase regulatory subunit  ARG5,6 - bifunctional acetylglutamate kinase/n-acetyl-gamma-glutamyl-phosphate reductase  MOT2 - ccr4-not core ubiquitin-protein ligase subunit mot2  BUD17 - putative pyridoxal kinase bud17  KAR2 - hsp70 family atpase kar2  GPI17 - gpi17p  CYM1 - cym1p  RPN9 - proteasome regulatory particle lid subunit rpn9  YDL119C - hypothetical protein  HUB1 - hub1p  ABZ1 - 4-amino-4-deoxychorismate synthase  ASN1 - asparagine synthase (glutamine-hydrolyzing) 1  HIS1 - atp phosphoribosyltransferase  GUP1 - gup1p  GUK1 - guanylate kinase  CWH43 - cwh43p  HOM3 - aspartate kinase  ATP7 - f1f0 atp synthase subunit d  LRO1 - phospholipid:diacylglycerol acyltransferase  HEM4 - uroporphyrinogen-iii synthase hem4  UFD4 - putative ubiquitin-protein ligase ufd4  TUM1 - tum1p  OCH1 - och1p  RPL31A - ribosomal 60s subunit protein l31a  PNC1 - nicotinamidase  HPT1 - hypoxanthine phosphoribosyltransferase  MDH3 - malate dehydrogenase mdh3  RPT4 - proteasome regulatory particle base subunit rpt4  PAI3 - pai3p  HEM2 - porphobilinogen synthase hem2  TRP2 - anthranilate synthase trp2  RPT3 - proteasome regulatory particle base subunit rpt3  GCD1 - gcd1p  MET6 - 5-methyltetrahydropteroyltriglutamate-homocysteine s-methyltransferase  RPN8 - proteasome regulatory particle lid subunit rpn8  BNA5 - kynureninase  ILV1 - threonine ammonia-lyase ilv1  UBA2 - e1 ubiquitin-activating protein uba2  ALD2 - aldehyde dehydrogenase (nad(+)) ald2  MRI1 - s-methyl-5-thioribose-1-phosphate isomerase mri1  MRPS12 - putative mitochondrial 37s ribosomal protein mrps12  RPL33B - ribosomal 60s subunit protein l33b  NDE1 - nadh-ubiquinone reductase (h(+)-translocating) nde1  MET7 - tetrahydrofolate synthase  TRM112 - trm112p  BRE5 - bre5p  CAB4 - putative pantetheine-phosphate adenylyltransferase  HSP82 - hsp90 family chaperone hsp82  CHS5 - chs5p  NMA1 - nicotinamide-nucleotide adenylyltransferase nma1  ACH1 - ach1p  NPT1 - nicotinate phosphoribosyltransferase  TIF11 - tif11p  RPL38 - ribosomal 60s subunit protein l38  HIS3 - imidazoleglycerol-phosphate dehydratase his3  FAA4 - long-chain fatty acid-coa ligase faa4  TAL1 - sedoheptulose-7-phosphate:d-glyceraldehyde-3-phosphate transaldolase tal1  ILV5 - ketol-acid reductoisomerase  GAD1 - glutamate decarboxylase gad1  ENO1 - phosphopyruvate hydratase eno1  UBC13 - e2 ubiquitin-conjugating protein ubc13  FOL2 - gtp cyclohydrolase i  RPL26A - ribosomal 60s subunit protein l26a  RPP0 - ribosomal protein p0  MET17 - bifunctional cysteine synthase/o-acetylhomoserine aminocarboxypropyltransferase met17  TUF1 - tuf1p  PEP5 - pep5p  MAE1 - malate dehydrogenase (oxaloacetate-decarboxylating)  COX6 - cytochrome c oxidase subunit vi  SAM4 - sam4p  LSC2 - succinate--coa ligase (gdp-forming) subunit beta  SER1 - o-phospho-l-serine:2-oxoglutarate transaminase  ATP15 - f1f0 atp synthase subunit epsilon  PRE7 - proteasome core particle subunit beta 6  GLN4 - glutamine--trna ligase  ARO1 - pentafunctional protein aro1p  KIN1 - kin1p  ADE5,7 - bifunctional aminoimidazole ribotide synthase/glycinamide ribotide synthase  YCF1 - atp-binding cassette glutathione s-conjugate transporter ycf1  AAP1 - aap1p  ERG12 - mevalonate kinase  SCJ1 - scj1p  RVB2 - ruvb family atp-dependent dna helicase reptin  RFA3 - rfa3p  MRP21 - mitochondrial 37s ribosomal protein mrp21  GCV1 - glycine decarboxylase subunit t  RPL17B - rpl17bp  ISN1 - imp 5'-nucleotidase  MAS2 - mas2p  RPL7B - ribosomal 60s subunit protein l7b  YPK1 - ypk1p  RPL32 - ribosomal 60s subunit protein l32  RPL22B - ribosomal 60s subunit protein l22b  SLT2 - slt2p  ARO8 - bifunctional 2-aminoadipate transaminase/aromatic-amino-acid:2-oxoglutarate transaminase  OCT1 - oct1p  ACT1 - actin  RPT1 - proteasome regulatory particle base subunit rpt1  LSC1 - succinate--coa ligase (gdp-forming) subunit alpha  YMR315W - hypothetical protein  MDJ1 - mdj1p  HXK2 - hexokinase 2  SSA3 - hsp70 family atpase ssa3  ARO3 - 3-deoxy-7-phosphoheptulonate synthase aro3  RPS22A - rps22ap  EHD3 - ehd3p  GON7 - gon7p  IDH2 - isocitrate dehydrogenase (nad(+)) idh2  KRS1 - lysine--trna ligase krs1  LPD1 - dihydrolipoyl dehydrogenase  GPM1 - phosphoglycerate mutase gpm1  YKL151C - nadhx dehydratase  GCG1 - gcg1p  DED81 - asparagine--trna ligase ded81  YHR020W - proline--trna ligase  ADE4 - amidophosphoribosyltransferase  ADE2 - phosphoribosylaminoimidazole carboxylase ade2  HEM13 - coproporphyrinogen oxidase  RPS22B - ribosomal 40s subunit protein s22b  ACO2 - aco2p  SSQ1 - hsp70 family atpase ssq1  CAB1 - pantothenate kinase  RPT5 - proteasome regulatory particle base subunit rpt5  APE1 - ape1p  GFA1 - glutamine--fructose-6-phosphate transaminase (isomerizing) gfa1  BLM10 - blm10p  LCB2 - serine c-palmitoyltransferase lcb2  POS5 - pos5p  AIM17 - aim17p  RPS20 - ribosomal 40s subunit protein s20  RPS13 - ribosomal 40s subunit protein s13  PGM3 - phosphoglucomutase pgm3  KTR1 - ktr1p  ATP1 - f1f0 atp synthase subunit alpha  BNA4 - kynurenine 3-monooxygenase  UBP6 - ubp6p  RPS7A - ribosomal 40s subunit protein s7a  CDC55 - cdc55p  RPL29 - ribosomal 60s subunit protein l29  FUN12 - fun12p  URA5 - orotate phosphoribosyltransferase ura5  ENO2 - phosphopyruvate hydratase eno2  WHI2 - whi2p  LYS2 - l-aminoadipate-semialdehyde dehydrogenase  CUE5 - cue5p  SOL3 - 6-phosphogluconolactonase sol3  MET5 - met5p  GRS1 - glycine--trna ligase  XPT1 - xpt1p  ALD6 - aldehyde dehydrogenase (nadp(+)) ald6  URM1 - ubiquitin-related modifier urm1  ASN2 - asparagine synthase (glutamine-hydrolyzing) 2  GPI16 - gpi16p  GCD6 - gcd6p  CKA2 - cka2p  FES1 - fes1p  CPR3 - peptidylprolyl isomerase cpr3  BAT2 - bat2p  GND1 - phosphogluconate dehydrogenase (decarboxylating) gnd1  RSM7 - rsm7p  BRO1 - bro1p  RPL3 - ribosomal 60s subunit protein l3  FRA1 - fra1p  CPA2 - cpa2p  PBY1 - pby1p  HSP104 - chaperone atpase hsp104  GLR1 - glutathione-disulfide reductase glr1  SSA2 - hsp70 family chaperone ssa2  URA8 - ura8p  RKM4 - rkm4p  RSM26 - rsm26p  FPR4 - peptidylprolyl isomerase fpr4  RPL6B - ribosomal 60s subunit protein l6b  PNG1 - png1p  RPG1 - rpg1p  OST3 - ost3p  SSM4 - e3 ubiquitin-protein ligase ssm4  YMR31 - mitochondrial 37s ribosomal protein ymr31  CYS4 - cystathionine beta-synthase cys4  RPN12 - proteasome regulatory particle lid subunit rpn12  ATP2 - atp2p  LYS1 - saccharopine dehydrogenase (nad+, l-lysine-forming)  HXK1 - hexokinase 1  CAR2 - ornithine-oxo-acid transaminase  THI4 - thi4p  CNA1 - cna1p  RPL24B - ribosomal 60s subunit protein l24b  PTC5 - ptc5p  ATP4 - atp4p  PRP19 - e3 ubiquitin-protein ligase prp19  YGR149W - hypothetical protein  RPS1B - ribosomal 40s subunit protein s1b  RPL17A - ribosomal 60s subunit protein l17a  CAB2 - phosphopantothenate--cysteine ligase cab2  YHR112C - putative cystathionine beta-lyase  CDC37 - cdc37p  MHT1 - mht1p  MRPS35 - mitochondrial 37s ribosomal protein mrps35  CMP2 - cmp2p  UBA1 - e1 ubiquitin-activating protein uba1  KGD2 - alpha-ketoglutarate dehydrogenase kgd2  SSE1 - sse1p  FUR1 - uracil phosphoribosyltransferase  SUI1 - sui1p  HOM2 - aspartate-semialdehyde dehydrogenase  ZWF1 - glucose-6-phosphate dehydrogenase  APE2 - ape2p  DPS1 - aspartate--trna ligase dps1  ATP3 - atp3p  MCD4 - mcd4p  COX17 - cox17p  RPL6A - ribosomal 60s subunit protein l6a  ELP2 - elongator subunit elp2  URE2 - ure2p  PEP4 - pep4p  HIS5 - histidinol-phosphate transaminase  ARO9 - aromatic-amino-acid:2-oxoglutarate transaminase  HMG1 - hydroxymethylglutaryl-coa reductase (nadph) hmg1  RVB1 - ruvb family atp-dependent dna helicase pontin  CRH1 - crh1p  ERG13 - hydroxymethylglutaryl-coa synthase  ADE12 - adenylosuccinate synthase  SGT2 - sgt2p  YIL108W - putative metalloendopeptidase  CDC60 - leucine--trna ligase cdc60  CAB5 - putative dephospho-coa kinase  NDI1 - nadh-ubiquinone reductase (h(+)-translocating) ndi1  SPP1 - spp1p  TRX2 - trx2p  RTS1 - rts1p  RPL5 - ribosomal 60s subunit protein l5  TPD3 - tpd3p  CTK3 - ctk3p  BUL2 - ubiquitin-ubiquitin ligase bul2  KES1 - kes1p  SFM1 - sfm1p  RPL24A - ribosomal 60s subunit protein l24a  RPS3 - ribosomal 40s subunit protein s3  RPL30 - ribosomal 60s subunit protein l30  TRP5 - tryptophan synthase trp5  MRPL19 - mitochondrial 54s ribosomal protein yml19  RIB5 - riboflavin synthase  MRP1 - mitochondrial 37s ribosomal protein mrp1  CWH41 - cwh41p  ILV3 - ilv3p  COX12 - cytochrome c oxidase subunit vib  PDC1 - indolepyruvate decarboxylase 1  SPC1 - spc1p  YOL098C - hypothetical protein  GLO1 - lactoylglutathione lyase glo1  CHS1 - chitin synthase chs1  CKB1 - ckb1p  TRP4 - anthranilate phosphoribosyltransferase  TRX1 - trx1p  BNA1 - bna1p  YDR333C - hypothetical protein  LEU1 - 3-isopropylmalate dehydratase leu1  UBX5 - ubx5p  YDR341C - arginine--trna ligase  HIS7 - imidazoleglycerol-phosphate synthase  VMA9 - vma9p  RPL15A - ribosomal 60s subunit protein l15a  ADE16 - bifunctional phosphoribosylaminoimidazolecarboxamide formyltransferase/imp cyclohydrolase ade16  CPR7 - cpr7p  ARO4 - 3-deoxy-7-phosphoheptulonate synthase aro4  ISC1 - inositol phosphosphingolipid phospholipase  AFG3 - aaa family atpase afg3  HTS1 - histidine--trna ligase  PRO3 - pyrroline-5-carboxylate reductase  GLN1 - glutamate--ammonia ligase  ARG1 - argininosuccinate synthase  YOL057W - dipeptidyl-peptidase iii  MCX1 - mcx1p  YCK2 - yck2p  RPL10 - ribosomal 60s subunit protein l10  GDH2 - glutamate dehydrogenase (nad(+))  UGA1 - 4-aminobutyrate transaminase  AMD1 - amp deaminase  RPS15 - ribosomal 40s subunit protein s15  YMC1 - ymc1p  NRK1 - ribosylnicotinamide kinase  RPP2A - ribosomal protein p2a  DUR1,2 - bifunctional urea carboxylase/allophanate hydrolase  GUT2 - glycerol-3-phosphate dehydrogenase  MEF1 - mef1p  FCY1 - cytosine deaminase  SHM2 - glycine hydroxymethyltransferase shm2  RPL22A - ribosomal 60s subunit protein l22a  LYS21 - homocitrate synthase lys21  RPS9B - ribosomal 40s subunit protein s9b  YPS1 - yps1p  ARG7 - glutamate n-acetyltransferase  RPL26B - ribosomal 60s subunit protein l26b  MHR1 - mhr1p  PRB1 - prb1p  DPL1 - sphinganine-1-phosphate aldolase dpl1  NMA111 - nma111p  PRO1 - glutamate 5-kinase  ATP5 - atp5p  PGI1 - glucose-6-phosphate isomerase  HRT1 - scf ubiquitin ligase complex subunit hrt1  YGR054W - hypothetical protein  PCM1 - phosphoacetylglucosamine mutase pcm1  TYR1 - pprephenate dehydrogenase (nadp(+))  YME1 - yme1p  SSE2 - sse2p  ADE6 - phosphoribosylformylglycinamidine synthase  LEU4 - 2-isopropylmalate synthase leu4  ALD4 - aldehyde dehydrogenase (nadp(+)) ald4  MET12 - methylenetetrahydrofolate reductase (nad(p)h) met12  GLT1 - glutamate synthase (nadh)  RPS12 - ribosomal 40s subunit protein s12  YMR027W - hypothetical protein  SPE4 - spermine synthase  CTR9 - ctr9p  CDC28 - cdc28p  YDJ1 - ydj1p  CPR5 - peptidylprolyl isomerase family protein cpr5  TAE2 - tae2p  PDC5 - indolepyruvate decarboxylase 5  LAT1 - dihydrolipoyllysine-residue acetyltransferase  TFB1 - tfb1p  CCS1 - ccs1p  SSB1 - hsp70 family atpase ssb1  RPL9B - ribosomal 60s subunit protein l9b  PTP1 - ptp1p  RPL16B - ribosomal 60s subunit protein l16b |
| GO:0009150 | purine ribonucleotide metabolic process | 5.59E-7 | 1.15E-4 | 4.05 (2298,86,132,20) | [+] Show genes  CAB2 - phosphopantothenate--cysteine ligase cab2  ADE2 - phosphoribosylaminoimidazole carboxylase ade2  ENO2 - phosphopyruvate hydratase eno2  CIT1 - citrate (si)-synthase cit1  ACH1 - ach1p  PDC1 - indolepyruvate decarboxylase 1  AMD1 - amp deaminase  LSC1 - succinate--coa ligase (gdp-forming) subunit alpha  GUK1 - guanylate kinase  ENO1 - phosphopyruvate hydratase eno1  HXK2 - hexokinase 2  NDE1 - nadh-ubiquinone reductase (h(+)-translocating) nde1  ATP1 - f1f0 atp synthase subunit alpha  ATP2 - atp2p  GLK1 - glucokinase  ADE6 - phosphoribosylformylglycinamidine synthase  ERG12 - mevalonate kinase  ATP3 - atp3p  ADE16 - bifunctional phosphoribosylaminoimidazolecarboxamide formyltransferase/imp cyclohydrolase ade16  EMI2 - putative glucokinase |
| GO:0055086 | nucleobase-containing small molecule metabolic process | 5.8E-7 | 1.14E-4 | 2.86 (2298,183,136,31) | [+] Show genes  CAB2 - phosphopantothenate--cysteine ligase cab2  PGM1 - phosphoglucomutase pgm1  ENO2 - phosphopyruvate hydratase eno2  ACH1 - ach1p  PDC1 - indolepyruvate decarboxylase 1  GPD1 - glycerol-3-phosphate dehydrogenase (nad(+)) gpd1  APA1 - apa1p  LSC1 - succinate--coa ligase (gdp-forming) subunit alpha  GUK1 - guanylate kinase  ENO1 - phosphopyruvate hydratase eno1  HXK2 - hexokinase 2  ZWF1 - glucose-6-phosphate dehydrogenase  GLK1 - glucokinase  ADE6 - phosphoribosylformylglycinamidine synthase  YKL151C - nadhx dehydratase  ATP3 - atp3p  EMI2 - putative glucokinase  ADE16 - bifunctional phosphoribosylaminoimidazolecarboxamide formyltransferase/imp cyclohydrolase ade16  ADE2 - phosphoribosylaminoimidazole carboxylase ade2  PNC1 - nicotinamidase  PGM2 - phosphoglucomutase pgm2  MDH3 - malate dehydrogenase mdh3  ALD4 - aldehyde dehydrogenase (nadp(+)) ald4  CIT1 - citrate (si)-synthase cit1  GFA1 - glutamine--fructose-6-phosphate transaminase (isomerizing) gfa1  UGP1 - utp glucose-1-phosphate uridylyltransferase  AMD1 - amp deaminase  NDE1 - nadh-ubiquinone reductase (h(+)-translocating) nde1  ATP1 - f1f0 atp synthase subunit alpha  ATP2 - atp2p  ERG12 - mevalonate kinase |
| GO:0009064 | glutamine family amino acid metabolic process | 6.88E-7 | 1.29E-4 | 3.55 (2298,39,332,20) | [+] Show genes  IDH1 - isocitrate dehydrogenase (nad(+)) idh1  ARG5,6 - bifunctional acetylglutamate kinase/n-acetyl-gamma-glutamyl-phosphate reductase  ADE4 - amidophosphoribosyltransferase  ARG7 - glutamate n-acetyltransferase  CPA2 - cpa2p  GLN1 - glutamate--ammonia ligase  ARG1 - argininosuccinate synthase  CIT1 - citrate (si)-synthase cit1  GLT1 - glutamate synthase (nadh)  PRO2 - glutamate-5-semialdehyde dehydrogenase  GFA1 - glutamine--fructose-6-phosphate transaminase (isomerizing) gfa1  GDH2 - glutamate dehydrogenase (nad(+))  ASN1 - asparagine synthase (glutamine-hydrolyzing) 1  GAD1 - glutamate decarboxylase gad1  ASN2 - asparagine synthase (glutamine-hydrolyzing) 2  DUR1,2 - bifunctional urea carboxylase/allophanate hydrolase  CPA1 - carbamoyl-phosphate synthase (glutamine-hydrolyzing) cpa1  CAR2 - ornithine-oxo-acid transaminase  IDH2 - isocitrate dehydrogenase (nad(+)) idh2  ADE6 - phosphoribosylformylglycinamidine synthase |
| GO:0044249 | cellular biosynthetic process | 8.36E-7 | 1.5E-4 | 1.25 (2298,726,737,290) | [+] Show genes  TFA2 - tfa2p  IDH1 - isocitrate dehydrogenase (nad(+)) idh1  HCR1 - hcr1p  PYK2 - pyruvate kinase pyk2  PHS1 - phs1p  GSH1 - gsh1p  YKR070W - hypothetical protein  ARO2 - bifunctional chorismate synthase/riboflavin reductase [nad(p)h] aro2  RPL9A - ribosomal 60s subunit protein l9a  ASC1 - asc1p  APA1 - apa1p  GLK1 - glucokinase  PRS2 - ribose phosphate diphosphokinase subunit prs2  EMI2 - putative glucokinase  ATP16 - f1f0 atp synthase subunit delta  PGM2 - phosphoglucomutase pgm2  RPC82 - rpc82p  HIS4 - trifunctional histidinol dehydrogenase/phosphoribosyl-amp cyclohydrolase/phosphoribosyl-atp diphosphatase  CIT1 - citrate (si)-synthase cit1  ILV2 - acetolactate synthase catalytic subunit  PRO2 - glutamate-5-semialdehyde dehydrogenase  RPS31 - ubiquitin-ribosomal 40s subunit protein s31 fusion protein  MRPL28 - mitochondrial 54s ribosomal protein yml28  RPO26 - rpo26p  HFD1 - hfd1p  RIB3 - 3,4-dihydroxy-2-butanone-4-phosphate synthase rib3  LEU2 - 3-isopropylmalate dehydrogenase  RPS2 - ribosomal 40s subunit protein s2  CPA1 - carbamoyl-phosphate synthase (glutamine-hydrolyzing) cpa1  COX15 - cox15p  ILV6 - acetolactate synthase regulatory subunit  ARG5,6 - bifunctional acetylglutamate kinase/n-acetyl-gamma-glutamyl-phosphate reductase  KRE6 - kre6p  BUD17 - putative pyridoxal kinase bud17  KAR2 - hsp70 family atpase kar2  GPI17 - gpi17p  GSY2 - glycogen (starch) synthase gsy2  HOR2 - glycerol-1-phosphatase hor2  THO1 - tho1p  YDL119C - hypothetical protein  ABZ1 - 4-amino-4-deoxychorismate synthase  ASN1 - asparagine synthase (glutamine-hydrolyzing) 1  GUP1 - gup1p  HIS1 - atp phosphoribosyltransferase  GUK1 - guanylate kinase  CWH43 - cwh43p  ATP7 - f1f0 atp synthase subunit d  HOM3 - aspartate kinase  LRO1 - phospholipid:diacylglycerol acyltransferase  HEM4 - uroporphyrinogen-iii synthase hem4  RPB7 - rpb7p  PNC1 - nicotinamidase  RPL31A - ribosomal 60s subunit protein l31a  HPT1 - hypoxanthine phosphoribosyltransferase  HEM2 - porphobilinogen synthase hem2  GCD1 - gcd1p  TRP2 - anthranilate synthase trp2  MET6 - 5-methyltetrahydropteroyltriglutamate-homocysteine s-methyltransferase  BNA5 - kynureninase  ALD2 - aldehyde dehydrogenase (nad(+)) ald2  ILV1 - threonine ammonia-lyase ilv1  MRI1 - s-methyl-5-thioribose-1-phosphate isomerase mri1  MRPS12 - putative mitochondrial 37s ribosomal protein mrps12  RPL33B - ribosomal 60s subunit protein l33b  OLE1 - stearoyl-coa 9-desaturase  MET7 - tetrahydrofolate synthase  ALD5 - aldehyde dehydrogenase (nad(p)(+)) ald5  CAB4 - putative pantetheine-phosphate adenylyltransferase  NMA1 - nicotinamide-nucleotide adenylyltransferase nma1  YDR089W - hypothetical protein  NPT1 - nicotinate phosphoribosyltransferase  TIF11 - tif11p  RPL38 - ribosomal 60s subunit protein l38  YGR169C-A - hypothetical protein  HIS3 - imidazoleglycerol-phosphate dehydratase his3  ILV5 - ketol-acid reductoisomerase  ERG7 - lanosterol synthase erg7  RET1 - ret1p  ENO1 - phosphopyruvate hydratase eno1  FOL2 - gtp cyclohydrolase i  RPL26A - ribosomal 60s subunit protein l26a  COQ1 - trans-hexaprenyltranstransferase  FKS1 - fks1p  RPP0 - ribosomal protein p0  MET17 - bifunctional cysteine synthase/o-acetylhomoserine aminocarboxypropyltransferase met17  TUF1 - tuf1p  DPB4 - dpb4p  SAM4 - sam4p  UGP1 - utp glucose-1-phosphate uridylyltransferase  ATP15 - f1f0 atp synthase subunit epsilon  SER1 - o-phospho-l-serine:2-oxoglutarate transaminase  GLN4 - glutamine--trna ligase  RTF1 - rtf1p  GAS3 - gas3p  ARO1 - pentafunctional protein aro1p  HEK2 - hek2p  ADE5,7 - bifunctional aminoimidazole ribotide synthase/glycinamide ribotide synthase  ERG12 - mevalonate kinase  NCP1 - ncp1p  RFA3 - rfa3p  PGM1 - phosphoglucomutase pgm1  RPL17B - rpl17bp  MRP21 - mitochondrial 37s ribosomal protein mrp21  ISN1 - imp 5'-nucleotidase  RPL7B - ribosomal 60s subunit protein l7b  RPL32 - ribosomal 60s subunit protein l32  RPB2 - rpb2p  RPL22B - ribosomal 60s subunit protein l22b  ARO8 - bifunctional 2-aminoadipate transaminase/aromatic-amino-acid:2-oxoglutarate transaminase  LSC1 - succinate--coa ligase (gdp-forming) subunit alpha  HXK2 - hexokinase 2  RPS22A - rps22ap  ARO3 - 3-deoxy-7-phosphoheptulonate synthase aro3  GON7 - gon7p  EHD3 - ehd3p  IDH2 - isocitrate dehydrogenase (nad(+)) idh2  LPD1 - dihydrolipoyl dehydrogenase  KRS1 - lysine--trna ligase krs1  GNA1 - glucosamine 6-phosphate n-acetyltransferase  GPM1 - phosphoglycerate mutase gpm1  MCR1 - mcr1p  DED81 - asparagine--trna ligase ded81  YHR020W - proline--trna ligase  ADE4 - amidophosphoribosyltransferase  ADE2 - phosphoribosylaminoimidazole carboxylase ade2  RPS22B - ribosomal 40s subunit protein s22b  HEM13 - coproporphyrinogen oxidase  ACO2 - aco2p  CAB1 - pantothenate kinase  RPO31 - rpo31p  GFA1 - glutamine--fructose-6-phosphate transaminase (isomerizing) gfa1  POS5 - pos5p  LCB2 - serine c-palmitoyltransferase lcb2  INP53 - phosphatidylinositol-3-/phosphoinositide 5-phosphatase inp53  TFC7 - tfc7p  AIM17 - aim17p  ABF1 - abf1p  RPS20 - ribosomal 40s subunit protein s20  PGM3 - phosphoglucomutase pgm3  RPS13 - ribosomal 40s subunit protein s13  ATP1 - f1f0 atp synthase subunit alpha  KTR1 - ktr1p  BNA4 - kynurenine 3-monooxygenase  TPS2 - trehalose-phosphatase tps2  RPS7A - ribosomal 40s subunit protein s7a  TPS1 - alpha,alpha-trehalose-phosphate synthase (udp-forming) tps1  RPL29 - ribosomal 60s subunit protein l29  FUN12 - fun12p  ENO2 - phosphopyruvate hydratase eno2  TSL1 - tsl1p  LYS2 - l-aminoadipate-semialdehyde dehydrogenase  MET5 - met5p  GRS1 - glycine--trna ligase  XPT1 - xpt1p  ALD6 - aldehyde dehydrogenase (nadp(+)) ald6  TFC1 - tfc1p  CMD1 - cmd1p  ASN2 - asparagine synthase (glutamine-hydrolyzing) 2  GPI16 - gpi16p  SPT6 - spt6p  GCD6 - gcd6p  BAT2 - bat2p  RSM7 - rsm7p  NHP6B - nhp6bp  RHR2 - glycerol-1-phosphatase rhr2  POL30 - pol30p  CPA2 - cpa2p  RFC3 - replication factor c subunit 3  RPL3 - ribosomal 60s subunit protein l3  URA8 - ura8p  RSM26 - rsm26p  RPL6B - ribosomal 60s subunit protein l6b  RPG1 - rpg1p  CYS4 - cystathionine beta-synthase cys4  YMR31 - mitochondrial 37s ribosomal protein ymr31  PTI1 - pti1p  ERG24 - delta(14)-sterol reductase  ATP2 - atp2p  CAR2 - ornithine-oxo-acid transaminase  HXK1 - hexokinase 1  LYS1 - saccharopine dehydrogenase (nad+, l-lysine-forming)  THI4 - thi4p  RPL24B - ribosomal 60s subunit protein l24b  ATP4 - atp4p  YGR149W - hypothetical protein  RPL17A - ribosomal 60s subunit protein l17a  RPS1B - ribosomal 40s subunit protein s1b  CAB2 - phosphopantothenate--cysteine ligase cab2  MHT1 - mht1p  ORC5 - origin recognition complex subunit 5  MRPS35 - mitochondrial 37s ribosomal protein mrps35  GAS5 - gas5p  RPA49 - rpa49p  FUR1 - uracil phosphoribosyltransferase  SUI1 - sui1p  RNR4 - ribonucleotide-diphosphate reductase subunit rnr4  HOM2 - aspartate-semialdehyde dehydrogenase  DPS1 - aspartate--trna ligase dps1  MCD4 - mcd4p  ATP3 - atp3p  PDR16 - pdr16p  RPL6A - ribosomal 60s subunit protein l6a  HIS5 - histidinol-phosphate transaminase  ARO9 - aromatic-amino-acid:2-oxoglutarate transaminase  HMG1 - hydroxymethylglutaryl-coa reductase (nadph) hmg1  ETR1 - etr1p  ERG13 - hydroxymethylglutaryl-coa synthase  ADE12 - adenylosuccinate synthase  CDC60 - leucine--trna ligase cdc60  CAB5 - putative dephospho-coa kinase  TRX2 - trx2p  RPL5 - ribosomal 60s subunit protein l5  HMO1 - hmo1p  GLC3 - 1,4-alpha-glucan branching enzyme  RPS3 - ribosomal 40s subunit protein s3  RPL24A - ribosomal 60s subunit protein l24a  RPL30 - ribosomal 60s subunit protein l30  MRPL19 - mitochondrial 54s ribosomal protein yml19  TRP5 - tryptophan synthase trp5  RIB5 - riboflavin synthase  MRP1 - mitochondrial 37s ribosomal protein mrp1  CWH41 - cwh41p  ILV3 - ilv3p  ERG6 - sterol 24-c-methyltransferase  TRP4 - anthranilate phosphoribosyltransferase  TRX1 - trx1p  ERG4 - delta(24(24(1)))-sterol reductase  BNA1 - bna1p  AYR1 - acylglycerone-phosphate reductase  YDR333C - hypothetical protein  LEU1 - 3-isopropylmalate dehydratase leu1  YDR341C - arginine--trna ligase  HIS7 - imidazoleglycerol-phosphate synthase  VMA9 - vma9p  RPL15A - ribosomal 60s subunit protein l15a  ERG26 - sterol-4-alpha-carboxylate 3-dehydrogenase (decarboxylating)  ADE16 - bifunctional phosphoribosylaminoimidazolecarboxamide formyltransferase/imp cyclohydrolase ade16  ARO4 - 3-deoxy-7-phosphoheptulonate synthase aro4  ISC1 - inositol phosphosphingolipid phospholipase  AFG3 - aaa family atpase afg3  HTS1 - histidine--trna ligase  PRO3 - pyrroline-5-carboxylate reductase  GLN1 - glutamate--ammonia ligase  ARG1 - argininosuccinate synthase  RPL10 - ribosomal 60s subunit protein l10  AMD1 - amp deaminase  RPS15 - ribosomal 40s subunit protein s15  YMC1 - ymc1p  NRK1 - ribosylnicotinamide kinase  RPP2A - ribosomal protein p2a  NHP6A - nhp6ap  MEF1 - mef1p  ERG3 - c-5 sterol desaturase  ERG28 - erg28p  SHM2 - glycine hydroxymethyltransferase shm2  FCY1 - cytosine deaminase  RPL22A - ribosomal 60s subunit protein l22a  LYS21 - homocitrate synthase lys21  RPS9B - ribosomal 40s subunit protein s9b  ACB1 - long-chain fatty acid transporter acb1  ARG7 - glutamate n-acetyltransferase  DLD3 - dld3p  RPL26B - ribosomal 60s subunit protein l26b  MHR1 - mhr1p  PRO1 - glutamate 5-kinase  ATP5 - atp5p  PGI1 - glucose-6-phosphate isomerase  YGR054W - hypothetical protein  PCM1 - phosphoacetylglucosamine mutase pcm1  TYR1 - pprephenate dehydrogenase (nadp(+))  RPO21 - rpo21p  LEU4 - 2-isopropylmalate synthase leu4  ADE6 - phosphoribosylformylglycinamidine synthase  POL1 - pol1p  EHT1 - eht1p  ERG10 - acetyl-coa c-acetyltransferase  STM1 - stm1p  ALD4 - aldehyde dehydrogenase (nadp(+)) ald4  MET12 - methylenetetrahydrofolate reductase (nad(p)h) met12  GLT1 - glutamate synthase (nadh)  RPS12 - ribosomal 40s subunit protein s12  SPE4 - spermine synthase  CTR9 - ctr9p  TAE2 - tae2p  LAT1 - dihydrolipoyllysine-residue acetyltransferase  ARA2 - d-arabinose 1-dehydrogenase (nad(p)(+)) ara2  TFB1 - tfb1p  SSB1 - hsp70 family atpase ssb1  RPL9B - ribosomal 60s subunit protein l9b  RPL16B - ribosomal 60s subunit protein l16b |
| GO:0006099 | tricarboxylic acid cycle | 8.97E-7 | 1.54E-4 | 3.63 (2298,18,527,15) | [+] Show genes  IDH1 - isocitrate dehydrogenase (nad(+)) idh1  FUM1 - fumarase fum1  ACO1 - aconitate hydratase aco1  ACO2 - aco2p  MDH3 - malate dehydrogenase mdh3  CIT1 - citrate (si)-synthase cit1  LSC2 - succinate--coa ligase (gdp-forming) subunit beta  KGD1 - alpha-ketoglutarate dehydrogenase kgd1  LSC1 - succinate--coa ligase (gdp-forming) subunit alpha  KGD2 - alpha-ketoglutarate dehydrogenase kgd2  SDH2 - succinate dehydrogenase iron-sulfur protein subunit sdh2  YMR31 - mitochondrial 37s ribosomal protein ymr31  IDH2 - isocitrate dehydrogenase (nad(+)) idh2  MDH1 - malate dehydrogenase mdh1  SDH1 - succinate dehydrogenase flavoprotein subunit sdh1 |
| GO:0006101 | citrate metabolic process | 8.97E-7 | 1.48E-4 | 3.63 (2298,18,527,15) | [+] Show genes  IDH1 - isocitrate dehydrogenase (nad(+)) idh1  FUM1 - fumarase fum1  ACO1 - aconitate hydratase aco1  ACO2 - aco2p  MDH3 - malate dehydrogenase mdh3  CIT1 - citrate (si)-synthase cit1  LSC2 - succinate--coa ligase (gdp-forming) subunit beta  KGD1 - alpha-ketoglutarate dehydrogenase kgd1  LSC1 - succinate--coa ligase (gdp-forming) subunit alpha  KGD2 - alpha-ketoglutarate dehydrogenase kgd2  SDH2 - succinate dehydrogenase iron-sulfur protein subunit sdh2  YMR31 - mitochondrial 37s ribosomal protein ymr31  IDH2 - isocitrate dehydrogenase (nad(+)) idh2  MDH1 - malate dehydrogenase mdh1  SDH1 - succinate dehydrogenase flavoprotein subunit sdh1 |
| GO:0016999 | antibiotic metabolic process | 1.53E-6 | 2.43E-4 | 2.75 (2298,38,527,24) | [+] Show genes  IDH1 - isocitrate dehydrogenase (nad(+)) idh1  IAH1 - iah1p  FUM1 - fumarase fum1  ACO1 - aconitate hydratase aco1  ACO2 - aco2p  MDH3 - malate dehydrogenase mdh3  ALD4 - aldehyde dehydrogenase (nadp(+)) ald4  CIT1 - citrate (si)-synthase cit1  ACH1 - ach1p  PDC1 - indolepyruvate decarboxylase 1  LSC2 - succinate--coa ligase (gdp-forming) subunit beta  ALD6 - aldehyde dehydrogenase (nadp(+)) ald6  YJL068C - hypothetical protein  KGD1 - alpha-ketoglutarate dehydrogenase kgd1  KGD2 - alpha-ketoglutarate dehydrogenase kgd2  LSC1 - succinate--coa ligase (gdp-forming) subunit alpha  SDH2 - succinate dehydrogenase iron-sulfur protein subunit sdh2  YMR31 - mitochondrial 37s ribosomal protein ymr31  NDE1 - nadh-ubiquinone reductase (h(+)-translocating) nde1  IDH2 - isocitrate dehydrogenase (nad(+)) idh2  MDH1 - malate dehydrogenase mdh1  ALD5 - aldehyde dehydrogenase (nad(p)(+)) ald5  SDH1 - succinate dehydrogenase flavoprotein subunit sdh1  PRX1 - prx1p |
| GO:0009311 | oligosaccharide metabolic process | 2.27E-6 | 3.46E-4 | 5.85 (2298,13,302,10) | [+] Show genes  TPS1 - alpha,alpha-trehalose-phosphate synthase (udp-forming) tps1  YMR196W - hypothetical protein  PGM1 - phosphoglucomutase pgm1  PGM2 - phosphoglucomutase pgm2  CWH41 - cwh41p  TSL1 - tsl1p  HSP104 - chaperone atpase hsp104  TPS2 - trehalose-phosphatase tps2  NTH1 - alpha,alpha-trehalase nth1  UGP1 - utp glucose-1-phosphate uridylyltransferase |
| GO:1901576 | organic substance biosynthetic process | 2.86E-6 | 4.21E-4 | 1.23 (2298,748,737,295) | [+] Show genes  TFA2 - tfa2p  IDH1 - isocitrate dehydrogenase (nad(+)) idh1  HCR1 - hcr1p  PHS1 - phs1p  PYK2 - pyruvate kinase pyk2  GSH1 - gsh1p  YKR070W - hypothetical protein  ARO2 - bifunctional chorismate synthase/riboflavin reductase [nad(p)h] aro2  RPL9A - ribosomal 60s subunit protein l9a  ASC1 - asc1p  APA1 - apa1p  GLK1 - glucokinase  PRS2 - ribose phosphate diphosphokinase subunit prs2  EMI2 - putative glucokinase  ATP16 - f1f0 atp synthase subunit delta  PGM2 - phosphoglucomutase pgm2  RPC82 - rpc82p  HIS4 - trifunctional histidinol dehydrogenase/phosphoribosyl-amp cyclohydrolase/phosphoribosyl-atp diphosphatase  CIT1 - citrate (si)-synthase cit1  ILV2 - acetolactate synthase catalytic subunit  PRO2 - glutamate-5-semialdehyde dehydrogenase  RPS31 - ubiquitin-ribosomal 40s subunit protein s31 fusion protein  MRPL28 - mitochondrial 54s ribosomal protein yml28  RPO26 - rpo26p  HFD1 - hfd1p  RIB3 - 3,4-dihydroxy-2-butanone-4-phosphate synthase rib3  LEU2 - 3-isopropylmalate dehydrogenase  RPS2 - ribosomal 40s subunit protein s2  CPA1 - carbamoyl-phosphate synthase (glutamine-hydrolyzing) cpa1  COX15 - cox15p  ILV6 - acetolactate synthase regulatory subunit  ARG5,6 - bifunctional acetylglutamate kinase/n-acetyl-gamma-glutamyl-phosphate reductase  KRE6 - kre6p  KAR2 - hsp70 family atpase kar2  BUD17 - putative pyridoxal kinase bud17  GSY2 - glycogen (starch) synthase gsy2  GPI17 - gpi17p  HOR2 - glycerol-1-phosphatase hor2  THO1 - tho1p  YDL119C - hypothetical protein  ABZ1 - 4-amino-4-deoxychorismate synthase  ASN1 - asparagine synthase (glutamine-hydrolyzing) 1  GUP1 - gup1p  HIS1 - atp phosphoribosyltransferase  GUK1 - guanylate kinase  CWH43 - cwh43p  ATP7 - f1f0 atp synthase subunit d  HOM3 - aspartate kinase  LRO1 - phospholipid:diacylglycerol acyltransferase  HEM4 - uroporphyrinogen-iii synthase hem4  RPB7 - rpb7p  PNC1 - nicotinamidase  RPL31A - ribosomal 60s subunit protein l31a  HPT1 - hypoxanthine phosphoribosyltransferase  HEM2 - porphobilinogen synthase hem2  GCD1 - gcd1p  TRP2 - anthranilate synthase trp2  BNA5 - kynureninase  MET6 - 5-methyltetrahydropteroyltriglutamate-homocysteine s-methyltransferase  ALD2 - aldehyde dehydrogenase (nad(+)) ald2  ILV1 - threonine ammonia-lyase ilv1  MRI1 - s-methyl-5-thioribose-1-phosphate isomerase mri1  MRPS12 - putative mitochondrial 37s ribosomal protein mrps12  RPL33B - ribosomal 60s subunit protein l33b  OLE1 - stearoyl-coa 9-desaturase  MET7 - tetrahydrofolate synthase  ALD5 - aldehyde dehydrogenase (nad(p)(+)) ald5  CAB4 - putative pantetheine-phosphate adenylyltransferase  NMA1 - nicotinamide-nucleotide adenylyltransferase nma1  YDR089W - hypothetical protein  NPT1 - nicotinate phosphoribosyltransferase  TIF11 - tif11p  RPL38 - ribosomal 60s subunit protein l38  YGR169C-A - hypothetical protein  HIS3 - imidazoleglycerol-phosphate dehydratase his3  ILV5 - ketol-acid reductoisomerase  ERG7 - lanosterol synthase erg7  ENO1 - phosphopyruvate hydratase eno1  RET1 - ret1p  FOL2 - gtp cyclohydrolase i  RPL26A - ribosomal 60s subunit protein l26a  COQ1 - trans-hexaprenyltranstransferase  FKS1 - fks1p  RPP0 - ribosomal protein p0  MET17 - bifunctional cysteine synthase/o-acetylhomoserine aminocarboxypropyltransferase met17  TUF1 - tuf1p  DPB4 - dpb4p  SAM4 - sam4p  UGP1 - utp glucose-1-phosphate uridylyltransferase  SER1 - o-phospho-l-serine:2-oxoglutarate transaminase  ATP15 - f1f0 atp synthase subunit epsilon  GLN4 - glutamine--trna ligase  RTF1 - rtf1p  GAS3 - gas3p  ARO1 - pentafunctional protein aro1p  HEK2 - hek2p  ADE5,7 - bifunctional aminoimidazole ribotide synthase/glycinamide ribotide synthase  ERG12 - mevalonate kinase  NCP1 - ncp1p  RFA3 - rfa3p  PGM1 - phosphoglucomutase pgm1  RPL17B - rpl17bp  MRP21 - mitochondrial 37s ribosomal protein mrp21  ISN1 - imp 5'-nucleotidase  RPL7B - ribosomal 60s subunit protein l7b  RPL32 - ribosomal 60s subunit protein l32  RPB2 - rpb2p  RPL22B - ribosomal 60s subunit protein l22b  ARO8 - bifunctional 2-aminoadipate transaminase/aromatic-amino-acid:2-oxoglutarate transaminase  LSC1 - succinate--coa ligase (gdp-forming) subunit alpha  HXK2 - hexokinase 2  RPS22A - rps22ap  ARO3 - 3-deoxy-7-phosphoheptulonate synthase aro3  GON7 - gon7p  EHD3 - ehd3p  IDH2 - isocitrate dehydrogenase (nad(+)) idh2  KRS1 - lysine--trna ligase krs1  LPD1 - dihydrolipoyl dehydrogenase  GPM1 - phosphoglycerate mutase gpm1  GNA1 - glucosamine 6-phosphate n-acetyltransferase  MCR1 - mcr1p  DED81 - asparagine--trna ligase ded81  YHR020W - proline--trna ligase  ADE4 - amidophosphoribosyltransferase  ADE2 - phosphoribosylaminoimidazole carboxylase ade2  RPS22B - ribosomal 40s subunit protein s22b  HEM13 - coproporphyrinogen oxidase  ACO2 - aco2p  CAB1 - pantothenate kinase  RPO31 - rpo31p  GFA1 - glutamine--fructose-6-phosphate transaminase (isomerizing) gfa1  POS5 - pos5p  LCB2 - serine c-palmitoyltransferase lcb2  INP53 - phosphatidylinositol-3-/phosphoinositide 5-phosphatase inp53  TFC7 - tfc7p  ABF1 - abf1p  AIM17 - aim17p  RPS20 - ribosomal 40s subunit protein s20  PGM3 - phosphoglucomutase pgm3  RPS13 - ribosomal 40s subunit protein s13  ATP1 - f1f0 atp synthase subunit alpha  KTR1 - ktr1p  BNA4 - kynurenine 3-monooxygenase  TPS2 - trehalose-phosphatase tps2  RPS7A - ribosomal 40s subunit protein s7a  TPS1 - alpha,alpha-trehalose-phosphate synthase (udp-forming) tps1  RPL29 - ribosomal 60s subunit protein l29  FUN12 - fun12p  ENO2 - phosphopyruvate hydratase eno2  TSL1 - tsl1p  LYS2 - l-aminoadipate-semialdehyde dehydrogenase  MET5 - met5p  XPT1 - xpt1p  GRS1 - glycine--trna ligase  ALD6 - aldehyde dehydrogenase (nadp(+)) ald6  TFC1 - tfc1p  CMD1 - cmd1p  ASN2 - asparagine synthase (glutamine-hydrolyzing) 2  GPI16 - gpi16p  SPT6 - spt6p  GCD6 - gcd6p  BAT2 - bat2p  RSM7 - rsm7p  NHP6B - nhp6bp  RHR2 - glycerol-1-phosphatase rhr2  POL30 - pol30p  CPA2 - cpa2p  RFC3 - replication factor c subunit 3  RPL3 - ribosomal 60s subunit protein l3  URA8 - ura8p  RSM26 - rsm26p  RPL6B - ribosomal 60s subunit protein l6b  RPG1 - rpg1p  CYS4 - cystathionine beta-synthase cys4  YMR31 - mitochondrial 37s ribosomal protein ymr31  PTI1 - pti1p  ERG24 - delta(14)-sterol reductase  ATP2 - atp2p  CAR2 - ornithine-oxo-acid transaminase  LYS1 - saccharopine dehydrogenase (nad+, l-lysine-forming)  HXK1 - hexokinase 1  THI4 - thi4p  RPL24B - ribosomal 60s subunit protein l24b  YGR149W - hypothetical protein  ATP4 - atp4p  RPL17A - ribosomal 60s subunit protein l17a  RPS1B - ribosomal 40s subunit protein s1b  CAB2 - phosphopantothenate--cysteine ligase cab2  MHT1 - mht1p  ORC5 - origin recognition complex subunit 5  MRPS35 - mitochondrial 37s ribosomal protein mrps35  GAS5 - gas5p  RPA49 - rpa49p  FUR1 - uracil phosphoribosyltransferase  SUI1 - sui1p  RNR4 - ribonucleotide-diphosphate reductase subunit rnr4  HOM2 - aspartate-semialdehyde dehydrogenase  DPS1 - aspartate--trna ligase dps1  ATP3 - atp3p  MCD4 - mcd4p  PDR16 - pdr16p  RPL6A - ribosomal 60s subunit protein l6a  HIS5 - histidinol-phosphate transaminase  ARO9 - aromatic-amino-acid:2-oxoglutarate transaminase  HMG1 - hydroxymethylglutaryl-coa reductase (nadph) hmg1  ETR1 - etr1p  BDH1 - (r,r)-butanediol dehydrogenase  ERG13 - hydroxymethylglutaryl-coa synthase  ADE12 - adenylosuccinate synthase  CDC60 - leucine--trna ligase cdc60  CAB5 - putative dephospho-coa kinase  TRX2 - trx2p  RPL5 - ribosomal 60s subunit protein l5  HMO1 - hmo1p  KES1 - kes1p  GLC3 - 1,4-alpha-glucan branching enzyme  RPS3 - ribosomal 40s subunit protein s3  RPL24A - ribosomal 60s subunit protein l24a  RPL30 - ribosomal 60s subunit protein l30  MRPL19 - mitochondrial 54s ribosomal protein yml19  TRP5 - tryptophan synthase trp5  RIB5 - riboflavin synthase  MRP1 - mitochondrial 37s ribosomal protein mrp1  CWH41 - cwh41p  ILV3 - ilv3p  ERG6 - sterol 24-c-methyltransferase  PDC1 - indolepyruvate decarboxylase 1  CHS1 - chitin synthase chs1  TRX1 - trx1p  TRP4 - anthranilate phosphoribosyltransferase  ERG4 - delta(24(24(1)))-sterol reductase  BNA1 - bna1p  AYR1 - acylglycerone-phosphate reductase  YDR333C - hypothetical protein  LEU1 - 3-isopropylmalate dehydratase leu1  YDR341C - arginine--trna ligase  HIS7 - imidazoleglycerol-phosphate synthase  VMA9 - vma9p  RPL15A - ribosomal 60s subunit protein l15a  ERG26 - sterol-4-alpha-carboxylate 3-dehydrogenase (decarboxylating)  ADE16 - bifunctional phosphoribosylaminoimidazolecarboxamide formyltransferase/imp cyclohydrolase ade16  ARO4 - 3-deoxy-7-phosphoheptulonate synthase aro4  ISC1 - inositol phosphosphingolipid phospholipase  AFG3 - aaa family atpase afg3  HTS1 - histidine--trna ligase  PRO3 - pyrroline-5-carboxylate reductase  ARG1 - argininosuccinate synthase  GLN1 - glutamate--ammonia ligase  RPL10 - ribosomal 60s subunit protein l10  AMD1 - amp deaminase  RPS15 - ribosomal 40s subunit protein s15  YMC1 - ymc1p  NRK1 - ribosylnicotinamide kinase  RPP2A - ribosomal protein p2a  NHP6A - nhp6ap  MEF1 - mef1p  ERG3 - c-5 sterol desaturase  ERG28 - erg28p  FCY1 - cytosine deaminase  SHM2 - glycine hydroxymethyltransferase shm2  RPL22A - ribosomal 60s subunit protein l22a  LYS21 - homocitrate synthase lys21  RPS9B - ribosomal 40s subunit protein s9b  ACB1 - long-chain fatty acid transporter acb1  ARG7 - glutamate n-acetyltransferase  DLD3 - dld3p  RPL26B - ribosomal 60s subunit protein l26b  MHR1 - mhr1p  PRO1 - glutamate 5-kinase  ATP5 - atp5p  PGI1 - glucose-6-phosphate isomerase  YGR054W - hypothetical protein  TYR1 - pprephenate dehydrogenase (nadp(+))  PCM1 - phosphoacetylglucosamine mutase pcm1  RPO21 - rpo21p  LEU4 - 2-isopropylmalate synthase leu4  ADE6 - phosphoribosylformylglycinamidine synthase  POL1 - pol1p  EHT1 - eht1p  ERG10 - acetyl-coa c-acetyltransferase  STM1 - stm1p  ALD4 - aldehyde dehydrogenase (nadp(+)) ald4  MET12 - methylenetetrahydrofolate reductase (nad(p)h) met12  GLT1 - glutamate synthase (nadh)  RPS12 - ribosomal 40s subunit protein s12  SPE4 - spermine synthase  CTR9 - ctr9p  TAE2 - tae2p  PDC5 - indolepyruvate decarboxylase 5  LAT1 - dihydrolipoyllysine-residue acetyltransferase  ARA2 - d-arabinose 1-dehydrogenase (nad(p)(+)) ara2  TFB1 - tfb1p  SSB1 - hsp70 family atpase ssb1  RPL9B - ribosomal 60s subunit protein l9b  RPL16B - ribosomal 60s subunit protein l16b |
| GO:0006163 | purine nucleotide metabolic process | 3.36E-6 | 4.78E-4 | 3.67 (2298,95,132,20) | [+] Show genes  CAB2 - phosphopantothenate--cysteine ligase cab2  ADE2 - phosphoribosylaminoimidazole carboxylase ade2  ENO2 - phosphopyruvate hydratase eno2  CIT1 - citrate (si)-synthase cit1  ACH1 - ach1p  PDC1 - indolepyruvate decarboxylase 1  AMD1 - amp deaminase  LSC1 - succinate--coa ligase (gdp-forming) subunit alpha  GUK1 - guanylate kinase  ENO1 - phosphopyruvate hydratase eno1  HXK2 - hexokinase 2  NDE1 - nadh-ubiquinone reductase (h(+)-translocating) nde1  ATP1 - f1f0 atp synthase subunit alpha  ATP2 - atp2p  GLK1 - glucokinase  ADE6 - phosphoribosylformylglycinamidine synthase  ERG12 - mevalonate kinase  ATP3 - atp3p  ADE16 - bifunctional phosphoribosylaminoimidazolecarboxamide formyltransferase/imp cyclohydrolase ade16  EMI2 - putative glucokinase |
| GO:0006091 | generation of precursor metabolites and energy | 4.12E-6 | 5.67E-4 | 2.60 (2298,89,298,30) | [+] Show genes  GLC3 - 1,4-alpha-glucan branching enzyme  PGM1 - phosphoglucomutase pgm1  GPH1 - gph1p  GSY2 - glycogen (starch) synthase gsy2  ENO2 - phosphopyruvate hydratase eno2  DLD3 - dld3p  PDC1 - indolepyruvate decarboxylase 1  GRX3 - grx3p  ENO1 - phosphopyruvate hydratase eno1  HXK2 - hexokinase 2  GLK1 - glucokinase  EMI2 - putative glucokinase  MDH1 - malate dehydrogenase mdh1  RMD9 - rmd9p  PGM2 - phosphoglucomutase pgm2  QCR2 - ubiquinol--cytochrome-c reductase subunit 2  GLR1 - glutathione-disulfide reductase glr1  COR1 - ubiquinol--cytochrome-c reductase subunit cor1  UGP1 - utp glucose-1-phosphate uridylyltransferase  RIB3 - 3,4-dihydroxy-2-butanone-4-phosphate synthase rib3  SDH2 - succinate dehydrogenase iron-sulfur protein subunit sdh2  RIP1 - ubiquinol--cytochrome-c reductase catalytic subunit rip1  DLD2 - dld2p  OLE1 - stearoyl-coa 9-desaturase  NDE1 - nadh-ubiquinone reductase (h(+)-translocating) nde1  GLC7 - glc7p  ATP2 - atp2p  PET9 - pet9p  HXK1 - hexokinase 1  AAP1 - aap1p |
| GO:0009259 | ribonucleotide metabolic process | 4.74E-6 | 6.3E-4 | 3.59 (2298,97,132,20) | [+] Show genes  CAB2 - phosphopantothenate--cysteine ligase cab2  ADE2 - phosphoribosylaminoimidazole carboxylase ade2  ENO2 - phosphopyruvate hydratase eno2  CIT1 - citrate (si)-synthase cit1  ACH1 - ach1p  PDC1 - indolepyruvate decarboxylase 1  AMD1 - amp deaminase  LSC1 - succinate--coa ligase (gdp-forming) subunit alpha  GUK1 - guanylate kinase  ENO1 - phosphopyruvate hydratase eno1  HXK2 - hexokinase 2  NDE1 - nadh-ubiquinone reductase (h(+)-translocating) nde1  ATP1 - f1f0 atp synthase subunit alpha  ATP2 - atp2p  GLK1 - glucokinase  ADE6 - phosphoribosylformylglycinamidine synthase  ERG12 - mevalonate kinase  EMI2 - putative glucokinase  ATP3 - atp3p  ADE16 - bifunctional phosphoribosylaminoimidazolecarboxamide formyltransferase/imp cyclohydrolase ade16 |
| GO:0005984 | disaccharide metabolic process | 6.25E-6 | 8.06E-4 | 7.56 (2298,11,221,8) | [+] Show genes  TPS1 - alpha,alpha-trehalose-phosphate synthase (udp-forming) tps1  PGM1 - phosphoglucomutase pgm1  PGM2 - phosphoglucomutase pgm2  TSL1 - tsl1p  HSP104 - chaperone atpase hsp104  UGP1 - utp glucose-1-phosphate uridylyltransferase  TPS2 - trehalose-phosphatase tps2  NTH1 - alpha,alpha-trehalase nth1 |
| GO:0009126 | purine nucleoside monophosphate metabolic process | 6.62E-6 | 8.28E-4 | 4.52 (2298,62,123,15) | [+] Show genes  ADE2 - phosphoribosylaminoimidazole carboxylase ade2  ENO2 - phosphopyruvate hydratase eno2  PDC1 - indolepyruvate decarboxylase 1  AMD1 - amp deaminase  ENO1 - phosphopyruvate hydratase eno1  GUK1 - guanylate kinase  HXK2 - hexokinase 2  NDE1 - nadh-ubiquinone reductase (h(+)-translocating) nde1  ATP1 - f1f0 atp synthase subunit alpha  ATP2 - atp2p  GLK1 - glucokinase  ADE6 - phosphoribosylformylglycinamidine synthase  ADE16 - bifunctional phosphoribosylaminoimidazolecarboxamide formyltransferase/imp cyclohydrolase ade16  EMI2 - putative glucokinase  ATP3 - atp3p |
| GO:0009167 | purine ribonucleoside monophosphate metabolic process | 6.62E-6 | 8.03E-4 | 4.52 (2298,62,123,15) | [+] Show genes  ADE2 - phosphoribosylaminoimidazole carboxylase ade2  ENO2 - phosphopyruvate hydratase eno2  PDC1 - indolepyruvate decarboxylase 1  AMD1 - amp deaminase  ENO1 - phosphopyruvate hydratase eno1  GUK1 - guanylate kinase  HXK2 - hexokinase 2  NDE1 - nadh-ubiquinone reductase (h(+)-translocating) nde1  ATP1 - f1f0 atp synthase subunit alpha  ATP2 - atp2p  GLK1 - glucokinase  ADE6 - phosphoribosylformylglycinamidine synthase  EMI2 - putative glucokinase  ADE16 - bifunctional phosphoribosylaminoimidazolecarboxamide formyltransferase/imp cyclohydrolase ade16  ATP3 - atp3p |
| GO:0009117 | nucleotide metabolic process | 7.07E-6 | 8.33E-4 | 2.82 (2298,162,136,27) | [+] Show genes  CAB2 - phosphopantothenate--cysteine ligase cab2  ENO2 - phosphopyruvate hydratase eno2  ACH1 - ach1p  PDC1 - indolepyruvate decarboxylase 1  GPD1 - glycerol-3-phosphate dehydrogenase (nad(+)) gpd1  APA1 - apa1p  LSC1 - succinate--coa ligase (gdp-forming) subunit alpha  GUK1 - guanylate kinase  ENO1 - phosphopyruvate hydratase eno1  HXK2 - hexokinase 2  ZWF1 - glucose-6-phosphate dehydrogenase  GLK1 - glucokinase  ADE6 - phosphoribosylformylglycinamidine synthase  YKL151C - nadhx dehydratase  ATP3 - atp3p  EMI2 - putative glucokinase  ADE16 - bifunctional phosphoribosylaminoimidazolecarboxamide formyltransferase/imp cyclohydrolase ade16  ADE2 - phosphoribosylaminoimidazole carboxylase ade2  PNC1 - nicotinamidase  MDH3 - malate dehydrogenase mdh3  ALD4 - aldehyde dehydrogenase (nadp(+)) ald4  CIT1 - citrate (si)-synthase cit1  AMD1 - amp deaminase  NDE1 - nadh-ubiquinone reductase (h(+)-translocating) nde1  ATP1 - f1f0 atp synthase subunit alpha  ATP2 - atp2p  ERG12 - mevalonate kinase |
| GO:0044262 | cellular carbohydrate metabolic process | 8.01E-6 | 9.18E-4 | 2.81 (2298,71,288,25) | [+] Show genes  TPS1 - alpha,alpha-trehalose-phosphate synthase (udp-forming) tps1  GLC3 - 1,4-alpha-glucan branching enzyme  PGM1 - phosphoglucomutase pgm1  GPH1 - gph1p  GSY2 - glycogen (starch) synthase gsy2  CWH41 - cwh41p  TSL1 - tsl1p  HOR2 - glycerol-1-phosphatase hor2  HXK2 - hexokinase 2  GLK1 - glucokinase  EMI2 - putative glucokinase  PGM2 - phosphoglucomutase pgm2  MDH3 - malate dehydrogenase mdh3  EXG2 - exg2p  HSP104 - chaperone atpase hsp104  GFA1 - glutamine--fructose-6-phosphate transaminase (isomerizing) gfa1  UGP1 - utp glucose-1-phosphate uridylyltransferase  INP53 - phosphatidylinositol-3-/phosphoinositide 5-phosphatase inp53  GAS3 - gas3p  GLC7 - glc7p  HXK1 - hexokinase 1  ARA1 - d-arabinose 1-dehydrogenase (nad(p)(+)) ara1  AAP1 - aap1p  NTH1 - alpha,alpha-trehalase nth1  TPS2 - trehalose-phosphatase tps2 |
| GO:0006753 | nucleoside phosphate metabolic process | 8.83E-6 | 9.85E-4 | 2.78 (2298,164,136,27) | [+] Show genes  CAB2 - phosphopantothenate--cysteine ligase cab2  ENO2 - phosphopyruvate hydratase eno2  ACH1 - ach1p  PDC1 - indolepyruvate decarboxylase 1  GPD1 - glycerol-3-phosphate dehydrogenase (nad(+)) gpd1  APA1 - apa1p  LSC1 - succinate--coa ligase (gdp-forming) subunit alpha  GUK1 - guanylate kinase  ENO1 - phosphopyruvate hydratase eno1  HXK2 - hexokinase 2  ZWF1 - glucose-6-phosphate dehydrogenase  GLK1 - glucokinase  ADE6 - phosphoribosylformylglycinamidine synthase  YKL151C - nadhx dehydratase  ATP3 - atp3p  EMI2 - putative glucokinase  ADE16 - bifunctional phosphoribosylaminoimidazolecarboxamide formyltransferase/imp cyclohydrolase ade16  ADE2 - phosphoribosylaminoimidazole carboxylase ade2  PNC1 - nicotinamidase  MDH3 - malate dehydrogenase mdh3  ALD4 - aldehyde dehydrogenase (nadp(+)) ald4  CIT1 - citrate (si)-synthase cit1  AMD1 - amp deaminase  NDE1 - nadh-ubiquinone reductase (h(+)-translocating) nde1  ATP1 - f1f0 atp synthase subunit alpha  ATP2 - atp2p  ERG12 - mevalonate kinase |
| GO:0019318 | hexose metabolic process | 9.48E-6 | 1.03E-3 | 6.67 (2298,28,123,10) | [+] Show genes  PGM1 - phosphoglucomutase pgm1  ENO1 - phosphopyruvate hydratase eno1  PGM2 - phosphoglucomutase pgm2  HXK2 - hexokinase 2  ZWF1 - glucose-6-phosphate dehydrogenase  NDE1 - nadh-ubiquinone reductase (h(+)-translocating) nde1  ENO2 - phosphopyruvate hydratase eno2  GLK1 - glucokinase  GRE3 - trifunctional aldehyde reductase/xylose reductase/glucose 1-dehydrogenase (nadp(+))  PDC1 - indolepyruvate decarboxylase 1 |
| GO:0009423 | chorismate biosynthetic process | 1.12E-5 | 1.18E-3 | 21.28 (2298,4,108,4) | [+] Show genes  ARO1 - pentafunctional protein aro1p  ARO3 - 3-deoxy-7-phosphoheptulonate synthase aro3  ARO2 - bifunctional chorismate synthase/riboflavin reductase [nad(p)h] aro2  ARO4 - 3-deoxy-7-phosphoheptulonate synthase aro4 |
| GO:0009058 | biosynthetic process | 1.2E-5 | 1.24E-3 | 1.22 (2298,757,737,295) | [+] Show genes  TFA2 - tfa2p  IDH1 - isocitrate dehydrogenase (nad(+)) idh1  HCR1 - hcr1p  PYK2 - pyruvate kinase pyk2  PHS1 - phs1p  GSH1 - gsh1p  YKR070W - hypothetical protein  ARO2 - bifunctional chorismate synthase/riboflavin reductase [nad(p)h] aro2  RPL9A - ribosomal 60s subunit protein l9a  ASC1 - asc1p  APA1 - apa1p  GLK1 - glucokinase  PRS2 - ribose phosphate diphosphokinase subunit prs2  EMI2 - putative glucokinase  ATP16 - f1f0 atp synthase subunit delta  PGM2 - phosphoglucomutase pgm2  RPC82 - rpc82p  HIS4 - trifunctional histidinol dehydrogenase/phosphoribosyl-amp cyclohydrolase/phosphoribosyl-atp diphosphatase  CIT1 - citrate (si)-synthase cit1  ILV2 - acetolactate synthase catalytic subunit  PRO2 - glutamate-5-semialdehyde dehydrogenase  MRPL28 - mitochondrial 54s ribosomal protein yml28  RPS31 - ubiquitin-ribosomal 40s subunit protein s31 fusion protein  RPO26 - rpo26p  HFD1 - hfd1p  RIB3 - 3,4-dihydroxy-2-butanone-4-phosphate synthase rib3  LEU2 - 3-isopropylmalate dehydrogenase  RPS2 - ribosomal 40s subunit protein s2  CPA1 - carbamoyl-phosphate synthase (glutamine-hydrolyzing) cpa1  COX15 - cox15p  ILV6 - acetolactate synthase regulatory subunit  ARG5,6 - bifunctional acetylglutamate kinase/n-acetyl-gamma-glutamyl-phosphate reductase  KRE6 - kre6p  KAR2 - hsp70 family atpase kar2  BUD17 - putative pyridoxal kinase bud17  GSY2 - glycogen (starch) synthase gsy2  GPI17 - gpi17p  HOR2 - glycerol-1-phosphatase hor2  THO1 - tho1p  YDL119C - hypothetical protein  ABZ1 - 4-amino-4-deoxychorismate synthase  ASN1 - asparagine synthase (glutamine-hydrolyzing) 1  GUP1 - gup1p  HIS1 - atp phosphoribosyltransferase  GUK1 - guanylate kinase  CWH43 - cwh43p  ATP7 - f1f0 atp synthase subunit d  HOM3 - aspartate kinase  LRO1 - phospholipid:diacylglycerol acyltransferase  HEM4 - uroporphyrinogen-iii synthase hem4  RPB7 - rpb7p  PNC1 - nicotinamidase  RPL31A - ribosomal 60s subunit protein l31a  HPT1 - hypoxanthine phosphoribosyltransferase  HEM2 - porphobilinogen synthase hem2  GCD1 - gcd1p  TRP2 - anthranilate synthase trp2  BNA5 - kynureninase  MET6 - 5-methyltetrahydropteroyltriglutamate-homocysteine s-methyltransferase  ALD2 - aldehyde dehydrogenase (nad(+)) ald2  ILV1 - threonine ammonia-lyase ilv1  MRI1 - s-methyl-5-thioribose-1-phosphate isomerase mri1  MRPS12 - putative mitochondrial 37s ribosomal protein mrps12  RPL33B - ribosomal 60s subunit protein l33b  OLE1 - stearoyl-coa 9-desaturase  MET7 - tetrahydrofolate synthase  ALD5 - aldehyde dehydrogenase (nad(p)(+)) ald5  CAB4 - putative pantetheine-phosphate adenylyltransferase  NMA1 - nicotinamide-nucleotide adenylyltransferase nma1  YDR089W - hypothetical protein  NPT1 - nicotinate phosphoribosyltransferase  TIF11 - tif11p  RPL38 - ribosomal 60s subunit protein l38  YGR169C-A - hypothetical protein  HIS3 - imidazoleglycerol-phosphate dehydratase his3  ILV5 - ketol-acid reductoisomerase  ERG7 - lanosterol synthase erg7  RET1 - ret1p  ENO1 - phosphopyruvate hydratase eno1  FOL2 - gtp cyclohydrolase i  RPL26A - ribosomal 60s subunit protein l26a  COQ1 - trans-hexaprenyltranstransferase  FKS1 - fks1p  RPP0 - ribosomal protein p0  MET17 - bifunctional cysteine synthase/o-acetylhomoserine aminocarboxypropyltransferase met17  TUF1 - tuf1p  DPB4 - dpb4p  SAM4 - sam4p  UGP1 - utp glucose-1-phosphate uridylyltransferase  SER1 - o-phospho-l-serine:2-oxoglutarate transaminase  ATP15 - f1f0 atp synthase subunit epsilon  GLN4 - glutamine--trna ligase  RTF1 - rtf1p  GAS3 - gas3p  ARO1 - pentafunctional protein aro1p  HEK2 - hek2p  ADE5,7 - bifunctional aminoimidazole ribotide synthase/glycinamide ribotide synthase  ERG12 - mevalonate kinase  NCP1 - ncp1p  RFA3 - rfa3p  PGM1 - phosphoglucomutase pgm1  MRP21 - mitochondrial 37s ribosomal protein mrp21  RPL17B - rpl17bp  ISN1 - imp 5'-nucleotidase  RPL7B - ribosomal 60s subunit protein l7b  RPL32 - ribosomal 60s subunit protein l32  RPB2 - rpb2p  RPL22B - ribosomal 60s subunit protein l22b  ARO8 - bifunctional 2-aminoadipate transaminase/aromatic-amino-acid:2-oxoglutarate transaminase  LSC1 - succinate--coa ligase (gdp-forming) subunit alpha  HXK2 - hexokinase 2  RPS22A - rps22ap  ARO3 - 3-deoxy-7-phosphoheptulonate synthase aro3  GON7 - gon7p  EHD3 - ehd3p  IDH2 - isocitrate dehydrogenase (nad(+)) idh2  LPD1 - dihydrolipoyl dehydrogenase  KRS1 - lysine--trna ligase krs1  GNA1 - glucosamine 6-phosphate n-acetyltransferase  GPM1 - phosphoglycerate mutase gpm1  MCR1 - mcr1p  DED81 - asparagine--trna ligase ded81  YHR020W - proline--trna ligase  ADE4 - amidophosphoribosyltransferase  ADE2 - phosphoribosylaminoimidazole carboxylase ade2  RPS22B - ribosomal 40s subunit protein s22b  HEM13 - coproporphyrinogen oxidase  ACO2 - aco2p  CAB1 - pantothenate kinase  RPO31 - rpo31p  GFA1 - glutamine--fructose-6-phosphate transaminase (isomerizing) gfa1  POS5 - pos5p  LCB2 - serine c-palmitoyltransferase lcb2  INP53 - phosphatidylinositol-3-/phosphoinositide 5-phosphatase inp53  TFC7 - tfc7p  AIM17 - aim17p  ABF1 - abf1p  RPS20 - ribosomal 40s subunit protein s20  RPS13 - ribosomal 40s subunit protein s13  PGM3 - phosphoglucomutase pgm3  KTR1 - ktr1p  ATP1 - f1f0 atp synthase subunit alpha  BNA4 - kynurenine 3-monooxygenase  TPS2 - trehalose-phosphatase tps2  RPS7A - ribosomal 40s subunit protein s7a  TPS1 - alpha,alpha-trehalose-phosphate synthase (udp-forming) tps1  RPL29 - ribosomal 60s subunit protein l29  FUN12 - fun12p  ENO2 - phosphopyruvate hydratase eno2  TSL1 - tsl1p  LYS2 - l-aminoadipate-semialdehyde dehydrogenase  MET5 - met5p  XPT1 - xpt1p  GRS1 - glycine--trna ligase  ALD6 - aldehyde dehydrogenase (nadp(+)) ald6  TFC1 - tfc1p  CMD1 - cmd1p  ASN2 - asparagine synthase (glutamine-hydrolyzing) 2  GPI16 - gpi16p  SPT6 - spt6p  GCD6 - gcd6p  BAT2 - bat2p  RSM7 - rsm7p  NHP6B - nhp6bp  RHR2 - glycerol-1-phosphatase rhr2  POL30 - pol30p  CPA2 - cpa2p  RFC3 - replication factor c subunit 3  RPL3 - ribosomal 60s subunit protein l3  URA8 - ura8p  RSM26 - rsm26p  RPL6B - ribosomal 60s subunit protein l6b  RPG1 - rpg1p  YMR31 - mitochondrial 37s ribosomal protein ymr31  CYS4 - cystathionine beta-synthase cys4  PTI1 - pti1p  ERG24 - delta(14)-sterol reductase  ATP2 - atp2p  CAR2 - ornithine-oxo-acid transaminase  HXK1 - hexokinase 1  LYS1 - saccharopine dehydrogenase (nad+, l-lysine-forming)  THI4 - thi4p  RPL24B - ribosomal 60s subunit protein l24b  YGR149W - hypothetical protein  ATP4 - atp4p  RPL17A - ribosomal 60s subunit protein l17a  RPS1B - ribosomal 40s subunit protein s1b  CAB2 - phosphopantothenate--cysteine ligase cab2  MHT1 - mht1p  ORC5 - origin recognition complex subunit 5  MRPS35 - mitochondrial 37s ribosomal protein mrps35  GAS5 - gas5p  RPA49 - rpa49p  FUR1 - uracil phosphoribosyltransferase  SUI1 - sui1p  RNR4 - ribonucleotide-diphosphate reductase subunit rnr4  HOM2 - aspartate-semialdehyde dehydrogenase  DPS1 - aspartate--trna ligase dps1  ATP3 - atp3p  MCD4 - mcd4p  PDR16 - pdr16p  RPL6A - ribosomal 60s subunit protein l6a  HIS5 - histidinol-phosphate transaminase  ARO9 - aromatic-amino-acid:2-oxoglutarate transaminase  HMG1 - hydroxymethylglutaryl-coa reductase (nadph) hmg1  ETR1 - etr1p  BDH1 - (r,r)-butanediol dehydrogenase  ERG13 - hydroxymethylglutaryl-coa synthase  ADE12 - adenylosuccinate synthase  CDC60 - leucine--trna ligase cdc60  CAB5 - putative dephospho-coa kinase  TRX2 - trx2p  RPL5 - ribosomal 60s subunit protein l5  HMO1 - hmo1p  KES1 - kes1p  RPS3 - ribosomal 40s subunit protein s3  GLC3 - 1,4-alpha-glucan branching enzyme  RPL24A - ribosomal 60s subunit protein l24a  RPL30 - ribosomal 60s subunit protein l30  MRPL19 - mitochondrial 54s ribosomal protein yml19  TRP5 - tryptophan synthase trp5  RIB5 - riboflavin synthase  CWH41 - cwh41p  MRP1 - mitochondrial 37s ribosomal protein mrp1  ILV3 - ilv3p  ERG6 - sterol 24-c-methyltransferase  PDC1 - indolepyruvate decarboxylase 1  CHS1 - chitin synthase chs1  TRX1 - trx1p  TRP4 - anthranilate phosphoribosyltransferase  ERG4 - delta(24(24(1)))-sterol reductase  BNA1 - bna1p  AYR1 - acylglycerone-phosphate reductase  YDR333C - hypothetical protein  LEU1 - 3-isopropylmalate dehydratase leu1  YDR341C - arginine--trna ligase  HIS7 - imidazoleglycerol-phosphate synthase  VMA9 - vma9p  RPL15A - ribosomal 60s subunit protein l15a  ADE16 - bifunctional phosphoribosylaminoimidazolecarboxamide formyltransferase/imp cyclohydrolase ade16  ERG26 - sterol-4-alpha-carboxylate 3-dehydrogenase (decarboxylating)  ARO4 - 3-deoxy-7-phosphoheptulonate synthase aro4  ISC1 - inositol phosphosphingolipid phospholipase  AFG3 - aaa family atpase afg3  HTS1 - histidine--trna ligase  PRO3 - pyrroline-5-carboxylate reductase  ARG1 - argininosuccinate synthase  GLN1 - glutamate--ammonia ligase  RPL10 - ribosomal 60s subunit protein l10  AMD1 - amp deaminase  YMC1 - ymc1p  RPS15 - ribosomal 40s subunit protein s15  NRK1 - ribosylnicotinamide kinase  RPP2A - ribosomal protein p2a  NHP6A - nhp6ap  MEF1 - mef1p  ERG3 - c-5 sterol desaturase  ERG28 - erg28p  FCY1 - cytosine deaminase  SHM2 - glycine hydroxymethyltransferase shm2  RPL22A - ribosomal 60s subunit protein l22a  LYS21 - homocitrate synthase lys21  RPS9B - ribosomal 40s subunit protein s9b  ARG7 - glutamate n-acetyltransferase  ACB1 - long-chain fatty acid transporter acb1  DLD3 - dld3p  RPL26B - ribosomal 60s subunit protein l26b  MHR1 - mhr1p  PRO1 - glutamate 5-kinase  ATP5 - atp5p  PGI1 - glucose-6-phosphate isomerase  YGR054W - hypothetical protein  TYR1 - pprephenate dehydrogenase (nadp(+))  PCM1 - phosphoacetylglucosamine mutase pcm1  RPO21 - rpo21p  LEU4 - 2-isopropylmalate synthase leu4  ADE6 - phosphoribosylformylglycinamidine synthase  POL1 - pol1p  EHT1 - eht1p  ERG10 - acetyl-coa c-acetyltransferase  STM1 - stm1p  ALD4 - aldehyde dehydrogenase (nadp(+)) ald4  MET12 - methylenetetrahydrofolate reductase (nad(p)h) met12  GLT1 - glutamate synthase (nadh)  RPS12 - ribosomal 40s subunit protein s12  SPE4 - spermine synthase  CTR9 - ctr9p  TAE2 - tae2p  PDC5 - indolepyruvate decarboxylase 5  LAT1 - dihydrolipoyllysine-residue acetyltransferase  ARA2 - d-arabinose 1-dehydrogenase (nad(p)(+)) ara2  TFB1 - tfb1p  RPL9B - ribosomal 60s subunit protein l9b  SSB1 - hsp70 family atpase ssb1  RPL16B - ribosomal 60s subunit protein l16b |
| GO:0019693 | ribose phosphate metabolic process | 1.43E-5 | 1.44E-3 | 3.24 (2298,113,132,21) | [+] Show genes  CAB2 - phosphopantothenate--cysteine ligase cab2  ADE2 - phosphoribosylaminoimidazole carboxylase ade2  ENO2 - phosphopyruvate hydratase eno2  CIT1 - citrate (si)-synthase cit1  ACH1 - ach1p  PDC1 - indolepyruvate decarboxylase 1  AMD1 - amp deaminase  LSC1 - succinate--coa ligase (gdp-forming) subunit alpha  GUK1 - guanylate kinase  ENO1 - phosphopyruvate hydratase eno1  HXK2 - hexokinase 2  ZWF1 - glucose-6-phosphate dehydrogenase  NDE1 - nadh-ubiquinone reductase (h(+)-translocating) nde1  ATP1 - f1f0 atp synthase subunit alpha  ATP2 - atp2p  GLK1 - glucokinase  ADE6 - phosphoribosylformylglycinamidine synthase  ERG12 - mevalonate kinase  ADE16 - bifunctional phosphoribosylaminoimidazolecarboxamide formyltransferase/imp cyclohydrolase ade16  EMI2 - putative glucokinase  ATP3 - atp3p |
| GO:0051186 | cofactor metabolic process | 1.43E-5 | 1.41E-3 | 2.60 (2298,170,151,29) | [+] Show genes  CAB2 - phosphopantothenate--cysteine ligase cab2  ENO2 - phosphopyruvate hydratase eno2  ACH1 - ach1p  PDC1 - indolepyruvate decarboxylase 1  YDL119C - hypothetical protein  GLO1 - lactoylglutathione lyase glo1  GPD1 - glycerol-3-phosphate dehydrogenase (nad(+)) gpd1  LSC1 - succinate--coa ligase (gdp-forming) subunit alpha  ENO1 - phosphopyruvate hydratase eno1  HXK2 - hexokinase 2  ZWF1 - glucose-6-phosphate dehydrogenase  GLK1 - glucokinase  YKL151C - nadhx dehydratase  EMI2 - putative glucokinase  PRX1 - prx1p  HEM13 - coproporphyrinogen oxidase  PNC1 - nicotinamidase  MDH3 - malate dehydrogenase mdh3  ALD4 - aldehyde dehydrogenase (nadp(+)) ald4  CIT1 - citrate (si)-synthase cit1  HEM2 - porphobilinogen synthase hem2  GLR1 - glutathione-disulfide reductase glr1  ERG13 - hydroxymethylglutaryl-coa synthase  ALD2 - aldehyde dehydrogenase (nad(+)) ald2  HFD1 - hfd1p  NFS1 - nfs1p  NDE1 - nadh-ubiquinone reductase (h(+)-translocating) nde1  ARA2 - d-arabinose 1-dehydrogenase (nad(p)(+)) ara2  ERG12 - mevalonate kinase |
| GO:0032787 | monocarboxylic acid metabolic process | 1.71E-5 | 1.65E-3 | 3.62 (2298,93,123,18) | [+] Show genes  MDH3 - malate dehydrogenase mdh3  ALD4 - aldehyde dehydrogenase (nadp(+)) ald4  ENO2 - phosphopyruvate hydratase eno2  DLD3 - dld3p  ACH1 - ach1p  DLD1 - dld1p  PDC1 - indolepyruvate decarboxylase 1  ALD2 - aldehyde dehydrogenase (nad(+)) ald2  GLO1 - lactoylglutathione lyase glo1  ASN1 - asparagine synthase (glutamine-hydrolyzing) 1  ASN2 - asparagine synthase (glutamine-hydrolyzing) 2  ENO1 - phosphopyruvate hydratase eno1  HXK2 - hexokinase 2  NDE1 - nadh-ubiquinone reductase (h(+)-translocating) nde1  GLK1 - glucokinase  EMI2 - putative glucokinase  ALD5 - aldehyde dehydrogenase (nad(p)(+)) ald5  EHT1 - eht1p |
| GO:0072521 | purine-containing compound metabolic process | 1.75E-5 | 1.64E-3 | 3.32 (2298,105,132,20) | [+] Show genes  CAB2 - phosphopantothenate--cysteine ligase cab2  ADE2 - phosphoribosylaminoimidazole carboxylase ade2  ENO2 - phosphopyruvate hydratase eno2  CIT1 - citrate (si)-synthase cit1  ACH1 - ach1p  PDC1 - indolepyruvate decarboxylase 1  AMD1 - amp deaminase  LSC1 - succinate--coa ligase (gdp-forming) subunit alpha  GUK1 - guanylate kinase  ENO1 - phosphopyruvate hydratase eno1  HXK2 - hexokinase 2  NDE1 - nadh-ubiquinone reductase (h(+)-translocating) nde1  ATP1 - f1f0 atp synthase subunit alpha  ATP2 - atp2p  GLK1 - glucokinase  ADE6 - phosphoribosylformylglycinamidine synthase  ERG12 - mevalonate kinase  EMI2 - putative glucokinase  ATP3 - atp3p  ADE16 - bifunctional phosphoribosylaminoimidazolecarboxamide formyltransferase/imp cyclohydrolase ade16 |
| GO:0009072 | aromatic amino acid family metabolic process | 1.82E-5 | 1.67E-3 | 2.47 (2298,32,639,22) | [+] Show genes  TRP5 - tryptophan synthase trp5  HIS5 - histidinol-phosphate transaminase  HIS4 - trifunctional histidinol dehydrogenase/phosphoribosyl-amp cyclohydrolase/phosphoribosyl-atp diphosphatase  ARO9 - aromatic-amino-acid:2-oxoglutarate transaminase  ARO2 - bifunctional chorismate synthase/riboflavin reductase [nad(p)h] aro2  TRP2 - anthranilate synthase trp2  ARO8 - bifunctional 2-aminoadipate transaminase/aromatic-amino-acid:2-oxoglutarate transaminase  BNA5 - kynureninase  PDC1 - indolepyruvate decarboxylase 1  HFD1 - hfd1p  ABZ1 - 4-amino-4-deoxychorismate synthase  TRP4 - anthranilate phosphoribosyltransferase  HIS3 - imidazoleglycerol-phosphate dehydratase his3  BNA1 - bna1p  HIS1 - atp phosphoribosyltransferase  ARO1 - pentafunctional protein aro1p  TYR1 - pprephenate dehydrogenase (nadp(+))  PDC5 - indolepyruvate decarboxylase 5  ARO3 - 3-deoxy-7-phosphoheptulonate synthase aro3  BNA4 - kynurenine 3-monooxygenase  HIS7 - imidazoleglycerol-phosphate synthase  ARO4 - 3-deoxy-7-phosphoheptulonate synthase aro4 |
| GO:0005992 | trehalose biosynthetic process | 2.09E-5 | 1.87E-3 | 8.91 (2298,7,221,6) | [+] Show genes  TPS1 - alpha,alpha-trehalose-phosphate synthase (udp-forming) tps1  PGM1 - phosphoglucomutase pgm1  PGM2 - phosphoglucomutase pgm2  TSL1 - tsl1p  UGP1 - utp glucose-1-phosphate uridylyltransferase  TPS2 - trehalose-phosphatase tps2 |
| GO:0046351 | disaccharide biosynthetic process | 2.09E-5 | 1.83E-3 | 8.91 (2298,7,221,6) | [+] Show genes  TPS1 - alpha,alpha-trehalose-phosphate synthase (udp-forming) tps1  PGM1 - phosphoglucomutase pgm1  PGM2 - phosphoglucomutase pgm2  TSL1 - tsl1p  TPS2 - trehalose-phosphatase tps2  UGP1 - utp glucose-1-phosphate uridylyltransferase |
| GO:0009312 | oligosaccharide biosynthetic process | 2.09E-5 | 1.8E-3 | 8.91 (2298,7,221,6) | [+] Show genes  TPS1 - alpha,alpha-trehalose-phosphate synthase (udp-forming) tps1  PGM1 - phosphoglucomutase pgm1  PGM2 - phosphoglucomutase pgm2  TSL1 - tsl1p  UGP1 - utp glucose-1-phosphate uridylyltransferase  TPS2 - trehalose-phosphatase tps2 |
| GO:0015980 | energy derivation by oxidation of organic compounds | 2.11E-5 | 1.77E-3 | 3.12 (2298,44,318,19) | [+] Show genes  PGM1 - phosphoglucomutase pgm1  GLC3 - 1,4-alpha-glucan branching enzyme  RMD9 - rmd9p  GPH1 - gph1p  PGM2 - phosphoglucomutase pgm2  GSY2 - glycogen (starch) synthase gsy2  QCR2 - ubiquinol--cytochrome-c reductase subunit 2  COR1 - ubiquinol--cytochrome-c reductase subunit cor1  PDC1 - indolepyruvate decarboxylase 1  UGP1 - utp glucose-1-phosphate uridylyltransferase  RIB3 - 3,4-dihydroxy-2-butanone-4-phosphate synthase rib3  SDH2 - succinate dehydrogenase iron-sulfur protein subunit sdh2  RIP1 - ubiquinol--cytochrome-c reductase catalytic subunit rip1  NDE1 - nadh-ubiquinone reductase (h(+)-translocating) nde1  GLC7 - glc7p  PET9 - pet9p  AAP1 - aap1p  MDH1 - malate dehydrogenase mdh1  SDH1 - succinate dehydrogenase flavoprotein subunit sdh1 |
| GO:0006006 | glucose metabolic process | 2.98E-5 | 2.46E-3 | 6.73 (2298,25,123,9) | [+] Show genes  PGM1 - phosphoglucomutase pgm1  ENO1 - phosphopyruvate hydratase eno1  PGM2 - phosphoglucomutase pgm2  HXK2 - hexokinase 2  ZWF1 - glucose-6-phosphate dehydrogenase  NDE1 - nadh-ubiquinone reductase (h(+)-translocating) nde1  ENO2 - phosphopyruvate hydratase eno2  GLK1 - glucokinase  PDC1 - indolepyruvate decarboxylase 1 |
| GO:0009084 | glutamine family amino acid biosynthetic process | 3.3E-5 | 2.67E-3 | 4.18 (2298,21,314,12) | [+] Show genes  IDH1 - isocitrate dehydrogenase (nad(+)) idh1  ARG5,6 - bifunctional acetylglutamate kinase/n-acetyl-gamma-glutamyl-phosphate reductase  ARG7 - glutamate n-acetyltransferase  CPA2 - cpa2p  CPA1 - carbamoyl-phosphate synthase (glutamine-hydrolyzing) cpa1  ARG1 - argininosuccinate synthase  GLN1 - glutamate--ammonia ligase  CIT1 - citrate (si)-synthase cit1  CAR2 - ornithine-oxo-acid transaminase  IDH2 - isocitrate dehydrogenase (nad(+)) idh2  PRO2 - glutamate-5-semialdehyde dehydrogenase  GLT1 - glutamate synthase (nadh) |
| GO:0009073 | aromatic amino acid family biosynthetic process | 5.11E-5 | 4.05E-3 | 2.74 (2298,21,639,16) | [+] Show genes  TRP5 - tryptophan synthase trp5  HIS5 - histidinol-phosphate transaminase  ARO9 - aromatic-amino-acid:2-oxoglutarate transaminase  HIS4 - trifunctional histidinol dehydrogenase/phosphoribosyl-amp cyclohydrolase/phosphoribosyl-atp diphosphatase  ARO2 - bifunctional chorismate synthase/riboflavin reductase [nad(p)h] aro2  ARO8 - bifunctional 2-aminoadipate transaminase/aromatic-amino-acid:2-oxoglutarate transaminase  TRP2 - anthranilate synthase trp2  ABZ1 - 4-amino-4-deoxychorismate synthase  TRP4 - anthranilate phosphoribosyltransferase  HIS3 - imidazoleglycerol-phosphate dehydratase his3  HIS1 - atp phosphoribosyltransferase  ARO1 - pentafunctional protein aro1p  TYR1 - pprephenate dehydrogenase (nadp(+))  ARO3 - 3-deoxy-7-phosphoheptulonate synthase aro3  HIS7 - imidazoleglycerol-phosphate synthase  ARO4 - 3-deoxy-7-phosphoheptulonate synthase aro4 |
| GO:0070981 | L-asparagine biosynthetic process | 5.15E-5 | 4.01E-3 | 135.18 (2298,2,17,2) | [+] Show genes  ASN1 - asparagine synthase (glutamine-hydrolyzing) 1  ASN2 - asparagine synthase (glutamine-hydrolyzing) 2 |
| GO:0070982 | L-asparagine metabolic process | 5.15E-5 | 3.94E-3 | 135.18 (2298,2,17,2) | [+] Show genes  ASN1 - asparagine synthase (glutamine-hydrolyzing) 1  ASN2 - asparagine synthase (glutamine-hydrolyzing) 2 |
| GO:0005996 | monosaccharide metabolic process | 5.49E-5 | 4.12E-3 | 5.66 (2298,33,123,10) | [+] Show genes  PGM1 - phosphoglucomutase pgm1  ENO1 - phosphopyruvate hydratase eno1  PGM2 - phosphoglucomutase pgm2  HXK2 - hexokinase 2  ZWF1 - glucose-6-phosphate dehydrogenase  NDE1 - nadh-ubiquinone reductase (h(+)-translocating) nde1  ENO2 - phosphopyruvate hydratase eno2  GLK1 - glucokinase  GRE3 - trifunctional aldehyde reductase/xylose reductase/glucose 1-dehydrogenase (nadp(+))  PDC1 - indolepyruvate decarboxylase 1 |
| GO:0009152 | purine ribonucleotide biosynthetic process | 5.79E-5 | 4.27E-3 | 4.09 (2298,64,123,14) | [+] Show genes  CAB2 - phosphopantothenate--cysteine ligase cab2  ADE2 - phosphoribosylaminoimidazole carboxylase ade2  ENO2 - phosphopyruvate hydratase eno2  AMD1 - amp deaminase  ENO1 - phosphopyruvate hydratase eno1  GUK1 - guanylate kinase  HXK2 - hexokinase 2  ATP1 - f1f0 atp synthase subunit alpha  ATP2 - atp2p  GLK1 - glucokinase  ADE6 - phosphoribosylformylglycinamidine synthase  ADE16 - bifunctional phosphoribosylaminoimidazolecarboxamide formyltransferase/imp cyclohydrolase ade16  EMI2 - putative glucokinase  ATP3 - atp3p |
| GO:0009132 | nucleoside diphosphate metabolic process | 6.05E-5 | 4.38E-3 | 6.23 (2298,27,123,9) | [+] Show genes  LSC1 - succinate--coa ligase (gdp-forming) subunit alpha  ENO1 - phosphopyruvate hydratase eno1  GUK1 - guanylate kinase  HXK2 - hexokinase 2  NDE1 - nadh-ubiquinone reductase (h(+)-translocating) nde1  ENO2 - phosphopyruvate hydratase eno2  GLK1 - glucokinase  PDC1 - indolepyruvate decarboxylase 1  EMI2 - putative glucokinase |
| GO:0046417 | chorismate metabolic process | 7.28E-5 | 5.18E-3 | 17.02 (2298,5,108,4) | [+] Show genes  ARO1 - pentafunctional protein aro1p  ARO3 - 3-deoxy-7-phosphoheptulonate synthase aro3  ARO2 - bifunctional chorismate synthase/riboflavin reductase [nad(p)h] aro2  ARO4 - 3-deoxy-7-phosphoheptulonate synthase aro4 |
| GO:0006011 | UDP-glucose metabolic process | 8.5E-5 | 5.94E-3 | 27.36 (2298,3,84,3) | [+] Show genes  PGM1 - phosphoglucomutase pgm1  PGM2 - phosphoglucomutase pgm2  UGP1 - utp glucose-1-phosphate uridylyltransferase |
| GO:2000431 | regulation of cytokinesis, actomyosin contractile ring assembly | 8.83E-5 | 6.07E-3 | 26.72 (2298,3,86,3) | [+] Show genes  GLC7 - glc7p  CDC28 - cdc28p  RHO1 - rho1p |
| GO:1903499 | regulation of mitotic actomyosin contractile ring assembly | 8.83E-5 | 5.98E-3 | 26.72 (2298,3,86,3) | [+] Show genes  GLC7 - glc7p  CDC28 - cdc28p  RHO1 - rho1p |
| GO:0019637 | organophosphate metabolic process | 9.22E-5 | 6.14E-3 | 2.28 (2298,230,136,31) | [+] Show genes  CAB2 - phosphopantothenate--cysteine ligase cab2  PGM1 - phosphoglucomutase pgm1  ENO2 - phosphopyruvate hydratase eno2  ACH1 - ach1p  PDC1 - indolepyruvate decarboxylase 1  GPD1 - glycerol-3-phosphate dehydrogenase (nad(+)) gpd1  APA1 - apa1p  LSC1 - succinate--coa ligase (gdp-forming) subunit alpha  ENO1 - phosphopyruvate hydratase eno1  GUK1 - guanylate kinase  HXK2 - hexokinase 2  ZWF1 - glucose-6-phosphate dehydrogenase  GLK1 - glucokinase  ADE6 - phosphoribosylformylglycinamidine synthase  YKL151C - nadhx dehydratase  ADE16 - bifunctional phosphoribosylaminoimidazolecarboxamide formyltransferase/imp cyclohydrolase ade16  ATP3 - atp3p  EMI2 - putative glucokinase  GDE1 - gde1p  ADE2 - phosphoribosylaminoimidazole carboxylase ade2  PNC1 - nicotinamidase  PGM2 - phosphoglucomutase pgm2  MDH3 - malate dehydrogenase mdh3  ALD4 - aldehyde dehydrogenase (nadp(+)) ald4  CIT1 - citrate (si)-synthase cit1  GFA1 - glutamine--fructose-6-phosphate transaminase (isomerizing) gfa1  AMD1 - amp deaminase  NDE1 - nadh-ubiquinone reductase (h(+)-translocating) nde1  ATP1 - f1f0 atp synthase subunit alpha  ATP2 - atp2p  ERG12 - mevalonate kinase |
| GO:0009161 | ribonucleoside monophosphate metabolic process | 1.02E-4 | 6.65E-3 | 3.69 (2298,76,123,15) | [+] Show genes  ADE2 - phosphoribosylaminoimidazole carboxylase ade2  ENO2 - phosphopyruvate hydratase eno2  PDC1 - indolepyruvate decarboxylase 1  AMD1 - amp deaminase  ENO1 - phosphopyruvate hydratase eno1  GUK1 - guanylate kinase  HXK2 - hexokinase 2  NDE1 - nadh-ubiquinone reductase (h(+)-translocating) nde1  ATP1 - f1f0 atp synthase subunit alpha  ATP2 - atp2p  GLK1 - glucokinase  ADE6 - phosphoribosylformylglycinamidine synthase  ADE16 - bifunctional phosphoribosylaminoimidazolecarboxamide formyltransferase/imp cyclohydrolase ade16  EMI2 - putative glucokinase  ATP3 - atp3p |
| GO:0044282 | small molecule catabolic process | 1.28E-4 | 8.23E-3 | 1.96 (2298,81,521,36) | [+] Show genes  AIM45 - aim45p  PGM1 - phosphoglucomutase pgm1  ARO8 - bifunctional 2-aminoadipate transaminase/aromatic-amino-acid:2-oxoglutarate transaminase  GRE3 - trifunctional aldehyde reductase/xylose reductase/glucose 1-dehydrogenase (nadp(+))  PDC1 - indolepyruvate decarboxylase 1  ALD6 - aldehyde dehydrogenase (nadp(+)) ald6  GLO1 - lactoylglutathione lyase glo1  BNA1 - bna1p  APA1 - apa1p  KGD2 - alpha-ketoglutarate dehydrogenase kgd2  GUP1 - gup1p  GAD1 - glutamate decarboxylase gad1  EHD3 - ehd3p  BAT2 - bat2p  GND1 - phosphogluconate dehydrogenase (decarboxylating) gnd1  EHT1 - eht1p  ERG10 - acetyl-coa c-acetyltransferase  PGM2 - phosphoglucomutase pgm2  MDH3 - malate dehydrogenase mdh3  ARO9 - aromatic-amino-acid:2-oxoglutarate transaminase  PCS60 - pcs60p  BNA5 - kynureninase  DLD1 - dld1p  ILV1 - threonine ammonia-lyase ilv1  GDH2 - glutamate dehydrogenase (nad(+))  YJL068C - hypothetical protein  UGA1 - 4-aminobutyrate transaminase  INP53 - phosphatidylinositol-3-/phosphoinositide 5-phosphatase inp53  DUR1,2 - bifunctional urea carboxylase/allophanate hydrolase  PGM3 - phosphoglucomutase pgm3  DLD2 - dld2p  GUT2 - glycerol-3-phosphate dehydrogenase  BNA4 - kynurenine 3-monooxygenase  CAR2 - ornithine-oxo-acid transaminase  SHM2 - glycine hydroxymethyltransferase shm2  ALD5 - aldehyde dehydrogenase (nad(p)(+)) ald5 |
| GO:1901135 | carbohydrate derivative metabolic process | 1.3E-4 | 8.28E-3 | 2.48 (2298,196,123,26) | [+] Show genes  CAB2 - phosphopantothenate--cysteine ligase cab2  PGM1 - phosphoglucomutase pgm1  ENO2 - phosphopyruvate hydratase eno2  ACH1 - ach1p  PDC1 - indolepyruvate decarboxylase 1  GPD1 - glycerol-3-phosphate dehydrogenase (nad(+)) gpd1  LSC1 - succinate--coa ligase (gdp-forming) subunit alpha  APA1 - apa1p  ENO1 - phosphopyruvate hydratase eno1  GUK1 - guanylate kinase  HXK2 - hexokinase 2  ZWF1 - glucose-6-phosphate dehydrogenase  GLK1 - glucokinase  ADE6 - phosphoribosylformylglycinamidine synthase  EMI2 - putative glucokinase  ATP3 - atp3p  ADE16 - bifunctional phosphoribosylaminoimidazolecarboxamide formyltransferase/imp cyclohydrolase ade16  ADE2 - phosphoribosylaminoimidazole carboxylase ade2  PGM2 - phosphoglucomutase pgm2  CIT1 - citrate (si)-synthase cit1  GFA1 - glutamine--fructose-6-phosphate transaminase (isomerizing) gfa1  UGP1 - utp glucose-1-phosphate uridylyltransferase  AMD1 - amp deaminase  NDE1 - nadh-ubiquinone reductase (h(+)-translocating) nde1  ATP1 - f1f0 atp synthase subunit alpha  ATP2 - atp2p |
| GO:0009127 | purine nucleoside monophosphate biosynthetic process | 1.44E-4 | 8.99E-3 | 4.31 (2298,52,123,12) | [+] Show genes  ADE2 - phosphoribosylaminoimidazole carboxylase ade2  ENO1 - phosphopyruvate hydratase eno1  HXK2 - hexokinase 2  ENO2 - phosphopyruvate hydratase eno2  ATP1 - f1f0 atp synthase subunit alpha  ATP2 - atp2p  GLK1 - glucokinase  ADE6 - phosphoribosylformylglycinamidine synthase  ATP3 - atp3p  ADE16 - bifunctional phosphoribosylaminoimidazolecarboxamide formyltransferase/imp cyclohydrolase ade16  EMI2 - putative glucokinase  AMD1 - amp deaminase |
| GO:0009168 | purine ribonucleoside monophosphate biosynthetic process | 1.44E-4 | 8.86E-3 | 4.31 (2298,52,123,12) | [+] Show genes  ADE2 - phosphoribosylaminoimidazole carboxylase ade2  ENO1 - phosphopyruvate hydratase eno1  HXK2 - hexokinase 2  ENO2 - phosphopyruvate hydratase eno2  ATP1 - f1f0 atp synthase subunit alpha  ATP2 - atp2p  GLK1 - glucokinase  ADE6 - phosphoribosylformylglycinamidine synthase  ATP3 - atp3p  ADE16 - bifunctional phosphoribosylaminoimidazolecarboxamide formyltransferase/imp cyclohydrolase ade16  EMI2 - putative glucokinase  AMD1 - amp deaminase |
| GO:0006525 | arginine metabolic process | 1.51E-4 | 9.18E-3 | 5.69 (2298,9,314,7) | [+] Show genes  ARG5,6 - bifunctional acetylglutamate kinase/n-acetyl-gamma-glutamyl-phosphate reductase  ARG7 - glutamate n-acetyltransferase  CPA2 - cpa2p  DUR1,2 - bifunctional urea carboxylase/allophanate hydrolase  ARG1 - argininosuccinate synthase  CPA1 - carbamoyl-phosphate synthase (glutamine-hydrolyzing) cpa1  CAR2 - ornithine-oxo-acid transaminase |
| GO:0006012 | galactose metabolic process | 1.52E-4 | 9.08E-3 | 22.98 (2298,3,100,3) | [+] Show genes  PGM1 - phosphoglucomutase pgm1  PGM2 - phosphoglucomutase pgm2  GRE3 - trifunctional aldehyde reductase/xylose reductase/glucose 1-dehydrogenase (nadp(+)) |
| GO:0019388 | galactose catabolic process | 1.52E-4 | 8.95E-3 | 22.98 (2298,3,100,3) | [+] Show genes  PGM1 - phosphoglucomutase pgm1  PGM2 - phosphoglucomutase pgm2  GRE3 - trifunctional aldehyde reductase/xylose reductase/glucose 1-dehydrogenase (nadp(+)) |
| GO:0019627 | urea metabolic process | 1.72E-4 | 1E-2 | 117.85 (2298,3,13,2) | [+] Show genes  DUR1,2 - bifunctional urea carboxylase/allophanate hydrolase  ARG1 - argininosuccinate synthase |
| GO:0034982 | mitochondrial protein processing | 1.99E-4 | 1.14E-2 | 5.23 (2298,6,439,6) | [+] Show genes  AFG3 - aaa family atpase afg3  MAS2 - mas2p  YTA12 - m-aaa protease subunit yta12  QCR2 - ubiquinol--cytochrome-c reductase subunit 2  OCT1 - oct1p  MAS1 - mas1p |
| GO:0006108 | malate metabolic process | 2.04E-4 | 1.15E-2 | 10.64 (2298,4,216,4) | [+] Show genes  FUM1 - fumarase fum1  MDH3 - malate dehydrogenase mdh3  MAE1 - malate dehydrogenase (oxaloacetate-decarboxylating)  MDH1 - malate dehydrogenase mdh1 |
| GO:0006733 | oxidoreduction coenzyme metabolic process | 2.09E-4 | 1.17E-2 | 3.64 (2298,65,136,14) | [+] Show genes  PNC1 - nicotinamidase  MDH3 - malate dehydrogenase mdh3  ENO2 - phosphopyruvate hydratase eno2  ALD4 - aldehyde dehydrogenase (nadp(+)) ald4  PDC1 - indolepyruvate decarboxylase 1  HFD1 - hfd1p  GPD1 - glycerol-3-phosphate dehydrogenase (nad(+)) gpd1  ENO1 - phosphopyruvate hydratase eno1  HXK2 - hexokinase 2  ZWF1 - glucose-6-phosphate dehydrogenase  NDE1 - nadh-ubiquinone reductase (h(+)-translocating) nde1  GLK1 - glucokinase  YKL151C - nadhx dehydratase  EMI2 - putative glucokinase |
| GO:0009097 | isoleucine biosynthetic process | 2.28E-4 | 1.25E-2 | 5.66 (2298,10,284,7) | [+] Show genes  ILV5 - ketol-acid reductoisomerase  HOM2 - aspartate-semialdehyde dehydrogenase  ILV2 - acetolactate synthase catalytic subunit  ILV3 - ilv3p  ILV6 - acetolactate synthase regulatory subunit  BAT2 - bat2p  ILV1 - threonine ammonia-lyase ilv1 |
| GO:0006732 | coenzyme metabolic process | 2.3E-4 | 1.25E-2 | 2.82 (2298,120,136,20) | [+] Show genes  CAB2 - phosphopantothenate--cysteine ligase cab2  PNC1 - nicotinamidase  MDH3 - malate dehydrogenase mdh3  ALD4 - aldehyde dehydrogenase (nadp(+)) ald4  ENO2 - phosphopyruvate hydratase eno2  CIT1 - citrate (si)-synthase cit1  ACH1 - ach1p  PDC1 - indolepyruvate decarboxylase 1  ALD2 - aldehyde dehydrogenase (nad(+)) ald2  HFD1 - hfd1p  GPD1 - glycerol-3-phosphate dehydrogenase (nad(+)) gpd1  LSC1 - succinate--coa ligase (gdp-forming) subunit alpha  ENO1 - phosphopyruvate hydratase eno1  HXK2 - hexokinase 2  ZWF1 - glucose-6-phosphate dehydrogenase  NDE1 - nadh-ubiquinone reductase (h(+)-translocating) nde1  GLK1 - glucokinase  ERG12 - mevalonate kinase  YKL151C - nadhx dehydratase  EMI2 - putative glucokinase |
| GO:0006164 | purine nucleotide biosynthetic process | 2.37E-4 | 1.27E-2 | 3.63 (2298,72,123,14) | [+] Show genes  CAB2 - phosphopantothenate--cysteine ligase cab2  ADE2 - phosphoribosylaminoimidazole carboxylase ade2  ENO2 - phosphopyruvate hydratase eno2  AMD1 - amp deaminase  ENO1 - phosphopyruvate hydratase eno1  GUK1 - guanylate kinase  HXK2 - hexokinase 2  ATP1 - f1f0 atp synthase subunit alpha  ATP2 - atp2p  GLK1 - glucokinase  ADE6 - phosphoribosylformylglycinamidine synthase  EMI2 - putative glucokinase  ADE16 - bifunctional phosphoribosylaminoimidazolecarboxamide formyltransferase/imp cyclohydrolase ade16  ATP3 - atp3p |
| GO:0009185 | ribonucleoside diphosphate metabolic process | 2.68E-4 | 1.42E-2 | 5.98 (2298,25,123,8) | [+] Show genes  GUK1 - guanylate kinase  ENO1 - phosphopyruvate hydratase eno1  HXK2 - hexokinase 2  NDE1 - nadh-ubiquinone reductase (h(+)-translocating) nde1  ENO2 - phosphopyruvate hydratase eno2  GLK1 - glucokinase  PDC1 - indolepyruvate decarboxylase 1  EMI2 - putative glucokinase |
| GO:0009179 | purine ribonucleoside diphosphate metabolic process | 2.68E-4 | 1.4E-2 | 5.98 (2298,25,123,8) | [+] Show genes  ENO1 - phosphopyruvate hydratase eno1  GUK1 - guanylate kinase  HXK2 - hexokinase 2  NDE1 - nadh-ubiquinone reductase (h(+)-translocating) nde1  ENO2 - phosphopyruvate hydratase eno2  GLK1 - glucokinase  PDC1 - indolepyruvate decarboxylase 1  EMI2 - putative glucokinase |
| GO:0009135 | purine nucleoside diphosphate metabolic process | 2.68E-4 | 1.38E-2 | 5.98 (2298,25,123,8) | [+] Show genes  GUK1 - guanylate kinase  ENO1 - phosphopyruvate hydratase eno1  HXK2 - hexokinase 2  NDE1 - nadh-ubiquinone reductase (h(+)-translocating) nde1  ENO2 - phosphopyruvate hydratase eno2  GLK1 - glucokinase  PDC1 - indolepyruvate decarboxylase 1  EMI2 - putative glucokinase |
| GO:0019362 | pyridine nucleotide metabolic process | 2.71E-4 | 1.38E-2 | 2.14 (2298,59,491,27) | [+] Show genes  PYK2 - pyruvate kinase pyk2  ENO2 - phosphopyruvate hydratase eno2  NPT1 - nicotinate phosphoribosyltransferase  PDC1 - indolepyruvate decarboxylase 1  ALD6 - aldehyde dehydrogenase (nadp(+)) ald6  TAL1 - sedoheptulose-7-phosphate:d-glyceraldehyde-3-phosphate transaldolase tal1  GPD1 - glycerol-3-phosphate dehydrogenase (nad(+)) gpd1  BNA1 - bna1p  YMR315W - hypothetical protein  ENO1 - phosphopyruvate hydratase eno1  HXK2 - hexokinase 2  ZWF1 - glucose-6-phosphate dehydrogenase  GLK1 - glucokinase  GPM1 - phosphoglycerate mutase gpm1  YKL151C - nadhx dehydratase  EMI2 - putative glucokinase  GND1 - phosphogluconate dehydrogenase (decarboxylating) gnd1  PNC1 - nicotinamidase  MDH3 - malate dehydrogenase mdh3  ALD4 - aldehyde dehydrogenase (nadp(+)) ald4  BNA5 - kynureninase  NDI1 - nadh-ubiquinone reductase (h(+)-translocating) ndi1  POS5 - pos5p  NDE1 - nadh-ubiquinone reductase (h(+)-translocating) nde1  GUT2 - glycerol-3-phosphate dehydrogenase  BNA4 - kynurenine 3-monooxygenase  HXK1 - hexokinase 1 |
| GO:0072330 | monocarboxylic acid biosynthetic process | 2.76E-4 | 1.39E-2 | 4.08 (2298,55,123,12) | [+] Show genes  ASN1 - asparagine synthase (glutamine-hydrolyzing) 1  ENO1 - phosphopyruvate hydratase eno1  ASN2 - asparagine synthase (glutamine-hydrolyzing) 2  HXK2 - hexokinase 2  ALD4 - aldehyde dehydrogenase (nadp(+)) ald4  DLD3 - dld3p  ENO2 - phosphopyruvate hydratase eno2  GLK1 - glucokinase  ALD2 - aldehyde dehydrogenase (nad(+)) ald2  EMI2 - putative glucokinase  ALD5 - aldehyde dehydrogenase (nad(p)(+)) ald5  EHT1 - eht1p |
| GO:0009123 | nucleoside monophosphate metabolic process | 2.93E-4 | 1.46E-2 | 3.38 (2298,83,123,15) | [+] Show genes  ADE2 - phosphoribosylaminoimidazole carboxylase ade2  ENO2 - phosphopyruvate hydratase eno2  PDC1 - indolepyruvate decarboxylase 1  AMD1 - amp deaminase  ENO1 - phosphopyruvate hydratase eno1  GUK1 - guanylate kinase  HXK2 - hexokinase 2  NDE1 - nadh-ubiquinone reductase (h(+)-translocating) nde1  ATP1 - f1f0 atp synthase subunit alpha  ATP2 - atp2p  GLK1 - glucokinase  ADE6 - phosphoribosylformylglycinamidine synthase  EMI2 - putative glucokinase  ADE16 - bifunctional phosphoribosylaminoimidazolecarboxamide formyltransferase/imp cyclohydrolase ade16  ATP3 - atp3p |
| GO:0006529 | asparagine biosynthetic process | 3E-4 | 1.47E-2 | 90.12 (2298,3,17,2) | [+] Show genes  ASN1 - asparagine synthase (glutamine-hydrolyzing) 1  ASN2 - asparagine synthase (glutamine-hydrolyzing) 2 |
| GO:0046034 | ATP metabolic process | 3.42E-4 | 1.66E-2 | 4.67 (2298,40,123,10) | [+] Show genes  ENO1 - phosphopyruvate hydratase eno1  HXK2 - hexokinase 2  NDE1 - nadh-ubiquinone reductase (h(+)-translocating) nde1  ENO2 - phosphopyruvate hydratase eno2  ATP1 - f1f0 atp synthase subunit alpha  ATP2 - atp2p  GLK1 - glucokinase  PDC1 - indolepyruvate decarboxylase 1  EMI2 - putative glucokinase  ATP3 - atp3p |
| GO:0001678 | cellular glucose homeostasis | 3.71E-4 | 1.78E-2 | 9.23 (2298,4,249,4) | [+] Show genes  HXK2 - hexokinase 2  HXK1 - hexokinase 1  GLK1 - glucokinase  EMI2 - putative glucokinase |
| GO:0033500 | carbohydrate homeostasis | 3.71E-4 | 1.76E-2 | 9.23 (2298,4,249,4) | [+] Show genes  HXK2 - hexokinase 2  HXK1 - hexokinase 1  GLK1 - glucokinase  EMI2 - putative glucokinase |
| GO:0042593 | glucose homeostasis | 3.71E-4 | 1.74E-2 | 9.23 (2298,4,249,4) | [+] Show genes  HXK2 - hexokinase 2  HXK1 - hexokinase 1  GLK1 - glucokinase  EMI2 - putative glucokinase |
| GO:0072524 | pyridine-containing compound metabolic process | 3.72E-4 | 1.73E-2 | 2.08 (2298,63,491,28) | [+] Show genes  PYK2 - pyruvate kinase pyk2  ISN1 - imp 5'-nucleotidase  ENO2 - phosphopyruvate hydratase eno2  NPT1 - nicotinate phosphoribosyltransferase  PDC1 - indolepyruvate decarboxylase 1  ALD6 - aldehyde dehydrogenase (nadp(+)) ald6  TAL1 - sedoheptulose-7-phosphate:d-glyceraldehyde-3-phosphate transaldolase tal1  GPD1 - glycerol-3-phosphate dehydrogenase (nad(+)) gpd1  BNA1 - bna1p  YMR315W - hypothetical protein  ENO1 - phosphopyruvate hydratase eno1  HXK2 - hexokinase 2  ZWF1 - glucose-6-phosphate dehydrogenase  GLK1 - glucokinase  GPM1 - phosphoglycerate mutase gpm1  YKL151C - nadhx dehydratase  EMI2 - putative glucokinase  GND1 - phosphogluconate dehydrogenase (decarboxylating) gnd1  PNC1 - nicotinamidase  MDH3 - malate dehydrogenase mdh3  ALD4 - aldehyde dehydrogenase (nadp(+)) ald4  BNA5 - kynureninase  POS5 - pos5p  NDI1 - nadh-ubiquinone reductase (h(+)-translocating) ndi1  NDE1 - nadh-ubiquinone reductase (h(+)-translocating) nde1  GUT2 - glycerol-3-phosphate dehydrogenase  BNA4 - kynurenine 3-monooxygenase  HXK1 - hexokinase 1 |
| GO:0009260 | ribonucleotide biosynthetic process | 3.72E-4 | 1.71E-2 | 3.49 (2298,75,123,14) | [+] Show genes  CAB2 - phosphopantothenate--cysteine ligase cab2  ADE2 - phosphoribosylaminoimidazole carboxylase ade2  ENO2 - phosphopyruvate hydratase eno2  AMD1 - amp deaminase  ENO1 - phosphopyruvate hydratase eno1  GUK1 - guanylate kinase  HXK2 - hexokinase 2  ATP1 - f1f0 atp synthase subunit alpha  ATP2 - atp2p  GLK1 - glucokinase  ADE6 - phosphoribosylformylglycinamidine synthase  EMI2 - putative glucokinase  ADE16 - bifunctional phosphoribosylaminoimidazolecarboxamide formyltransferase/imp cyclohydrolase ade16  ATP3 - atp3p |
| GO:0043650 | dicarboxylic acid biosynthetic process | 4.29E-4 | 1.95E-2 | 4.73 (2298,24,182,9) | [+] Show genes  IDH1 - isocitrate dehydrogenase (nad(+)) idh1  ARO1 - pentafunctional protein aro1p  ARO3 - 3-deoxy-7-phosphoheptulonate synthase aro3  CIT1 - citrate (si)-synthase cit1  IDH2 - isocitrate dehydrogenase (nad(+)) idh2  ARO2 - bifunctional chorismate synthase/riboflavin reductase [nad(p)h] aro2  GLT1 - glutamate synthase (nadh)  BNA5 - kynureninase  ARO4 - 3-deoxy-7-phosphoheptulonate synthase aro4 |
| GO:0071941 | nitrogen cycle metabolic process | 4.7E-4 | 2.11E-2 | 88.38 (2298,4,13,2) | [+] Show genes  DUR1,2 - bifunctional urea carboxylase/allophanate hydrolase  ARG1 - argininosuccinate synthase |
| GO:0072522 | purine-containing compound biosynthetic process | 5E-4 | 2.22E-2 | 3.40 (2298,77,123,14) | [+] Show genes  CAB2 - phosphopantothenate--cysteine ligase cab2  ADE2 - phosphoribosylaminoimidazole carboxylase ade2  ENO2 - phosphopyruvate hydratase eno2  AMD1 - amp deaminase  ENO1 - phosphopyruvate hydratase eno1  GUK1 - guanylate kinase  HXK2 - hexokinase 2  ATP1 - f1f0 atp synthase subunit alpha  ATP2 - atp2p  GLK1 - glucokinase  ADE6 - phosphoribosylformylglycinamidine synthase  ADE16 - bifunctional phosphoribosylaminoimidazolecarboxamide formyltransferase/imp cyclohydrolase ade16  EMI2 - putative glucokinase  ATP3 - atp3p |
| GO:0006796 | phosphate-containing compound metabolic process | 5.07E-4 | 2.23E-2 | 1.87 (2298,343,136,38) | [+] Show genes  CAB2 - phosphopantothenate--cysteine ligase cab2  PGM1 - phosphoglucomutase pgm1  SIT4 - sit4p  ENO2 - phosphopyruvate hydratase eno2  TSL1 - tsl1p  HOR2 - glycerol-1-phosphatase hor2  ACH1 - ach1p  PDC1 - indolepyruvate decarboxylase 1  GPD1 - glycerol-3-phosphate dehydrogenase (nad(+)) gpd1  APA1 - apa1p  LSC1 - succinate--coa ligase (gdp-forming) subunit alpha  GUK1 - guanylate kinase  ENO1 - phosphopyruvate hydratase eno1  HXK2 - hexokinase 2  ZWF1 - glucose-6-phosphate dehydrogenase  GLK1 - glucokinase  ADE6 - phosphoribosylformylglycinamidine synthase  YKL151C - nadhx dehydratase  ADE16 - bifunctional phosphoribosylaminoimidazolecarboxamide formyltransferase/imp cyclohydrolase ade16  ATP3 - atp3p  GDE1 - gde1p  EMI2 - putative glucokinase  ADE2 - phosphoribosylaminoimidazole carboxylase ade2  PNC1 - nicotinamidase  PGM2 - phosphoglucomutase pgm2  MDH3 - malate dehydrogenase mdh3  ALD4 - aldehyde dehydrogenase (nadp(+)) ald4  CIT1 - citrate (si)-synthase cit1  GFA1 - glutamine--fructose-6-phosphate transaminase (isomerizing) gfa1  CDC28 - cdc28p  AMD1 - amp deaminase  ARO1 - pentafunctional protein aro1p  NDE1 - nadh-ubiquinone reductase (h(+)-translocating) nde1  ATP2 - atp2p  GLC7 - glc7p  ATP1 - f1f0 atp synthase subunit alpha  ERG12 - mevalonate kinase  PTP1 - ptp1p |
| GO:0006793 | phosphorus metabolic process | 5.15E-4 | 2.24E-2 | 1.85 (2298,356,136,39) | [+] Show genes  CAB2 - phosphopantothenate--cysteine ligase cab2  PGM1 - phosphoglucomutase pgm1  SIT4 - sit4p  ENO2 - phosphopyruvate hydratase eno2  TSL1 - tsl1p  HOR2 - glycerol-1-phosphatase hor2  ACH1 - ach1p  PDC1 - indolepyruvate decarboxylase 1  GPD1 - glycerol-3-phosphate dehydrogenase (nad(+)) gpd1  APA1 - apa1p  LSC1 - succinate--coa ligase (gdp-forming) subunit alpha  GUK1 - guanylate kinase  ENO1 - phosphopyruvate hydratase eno1  HXK2 - hexokinase 2  ZWF1 - glucose-6-phosphate dehydrogenase  GLK1 - glucokinase  ADE6 - phosphoribosylformylglycinamidine synthase  YKL151C - nadhx dehydratase  ADE16 - bifunctional phosphoribosylaminoimidazolecarboxamide formyltransferase/imp cyclohydrolase ade16  ATP3 - atp3p  GDE1 - gde1p  EMI2 - putative glucokinase  ADE2 - phosphoribosylaminoimidazole carboxylase ade2  PNC1 - nicotinamidase  PGM2 - phosphoglucomutase pgm2  MDH3 - malate dehydrogenase mdh3  ALD4 - aldehyde dehydrogenase (nadp(+)) ald4  CIT1 - citrate (si)-synthase cit1  GFA1 - glutamine--fructose-6-phosphate transaminase (isomerizing) gfa1  UGP1 - utp glucose-1-phosphate uridylyltransferase  CDC28 - cdc28p  AMD1 - amp deaminase  ARO1 - pentafunctional protein aro1p  NDE1 - nadh-ubiquinone reductase (h(+)-translocating) nde1  ATP2 - atp2p  GLC7 - glc7p  ATP1 - f1f0 atp synthase subunit alpha  ERG12 - mevalonate kinase  PTP1 - ptp1p |
| GO:0110020 | regulation of actomyosin structure organization | 5.38E-4 | 2.31E-2 | 20.04 (2298,4,86,3) | [+] Show genes  GLC7 - glc7p  RHO1 - rho1p  CDC28 - cdc28p |
| GO:1901575 | organic substance catabolic process | 5.5E-4 | 2.34E-2 | 2.00 (2298,335,110,32) | [+] Show genes  PGM1 - phosphoglucomutase pgm1  GPH1 - gph1p  ARO8 - bifunctional 2-aminoadipate transaminase/aromatic-amino-acid:2-oxoglutarate transaminase  GRE3 - trifunctional aldehyde reductase/xylose reductase/glucose 1-dehydrogenase (nadp(+))  PDC1 - indolepyruvate decarboxylase 1  DCS1 - dcs1p  GLO1 - lactoylglutathione lyase glo1  GPD1 - glycerol-3-phosphate dehydrogenase (nad(+)) gpd1  APA1 - apa1p  SSE1 - sse1p  ENO1 - phosphopyruvate hydratase eno1  HXK2 - hexokinase 2  EMI2 - putative glucokinase  GDE1 - gde1p  EHT1 - eht1p  ISC1 - inositol phosphosphingolipid phospholipase  PGM2 - phosphoglucomutase pgm2  MDH3 - malate dehydrogenase mdh3  PCS60 - pcs60p  CIT1 - citrate (si)-synthase cit1  SSA2 - hsp70 family chaperone ssa2  DLD1 - dld1p  ALD2 - aldehyde dehydrogenase (nad(+)) ald2  CDC28 - cdc28p  HFD1 - hfd1p  YDJ1 - ydj1p  DCS2 - dcs2p  DUR1,2 - bifunctional urea carboxylase/allophanate hydrolase  CAR2 - ornithine-oxo-acid transaminase  NTH1 - alpha,alpha-trehalase nth1  SCJ1 - scj1p  ALD5 - aldehyde dehydrogenase (nad(p)(+)) ald5 |
| GO:0046496 | nicotinamide nucleotide metabolic process | 5.68E-4 | 2.39E-2 | 2.10 (2298,58,491,26) | [+] Show genes  PYK2 - pyruvate kinase pyk2  ENO2 - phosphopyruvate hydratase eno2  NPT1 - nicotinate phosphoribosyltransferase  PDC1 - indolepyruvate decarboxylase 1  ALD6 - aldehyde dehydrogenase (nadp(+)) ald6  TAL1 - sedoheptulose-7-phosphate:d-glyceraldehyde-3-phosphate transaldolase tal1  GPD1 - glycerol-3-phosphate dehydrogenase (nad(+)) gpd1  BNA1 - bna1p  YMR315W - hypothetical protein  ENO1 - phosphopyruvate hydratase eno1  HXK2 - hexokinase 2  ZWF1 - glucose-6-phosphate dehydrogenase  GLK1 - glucokinase  GPM1 - phosphoglycerate mutase gpm1  YKL151C - nadhx dehydratase  EMI2 - putative glucokinase  GND1 - phosphogluconate dehydrogenase (decarboxylating) gnd1  MDH3 - malate dehydrogenase mdh3  ALD4 - aldehyde dehydrogenase (nadp(+)) ald4  BNA5 - kynureninase  NDI1 - nadh-ubiquinone reductase (h(+)-translocating) ndi1  POS5 - pos5p  GUT2 - glycerol-3-phosphate dehydrogenase  NDE1 - nadh-ubiquinone reductase (h(+)-translocating) nde1  BNA4 - kynurenine 3-monooxygenase  HXK1 - hexokinase 1 |
| GO:0006549 | isoleucine metabolic process | 5.83E-4 | 2.43E-2 | 5.15 (2298,11,284,7) | [+] Show genes  ILV5 - ketol-acid reductoisomerase  HOM2 - aspartate-semialdehyde dehydrogenase  ILV2 - acetolactate synthase catalytic subunit  ILV3 - ilv3p  ILV6 - acetolactate synthase regulatory subunit  ILV1 - threonine ammonia-lyase ilv1  BAT2 - bat2p |
| GO:0042026 | protein refolding | 5.9E-4 | 2.44E-2 | 8.20 (2298,29,58,6) | [+] Show genes  HSP82 - hsp90 family chaperone hsp82  YDJ1 - ydj1p  SSE1 - sse1p  SSA2 - hsp70 family chaperone ssa2  CPR6 - peptidylprolyl isomerase cpr6  HSC82 - hsp90 family chaperone hsc82 |
| GO:0009099 | valine biosynthetic process | 6.35E-4 | 2.59E-2 | 6.74 (2298,6,284,5) | [+] Show genes  ILV5 - ketol-acid reductoisomerase  ILV2 - acetolactate synthase catalytic subunit  ILV3 - ilv3p  ILV6 - acetolactate synthase regulatory subunit  BAT2 - bat2p |
| GO:0071704 | organic substance metabolic process | 7.22E-4 | 2.92E-2 | 1.15 (2298,1498,315,236) | [+] Show genes  IDH1 - isocitrate dehydrogenase (nad(+)) idh1  PHS1 - phs1p  RSP5 - nedd4 family e3 ubiquitin-protein ligase  SIT4 - sit4p  MSC6 - msc6p  ARO2 - bifunctional chorismate synthase/riboflavin reductase [nad(p)h] aro2  RPL9A - ribosomal 60s subunit protein l9a  GPD1 - glycerol-3-phosphate dehydrogenase (nad(+)) gpd1  APA1 - apa1p  LAP2 - lap2p  GLK1 - glucokinase  CPR6 - peptidylprolyl isomerase cpr6  EMI2 - putative glucokinase  SIS1 - sis1p  FAA1 - long-chain fatty acid-coa ligase faa1  PGM2 - phosphoglucomutase pgm2  HIS4 - trifunctional histidinol dehydrogenase/phosphoribosyl-amp cyclohydrolase/phosphoribosyl-atp diphosphatase  QCR2 - ubiquinol--cytochrome-c reductase subunit 2  CIT1 - citrate (si)-synthase cit1  ILV2 - acetolactate synthase catalytic subunit  PRO2 - glutamate-5-semialdehyde dehydrogenase  MAS1 - mas1p  HFD1 - hfd1p  NFS1 - nfs1p  RIB3 - 3,4-dihydroxy-2-butanone-4-phosphate synthase rib3  YTA12 - m-aaa protease subunit yta12  CPA1 - carbamoyl-phosphate synthase (glutamine-hydrolyzing) cpa1  GLC7 - glc7p  ILV6 - acetolactate synthase regulatory subunit  NTH1 - alpha,alpha-trehalase nth1  ARG5,6 - bifunctional acetylglutamate kinase/n-acetyl-gamma-glutamyl-phosphate reductase  YMR196W - hypothetical protein  GPH1 - gph1p  GSY2 - glycogen (starch) synthase gsy2  YDL124W - aldo-keto reductase superfamily protein  HOR2 - glycerol-1-phosphatase hor2  CYM1 - cym1p  PAP1 - pap1p  OSH6 - oxysterol-binding protein osh6  DCS1 - dcs1p  YDL119C - hypothetical protein  HUB1 - hub1p  ASN1 - asparagine synthase (glutamine-hydrolyzing) 1  HIS1 - atp phosphoribosyltransferase  GUK1 - guanylate kinase  ATP7 - f1f0 atp synthase subunit d  HOM3 - aspartate kinase  HEM4 - uroporphyrinogen-iii synthase hem4  HSC82 - hsp90 family chaperone hsc82  RPB7 - rpb7p  PNC1 - nicotinamidase  MDH3 - malate dehydrogenase mdh3  YGL039W - carbonyl reductase (nadph-dependent)  HEM2 - porphobilinogen synthase hem2  PAI3 - pai3p  TRP2 - anthranilate synthase trp2  BNA5 - kynureninase  ALD2 - aldehyde dehydrogenase (nad(+)) ald2  ILV1 - threonine ammonia-lyase ilv1  MRPS12 - putative mitochondrial 37s ribosomal protein mrps12  RPL33B - ribosomal 60s subunit protein l33b  OLE1 - stearoyl-coa 9-desaturase  NDE1 - nadh-ubiquinone reductase (h(+)-translocating) nde1  ALD5 - aldehyde dehydrogenase (nad(p)(+)) ald5  HSP82 - hsp90 family chaperone hsp82  TRX3 - trx3p  RRP1 - rrp1p  ACH1 - ach1p  BGL2 - bgl2p  TIF11 - tif11p  SCW4 - scw4p  HIS3 - imidazoleglycerol-phosphate dehydratase his3  TAL1 - sedoheptulose-7-phosphate:d-glyceraldehyde-3-phosphate transaldolase tal1  ILV5 - ketol-acid reductoisomerase  GAD1 - glutamate decarboxylase gad1  ERG7 - lanosterol synthase erg7  ENO1 - phosphopyruvate hydratase eno1  HSM3 - hsm3p  UBC13 - e2 ubiquitin-conjugating protein ubc13  RPL26A - ribosomal 60s subunit protein l26a  COQ1 - trans-hexaprenyltranstransferase  RPP0 - ribosomal protein p0  MDH1 - malate dehydrogenase mdh1  MET17 - bifunctional cysteine synthase/o-acetylhomoserine aminocarboxypropyltransferase met17  FUM1 - fumarase fum1  TUF1 - tuf1p  MAE1 - malate dehydrogenase (oxaloacetate-decarboxylating)  UGP1 - utp glucose-1-phosphate uridylyltransferase  ATP15 - f1f0 atp synthase subunit epsilon  GAS3 - gas3p  DCS2 - dcs2p  ARO1 - pentafunctional protein aro1p  ADE5,7 - bifunctional aminoimidazole ribotide synthase/glycinamide ribotide synthase  YCF1 - atp-binding cassette glutathione s-conjugate transporter ycf1  ERG12 - mevalonate kinase  AAP1 - aap1p  SCJ1 - scj1p  RVB2 - ruvb family atp-dependent dna helicase reptin  PGM1 - phosphoglucomutase pgm1  ISN1 - imp 5'-nucleotidase  ARO8 - bifunctional 2-aminoadipate transaminase/aromatic-amino-acid:2-oxoglutarate transaminase  SLT2 - slt2p  ACT1 - actin  LSC1 - succinate--coa ligase (gdp-forming) subunit alpha  YMR315W - hypothetical protein  HXK2 - hexokinase 2  ARO3 - 3-deoxy-7-phosphoheptulonate synthase aro3  EHD3 - ehd3p  IDH2 - isocitrate dehydrogenase (nad(+)) idh2  KRS1 - lysine--trna ligase krs1  CCA1 - cca1p  GNA1 - glucosamine 6-phosphate n-acetyltransferase  YKL151C - nadhx dehydratase  MCR1 - mcr1p  YHR020W - proline--trna ligase  ADE2 - phosphoribosylaminoimidazole carboxylase ade2  HEM13 - coproporphyrinogen oxidase  APE1 - ape1p  GFA1 - glutamine--fructose-6-phosphate transaminase (isomerizing) gfa1  BLM10 - blm10p  LCB2 - serine c-palmitoyltransferase lcb2  INP53 - phosphatidylinositol-3-/phosphoinositide 5-phosphatase inp53  AIM17 - aim17p  ATP1 - f1f0 atp synthase subunit alpha  TPS2 - trehalose-phosphatase tps2  CDC55 - cdc55p  TPS1 - alpha,alpha-trehalose-phosphate synthase (udp-forming) tps1  ENO2 - phosphopyruvate hydratase eno2  TSL1 - tsl1p  MET5 - met5p  TFC1 - tfc1p  ASN2 - asparagine synthase (glutamine-hydrolyzing) 2  YSA1 - ysa1p  GCD6 - gcd6p  FES1 - fes1p  BAT2 - bat2p  RSM7 - rsm7p  PPX1 - ppx1p  NHP6B - nhp6bp  POL30 - pol30p  CPA2 - cpa2p  FRA1 - fra1p  RPL3 - ribosomal 60s subunit protein l3  EXG2 - exg2p  HSP104 - chaperone atpase hsp104  SSA2 - hsp70 family chaperone ssa2  GLR1 - glutathione-disulfide reductase glr1  RSM26 - rsm26p  ERG24 - delta(14)-sterol reductase  RPN12 - proteasome regulatory particle lid subunit rpn12  ATP2 - atp2p  HXK1 - hexokinase 1  CAR2 - ornithine-oxo-acid transaminase  THI4 - thi4p  CAB2 - phosphopantothenate--cysteine ligase cab2  ORC5 - origin recognition complex subunit 5  GRE3 - trifunctional aldehyde reductase/xylose reductase/glucose 1-dehydrogenase (nadp(+))  SSE1 - sse1p  KGD2 - alpha-ketoglutarate dehydrogenase kgd2  HOM2 - aspartate-semialdehyde dehydrogenase  APE2 - ape2p  ZWF1 - glucose-6-phosphate dehydrogenase  ATP3 - atp3p  GDE1 - gde1p  RPL6A - ribosomal 60s subunit protein l6a  ELP2 - elongator subunit elp2  PEP4 - pep4p  ARO9 - aromatic-amino-acid:2-oxoglutarate transaminase  CRH1 - crh1p  ERG13 - hydroxymethylglutaryl-coa synthase  BDH1 - (r,r)-butanediol dehydrogenase  SGT2 - sgt2p  YIL108W - putative metalloendopeptidase  CDC60 - leucine--trna ligase cdc60  SDH2 - succinate dehydrogenase iron-sulfur protein subunit sdh2  TPD3 - tpd3p  RPL5 - ribosomal 60s subunit protein l5  KES1 - kes1p  SFM1 - sfm1p  GLC3 - 1,4-alpha-glucan branching enzyme  RPL24A - ribosomal 60s subunit protein l24a  RPL30 - ribosomal 60s subunit protein l30  TRP5 - tryptophan synthase trp5  MRPL19 - mitochondrial 54s ribosomal protein yml19  RIB5 - riboflavin synthase  CWH41 - cwh41p  ILV3 - ilv3p  PDC1 - indolepyruvate decarboxylase 1  GLO1 - lactoylglutathione lyase glo1  KGD1 - alpha-ketoglutarate dehydrogenase kgd1  AYR1 - acylglycerone-phosphate reductase  YDR341C - arginine--trna ligase  RPL15A - ribosomal 60s subunit protein l15a  ADE16 - bifunctional phosphoribosylaminoimidazolecarboxamide formyltransferase/imp cyclohydrolase ade16  ARO4 - 3-deoxy-7-phosphoheptulonate synthase aro4  ISC1 - inositol phosphosphingolipid phospholipase  AFG3 - aaa family atpase afg3  HTS1 - histidine--trna ligase  GLN1 - glutamate--ammonia ligase  ARG1 - argininosuccinate synthase  PCS60 - pcs60p  LSM6 - lsm6p  RPL10 - ribosomal 60s subunit protein l10  AMD1 - amp deaminase  UGA1 - 4-aminobutyrate transaminase  RPP2A - ribosomal protein p2a  DUR1,2 - bifunctional urea carboxylase/allophanate hydrolase  NHP6A - nhp6ap  ERG3 - c-5 sterol desaturase  SHM2 - glycine hydroxymethyltransferase shm2  FCY1 - cytosine deaminase  ARG7 - glutamate n-acetyltransferase  ACB1 - long-chain fatty acid transporter acb1  DLD3 - dld3p  PRB1 - prb1p  ATP5 - atp5p  SSE2 - sse2p  RPO21 - rpo21p  ADE6 - phosphoribosylformylglycinamidine synthase  DPP1 - bifunctional diacylglycerol diphosphate phospatase/phosphatidate phosphatase  EHT1 - eht1p  STM1 - stm1p  ALD4 - aldehyde dehydrogenase (nadp(+)) ald4  GLT1 - glutamate synthase (nadh)  YMR027W - hypothetical protein  DLD1 - dld1p  CDC28 - cdc28p  MKT1 - mkt1p  YDJ1 - ydj1p  CPR5 - peptidylprolyl isomerase family protein cpr5  DLD2 - dld2p  ARA1 - d-arabinose 1-dehydrogenase (nad(p)(+)) ara1  ARA2 - d-arabinose 1-dehydrogenase (nad(p)(+)) ara2  RRP46 - rrp46p  PTP1 - ptp1p  RPL16B - ribosomal 60s subunit protein l16b |
| GO:0009141 | nucleoside triphosphate metabolic process | 7.57E-4 | 3.03E-2 | 3.95 (2298,52,123,11) | [+] Show genes  LSC1 - succinate--coa ligase (gdp-forming) subunit alpha  ENO1 - phosphopyruvate hydratase eno1  HXK2 - hexokinase 2  NDE1 - nadh-ubiquinone reductase (h(+)-translocating) nde1  ENO2 - phosphopyruvate hydratase eno2  ATP1 - f1f0 atp synthase subunit alpha  ATP2 - atp2p  GLK1 - glucokinase  PDC1 - indolepyruvate decarboxylase 1  ATP3 - atp3p  EMI2 - putative glucokinase |
| GO:0034637 | cellular carbohydrate biosynthetic process | 7.8E-4 | 3.09E-2 | 3.69 (2298,31,221,11) | [+] Show genes  TPS1 - alpha,alpha-trehalose-phosphate synthase (udp-forming) tps1  GAS3 - gas3p  PGM1 - phosphoglucomutase pgm1  PGM2 - phosphoglucomutase pgm2  GSY2 - glycogen (starch) synthase gsy2  CWH41 - cwh41p  HOR2 - glycerol-1-phosphatase hor2  TSL1 - tsl1p  GFA1 - glutamine--fructose-6-phosphate transaminase (isomerizing) gfa1  TPS2 - trehalose-phosphatase tps2  UGP1 - utp glucose-1-phosphate uridylyltransferase |
| GO:0051156 | glucose 6-phosphate metabolic process | 8E-4 | 3.14E-2 | 4.61 (2298,16,249,8) | [+] Show genes  TAL1 - sedoheptulose-7-phosphate:d-glyceraldehyde-3-phosphate transaldolase tal1  PGM1 - phosphoglucomutase pgm1  HXK2 - hexokinase 2  PGM2 - phosphoglucomutase pgm2  ZWF1 - glucose-6-phosphate dehydrogenase  HXK1 - hexokinase 1  GLK1 - glucokinase  EMI2 - putative glucokinase |
| GO:0016052 | carbohydrate catabolic process | 8.13E-4 | 3.16E-2 | 4.25 (2298,44,123,10) | [+] Show genes  PGM1 - phosphoglucomutase pgm1  GPH1 - gph1p  ENO1 - phosphopyruvate hydratase eno1  PGM2 - phosphoglucomutase pgm2  HXK2 - hexokinase 2  ENO2 - phosphopyruvate hydratase eno2  GLK1 - glucokinase  GRE3 - trifunctional aldehyde reductase/xylose reductase/glucose 1-dehydrogenase (nadp(+))  NTH1 - alpha,alpha-trehalase nth1  EMI2 - putative glucokinase |
| GO:0009205 | purine ribonucleoside triphosphate metabolic process | 8.13E-4 | 3.13E-2 | 4.25 (2298,44,123,10) | [+] Show genes  ENO1 - phosphopyruvate hydratase eno1  HXK2 - hexokinase 2  NDE1 - nadh-ubiquinone reductase (h(+)-translocating) nde1  ENO2 - phosphopyruvate hydratase eno2  ATP2 - atp2p  ATP1 - f1f0 atp synthase subunit alpha  GLK1 - glucokinase  PDC1 - indolepyruvate decarboxylase 1  EMI2 - putative glucokinase  ATP3 - atp3p |
| GO:0000255 | allantoin metabolic process | 8.7E-4 | 3.32E-2 | 1,149.00 (2298,1,2,1) | [+] Show genes  DUR1,2 - bifunctional urea carboxylase/allophanate hydrolase |
| GO:0000256 | allantoin catabolic process | 8.7E-4 | 3.29E-2 | 1,149.00 (2298,1,2,1) | [+] Show genes  DUR1,2 - bifunctional urea carboxylase/allophanate hydrolase |
| GO:0043605 | cellular amide catabolic process | 8.7E-4 | 3.26E-2 | 1,149.00 (2298,1,2,1) | [+] Show genes  DUR1,2 - bifunctional urea carboxylase/allophanate hydrolase |
| GO:0043419 | urea catabolic process | 8.7E-4 | 3.24E-2 | 1,149.00 (2298,1,2,1) | [+] Show genes  DUR1,2 - bifunctional urea carboxylase/allophanate hydrolase |
| GO:0046390 | ribose phosphate biosynthetic process | 9.17E-4 | 3.38E-2 | 3.23 (2298,81,123,14) | [+] Show genes  CAB2 - phosphopantothenate--cysteine ligase cab2  ADE2 - phosphoribosylaminoimidazole carboxylase ade2  ENO2 - phosphopyruvate hydratase eno2  AMD1 - amp deaminase  ENO1 - phosphopyruvate hydratase eno1  GUK1 - guanylate kinase  HXK2 - hexokinase 2  ATP1 - f1f0 atp synthase subunit alpha  ATP2 - atp2p  GLK1 - glucokinase  ADE6 - phosphoribosylformylglycinamidine synthase  ADE16 - bifunctional phosphoribosylaminoimidazolecarboxamide formyltransferase/imp cyclohydrolase ade16  EMI2 - putative glucokinase  ATP3 - atp3p |

Species used: Saccharomyces cerevisiae

The system has recognized 2316 genes out of 2316 gene terms entered by the user.  
 136 genes were recognized by gene symbol and 2180 genes by other gene IDs .  
Only 2298 of these genes are associated with a GO term.

  
 Output in Microsoft Excel format  

% List genereted using GOrilla
% http://cbl-gorilla.cs.technion.ac.il/
% GO term pValue
GO:0044281 1.27E-18
GO:0019752 4.6E-14
GO:0043436 5.41E-14
GO:0006082 6.92E-14
GO:0044283 3.44E-11
GO:0016053 2.24E-10
GO:0046394 2.24E-10
GO:0008652 1.04E-9
GO:0006520 2.31E-9
GO:0043648 2.44E-9
GO:1901605 2.24E-8
GO:0017144 3.14E-8
GO:1901607 6.73E-8
GO:1901566 1.05E-7
GO:0005975 1.46E-7
GO:0072350 2.65E-7
GO:0055114 3.15E-7
GO:0005991 3.27E-7
GO:1901564 4.37E-7
GO:0009150 5.59E-7
GO:0055086 5.8E-7
GO:0009064 6.88E-7
GO:0044249 8.36E-7
GO:0006099 8.97E-7
GO:0006101 8.97E-7
GO:0016999 1.53E-6
GO:0009311 2.27E-6
GO:1901576 2.86E-6
GO:0006163 3.36E-6
GO:0006091 4.12E-6
GO:0009259 4.74E-6
GO:0005984 6.25E-6
GO:0009126 6.62E-6
GO:0009167 6.62E-6
GO:0009117 7.07E-6
GO:0044262 8.01E-6
GO:0006753 8.83E-6
GO:0019318 9.48E-6
GO:0009423 1.12E-5
GO:0009058 1.2E-5
GO:0019693 1.43E-5
GO:0051186 1.43E-5
GO:0032787 1.71E-5
GO:0072521 1.75E-5
GO:0009072 1.82E-5
GO:0005992 2.09E-5
GO:0046351 2.09E-5
GO:0009312 2.09E-5
GO:0015980 2.11E-5
GO:0006006 2.98E-5
GO:0009084 3.3E-5
GO:0009073 5.11E-5
GO:0070981 5.15E-5
GO:0070982 5.15E-5
GO:0005996 5.49E-5
GO:0009152 5.79E-5
GO:0009132 6.05E-5
GO:0046417 7.28E-5
GO:0006011 8.5E-5
GO:2000431 8.83E-5
GO:1903499 8.83E-5
GO:0019637 9.22E-5
GO:0009161 1.02E-4
GO:0044282 1.28E-4
GO:1901135 1.3E-4
GO:0009127 1.44E-4
GO:0009168 1.44E-4
GO:0006525 1.51E-4
GO:0006012 1.52E-4
GO:0019388 1.52E-4
GO:0019627 1.72E-4
GO:0034982 1.99E-4
GO:0006108 2.04E-4
GO:0006733 2.09E-4
GO:0009097 2.28E-4
GO:0006732 2.3E-4
GO:0006164 2.37E-4
GO:0009185 2.68E-4
GO:0009179 2.68E-4
GO:0009135 2.68E-4
GO:0019362 2.71E-4
GO:0072330 2.76E-4
GO:0009123 2.93E-4
GO:0006529 3E-4
GO:0046034 3.42E-4
GO:0001678 3.71E-4
GO:0033500 3.71E-4
GO:0042593 3.71E-4
GO:0072524 3.72E-4
GO:0009260 3.72E-4
GO:0043650 4.29E-4
GO:0071941 4.7E-4
GO:0072522 5E-4
GO:0006796 5.07E-4
GO:0006793 5.15E-4
GO:0110020 5.38E-4
GO:1901575 5.5E-4
GO:0046496 5.68E-4
GO:0006549 5.83E-4
GO:0042026 5.9E-4
GO:0009099 6.35E-4
GO:0071704 7.22E-4
GO:0009141 7.57E-4
GO:0034637 7.8E-4
GO:0051156 8E-4
GO:0016052 8.13E-4
GO:0009205 8.13E-4
GO:0000255 8.7E-4
GO:0000256 8.7E-4
GO:0043605 8.7E-4
GO:0043419 8.7E-4
GO:0046390 9.17E-4
 Visualize output in REViGO   

The GOrilla database is periodically updated using the GO database and other sources.  
The GOrilla database was last updated on Sep 21, 2019

This results page will be available on this site for one month from now (until
Oct 23, 2019
). You can bookmark this page and come back to it later.

  
**'P-value'** is the enrichment
p-value computed according to the mHG or HG model. This p-value is not
corrected for multiple testing of 4126 GO terms.  
  
**'FDR q-value'** is the correction of the above p-value for multiple testing using the Benjamini and Hochberg (1995) method.   
Namely, for the i^th^ term (ranked according to p-value) the FDR q-value is (p-value \* number of GO terms) / i.   
  
**Enrichment (N, B, n, b)** is defined as follows:  
N - is the total number of genes  
B - is the total number of genes associated with a specific GO term  
n - is the number of genes in the top of the user's input list or in the target set when appropriate  
b - is the number of genes in the intersection  
Enrichment = (b/n) / (B/N)  
  
**Genes:** For each GO term you can see the list of associated genes that appear in the optimal top of the list.  
Each gene name is specified by gene symbol followed by a short description of the gene   

Back to GOrilla analysis results
