## Supplemental Data S1 for "A simple mass-action model predicts genome-wide protein timecourses from mRNA trajectories during a dynamic response in two strains of *Saccharomyces cerevisiae*": GOPROCESS_protein.html

Results


 **protein** 

*P-value color scale*

|  |  |  |  |  |
| --- | --- | --- | --- | --- |
| > 10^-3^ | 10^-3^ to 10^-5^ | 10^-5^ to 10^-7^ | 10^-7^ to 10^-9^ | < 10^-9^ |


|  |  |  |  |  |  |
| --- | --- | --- | --- | --- | --- |
| **GO term** | **Description** | **P-value** | **FDR q-value** | **Enrichment (N, B, n, b)** | **Genes** |
| GO:0008652 | cellular amino acid biosynthetic process | 2.15E-18 | 8.89E-15 | 7.57 (2325,94,98,30) | [+] Show genes  IDH1 - isocitrate dehydrogenase (nad(+)) idh1  ARG5,6 - bifunctional acetylglutamate kinase/n-acetyl-gamma-glutamyl-phosphate reductase  ARG7 - glutamate n-acetyltransferase  ARO2 - bifunctional chorismate synthase/riboflavin reductase [nad(p)h] aro2  ILV3 - ilv3p  ARO8 - bifunctional 2-aminoadipate transaminase/aromatic-amino-acid:2-oxoglutarate transaminase  TRP3 - bifunctional anthranilate synthase/indole-3-glycerol-phosphate synthase  HIS3 - imidazoleglycerol-phosphate dehydratase his3  ILV5 - ketol-acid reductoisomerase  ASN1 - asparagine synthase (glutamine-hydrolyzing) 1  HOM2 - aspartate-semialdehyde dehydrogenase  HOM3 - aspartate kinase  ARO3 - 3-deoxy-7-phosphoheptulonate synthase aro3  IDH2 - isocitrate dehydrogenase (nad(+)) idh2  LEU4 - 2-isopropylmalate synthase leu4  HIS7 - imidazoleglycerol-phosphate synthase  ARG4 - argininosuccinate lyase arg4  MET10 - sulfite reductase subunit alpha  ARO4 - 3-deoxy-7-phosphoheptulonate synthase aro4  CPA2 - cpa2p  GLN1 - glutamate--ammonia ligase  ARG1 - argininosuccinate synthase  HIS4 - trifunctional histidinol dehydrogenase/phosphoribosyl-amp cyclohydrolase/phosphoribosyl-atp diphosphatase  GLT1 - glutamate synthase (nadh)  ILV2 - acetolactate synthase catalytic subunit  TRP2 - anthranilate synthase trp2  ILV1 - threonine ammonia-lyase ilv1  ARO1 - pentafunctional protein aro1p  CYS4 - cystathionine beta-synthase cys4  ILV6 - acetolactate synthase regulatory subunit |
| GO:0016053 | organic acid biosynthetic process | 2.79E-16 | 5.76E-13 | 5.39 (2325,154,98,35) | [+] Show genes  ARG5,6 - bifunctional acetylglutamate kinase/n-acetyl-gamma-glutamyl-phosphate reductase  IDH1 - isocitrate dehydrogenase (nad(+)) idh1  ARG7 - glutamate n-acetyltransferase  DLD3 - dld3p  ARO2 - bifunctional chorismate synthase/riboflavin reductase [nad(p)h] aro2  ARO8 - bifunctional 2-aminoadipate transaminase/aromatic-amino-acid:2-oxoglutarate transaminase  ILV3 - ilv3p  TRP3 - bifunctional anthranilate synthase/indole-3-glycerol-phosphate synthase  HIS3 - imidazoleglycerol-phosphate dehydratase his3  ASN1 - asparagine synthase (glutamine-hydrolyzing) 1  ILV5 - ketol-acid reductoisomerase  HOM2 - aspartate-semialdehyde dehydrogenase  HOM3 - aspartate kinase  ARO3 - 3-deoxy-7-phosphoheptulonate synthase aro3  IDH2 - isocitrate dehydrogenase (nad(+)) idh2  LEU4 - 2-isopropylmalate synthase leu4  HIS7 - imidazoleglycerol-phosphate synthase  ARG4 - argininosuccinate lyase arg4  MET10 - sulfite reductase subunit alpha  ARO4 - 3-deoxy-7-phosphoheptulonate synthase aro4  CPA2 - cpa2p  ARG1 - argininosuccinate synthase  GLN1 - glutamate--ammonia ligase  ALD4 - aldehyde dehydrogenase (nadp(+)) ald4  HIS4 - trifunctional histidinol dehydrogenase/phosphoribosyl-amp cyclohydrolase/phosphoribosyl-atp diphosphatase  GLT1 - glutamate synthase (nadh)  ILV2 - acetolactate synthase catalytic subunit  TRP2 - anthranilate synthase trp2  BNA5 - kynureninase  ILV1 - threonine ammonia-lyase ilv1  ARO1 - pentafunctional protein aro1p  CYS4 - cystathionine beta-synthase cys4  OLE1 - stearoyl-coa 9-desaturase  ILV6 - acetolactate synthase regulatory subunit  ALD5 - aldehyde dehydrogenase (nad(p)(+)) ald5 |
| GO:0046394 | carboxylic acid biosynthetic process | 2.79E-16 | 3.84E-13 | 5.39 (2325,154,98,35) | [+] Show genes  ARG5,6 - bifunctional acetylglutamate kinase/n-acetyl-gamma-glutamyl-phosphate reductase  IDH1 - isocitrate dehydrogenase (nad(+)) idh1  ARG7 - glutamate n-acetyltransferase  DLD3 - dld3p  ARO2 - bifunctional chorismate synthase/riboflavin reductase [nad(p)h] aro2  ARO8 - bifunctional 2-aminoadipate transaminase/aromatic-amino-acid:2-oxoglutarate transaminase  ILV3 - ilv3p  TRP3 - bifunctional anthranilate synthase/indole-3-glycerol-phosphate synthase  HIS3 - imidazoleglycerol-phosphate dehydratase his3  ASN1 - asparagine synthase (glutamine-hydrolyzing) 1  ILV5 - ketol-acid reductoisomerase  HOM2 - aspartate-semialdehyde dehydrogenase  HOM3 - aspartate kinase  ARO3 - 3-deoxy-7-phosphoheptulonate synthase aro3  IDH2 - isocitrate dehydrogenase (nad(+)) idh2  LEU4 - 2-isopropylmalate synthase leu4  HIS7 - imidazoleglycerol-phosphate synthase  ARG4 - argininosuccinate lyase arg4  MET10 - sulfite reductase subunit alpha  ARO4 - 3-deoxy-7-phosphoheptulonate synthase aro4  CPA2 - cpa2p  GLN1 - glutamate--ammonia ligase  ARG1 - argininosuccinate synthase  ALD4 - aldehyde dehydrogenase (nadp(+)) ald4  HIS4 - trifunctional histidinol dehydrogenase/phosphoribosyl-amp cyclohydrolase/phosphoribosyl-atp diphosphatase  GLT1 - glutamate synthase (nadh)  ILV2 - acetolactate synthase catalytic subunit  TRP2 - anthranilate synthase trp2  BNA5 - kynureninase  ILV1 - threonine ammonia-lyase ilv1  ARO1 - pentafunctional protein aro1p  CYS4 - cystathionine beta-synthase cys4  OLE1 - stearoyl-coa 9-desaturase  ILV6 - acetolactate synthase regulatory subunit  ALD5 - aldehyde dehydrogenase (nad(p)(+)) ald5 |
| GO:0006520 | cellular amino acid metabolic process | 2.84E-16 | 2.94E-13 | 5.21 (2325,164,98,36) | [+] Show genes  ARG5,6 - bifunctional acetylglutamate kinase/n-acetyl-gamma-glutamyl-phosphate reductase  IDH1 - isocitrate dehydrogenase (nad(+)) idh1  ARG7 - glutamate n-acetyltransferase  ARO2 - bifunctional chorismate synthase/riboflavin reductase [nad(p)h] aro2  ARO8 - bifunctional 2-aminoadipate transaminase/aromatic-amino-acid:2-oxoglutarate transaminase  ILV3 - ilv3p  PDC1 - indolepyruvate decarboxylase 1  TRP3 - bifunctional anthranilate synthase/indole-3-glycerol-phosphate synthase  HIS3 - imidazoleglycerol-phosphate dehydratase his3  ASN1 - asparagine synthase (glutamine-hydrolyzing) 1  ILV5 - ketol-acid reductoisomerase  GAD1 - glutamate decarboxylase gad1  HOM2 - aspartate-semialdehyde dehydrogenase  HOM3 - aspartate kinase  ARO3 - 3-deoxy-7-phosphoheptulonate synthase aro3  EHD3 - ehd3p  IDH2 - isocitrate dehydrogenase (nad(+)) idh2  LEU4 - 2-isopropylmalate synthase leu4  HIS7 - imidazoleglycerol-phosphate synthase  ARG4 - argininosuccinate lyase arg4  MET10 - sulfite reductase subunit alpha  YHR020W - proline--trna ligase  ARO4 - 3-deoxy-7-phosphoheptulonate synthase aro4  CPA2 - cpa2p  GLN1 - glutamate--ammonia ligase  ARG1 - argininosuccinate synthase  HIS4 - trifunctional histidinol dehydrogenase/phosphoribosyl-amp cyclohydrolase/phosphoribosyl-atp diphosphatase  GLT1 - glutamate synthase (nadh)  ILV2 - acetolactate synthase catalytic subunit  TRP2 - anthranilate synthase trp2  BNA5 - kynureninase  ILV1 - threonine ammonia-lyase ilv1  ARO1 - pentafunctional protein aro1p  DUR1,2 - bifunctional urea carboxylase/allophanate hydrolase  CYS4 - cystathionine beta-synthase cys4  ILV6 - acetolactate synthase regulatory subunit |
| GO:0019752 | carboxylic acid metabolic process | 2.85E-15 | 2.36E-12 | 4.01 (2325,252,99,43) | [+] Show genes  ARG5,6 - bifunctional acetylglutamate kinase/n-acetyl-gamma-glutamyl-phosphate reductase  IDH1 - isocitrate dehydrogenase (nad(+)) idh1  ARG7 - glutamate n-acetyltransferase  DLD3 - dld3p  ARO2 - bifunctional chorismate synthase/riboflavin reductase [nad(p)h] aro2  ILV3 - ilv3p  ARO8 - bifunctional 2-aminoadipate transaminase/aromatic-amino-acid:2-oxoglutarate transaminase  PDC1 - indolepyruvate decarboxylase 1  TRP3 - bifunctional anthranilate synthase/indole-3-glycerol-phosphate synthase  FAA4 - long-chain fatty acid-coa ligase faa4  HIS3 - imidazoleglycerol-phosphate dehydratase his3  ASN1 - asparagine synthase (glutamine-hydrolyzing) 1  ILV5 - ketol-acid reductoisomerase  GAD1 - glutamate decarboxylase gad1  HOM2 - aspartate-semialdehyde dehydrogenase  HOM3 - aspartate kinase  ARO3 - 3-deoxy-7-phosphoheptulonate synthase aro3  EHD3 - ehd3p  IDH2 - isocitrate dehydrogenase (nad(+)) idh2  LEU4 - 2-isopropylmalate synthase leu4  HIS7 - imidazoleglycerol-phosphate synthase  MET10 - sulfite reductase subunit alpha  ARG4 - argininosuccinate lyase arg4  ARO4 - 3-deoxy-7-phosphoheptulonate synthase aro4  YHR020W - proline--trna ligase  FUM1 - fumarase fum1  CPA2 - cpa2p  ARG1 - argininosuccinate synthase  GLN1 - glutamate--ammonia ligase  ALD4 - aldehyde dehydrogenase (nadp(+)) ald4  HIS4 - trifunctional histidinol dehydrogenase/phosphoribosyl-amp cyclohydrolase/phosphoribosyl-atp diphosphatase  GLT1 - glutamate synthase (nadh)  ILV2 - acetolactate synthase catalytic subunit  TRP2 - anthranilate synthase trp2  DLD1 - dld1p  BNA5 - kynureninase  ILV1 - threonine ammonia-lyase ilv1  ARO1 - pentafunctional protein aro1p  DUR1,2 - bifunctional urea carboxylase/allophanate hydrolase  CYS4 - cystathionine beta-synthase cys4  OLE1 - stearoyl-coa 9-desaturase  ILV6 - acetolactate synthase regulatory subunit  ALD5 - aldehyde dehydrogenase (nad(p)(+)) ald5 |
| GO:1901607 | alpha-amino acid biosynthetic process | 9.92E-15 | 6.83E-12 | 7.01 (2325,88,98,26) | [+] Show genes  IDH1 - isocitrate dehydrogenase (nad(+)) idh1  ARG5,6 - bifunctional acetylglutamate kinase/n-acetyl-gamma-glutamyl-phosphate reductase  ARG7 - glutamate n-acetyltransferase  ILV3 - ilv3p  ARO8 - bifunctional 2-aminoadipate transaminase/aromatic-amino-acid:2-oxoglutarate transaminase  TRP3 - bifunctional anthranilate synthase/indole-3-glycerol-phosphate synthase  HIS3 - imidazoleglycerol-phosphate dehydratase his3  ILV5 - ketol-acid reductoisomerase  ASN1 - asparagine synthase (glutamine-hydrolyzing) 1  HOM2 - aspartate-semialdehyde dehydrogenase  HOM3 - aspartate kinase  IDH2 - isocitrate dehydrogenase (nad(+)) idh2  LEU4 - 2-isopropylmalate synthase leu4  HIS7 - imidazoleglycerol-phosphate synthase  MET10 - sulfite reductase subunit alpha  ARG4 - argininosuccinate lyase arg4  CPA2 - cpa2p  GLN1 - glutamate--ammonia ligase  ARG1 - argininosuccinate synthase  HIS4 - trifunctional histidinol dehydrogenase/phosphoribosyl-amp cyclohydrolase/phosphoribosyl-atp diphosphatase  ILV2 - acetolactate synthase catalytic subunit  GLT1 - glutamate synthase (nadh)  TRP2 - anthranilate synthase trp2  ILV1 - threonine ammonia-lyase ilv1  CYS4 - cystathionine beta-synthase cys4  ILV6 - acetolactate synthase regulatory subunit |
| GO:0043436 | oxoacid metabolic process | 1.93E-14 | 1.14E-11 | 3.83 (2325,264,99,43) | [+] Show genes  ARG5,6 - bifunctional acetylglutamate kinase/n-acetyl-gamma-glutamyl-phosphate reductase  IDH1 - isocitrate dehydrogenase (nad(+)) idh1  ARG7 - glutamate n-acetyltransferase  DLD3 - dld3p  ARO2 - bifunctional chorismate synthase/riboflavin reductase [nad(p)h] aro2  ILV3 - ilv3p  ARO8 - bifunctional 2-aminoadipate transaminase/aromatic-amino-acid:2-oxoglutarate transaminase  PDC1 - indolepyruvate decarboxylase 1  TRP3 - bifunctional anthranilate synthase/indole-3-glycerol-phosphate synthase  HIS3 - imidazoleglycerol-phosphate dehydratase his3  FAA4 - long-chain fatty acid-coa ligase faa4  ASN1 - asparagine synthase (glutamine-hydrolyzing) 1  ILV5 - ketol-acid reductoisomerase  GAD1 - glutamate decarboxylase gad1  HOM2 - aspartate-semialdehyde dehydrogenase  HOM3 - aspartate kinase  ARO3 - 3-deoxy-7-phosphoheptulonate synthase aro3  EHD3 - ehd3p  IDH2 - isocitrate dehydrogenase (nad(+)) idh2  LEU4 - 2-isopropylmalate synthase leu4  HIS7 - imidazoleglycerol-phosphate synthase  MET10 - sulfite reductase subunit alpha  ARG4 - argininosuccinate lyase arg4  ARO4 - 3-deoxy-7-phosphoheptulonate synthase aro4  YHR020W - proline--trna ligase  FUM1 - fumarase fum1  CPA2 - cpa2p  GLN1 - glutamate--ammonia ligase  ARG1 - argininosuccinate synthase  ALD4 - aldehyde dehydrogenase (nadp(+)) ald4  HIS4 - trifunctional histidinol dehydrogenase/phosphoribosyl-amp cyclohydrolase/phosphoribosyl-atp diphosphatase  GLT1 - glutamate synthase (nadh)  ILV2 - acetolactate synthase catalytic subunit  TRP2 - anthranilate synthase trp2  DLD1 - dld1p  BNA5 - kynureninase  ILV1 - threonine ammonia-lyase ilv1  ARO1 - pentafunctional protein aro1p  DUR1,2 - bifunctional urea carboxylase/allophanate hydrolase  CYS4 - cystathionine beta-synthase cys4  OLE1 - stearoyl-coa 9-desaturase  ILV6 - acetolactate synthase regulatory subunit  ALD5 - aldehyde dehydrogenase (nad(p)(+)) ald5 |
| GO:0006082 | organic acid metabolic process | 2.25E-14 | 1.16E-11 | 3.81 (2325,265,99,43) | [+] Show genes  ARG5,6 - bifunctional acetylglutamate kinase/n-acetyl-gamma-glutamyl-phosphate reductase  IDH1 - isocitrate dehydrogenase (nad(+)) idh1  ARG7 - glutamate n-acetyltransferase  DLD3 - dld3p  ARO2 - bifunctional chorismate synthase/riboflavin reductase [nad(p)h] aro2  ILV3 - ilv3p  ARO8 - bifunctional 2-aminoadipate transaminase/aromatic-amino-acid:2-oxoglutarate transaminase  PDC1 - indolepyruvate decarboxylase 1  TRP3 - bifunctional anthranilate synthase/indole-3-glycerol-phosphate synthase  FAA4 - long-chain fatty acid-coa ligase faa4  HIS3 - imidazoleglycerol-phosphate dehydratase his3  ASN1 - asparagine synthase (glutamine-hydrolyzing) 1  ILV5 - ketol-acid reductoisomerase  GAD1 - glutamate decarboxylase gad1  HOM2 - aspartate-semialdehyde dehydrogenase  HOM3 - aspartate kinase  ARO3 - 3-deoxy-7-phosphoheptulonate synthase aro3  EHD3 - ehd3p  IDH2 - isocitrate dehydrogenase (nad(+)) idh2  LEU4 - 2-isopropylmalate synthase leu4  HIS7 - imidazoleglycerol-phosphate synthase  MET10 - sulfite reductase subunit alpha  ARG4 - argininosuccinate lyase arg4  ARO4 - 3-deoxy-7-phosphoheptulonate synthase aro4  YHR020W - proline--trna ligase  FUM1 - fumarase fum1  CPA2 - cpa2p  GLN1 - glutamate--ammonia ligase  ARG1 - argininosuccinate synthase  ALD4 - aldehyde dehydrogenase (nadp(+)) ald4  HIS4 - trifunctional histidinol dehydrogenase/phosphoribosyl-amp cyclohydrolase/phosphoribosyl-atp diphosphatase  GLT1 - glutamate synthase (nadh)  ILV2 - acetolactate synthase catalytic subunit  TRP2 - anthranilate synthase trp2  DLD1 - dld1p  BNA5 - kynureninase  ILV1 - threonine ammonia-lyase ilv1  ARO1 - pentafunctional protein aro1p  DUR1,2 - bifunctional urea carboxylase/allophanate hydrolase  CYS4 - cystathionine beta-synthase cys4  OLE1 - stearoyl-coa 9-desaturase  ILV6 - acetolactate synthase regulatory subunit  ALD5 - aldehyde dehydrogenase (nad(p)(+)) ald5 |
| GO:1901605 | alpha-amino acid metabolic process | 3.94E-14 | 1.81E-11 | 5.60 (2325,127,98,30) | [+] Show genes  ARG5,6 - bifunctional acetylglutamate kinase/n-acetyl-gamma-glutamyl-phosphate reductase  IDH1 - isocitrate dehydrogenase (nad(+)) idh1  ARG7 - glutamate n-acetyltransferase  ARO8 - bifunctional 2-aminoadipate transaminase/aromatic-amino-acid:2-oxoglutarate transaminase  ILV3 - ilv3p  PDC1 - indolepyruvate decarboxylase 1  TRP3 - bifunctional anthranilate synthase/indole-3-glycerol-phosphate synthase  HIS3 - imidazoleglycerol-phosphate dehydratase his3  ASN1 - asparagine synthase (glutamine-hydrolyzing) 1  ILV5 - ketol-acid reductoisomerase  GAD1 - glutamate decarboxylase gad1  HOM2 - aspartate-semialdehyde dehydrogenase  HOM3 - aspartate kinase  IDH2 - isocitrate dehydrogenase (nad(+)) idh2  LEU4 - 2-isopropylmalate synthase leu4  HIS7 - imidazoleglycerol-phosphate synthase  ARG4 - argininosuccinate lyase arg4  MET10 - sulfite reductase subunit alpha  CPA2 - cpa2p  GLN1 - glutamate--ammonia ligase  ARG1 - argininosuccinate synthase  HIS4 - trifunctional histidinol dehydrogenase/phosphoribosyl-amp cyclohydrolase/phosphoribosyl-atp diphosphatase  GLT1 - glutamate synthase (nadh)  ILV2 - acetolactate synthase catalytic subunit  TRP2 - anthranilate synthase trp2  BNA5 - kynureninase  ILV1 - threonine ammonia-lyase ilv1  CYS4 - cystathionine beta-synthase cys4  DUR1,2 - bifunctional urea carboxylase/allophanate hydrolase  ILV6 - acetolactate synthase regulatory subunit |
| GO:0044281 | small molecule metabolic process | 4.06E-14 | 1.68E-11 | 2.93 (2325,441,99,55) | [+] Show genes  IDH1 - isocitrate dehydrogenase (nad(+)) idh1  ARO2 - bifunctional chorismate synthase/riboflavin reductase [nad(p)h] aro2  ILV3 - ilv3p  PDC1 - indolepyruvate decarboxylase 1  FAA4 - long-chain fatty acid-coa ligase faa4  HIS3 - imidazoleglycerol-phosphate dehydratase his3  ILV5 - ketol-acid reductoisomerase  GAD1 - glutamate decarboxylase gad1  YSA1 - ysa1p  HIS7 - imidazoleglycerol-phosphate synthase  MET10 - sulfite reductase subunit alpha  ADE16 - bifunctional phosphoribosylaminoimidazolecarboxamide formyltransferase/imp cyclohydrolase ade16  ARO4 - 3-deoxy-7-phosphoheptulonate synthase aro4  FUM1 - fumarase fum1  CPA2 - cpa2p  PGM2 - phosphoglucomutase pgm2  GLN1 - glutamate--ammonia ligase  ARG1 - argininosuccinate synthase  HIS4 - trifunctional histidinol dehydrogenase/phosphoribosyl-amp cyclohydrolase/phosphoribosyl-atp diphosphatase  ILV2 - acetolactate synthase catalytic subunit  ARO1 - pentafunctional protein aro1p  DUR1,2 - bifunctional urea carboxylase/allophanate hydrolase  CYS4 - cystathionine beta-synthase cys4  ILV6 - acetolactate synthase regulatory subunit  ARG5,6 - bifunctional acetylglutamate kinase/n-acetyl-gamma-glutamyl-phosphate reductase  CAB2 - phosphopantothenate--cysteine ligase cab2  PGM1 - phosphoglucomutase pgm1  ARG7 - glutamate n-acetyltransferase  DLD3 - dld3p  YDL124W - aldo-keto reductase superfamily protein  ARO8 - bifunctional 2-aminoadipate transaminase/aromatic-amino-acid:2-oxoglutarate transaminase  TRP3 - bifunctional anthranilate synthase/indole-3-glycerol-phosphate synthase  ASN1 - asparagine synthase (glutamine-hydrolyzing) 1  GUK1 - guanylate kinase  HOM2 - aspartate-semialdehyde dehydrogenase  HOM3 - aspartate kinase  ZWF1 - glucose-6-phosphate dehydrogenase  ARO3 - 3-deoxy-7-phosphoheptulonate synthase aro3  EHD3 - ehd3p  IDH2 - isocitrate dehydrogenase (nad(+)) idh2  LEU4 - 2-isopropylmalate synthase leu4  ARG4 - argininosuccinate lyase arg4  YHR020W - proline--trna ligase  APA2 - apa2p  ALD4 - aldehyde dehydrogenase (nadp(+)) ald4  GLT1 - glutamate synthase (nadh)  TRP2 - anthranilate synthase trp2  BDH1 - (r,r)-butanediol dehydrogenase  DLD1 - dld1p  BNA5 - kynureninase  ILV1 - threonine ammonia-lyase ilv1  INP53 - phosphatidylinositol-3-/phosphoinositide 5-phosphatase inp53  PGM3 - phosphoglucomutase pgm3  OLE1 - stearoyl-coa 9-desaturase  ALD5 - aldehyde dehydrogenase (nad(p)(+)) ald5 |
| GO:0044283 | small molecule biosynthetic process | 1.68E-13 | 6.3E-11 | 4.01 (2325,231,98,39) | [+] Show genes  ARG5,6 - bifunctional acetylglutamate kinase/n-acetyl-gamma-glutamyl-phosphate reductase  IDH1 - isocitrate dehydrogenase (nad(+)) idh1  ARG7 - glutamate n-acetyltransferase  DLD3 - dld3p  ARO2 - bifunctional chorismate synthase/riboflavin reductase [nad(p)h] aro2  ILV3 - ilv3p  ARO8 - bifunctional 2-aminoadipate transaminase/aromatic-amino-acid:2-oxoglutarate transaminase  PDC1 - indolepyruvate decarboxylase 1  TRP3 - bifunctional anthranilate synthase/indole-3-glycerol-phosphate synthase  HIS3 - imidazoleglycerol-phosphate dehydratase his3  ASN1 - asparagine synthase (glutamine-hydrolyzing) 1  ILV5 - ketol-acid reductoisomerase  GUK1 - guanylate kinase  HOM2 - aspartate-semialdehyde dehydrogenase  HOM3 - aspartate kinase  ARO3 - 3-deoxy-7-phosphoheptulonate synthase aro3  IDH2 - isocitrate dehydrogenase (nad(+)) idh2  LEU4 - 2-isopropylmalate synthase leu4  HIS7 - imidazoleglycerol-phosphate synthase  ARG4 - argininosuccinate lyase arg4  MET10 - sulfite reductase subunit alpha  ARO4 - 3-deoxy-7-phosphoheptulonate synthase aro4  CPA2 - cpa2p  GLN1 - glutamate--ammonia ligase  ARG1 - argininosuccinate synthase  ALD4 - aldehyde dehydrogenase (nadp(+)) ald4  HIS4 - trifunctional histidinol dehydrogenase/phosphoribosyl-amp cyclohydrolase/phosphoribosyl-atp diphosphatase  GLT1 - glutamate synthase (nadh)  ILV2 - acetolactate synthase catalytic subunit  BDH1 - (r,r)-butanediol dehydrogenase  TRP2 - anthranilate synthase trp2  BNA5 - kynureninase  ILV1 - threonine ammonia-lyase ilv1  ARO1 - pentafunctional protein aro1p  CYS4 - cystathionine beta-synthase cys4  OLE1 - stearoyl-coa 9-desaturase  PGM3 - phosphoglucomutase pgm3  ILV6 - acetolactate synthase regulatory subunit  ALD5 - aldehyde dehydrogenase (nad(p)(+)) ald5 |
| GO:1901576 | organic substance biosynthetic process | 4.37E-11 | 1.51E-8 | 1.45 (2325,756,438,206) | [+] Show genes  TFA2 - tfa2p  IDH1 - isocitrate dehydrogenase (nad(+)) idh1  GSH1 - gsh1p  ARO2 - bifunctional chorismate synthase/riboflavin reductase [nad(p)h] aro2  RPL9A - ribosomal 60s subunit protein l9a  RPL8A - ribosomal 60s subunit protein l8a  ASC1 - asc1p  APA1 - apa1p  GLK1 - glucokinase  PRS2 - ribose phosphate diphosphokinase subunit prs2  EMI2 - putative glucokinase  PGM2 - phosphoglucomutase pgm2  HIS4 - trifunctional histidinol dehydrogenase/phosphoribosyl-amp cyclohydrolase/phosphoribosyl-atp diphosphatase  CIT1 - citrate (si)-synthase cit1  ILV2 - acetolactate synthase catalytic subunit  SPT15 - spt15p  RPS2 - ribosomal 40s subunit protein s2  CPA1 - carbamoyl-phosphate synthase (glutamine-hydrolyzing) cpa1  ILV6 - acetolactate synthase regulatory subunit  ARG5,6 - bifunctional acetylglutamate kinase/n-acetyl-gamma-glutamyl-phosphate reductase  KRE6 - kre6p  GPI17 - gpi17p  RPB9 - rpb9p  ABZ1 - 4-amino-4-deoxychorismate synthase  ASN1 - asparagine synthase (glutamine-hydrolyzing) 1  HIS1 - atp phosphoribosyltransferase  YSH1 - ysh1p  GUK1 - guanylate kinase  HOM3 - aspartate kinase  MCM5 - mcm5p  LRO1 - phospholipid:diacylglycerol acyltransferase  RSM18 - mitochondrial 37s ribosomal protein rsm18  TDH1 - tdh1p  LAC1 - sphingosine n-acyltransferase lac1  RPB7 - rpb7p  RPL31A - ribosomal 60s subunit protein l31a  TRP2 - anthranilate synthase trp2  BNA5 - kynureninase  SUB2 - sub2p  ALD2 - aldehyde dehydrogenase (nad(+)) ald2  ILV1 - threonine ammonia-lyase ilv1  TIF4632 - tif4632p  MRI1 - s-methyl-5-thioribose-1-phosphate isomerase mri1  MRPS12 - putative mitochondrial 37s ribosomal protein mrps12  RPL33B - ribosomal 60s subunit protein l33b  OLE1 - stearoyl-coa 9-desaturase  MVD1 - diphosphomevalonate decarboxylase mvd1  ALD5 - aldehyde dehydrogenase (nad(p)(+)) ald5  LYS9 - saccharopine dehydrogenase (nadp+, l-glutamate-forming)  YDR089W - hypothetical protein  TPS3 - tps3p  HIS3 - imidazoleglycerol-phosphate dehydratase his3  RKR1 - ubiquitin-protein ligase rkr1  ILV5 - ketol-acid reductoisomerase  ENO1 - phosphopyruvate hydratase eno1  FOL2 - gtp cyclohydrolase i  RPL26A - ribosomal 60s subunit protein l26a  COQ1 - trans-hexaprenyltranstransferase  RPP0 - ribosomal protein p0  TUF1 - tuf1p  DPB4 - dpb4p  SAM4 - sam4p  GLN4 - glutamine--trna ligase  ARO1 - pentafunctional protein aro1p  HEK2 - hek2p  FAS2 - trifunctional fatty acid synthase subunit fas2  NCP1 - ncp1p  PGM1 - phosphoglucomutase pgm1  RPL17B - rpl17bp  RPL7B - ribosomal 60s subunit protein l7b  RPL32 - ribosomal 60s subunit protein l32  ARO8 - bifunctional 2-aminoadipate transaminase/aromatic-amino-acid:2-oxoglutarate transaminase  FLC1 - flc1p  GLC8 - glc8p  ARO3 - 3-deoxy-7-phosphoheptulonate synthase aro3  EHD3 - ehd3p  GON7 - gon7p  KRS1 - lysine--trna ligase krs1  IDH2 - isocitrate dehydrogenase (nad(+)) idh2  ARG4 - argininosuccinate lyase arg4  YHR020W - proline--trna ligase  HSP31 - hsp31p  HEM13 - coproporphyrinogen oxidase  LCB1 - serine c-palmitoyltransferase lcb1  APA2 - apa2p  RPL14B - ribosomal 60s subunit protein l14b  LEO1 - leo1p  CAB1 - pantothenate kinase  LCB2 - serine c-palmitoyltransferase lcb2  POS5 - pos5p  INP53 - phosphatidylinositol-3-/phosphoinositide 5-phosphatase inp53  TFC7 - tfc7p  RPS13 - ribosomal 40s subunit protein s13  PGM3 - phosphoglucomutase pgm3  RPS7A - ribosomal 40s subunit protein s7a  RPL29 - ribosomal 60s subunit protein l29  ENO2 - phosphopyruvate hydratase eno2  TSL1 - tsl1p  PCF11 - pcf11p  ALD6 - aldehyde dehydrogenase (nadp(+)) ald6  ASN2 - asparagine synthase (glutamine-hydrolyzing) 2  ERG9 - bifunctional farnesyl-diphosphate farnesyltransferase/squalene synthase  SPT6 - spt6p  GCD6 - gcd6p  MET10 - sulfite reductase subunit alpha  RSM7 - rsm7p  NHP6B - nhp6bp  POL30 - pol30p  RPL3 - ribosomal 60s subunit protein l3  RFC3 - replication factor c subunit 3  CPA2 - cpa2p  GPI13 - gpi13p  RSM26 - rsm26p  YNL284C-B - gag-pol fusion protein  RPL6B - ribosomal 60s subunit protein l6b  CYS4 - cystathionine beta-synthase cys4  PTI1 - pti1p  RPS5 - rps5p  CAR2 - ornithine-oxo-acid transaminase  THI4 - thi4p  BAT1 - branched-chain-amino-acid transaminase bat1  RPS1B - ribosomal 40s subunit protein s1b  CAB2 - phosphopantothenate--cysteine ligase cab2  SER33 - phosphoglycerate dehydrogenase ser33  TRP3 - bifunctional anthranilate synthase/indole-3-glycerol-phosphate synthase  RPA49 - rpa49p  FUR1 - uracil phosphoribosyltransferase  RNR4 - ribonucleotide-diphosphate reductase subunit rnr4  HOM2 - aspartate-semialdehyde dehydrogenase  DPS1 - aspartate--trna ligase dps1  ERG1 - squalene monooxygenase  PDR16 - pdr16p  TOP1 - dna topoisomerase 1  RPL6A - ribosomal 60s subunit protein l6a  PDX3 - pyridoxamine-phosphate oxidase pdx3  SSU72 - ssu72p  ARO9 - aromatic-amino-acid:2-oxoglutarate transaminase  BDH1 - (r,r)-butanediol dehydrogenase  CDC60 - leucine--trna ligase cdc60  RPL5 - ribosomal 60s subunit protein l5  HMO1 - hmo1p  COQ5 - 2-hexaprenyl-6-methoxy-1,4-benzoquinone methyltransferase  KES1 - kes1p  SER2 - phosphoserine phosphatase  FAS1 - tetrafunctional fatty acid synthase subunit fas1  VAN1 - van1p  RPL33A - ribosomal 60s subunit protein l33a  MRPL6 - mitochondrial 54s ribosomal protein yml16  RPL24A - ribosomal 60s subunit protein l24a  RPS3 - ribosomal 40s subunit protein s3  GLC3 - 1,4-alpha-glucan branching enzyme  RPL30 - ribosomal 60s subunit protein l30  MRPL19 - mitochondrial 54s ribosomal protein yml19  CWH41 - cwh41p  POL31 - pol31p  ILV3 - ilv3p  PDC1 - indolepyruvate decarboxylase 1  CHS1 - chitin synthase chs1  TRP4 - anthranilate phosphoribosyltransferase  STH1 - sth1p  AYR1 - acylglycerone-phosphate reductase  LEU1 - 3-isopropylmalate dehydratase leu1  HIS7 - imidazoleglycerol-phosphate synthase  RPL15A - ribosomal 60s subunit protein l15a  ADE16 - bifunctional phosphoribosylaminoimidazolecarboxamide formyltransferase/imp cyclohydrolase ade16  ARO4 - 3-deoxy-7-phosphoheptulonate synthase aro4  HEM3 - hydroxymethylbilane synthase  ISC1 - inositol phosphosphingolipid phospholipase  HTS1 - histidine--trna ligase  PRO3 - pyrroline-5-carboxylate reductase  ARG1 - argininosuccinate synthase  GLN1 - glutamate--ammonia ligase  PRS5 - ribose phosphate diphosphokinase subunit prs5  RPL10 - ribosomal 60s subunit protein l10  AMD1 - amp deaminase  YMC1 - ymc1p  NRK1 - ribosylnicotinamide kinase  RPP2A - ribosomal protein p2a  NHP6A - nhp6ap  MEF1 - mef1p  ERG3 - c-5 sterol desaturase  FCY1 - cytosine deaminase  RPL25 - ribosomal 60s subunit protein l25  ARG7 - glutamate n-acetyltransferase  DLD3 - dld3p  RPL26B - ribosomal 60s subunit protein l26b  ERG27 - 3-keto-steroid reductase  YGR054W - hypothetical protein  GRE2 - methylglyoxal reductase (nadph-dependent) gre2  RCO1 - rco1p  RPO21 - rpo21p  LEU4 - 2-isopropylmalate synthase leu4  POL1 - pol1p  EHT1 - eht1p  ARP9 - arp9p  UTR4 - putative acireductone synthase utr4  STM1 - stm1p  ALD4 - aldehyde dehydrogenase (nadp(+)) ald4  MET12 - methylenetetrahydrofolate reductase (nad(p)h) met12  GLT1 - glutamate synthase (nadh)  MCM3 - mcm3p  PDC5 - indolepyruvate decarboxylase 5  ARA2 - d-arabinose 1-dehydrogenase (nad(p)(+)) ara2  SSB1 - hsp70 family atpase ssb1  RPL16B - ribosomal 60s subunit protein l16b  SUB1 - sub1p |
| GO:0044249 | cellular biosynthetic process | 5.38E-11 | 1.71E-8 | 1.46 (2325,733,438,201) | [+] Show genes  TFA2 - tfa2p  IDH1 - isocitrate dehydrogenase (nad(+)) idh1  GSH1 - gsh1p  ARO2 - bifunctional chorismate synthase/riboflavin reductase [nad(p)h] aro2  RPL9A - ribosomal 60s subunit protein l9a  RPL8A - ribosomal 60s subunit protein l8a  ASC1 - asc1p  APA1 - apa1p  GLK1 - glucokinase  PRS2 - ribose phosphate diphosphokinase subunit prs2  EMI2 - putative glucokinase  PGM2 - phosphoglucomutase pgm2  HIS4 - trifunctional histidinol dehydrogenase/phosphoribosyl-amp cyclohydrolase/phosphoribosyl-atp diphosphatase  CIT1 - citrate (si)-synthase cit1  ILV2 - acetolactate synthase catalytic subunit  SPT15 - spt15p  RPS2 - ribosomal 40s subunit protein s2  CPA1 - carbamoyl-phosphate synthase (glutamine-hydrolyzing) cpa1  ILV6 - acetolactate synthase regulatory subunit  ARG5,6 - bifunctional acetylglutamate kinase/n-acetyl-gamma-glutamyl-phosphate reductase  KRE6 - kre6p  GPI17 - gpi17p  RPB9 - rpb9p  ABZ1 - 4-amino-4-deoxychorismate synthase  ASN1 - asparagine synthase (glutamine-hydrolyzing) 1  HIS1 - atp phosphoribosyltransferase  YSH1 - ysh1p  GUK1 - guanylate kinase  HOM3 - aspartate kinase  MCM5 - mcm5p  LRO1 - phospholipid:diacylglycerol acyltransferase  RSM18 - mitochondrial 37s ribosomal protein rsm18  TDH1 - tdh1p  LAC1 - sphingosine n-acyltransferase lac1  RPB7 - rpb7p  RPL31A - ribosomal 60s subunit protein l31a  TRP2 - anthranilate synthase trp2  BNA5 - kynureninase  SUB2 - sub2p  ALD2 - aldehyde dehydrogenase (nad(+)) ald2  ILV1 - threonine ammonia-lyase ilv1  TIF4632 - tif4632p  MRI1 - s-methyl-5-thioribose-1-phosphate isomerase mri1  MRPS12 - putative mitochondrial 37s ribosomal protein mrps12  RPL33B - ribosomal 60s subunit protein l33b  OLE1 - stearoyl-coa 9-desaturase  MVD1 - diphosphomevalonate decarboxylase mvd1  LYS9 - saccharopine dehydrogenase (nadp+, l-glutamate-forming)  ALD5 - aldehyde dehydrogenase (nad(p)(+)) ald5  YDR089W - hypothetical protein  TPS3 - tps3p  HIS3 - imidazoleglycerol-phosphate dehydratase his3  RKR1 - ubiquitin-protein ligase rkr1  ILV5 - ketol-acid reductoisomerase  ENO1 - phosphopyruvate hydratase eno1  FOL2 - gtp cyclohydrolase i  RPL26A - ribosomal 60s subunit protein l26a  COQ1 - trans-hexaprenyltranstransferase  RPP0 - ribosomal protein p0  TUF1 - tuf1p  DPB4 - dpb4p  SAM4 - sam4p  GLN4 - glutamine--trna ligase  ARO1 - pentafunctional protein aro1p  HEK2 - hek2p  FAS2 - trifunctional fatty acid synthase subunit fas2  NCP1 - ncp1p  PGM1 - phosphoglucomutase pgm1  RPL17B - rpl17bp  RPL7B - ribosomal 60s subunit protein l7b  RPL32 - ribosomal 60s subunit protein l32  ARO8 - bifunctional 2-aminoadipate transaminase/aromatic-amino-acid:2-oxoglutarate transaminase  FLC1 - flc1p  GLC8 - glc8p  ARO3 - 3-deoxy-7-phosphoheptulonate synthase aro3  EHD3 - ehd3p  GON7 - gon7p  KRS1 - lysine--trna ligase krs1  IDH2 - isocitrate dehydrogenase (nad(+)) idh2  ARG4 - argininosuccinate lyase arg4  YHR020W - proline--trna ligase  HSP31 - hsp31p  HEM13 - coproporphyrinogen oxidase  LCB1 - serine c-palmitoyltransferase lcb1  APA2 - apa2p  RPL14B - ribosomal 60s subunit protein l14b  LEO1 - leo1p  CAB1 - pantothenate kinase  LCB2 - serine c-palmitoyltransferase lcb2  POS5 - pos5p  INP53 - phosphatidylinositol-3-/phosphoinositide 5-phosphatase inp53  TFC7 - tfc7p  RPS13 - ribosomal 40s subunit protein s13  PGM3 - phosphoglucomutase pgm3  RPS7A - ribosomal 40s subunit protein s7a  RPL29 - ribosomal 60s subunit protein l29  ENO2 - phosphopyruvate hydratase eno2  TSL1 - tsl1p  PCF11 - pcf11p  ALD6 - aldehyde dehydrogenase (nadp(+)) ald6  ASN2 - asparagine synthase (glutamine-hydrolyzing) 2  ERG9 - bifunctional farnesyl-diphosphate farnesyltransferase/squalene synthase  GCD6 - gcd6p  SPT6 - spt6p  MET10 - sulfite reductase subunit alpha  RSM7 - rsm7p  NHP6B - nhp6bp  POL30 - pol30p  RPL3 - ribosomal 60s subunit protein l3  RFC3 - replication factor c subunit 3  CPA2 - cpa2p  GPI13 - gpi13p  RSM26 - rsm26p  RPL6B - ribosomal 60s subunit protein l6b  YNL284C-B - gag-pol fusion protein  CYS4 - cystathionine beta-synthase cys4  PTI1 - pti1p  RPS5 - rps5p  CAR2 - ornithine-oxo-acid transaminase  THI4 - thi4p  BAT1 - branched-chain-amino-acid transaminase bat1  RPS1B - ribosomal 40s subunit protein s1b  CAB2 - phosphopantothenate--cysteine ligase cab2  SER33 - phosphoglycerate dehydrogenase ser33  TRP3 - bifunctional anthranilate synthase/indole-3-glycerol-phosphate synthase  RPA49 - rpa49p  FUR1 - uracil phosphoribosyltransferase  RNR4 - ribonucleotide-diphosphate reductase subunit rnr4  HOM2 - aspartate-semialdehyde dehydrogenase  DPS1 - aspartate--trna ligase dps1  ERG1 - squalene monooxygenase  PDR16 - pdr16p  TOP1 - dna topoisomerase 1  RPL6A - ribosomal 60s subunit protein l6a  PDX3 - pyridoxamine-phosphate oxidase pdx3  SSU72 - ssu72p  ARO9 - aromatic-amino-acid:2-oxoglutarate transaminase  DPH1 - dph1p  CDC60 - leucine--trna ligase cdc60  RPL5 - ribosomal 60s subunit protein l5  HMO1 - hmo1p  COQ5 - 2-hexaprenyl-6-methoxy-1,4-benzoquinone methyltransferase  SER2 - phosphoserine phosphatase  FAS1 - tetrafunctional fatty acid synthase subunit fas1  VAN1 - van1p  RPL33A - ribosomal 60s subunit protein l33a  MRPL6 - mitochondrial 54s ribosomal protein yml16  RPL24A - ribosomal 60s subunit protein l24a  RPS3 - ribosomal 40s subunit protein s3  GLC3 - 1,4-alpha-glucan branching enzyme  RPL30 - ribosomal 60s subunit protein l30  MRPL19 - mitochondrial 54s ribosomal protein yml19  CWH41 - cwh41p  POL31 - pol31p  ILV3 - ilv3p  TRP4 - anthranilate phosphoribosyltransferase  STH1 - sth1p  AYR1 - acylglycerone-phosphate reductase  LEU1 - 3-isopropylmalate dehydratase leu1  HIS7 - imidazoleglycerol-phosphate synthase  RPL15A - ribosomal 60s subunit protein l15a  ADE16 - bifunctional phosphoribosylaminoimidazolecarboxamide formyltransferase/imp cyclohydrolase ade16  ARO4 - 3-deoxy-7-phosphoheptulonate synthase aro4  ISC1 - inositol phosphosphingolipid phospholipase  HEM3 - hydroxymethylbilane synthase  HTS1 - histidine--trna ligase  PRO3 - pyrroline-5-carboxylate reductase  ARG1 - argininosuccinate synthase  GLN1 - glutamate--ammonia ligase  PRS5 - ribose phosphate diphosphokinase subunit prs5  RPL10 - ribosomal 60s subunit protein l10  AMD1 - amp deaminase  YMC1 - ymc1p  NRK1 - ribosylnicotinamide kinase  RPP2A - ribosomal protein p2a  NHP6A - nhp6ap  MEF1 - mef1p  ERG3 - c-5 sterol desaturase  FCY1 - cytosine deaminase  RPL25 - ribosomal 60s subunit protein l25  ARG7 - glutamate n-acetyltransferase  RPL26B - ribosomal 60s subunit protein l26b  DLD3 - dld3p  ERG27 - 3-keto-steroid reductase  YGR054W - hypothetical protein  RCO1 - rco1p  RPO21 - rpo21p  LEU4 - 2-isopropylmalate synthase leu4  POL1 - pol1p  EHT1 - eht1p  ARP9 - arp9p  UTR4 - putative acireductone synthase utr4  STM1 - stm1p  ALD4 - aldehyde dehydrogenase (nadp(+)) ald4  MET12 - methylenetetrahydrofolate reductase (nad(p)h) met12  GLT1 - glutamate synthase (nadh)  MCM3 - mcm3p  ARA2 - d-arabinose 1-dehydrogenase (nad(p)(+)) ara2  SSB1 - hsp70 family atpase ssb1  RPL16B - ribosomal 60s subunit protein l16b  SUB1 - sub1p |
| GO:0009058 | biosynthetic process | 8.61E-11 | 2.54E-8 | 1.44 (2325,765,438,207) | [+] Show genes  TFA2 - tfa2p  IDH1 - isocitrate dehydrogenase (nad(+)) idh1  GSH1 - gsh1p  ARO2 - bifunctional chorismate synthase/riboflavin reductase [nad(p)h] aro2  RPL9A - ribosomal 60s subunit protein l9a  RPL8A - ribosomal 60s subunit protein l8a  APA1 - apa1p  ASC1 - asc1p  GLK1 - glucokinase  PRS2 - ribose phosphate diphosphokinase subunit prs2  EMI2 - putative glucokinase  PGM2 - phosphoglucomutase pgm2  HIS4 - trifunctional histidinol dehydrogenase/phosphoribosyl-amp cyclohydrolase/phosphoribosyl-atp diphosphatase  CIT1 - citrate (si)-synthase cit1  ILV2 - acetolactate synthase catalytic subunit  SPT15 - spt15p  RPS2 - ribosomal 40s subunit protein s2  CPA1 - carbamoyl-phosphate synthase (glutamine-hydrolyzing) cpa1  ILV6 - acetolactate synthase regulatory subunit  ARG5,6 - bifunctional acetylglutamate kinase/n-acetyl-gamma-glutamyl-phosphate reductase  KRE6 - kre6p  GPI17 - gpi17p  RPB9 - rpb9p  ABZ1 - 4-amino-4-deoxychorismate synthase  ASN1 - asparagine synthase (glutamine-hydrolyzing) 1  HIS1 - atp phosphoribosyltransferase  GUK1 - guanylate kinase  YSH1 - ysh1p  HOM3 - aspartate kinase  LRO1 - phospholipid:diacylglycerol acyltransferase  MCM5 - mcm5p  RSM18 - mitochondrial 37s ribosomal protein rsm18  TDH1 - tdh1p  LAC1 - sphingosine n-acyltransferase lac1  RPB7 - rpb7p  RPL31A - ribosomal 60s subunit protein l31a  TRP2 - anthranilate synthase trp2  BNA5 - kynureninase  SUB2 - sub2p  ALD2 - aldehyde dehydrogenase (nad(+)) ald2  ILV1 - threonine ammonia-lyase ilv1  TIF4632 - tif4632p  MRI1 - s-methyl-5-thioribose-1-phosphate isomerase mri1  MRPS12 - putative mitochondrial 37s ribosomal protein mrps12  RPL33B - ribosomal 60s subunit protein l33b  OLE1 - stearoyl-coa 9-desaturase  MVD1 - diphosphomevalonate decarboxylase mvd1  LYS9 - saccharopine dehydrogenase (nadp+, l-glutamate-forming)  ALD5 - aldehyde dehydrogenase (nad(p)(+)) ald5  YDR089W - hypothetical protein  TPS3 - tps3p  HIS3 - imidazoleglycerol-phosphate dehydratase his3  RKR1 - ubiquitin-protein ligase rkr1  ILV5 - ketol-acid reductoisomerase  ENO1 - phosphopyruvate hydratase eno1  FOL2 - gtp cyclohydrolase i  RPL26A - ribosomal 60s subunit protein l26a  COQ1 - trans-hexaprenyltranstransferase  RPP0 - ribosomal protein p0  TUF1 - tuf1p  DPB4 - dpb4p  SAM4 - sam4p  GLN4 - glutamine--trna ligase  ARO1 - pentafunctional protein aro1p  HEK2 - hek2p  FAS2 - trifunctional fatty acid synthase subunit fas2  NCP1 - ncp1p  PGM1 - phosphoglucomutase pgm1  RPL17B - rpl17bp  RPL7B - ribosomal 60s subunit protein l7b  RPL32 - ribosomal 60s subunit protein l32  ARO8 - bifunctional 2-aminoadipate transaminase/aromatic-amino-acid:2-oxoglutarate transaminase  FLC1 - flc1p  GLC8 - glc8p  ARO3 - 3-deoxy-7-phosphoheptulonate synthase aro3  EHD3 - ehd3p  GON7 - gon7p  KRS1 - lysine--trna ligase krs1  IDH2 - isocitrate dehydrogenase (nad(+)) idh2  ARG4 - argininosuccinate lyase arg4  YHR020W - proline--trna ligase  HSP31 - hsp31p  HEM13 - coproporphyrinogen oxidase  LCB1 - serine c-palmitoyltransferase lcb1  APA2 - apa2p  RPL14B - ribosomal 60s subunit protein l14b  CAB1 - pantothenate kinase  LEO1 - leo1p  LCB2 - serine c-palmitoyltransferase lcb2  POS5 - pos5p  INP53 - phosphatidylinositol-3-/phosphoinositide 5-phosphatase inp53  TFC7 - tfc7p  PGM3 - phosphoglucomutase pgm3  RPS13 - ribosomal 40s subunit protein s13  RPS7A - ribosomal 40s subunit protein s7a  RPL29 - ribosomal 60s subunit protein l29  ENO2 - phosphopyruvate hydratase eno2  TSL1 - tsl1p  PCF11 - pcf11p  ALD6 - aldehyde dehydrogenase (nadp(+)) ald6  ASN2 - asparagine synthase (glutamine-hydrolyzing) 2  ERG9 - bifunctional farnesyl-diphosphate farnesyltransferase/squalene synthase  SPT6 - spt6p  GCD6 - gcd6p  MET10 - sulfite reductase subunit alpha  RSM7 - rsm7p  NHP6B - nhp6bp  POL30 - pol30p  RPL3 - ribosomal 60s subunit protein l3  CPA2 - cpa2p  RFC3 - replication factor c subunit 3  GPI13 - gpi13p  RSM26 - rsm26p  RPL6B - ribosomal 60s subunit protein l6b  YNL284C-B - gag-pol fusion protein  CYS4 - cystathionine beta-synthase cys4  PTI1 - pti1p  RPS5 - rps5p  CAR2 - ornithine-oxo-acid transaminase  THI4 - thi4p  BAT1 - branched-chain-amino-acid transaminase bat1  RPS1B - ribosomal 40s subunit protein s1b  CAB2 - phosphopantothenate--cysteine ligase cab2  SER33 - phosphoglycerate dehydrogenase ser33  TRP3 - bifunctional anthranilate synthase/indole-3-glycerol-phosphate synthase  RPA49 - rpa49p  FUR1 - uracil phosphoribosyltransferase  RNR4 - ribonucleotide-diphosphate reductase subunit rnr4  HOM2 - aspartate-semialdehyde dehydrogenase  DPS1 - aspartate--trna ligase dps1  ERG1 - squalene monooxygenase  PDR16 - pdr16p  TOP1 - dna topoisomerase 1  RPL6A - ribosomal 60s subunit protein l6a  PDX3 - pyridoxamine-phosphate oxidase pdx3  SSU72 - ssu72p  ARO9 - aromatic-amino-acid:2-oxoglutarate transaminase  DPH1 - dph1p  BDH1 - (r,r)-butanediol dehydrogenase  CDC60 - leucine--trna ligase cdc60  RPL5 - ribosomal 60s subunit protein l5  HMO1 - hmo1p  COQ5 - 2-hexaprenyl-6-methoxy-1,4-benzoquinone methyltransferase  KES1 - kes1p  SER2 - phosphoserine phosphatase  FAS1 - tetrafunctional fatty acid synthase subunit fas1  VAN1 - van1p  RPL33A - ribosomal 60s subunit protein l33a  MRPL6 - mitochondrial 54s ribosomal protein yml16  RPL24A - ribosomal 60s subunit protein l24a  GLC3 - 1,4-alpha-glucan branching enzyme  RPS3 - ribosomal 40s subunit protein s3  RPL30 - ribosomal 60s subunit protein l30  MRPL19 - mitochondrial 54s ribosomal protein yml19  CWH41 - cwh41p  POL31 - pol31p  ILV3 - ilv3p  PDC1 - indolepyruvate decarboxylase 1  CHS1 - chitin synthase chs1  TRP4 - anthranilate phosphoribosyltransferase  STH1 - sth1p  AYR1 - acylglycerone-phosphate reductase  LEU1 - 3-isopropylmalate dehydratase leu1  HIS7 - imidazoleglycerol-phosphate synthase  RPL15A - ribosomal 60s subunit protein l15a  ADE16 - bifunctional phosphoribosylaminoimidazolecarboxamide formyltransferase/imp cyclohydrolase ade16  ARO4 - 3-deoxy-7-phosphoheptulonate synthase aro4  HEM3 - hydroxymethylbilane synthase  ISC1 - inositol phosphosphingolipid phospholipase  HTS1 - histidine--trna ligase  PRO3 - pyrroline-5-carboxylate reductase  ARG1 - argininosuccinate synthase  GLN1 - glutamate--ammonia ligase  PRS5 - ribose phosphate diphosphokinase subunit prs5  RPL10 - ribosomal 60s subunit protein l10  AMD1 - amp deaminase  YMC1 - ymc1p  NRK1 - ribosylnicotinamide kinase  RPP2A - ribosomal protein p2a  NHP6A - nhp6ap  MEF1 - mef1p  ERG3 - c-5 sterol desaturase  FCY1 - cytosine deaminase  RPL25 - ribosomal 60s subunit protein l25  ARG7 - glutamate n-acetyltransferase  DLD3 - dld3p  RPL26B - ribosomal 60s subunit protein l26b  ERG27 - 3-keto-steroid reductase  YGR054W - hypothetical protein  GRE2 - methylglyoxal reductase (nadph-dependent) gre2  RCO1 - rco1p  RPO21 - rpo21p  LEU4 - 2-isopropylmalate synthase leu4  POL1 - pol1p  EHT1 - eht1p  ARP9 - arp9p  UTR4 - putative acireductone synthase utr4  STM1 - stm1p  ALD4 - aldehyde dehydrogenase (nadp(+)) ald4  MET12 - methylenetetrahydrofolate reductase (nad(p)h) met12  GLT1 - glutamate synthase (nadh)  MCM3 - mcm3p  PDC5 - indolepyruvate decarboxylase 5  ARA2 - d-arabinose 1-dehydrogenase (nad(p)(+)) ara2  SSB1 - hsp70 family atpase ssb1  RPL16B - ribosomal 60s subunit protein l16b  SUB1 - sub1p |
| GO:0009064 | glutamine family amino acid metabolic process | 2.45E-9 | 6.76E-7 | 8.52 (2325,39,98,14) | [+] Show genes  ARG5,6 - bifunctional acetylglutamate kinase/n-acetyl-gamma-glutamyl-phosphate reductase  IDH1 - isocitrate dehydrogenase (nad(+)) idh1  ARG7 - glutamate n-acetyltransferase  CPA2 - cpa2p  ARG1 - argininosuccinate synthase  GLN1 - glutamate--ammonia ligase  GLT1 - glutamate synthase (nadh)  TRP3 - bifunctional anthranilate synthase/indole-3-glycerol-phosphate synthase  ASN1 - asparagine synthase (glutamine-hydrolyzing) 1  GAD1 - glutamate decarboxylase gad1  DUR1,2 - bifunctional urea carboxylase/allophanate hydrolase  IDH2 - isocitrate dehydrogenase (nad(+)) idh2  HIS7 - imidazoleglycerol-phosphate synthase  ARG4 - argininosuccinate lyase arg4 |
| GO:0009072 | aromatic amino acid family metabolic process | 1E-8 | 2.59E-6 | 9.69 (2325,32,90,12) | [+] Show genes  HIS3 - imidazoleglycerol-phosphate dehydratase his3  ARO1 - pentafunctional protein aro1p  HIS4 - trifunctional histidinol dehydrogenase/phosphoribosyl-amp cyclohydrolase/phosphoribosyl-atp diphosphatase  ARO3 - 3-deoxy-7-phosphoheptulonate synthase aro3  ARO2 - bifunctional chorismate synthase/riboflavin reductase [nad(p)h] aro2  TRP2 - anthranilate synthase trp2  ARO8 - bifunctional 2-aminoadipate transaminase/aromatic-amino-acid:2-oxoglutarate transaminase  BNA5 - kynureninase  HIS7 - imidazoleglycerol-phosphate synthase  PDC1 - indolepyruvate decarboxylase 1  TRP3 - bifunctional anthranilate synthase/indole-3-glycerol-phosphate synthase  ARO4 - 3-deoxy-7-phosphoheptulonate synthase aro4 |
| GO:0009073 | aromatic amino acid family biosynthetic process | 1.36E-8 | 3.32E-6 | 12.30 (2325,21,90,10) | [+] Show genes  HIS3 - imidazoleglycerol-phosphate dehydratase his3  ARO1 - pentafunctional protein aro1p  ARO3 - 3-deoxy-7-phosphoheptulonate synthase aro3  HIS4 - trifunctional histidinol dehydrogenase/phosphoribosyl-amp cyclohydrolase/phosphoribosyl-atp diphosphatase  ARO2 - bifunctional chorismate synthase/riboflavin reductase [nad(p)h] aro2  TRP2 - anthranilate synthase trp2  ARO8 - bifunctional 2-aminoadipate transaminase/aromatic-amino-acid:2-oxoglutarate transaminase  HIS7 - imidazoleglycerol-phosphate synthase  TRP3 - bifunctional anthranilate synthase/indole-3-glycerol-phosphate synthase  ARO4 - 3-deoxy-7-phosphoheptulonate synthase aro4 |
| GO:1901566 | organonitrogen compound biosynthetic process | 2.24E-7 | 5.14E-5 | 1.50 (2325,465,451,135) | [+] Show genes  RPL29 - ribosomal 60s subunit protein l29  IDH1 - isocitrate dehydrogenase (nad(+)) idh1  GSH1 - gsh1p  ENO2 - phosphopyruvate hydratase eno2  ARO2 - bifunctional chorismate synthase/riboflavin reductase [nad(p)h] aro2  RPL9A - ribosomal 60s subunit protein l9a  RPL8A - ribosomal 60s subunit protein l8a  ASN2 - asparagine synthase (glutamine-hydrolyzing) 2  GCD6 - gcd6p  GLK1 - glucokinase  PRS2 - ribose phosphate diphosphokinase subunit prs2  MET10 - sulfite reductase subunit alpha  EMI2 - putative glucokinase  RSM7 - rsm7p  CPA2 - cpa2p  RPL3 - ribosomal 60s subunit protein l3  HIS4 - trifunctional histidinol dehydrogenase/phosphoribosyl-amp cyclohydrolase/phosphoribosyl-atp diphosphatase  CIT1 - citrate (si)-synthase cit1  ILV2 - acetolactate synthase catalytic subunit  RSM26 - rsm26p  RPL6B - ribosomal 60s subunit protein l6b  RPS2 - ribosomal 40s subunit protein s2  CYS4 - cystathionine beta-synthase cys4  RPS5 - rps5p  CPA1 - carbamoyl-phosphate synthase (glutamine-hydrolyzing) cpa1  CAR2 - ornithine-oxo-acid transaminase  THI4 - thi4p  BAT1 - branched-chain-amino-acid transaminase bat1  ILV6 - acetolactate synthase regulatory subunit  RPS1B - ribosomal 40s subunit protein s1b  ARG5,6 - bifunctional acetylglutamate kinase/n-acetyl-gamma-glutamyl-phosphate reductase  CAB2 - phosphopantothenate--cysteine ligase cab2  SER33 - phosphoglycerate dehydrogenase ser33  TRP3 - bifunctional anthranilate synthase/indole-3-glycerol-phosphate synthase  ABZ1 - 4-amino-4-deoxychorismate synthase  ASN1 - asparagine synthase (glutamine-hydrolyzing) 1  HIS1 - atp phosphoribosyltransferase  FUR1 - uracil phosphoribosyltransferase  GUK1 - guanylate kinase  HOM2 - aspartate-semialdehyde dehydrogenase  HOM3 - aspartate kinase  DPS1 - aspartate--trna ligase dps1  RSM18 - mitochondrial 37s ribosomal protein rsm18  TDH1 - tdh1p  LAC1 - sphingosine n-acyltransferase lac1  PDX3 - pyridoxamine-phosphate oxidase pdx3  RPL6A - ribosomal 60s subunit protein l6a  RPL31A - ribosomal 60s subunit protein l31a  ARO9 - aromatic-amino-acid:2-oxoglutarate transaminase  TRP2 - anthranilate synthase trp2  BNA5 - kynureninase  CDC60 - leucine--trna ligase cdc60  ALD2 - aldehyde dehydrogenase (nad(+)) ald2  ILV1 - threonine ammonia-lyase ilv1  TIF4632 - tif4632p  MRI1 - s-methyl-5-thioribose-1-phosphate isomerase mri1  MRPS12 - putative mitochondrial 37s ribosomal protein mrps12  RPL33B - ribosomal 60s subunit protein l33b  RPL5 - ribosomal 60s subunit protein l5  SER2 - phosphoserine phosphatase  RSM25 - mitochondrial 37s ribosomal protein rsm25  MRPL6 - mitochondrial 54s ribosomal protein yml16  PRS1 - ribose phosphate diphosphokinase subunit prs1  RPL33A - ribosomal 60s subunit protein l33a  VAN1 - van1p  LYS9 - saccharopine dehydrogenase (nadp+, l-glutamate-forming)  RPS3 - ribosomal 40s subunit protein s3  RPL24A - ribosomal 60s subunit protein l24a  RPL30 - ribosomal 60s subunit protein l30  MRPL19 - mitochondrial 54s ribosomal protein yml19  ILV3 - ilv3p  TIF11 - tif11p  CHS1 - chitin synthase chs1  TRP4 - anthranilate phosphoribosyltransferase  HIS3 - imidazoleglycerol-phosphate dehydratase his3  ILV5 - ketol-acid reductoisomerase  ENO1 - phosphopyruvate hydratase eno1  LEU1 - 3-isopropylmalate dehydratase leu1  FOL2 - gtp cyclohydrolase i  RPL26A - ribosomal 60s subunit protein l26a  HIS7 - imidazoleglycerol-phosphate synthase  RPL15A - ribosomal 60s subunit protein l15a  ADE16 - bifunctional phosphoribosylaminoimidazolecarboxamide formyltransferase/imp cyclohydrolase ade16  RPP0 - ribosomal protein p0  ARO4 - 3-deoxy-7-phosphoheptulonate synthase aro4  ISC1 - inositol phosphosphingolipid phospholipase  HEM3 - hydroxymethylbilane synthase  TUF1 - tuf1p  HTS1 - histidine--trna ligase  PRO3 - pyrroline-5-carboxylate reductase  GLN1 - glutamate--ammonia ligase  ARG1 - argininosuccinate synthase  PRS5 - ribose phosphate diphosphokinase subunit prs5  RPL10 - ribosomal 60s subunit protein l10  SAM4 - sam4p  AMD1 - amp deaminase  YMC1 - ymc1p  GLN4 - glutamine--trna ligase  NRK1 - ribosylnicotinamide kinase  ARO1 - pentafunctional protein aro1p  RPP2A - ribosomal protein p2a  MEF1 - mef1p  FCY1 - cytosine deaminase  RPS9B - ribosomal 40s subunit protein s9b  RPL25 - ribosomal 60s subunit protein l25  ARG7 - glutamate n-acetyltransferase  RPL17B - rpl17bp  RPL7B - ribosomal 60s subunit protein l7b  RPL32 - ribosomal 60s subunit protein l32  RPL26B - ribosomal 60s subunit protein l26b  ARO8 - bifunctional 2-aminoadipate transaminase/aromatic-amino-acid:2-oxoglutarate transaminase  YGR054W - hypothetical protein  FLC1 - flc1p  ARO3 - 3-deoxy-7-phosphoheptulonate synthase aro3  GON7 - gon7p  EHD3 - ehd3p  IDH2 - isocitrate dehydrogenase (nad(+)) idh2  KRS1 - lysine--trna ligase krs1  LEU4 - 2-isopropylmalate synthase leu4  ARG4 - argininosuccinate lyase arg4  YHR020W - proline--trna ligase  UTR4 - putative acireductone synthase utr4  HEM13 - coproporphyrinogen oxidase  LCB1 - serine c-palmitoyltransferase lcb1  RPL14B - ribosomal 60s subunit protein l14b  CAB1 - pantothenate kinase  MET12 - methylenetetrahydrofolate reductase (nad(p)h) met12  GLT1 - glutamate synthase (nadh)  POS5 - pos5p  LCB2 - serine c-palmitoyltransferase lcb2  RPS13 - ribosomal 40s subunit protein s13  PGM3 - phosphoglucomutase pgm3  SSB1 - hsp70 family atpase ssb1  RPS7A - ribosomal 40s subunit protein s7a  RPL16B - ribosomal 60s subunit protein l16b |
| GO:0006525 | arginine metabolic process | 2.59E-7 | 5.64E-5 | 14.13 (2325,9,128,7) | [+] Show genes  ARG5,6 - bifunctional acetylglutamate kinase/n-acetyl-gamma-glutamyl-phosphate reductase  ARG7 - glutamate n-acetyltransferase  CPA2 - cpa2p  DUR1,2 - bifunctional urea carboxylase/allophanate hydrolase  ARG1 - argininosuccinate synthase  CAR2 - ornithine-oxo-acid transaminase  ARG4 - argininosuccinate lyase arg4 |
| GO:0009423 | chorismate biosynthetic process | 3.32E-7 | 6.86E-5 | 46.50 (2325,4,50,4) | [+] Show genes  ARO1 - pentafunctional protein aro1p  ARO3 - 3-deoxy-7-phosphoheptulonate synthase aro3  ARO2 - bifunctional chorismate synthase/riboflavin reductase [nad(p)h] aro2  ARO4 - 3-deoxy-7-phosphoheptulonate synthase aro4 |
| GO:0009084 | glutamine family amino acid biosynthetic process | 3.6E-7 | 7.09E-5 | 5.51 (2325,21,261,13) | [+] Show genes  ARG5,6 - bifunctional acetylglutamate kinase/n-acetyl-gamma-glutamyl-phosphate reductase  IDH1 - isocitrate dehydrogenase (nad(+)) idh1  ARG7 - glutamate n-acetyltransferase  CPA2 - cpa2p  PRO3 - pyrroline-5-carboxylate reductase  ARG1 - argininosuccinate synthase  GLN1 - glutamate--ammonia ligase  CIT1 - citrate (si)-synthase cit1  GLT1 - glutamate synthase (nadh)  CPA1 - carbamoyl-phosphate synthase (glutamine-hydrolyzing) cpa1  CAR2 - ornithine-oxo-acid transaminase  IDH2 - isocitrate dehydrogenase (nad(+)) idh2  ARG4 - argininosuccinate lyase arg4 |
| GO:0044282 | small molecule catabolic process | 6.73E-7 | 1.26E-4 | 4.21 (2325,82,128,19) | [+] Show genes  PGM1 - phosphoglucomutase pgm1  APA2 - apa2p  PGM2 - phosphoglucomutase pgm2  PCS60 - pcs60p  ARO8 - bifunctional 2-aminoadipate transaminase/aromatic-amino-acid:2-oxoglutarate transaminase  DLD1 - dld1p  BNA5 - kynureninase  PDC1 - indolepyruvate decarboxylase 1  ILV1 - threonine ammonia-lyase ilv1  YPR127W - pyridoxine 4-dehydrogenase  INP53 - phosphatidylinositol-3-/phosphoinositide 5-phosphatase inp53  GAD1 - glutamate decarboxylase gad1  DUR1,2 - bifunctional urea carboxylase/allophanate hydrolase  PGM3 - phosphoglucomutase pgm3  EHD3 - ehd3p  LAP3 - lap3p  CAR2 - ornithine-oxo-acid transaminase  ALD5 - aldehyde dehydrogenase (nad(p)(+)) ald5  EHT1 - eht1p |
| GO:0043648 | dicarboxylic acid metabolic process | 7.53E-7 | 1.35E-4 | 5.65 (2325,45,128,14) | [+] Show genes  IDH1 - isocitrate dehydrogenase (nad(+)) idh1  FUM1 - fumarase fum1  PCS60 - pcs60p  ARO2 - bifunctional chorismate synthase/riboflavin reductase [nad(p)h] aro2  GLT1 - glutamate synthase (nadh)  BNA5 - kynureninase  GAD1 - glutamate decarboxylase gad1  ARO1 - pentafunctional protein aro1p  HOM2 - aspartate-semialdehyde dehydrogenase  HOM3 - aspartate kinase  ARO3 - 3-deoxy-7-phosphoheptulonate synthase aro3  CAR2 - ornithine-oxo-acid transaminase  IDH2 - isocitrate dehydrogenase (nad(+)) idh2  ARO4 - 3-deoxy-7-phosphoheptulonate synthase aro4 |
| GO:0009081 | branched-chain amino acid metabolic process | 7.98E-7 | 1.37E-4 | 10.01 (2325,22,95,9) | [+] Show genes  ILV5 - ketol-acid reductoisomerase  HOM2 - aspartate-semialdehyde dehydrogenase  EHD3 - ehd3p  ILV2 - acetolactate synthase catalytic subunit  ILV3 - ilv3p  ILV6 - acetolactate synthase regulatory subunit  LEU4 - 2-isopropylmalate synthase leu4  PDC1 - indolepyruvate decarboxylase 1  ILV1 - threonine ammonia-lyase ilv1 |
| GO:0006526 | arginine biosynthetic process | 1.51E-6 | 2.5E-4 | 48.44 (2325,6,32,4) | [+] Show genes  ARG5,6 - bifunctional acetylglutamate kinase/n-acetyl-gamma-glutamyl-phosphate reductase  ARG7 - glutamate n-acetyltransferase  CPA2 - cpa2p  ARG1 - argininosuccinate synthase |
| GO:0043650 | dicarboxylic acid biosynthetic process | 1.69E-6 | 2.68E-4 | 11.74 (2325,24,66,8) | [+] Show genes  IDH1 - isocitrate dehydrogenase (nad(+)) idh1  ARO1 - pentafunctional protein aro1p  ARO3 - 3-deoxy-7-phosphoheptulonate synthase aro3  IDH2 - isocitrate dehydrogenase (nad(+)) idh2  GLT1 - glutamate synthase (nadh)  ARO2 - bifunctional chorismate synthase/riboflavin reductase [nad(p)h] aro2  BNA5 - kynureninase  ARO4 - 3-deoxy-7-phosphoheptulonate synthase aro4 |
| GO:0055114 | oxidation-reduction process | 2.19E-6 | 3.36E-4 | 2.68 (2325,243,107,30) | [+] Show genes  ARG5,6 - bifunctional acetylglutamate kinase/n-acetyl-gamma-glutamyl-phosphate reductase  IDH1 - isocitrate dehydrogenase (nad(+)) idh1  PGM1 - phosphoglucomutase pgm1  GLC3 - 1,4-alpha-glucan branching enzyme  GPH1 - gph1p  PPA2 - ppa2p  DLD3 - dld3p  YDL124W - aldo-keto reductase superfamily protein  ARO2 - bifunctional chorismate synthase/riboflavin reductase [nad(p)h] aro2  PDC1 - indolepyruvate decarboxylase 1  ILV5 - ketol-acid reductoisomerase  HOM2 - aspartate-semialdehyde dehydrogenase  ZWF1 - glucose-6-phosphate dehydrogenase  EHD3 - ehd3p  IDH2 - isocitrate dehydrogenase (nad(+)) idh2  MET10 - sulfite reductase subunit alpha  HEM13 - coproporphyrinogen oxidase  PGM2 - phosphoglucomutase pgm2  ALD4 - aldehyde dehydrogenase (nadp(+)) ald4  HIS4 - trifunctional histidinol dehydrogenase/phosphoribosyl-amp cyclohydrolase/phosphoribosyl-atp diphosphatase  GLT1 - glutamate synthase (nadh)  GLR1 - glutathione-disulfide reductase glr1  BDH1 - (r,r)-butanediol dehydrogenase  DLD1 - dld1p  YPR127W - pyridoxine 4-dehydrogenase  ARO1 - pentafunctional protein aro1p  OLE1 - stearoyl-coa 9-desaturase  GLC7 - glc7p  YHB1 - yhb1p  ALD5 - aldehyde dehydrogenase (nad(p)(+)) ald5 |
| GO:0046417 | chorismate metabolic process | 2.56E-6 | 3.78E-4 | 37.20 (2325,5,50,4) | [+] Show genes  ARO1 - pentafunctional protein aro1p  ARO3 - 3-deoxy-7-phosphoheptulonate synthase aro3  ARO2 - bifunctional chorismate synthase/riboflavin reductase [nad(p)h] aro2  ARO4 - 3-deoxy-7-phosphoheptulonate synthase aro4 |
| GO:0009082 | branched-chain amino acid biosynthetic process | 3.71E-6 | 5.29E-4 | 12.24 (2325,14,95,7) | [+] Show genes  ILV5 - ketol-acid reductoisomerase  HOM2 - aspartate-semialdehyde dehydrogenase  ILV2 - acetolactate synthase catalytic subunit  ILV3 - ilv3p  ILV6 - acetolactate synthase regulatory subunit  LEU4 - 2-isopropylmalate synthase leu4  ILV1 - threonine ammonia-lyase ilv1 |
| GO:0009097 | isoleucine biosynthetic process | 4.92E-6 | 6.78E-4 | 14.68 (2325,10,95,6) | [+] Show genes  ILV5 - ketol-acid reductoisomerase  HOM2 - aspartate-semialdehyde dehydrogenase  ILV2 - acetolactate synthase catalytic subunit  ILV3 - ilv3p  ILV6 - acetolactate synthase regulatory subunit  ILV1 - threonine ammonia-lyase ilv1 |
| GO:0006549 | isoleucine metabolic process | 1.09E-5 | 1.46E-3 | 13.35 (2325,11,95,6) | [+] Show genes  ILV5 - ketol-acid reductoisomerase  HOM2 - aspartate-semialdehyde dehydrogenase  ILV2 - acetolactate synthase catalytic subunit  ILV3 - ilv3p  ILV6 - acetolactate synthase regulatory subunit  ILV1 - threonine ammonia-lyase ilv1 |
| GO:0009099 | valine biosynthetic process | 2.02E-5 | 2.61E-3 | 26.27 (2325,6,59,4) | [+] Show genes  ILV5 - ketol-acid reductoisomerase  ILV2 - acetolactate synthase catalytic subunit  ILV3 - ilv3p  ILV6 - acetolactate synthase regulatory subunit |
| GO:0006073 | cellular glucan metabolic process | 1.42E-4 | 1.78E-2 | 6.62 (2325,27,104,8) | [+] Show genes  KRE6 - kre6p  GLC3 - 1,4-alpha-glucan branching enzyme  PGM1 - phosphoglucomutase pgm1  GPH1 - gph1p  PGM2 - phosphoglucomutase pgm2  EXG1 - exg1p  CWH41 - cwh41p  GLC7 - glc7p |
| GO:0044042 | glucan metabolic process | 1.42E-4 | 1.73E-2 | 6.62 (2325,27,104,8) | [+] Show genes  KRE6 - kre6p  PGM1 - phosphoglucomutase pgm1  GLC3 - 1,4-alpha-glucan branching enzyme  GPH1 - gph1p  PGM2 - phosphoglucomutase pgm2  EXG1 - exg1p  CWH41 - cwh41p  GLC7 - glc7p |
| GO:0006573 | valine metabolic process | 2.81E-4 | 3.32E-2 | 17.51 (2325,9,59,4) | [+] Show genes  ILV5 - ketol-acid reductoisomerase  ILV2 - acetolactate synthase catalytic subunit  ILV3 - ilv3p  ILV6 - acetolactate synthase regulatory subunit |
| GO:1901564 | organonitrogen compound metabolic process | 3.53E-4 | 4.06E-2 | 1.30 (2325,952,251,134) | [+] Show genes  IDH1 - isocitrate dehydrogenase (nad(+)) idh1  RPL29 - ribosomal 60s subunit protein l29  RSP5 - nedd4 family e3 ubiquitin-protein ligase  SIT4 - sit4p  GSH1 - gsh1p  WHI2 - whi2p  ARO2 - bifunctional chorismate synthase/riboflavin reductase [nad(p)h] aro2  RPL8A - ribosomal 60s subunit protein l8a  APA1 - apa1p  GCD6 - gcd6p  GLK1 - glucokinase  MET10 - sulfite reductase subunit alpha  EMI2 - putative glucokinase  CPA2 - cpa2p  RPL3 - ribosomal 60s subunit protein l3  GPI13 - gpi13p  HIS4 - trifunctional histidinol dehydrogenase/phosphoribosyl-amp cyclohydrolase/phosphoribosyl-atp diphosphatase  CIT1 - citrate (si)-synthase cit1  ILV2 - acetolactate synthase catalytic subunit  SSA2 - hsp70 family chaperone ssa2  GLR1 - glutathione-disulfide reductase glr1  RSM26 - rsm26p  CYS4 - cystathionine beta-synthase cys4  GLC7 - glc7p  CAR2 - ornithine-oxo-acid transaminase  SET5 - set5p  BAT1 - branched-chain-amino-acid transaminase bat1  ILV6 - acetolactate synthase regulatory subunit  PTC5 - ptc5p  ARG5,6 - bifunctional acetylglutamate kinase/n-acetyl-gamma-glutamyl-phosphate reductase  RPS1B - ribosomal 40s subunit protein s1b  CAB2 - phosphopantothenate--cysteine ligase cab2  CDC37 - cdc37p  TRP3 - bifunctional anthranilate synthase/indole-3-glycerol-phosphate synthase  HUB1 - hub1p  TRA1 - tra1p  ASN1 - asparagine synthase (glutamine-hydrolyzing) 1  HIS1 - atp phosphoribosyltransferase  SSE1 - sse1p  GUK1 - guanylate kinase  HOM2 - aspartate-semialdehyde dehydrogenase  HOM3 - aspartate kinase  ZWF1 - glucose-6-phosphate dehydrogenase  DPS1 - aspartate--trna ligase dps1  CPR1 - peptidylprolyl isomerase cpr1  LAP3 - lap3p  LAC1 - sphingosine n-acyltransferase lac1  COX17 - cox17p  TUM1 - tum1p  RPL6A - ribosomal 60s subunit protein l6a  CRH1 - crh1p  TRP2 - anthranilate synthase trp2  BNA5 - kynureninase  ILV1 - threonine ammonia-lyase ilv1  ALD2 - aldehyde dehydrogenase (nad(+)) ald2  YPR127W - pyridoxine 4-dehydrogenase  SPP1 - spp1p  MVD1 - diphosphomevalonate decarboxylase mvd1  TPD3 - tpd3p  RPL5 - ribosomal 60s subunit protein l5  TRM112 - trm112p  KES1 - kes1p  SFM1 - sfm1p  RPL33A - ribosomal 60s subunit protein l33a  RPL24A - ribosomal 60s subunit protein l24a  RPL30 - ribosomal 60s subunit protein l30  MRPL19 - mitochondrial 54s ribosomal protein yml19  CHS5 - chs5p  CWH41 - cwh41p  ILV3 - ilv3p  FUS3 - fus3p  PDC1 - indolepyruvate decarboxylase 1  TRP4 - anthranilate phosphoribosyltransferase  HIS3 - imidazoleglycerol-phosphate dehydratase his3  TAL1 - sedoheptulose-7-phosphate:d-glyceraldehyde-3-phosphate transaldolase tal1  FAA4 - long-chain fatty acid-coa ligase faa4  ILV5 - ketol-acid reductoisomerase  RKR1 - ubiquitin-protein ligase rkr1  GAD1 - glutamate decarboxylase gad1  ENO1 - phosphopyruvate hydratase eno1  FOL2 - gtp cyclohydrolase i  HIS7 - imidazoleglycerol-phosphate synthase  RPP0 - ribosomal protein p0  ADE16 - bifunctional phosphoribosylaminoimidazolecarboxamide formyltransferase/imp cyclohydrolase ade16  CPR7 - cpr7p  ARO4 - 3-deoxy-7-phosphoheptulonate synthase aro4  ISC1 - inositol phosphosphingolipid phospholipase  TUF1 - tuf1p  PEP5 - pep5p  PRO3 - pyrroline-5-carboxylate reductase  GLN1 - glutamate--ammonia ligase  ARG1 - argininosuccinate synthase  YCK2 - yck2p  RPL10 - ribosomal 60s subunit protein l10  SAM4 - sam4p  AMD1 - amp deaminase  YMC1 - ymc1p  ARO1 - pentafunctional protein aro1p  DUR1,2 - bifunctional urea carboxylase/allophanate hydrolase  KIN1 - kin1p  MEF1 - mef1p  YCF1 - atp-binding cassette glutathione s-conjugate transporter ycf1  FCY1 - cytosine deaminase  SCJ1 - scj1p  RVB2 - ruvb family atp-dependent dna helicase reptin  ARG7 - glutamate n-acetyltransferase  RPL32 - ribosomal 60s subunit protein l32  RPL26B - ribosomal 60s subunit protein l26b  ARO8 - bifunctional 2-aminoadipate transaminase/aromatic-amino-acid:2-oxoglutarate transaminase  BPL1 - biotin--[acetyl-coa-carboxylase] ligase bpl1  SSE2 - sse2p  ARO3 - 3-deoxy-7-phosphoheptulonate synthase aro3  EHD3 - ehd3p  IDH2 - isocitrate dehydrogenase (nad(+)) idh2  LEU4 - 2-isopropylmalate synthase leu4  ARG4 - argininosuccinate lyase arg4  YHR020W - proline--trna ligase  HEM13 - coproporphyrinogen oxidase  APA2 - apa2p  RPL14B - ribosomal 60s subunit protein l14b  UBP2 - ubp2p  ALD4 - aldehyde dehydrogenase (nadp(+)) ald4  CAB1 - pantothenate kinase  LEO1 - leo1p  GLT1 - glutamate synthase (nadh)  UBC7 - e2 ubiquitin-conjugating protein ubc7  POS5 - pos5p  LCB2 - serine c-palmitoyltransferase lcb2  CPR5 - peptidylprolyl isomerase family protein cpr5  PGM3 - phosphoglucomutase pgm3  RPS13 - ribosomal 40s subunit protein s13  COX13 - cytochrome c oxidase subunit via  RPS7A - ribosomal 40s subunit protein s7a  NTA1 - nta1p |
| GO:0006592 | ornithine biosynthetic process | 3.67E-4 | 4.1E-2 | 72.66 (2325,2,32,2) | [+] Show genes  ARG5,6 - bifunctional acetylglutamate kinase/n-acetyl-gamma-glutamyl-phosphate reductase  ARG7 - glutamate n-acetyltransferase |
| GO:0000255 | allantoin metabolic process | 4.3E-4 | 4.68E-2 | 2,325.00 (2325,1,1,1) | [+] Show genes  DUR1,2 - bifunctional urea carboxylase/allophanate hydrolase |
| GO:0000256 | allantoin catabolic process | 4.3E-4 | 4.56E-2 | 2,325.00 (2325,1,1,1) | [+] Show genes  DUR1,2 - bifunctional urea carboxylase/allophanate hydrolase |
| GO:0043605 | cellular amide catabolic process | 4.3E-4 | 4.45E-2 | 2,325.00 (2325,1,1,1) | [+] Show genes  DUR1,2 - bifunctional urea carboxylase/allophanate hydrolase |
| GO:0043419 | urea catabolic process | 4.3E-4 | 4.34E-2 | 2,325.00 (2325,1,1,1) | [+] Show genes  DUR1,2 - bifunctional urea carboxylase/allophanate hydrolase |
| GO:0005975 | carbohydrate metabolic process | 5.22E-4 | 5.14E-2 | 2.38 (2325,126,186,24) | [+] Show genes  KRE6 - kre6p  YMR196W - hypothetical protein  PGM1 - phosphoglucomutase pgm1  GLC3 - 1,4-alpha-glucan branching enzyme  GPH1 - gph1p  PGM2 - phosphoglucomutase pgm2  EXG1 - exg1p  CWH41 - cwh41p  TSL1 - tsl1p  CRH1 - crh1p  GRE3 - trifunctional aldehyde reductase/xylose reductase/glucose 1-dehydrogenase (nadp(+))  PDC1 - indolepyruvate decarboxylase 1  SCW4 - scw4p  TPS3 - tps3p  TAL1 - sedoheptulose-7-phosphate:d-glyceraldehyde-3-phosphate transaldolase tal1  INP53 - phosphatidylinositol-3-/phosphoinositide 5-phosphatase inp53  ENO1 - phosphopyruvate hydratase eno1  YDR248C - gluconokinase  PGM3 - phosphoglucomutase pgm3  ZWF1 - glucose-6-phosphate dehydrogenase  SCW10 - scw10p  GLC7 - glc7p  MDH1 - malate dehydrogenase mdh1  EMI2 - putative glucokinase |
| GO:0006591 | ornithine metabolic process | 5.29E-4 | 5.09E-2 | 12.11 (2325,6,128,4) | [+] Show genes  ARG5,6 - bifunctional acetylglutamate kinase/n-acetyl-gamma-glutamyl-phosphate reductase  ARG7 - glutamate n-acetyltransferase  CAR2 - ornithine-oxo-acid transaminase  ARG4 - argininosuccinate lyase arg4 |
| GO:0006012 | galactose metabolic process | 5.88E-4 | 5.53E-2 | 14.72 (2325,3,158,3) | [+] Show genes  PGM1 - phosphoglucomutase pgm1  PGM2 - phosphoglucomutase pgm2  GRE3 - trifunctional aldehyde reductase/xylose reductase/glucose 1-dehydrogenase (nadp(+)) |
| GO:0019388 | galactose catabolic process | 5.88E-4 | 5.41E-2 | 14.72 (2325,3,158,3) | [+] Show genes  PGM1 - phosphoglucomutase pgm1  PGM2 - phosphoglucomutase pgm2  GRE3 - trifunctional aldehyde reductase/xylose reductase/glucose 1-dehydrogenase (nadp(+)) |
| GO:0019413 | acetate biosynthetic process | 6.37E-4 | 5.72E-2 | 62.00 (2325,3,25,2) | [+] Show genes  ALD4 - aldehyde dehydrogenase (nadp(+)) ald4  ALD5 - aldehyde dehydrogenase (nad(p)(+)) ald5 |
| GO:0034219 | carbohydrate transmembrane transport | 6.49E-4 | 5.71E-2 | 3.59 (2325,7,648,7) | [+] Show genes  HXK2 - hexokinase 2  HXT4 - hxt4p  HXT3 - hxt3p  HXK1 - hexokinase 1  GLK1 - glucokinase  HXT1 - hxt1p  HXT2 - hxt2p |
| GO:0008645 | hexose transmembrane transport | 6.49E-4 | 5.59E-2 | 3.59 (2325,7,648,7) | [+] Show genes  HXK2 - hexokinase 2  HXT3 - hxt3p  HXT4 - hxt4p  HXK1 - hexokinase 1  GLK1 - glucokinase  HXT1 - hxt1p  HXT2 - hxt2p |
| GO:1904659 | glucose transmembrane transport | 6.49E-4 | 5.48E-2 | 3.59 (2325,7,648,7) | [+] Show genes  HXK2 - hexokinase 2  HXT4 - hxt4p  HXT3 - hxt3p  HXK1 - hexokinase 1  GLK1 - glucokinase  HXT1 - hxt1p  HXT2 - hxt2p |
| GO:0015749 | monosaccharide transmembrane transport | 6.49E-4 | 5.37E-2 | 3.59 (2325,7,648,7) | [+] Show genes  HXK2 - hexokinase 2  HXT4 - hxt4p  HXT3 - hxt3p  HXK1 - hexokinase 1  GLK1 - glucokinase  HXT1 - hxt1p  HXT2 - hxt2p |
| GO:0043603 | cellular amide metabolic process | 6.5E-4 | 5.27E-2 | 1.48 (2325,260,458,76) | [+] Show genes  RPL29 - ribosomal 60s subunit protein l29  RPL24A - ribosomal 60s subunit protein l24a  RPS3 - ribosomal 40s subunit protein s3  RPL30 - ribosomal 60s subunit protein l30  MRPL19 - mitochondrial 54s ribosomal protein yml19  GSH1 - gsh1p  RPL9A - ribosomal 60s subunit protein l9a  TIF11 - tif11p  RPL8A - ribosomal 60s subunit protein l8a  FAA4 - long-chain fatty acid-coa ligase faa4  ASN2 - asparagine synthase (glutamine-hydrolyzing) 2  FOL2 - gtp cyclohydrolase i  GCD6 - gcd6p  RPL26A - ribosomal 60s subunit protein l26a  RPL15A - ribosomal 60s subunit protein l15a  RPP0 - ribosomal protein p0  ISC1 - inositol phosphosphingolipid phospholipase  RSM7 - rsm7p  TUF1 - tuf1p  HTS1 - histidine--trna ligase  FAA1 - long-chain fatty acid-coa ligase faa1  RPL3 - ribosomal 60s subunit protein l3  ARG1 - argininosuccinate synthase  CIT1 - citrate (si)-synthase cit1  GLR1 - glutathione-disulfide reductase glr1  RPL10 - ribosomal 60s subunit protein l10  RSM26 - rsm26p  RPL6B - ribosomal 60s subunit protein l6b  GLN4 - glutamine--trna ligase  RPS2 - ribosomal 40s subunit protein s2  RPP2A - ribosomal protein p2a  DUR1,2 - bifunctional urea carboxylase/allophanate hydrolase  RPS5 - rps5p  MEF1 - mef1p  YCF1 - atp-binding cassette glutathione s-conjugate transporter ycf1  GTT1 - bifunctional glutathione transferase/peroxidase  RPS1B - ribosomal 40s subunit protein s1b  RPL25 - ribosomal 60s subunit protein l25  RPS9B - ribosomal 40s subunit protein s9b  RPL17B - rpl17bp  RPL7B - ribosomal 60s subunit protein l7b  RPL32 - ribosomal 60s subunit protein l32  RPL26B - ribosomal 60s subunit protein l26b  YDL124W - aldo-keto reductase superfamily protein  OXP1 - oxp1p  ABZ1 - 4-amino-4-deoxychorismate synthase  ASN1 - asparagine synthase (glutamine-hydrolyzing) 1  YGR054W - hypothetical protein  KGD2 - alpha-ketoglutarate dehydrogenase kgd2  SUI1 - sui1p  APE2 - ape2p  DPS1 - aspartate--trna ligase dps1  LRO1 - phospholipid:diacylglycerol acyltransferase  RSM18 - mitochondrial 37s ribosomal protein rsm18  EHD3 - ehd3p  KRS1 - lysine--trna ligase krs1  LAC1 - sphingosine n-acyltransferase lac1  YHR020W - proline--trna ligase  RPL6A - ribosomal 60s subunit protein l6a  RPL31A - ribosomal 60s subunit protein l31a  RPL14B - ribosomal 60s subunit protein l14b  MET12 - methylenetetrahydrofolate reductase (nad(p)h) met12  CDC60 - leucine--trna ligase cdc60  ALD2 - aldehyde dehydrogenase (nad(+)) ald2  TIF4632 - tif4632p  MRPS12 - putative mitochondrial 37s ribosomal protein mrps12  RPL33B - ribosomal 60s subunit protein l33b  MVD1 - diphosphomevalonate decarboxylase mvd1  RPS13 - ribosomal 40s subunit protein s13  RPL5 - ribosomal 60s subunit protein l5  SSB1 - hsp70 family atpase ssb1  RSM25 - mitochondrial 37s ribosomal protein rsm25  RPL33A - ribosomal 60s subunit protein l33a  MRPL6 - mitochondrial 54s ribosomal protein yml16  RPS7A - ribosomal 40s subunit protein s7a  RPL16B - ribosomal 60s subunit protein l16b |
| GO:0019627 | urea metabolic process | 6.93E-4 | 5.51E-2 | 59.62 (2325,3,26,2) | [+] Show genes  DUR1,2 - bifunctional urea carboxylase/allophanate hydrolase  ARG1 - argininosuccinate synthase |
| GO:0002181 | cytoplasmic translation | 7.78E-4 | 6.07E-2 | 1.99 (2325,80,438,30) | [+] Show genes  RPL29 - ribosomal 60s subunit protein l29  RPS1B - ribosomal 40s subunit protein s1b  RPL25 - ribosomal 60s subunit protein l25  RPL24A - ribosomal 60s subunit protein l24a  RPS3 - ribosomal 40s subunit protein s3  RPL30 - ribosomal 60s subunit protein l30  RPL17B - rpl17bp  RPL7B - ribosomal 60s subunit protein l7b  RPL32 - ribosomal 60s subunit protein l32  RPL26B - ribosomal 60s subunit protein l26b  RPL9A - ribosomal 60s subunit protein l9a  RPL8A - ribosomal 60s subunit protein l8a  RPL26A - ribosomal 60s subunit protein l26a  RPL15A - ribosomal 60s subunit protein l15a  RPP0 - ribosomal protein p0  RPL6A - ribosomal 60s subunit protein l6a  RPL3 - ribosomal 60s subunit protein l3  RPL14B - ribosomal 60s subunit protein l14b  RPL31A - ribosomal 60s subunit protein l31a  RPL10 - ribosomal 60s subunit protein l10  RPL6B - ribosomal 60s subunit protein l6b  RPL33B - ribosomal 60s subunit protein l33b  RPP2A - ribosomal protein p2a  RPS5 - rps5p  RPS13 - ribosomal 40s subunit protein s13  RPL5 - ribosomal 60s subunit protein l5  SSB1 - hsp70 family atpase ssb1  RPS7A - ribosomal 40s subunit protein s7a  RPL33A - ribosomal 60s subunit protein l33a  RPL16B - ribosomal 60s subunit protein l16b |
| GO:0044264 | cellular polysaccharide metabolic process | 8.58E-4 | 6.57E-2 | 5.26 (2325,34,104,8) | [+] Show genes  KRE6 - kre6p  PGM1 - phosphoglucomutase pgm1  GLC3 - 1,4-alpha-glucan branching enzyme  GPH1 - gph1p  PGM2 - phosphoglucomutase pgm2  EXG1 - exg1p  CWH41 - cwh41p  GLC7 - glc7p |
| GO:0016054 | organic acid catabolic process | 9.61E-4 | 7.23E-2 | 3.84 (2325,52,128,11) | [+] Show genes  GAD1 - glutamate decarboxylase gad1  PCS60 - pcs60p  LAP3 - lap3p  EHD3 - ehd3p  CAR2 - ornithine-oxo-acid transaminase  ARO8 - bifunctional 2-aminoadipate transaminase/aromatic-amino-acid:2-oxoglutarate transaminase  BNA5 - kynureninase  DLD1 - dld1p  PDC1 - indolepyruvate decarboxylase 1  ILV1 - threonine ammonia-lyase ilv1  EHT1 - eht1p |
| GO:0046395 | carboxylic acid catabolic process | 9.61E-4 | 7.1E-2 | 3.84 (2325,52,128,11) | [+] Show genes  GAD1 - glutamate decarboxylase gad1  PCS60 - pcs60p  LAP3 - lap3p  EHD3 - ehd3p  CAR2 - ornithine-oxo-acid transaminase  ARO8 - bifunctional 2-aminoadipate transaminase/aromatic-amino-acid:2-oxoglutarate transaminase  BNA5 - kynureninase  DLD1 - dld1p  PDC1 - indolepyruvate decarboxylase 1  ILV1 - threonine ammonia-lyase ilv1  EHT1 - eht1p |

Species used: Saccharomyces cerevisiae

The system has recognized 2343 genes out of 2345 gene terms entered by the user.  
 141 genes were recognized by gene symbol and 2202 genes by other gene IDs .  
Only 2325 of these genes are associated with a GO term.

  
 Output in Microsoft Excel format  

% List genereted using GOrilla
% http://cbl-gorilla.cs.technion.ac.il/
% GO term pValue
GO:0008652 2.15E-18
GO:0016053 2.79E-16
GO:0046394 2.79E-16
GO:0006520 2.84E-16
GO:0019752 2.85E-15
GO:1901607 9.92E-15
GO:0043436 1.93E-14
GO:0006082 2.25E-14
GO:1901605 3.94E-14
GO:0044281 4.06E-14
GO:0044283 1.68E-13
GO:1901576 4.37E-11
GO:0044249 5.38E-11
GO:0009058 8.61E-11
GO:0009064 2.45E-9
GO:0009072 1E-8
GO:0009073 1.36E-8
GO:1901566 2.24E-7
GO:0006525 2.59E-7
GO:0009423 3.32E-7
GO:0009084 3.6E-7
GO:0044282 6.73E-7
GO:0043648 7.53E-7
GO:0009081 7.98E-7
GO:0006526 1.51E-6
GO:0043650 1.69E-6
GO:0055114 2.19E-6
GO:0046417 2.56E-6
GO:0009082 3.71E-6
GO:0009097 4.92E-6
GO:0006549 1.09E-5
GO:0009099 2.02E-5
GO:0006073 1.42E-4
GO:0044042 1.42E-4
GO:0006573 2.81E-4
GO:1901564 3.53E-4
GO:0006592 3.67E-4
GO:0000255 4.3E-4
GO:0000256 4.3E-4
GO:0043605 4.3E-4
GO:0043419 4.3E-4
GO:0005975 5.22E-4
GO:0006591 5.29E-4
GO:0006012 5.88E-4
GO:0019388 5.88E-4
GO:0019413 6.37E-4
GO:0034219 6.49E-4
GO:0008645 6.49E-4
GO:1904659 6.49E-4
GO:0015749 6.49E-4
GO:0043603 6.5E-4
GO:0019627 6.93E-4
GO:0002181 7.78E-4
GO:0044264 8.58E-4
GO:0016054 9.61E-4
GO:0046395 9.61E-4
 Visualize output in REViGO   

The GOrilla database is periodically updated using the GO database and other sources.  
The GOrilla database was last updated on Sep 21, 2019

This results page will be available on this site for one month from now (until
Oct 23, 2019
). You can bookmark this page and come back to it later.

  
**'P-value'** is the enrichment
p-value computed according to the mHG or HG model. This p-value is not
corrected for multiple testing of 4135 GO terms.  
  
**'FDR q-value'** is the correction of the above p-value for multiple testing using the Benjamini and Hochberg (1995) method.   
Namely, for the i^th^ term (ranked according to p-value) the FDR q-value is (p-value \* number of GO terms) / i.   
  
**Enrichment (N, B, n, b)** is defined as follows:  
N - is the total number of genes  
B - is the total number of genes associated with a specific GO term  
n - is the number of genes in the top of the user's input list or in the target set when appropriate  
b - is the number of genes in the intersection  
Enrichment = (b/n) / (B/N)  
  
**Genes:** For each GO term you can see the list of associated genes that appear in the optimal top of the list.  
Each gene name is specified by gene symbol followed by a short description of the gene   

Back to GOrilla analysis results
