## Supplemental Data S1 for "A simple mass-action model predicts genome-wide protein timecourses from mRNA trajectories during a dynamic response in two strains of *Saccharomyces cerevisiae*": GOPROCESS_rna.html

Results


 **rna** 

*P-value color scale*

|  |  |  |  |  |
| --- | --- | --- | --- | --- |
| > 10^-3^ | 10^-3^ to 10^-5^ | 10^-5^ to 10^-7^ | 10^-7^ to 10^-9^ | < 10^-9^ |


|  |  |  |  |  |  |
| --- | --- | --- | --- | --- | --- |
| **GO term** | **Description** | **P-value** | **FDR q-value** | **Enrichment (N, B, n, b)** | **Genes** |
| GO:0002181 | cytoplasmic translation | 5.22E-56 | 2.72E-52 | 12.98 (5247,132,193,63) | [+] Show genes  RPL29 - ribosomal 60s subunit protein l29  RPL2B - ribosomal 60s subunit protein l2b  RPS0B - ribosomal 40s subunit protein s0b  RPL24A - ribosomal 60s subunit protein l24a  RPL30 - ribosomal 60s subunit protein l30  RPL9A - ribosomal 60s subunit protein l9a  RPL23B - ribosomal 60s subunit protein l23b  RPL40A - ubiquitin-ribosomal 60s subunit protein l40a fusion protein  RPL13B - ribosomal 60s subunit protein l13b  RPL38 - ribosomal 60s subunit protein l38  RPS16A - ribosomal 40s subunit protein s16a  RPL8A - ribosomal 60s subunit protein l8a  RPS8B - ribosomal 40s subunit protein s8b  RPL16A - ribosomal 60s subunit protein l16a  RPL26A - ribosomal 60s subunit protein l26a  RPL11A - ribosomal 60s subunit protein l11a  RPL15A - ribosomal 60s subunit protein l15a  RPP0 - ribosomal protein p0  RPL27B - ribosomal 60s subunit protein l27b  RPL34B - ribosomal 60s subunit protein l34b  RPL3 - ribosomal 60s subunit protein l3  RPS6A - ribosomal 40s subunit protein s6a  RPS18B - ribosomal 40s subunit protein s18b  RPL19A - ribosomal 60s subunit protein l19a  RPS21B - rps21bp  RPS10B - ribosomal 40s subunit protein s10b  RPP2B - ribosomal protein p2b  RPL6B - ribosomal 60s subunit protein l6b  RPS15 - ribosomal 40s subunit protein s15  RPP2A - ribosomal protein p2a  RPS5 - rps5p  RPL37A - ribosomal 60s subunit protein l37a  RPS17A - ribosomal 40s subunit protein s17a  RPL24B - ribosomal 60s subunit protein l24b  RPS28A - ribosomal 40s subunit protein s28a  RPS1B - ribosomal 40s subunit protein s1b  RPL17A - ribosomal 60s subunit protein l17a  RPS27B - ribosomal 40s subunit protein s27b  RPL14A - ribosomal 60s subunit protein l14a  RPL21A - ribosomal 60s subunit protein l21a  RPP1B - ribosomal protein p1b  RPL17B - rpl17bp  RPL31B - ribosomal 60s subunit protein l31b  RPL32 - ribosomal 60s subunit protein l32  RPL26B - ribosomal 60s subunit protein l26b  RPL22B - ribosomal 60s subunit protein l22b  RPL35B - ribosomal 60s subunit protein l35b  RPS10A - ribosomal 40s subunit protein s10a  RPL41B - ribosomal 60s subunit protein l41b  RPL34A - ribosomal 60s subunit protein l34a  RPL27A - ribosomal 60s subunit protein l27a  RPS22A - rps22ap  RPS17B - ribosomal 40s subunit protein s17b  RPL6A - ribosomal 60s subunit protein l6a  RPL42B - ribosomal 60s subunit protein l42b  RPS14A - ribosomal 40s subunit protein s14a  RPL31A - ribosomal 60s subunit protein l31a  RPL11B - ribosomal 60s subunit protein l11b  RPS13 - ribosomal 40s subunit protein s13  RPS26A - ribosomal 40s subunit protein s26a  RPS7A - ribosomal 40s subunit protein s7a  RPL16B - ribosomal 60s subunit protein l16b  RPS24A - ribosomal 40s subunit protein s24a |
| GO:0006412 | translation | 1.14E-34 | 2.98E-31 | 6.71 (5247,313,155,62) | [+] Show genes  RPL29 - ribosomal 60s subunit protein l29  RPL2B - ribosomal 60s subunit protein l2b  RPS0B - ribosomal 40s subunit protein s0b  RPL24A - ribosomal 60s subunit protein l24a  RPL30 - ribosomal 60s subunit protein l30  RPL9A - ribosomal 60s subunit protein l9a  RPL23B - ribosomal 60s subunit protein l23b  RPL40A - ubiquitin-ribosomal 60s subunit protein l40a fusion protein  RPL38 - ribosomal 60s subunit protein l38  RPL13B - ribosomal 60s subunit protein l13b  TEF1 - tef1p  RPS16A - ribosomal 40s subunit protein s16a  RPL8A - ribosomal 60s subunit protein l8a  RPS8B - ribosomal 40s subunit protein s8b  RPL26A - ribosomal 60s subunit protein l26a  RPL11A - ribosomal 60s subunit protein l11a  RPL15A - ribosomal 60s subunit protein l15a  RPP0 - ribosomal protein p0  RPL27B - ribosomal 60s subunit protein l27b  RPL34B - ribosomal 60s subunit protein l34b  RPL3 - ribosomal 60s subunit protein l3  RPS18B - ribosomal 40s subunit protein s18b  RPL19A - ribosomal 60s subunit protein l19a  RPS21B - rps21bp  RPS10B - ribosomal 40s subunit protein s10b  RPP2B - ribosomal protein p2b  RPL6B - ribosomal 60s subunit protein l6b  RPS15 - ribosomal 40s subunit protein s15  RPP2A - ribosomal protein p2a  RPS5 - rps5p  RPS17A - ribosomal 40s subunit protein s17a  RPL37A - ribosomal 60s subunit protein l37a  RPL24B - ribosomal 60s subunit protein l24b  RPS28A - ribosomal 40s subunit protein s28a  RPS27B - ribosomal 40s subunit protein s27b  RPL17A - ribosomal 60s subunit protein l17a  RPS1B - ribosomal 40s subunit protein s1b  RPP1B - ribosomal protein p1b  RPL21A - ribosomal 60s subunit protein l21a  RPL14A - ribosomal 60s subunit protein l14a  RPL17B - rpl17bp  MRPL24 - mitochondrial 54s ribosomal protein yml24/yml14  RPL32 - ribosomal 60s subunit protein l32  RPL26B - ribosomal 60s subunit protein l26b  RPL22B - ribosomal 60s subunit protein l22b  RPL35B - ribosomal 60s subunit protein l35b  RPS10A - ribosomal 40s subunit protein s10a  RPL34A - ribosomal 60s subunit protein l34a  RPL27A - ribosomal 60s subunit protein l27a  RPS22A - rps22ap  RPS17B - ribosomal 40s subunit protein s17b  YHR020W - proline--trna ligase  RPL6A - ribosomal 60s subunit protein l6a  RPS23B - ribosomal 40s subunit protein s23b  RPL42B - ribosomal 60s subunit protein l42b  RPS14A - ribosomal 40s subunit protein s14a  RPL31A - ribosomal 60s subunit protein l31a  RPS13 - ribosomal 40s subunit protein s13  RPS26A - ribosomal 40s subunit protein s26a  RPS7A - ribosomal 40s subunit protein s7a  RPL16B - ribosomal 60s subunit protein l16b  RPS24A - ribosomal 40s subunit protein s24a |
| GO:0043043 | peptide biosynthetic process | 2.11E-34 | 3.66E-31 | 6.64 (5247,316,155,62) | [+] Show genes  RPL29 - ribosomal 60s subunit protein l29  RPL2B - ribosomal 60s subunit protein l2b  RPS0B - ribosomal 40s subunit protein s0b  RPL24A - ribosomal 60s subunit protein l24a  RPL30 - ribosomal 60s subunit protein l30  RPL9A - ribosomal 60s subunit protein l9a  RPL23B - ribosomal 60s subunit protein l23b  RPL40A - ubiquitin-ribosomal 60s subunit protein l40a fusion protein  RPL38 - ribosomal 60s subunit protein l38  RPL13B - ribosomal 60s subunit protein l13b  TEF1 - tef1p  RPS16A - ribosomal 40s subunit protein s16a  RPL8A - ribosomal 60s subunit protein l8a  RPS8B - ribosomal 40s subunit protein s8b  RPL26A - ribosomal 60s subunit protein l26a  RPL15A - ribosomal 60s subunit protein l15a  RPL11A - ribosomal 60s subunit protein l11a  RPP0 - ribosomal protein p0  RPL27B - ribosomal 60s subunit protein l27b  RPL34B - ribosomal 60s subunit protein l34b  RPL3 - ribosomal 60s subunit protein l3  RPS18B - ribosomal 40s subunit protein s18b  RPL19A - ribosomal 60s subunit protein l19a  RPS21B - rps21bp  RPS10B - ribosomal 40s subunit protein s10b  RPP2B - ribosomal protein p2b  RPL6B - ribosomal 60s subunit protein l6b  RPS15 - ribosomal 40s subunit protein s15  RPP2A - ribosomal protein p2a  RPS5 - rps5p  RPS17A - ribosomal 40s subunit protein s17a  RPL37A - ribosomal 60s subunit protein l37a  RPL24B - ribosomal 60s subunit protein l24b  RPS28A - ribosomal 40s subunit protein s28a  RPS27B - ribosomal 40s subunit protein s27b  RPS1B - ribosomal 40s subunit protein s1b  RPL17A - ribosomal 60s subunit protein l17a  RPL21A - ribosomal 60s subunit protein l21a  RPP1B - ribosomal protein p1b  RPL14A - ribosomal 60s subunit protein l14a  RPL17B - rpl17bp  MRPL24 - mitochondrial 54s ribosomal protein yml24/yml14  RPL32 - ribosomal 60s subunit protein l32  RPL26B - ribosomal 60s subunit protein l26b  RPL22B - ribosomal 60s subunit protein l22b  RPL35B - ribosomal 60s subunit protein l35b  RPS10A - ribosomal 40s subunit protein s10a  RPL34A - ribosomal 60s subunit protein l34a  RPL27A - ribosomal 60s subunit protein l27a  RPS22A - rps22ap  RPS17B - ribosomal 40s subunit protein s17b  YHR020W - proline--trna ligase  RPL6A - ribosomal 60s subunit protein l6a  RPS23B - ribosomal 40s subunit protein s23b  RPL42B - ribosomal 60s subunit protein l42b  RPS14A - ribosomal 40s subunit protein s14a  RPL31A - ribosomal 60s subunit protein l31a  RPS13 - ribosomal 40s subunit protein s13  RPS26A - ribosomal 40s subunit protein s26a  RPS7A - ribosomal 40s subunit protein s7a  RPL16B - ribosomal 60s subunit protein l16b  RPS24A - ribosomal 40s subunit protein s24a |
| GO:0006518 | peptide metabolic process | 6.59E-32 | 8.58E-29 | 6.07 (5247,346,155,62) | [+] Show genes  RPL29 - ribosomal 60s subunit protein l29  RPL2B - ribosomal 60s subunit protein l2b  RPS0B - ribosomal 40s subunit protein s0b  RPL24A - ribosomal 60s subunit protein l24a  RPL30 - ribosomal 60s subunit protein l30  RPL9A - ribosomal 60s subunit protein l9a  RPL23B - ribosomal 60s subunit protein l23b  RPL40A - ubiquitin-ribosomal 60s subunit protein l40a fusion protein  RPL38 - ribosomal 60s subunit protein l38  RPL13B - ribosomal 60s subunit protein l13b  TEF1 - tef1p  RPS16A - ribosomal 40s subunit protein s16a  RPL8A - ribosomal 60s subunit protein l8a  RPS8B - ribosomal 40s subunit protein s8b  RPL26A - ribosomal 60s subunit protein l26a  RPL11A - ribosomal 60s subunit protein l11a  RPL15A - ribosomal 60s subunit protein l15a  RPP0 - ribosomal protein p0  RPL27B - ribosomal 60s subunit protein l27b  RPL34B - ribosomal 60s subunit protein l34b  RPL3 - ribosomal 60s subunit protein l3  RPS18B - ribosomal 40s subunit protein s18b  RPL19A - ribosomal 60s subunit protein l19a  RPS21B - rps21bp  RPS10B - ribosomal 40s subunit protein s10b  RPP2B - ribosomal protein p2b  RPL6B - ribosomal 60s subunit protein l6b  RPS15 - ribosomal 40s subunit protein s15  RPP2A - ribosomal protein p2a  RPS5 - rps5p  RPS17A - ribosomal 40s subunit protein s17a  RPL37A - ribosomal 60s subunit protein l37a  RPL24B - ribosomal 60s subunit protein l24b  RPS28A - ribosomal 40s subunit protein s28a  RPS27B - ribosomal 40s subunit protein s27b  RPL17A - ribosomal 60s subunit protein l17a  RPS1B - ribosomal 40s subunit protein s1b  RPP1B - ribosomal protein p1b  RPL21A - ribosomal 60s subunit protein l21a  RPL14A - ribosomal 60s subunit protein l14a  RPL17B - rpl17bp  MRPL24 - mitochondrial 54s ribosomal protein yml24/yml14  RPL32 - ribosomal 60s subunit protein l32  RPL26B - ribosomal 60s subunit protein l26b  RPL22B - ribosomal 60s subunit protein l22b  RPL35B - ribosomal 60s subunit protein l35b  RPS10A - ribosomal 40s subunit protein s10a  RPL34A - ribosomal 60s subunit protein l34a  RPL27A - ribosomal 60s subunit protein l27a  RPS22A - rps22ap  RPS17B - ribosomal 40s subunit protein s17b  YHR020W - proline--trna ligase  RPL6A - ribosomal 60s subunit protein l6a  RPS23B - ribosomal 40s subunit protein s23b  RPL42B - ribosomal 60s subunit protein l42b  RPS14A - ribosomal 40s subunit protein s14a  RPL31A - ribosomal 60s subunit protein l31a  RPS13 - ribosomal 40s subunit protein s13  RPS26A - ribosomal 40s subunit protein s26a  RPS7A - ribosomal 40s subunit protein s7a  RPL16B - ribosomal 60s subunit protein l16b  RPS24A - ribosomal 40s subunit protein s24a |
| GO:0043604 | amide biosynthetic process | 1.44E-31 | 1.5E-28 | 5.86 (5247,364,155,63) | [+] Show genes  RPL29 - ribosomal 60s subunit protein l29  RPL2B - ribosomal 60s subunit protein l2b  RPS0B - ribosomal 40s subunit protein s0b  RPL24A - ribosomal 60s subunit protein l24a  RPL30 - ribosomal 60s subunit protein l30  RPL9A - ribosomal 60s subunit protein l9a  RPL40A - ubiquitin-ribosomal 60s subunit protein l40a fusion protein  RPL23B - ribosomal 60s subunit protein l23b  RPL38 - ribosomal 60s subunit protein l38  RPL13B - ribosomal 60s subunit protein l13b  TEF1 - tef1p  RPS16A - ribosomal 40s subunit protein s16a  RPL8A - ribosomal 60s subunit protein l8a  RPS8B - ribosomal 40s subunit protein s8b  RPL26A - ribosomal 60s subunit protein l26a  RPL11A - ribosomal 60s subunit protein l11a  RPL15A - ribosomal 60s subunit protein l15a  RPP0 - ribosomal protein p0  RPL27B - ribosomal 60s subunit protein l27b  RPL34B - ribosomal 60s subunit protein l34b  RPL3 - ribosomal 60s subunit protein l3  RPS18B - ribosomal 40s subunit protein s18b  RPL19A - ribosomal 60s subunit protein l19a  RPS21B - rps21bp  RPS10B - ribosomal 40s subunit protein s10b  RPP2B - ribosomal protein p2b  RPL6B - ribosomal 60s subunit protein l6b  RPS15 - ribosomal 40s subunit protein s15  RPP2A - ribosomal protein p2a  RPS5 - rps5p  RPS17A - ribosomal 40s subunit protein s17a  RPL37A - ribosomal 60s subunit protein l37a  RPL24B - ribosomal 60s subunit protein l24b  RPS28A - ribosomal 40s subunit protein s28a  RPS27B - ribosomal 40s subunit protein s27b  RPL17A - ribosomal 60s subunit protein l17a  RPS1B - ribosomal 40s subunit protein s1b  RPP1B - ribosomal protein p1b  RPL21A - ribosomal 60s subunit protein l21a  RPL14A - ribosomal 60s subunit protein l14a  RPL17B - rpl17bp  MRPL24 - mitochondrial 54s ribosomal protein yml24/yml14  RPL32 - ribosomal 60s subunit protein l32  RPL26B - ribosomal 60s subunit protein l26b  RPL22B - ribosomal 60s subunit protein l22b  RPL35B - ribosomal 60s subunit protein l35b  RPS10A - ribosomal 40s subunit protein s10a  ABZ1 - 4-amino-4-deoxychorismate synthase  RPL34A - ribosomal 60s subunit protein l34a  RPL27A - ribosomal 60s subunit protein l27a  RPS22A - rps22ap  RPS17B - ribosomal 40s subunit protein s17b  YHR020W - proline--trna ligase  RPL6A - ribosomal 60s subunit protein l6a  RPS23B - ribosomal 40s subunit protein s23b  RPL42B - ribosomal 60s subunit protein l42b  RPS14A - ribosomal 40s subunit protein s14a  RPL31A - ribosomal 60s subunit protein l31a  RPS13 - ribosomal 40s subunit protein s13  RPS26A - ribosomal 40s subunit protein s26a  RPS7A - ribosomal 40s subunit protein s7a  RPL16B - ribosomal 60s subunit protein l16b  RPS24A - ribosomal 40s subunit protein s24a |
| GO:0043603 | cellular amide metabolic process | 2.8E-28 | 2.43E-25 | 5.09 (5247,426,155,64) | [+] Show genes  RPL2B - ribosomal 60s subunit protein l2b  RPS0B - ribosomal 40s subunit protein s0b  RPL29 - ribosomal 60s subunit protein l29  RPL24A - ribosomal 60s subunit protein l24a  RPL30 - ribosomal 60s subunit protein l30  RPL9A - ribosomal 60s subunit protein l9a  RPL40A - ubiquitin-ribosomal 60s subunit protein l40a fusion protein  RPL23B - ribosomal 60s subunit protein l23b  RPL13B - ribosomal 60s subunit protein l13b  RPL38 - ribosomal 60s subunit protein l38  TEF1 - tef1p  RPS16A - ribosomal 40s subunit protein s16a  RPL8A - ribosomal 60s subunit protein l8a  RPS8B - ribosomal 40s subunit protein s8b  RPL26A - ribosomal 60s subunit protein l26a  RPL15A - ribosomal 60s subunit protein l15a  RPL11A - ribosomal 60s subunit protein l11a  RPP0 - ribosomal protein p0  RPL34B - ribosomal 60s subunit protein l34b  RPL27B - ribosomal 60s subunit protein l27b  RPL3 - ribosomal 60s subunit protein l3  RPS18B - ribosomal 40s subunit protein s18b  RPL19A - ribosomal 60s subunit protein l19a  RPS21B - rps21bp  RPS10B - ribosomal 40s subunit protein s10b  RPP2B - ribosomal protein p2b  RPL6B - ribosomal 60s subunit protein l6b  RPS15 - ribosomal 40s subunit protein s15  RPP2A - ribosomal protein p2a  DUR1,2 - bifunctional urea carboxylase/allophanate hydrolase  RPS5 - rps5p  RPS17A - ribosomal 40s subunit protein s17a  RPL37A - ribosomal 60s subunit protein l37a  RPL24B - ribosomal 60s subunit protein l24b  RPS28A - ribosomal 40s subunit protein s28a  RPL17A - ribosomal 60s subunit protein l17a  RPS1B - ribosomal 40s subunit protein s1b  RPS27B - ribosomal 40s subunit protein s27b  RPP1B - ribosomal protein p1b  RPL21A - ribosomal 60s subunit protein l21a  RPL14A - ribosomal 60s subunit protein l14a  RPL17B - rpl17bp  MRPL24 - mitochondrial 54s ribosomal protein yml24/yml14  RPL32 - ribosomal 60s subunit protein l32  RPL26B - ribosomal 60s subunit protein l26b  RPL22B - ribosomal 60s subunit protein l22b  RPL35B - ribosomal 60s subunit protein l35b  RPS10A - ribosomal 40s subunit protein s10a  ABZ1 - 4-amino-4-deoxychorismate synthase  RPL34A - ribosomal 60s subunit protein l34a  RPL27A - ribosomal 60s subunit protein l27a  RPS22A - rps22ap  RPS17B - ribosomal 40s subunit protein s17b  YHR020W - proline--trna ligase  RPL6A - ribosomal 60s subunit protein l6a  RPS23B - ribosomal 40s subunit protein s23b  RPL42B - ribosomal 60s subunit protein l42b  RPS14A - ribosomal 40s subunit protein s14a  RPL31A - ribosomal 60s subunit protein l31a  RPS13 - ribosomal 40s subunit protein s13  RPS26A - ribosomal 40s subunit protein s26a  RPS7A - ribosomal 40s subunit protein s7a  RPL16B - ribosomal 60s subunit protein l16b  RPS24A - ribosomal 40s subunit protein s24a |
| GO:0034645 | cellular macromolecule biosynthetic process | 5.02E-23 | 3.73E-20 | 3.72 (5247,642,156,71) | [+] Show genes  RPL2B - ribosomal 60s subunit protein l2b  RPL29 - ribosomal 60s subunit protein l29  RPS0B - ribosomal 40s subunit protein s0b  RPL24A - ribosomal 60s subunit protein l24a  RPL30 - ribosomal 60s subunit protein l30  KTR2 - ktr2p  RPL9A - ribosomal 60s subunit protein l9a  RPL40A - ubiquitin-ribosomal 60s subunit protein l40a fusion protein  RPL23B - ribosomal 60s subunit protein l23b  RPL38 - ribosomal 60s subunit protein l38  RPL13B - ribosomal 60s subunit protein l13b  RPS16A - ribosomal 40s subunit protein s16a  TEF1 - tef1p  RPL8A - ribosomal 60s subunit protein l8a  ASC1 - asc1p  RPS8B - ribosomal 40s subunit protein s8b  DPB3 - dpb3p  RPL26A - ribosomal 60s subunit protein l26a  RPL11A - ribosomal 60s subunit protein l11a  RPL15A - ribosomal 60s subunit protein l15a  RPP0 - ribosomal protein p0  RPL34B - ribosomal 60s subunit protein l34b  RPL27B - ribosomal 60s subunit protein l27b  RPL3 - ribosomal 60s subunit protein l3  RPS18B - ribosomal 40s subunit protein s18b  RPL19A - ribosomal 60s subunit protein l19a  RPS21B - rps21bp  SLX1 - slx1p  RPS10B - ribosomal 40s subunit protein s10b  RPP2B - ribosomal protein p2b  RPL6B - ribosomal 60s subunit protein l6b  RPS15 - ribosomal 40s subunit protein s15  RPP2A - ribosomal protein p2a  RPS5 - rps5p  RPS17A - ribosomal 40s subunit protein s17a  RPL37A - ribosomal 60s subunit protein l37a  RPL24B - ribosomal 60s subunit protein l24b  GSC2 - gsc2p  RPS28A - ribosomal 40s subunit protein s28a  RPL17A - ribosomal 60s subunit protein l17a  RPS1B - ribosomal 40s subunit protein s1b  RPS27B - ribosomal 40s subunit protein s27b  RPL21A - ribosomal 60s subunit protein l21a  RPP1B - ribosomal protein p1b  RPL14A - ribosomal 60s subunit protein l14a  RPL17B - rpl17bp  RPL32 - ribosomal 60s subunit protein l32  MRPL24 - mitochondrial 54s ribosomal protein yml24/yml14  RPL26B - ribosomal 60s subunit protein l26b  RPL22B - ribosomal 60s subunit protein l22b  RPL35B - ribosomal 60s subunit protein l35b  RPS10A - ribosomal 40s subunit protein s10a  RPL34A - ribosomal 60s subunit protein l34a  RPL27A - ribosomal 60s subunit protein l27a  RNR4 - ribonucleotide-diphosphate reductase subunit rnr4  RPS22A - rps22ap  RPS17B - ribosomal 40s subunit protein s17b  YHR020W - proline--trna ligase  RPL6A - ribosomal 60s subunit protein l6a  RPC37 - rpc37p  RPS23B - ribosomal 40s subunit protein s23b  RPL42B - ribosomal 60s subunit protein l42b  RPS14A - ribosomal 40s subunit protein s14a  RPL31A - ribosomal 60s subunit protein l31a  STM1 - stm1p  CHS3 - chitin synthase chs3  RPS13 - ribosomal 40s subunit protein s13  RPS26A - ribosomal 40s subunit protein s26a  RPS7A - ribosomal 40s subunit protein s7a  RPL16B - ribosomal 60s subunit protein l16b  RPS24A - ribosomal 40s subunit protein s24a |
| GO:1901566 | organonitrogen compound biosynthetic process | 5.04E-21 | 3.28E-18 | 3.51 (5247,716,144,69) | [+] Show genes  RPL2B - ribosomal 60s subunit protein l2b  RPL29 - ribosomal 60s subunit protein l29  RPS0B - ribosomal 40s subunit protein s0b  RPL30 - ribosomal 60s subunit protein l30  KTR2 - ktr2p  RPL40A - ubiquitin-ribosomal 60s subunit protein l40a fusion protein  RPL23B - ribosomal 60s subunit protein l23b  CHS1 - chitin synthase chs1  RPL13B - ribosomal 60s subunit protein l13b  RPL38 - ribosomal 60s subunit protein l38  RPS16A - ribosomal 40s subunit protein s16a  TEF1 - tef1p  RPL8A - ribosomal 60s subunit protein l8a  ILV5 - ketol-acid reductoisomerase  RPS8B - ribosomal 40s subunit protein s8b  ATP18 - atp18p  RPL26A - ribosomal 60s subunit protein l26a  RPL15A - ribosomal 60s subunit protein l15a  RPL11A - ribosomal 60s subunit protein l11a  RPP0 - ribosomal protein p0  RPL34B - ribosomal 60s subunit protein l34b  RPL27B - ribosomal 60s subunit protein l27b  RPL3 - ribosomal 60s subunit protein l3  CYT1 - ubiquinol--cytochrome-c reductase catalytic subunit cyt1  RPS18B - ribosomal 40s subunit protein s18b  RPL19A - ribosomal 60s subunit protein l19a  RPS21B - rps21bp  RPS10B - ribosomal 40s subunit protein s10b  RPP2B - ribosomal protein p2b  RPL6B - ribosomal 60s subunit protein l6b  RPS15 - ribosomal 40s subunit protein s15  RPP2A - ribosomal protein p2a  RPS5 - rps5p  RPS17A - ribosomal 40s subunit protein s17a  RPL37A - ribosomal 60s subunit protein l37a  RPL24B - ribosomal 60s subunit protein l24b  RPS28A - ribosomal 40s subunit protein s28a  RPL17A - ribosomal 60s subunit protein l17a  RPS1B - ribosomal 40s subunit protein s1b  RPS27B - ribosomal 40s subunit protein s27b  RPP1B - ribosomal protein p1b  RPL21A - ribosomal 60s subunit protein l21a  RPL14A - ribosomal 60s subunit protein l14a  RPL17B - rpl17bp  RPL32 - ribosomal 60s subunit protein l32  MRPL24 - mitochondrial 54s ribosomal protein yml24/yml14  RPL26B - ribosomal 60s subunit protein l26b  RPL22B - ribosomal 60s subunit protein l22b  RPS10A - ribosomal 40s subunit protein s10a  PGK1 - phosphoglycerate kinase  ABZ1 - 4-amino-4-deoxychorismate synthase  RPL34A - ribosomal 60s subunit protein l34a  RPL27A - ribosomal 60s subunit protein l27a  RPS22A - rps22ap  RPS17B - ribosomal 40s subunit protein s17b  YHR020W - proline--trna ligase  RPL6A - ribosomal 60s subunit protein l6a  RPS23B - ribosomal 40s subunit protein s23b  RPL42B - ribosomal 60s subunit protein l42b  RPS14A - ribosomal 40s subunit protein s14a  RPL31A - ribosomal 60s subunit protein l31a  CHS3 - chitin synthase chs3  BNA5 - kynureninase  THR4 - threonine synthase thr4  RPS13 - ribosomal 40s subunit protein s13  RPS26A - ribosomal 40s subunit protein s26a  RPS7A - ribosomal 40s subunit protein s7a  RPL16B - ribosomal 60s subunit protein l16b  RPS24A - ribosomal 40s subunit protein s24a |
| GO:0009059 | macromolecule biosynthetic process | 1.47E-20 | 8.52E-18 | 3.35 (5247,722,156,72) | [+] Show genes  RPL2B - ribosomal 60s subunit protein l2b  RPL29 - ribosomal 60s subunit protein l29  RPS0B - ribosomal 40s subunit protein s0b  RPL24A - ribosomal 60s subunit protein l24a  RPL30 - ribosomal 60s subunit protein l30  KTR2 - ktr2p  RPL9A - ribosomal 60s subunit protein l9a  RPL40A - ubiquitin-ribosomal 60s subunit protein l40a fusion protein  RPL23B - ribosomal 60s subunit protein l23b  CHS1 - chitin synthase chs1  RPL38 - ribosomal 60s subunit protein l38  RPL13B - ribosomal 60s subunit protein l13b  RPS16A - ribosomal 40s subunit protein s16a  TEF1 - tef1p  RPL8A - ribosomal 60s subunit protein l8a  ASC1 - asc1p  RPS8B - ribosomal 40s subunit protein s8b  DPB3 - dpb3p  RPL26A - ribosomal 60s subunit protein l26a  RPL15A - ribosomal 60s subunit protein l15a  RPL11A - ribosomal 60s subunit protein l11a  RPP0 - ribosomal protein p0  RPL34B - ribosomal 60s subunit protein l34b  RPL27B - ribosomal 60s subunit protein l27b  RPL3 - ribosomal 60s subunit protein l3  RPS18B - ribosomal 40s subunit protein s18b  RPL19A - ribosomal 60s subunit protein l19a  RPS21B - rps21bp  SLX1 - slx1p  RPS10B - ribosomal 40s subunit protein s10b  RPP2B - ribosomal protein p2b  RPL6B - ribosomal 60s subunit protein l6b  RPS15 - ribosomal 40s subunit protein s15  RPP2A - ribosomal protein p2a  RPS5 - rps5p  RPS17A - ribosomal 40s subunit protein s17a  RPL37A - ribosomal 60s subunit protein l37a  RPL24B - ribosomal 60s subunit protein l24b  GSC2 - gsc2p  RPS28A - ribosomal 40s subunit protein s28a  RPL17A - ribosomal 60s subunit protein l17a  RPS1B - ribosomal 40s subunit protein s1b  RPS27B - ribosomal 40s subunit protein s27b  RPP1B - ribosomal protein p1b  RPL21A - ribosomal 60s subunit protein l21a  RPL14A - ribosomal 60s subunit protein l14a  RPL17B - rpl17bp  RPL32 - ribosomal 60s subunit protein l32  MRPL24 - mitochondrial 54s ribosomal protein yml24/yml14  RPL26B - ribosomal 60s subunit protein l26b  RPL22B - ribosomal 60s subunit protein l22b  RPL35B - ribosomal 60s subunit protein l35b  RPS10A - ribosomal 40s subunit protein s10a  RPL34A - ribosomal 60s subunit protein l34a  RPL27A - ribosomal 60s subunit protein l27a  RNR4 - ribonucleotide-diphosphate reductase subunit rnr4  RPS22A - rps22ap  RPS17B - ribosomal 40s subunit protein s17b  YHR020W - proline--trna ligase  RPL6A - ribosomal 60s subunit protein l6a  RPC37 - rpc37p  RPS23B - ribosomal 40s subunit protein s23b  RPL42B - ribosomal 60s subunit protein l42b  RPS14A - ribosomal 40s subunit protein s14a  RPL31A - ribosomal 60s subunit protein l31a  STM1 - stm1p  CHS3 - chitin synthase chs3  RPS13 - ribosomal 40s subunit protein s13  RPS26A - ribosomal 40s subunit protein s26a  RPS7A - ribosomal 40s subunit protein s7a  RPL16B - ribosomal 60s subunit protein l16b  RPS24A - ribosomal 40s subunit protein s24a |
| GO:0044271 | cellular nitrogen compound biosynthetic process | 2.49E-16 | 1.3E-13 | 2.90 (5247,823,156,71) | [+] Show genes  RPL2B - ribosomal 60s subunit protein l2b  RPS0B - ribosomal 40s subunit protein s0b  RPL29 - ribosomal 60s subunit protein l29  RPL24A - ribosomal 60s subunit protein l24a  RPL30 - ribosomal 60s subunit protein l30  RPL9A - ribosomal 60s subunit protein l9a  RPL40A - ubiquitin-ribosomal 60s subunit protein l40a fusion protein  RPL23B - ribosomal 60s subunit protein l23b  RPL13B - ribosomal 60s subunit protein l13b  RPL38 - ribosomal 60s subunit protein l38  TEF1 - tef1p  RPS16A - ribosomal 40s subunit protein s16a  RPL8A - ribosomal 60s subunit protein l8a  RPS8B - ribosomal 40s subunit protein s8b  ATP18 - atp18p  DPB3 - dpb3p  RPL26A - ribosomal 60s subunit protein l26a  RPL11A - ribosomal 60s subunit protein l11a  RPL15A - ribosomal 60s subunit protein l15a  RPP0 - ribosomal protein p0  RPL34B - ribosomal 60s subunit protein l34b  RPL27B - ribosomal 60s subunit protein l27b  RPL3 - ribosomal 60s subunit protein l3  CYT1 - ubiquinol--cytochrome-c reductase catalytic subunit cyt1  RPS18B - ribosomal 40s subunit protein s18b  RPL19A - ribosomal 60s subunit protein l19a  RPS21B - rps21bp  RPS10B - ribosomal 40s subunit protein s10b  RPP2B - ribosomal protein p2b  RPL6B - ribosomal 60s subunit protein l6b  RPS15 - ribosomal 40s subunit protein s15  RPP2A - ribosomal protein p2a  RPS5 - rps5p  RPL37A - ribosomal 60s subunit protein l37a  RPS17A - ribosomal 40s subunit protein s17a  RPL24B - ribosomal 60s subunit protein l24b  RPS28A - ribosomal 40s subunit protein s28a  RPS1B - ribosomal 40s subunit protein s1b  RPL17A - ribosomal 60s subunit protein l17a  RPS27B - ribosomal 40s subunit protein s27b  RPL21A - ribosomal 60s subunit protein l21a  RPP1B - ribosomal protein p1b  RPL14A - ribosomal 60s subunit protein l14a  RPL17B - rpl17bp  RPL32 - ribosomal 60s subunit protein l32  MRPL24 - mitochondrial 54s ribosomal protein yml24/yml14  RPL26B - ribosomal 60s subunit protein l26b  RPL22B - ribosomal 60s subunit protein l22b  RPL35B - ribosomal 60s subunit protein l35b  RPS10A - ribosomal 40s subunit protein s10a  PGK1 - phosphoglycerate kinase  ABZ1 - 4-amino-4-deoxychorismate synthase  RPL34A - ribosomal 60s subunit protein l34a  RPL27A - ribosomal 60s subunit protein l27a  RNR4 - ribonucleotide-diphosphate reductase subunit rnr4  RPS22A - rps22ap  RPS17B - ribosomal 40s subunit protein s17b  YHR020W - proline--trna ligase  RPL6A - ribosomal 60s subunit protein l6a  RPC37 - rpc37p  RPS23B - ribosomal 40s subunit protein s23b  RPL42B - ribosomal 60s subunit protein l42b  APA2 - apa2p  RPS14A - ribosomal 40s subunit protein s14a  RPL31A - ribosomal 60s subunit protein l31a  BNA5 - kynureninase  RPS13 - ribosomal 40s subunit protein s13  RPS26A - ribosomal 40s subunit protein s26a  RPS7A - ribosomal 40s subunit protein s7a  RPS24A - ribosomal 40s subunit protein s24a  RPL16B - ribosomal 60s subunit protein l16b |
| GO:0044267 | cellular protein metabolic process | 4.73E-13 | 2.24E-10 | 2.34 (5247,1056,155,73) | [+] Show genes  RPL2B - ribosomal 60s subunit protein l2b  RPS0B - ribosomal 40s subunit protein s0b  RPL29 - ribosomal 60s subunit protein l29  RPL24A - ribosomal 60s subunit protein l24a  RPL30 - ribosomal 60s subunit protein l30  PFA3 - pfa3p  KTR2 - ktr2p  RPL9A - ribosomal 60s subunit protein l9a  RPL40A - ubiquitin-ribosomal 60s subunit protein l40a fusion protein  RPL23B - ribosomal 60s subunit protein l23b  RPL38 - ribosomal 60s subunit protein l38  RPL13B - ribosomal 60s subunit protein l13b  TEF1 - tef1p  RPS16A - ribosomal 40s subunit protein s16a  RPL8A - ribosomal 60s subunit protein l8a  PPT1 - ppt1p  RPS8B - ribosomal 40s subunit protein s8b  RPL26A - ribosomal 60s subunit protein l26a  IRE1 - ire1p  RPL11A - ribosomal 60s subunit protein l11a  RPL15A - ribosomal 60s subunit protein l15a  RPP0 - ribosomal protein p0  RPL34B - ribosomal 60s subunit protein l34b  RPL27B - ribosomal 60s subunit protein l27b  RPL3 - ribosomal 60s subunit protein l3  RPS18B - ribosomal 40s subunit protein s18b  RPL19A - ribosomal 60s subunit protein l19a  RPS21B - rps21bp  RPS10B - ribosomal 40s subunit protein s10b  RPP2B - ribosomal protein p2b  FPR4 - peptidylprolyl isomerase fpr4  RPL6B - ribosomal 60s subunit protein l6b  RPS15 - ribosomal 40s subunit protein s15  RPP2A - ribosomal protein p2a  RPS5 - rps5p  RPL37A - ribosomal 60s subunit protein l37a  RPS17A - ribosomal 40s subunit protein s17a  RPL24B - ribosomal 60s subunit protein l24b  RPS28A - ribosomal 40s subunit protein s28a  RPS1B - ribosomal 40s subunit protein s1b  RPL17A - ribosomal 60s subunit protein l17a  RPS27B - ribosomal 40s subunit protein s27b  RPL21A - ribosomal 60s subunit protein l21a  RPP1B - ribosomal protein p1b  RPL14A - ribosomal 60s subunit protein l14a  RPL17B - rpl17bp  RPL32 - ribosomal 60s subunit protein l32  MRPL24 - mitochondrial 54s ribosomal protein yml24/yml14  RPL26B - ribosomal 60s subunit protein l26b  RPL22B - ribosomal 60s subunit protein l22b  PRB1 - prb1p  RPL35B - ribosomal 60s subunit protein l35b  SLT2 - slt2p  RPS10A - ribosomal 40s subunit protein s10a  ACT1 - actin  NCS2 - ncs2p  RPL34A - ribosomal 60s subunit protein l34a  RPL27A - ribosomal 60s subunit protein l27a  RPS22A - rps22ap  RPS17B - ribosomal 40s subunit protein s17b  YHR020W - proline--trna ligase  RPL6A - ribosomal 60s subunit protein l6a  RPS23B - ribosomal 40s subunit protein s23b  RPL42B - ribosomal 60s subunit protein l42b  RPS14A - ribosomal 40s subunit protein s14a  RPL31A - ribosomal 60s subunit protein l31a  HSL1 - hsl1p  RIO1 - rio1p  RPS13 - ribosomal 40s subunit protein s13  RPS26A - ribosomal 40s subunit protein s26a  RPS7A - ribosomal 40s subunit protein s7a  RPS24A - ribosomal 40s subunit protein s24a  RPL16B - ribosomal 60s subunit protein l16b |
| GO:0044249 | cellular biosynthetic process | 6.48E-13 | 2.81E-10 | 2.16 (5247,1261,156,81) | [+] Show genes  RPL2B - ribosomal 60s subunit protein l2b  RPS0B - ribosomal 40s subunit protein s0b  RPL29 - ribosomal 60s subunit protein l29  RPL24A - ribosomal 60s subunit protein l24a  RPL30 - ribosomal 60s subunit protein l30  KTR2 - ktr2p  TSL1 - tsl1p  RPL9A - ribosomal 60s subunit protein l9a  RPL40A - ubiquitin-ribosomal 60s subunit protein l40a fusion protein  RPL23B - ribosomal 60s subunit protein l23b  RPL13B - ribosomal 60s subunit protein l13b  RPL38 - ribosomal 60s subunit protein l38  TEF1 - tef1p  RPS16A - ribosomal 40s subunit protein s16a  RPL8A - ribosomal 60s subunit protein l8a  ILV5 - ketol-acid reductoisomerase  ASC1 - asc1p  RPS8B - ribosomal 40s subunit protein s8b  ATP18 - atp18p  DPB3 - dpb3p  RPL26A - ribosomal 60s subunit protein l26a  RPL11A - ribosomal 60s subunit protein l11a  RPL15A - ribosomal 60s subunit protein l15a  RPP0 - ribosomal protein p0  RPL34B - ribosomal 60s subunit protein l34b  RPL27B - ribosomal 60s subunit protein l27b  RPL3 - ribosomal 60s subunit protein l3  CYT1 - ubiquinol--cytochrome-c reductase catalytic subunit cyt1  RPS18B - ribosomal 40s subunit protein s18b  RPL19A - ribosomal 60s subunit protein l19a  RPS21B - rps21bp  SLX1 - slx1p  RPS10B - ribosomal 40s subunit protein s10b  RPP2B - ribosomal protein p2b  RPL6B - ribosomal 60s subunit protein l6b  RPS15 - ribosomal 40s subunit protein s15  RPP2A - ribosomal protein p2a  RPS5 - rps5p  RPS17A - ribosomal 40s subunit protein s17a  RPL37A - ribosomal 60s subunit protein l37a  RPL24B - ribosomal 60s subunit protein l24b  GSC2 - gsc2p  RPS28A - ribosomal 40s subunit protein s28a  RPS27B - ribosomal 40s subunit protein s27b  RPS1B - ribosomal 40s subunit protein s1b  RPL17A - ribosomal 60s subunit protein l17a  RPL21A - ribosomal 60s subunit protein l21a  RPP1B - ribosomal protein p1b  RPL14A - ribosomal 60s subunit protein l14a  RPL17B - rpl17bp  RPL32 - ribosomal 60s subunit protein l32  MRPL24 - mitochondrial 54s ribosomal protein yml24/yml14  RPL26B - ribosomal 60s subunit protein l26b  RPL22B - ribosomal 60s subunit protein l22b  RPL35B - ribosomal 60s subunit protein l35b  RPS10A - ribosomal 40s subunit protein s10a  PGK1 - phosphoglycerate kinase  ABZ1 - 4-amino-4-deoxychorismate synthase  RPL34A - ribosomal 60s subunit protein l34a  RPL27A - ribosomal 60s subunit protein l27a  RNR4 - ribonucleotide-diphosphate reductase subunit rnr4  RPS22A - rps22ap  RPS17B - ribosomal 40s subunit protein s17b  YHR020W - proline--trna ligase  PAH1 - phosphatidate phosphatase pah1  RPL6A - ribosomal 60s subunit protein l6a  RPC37 - rpc37p  RPS23B - ribosomal 40s subunit protein s23b  RPL42B - ribosomal 60s subunit protein l42b  APA2 - apa2p  RPS14A - ribosomal 40s subunit protein s14a  RPL31A - ribosomal 60s subunit protein l31a  STM1 - stm1p  CHS3 - chitin synthase chs3  BNA5 - kynureninase  THR4 - threonine synthase thr4  RPS13 - ribosomal 40s subunit protein s13  RPS26A - ribosomal 40s subunit protein s26a  RPS7A - ribosomal 40s subunit protein s7a  RPS24A - ribosomal 40s subunit protein s24a  RPL16B - ribosomal 60s subunit protein l16b |
| GO:1901576 | organic substance biosynthetic process | 1.09E-12 | 4.37E-10 | 2.12 (5247,1298,156,82) | [+] Show genes  RPL2B - ribosomal 60s subunit protein l2b  RPS0B - ribosomal 40s subunit protein s0b  RPL29 - ribosomal 60s subunit protein l29  RPL24A - ribosomal 60s subunit protein l24a  RPL30 - ribosomal 60s subunit protein l30  KTR2 - ktr2p  TSL1 - tsl1p  RPL9A - ribosomal 60s subunit protein l9a  RPL40A - ubiquitin-ribosomal 60s subunit protein l40a fusion protein  RPL23B - ribosomal 60s subunit protein l23b  RPL13B - ribosomal 60s subunit protein l13b  RPL38 - ribosomal 60s subunit protein l38  CHS1 - chitin synthase chs1  TEF1 - tef1p  RPS16A - ribosomal 40s subunit protein s16a  RPL8A - ribosomal 60s subunit protein l8a  ILV5 - ketol-acid reductoisomerase  ASC1 - asc1p  RPS8B - ribosomal 40s subunit protein s8b  ATP18 - atp18p  DPB3 - dpb3p  RPL26A - ribosomal 60s subunit protein l26a  RPL11A - ribosomal 60s subunit protein l11a  RPL15A - ribosomal 60s subunit protein l15a  RPP0 - ribosomal protein p0  RPL34B - ribosomal 60s subunit protein l34b  RPL27B - ribosomal 60s subunit protein l27b  RPL3 - ribosomal 60s subunit protein l3  CYT1 - ubiquinol--cytochrome-c reductase catalytic subunit cyt1  RPS18B - ribosomal 40s subunit protein s18b  RPL19A - ribosomal 60s subunit protein l19a  RPS21B - rps21bp  SLX1 - slx1p  RPS10B - ribosomal 40s subunit protein s10b  RPP2B - ribosomal protein p2b  RPL6B - ribosomal 60s subunit protein l6b  RPS15 - ribosomal 40s subunit protein s15  RPP2A - ribosomal protein p2a  RPS5 - rps5p  RPS17A - ribosomal 40s subunit protein s17a  RPL37A - ribosomal 60s subunit protein l37a  RPL24B - ribosomal 60s subunit protein l24b  GSC2 - gsc2p  RPS28A - ribosomal 40s subunit protein s28a  RPS27B - ribosomal 40s subunit protein s27b  RPS1B - ribosomal 40s subunit protein s1b  RPL17A - ribosomal 60s subunit protein l17a  RPL21A - ribosomal 60s subunit protein l21a  RPP1B - ribosomal protein p1b  RPL14A - ribosomal 60s subunit protein l14a  RPL17B - rpl17bp  RPL32 - ribosomal 60s subunit protein l32  MRPL24 - mitochondrial 54s ribosomal protein yml24/yml14  RPL26B - ribosomal 60s subunit protein l26b  RPL22B - ribosomal 60s subunit protein l22b  RPL35B - ribosomal 60s subunit protein l35b  RPS10A - ribosomal 40s subunit protein s10a  PGK1 - phosphoglycerate kinase  ABZ1 - 4-amino-4-deoxychorismate synthase  RPL34A - ribosomal 60s subunit protein l34a  RPL27A - ribosomal 60s subunit protein l27a  RNR4 - ribonucleotide-diphosphate reductase subunit rnr4  RPS22A - rps22ap  RPS17B - ribosomal 40s subunit protein s17b  YHR020W - proline--trna ligase  PAH1 - phosphatidate phosphatase pah1  RPL6A - ribosomal 60s subunit protein l6a  RPC37 - rpc37p  RPS23B - ribosomal 40s subunit protein s23b  RPL42B - ribosomal 60s subunit protein l42b  APA2 - apa2p  RPS14A - ribosomal 40s subunit protein s14a  RPL31A - ribosomal 60s subunit protein l31a  STM1 - stm1p  CHS3 - chitin synthase chs3  BNA5 - kynureninase  THR4 - threonine synthase thr4  RPS13 - ribosomal 40s subunit protein s13  RPS26A - ribosomal 40s subunit protein s26a  RPS7A - ribosomal 40s subunit protein s7a  RPS24A - ribosomal 40s subunit protein s24a  RPL16B - ribosomal 60s subunit protein l16b |
| GO:0009058 | biosynthetic process | 2.27E-12 | 8.46E-10 | 2.10 (5247,1314,156,82) | [+] Show genes  RPL2B - ribosomal 60s subunit protein l2b  RPS0B - ribosomal 40s subunit protein s0b  RPL29 - ribosomal 60s subunit protein l29  RPL24A - ribosomal 60s subunit protein l24a  RPL30 - ribosomal 60s subunit protein l30  KTR2 - ktr2p  TSL1 - tsl1p  RPL9A - ribosomal 60s subunit protein l9a  RPL40A - ubiquitin-ribosomal 60s subunit protein l40a fusion protein  RPL23B - ribosomal 60s subunit protein l23b  RPL13B - ribosomal 60s subunit protein l13b  RPL38 - ribosomal 60s subunit protein l38  CHS1 - chitin synthase chs1  TEF1 - tef1p  RPS16A - ribosomal 40s subunit protein s16a  RPL8A - ribosomal 60s subunit protein l8a  ILV5 - ketol-acid reductoisomerase  ASC1 - asc1p  RPS8B - ribosomal 40s subunit protein s8b  ATP18 - atp18p  DPB3 - dpb3p  RPL26A - ribosomal 60s subunit protein l26a  RPL11A - ribosomal 60s subunit protein l11a  RPL15A - ribosomal 60s subunit protein l15a  RPP0 - ribosomal protein p0  RPL34B - ribosomal 60s subunit protein l34b  RPL27B - ribosomal 60s subunit protein l27b  RPL3 - ribosomal 60s subunit protein l3  CYT1 - ubiquinol--cytochrome-c reductase catalytic subunit cyt1  RPS18B - ribosomal 40s subunit protein s18b  RPL19A - ribosomal 60s subunit protein l19a  RPS21B - rps21bp  SLX1 - slx1p  RPS10B - ribosomal 40s subunit protein s10b  RPP2B - ribosomal protein p2b  RPL6B - ribosomal 60s subunit protein l6b  RPS15 - ribosomal 40s subunit protein s15  RPP2A - ribosomal protein p2a  RPS5 - rps5p  RPS17A - ribosomal 40s subunit protein s17a  RPL37A - ribosomal 60s subunit protein l37a  RPL24B - ribosomal 60s subunit protein l24b  GSC2 - gsc2p  RPS28A - ribosomal 40s subunit protein s28a  RPS27B - ribosomal 40s subunit protein s27b  RPS1B - ribosomal 40s subunit protein s1b  RPL17A - ribosomal 60s subunit protein l17a  RPL21A - ribosomal 60s subunit protein l21a  RPP1B - ribosomal protein p1b  RPL14A - ribosomal 60s subunit protein l14a  RPL17B - rpl17bp  RPL32 - ribosomal 60s subunit protein l32  MRPL24 - mitochondrial 54s ribosomal protein yml24/yml14  RPL26B - ribosomal 60s subunit protein l26b  RPL22B - ribosomal 60s subunit protein l22b  RPL35B - ribosomal 60s subunit protein l35b  RPS10A - ribosomal 40s subunit protein s10a  PGK1 - phosphoglycerate kinase  ABZ1 - 4-amino-4-deoxychorismate synthase  RPL34A - ribosomal 60s subunit protein l34a  RPL27A - ribosomal 60s subunit protein l27a  RNR4 - ribonucleotide-diphosphate reductase subunit rnr4  RPS22A - rps22ap  RPS17B - ribosomal 40s subunit protein s17b  YHR020W - proline--trna ligase  PAH1 - phosphatidate phosphatase pah1  RPL6A - ribosomal 60s subunit protein l6a  RPS23B - ribosomal 40s subunit protein s23b  RPC37 - rpc37p  RPL42B - ribosomal 60s subunit protein l42b  APA2 - apa2p  RPS14A - ribosomal 40s subunit protein s14a  RPL31A - ribosomal 60s subunit protein l31a  STM1 - stm1p  CHS3 - chitin synthase chs3  BNA5 - kynureninase  THR4 - threonine synthase thr4  RPS13 - ribosomal 40s subunit protein s13  RPS26A - ribosomal 40s subunit protein s26a  RPS7A - ribosomal 40s subunit protein s7a  RPS24A - ribosomal 40s subunit protein s24a  RPL16B - ribosomal 60s subunit protein l16b |
| GO:0000028 | ribosomal small subunit assembly | 1.04E-11 | 3.61E-9 | 11.71 (5247,25,251,14) | [+] Show genes  RPS0B - ribosomal 40s subunit protein s0b  RPS27B - ribosomal 40s subunit protein s27b  RPS14A - ribosomal 40s subunit protein s14a  RPS10A - ribosomal 40s subunit protein s10a  RPS10B - ribosomal 40s subunit protein s10b  NSR1 - nsr1p  RPS11B - ribosomal 40s subunit protein s11b  RPS15 - ribosomal 40s subunit protein s15  RPS0A - ribosomal 40s subunit protein s0a  RPS5 - rps5p  RPS17A - ribosomal 40s subunit protein s17a  RPS17B - ribosomal 40s subunit protein s17b  RPS28A - ribosomal 40s subunit protein s28a  PWP2 - pwp2p |
| GO:0022613 | ribonucleoprotein complex biogenesis | 5.62E-11 | 1.83E-8 | 1.70 (5247,221,1630,117) | [+] Show genes  NSA2 - nsa2p  ECM16 - ecm16p  LCP5 - lcp5p  FAF1 - faf1p  RRB1 - rrb1p  UTP30 - utp30p  KRR1 - krr1p  RPL8A - ribosomal 60s subunit protein l8a  NOP7 - nop7p  UTP10 - utp10p  UTP14 - utp14p  MDV1 - mdv1p  KRI1 - kri1p  RPA190 - rpa190p  ALB1 - alb1p  CAM1 - cam1p  RPL34B - ribosomal 60s subunit protein l34b  RPS6A - ribosomal 40s subunit protein s6a  RPS31 - ubiquitin-ribosomal 40s subunit protein s31 fusion protein  RPO26 - rpo26p  UTP8 - utp8p  RIX1 - rix1p  RLP7 - rlp7p  UTP9 - utp9p  NSA1 - nsa1p  CBF5 - pseudouridine synthase cbf5  RPS2 - ribosomal 40s subunit protein s2  BUD21 - bud21p  RPA34 - rpa34p  JIP5 - jip5p  ENP2 - enp2p  RPS9A - ribosomal 40s subunit protein s9a  NOP58 - nop58p  RRP36 - rrp36p  RPL14A - ribosomal 60s subunit protein l14a  HCA4 - hca4p  TIF4631 - tif4631p  RRS1 - rrs1p  YDR161W - hypothetical protein  RPL7A - ribosomal 60s subunit protein l7a  RPL34A - ribosomal 60s subunit protein l34a  MAK11 - mak11p  RPA14 - rpa14p  DBP9 - dbp9p  EBP2 - ebp2p  DRS1 - drs1p  YTM1 - ytm1p  RCL1 - rcl1p  RPC10 - rpc10p  NOG2 - nog2p  RPS14A - ribosomal 40s subunit protein s14a  UTP5 - utp5p  ECM1 - ecm1p  SDO1 - sdo1p  RPS0A - ribosomal 40s subunit protein s0a  RPL33B - ribosomal 60s subunit protein l33b  RIO2 - rio2p  RPC40 - rpc40p  NOP53 - nop53p  TRM112 - trm112p  DBP7 - dbp7p  RPL33A - ribosomal 60s subunit protein l33a  PWP2 - pwp2p  RPS0B - ribosomal 40s subunit protein s0b  RNH70 - rnh70p  IPI3 - ipi3p  RPL40A - ubiquitin-ribosomal 60s subunit protein l40a fusion protein  RPF1 - rpf1p  RTC3 - rtc3p  RSA4 - rsa4p  PXR1 - pxr1p  RRP8 - rrp8p  NOP16 - nop16p  GAR1 - gar1p  REI1 - rei1p  NUG1 - nug1p  BUD22 - bud22p  SSF1 - ssf1p  DHR2 - dhr2p  RLI1 - rli1p  RIA1 - ria1p  BRX1 - brx1p  NOC2 - noc2p  UTP20 - utp20p  RPL26A - ribosomal 60s subunit protein l26a  MRPL11 - mitochondrial 54s ribosomal protein yml11  RPP0 - ribosomal protein p0  CIC1 - cic1p  RRP3 - rna-dependent atpase rrp3  SDA1 - sda1p  RPS15 - ribosomal 40s subunit protein s15  ESF1 - esf1p  NOP9 - nop9p  MPP10 - mpp10p  HRR25 - hrr25p  RPC19 - rpc19p  DBP2 - dbp2p  RPL7B - ribosomal 60s subunit protein l7b  NOP15 - nop15p  RPL26B - ribosomal 60s subunit protein l26b  NIP7 - nip7p  PNO1 - pno1p  RPS14B - rps14bp  BMS1 - bms1p  SAS10 - sas10p  NEW1 - new1p  NOP14 - nop14p  SLX9 - slx9p  HAS1 - atp-dependent rna helicase has1  RIO1 - rio1p  ROK1 - rna-dependent atpase rok1  UTP11 - utp11p  MAK21 - mak21p  RRP12 - rrp12p  REH1 - reh1p  RPA12 - rpa12p  RPS7A - ribosomal 40s subunit protein s7a |
| GO:0044085 | cellular component biogenesis | 7.64E-11 | 2.34E-8 | 1.66 (5247,244,1630,126) | [+] Show genes  ECM16 - ecm16p  NSA2 - nsa2p  LCP5 - lcp5p  FAF1 - faf1p  RRB1 - rrb1p  UTP30 - utp30p  KRR1 - krr1p  RPL8A - ribosomal 60s subunit protein l8a  NOP7 - nop7p  UTP10 - utp10p  UTP14 - utp14p  MDV1 - mdv1p  KRI1 - kri1p  RPA190 - rpa190p  ALB1 - alb1p  CAM1 - cam1p  RPL34B - ribosomal 60s subunit protein l34b  RPS6A - ribosomal 40s subunit protein s6a  RPS31 - ubiquitin-ribosomal 40s subunit protein s31 fusion protein  RPO26 - rpo26p  UTP8 - utp8p  RIX1 - rix1p  RLP7 - rlp7p  UTP9 - utp9p  NSA1 - nsa1p  CBF5 - pseudouridine synthase cbf5  RPS2 - ribosomal 40s subunit protein s2  BUD21 - bud21p  RPA34 - rpa34p  JIP5 - jip5p  ENP2 - enp2p  RPS9A - ribosomal 40s subunit protein s9a  NOP58 - nop58p  RRP36 - rrp36p  RPL14A - ribosomal 60s subunit protein l14a  HCA4 - hca4p  TIF4631 - tif4631p  RRS1 - rrs1p  BIG1 - big1p  YDR161W - hypothetical protein  RPL7A - ribosomal 60s subunit protein l7a  MAK11 - mak11p  RPL34A - ribosomal 60s subunit protein l34a  RPA14 - rpa14p  DBP9 - dbp9p  EBP2 - ebp2p  DRS1 - drs1p  YTM1 - ytm1p  RCL1 - rcl1p  RPC10 - rpc10p  NOG2 - nog2p  RPS14A - ribosomal 40s subunit protein s14a  UTP5 - utp5p  ECM1 - ecm1p  SDO1 - sdo1p  TGS1 - tgs1p  RPS0A - ribosomal 40s subunit protein s0a  RPL33B - ribosomal 60s subunit protein l33b  RIO2 - rio2p  RPC40 - rpc40p  NOP53 - nop53p  TRM112 - trm112p  DBP7 - dbp7p  RPL33A - ribosomal 60s subunit protein l33a  PWP2 - pwp2p  RPS0B - ribosomal 40s subunit protein s0b  LRG1 - lrg1p  RNH70 - rnh70p  IPI3 - ipi3p  CWH41 - cwh41p  RPL40A - ubiquitin-ribosomal 60s subunit protein l40a fusion protein  RPF1 - rpf1p  RRP8 - rrp8p  PXR1 - pxr1p  RSA4 - rsa4p  RTC3 - rtc3p  NOP16 - nop16p  GAR1 - gar1p  REI1 - rei1p  BUD22 - bud22p  NUG1 - nug1p  SSF1 - ssf1p  DHR2 - dhr2p  RLI1 - rli1p  RPI1 - rpi1p  BRX1 - brx1p  RIA1 - ria1p  NOC2 - noc2p  UTP20 - utp20p  RPL26A - ribosomal 60s subunit protein l26a  MRPL11 - mitochondrial 54s ribosomal protein yml11  RPP0 - ribosomal protein p0  DFG5 - dfg5p  CIC1 - cic1p  INN1 - inn1p  RRP3 - rna-dependent atpase rrp3  SDA1 - sda1p  RPS15 - ribosomal 40s subunit protein s15  ESF1 - esf1p  NOP9 - nop9p  MPP10 - mpp10p  HRR25 - hrr25p  RPC19 - rpc19p  DBP2 - dbp2p  RPL7B - ribosomal 60s subunit protein l7b  NOP15 - nop15p  RPL26B - ribosomal 60s subunit protein l26b  NIP7 - nip7p  SLT2 - slt2p  PNO1 - pno1p  RPS14B - rps14bp  BMS1 - bms1p  SAS10 - sas10p  NEW1 - new1p  NOP14 - nop14p  SLX9 - slx9p  HAS1 - atp-dependent rna helicase has1  RIO1 - rio1p  ROK1 - rna-dependent atpase rok1  UTP11 - utp11p  PKC1 - pkc1p  MAK21 - mak21p  RRP12 - rrp12p  REH1 - reh1p  RPA12 - rpa12p  RPS7A - ribosomal 40s subunit protein s7a |
| GO:0034641 | cellular nitrogen compound metabolic process | 1.51E-10 | 4.38E-8 | 1.80 (5247,1734,156,93) | [+] Show genes  RPL2B - ribosomal 60s subunit protein l2b  MGM101 - mgm101p  RPS0B - ribosomal 40s subunit protein s0b  ECM16 - ecm16p  RPL29 - ribosomal 60s subunit protein l29  RPL24A - ribosomal 60s subunit protein l24a  RPL30 - ribosomal 60s subunit protein l30  VMA2 - vma2p  RRB1 - rrb1p  EFG1 - efg1p  RPL9A - ribosomal 60s subunit protein l9a  RPL40A - ubiquitin-ribosomal 60s subunit protein l40a fusion protein  MSS51 - mss51p  RPL23B - ribosomal 60s subunit protein l23b  RPL13B - ribosomal 60s subunit protein l13b  RPL38 - ribosomal 60s subunit protein l38  RPS16A - ribosomal 40s subunit protein s16a  TEF1 - tef1p  RPL8A - ribosomal 60s subunit protein l8a  UTP14 - utp14p  RAD9 - rad9p  RPS8B - ribosomal 40s subunit protein s8b  ATP18 - atp18p  DPB3 - dpb3p  RPL26A - ribosomal 60s subunit protein l26a  IRE1 - ire1p  RPL15A - ribosomal 60s subunit protein l15a  RPL11A - ribosomal 60s subunit protein l11a  RPP0 - ribosomal protein p0  FUM1 - fumarase fum1  RPL34B - ribosomal 60s subunit protein l34b  RPL27B - ribosomal 60s subunit protein l27b  RPL3 - ribosomal 60s subunit protein l3  CYT1 - ubiquinol--cytochrome-c reductase catalytic subunit cyt1  RPS18B - ribosomal 40s subunit protein s18b  RPL19A - ribosomal 60s subunit protein l19a  RPS21B - rps21bp  SLX1 - slx1p  SMB1 - smb1p  RPS10B - ribosomal 40s subunit protein s10b  RPP2B - ribosomal protein p2b  RLP7 - rlp7p  RPL6B - ribosomal 60s subunit protein l6b  RPS15 - ribosomal 40s subunit protein s15  RPP2A - ribosomal protein p2a  DUR1,2 - bifunctional urea carboxylase/allophanate hydrolase  RPS5 - rps5p  RPL37A - ribosomal 60s subunit protein l37a  RPS17A - ribosomal 40s subunit protein s17a  RPL24B - ribosomal 60s subunit protein l24b  RPS28A - ribosomal 40s subunit protein s28a  RPL17A - ribosomal 60s subunit protein l17a  RPS1B - ribosomal 40s subunit protein s1b  RPS27B - ribosomal 40s subunit protein s27b  RPL14A - ribosomal 60s subunit protein l14a  RPL21A - ribosomal 60s subunit protein l21a  RPP1B - ribosomal protein p1b  RPL17B - rpl17bp  HCA4 - hca4p  RPL32 - ribosomal 60s subunit protein l32  MRPL24 - mitochondrial 54s ribosomal protein yml24/yml14  RPL26B - ribosomal 60s subunit protein l26b  RPL22B - ribosomal 60s subunit protein l22b  RPL35B - ribosomal 60s subunit protein l35b  RPS10A - ribosomal 40s subunit protein s10a  ACT1 - actin  NSR1 - nsr1p  NCS2 - ncs2p  ABZ1 - 4-amino-4-deoxychorismate synthase  PGK1 - phosphoglycerate kinase  RPL34A - ribosomal 60s subunit protein l34a  RPL27A - ribosomal 60s subunit protein l27a  RNR4 - ribonucleotide-diphosphate reductase subunit rnr4  SRD1 - srd1p  RPS22A - rps22ap  RPS17B - ribosomal 40s subunit protein s17b  YHR020W - proline--trna ligase  RPL6A - ribosomal 60s subunit protein l6a  RPC37 - rpc37p  RPS23B - ribosomal 40s subunit protein s23b  RPL42B - ribosomal 60s subunit protein l42b  APA2 - apa2p  RPS14A - ribosomal 40s subunit protein s14a  RPL31A - ribosomal 60s subunit protein l31a  STM1 - stm1p  SUA5 - sua5p  RIO1 - rio1p  BNA5 - kynureninase  RPS13 - ribosomal 40s subunit protein s13  RPS26A - ribosomal 40s subunit protein s26a  RPS7A - ribosomal 40s subunit protein s7a  RPS24A - ribosomal 40s subunit protein s24a  RPL16B - ribosomal 60s subunit protein l16b |
| GO:0042254 | ribosome biogenesis | 1.56E-10 | 4.27E-8 | 1.94 (5247,194,1201,86) | [+] Show genes  RPS0B - ribosomal 40s subunit protein s0b  ECM16 - ecm16p  NSA2 - nsa2p  FAF1 - faf1p  LCP5 - lcp5p  RRB1 - rrb1p  UTP30 - utp30p  KRR1 - krr1p  RPL40A - ubiquitin-ribosomal 60s subunit protein l40a fusion protein  RPF1 - rpf1p  PXR1 - pxr1p  RTC3 - rtc3p  NOP16 - nop16p  NOP7 - nop7p  RPL8A - ribosomal 60s subunit protein l8a  GAR1 - gar1p  UTP14 - utp14p  UTP10 - utp10p  REI1 - rei1p  NUG1 - nug1p  DHR2 - dhr2p  SSF1 - ssf1p  RLI1 - rli1p  RIA1 - ria1p  KRI1 - kri1p  NOC2 - noc2p  UTP20 - utp20p  MRPL11 - mitochondrial 54s ribosomal protein yml11  ALB1 - alb1p  RPP0 - ribosomal protein p0  CAM1 - cam1p  RPL34B - ribosomal 60s subunit protein l34b  RPS6A - ribosomal 40s subunit protein s6a  CIC1 - cic1p  RRP3 - rna-dependent atpase rrp3  RPS31 - ubiquitin-ribosomal 40s subunit protein s31 fusion protein  UTP8 - utp8p  RPO26 - rpo26p  SDA1 - sda1p  UTP9 - utp9p  RLP7 - rlp7p  NSA1 - nsa1p  RPS15 - ribosomal 40s subunit protein s15  ESF1 - esf1p  CBF5 - pseudouridine synthase cbf5  RPS2 - ribosomal 40s subunit protein s2  BUD21 - bud21p  MPP10 - mpp10p  ENP2 - enp2p  RPS9A - ribosomal 40s subunit protein s9a  NOP58 - nop58p  DBP2 - dbp2p  HCA4 - hca4p  RRS1 - rrs1p  NIP7 - nip7p  YDR161W - hypothetical protein  PNO1 - pno1p  RPL34A - ribosomal 60s subunit protein l34a  RPA14 - rpa14p  DBP9 - dbp9p  RPS14B - rps14bp  BMS1 - bms1p  EBP2 - ebp2p  SAS10 - sas10p  DRS1 - drs1p  YTM1 - ytm1p  NOP14 - nop14p  NOG2 - nog2p  RPS14A - ribosomal 40s subunit protein s14a  UTP5 - utp5p  SLX9 - slx9p  ECM1 - ecm1p  SDO1 - sdo1p  RIO1 - rio1p  HAS1 - atp-dependent rna helicase has1  ROK1 - rna-dependent atpase rok1  UTP11 - utp11p  RPS0A - ribosomal 40s subunit protein s0a  MAK21 - mak21p  RRP12 - rrp12p  RIO2 - rio2p  NOP53 - nop53p  DBP7 - dbp7p  RPA12 - rpa12p  RPS7A - ribosomal 40s subunit protein s7a  PWP2 - pwp2p |
| GO:0030490 | maturation of SSU-rRNA | 3.04E-10 | 7.93E-8 | 2.55 (5247,80,1184,46) | [+] Show genes  ECM16 - ecm16p  RPS27B - ribosomal 40s subunit protein s27b  RPS1B - ribosomal 40s subunit protein s1b  FAF1 - faf1p  LCP5 - lcp5p  EFG1 - efg1p  UTP30 - utp30p  RPS28B - ribosomal 40s subunit protein s28b  RPS16A - ribosomal 40s subunit protein s16a  NOP7 - nop7p  RPS11B - ribosomal 40s subunit protein s11b  UTP10 - utp10p  BUD22 - bud22p  DHR2 - dhr2p  RPS14B - rps14bp  RPS27A - ribosomal 40s subunit protein s27a  BMS1 - bms1p  RPS8B - ribosomal 40s subunit protein s8b  SAS10 - sas10p  NOP14 - nop14p  RPS23B - ribosomal 40s subunit protein s23b  RPS14A - ribosomal 40s subunit protein s14a  RPS1A - ribosomal 40s subunit protein s1a  UTP5 - utp5p  RPS6A - ribosomal 40s subunit protein s6a  RRP3 - rna-dependent atpase rrp3  SLX9 - slx9p  RIO1 - rio1p  HAS1 - atp-dependent rna helicase has1  RPS31 - ubiquitin-ribosomal 40s subunit protein s31 fusion protein  UTP8 - utp8p  ROK1 - rna-dependent atpase rok1  UTP9 - utp9p  RRP12 - rrp12p  RPS20 - ribosomal 40s subunit protein s20  BUD21 - bud21p  FYV7 - fyv7p  RPS13 - ribosomal 40s subunit protein s13  RIO2 - rio2p  TSR2 - tsr2p  TRM112 - trm112p  ENP2 - enp2p  RPS9A - ribosomal 40s subunit protein s9a  RPS24A - ribosomal 40s subunit protein s24a  PWP2 - pwp2p  RPS28A - ribosomal 40s subunit protein s28a |
| GO:0000462 | maturation of SSU-rRNA from tricistronic rRNA transcript (SSU-rRNA, 5.8S rRNA, LSU-rRNA) | 3.89E-10 | 9.65E-8 | 2.97 (5247,67,974,37) | [+] Show genes  ECM16 - ecm16p  RPS27B - ribosomal 40s subunit protein s27b  RPS1B - ribosomal 40s subunit protein s1b  FAF1 - faf1p  LCP5 - lcp5p  EFG1 - efg1p  UTP30 - utp30p  RPS16A - ribosomal 40s subunit protein s16a  NOP7 - nop7p  UTP10 - utp10p  RPS11B - ribosomal 40s subunit protein s11b  DHR2 - dhr2p  RPS14B - rps14bp  BMS1 - bms1p  RPS27A - ribosomal 40s subunit protein s27a  RPS8B - ribosomal 40s subunit protein s8b  SAS10 - sas10p  NOP14 - nop14p  RPS23B - ribosomal 40s subunit protein s23b  RPS14A - ribosomal 40s subunit protein s14a  RPS1A - ribosomal 40s subunit protein s1a  RPS6A - ribosomal 40s subunit protein s6a  RRP3 - rna-dependent atpase rrp3  SLX9 - slx9p  HAS1 - atp-dependent rna helicase has1  RIO1 - rio1p  UTP8 - utp8p  UTP9 - utp9p  RRP12 - rrp12p  RPS20 - ribosomal 40s subunit protein s20  FYV7 - fyv7p  BUD21 - bud21p  RPS13 - ribosomal 40s subunit protein s13  TSR2 - tsr2p  RPS9A - ribosomal 40s subunit protein s9a  RPS24A - ribosomal 40s subunit protein s24a  PWP2 - pwp2p |
| GO:0019538 | protein metabolic process | 9.66E-10 | 2.29E-7 | 2.01 (5247,1247,155,74) | [+] Show genes  RPL2B - ribosomal 60s subunit protein l2b  RPS0B - ribosomal 40s subunit protein s0b  RPL29 - ribosomal 60s subunit protein l29  RPL24A - ribosomal 60s subunit protein l24a  RPL30 - ribosomal 60s subunit protein l30  PFA3 - pfa3p  KTR2 - ktr2p  RPL9A - ribosomal 60s subunit protein l9a  RPL40A - ubiquitin-ribosomal 60s subunit protein l40a fusion protein  RPL23B - ribosomal 60s subunit protein l23b  RPL38 - ribosomal 60s subunit protein l38  RPL13B - ribosomal 60s subunit protein l13b  TEF1 - tef1p  RPS16A - ribosomal 40s subunit protein s16a  RPL8A - ribosomal 60s subunit protein l8a  PPT1 - ppt1p  RPS8B - ribosomal 40s subunit protein s8b  RPL26A - ribosomal 60s subunit protein l26a  IRE1 - ire1p  RPL11A - ribosomal 60s subunit protein l11a  RPL15A - ribosomal 60s subunit protein l15a  RPP0 - ribosomal protein p0  RPL34B - ribosomal 60s subunit protein l34b  RPL27B - ribosomal 60s subunit protein l27b  RPL3 - ribosomal 60s subunit protein l3  RPS18B - ribosomal 40s subunit protein s18b  RPL19A - ribosomal 60s subunit protein l19a  RPS21B - rps21bp  RPS10B - ribosomal 40s subunit protein s10b  RPP2B - ribosomal protein p2b  FPR4 - peptidylprolyl isomerase fpr4  RPL6B - ribosomal 60s subunit protein l6b  RPS15 - ribosomal 40s subunit protein s15  RPP2A - ribosomal protein p2a  RPS5 - rps5p  RPL37A - ribosomal 60s subunit protein l37a  RPS17A - ribosomal 40s subunit protein s17a  RPL24B - ribosomal 60s subunit protein l24b  RPS28A - ribosomal 40s subunit protein s28a  RPS1B - ribosomal 40s subunit protein s1b  RPL17A - ribosomal 60s subunit protein l17a  RPS27B - ribosomal 40s subunit protein s27b  RPL21A - ribosomal 60s subunit protein l21a  RPP1B - ribosomal protein p1b  RPL14A - ribosomal 60s subunit protein l14a  RPL17B - rpl17bp  RPL32 - ribosomal 60s subunit protein l32  MRPL24 - mitochondrial 54s ribosomal protein yml24/yml14  RPL26B - ribosomal 60s subunit protein l26b  RPL22B - ribosomal 60s subunit protein l22b  PRB1 - prb1p  RPL35B - ribosomal 60s subunit protein l35b  SLT2 - slt2p  RPS10A - ribosomal 40s subunit protein s10a  ACT1 - actin  NCS2 - ncs2p  RPL34A - ribosomal 60s subunit protein l34a  SSE1 - sse1p  RPL27A - ribosomal 60s subunit protein l27a  RPS22A - rps22ap  RPS17B - ribosomal 40s subunit protein s17b  YHR020W - proline--trna ligase  RPL6A - ribosomal 60s subunit protein l6a  RPS23B - ribosomal 40s subunit protein s23b  RPL42B - ribosomal 60s subunit protein l42b  RPS14A - ribosomal 40s subunit protein s14a  RPL31A - ribosomal 60s subunit protein l31a  HSL1 - hsl1p  RIO1 - rio1p  RPS13 - ribosomal 40s subunit protein s13  RPS26A - ribosomal 40s subunit protein s26a  RPS7A - ribosomal 40s subunit protein s7a  RPS24A - ribosomal 40s subunit protein s24a  RPL16B - ribosomal 60s subunit protein l16b |
| GO:0006364 | rRNA processing | 1.2E-9 | 2.71E-7 | 1.87 (5247,256,1009,92) | [+] Show genes  RPS0B - ribosomal 40s subunit protein s0b  ECM16 - ecm16p  NSA2 - nsa2p  FAF1 - faf1p  LCP5 - lcp5p  RPL30 - ribosomal 60s subunit protein l30  RRB1 - rrb1p  EFG1 - efg1p  UTP30 - utp30p  ZUO1 - zuo1p  RRP1 - rrp1p  KRR1 - krr1p  RPF1 - rpf1p  RTC3 - rtc3p  PXR1 - pxr1p  NOP16 - nop16p  RPS16A - ribosomal 40s subunit protein s16a  NOP7 - nop7p  RPL8A - ribosomal 60s subunit protein l8a  GAR1 - gar1p  UTP14 - utp14p  UTP10 - utp10p  RAT1 - rat1p  BUD22 - bud22p  NUG1 - nug1p  SSF1 - ssf1p  DHR2 - dhr2p  KRI1 - kri1p  RPS8B - ribosomal 40s subunit protein s8b  RPS21A - ribosomal 40s subunit protein s21a  UTP20 - utp20p  RPS1A - ribosomal 40s subunit protein s1a  RPL3 - ribosomal 60s subunit protein l3  RPS6A - ribosomal 40s subunit protein s6a  CIC1 - cic1p  RRP3 - rna-dependent atpase rrp3  RPS21B - rps21bp  UTP8 - utp8p  TMA64 - tma64p  RIB2 - bifunctional drap deaminase/trna pseudouridine synthase rib2  UTP9 - utp9p  RLP7 - rlp7p  NSA1 - nsa1p  ESF1 - esf1p  CBF5 - pseudouridine synthase cbf5  RPS2 - ribosomal 40s subunit protein s2  FYV7 - fyv7p  BUD21 - bud21p  MPP10 - mpp10p  TSR2 - tsr2p  RPS9A - ribosomal 40s subunit protein s9a  RPS28A - ribosomal 40s subunit protein s28a  RPS1B - ribosomal 40s subunit protein s1b  RPS27B - ribosomal 40s subunit protein s27b  DBP2 - dbp2p  HCA4 - hca4p  RPS28B - ribosomal 40s subunit protein s28b  RPL35B - ribosomal 60s subunit protein l35b  RPL7A - ribosomal 60s subunit protein l7a  PNO1 - pno1p  NSR1 - nsr1p  RPS11B - ribosomal 40s subunit protein s11b  SRD1 - srd1p  DBP9 - dbp9p  RPS14B - rps14bp  RPS27A - ribosomal 40s subunit protein s27a  BMS1 - bms1p  EBP2 - ebp2p  SAS10 - sas10p  DRS1 - drs1p  YTM1 - ytm1p  NOP14 - nop14p  RPS23B - ribosomal 40s subunit protein s23b  RPS14A - ribosomal 40s subunit protein s14a  SLX9 - slx9p  HAS1 - atp-dependent rna helicase has1  RIO1 - rio1p  RSC9 - rsc9p  UTP11 - utp11p  YBR141C - hypothetical protein  YIL096C - hypothetical protein  RPS0A - ribosomal 40s subunit protein s0a  RRP12 - rrp12p  RPS20 - ribosomal 40s subunit protein s20  RPS13 - ribosomal 40s subunit protein s13  BUD23 - bud23p  RRP46 - rrp46p  DBP7 - dbp7p  RPS7A - ribosomal 40s subunit protein s7a  RPS24A - ribosomal 40s subunit protein s24a  MRD1 - mrd1p  PWP2 - pwp2p |
| GO:0006407 | rRNA export from nucleus | 3.49E-9 | 7.56E-7 | 13.48 (5247,17,229,10) | [+] Show genes  RPS15 - ribosomal 40s subunit protein s15  RPS0B - ribosomal 40s subunit protein s0b  RPS0A - ribosomal 40s subunit protein s0a  RPS2 - ribosomal 40s subunit protein s2  RPS5 - rps5p  RPS18B - ribosomal 40s subunit protein s18b  RPS26A - ribosomal 40s subunit protein s26a  RPS10A - ribosomal 40s subunit protein s10a  RPS10B - ribosomal 40s subunit protein s10b  RPS28A - ribosomal 40s subunit protein s28a |
| GO:0051029 | rRNA transport | 3.49E-9 | 7.26E-7 | 13.48 (5247,17,229,10) | [+] Show genes  RPS15 - ribosomal 40s subunit protein s15  RPS0B - ribosomal 40s subunit protein s0b  RPS0A - ribosomal 40s subunit protein s0a  RPS2 - ribosomal 40s subunit protein s2  RPS5 - rps5p  RPS18B - ribosomal 40s subunit protein s18b  RPS26A - ribosomal 40s subunit protein s26a  RPS10A - ribosomal 40s subunit protein s10a  RPS10B - ribosomal 40s subunit protein s10b  RPS28A - ribosomal 40s subunit protein s28a |
| GO:0044260 | cellular macromolecule metabolic process | 4.14E-8 | 8.29E-6 | 1.70 (5247,1744,156,88) | [+] Show genes  RPL2B - ribosomal 60s subunit protein l2b  MGM101 - mgm101p  RPL29 - ribosomal 60s subunit protein l29  RPS0B - ribosomal 40s subunit protein s0b  RPL24A - ribosomal 60s subunit protein l24a  RPL30 - ribosomal 60s subunit protein l30  EFG1 - efg1p  PFA3 - pfa3p  KTR2 - ktr2p  RPL9A - ribosomal 60s subunit protein l9a  RPL40A - ubiquitin-ribosomal 60s subunit protein l40a fusion protein  RPL23B - ribosomal 60s subunit protein l23b  RPL38 - ribosomal 60s subunit protein l38  RPL13B - ribosomal 60s subunit protein l13b  RPS16A - ribosomal 40s subunit protein s16a  TEF1 - tef1p  RPL8A - ribosomal 60s subunit protein l8a  ASC1 - asc1p  RAD9 - rad9p  PPT1 - ppt1p  RPS8B - ribosomal 40s subunit protein s8b  DPB3 - dpb3p  RPL26A - ribosomal 60s subunit protein l26a  IRE1 - ire1p  RPL15A - ribosomal 60s subunit protein l15a  RPL11A - ribosomal 60s subunit protein l11a  RPP0 - ribosomal protein p0  FUM1 - fumarase fum1  RPL34B - ribosomal 60s subunit protein l34b  RPL27B - ribosomal 60s subunit protein l27b  RPL3 - ribosomal 60s subunit protein l3  RPS18B - ribosomal 40s subunit protein s18b  RPL19A - ribosomal 60s subunit protein l19a  RPS21B - rps21bp  SLX1 - slx1p  RPS10B - ribosomal 40s subunit protein s10b  FPR4 - peptidylprolyl isomerase fpr4  RPP2B - ribosomal protein p2b  RPL6B - ribosomal 60s subunit protein l6b  RPS15 - ribosomal 40s subunit protein s15  RPP2A - ribosomal protein p2a  RPS5 - rps5p  RPL37A - ribosomal 60s subunit protein l37a  RPS17A - ribosomal 40s subunit protein s17a  GSC2 - gsc2p  RPL24B - ribosomal 60s subunit protein l24b  RPS28A - ribosomal 40s subunit protein s28a  RPL17A - ribosomal 60s subunit protein l17a  RPS1B - ribosomal 40s subunit protein s1b  RPS27B - ribosomal 40s subunit protein s27b  RPL14A - ribosomal 60s subunit protein l14a  RPL21A - ribosomal 60s subunit protein l21a  RPP1B - ribosomal protein p1b  RPL17B - rpl17bp  MRPL24 - mitochondrial 54s ribosomal protein yml24/yml14  RPL32 - ribosomal 60s subunit protein l32  RPL26B - ribosomal 60s subunit protein l26b  RPL22B - ribosomal 60s subunit protein l22b  PRB1 - prb1p  RPL35B - ribosomal 60s subunit protein l35b  SLT2 - slt2p  RPS10A - ribosomal 40s subunit protein s10a  ACT1 - actin  NCS2 - ncs2p  RPL34A - ribosomal 60s subunit protein l34a  SSE1 - sse1p  RPL27A - ribosomal 60s subunit protein l27a  RNR4 - ribonucleotide-diphosphate reductase subunit rnr4  RPS22A - rps22ap  RPS17B - ribosomal 40s subunit protein s17b  YHR020W - proline--trna ligase  RPL6A - ribosomal 60s subunit protein l6a  RPC37 - rpc37p  RPS23B - ribosomal 40s subunit protein s23b  RPL42B - ribosomal 60s subunit protein l42b  APA2 - apa2p  RPS14A - ribosomal 40s subunit protein s14a  RPL31A - ribosomal 60s subunit protein l31a  STM1 - stm1p  HSL1 - hsl1p  SUA5 - sua5p  RIO1 - rio1p  CHS3 - chitin synthase chs3  RPS13 - ribosomal 40s subunit protein s13  RPS26A - ribosomal 40s subunit protein s26a  RPS7A - ribosomal 40s subunit protein s7a  RPL16B - ribosomal 60s subunit protein l16b  RPS24A - ribosomal 40s subunit protein s24a |
| GO:0022618 | ribonucleoprotein complex assembly | 1.3E-7 | 2.51E-5 | 3.90 (5247,176,176,23) | [+] Show genes  RPS27B - ribosomal 40s subunit protein s27b  PRP45 - prp45p  RPS0B - ribosomal 40s subunit protein s0b  RPL6A - ribosomal 60s subunit protein l6a  RPL24A - ribosomal 60s subunit protein l24a  RPL3 - ribosomal 60s subunit protein l3  RPS14A - ribosomal 40s subunit protein s14a  RPL40A - ubiquitin-ribosomal 60s subunit protein l40a fusion protein  RPS10A - ribosomal 40s subunit protein s10a  RPS10B - ribosomal 40s subunit protein s10b  RPL6B - ribosomal 60s subunit protein l6b  NSR1 - nsr1p  RPS15 - ribosomal 40s subunit protein s15  RPL11B - ribosomal 60s subunit protein l11b  RPS5 - rps5p  RPS17A - ribosomal 40s subunit protein s17a  RPS26A - ribosomal 40s subunit protein s26a  RPS17B - ribosomal 40s subunit protein s17b  RPL11A - ribosomal 60s subunit protein l11a  RPP0 - ribosomal protein p0  RPL24B - ribosomal 60s subunit protein l24b  RPS28A - ribosomal 40s subunit protein s28a  PWP2 - pwp2p |
| GO:1901564 | organonitrogen compound metabolic process | 1.39E-7 | 2.59E-5 | 1.70 (5247,1762,144,82) | [+] Show genes  RPL2B - ribosomal 60s subunit protein l2b  RPS0B - ribosomal 40s subunit protein s0b  RPL29 - ribosomal 60s subunit protein l29  RPL30 - ribosomal 60s subunit protein l30  VMA2 - vma2p  PFA3 - pfa3p  KTR2 - ktr2p  RPL40A - ubiquitin-ribosomal 60s subunit protein l40a fusion protein  RPL23B - ribosomal 60s subunit protein l23b  RPL38 - ribosomal 60s subunit protein l38  RPL13B - ribosomal 60s subunit protein l13b  CHS1 - chitin synthase chs1  RPS16A - ribosomal 40s subunit protein s16a  TEF1 - tef1p  RPL8A - ribosomal 60s subunit protein l8a  ILV5 - ketol-acid reductoisomerase  PPT1 - ppt1p  RPS8B - ribosomal 40s subunit protein s8b  ATP18 - atp18p  RPL26A - ribosomal 60s subunit protein l26a  RPL15A - ribosomal 60s subunit protein l15a  RPL11A - ribosomal 60s subunit protein l11a  RPP0 - ribosomal protein p0  RPL34B - ribosomal 60s subunit protein l34b  RPL27B - ribosomal 60s subunit protein l27b  RPL3 - ribosomal 60s subunit protein l3  CYT1 - ubiquinol--cytochrome-c reductase catalytic subunit cyt1  RPS18B - ribosomal 40s subunit protein s18b  RPL19A - ribosomal 60s subunit protein l19a  RPS21B - rps21bp  RPS10B - ribosomal 40s subunit protein s10b  FPR4 - peptidylprolyl isomerase fpr4  RPP2B - ribosomal protein p2b  RPL6B - ribosomal 60s subunit protein l6b  RPS15 - ribosomal 40s subunit protein s15  RPP2A - ribosomal protein p2a  DUR1,2 - bifunctional urea carboxylase/allophanate hydrolase  RPS5 - rps5p  RPL37A - ribosomal 60s subunit protein l37a  RPS17A - ribosomal 40s subunit protein s17a  RPL24B - ribosomal 60s subunit protein l24b  RPS28A - ribosomal 40s subunit protein s28a  RPL17A - ribosomal 60s subunit protein l17a  RPS1B - ribosomal 40s subunit protein s1b  RPS27B - ribosomal 40s subunit protein s27b  RPL14A - ribosomal 60s subunit protein l14a  RPL21A - ribosomal 60s subunit protein l21a  RPP1B - ribosomal protein p1b  RPL17B - rpl17bp  RPL32 - ribosomal 60s subunit protein l32  MRPL24 - mitochondrial 54s ribosomal protein yml24/yml14  RPL26B - ribosomal 60s subunit protein l26b  RPL22B - ribosomal 60s subunit protein l22b  PRB1 - prb1p  SLT2 - slt2p  RPS10A - ribosomal 40s subunit protein s10a  ACT1 - actin  NCS2 - ncs2p  ABZ1 - 4-amino-4-deoxychorismate synthase  PGK1 - phosphoglycerate kinase  RPL34A - ribosomal 60s subunit protein l34a  SSE1 - sse1p  RPL27A - ribosomal 60s subunit protein l27a  RPS22A - rps22ap  RPS17B - ribosomal 40s subunit protein s17b  YHR020W - proline--trna ligase  RPL6A - ribosomal 60s subunit protein l6a  RPS23B - ribosomal 40s subunit protein s23b  RPL42B - ribosomal 60s subunit protein l42b  APA2 - apa2p  RPS14A - ribosomal 40s subunit protein s14a  RPL31A - ribosomal 60s subunit protein l31a  HSL1 - hsl1p  RIO1 - rio1p  CHS3 - chitin synthase chs3  BNA5 - kynureninase  THR4 - threonine synthase thr4  RPS13 - ribosomal 40s subunit protein s13  RPS26A - ribosomal 40s subunit protein s26a  RPS7A - ribosomal 40s subunit protein s7a  RPL16B - ribosomal 60s subunit protein l16b  RPS24A - ribosomal 40s subunit protein s24a |
| GO:0016072 | rRNA metabolic process | 1.54E-7 | 2.77E-5 | 1.69 (5247,298,1009,97) | [+] Show genes  ECM16 - ecm16p  NSA2 - nsa2p  LCP5 - lcp5p  FAF1 - faf1p  RRB1 - rrb1p  UTP30 - utp30p  KRR1 - krr1p  RPS16A - ribosomal 40s subunit protein s16a  RPL8A - ribosomal 60s subunit protein l8a  NOP7 - nop7p  UTP10 - utp10p  UTP14 - utp14p  RAT1 - rat1p  KRI1 - kri1p  RPS8B - ribosomal 40s subunit protein s8b  RPS21A - ribosomal 40s subunit protein s21a  RPS1A - ribosomal 40s subunit protein s1a  RPL3 - ribosomal 60s subunit protein l3  RPS6A - ribosomal 40s subunit protein s6a  RPS21B - rps21bp  UTP8 - utp8p  RLP7 - rlp7p  UTP9 - utp9p  NSA1 - nsa1p  CBF5 - pseudouridine synthase cbf5  RPS2 - ribosomal 40s subunit protein s2  BUD21 - bud21p  TSR2 - tsr2p  RPS9A - ribosomal 40s subunit protein s9a  RPS1B - ribosomal 40s subunit protein s1b  HCA4 - hca4p  RPS28B - ribosomal 40s subunit protein s28b  RPL7A - ribosomal 60s subunit protein l7a  NSR1 - nsr1p  RPS11B - ribosomal 40s subunit protein s11b  SRD1 - srd1p  DBP9 - dbp9p  RPS27A - ribosomal 40s subunit protein s27a  EBP2 - ebp2p  DRS1 - drs1p  TFC3 - tfc3p  YTM1 - ytm1p  RPS23B - ribosomal 40s subunit protein s23b  RPS14A - ribosomal 40s subunit protein s14a  RSC9 - rsc9p  YIL096C - hypothetical protein  RPS0A - ribosomal 40s subunit protein s0a  BUD23 - bud23p  DBP7 - dbp7p  MRD1 - mrd1p  RPS24A - ribosomal 40s subunit protein s24a  PWP2 - pwp2p  RPS0B - ribosomal 40s subunit protein s0b  RPL30 - ribosomal 60s subunit protein l30  EFG1 - efg1p  ZUO1 - zuo1p  RRP1 - rrp1p  RPF1 - rpf1p  PXR1 - pxr1p  RTC3 - rtc3p  NOP16 - nop16p  GAR1 - gar1p  BUD22 - bud22p  NUG1 - nug1p  DHR2 - dhr2p  SSF1 - ssf1p  UTP20 - utp20p  CIC1 - cic1p  RRP3 - rna-dependent atpase rrp3  TMA64 - tma64p  RIB2 - bifunctional drap deaminase/trna pseudouridine synthase rib2  ESF1 - esf1p  FYV7 - fyv7p  MPP10 - mpp10p  RPS28A - ribosomal 40s subunit protein s28a  RPS27B - ribosomal 40s subunit protein s27b  DBP2 - dbp2p  RPL35B - ribosomal 60s subunit protein l35b  PNO1 - pno1p  RPS14B - rps14bp  POL5 - pol5p  BMS1 - bms1p  UFD1 - polyubiquitin-binding protein ufd1  SAS10 - sas10p  NOP14 - nop14p  SLX9 - slx9p  RIO1 - rio1p  HAS1 - atp-dependent rna helicase has1  AIR2 - air2p  UTP11 - utp11p  YBR141C - hypothetical protein  RRP12 - rrp12p  RPS20 - ribosomal 40s subunit protein s20  RPS13 - ribosomal 40s subunit protein s13  RRP46 - rrp46p  RPS7A - ribosomal 40s subunit protein s7a  RPA12 - rpa12p |
| GO:0071826 | ribonucleoprotein complex subunit organization | 4.14E-7 | 7.19E-5 | 3.67 (5247,187,176,23) | [+] Show genes  RPS27B - ribosomal 40s subunit protein s27b  PRP45 - prp45p  RPS0B - ribosomal 40s subunit protein s0b  RPL6A - ribosomal 60s subunit protein l6a  RPL24A - ribosomal 60s subunit protein l24a  RPL3 - ribosomal 60s subunit protein l3  RPS14A - ribosomal 40s subunit protein s14a  RPL40A - ubiquitin-ribosomal 60s subunit protein l40a fusion protein  RPS10A - ribosomal 40s subunit protein s10a  RPS10B - ribosomal 40s subunit protein s10b  RPL6B - ribosomal 60s subunit protein l6b  NSR1 - nsr1p  RPS15 - ribosomal 40s subunit protein s15  RPL11B - ribosomal 60s subunit protein l11b  RPS5 - rps5p  RPS17A - ribosomal 40s subunit protein s17a  RPS26A - ribosomal 40s subunit protein s26a  RPS17B - ribosomal 40s subunit protein s17b  RPL11A - ribosomal 60s subunit protein l11a  RPP0 - ribosomal protein p0  RPL24B - ribosomal 60s subunit protein l24b  RPS28A - ribosomal 40s subunit protein s28a  PWP2 - pwp2p |
| GO:0097064 | ncRNA export from nucleus | 4.98E-7 | 8.37E-5 | 8.13 (5247,31,229,11) | [+] Show genes  RPS15 - ribosomal 40s subunit protein s15  RPS0B - ribosomal 40s subunit protein s0b  RPS0A - ribosomal 40s subunit protein s0a  RPS2 - ribosomal 40s subunit protein s2  RPS5 - rps5p  RPS18B - ribosomal 40s subunit protein s18b  RPS26A - ribosomal 40s subunit protein s26a  RPS10A - ribosomal 40s subunit protein s10a  RPS10B - ribosomal 40s subunit protein s10b  TEF1 - tef1p  RPS28A - ribosomal 40s subunit protein s28a |
| GO:0043170 | macromolecule metabolic process | 1.72E-6 | 2.8E-4 | 1.56 (5247,2203,125,82) | [+] Show genes  RPL2B - ribosomal 60s subunit protein l2b  MGM101 - mgm101p  RPS0B - ribosomal 40s subunit protein s0b  ECM16 - ecm16p  RPL29 - ribosomal 60s subunit protein l29  RPL30 - ribosomal 60s subunit protein l30  EFG1 - efg1p  PFA3 - pfa3p  MSS51 - mss51p  RPL23B - ribosomal 60s subunit protein l23b  RPL38 - ribosomal 60s subunit protein l38  RPL13B - ribosomal 60s subunit protein l13b  CHS1 - chitin synthase chs1  RPS16A - ribosomal 40s subunit protein s16a  TEF1 - tef1p  RPL8A - ribosomal 60s subunit protein l8a  UTP14 - utp14p  ASC1 - asc1p  PPT1 - ppt1p  RPS8B - ribosomal 40s subunit protein s8b  DPB3 - dpb3p  RPL26A - ribosomal 60s subunit protein l26a  RPL15A - ribosomal 60s subunit protein l15a  RPL11A - ribosomal 60s subunit protein l11a  RPP0 - ribosomal protein p0  FUM1 - fumarase fum1  RPL34B - ribosomal 60s subunit protein l34b  RPL27B - ribosomal 60s subunit protein l27b  RPL3 - ribosomal 60s subunit protein l3  RPS21B - rps21bp  SMB1 - smb1p  FPR4 - peptidylprolyl isomerase fpr4  RPP2B - ribosomal protein p2b  RLP7 - rlp7p  RPL6B - ribosomal 60s subunit protein l6b  RPS15 - ribosomal 40s subunit protein s15  RPP2A - ribosomal protein p2a  RPS5 - rps5p  RPS17A - ribosomal 40s subunit protein s17a  GSC2 - gsc2p  RPL24B - ribosomal 60s subunit protein l24b  RPS28A - ribosomal 40s subunit protein s28a  RPL17A - ribosomal 60s subunit protein l17a  RPS1B - ribosomal 40s subunit protein s1b  RPS27B - ribosomal 40s subunit protein s27b  RPL14A - ribosomal 60s subunit protein l14a  RPP1B - ribosomal protein p1b  RPL17B - rpl17bp  HCA4 - hca4p  MRPL24 - mitochondrial 54s ribosomal protein yml24/yml14  RPL32 - ribosomal 60s subunit protein l32  RPL26B - ribosomal 60s subunit protein l26b  RPL22B - ribosomal 60s subunit protein l22b  PRB1 - prb1p  SLT2 - slt2p  RPS10A - ribosomal 40s subunit protein s10a  ACT1 - actin  NSR1 - nsr1p  NCS2 - ncs2p  RPL34A - ribosomal 60s subunit protein l34a  SSE1 - sse1p  RPL27A - ribosomal 60s subunit protein l27a  SRD1 - srd1p  RPS22A - rps22ap  RPS17B - ribosomal 40s subunit protein s17b  YHR020W - proline--trna ligase  RPL6A - ribosomal 60s subunit protein l6a  RPC37 - rpc37p  RPS23B - ribosomal 40s subunit protein s23b  RPL42B - ribosomal 60s subunit protein l42b  APA2 - apa2p  RPS14A - ribosomal 40s subunit protein s14a  RPL31A - ribosomal 60s subunit protein l31a  STM1 - stm1p  HSL1 - hsl1p  RIO1 - rio1p  CHS3 - chitin synthase chs3  RPS13 - ribosomal 40s subunit protein s13  RPS26A - ribosomal 40s subunit protein s26a  RPS7A - ribosomal 40s subunit protein s7a  RPL16B - ribosomal 60s subunit protein l16b  RPS24A - ribosomal 40s subunit protein s24a |
| GO:0034470 | ncRNA processing | 2.69E-6 | 4.24E-4 | 1.55 (5247,372,1009,111) | [+] Show genes  NSA2 - nsa2p  ECM16 - ecm16p  LCP5 - lcp5p  FAF1 - faf1p  RRB1 - rrb1p  UTP30 - utp30p  LHP1 - lhp1p  KRR1 - krr1p  RPS16A - ribosomal 40s subunit protein s16a  RPL8A - ribosomal 60s subunit protein l8a  NOP7 - nop7p  UTP10 - utp10p  UTP14 - utp14p  RAT1 - rat1p  IKI1 - elongator subunit iki1  KRI1 - kri1p  RPS8B - ribosomal 40s subunit protein s8b  RPS21A - ribosomal 40s subunit protein s21a  RPS1A - ribosomal 40s subunit protein s1a  RPL3 - ribosomal 60s subunit protein l3  RPS6A - ribosomal 40s subunit protein s6a  RPS21B - rps21bp  STP1 - stp1p  UTP8 - utp8p  RLP7 - rlp7p  UTP9 - utp9p  NSA1 - nsa1p  NFS1 - nfs1p  CBF5 - pseudouridine synthase cbf5  RPS2 - ribosomal 40s subunit protein s2  BUD21 - bud21p  TSR2 - tsr2p  RPS9A - ribosomal 40s subunit protein s9a  RPS1B - ribosomal 40s subunit protein s1b  HCA4 - hca4p  RPS28B - ribosomal 40s subunit protein s28b  RPL7A - ribosomal 60s subunit protein l7a  NSR1 - nsr1p  RPS11B - ribosomal 40s subunit protein s11b  SRD1 - srd1p  DBP9 - dbp9p  RPS27A - ribosomal 40s subunit protein s27a  EBP2 - ebp2p  DRS1 - drs1p  YTM1 - ytm1p  TUM1 - tum1p  RPS23B - ribosomal 40s subunit protein s23b  ELP2 - elongator subunit elp2  RPS14A - ribosomal 40s subunit protein s14a  RSC9 - rsc9p  TGS1 - tgs1p  TYW3 - tyw3p  YIL096C - hypothetical protein  RPS0A - ribosomal 40s subunit protein s0a  BUD23 - bud23p  DBP7 - dbp7p  ABP140 - abp140p  MRD1 - mrd1p  RPS24A - ribosomal 40s subunit protein s24a  PWP2 - pwp2p  RPS0B - ribosomal 40s subunit protein s0b  NCL1 - ncl1p  RPL30 - ribosomal 60s subunit protein l30  EFG1 - efg1p  ZUO1 - zuo1p  RRP1 - rrp1p  RPF1 - rpf1p  RTC3 - rtc3p  PXR1 - pxr1p  NOP16 - nop16p  GAR1 - gar1p  BUD22 - bud22p  NUG1 - nug1p  SSF1 - ssf1p  DHR2 - dhr2p  UTP20 - utp20p  CIC1 - cic1p  RRP3 - rna-dependent atpase rrp3  TMA64 - tma64p  RIB2 - bifunctional drap deaminase/trna pseudouridine synthase rib2  ESF1 - esf1p  TRM9 - trm9p  FYV7 - fyv7p  MPP10 - mpp10p  RPS28A - ribosomal 40s subunit protein s28a  TAN1 - tan1p  RPS27B - ribosomal 40s subunit protein s27b  TRM11 - trm11p  DBP2 - dbp2p  RPL35B - ribosomal 60s subunit protein l35b  PNO1 - pno1p  NCS2 - ncs2p  CIA1 - cia1p  RPS14B - rps14bp  BMS1 - bms1p  SAS10 - sas10p  NOP14 - nop14p  SLX9 - slx9p  SUA5 - sua5p  AIR2 - air2p  RIO1 - rio1p  HAS1 - atp-dependent rna helicase has1  MRPL3 - mitochondrial 54s ribosomal protein yml3  UTP11 - utp11p  YBR141C - hypothetical protein  RRP12 - rrp12p  RPS20 - ribosomal 40s subunit protein s20  RPS13 - ribosomal 40s subunit protein s13  RRP46 - rrp46p  RPS7A - ribosomal 40s subunit protein s7a  NAB3 - nab3p |
| GO:0000478 | endonucleolytic cleavage involved in rRNA processing | 6.13E-6 | 9.39E-4 | 2.59 (5247,55,993,27) | [+] Show genes  RPS0B - ribosomal 40s subunit protein s0b  RPS27B - ribosomal 40s subunit protein s27b  RPL17A - ribosomal 60s subunit protein l17a  LCP5 - lcp5p  RPL17B - rpl17bp  RRS1 - rrs1p  KRR1 - krr1p  PNO1 - pno1p  UTP14 - utp14p  UTP10 - utp10p  RAT1 - rat1p  KRI1 - kri1p  BMS1 - bms1p  UTP20 - utp20p  RPS21A - ribosomal 40s subunit protein s21a  SAS10 - sas10p  NOP14 - nop14p  RPS18B - ribosomal 40s subunit protein s18b  RPS21B - rps21bp  UTP11 - utp11p  RPS0A - ribosomal 40s subunit protein s0a  BUD21 - bud21p  BUD23 - bud23p  RPL37A - ribosomal 60s subunit protein l37a  MPP10 - mpp10p  MRD1 - mrd1p  PWP2 - pwp2p |
| GO:0000479 | endonucleolytic cleavage of tricistronic rRNA transcript (SSU-rRNA, 5.8S rRNA, LSU-rRNA) | 6.13E-6 | 9.12E-4 | 2.59 (5247,55,993,27) | [+] Show genes  RPS0B - ribosomal 40s subunit protein s0b  RPS27B - ribosomal 40s subunit protein s27b  RPL17A - ribosomal 60s subunit protein l17a  LCP5 - lcp5p  RPL17B - rpl17bp  RRS1 - rrs1p  KRR1 - krr1p  PNO1 - pno1p  UTP14 - utp14p  UTP10 - utp10p  RAT1 - rat1p  KRI1 - kri1p  BMS1 - bms1p  RPS21A - ribosomal 40s subunit protein s21a  UTP20 - utp20p  SAS10 - sas10p  NOP14 - nop14p  RPS18B - ribosomal 40s subunit protein s18b  RPS21B - rps21bp  UTP11 - utp11p  RPS0A - ribosomal 40s subunit protein s0a  BUD21 - bud21p  BUD23 - bud23p  RPL37A - ribosomal 60s subunit protein l37a  MPP10 - mpp10p  MRD1 - mrd1p  PWP2 - pwp2p |
| GO:0071555 | cell wall organization | 1.15E-5 | 1.67E-3 | 6.69 (5247,201,39,10) | [+] Show genes  YLR194C - hypothetical protein  SED1 - sed1p  FIG2 - fig2p  CHS3 - chitin synthase chs3  HSP150 - hsp150p  CHS1 - chitin synthase chs1  CWP2 - cwp2p  ACT1 - actin  SCW4 - scw4p  CIS3 - cis3p |
| GO:0045229 | external encapsulating structure organization | 1.15E-5 | 1.62E-3 | 6.69 (5247,201,39,10) | [+] Show genes  YLR194C - hypothetical protein  SED1 - sed1p  FIG2 - fig2p  CHS3 - chitin synthase chs3  HSP150 - hsp150p  CHS1 - chitin synthase chs1  CWP2 - cwp2p  ACT1 - actin  SCW4 - scw4p  CIS3 - cis3p |
| GO:0000027 | ribosomal large subunit assembly | 1.54E-5 | 2.11E-3 | 2.28 (5247,38,1575,26) | [+] Show genes  MRPL7 - mitochondrial 54s ribosomal protein yml7/yml5  RPL25 - ribosomal 60s subunit protein l25  RPL24A - ribosomal 60s subunit protein l24a  IPI3 - ipi3p  RPL40A - ubiquitin-ribosomal 60s subunit protein l40a fusion protein  RSA4 - rsa4p  MAK11 - mak11p  RPL12A - ribosomal 60s subunit protein l12a  SSF1 - ssf1p  BRX1 - brx1p  DRS1 - drs1p  RPL11A - ribosomal 60s subunit protein l11a  RPP0 - ribosomal protein p0  RPL6A - ribosomal 60s subunit protein l6a  RPL3 - ribosomal 60s subunit protein l3  LSG1 - lsg1p  RIX1 - rix1p  RPL6B - ribosomal 60s subunit protein l6b  HPM1 - hpm1p  RPF2 - rpf2p  YVH1 - yvh1p  MAK21 - mak21p  RPL11B - ribosomal 60s subunit protein l11b  RPL5 - ribosomal 60s subunit protein l5  NOP53 - nop53p  RPL24B - ribosomal 60s subunit protein l24b |
| GO:0071554 | cell wall organization or biogenesis | 1.92E-5 | 2.57E-3 | 6.35 (5247,212,39,10) | [+] Show genes  YLR194C - hypothetical protein  SED1 - sed1p  FIG2 - fig2p  CHS3 - chitin synthase chs3  HSP150 - hsp150p  CHS1 - chitin synthase chs1  CWP2 - cwp2p  ACT1 - actin  SCW4 - scw4p  CIS3 - cis3p |
| GO:0006450 | regulation of translational fidelity | 2.89E-5 | 3.76E-3 | 7.11 (5247,29,229,9) | [+] Show genes  RPS2 - ribosomal 40s subunit protein s2  RPS23B - ribosomal 40s subunit protein s23b  RPL31A - ribosomal 60s subunit protein l31a  RPL31B - ribosomal 60s subunit protein l31b  YNL040W - putative alanine--trna ligase  RPS5 - rps5p  SUA5 - sua5p  CDC60 - leucine--trna ligase cdc60  YHR020W - proline--trna ligase |
| GO:0034660 | ncRNA metabolic process | 3.64E-5 | 4.63E-3 | 1.31 (5247,476,1589,189) | [+] Show genes  NSA2 - nsa2p  ECM16 - ecm16p  FAF1 - faf1p  LCP5 - lcp5p  RRB1 - rrb1p  SAP185 - sap185p  YNL040W - putative alanine--trna ligase  UTP30 - utp30p  SNF6 - snf6p  GTF1 - glutamyl-trna(gln) amidotransferase subunit f  LHP1 - lhp1p  KRR1 - krr1p  GRS1 - glycine--trna ligase  RPS16A - ribosomal 40s subunit protein s16a  RPL8A - ribosomal 60s subunit protein l8a  NOP7 - nop7p  UTP10 - utp10p  UTP14 - utp14p  RAT1 - rat1p  IKI1 - elongator subunit iki1  KRI1 - kri1p  RPS8B - ribosomal 40s subunit protein s8b  SLM3 - slm3p  RPS21A - ribosomal 40s subunit protein s21a  RPA190 - rpa190p  TRF5 - non-canonical poly(a) polymerase trf5  GRC3 - grc3p  RPT2 - proteasome regulatory particle base subunit rpt2  RPS1A - ribosomal 40s subunit protein s1a  RPL3 - ribosomal 60s subunit protein l3  RPS6A - ribosomal 40s subunit protein s6a  PUS4 - pseudouridine synthase pus4  RPR2 - rpr2p  RPS21B - rps21bp  RPS31 - ubiquitin-ribosomal 40s subunit protein s31 fusion protein  STP1 - stp1p  RPO26 - rpo26p  UTP8 - utp8p  RIX1 - rix1p  UTP9 - utp9p  RLP7 - rlp7p  NSA1 - nsa1p  NFS1 - nfs1p  RPF2 - rpf2p  CBF5 - pseudouridine synthase cbf5  YNL022C - hypothetical protein  RPS2 - ribosomal 40s subunit protein s2  BUD21 - bud21p  RPA34 - rpa34p  TSR2 - tsr2p  ENP2 - enp2p  MOT1 - mot1p  RPS9A - ribosomal 40s subunit protein s9a  NOP58 - nop58p  RRP36 - rrp36p  RPS1B - ribosomal 40s subunit protein s1b  MSE1 - glutamate--trna ligase mse1  HCA4 - hca4p  TAD2 - tad2p  RPS28B - ribosomal 40s subunit protein s28b  PAP1 - pap1p  RPL7A - ribosomal 60s subunit protein l7a  NSR1 - nsr1p  RPS11B - ribosomal 40s subunit protein s11b  YNL247W - cysteine--trna ligase  MAK11 - mak11p  RPC34 - rpc34p  SRD1 - srd1p  RPA14 - rpa14p  DBP9 - dbp9p  RPS27A - ribosomal 40s subunit protein s27a  EBP2 - ebp2p  RRN11 - rrn11p  TFC3 - tfc3p  DRS1 - drs1p  YTM1 - ytm1p  RCL1 - rcl1p  TUM1 - tum1p  MST1 - threonine--trna ligase mst1  RPC37 - rpc37p  RPS23B - ribosomal 40s subunit protein s23b  ELP2 - elongator subunit elp2  RPS14A - ribosomal 40s subunit protein s14a  UTP5 - utp5p  RSC9 - rsc9p  CDC60 - leucine--trna ligase cdc60  TGS1 - tgs1p  YIL096C - hypothetical protein  TYW3 - tyw3p  RPS0A - ribosomal 40s subunit protein s0a  BUD23 - bud23p  RIO2 - rio2p  RPC40 - rpc40p  NOP53 - nop53p  TRM112 - trm112p  DBP7 - dbp7p  RPS24A - ribosomal 40s subunit protein s24a  ABP140 - abp140p  MRD1 - mrd1p  PWP2 - pwp2p  HST3 - hst3p  RPS0B - ribosomal 40s subunit protein s0b  RNH70 - rnh70p  NCL1 - ncl1p  RPL30 - ribosomal 60s subunit protein l30  EFG1 - efg1p  RRN9 - rrn9p  IPI3 - ipi3p  ZUO1 - zuo1p  RRP1 - rrp1p  RPF1 - rpf1p  TRM12 - trm12p  PXR1 - pxr1p  RTC3 - rtc3p  RRP8 - rrp8p  NOP16 - nop16p  GAR1 - gar1p  NUG1 - nug1p  MRM1 - mrm1p  BUD22 - bud22p  SSF1 - ssf1p  DHR2 - dhr2p  RLI1 - rli1p  BRX1 - brx1p  UTP20 - utp20p  PCC1 - pcc1p  LSM7 - lsm7p  CIC1 - cic1p  RRP3 - rna-dependent atpase rrp3  MSF1 - phenylalanine--trna ligase  TMA64 - tma64p  CGI121 - cgi121p  RIB2 - bifunctional drap deaminase/trna pseudouridine synthase rib2  ESF1 - esf1p  TRM9 - trm9p  NOP9 - nop9p  FYV7 - fyv7p  MPP10 - mpp10p  MTO1 - mto1p  RPS28A - ribosomal 40s subunit protein s28a  TAN1 - tan1p  FRS1 - phenylalanine--trna ligase subunit beta  RPS27B - ribosomal 40s subunit protein s27b  DUS3 - dus3p  BDF1 - bdf1p  RPC19 - rpc19p  DBP2 - dbp2p  RPL7B - ribosomal 60s subunit protein l7b  TRM11 - trm11p  NOP15 - nop15p  SEN15 - sen15p  TRM13 - trm13p  NIP7 - nip7p  PUS1 - pseudouridine synthase pus1  RPL35B - ribosomal 60s subunit protein l35b  PET112 - glutamyl-trna(gln) amidotransferase subunit pet112  PNO1 - pno1p  NCS2 - ncs2p  CIA1 - cia1p  DIA4 - putative serine--trna ligase dia4  POL5 - pol5p  RPS14B - rps14bp  BMS1 - bms1p  UFD1 - polyubiquitin-binding protein ufd1  GON7 - gon7p  SAS10 - sas10p  DBR1 - dbr1p  NOP14 - nop14p  YHR020W - proline--trna ligase  HER2 - glutamyl-trna(gln) amidotransferase subunit her2  SLX9 - slx9p  YHR003C - hypothetical protein  SUA5 - sua5p  AIR2 - air2p  RIO1 - rio1p  HAS1 - atp-dependent rna helicase has1  ROK1 - rna-dependent atpase rok1  MRPL3 - mitochondrial 54s ribosomal protein yml3  UTP11 - utp11p  YBR141C - hypothetical protein  RRP12 - rrp12p  RPS20 - ribosomal 40s subunit protein s20  RPS13 - ribosomal 40s subunit protein s13  RRG1 - rrg1p  RRP46 - rrp46p  RPA12 - rpa12p  RPS7A - ribosomal 40s subunit protein s7a  NAB3 - nab3p  VAS1 - valine--trna ligase |
| GO:0000447 | endonucleolytic cleavage in ITS1 to separate SSU-rRNA from 5.8S rRNA and LSU-rRNA from tricistronic rRNA transcript (SSU-rRNA, 5.8S rRNA, LSU-rRNA) | 7.15E-5 | 8.87E-3 | 2.64 (5247,42,993,21) | [+] Show genes  RPS0B - ribosomal 40s subunit protein s0b  LCP5 - lcp5p  RPS18B - ribosomal 40s subunit protein s18b  RRS1 - rrs1p  RPS21B - rps21bp  KRR1 - krr1p  PNO1 - pno1p  UTP11 - utp11p  UTP14 - utp14p  UTP10 - utp10p  RPS0A - ribosomal 40s subunit protein s0a  BUD21 - bud21p  BUD23 - bud23p  KRI1 - kri1p  MPP10 - mpp10p  RPS21A - ribosomal 40s subunit protein s21a  SAS10 - sas10p  UTP20 - utp20p  MRD1 - mrd1p  PWP2 - pwp2p  NOP14 - nop14p |
| GO:0022610 | biological adhesion | 7.53E-5 | 9.11E-3 | 27.84 (5247,13,58,4) | [+] Show genes  AGA2 - aga2p  NRG2 - nrg2p  FIG2 - fig2p  FLO11 - flo11p |
| GO:0007155 | cell adhesion | 7.53E-5 | 8.91E-3 | 27.84 (5247,13,58,4) | [+] Show genes  AGA2 - aga2p  NRG2 - nrg2p  FIG2 - fig2p  FLO11 - flo11p |
| GO:0042273 | ribosomal large subunit biogenesis | 7.79E-5 | 9.02E-3 | 1.98 (5247,52,1630,32) | [+] Show genes  NSA2 - nsa2p  RPL14A - ribosomal 60s subunit protein l14a  HRR25 - hrr25p  RPL7B - ribosomal 60s subunit protein l7b  NOP15 - nop15p  TIF4631 - tif4631p  RPL26B - ribosomal 60s subunit protein l26b  RRS1 - rrs1p  NIP7 - nip7p  RPL7A - ribosomal 60s subunit protein l7a  RRP8 - rrp8p  NOP16 - nop16p  NOP7 - nop7p  REI1 - rei1p  MAK11 - mak11p  MDV1 - mdv1p  RLI1 - rli1p  NOC2 - noc2p  EBP2 - ebp2p  RPL26A - ribosomal 60s subunit protein l26a  ALB1 - alb1p  YTM1 - ytm1p  HAS1 - atp-dependent rna helicase has1  SDA1 - sda1p  RLP7 - rlp7p  NSA1 - nsa1p  RPL33B - ribosomal 60s subunit protein l33b  MAK21 - mak21p  REH1 - reh1p  TRM112 - trm112p  JIP5 - jip5p  RPL33A - ribosomal 60s subunit protein l33a |
| GO:0071852 | fungal-type cell wall organization or biogenesis | 7.85E-5 | 8.89E-3 | 4.52 (5247,170,82,12) | [+] Show genes  YLR194C - hypothetical protein  DFG5 - dfg5p  SED1 - sed1p  PAU7 - pau7p  SLT2 - slt2p  CHS3 - chitin synthase chs3  HSP150 - hsp150p  CWP2 - cwp2p  ACT1 - actin  CHS1 - chitin synthase chs1  GSC2 - gsc2p  CIS3 - cis3p |
| GO:0044764 | multi-organism cellular process | 1.49E-4 | 1.65E-2 | 25.72 (5247,24,34,4) | [+] Show genes  ASG7 - asg7p  FIG2 - fig2p  FLO11 - flo11p  SCW4 - scw4p |
| GO:0006414 | translational elongation | 1.99E-4 | 2.15E-2 | 10.32 (5247,28,109,6) | [+] Show genes  ASC1 - asc1p  RPP1B - ribosomal protein p1b  RPP2A - ribosomal protein p2a  STM1 - stm1p  RPP2B - ribosomal protein p2b  TEF1 - tef1p |
| GO:0006807 | nitrogen compound metabolic process | 2.54E-4 | 2.69E-2 | 1.33 (5247,2646,156,105) | [+] Show genes  RPL2B - ribosomal 60s subunit protein l2b  RPL29 - ribosomal 60s subunit protein l29  ECM16 - ecm16p  MGM101 - mgm101p  RRB1 - rrb1p  VMA2 - vma2p  PFA3 - pfa3p  KTR2 - ktr2p  RPL9A - ribosomal 60s subunit protein l9a  MSS51 - mss51p  RPL23B - ribosomal 60s subunit protein l23b  RPL13B - ribosomal 60s subunit protein l13b  RPS16A - ribosomal 40s subunit protein s16a  RPL8A - ribosomal 60s subunit protein l8a  UTP14 - utp14p  RAD9 - rad9p  PPT1 - ppt1p  RPS8B - ribosomal 40s subunit protein s8b  ATP18 - atp18p  RPL34B - ribosomal 60s subunit protein l34b  RPL27B - ribosomal 60s subunit protein l27b  RPL3 - ribosomal 60s subunit protein l3  CYT1 - ubiquinol--cytochrome-c reductase catalytic subunit cyt1  RPL19A - ribosomal 60s subunit protein l19a  RPS21B - rps21bp  FPR4 - peptidylprolyl isomerase fpr4  RPL6B - ribosomal 60s subunit protein l6b  RLP7 - rlp7p  RPS5 - rps5p  RPL37A - ribosomal 60s subunit protein l37a  RPL24B - ribosomal 60s subunit protein l24b  RPS1B - ribosomal 40s subunit protein s1b  RPL17A - ribosomal 60s subunit protein l17a  RPL14A - ribosomal 60s subunit protein l14a  HCA4 - hca4p  MRPL24 - mitochondrial 54s ribosomal protein yml24/yml14  RPS10A - ribosomal 40s subunit protein s10a  NSR1 - nsr1p  ABZ1 - 4-amino-4-deoxychorismate synthase  PGK1 - phosphoglycerate kinase  RPL34A - ribosomal 60s subunit protein l34a  SSE1 - sse1p  SRD1 - srd1p  RNR4 - ribonucleotide-diphosphate reductase subunit rnr4  RPS17B - ribosomal 40s subunit protein s17b  RPL6A - ribosomal 60s subunit protein l6a  RPS23B - ribosomal 40s subunit protein s23b  RPC37 - rpc37p  RPL42B - ribosomal 60s subunit protein l42b  RPS14A - ribosomal 40s subunit protein s14a  RPL31A - ribosomal 60s subunit protein l31a  BNA5 - kynureninase  CHS3 - chitin synthase chs3  THR4 - threonine synthase thr4  RPS24A - ribosomal 40s subunit protein s24a  RPS0B - ribosomal 40s subunit protein s0b  RPL24A - ribosomal 60s subunit protein l24a  RPL30 - ribosomal 60s subunit protein l30  EFG1 - efg1p  RPL40A - ubiquitin-ribosomal 60s subunit protein l40a fusion protein  CHS1 - chitin synthase chs1  RPL38 - ribosomal 60s subunit protein l38  TEF1 - tef1p  ILV5 - ketol-acid reductoisomerase  DPB3 - dpb3p  IRE1 - ire1p  RPL26A - ribosomal 60s subunit protein l26a  RPL15A - ribosomal 60s subunit protein l15a  RPL11A - ribosomal 60s subunit protein l11a  RPP0 - ribosomal protein p0  FUM1 - fumarase fum1  RPS18B - ribosomal 40s subunit protein s18b  SLX1 - slx1p  RPS10B - ribosomal 40s subunit protein s10b  SMB1 - smb1p  RPP2B - ribosomal protein p2b  RPS15 - ribosomal 40s subunit protein s15  RPP2A - ribosomal protein p2a  DUR1,2 - bifunctional urea carboxylase/allophanate hydrolase  RPS17A - ribosomal 40s subunit protein s17a  RPS28A - ribosomal 40s subunit protein s28a  RPS27B - ribosomal 40s subunit protein s27b  RPP1B - ribosomal protein p1b  RPL21A - ribosomal 60s subunit protein l21a  RPL17B - rpl17bp  RPL32 - ribosomal 60s subunit protein l32  RPL26B - ribosomal 60s subunit protein l26b  RPL22B - ribosomal 60s subunit protein l22b  SLT2 - slt2p  RPL35B - ribosomal 60s subunit protein l35b  PRB1 - prb1p  ACT1 - actin  NCS2 - ncs2p  RPL27A - ribosomal 60s subunit protein l27a  RPS22A - rps22ap  YHR020W - proline--trna ligase  APA2 - apa2p  STM1 - stm1p  SUA5 - sua5p  HSL1 - hsl1p  RIO1 - rio1p  RPS13 - ribosomal 40s subunit protein s13  RPS26A - ribosomal 40s subunit protein s26a  RPS7A - ribosomal 40s subunit protein s7a  RPL16B - ribosomal 60s subunit protein l16b |
| GO:0031505 | fungal-type cell wall organization | 2.83E-4 | 2.95E-2 | 6.90 (5247,156,39,8) | [+] Show genes  YLR194C - hypothetical protein  SED1 - sed1p  CHS3 - chitin synthase chs3  HSP150 - hsp150p  CWP2 - cwp2p  ACT1 - actin  CHS1 - chitin synthase chs1  CIS3 - cis3p |
| GO:0006122 | mitochondrial electron transport, ubiquinol to cytochrome c | 4.28E-4 | 4.37E-2 | 5.17 (5247,10,710,7) | [+] Show genes  RIP1 - ubiquinol--cytochrome-c reductase catalytic subunit rip1  QCR7 - ubiquinol--cytochrome-c reductase subunit 7  CYT1 - ubiquinol--cytochrome-c reductase catalytic subunit cyt1  QCR6 - ubiquinol--cytochrome-c reductase subunit 6  QCR2 - ubiquinol--cytochrome-c reductase subunit 2  QCR8 - qcr8p  QCR10 - ubiquinol--cytochrome-c reductase subunit 10 |
| GO:0071704 | organic substance metabolic process | 4.69E-4 | 4.69E-2 | 1.31 (5247,2970,125,93) | [+] Show genes  RPL2B - ribosomal 60s subunit protein l2b  MGM101 - mgm101p  RPS0B - ribosomal 40s subunit protein s0b  ECM16 - ecm16p  RPL29 - ribosomal 60s subunit protein l29  RPL30 - ribosomal 60s subunit protein l30  EFG1 - efg1p  PFA3 - pfa3p  TSL1 - tsl1p  RPL23B - ribosomal 60s subunit protein l23b  MSS51 - mss51p  RPL13B - ribosomal 60s subunit protein l13b  RPL38 - ribosomal 60s subunit protein l38  CHS1 - chitin synthase chs1  SCW4 - scw4p  RPS16A - ribosomal 40s subunit protein s16a  TEF1 - tef1p  RPL8A - ribosomal 60s subunit protein l8a  UTP14 - utp14p  ASC1 - asc1p  PPT1 - ppt1p  RPS8B - ribosomal 40s subunit protein s8b  ATP18 - atp18p  DPB3 - dpb3p  RPL26A - ribosomal 60s subunit protein l26a  RPL11A - ribosomal 60s subunit protein l11a  RPL15A - ribosomal 60s subunit protein l15a  RPP0 - ribosomal protein p0  FUM1 - fumarase fum1  RPL34B - ribosomal 60s subunit protein l34b  RPL27B - ribosomal 60s subunit protein l27b  RPL3 - ribosomal 60s subunit protein l3  DFG5 - dfg5p  YPL088W - aldo-keto reductase superfamily protein  RPS21B - rps21bp  SMB1 - smb1p  RPP2B - ribosomal protein p2b  FPR4 - peptidylprolyl isomerase fpr4  RLP7 - rlp7p  RPL6B - ribosomal 60s subunit protein l6b  RPS15 - ribosomal 40s subunit protein s15  RPP2A - ribosomal protein p2a  DUR1,2 - bifunctional urea carboxylase/allophanate hydrolase  RPS5 - rps5p  RPS17A - ribosomal 40s subunit protein s17a  GSC2 - gsc2p  RPL24B - ribosomal 60s subunit protein l24b  RPS28A - ribosomal 40s subunit protein s28a  RPL17A - ribosomal 60s subunit protein l17a  RPS1B - ribosomal 40s subunit protein s1b  RPS27B - ribosomal 40s subunit protein s27b  RPL14A - ribosomal 60s subunit protein l14a  RPP1B - ribosomal protein p1b  RPL17B - rpl17bp  HCA4 - hca4p  RPL32 - ribosomal 60s subunit protein l32  MRPL24 - mitochondrial 54s ribosomal protein yml24/yml14  RPL26B - ribosomal 60s subunit protein l26b  RPL22B - ribosomal 60s subunit protein l22b  PRB1 - prb1p  SLT2 - slt2p  RPS10A - ribosomal 40s subunit protein s10a  ACT1 - actin  NSR1 - nsr1p  NCS2 - ncs2p  ABZ1 - 4-amino-4-deoxychorismate synthase  RPL34A - ribosomal 60s subunit protein l34a  SSE1 - sse1p  RPL27A - ribosomal 60s subunit protein l27a  SRD1 - srd1p  RPS22A - rps22ap  RPS17B - ribosomal 40s subunit protein s17b  YHR020W - proline--trna ligase  PAH1 - phosphatidate phosphatase pah1  RPL6A - ribosomal 60s subunit protein l6a  RPC37 - rpc37p  RPS23B - ribosomal 40s subunit protein s23b  RPL42B - ribosomal 60s subunit protein l42b  APA2 - apa2p  RPS14A - ribosomal 40s subunit protein s14a  RPL31A - ribosomal 60s subunit protein l31a  STM1 - stm1p  HSL1 - hsl1p  RIO1 - rio1p  CHS3 - chitin synthase chs3  BNA5 - kynureninase  DLD1 - dld1p  THR4 - threonine synthase thr4  RPS13 - ribosomal 40s subunit protein s13  RPS26A - ribosomal 40s subunit protein s26a  RPS7A - ribosomal 40s subunit protein s7a  RPS24A - ribosomal 40s subunit protein s24a  RPL16B - ribosomal 60s subunit protein l16b |
| GO:0051704 | multi-organism process | 5.08E-4 | 4.99E-2 | 2.10 (5247,38,1510,23) | [+] Show genes  AGA2 - aga2p  FUS2 - fus2p  SKI2 - ski2p  LSG1 - lsg1p  ATF1 - atf1p  ASG7 - asg7p  FLO9 - flo9p  CHS5 - chs5p  CDC42 - cdc42p  FLO11 - flo11p  FUS3 - fus3p  RVS161 - rvs161p  SCW4 - scw4p  MKT1 - mkt1p  MAK32 - mak32p  HBT1 - hbt1p  PRM1 - prm1p  SCW10 - scw10p  AGA1 - aga1p  FIG2 - fig2p  FLO1 - flo1p  TPS2 - trehalose-phosphatase tps2  NRG1 - nrg1p |
| GO:0043709 | cell adhesion involved in single-species biofilm formation | 6.3E-4 | 6.07E-2 | 79.50 (5247,4,33,2) | [+] Show genes  NRG2 - nrg2p  FLO11 - flo11p |
| GO:0043708 | cell adhesion involved in biofilm formation | 6.3E-4 | 5.96E-2 | 79.50 (5247,4,33,2) | [+] Show genes  NRG2 - nrg2p  FLO11 - flo11p |
| GO:0044238 | primary metabolic process | 6.36E-4 | 5.91E-2 | 1.29 (5247,2836,156,109) | [+] Show genes  RPL2B - ribosomal 60s subunit protein l2b  ECM16 - ecm16p  RPL29 - ribosomal 60s subunit protein l29  MGM101 - mgm101p  RRB1 - rrb1p  VMA2 - vma2p  PFA3 - pfa3p  KTR2 - ktr2p  TSL1 - tsl1p  RPL9A - ribosomal 60s subunit protein l9a  MSS51 - mss51p  RPL23B - ribosomal 60s subunit protein l23b  RPL13B - ribosomal 60s subunit protein l13b  RPS16A - ribosomal 40s subunit protein s16a  RPL8A - ribosomal 60s subunit protein l8a  UTP14 - utp14p  RAD9 - rad9p  PPT1 - ppt1p  RPS8B - ribosomal 40s subunit protein s8b  ATP18 - atp18p  RPL34B - ribosomal 60s subunit protein l34b  RPL27B - ribosomal 60s subunit protein l27b  RPL3 - ribosomal 60s subunit protein l3  CYT1 - ubiquinol--cytochrome-c reductase catalytic subunit cyt1  RPL19A - ribosomal 60s subunit protein l19a  RPS21B - rps21bp  FPR4 - peptidylprolyl isomerase fpr4  RPL6B - ribosomal 60s subunit protein l6b  RLP7 - rlp7p  RPS5 - rps5p  RPL37A - ribosomal 60s subunit protein l37a  RPL24B - ribosomal 60s subunit protein l24b  RPS1B - ribosomal 40s subunit protein s1b  RPL17A - ribosomal 60s subunit protein l17a  RPL14A - ribosomal 60s subunit protein l14a  HCA4 - hca4p  MRPL24 - mitochondrial 54s ribosomal protein yml24/yml14  RPS10A - ribosomal 40s subunit protein s10a  NSR1 - nsr1p  ABZ1 - 4-amino-4-deoxychorismate synthase  PGK1 - phosphoglycerate kinase  RPL34A - ribosomal 60s subunit protein l34a  SSE1 - sse1p  SRD1 - srd1p  RNR4 - ribonucleotide-diphosphate reductase subunit rnr4  RPS17B - ribosomal 40s subunit protein s17b  PAH1 - phosphatidate phosphatase pah1  RPL6A - ribosomal 60s subunit protein l6a  RPS23B - ribosomal 40s subunit protein s23b  RPC37 - rpc37p  RPL42B - ribosomal 60s subunit protein l42b  RPS14A - ribosomal 40s subunit protein s14a  RPL31A - ribosomal 60s subunit protein l31a  BNA5 - kynureninase  CHS3 - chitin synthase chs3  THR4 - threonine synthase thr4  RPS24A - ribosomal 40s subunit protein s24a  RPS0B - ribosomal 40s subunit protein s0b  RPL24A - ribosomal 60s subunit protein l24a  RPL30 - ribosomal 60s subunit protein l30  EFG1 - efg1p  RPL40A - ubiquitin-ribosomal 60s subunit protein l40a fusion protein  RPL38 - ribosomal 60s subunit protein l38  TEF1 - tef1p  SCW4 - scw4p  ILV5 - ketol-acid reductoisomerase  DPB3 - dpb3p  IRE1 - ire1p  RPL26A - ribosomal 60s subunit protein l26a  RPL15A - ribosomal 60s subunit protein l15a  RPL11A - ribosomal 60s subunit protein l11a  RPP0 - ribosomal protein p0  FUM1 - fumarase fum1  DFG5 - dfg5p  RPS18B - ribosomal 40s subunit protein s18b  SLX1 - slx1p  RPS10B - ribosomal 40s subunit protein s10b  SMB1 - smb1p  RPP2B - ribosomal protein p2b  RPS15 - ribosomal 40s subunit protein s15  RPP2A - ribosomal protein p2a  DUR1,2 - bifunctional urea carboxylase/allophanate hydrolase  RPS17A - ribosomal 40s subunit protein s17a  GSC2 - gsc2p  RPS28A - ribosomal 40s subunit protein s28a  RPS27B - ribosomal 40s subunit protein s27b  RPP1B - ribosomal protein p1b  RPL21A - ribosomal 60s subunit protein l21a  RPL17B - rpl17bp  RPL32 - ribosomal 60s subunit protein l32  RPL26B - ribosomal 60s subunit protein l26b  RPL22B - ribosomal 60s subunit protein l22b  SLT2 - slt2p  RPL35B - ribosomal 60s subunit protein l35b  PRB1 - prb1p  ACT1 - actin  NCS2 - ncs2p  RPL27A - ribosomal 60s subunit protein l27a  RPS22A - rps22ap  YHR020W - proline--trna ligase  APA2 - apa2p  STM1 - stm1p  HSL1 - hsl1p  SUA5 - sua5p  RIO1 - rio1p  RPS13 - ribosomal 40s subunit protein s13  RPS26A - ribosomal 40s subunit protein s26a  RPS7A - ribosomal 40s subunit protein s7a  RPL16B - ribosomal 60s subunit protein l16b |
| GO:0044237 | cellular metabolic process | 6.69E-4 | 6.12E-2 | 1.23 (5247,3029,202,144) | [+] Show genes  RPL2B - ribosomal 60s subunit protein l2b  MGM101 - mgm101p  PRP45 - prp45p  ECM16 - ecm16p  RPL29 - ribosomal 60s subunit protein l29  RRB1 - rrb1p  VMA2 - vma2p  UTP30 - utp30p  PFA3 - pfa3p  KTR2 - ktr2p  TSL1 - tsl1p  RPL9A - ribosomal 60s subunit protein l9a  MSS51 - mss51p  RPL23B - ribosomal 60s subunit protein l23b  RPL13B - ribosomal 60s subunit protein l13b  RPS16A - ribosomal 40s subunit protein s16a  RPL8A - ribosomal 60s subunit protein l8a  UTP14 - utp14p  ASC1 - asc1p  RAD9 - rad9p  SHH3 - protein shh3  GPI16 - gpi16p  PPT1 - ppt1p  RPS8B - ribosomal 40s subunit protein s8b  ATP18 - atp18p  TDA9 - tda9p  RPL34B - ribosomal 60s subunit protein l34b  RPL27B - ribosomal 60s subunit protein l27b  RPL3 - ribosomal 60s subunit protein l3  SPO1 - spo1p  RPS6A - ribosomal 40s subunit protein s6a  CYT1 - ubiquinol--cytochrome-c reductase catalytic subunit cyt1  YPL088W - aldo-keto reductase superfamily protein  RPL19A - ribosomal 60s subunit protein l19a  RPS21B - rps21bp  PDR10 - atp-binding cassette multidrug transporter pdr10  FPR4 - peptidylprolyl isomerase fpr4  RPL6B - ribosomal 60s subunit protein l6b  RLP7 - rlp7p  RPS5 - rps5p  RPL37A - ribosomal 60s subunit protein l37a  YJR120W - hypothetical protein  YRB1 - yrb1p  RPL24B - ribosomal 60s subunit protein l24b  RPL17A - ribosomal 60s subunit protein l17a  RPS1B - ribosomal 40s subunit protein s1b  CAB2 - phosphopantothenate--cysteine ligase cab2  RPL14A - ribosomal 60s subunit protein l14a  HCA4 - hca4p  SNX4 - snx4p  MRPL24 - mitochondrial 54s ribosomal protein yml24/yml14  RPS10A - ribosomal 40s subunit protein s10a  NSR1 - nsr1p  ABZ1 - 4-amino-4-deoxychorismate synthase  PGK1 - phosphoglycerate kinase  RPL34A - ribosomal 60s subunit protein l34a  SSE1 - sse1p  RNR4 - ribonucleotide-diphosphate reductase subunit rnr4  SRD1 - srd1p  CST26 - cst26p  MTQ2 - mtq2p  RPS17B - ribosomal 40s subunit protein s17b  PAH1 - phosphatidate phosphatase pah1  RPL6A - ribosomal 60s subunit protein l6a  PRM4 - prm4p  RPC37 - rpc37p  RPS23B - ribosomal 40s subunit protein s23b  RPL42B - ribosomal 60s subunit protein l42b  RPS14A - ribosomal 40s subunit protein s14a  RPL31A - ribosomal 60s subunit protein l31a  HPT1 - hypoxanthine phosphoribosyltransferase  BNA5 - kynureninase  CHS3 - chitin synthase chs3  THR4 - threonine synthase thr4  RPS24A - ribosomal 40s subunit protein s24a  PWP2 - pwp2p  RPS0B - ribosomal 40s subunit protein s0b  RPL24A - ribosomal 60s subunit protein l24a  RFC1 - replication factor c subunit 1  RPL30 - ribosomal 60s subunit protein l30  EFG1 - efg1p  RPL40A - ubiquitin-ribosomal 60s subunit protein l40a fusion protein  SSK22 - ssk22p  CHS1 - chitin synthase chs1  RPL38 - ribosomal 60s subunit protein l38  TEF1 - tef1p  ILV5 - ketol-acid reductoisomerase  RPL16A - ribosomal 60s subunit protein l16a  DPB3 - dpb3p  RPL26A - ribosomal 60s subunit protein l26a  IRE1 - ire1p  RPL15A - ribosomal 60s subunit protein l15a  RPL11A - ribosomal 60s subunit protein l11a  RPP0 - ribosomal protein p0  FUM1 - fumarase fum1  RPS18B - ribosomal 40s subunit protein s18b  SLX1 - slx1p  UGP1 - utp glucose-1-phosphate uridylyltransferase  RPS10B - ribosomal 40s subunit protein s10b  SMB1 - smb1p  RPP2B - ribosomal protein p2b  RPS15 - ribosomal 40s subunit protein s15  TRM9 - trm9p  RPP2A - ribosomal protein p2a  DUR1,2 - bifunctional urea carboxylase/allophanate hydrolase  RPS17A - ribosomal 40s subunit protein s17a  GSC2 - gsc2p  CDA1 - chitin deacetylase cda1  RPS28A - ribosomal 40s subunit protein s28a  RPS27B - ribosomal 40s subunit protein s27b  RPP1B - ribosomal protein p1b  RPL21A - ribosomal 60s subunit protein l21a  RPL17B - rpl17bp  DBP2 - dbp2p  RPL31B - ribosomal 60s subunit protein l31b  RPL32 - ribosomal 60s subunit protein l32  RPL26B - ribosomal 60s subunit protein l26b  RPL22B - ribosomal 60s subunit protein l22b  ARF2 - arf2p  QCR8 - qcr8p  RPL35B - ribosomal 60s subunit protein l35b  SLT2 - slt2p  PRB1 - prb1p  ACT1 - actin  RPL41B - ribosomal 60s subunit protein l41b  NCS2 - ncs2p  YHR033W - putative glutamate 5-kinase  RPL27A - ribosomal 60s subunit protein l27a  RPS22A - rps22ap  YJL185C - hypothetical protein  YHR020W - proline--trna ligase  APA2 - apa2p  STM1 - stm1p  HSL1 - hsl1p  SUA5 - sua5p  RIO1 - rio1p  DLD1 - dld1p  RPL11B - ribosomal 60s subunit protein l11b  RIP1 - ubiquinol--cytochrome-c reductase catalytic subunit rip1  RPS13 - ribosomal 40s subunit protein s13  ATP1 - f1f0 atp synthase subunit alpha  RPS26A - ribosomal 40s subunit protein s26a  RPS7A - ribosomal 40s subunit protein s7a  RPL16B - ribosomal 60s subunit protein l16b |

Species used: Saccharomyces cerevisiae

The system has recognized 5321 genes out of 5595 gene terms entered by the user.  
 628 genes were recognized by gene symbol and 4693 genes by other gene IDs .  
Only 5247 of these genes are associated with a GO term.

  
 Output in Microsoft Excel format  

% List genereted using GOrilla
% http://cbl-gorilla.cs.technion.ac.il/
% GO term pValue
GO:0002181 5.22E-56
GO:0006412 1.14E-34
GO:0043043 2.11E-34
GO:0006518 6.59E-32
GO:0043604 1.44E-31
GO:0043603 2.8E-28
GO:0034645 5.02E-23
GO:1901566 5.04E-21
GO:0009059 1.47E-20
GO:0044271 2.49E-16
GO:0044267 4.73E-13
GO:0044249 6.48E-13
GO:1901576 1.09E-12
GO:0009058 2.27E-12
GO:0000028 1.04E-11
GO:0022613 5.62E-11
GO:0044085 7.64E-11
GO:0034641 1.51E-10
GO:0042254 1.56E-10
GO:0030490 3.04E-10
GO:0000462 3.89E-10
GO:0019538 9.66E-10
GO:0006364 1.2E-9
GO:0006407 3.49E-9
GO:0051029 3.49E-9
GO:0044260 4.14E-8
GO:0022618 1.3E-7
GO:1901564 1.39E-7
GO:0016072 1.54E-7
GO:0071826 4.14E-7
GO:0097064 4.98E-7
GO:0043170 1.72E-6
GO:0034470 2.69E-6
GO:0000478 6.13E-6
GO:0000479 6.13E-6
GO:0071555 1.15E-5
GO:0045229 1.15E-5
GO:0000027 1.54E-5
GO:0071554 1.92E-5
GO:0006450 2.89E-5
GO:0034660 3.64E-5
GO:0000447 7.15E-5
GO:0022610 7.53E-5
GO:0007155 7.53E-5
GO:0042273 7.79E-5
GO:0071852 7.85E-5
GO:0044764 1.49E-4
GO:0006414 1.99E-4
GO:0006807 2.54E-4
GO:0031505 2.83E-4
GO:0006122 4.28E-4
GO:0071704 4.69E-4
GO:0051704 5.08E-4
GO:0043709 6.3E-4
GO:0043708 6.3E-4
GO:0044238 6.36E-4
GO:0044237 6.69E-4
 Visualize output in REViGO   

The GOrilla database is periodically updated using the GO database and other sources.  
The GOrilla database was last updated on Sep 21, 2019

This results page will be available on this site for one month from now (until
Oct 23, 2019
). You can bookmark this page and come back to it later.

  
**'P-value'** is the enrichment
p-value computed according to the mHG or HG model. This p-value is not
corrected for multiple testing of 5208 GO terms.  
  
**'FDR q-value'** is the correction of the above p-value for multiple testing using the Benjamini and Hochberg (1995) method.   
Namely, for the i^th^ term (ranked according to p-value) the FDR q-value is (p-value \* number of GO terms) / i.   
  
**Enrichment (N, B, n, b)** is defined as follows:  
N - is the total number of genes  
B - is the total number of genes associated with a specific GO term  
n - is the number of genes in the top of the user's input list or in the target set when appropriate  
b - is the number of genes in the intersection  
Enrichment = (b/n) / (B/N)  
  
**Genes:** For each GO term you can see the list of associated genes that appear in the optimal top of the list.  
Each gene name is specified by gene symbol followed by a short description of the gene   

Back to GOrilla analysis results
