## Supplementary figures and images for "A simple mass-action model predicts genome-wide protein timecourses from mRNA trajectories during a dynamic response in two strains of *Saccharomyces cerevisiae*"

### GOCOMPONENT.png

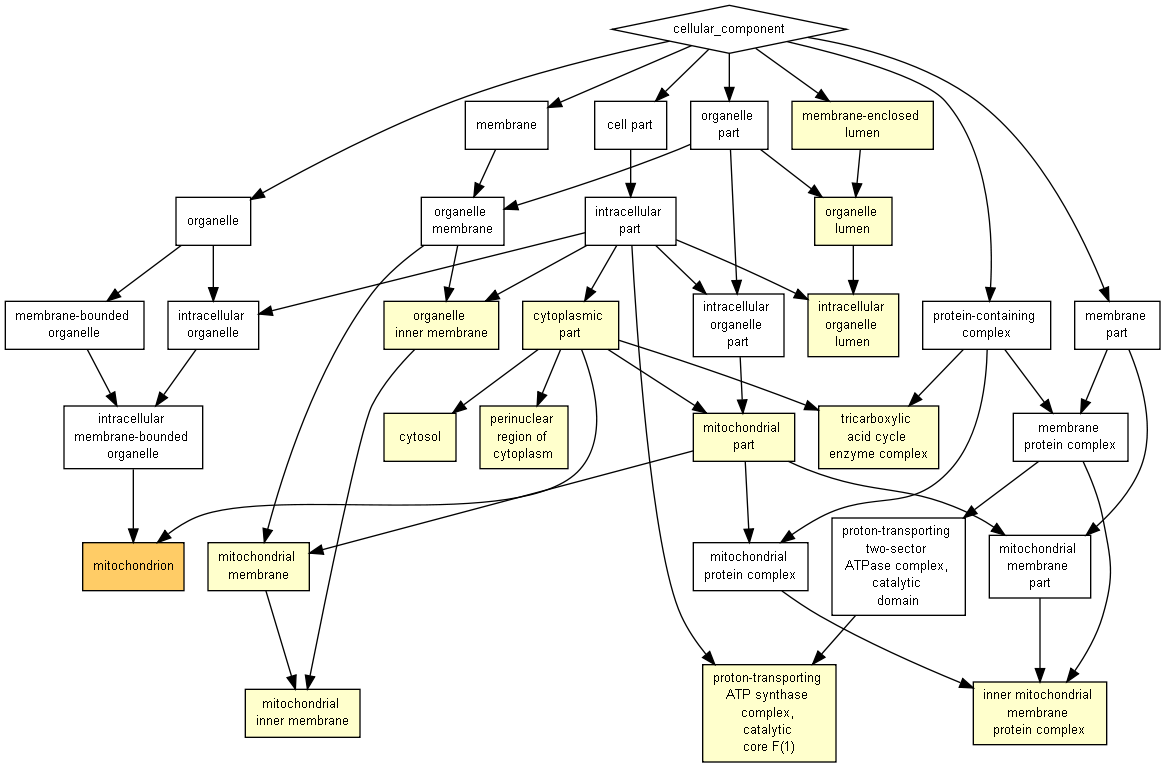

### GOCOMPONENT.png

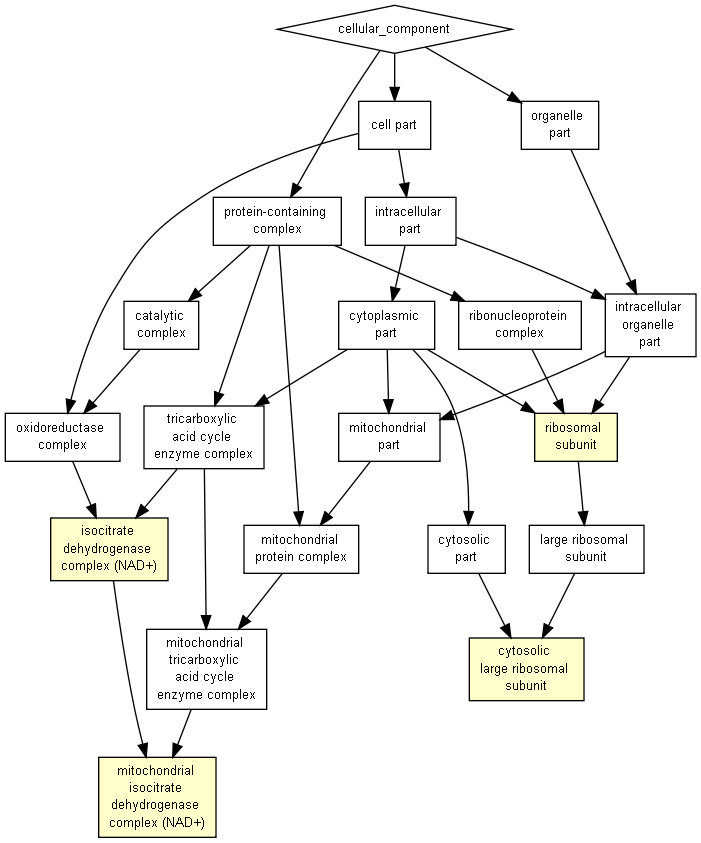

### GOCOMPONENT.png

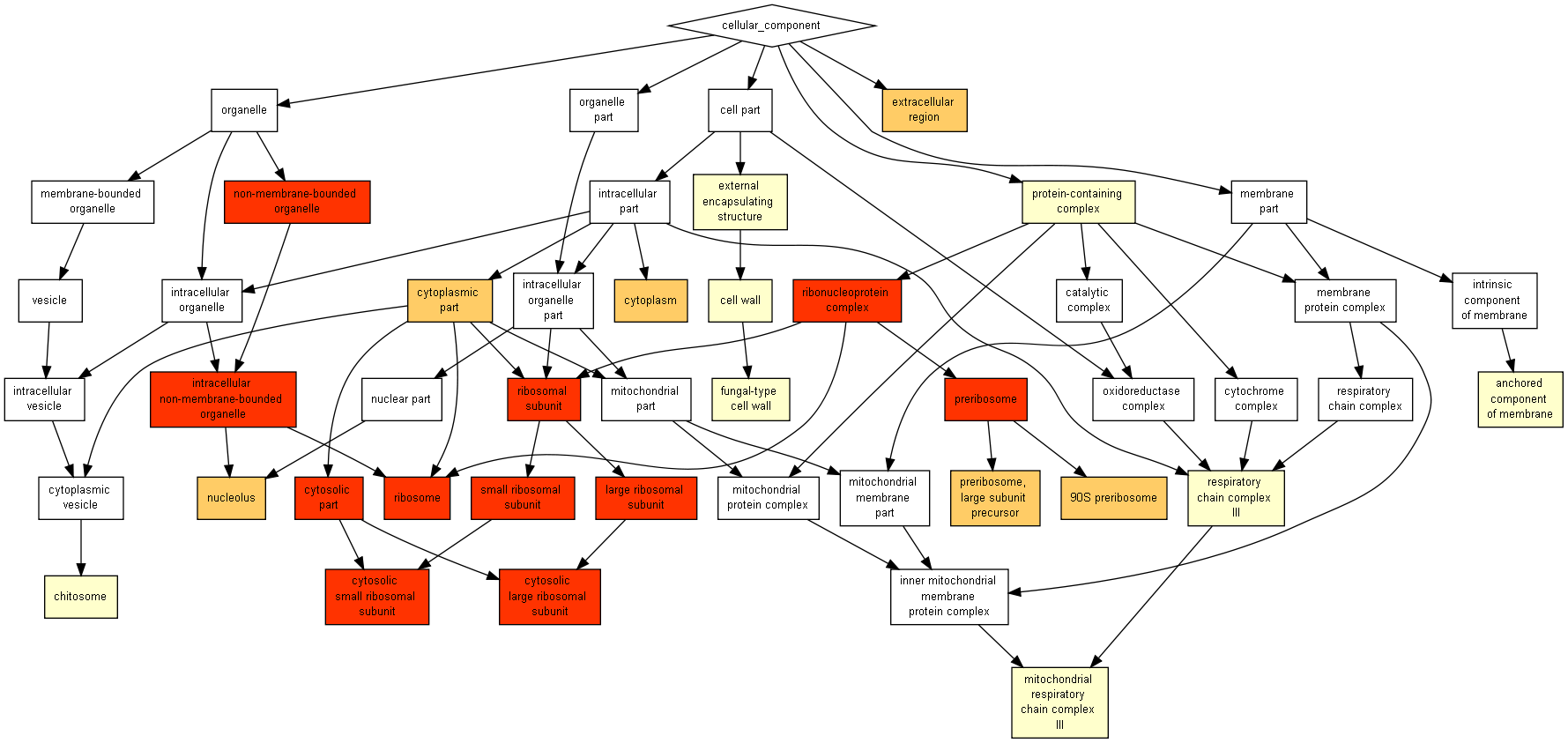

### GOFUNCTION.png

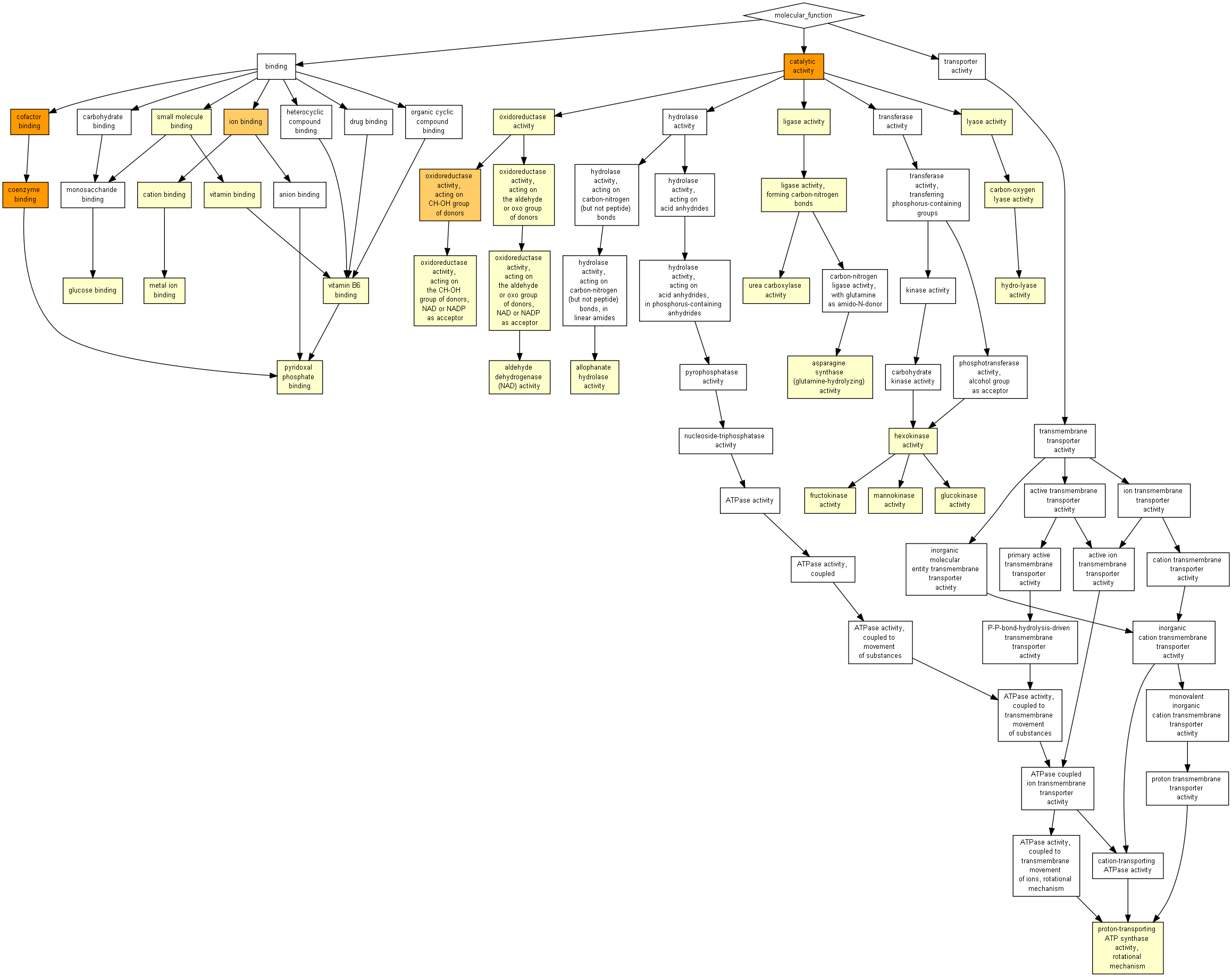

### GOFUNCTION.png

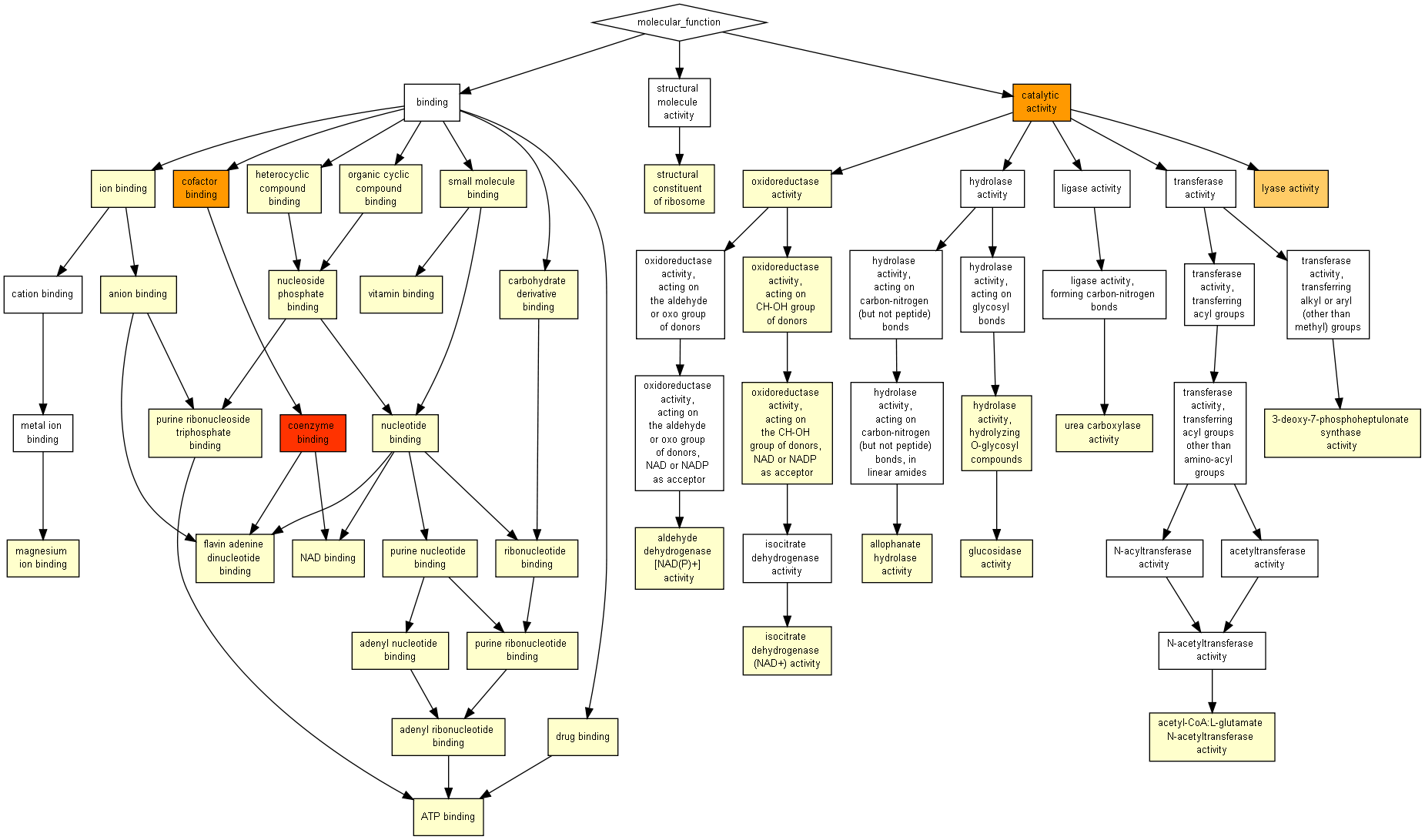

### GOFUNCTION.png

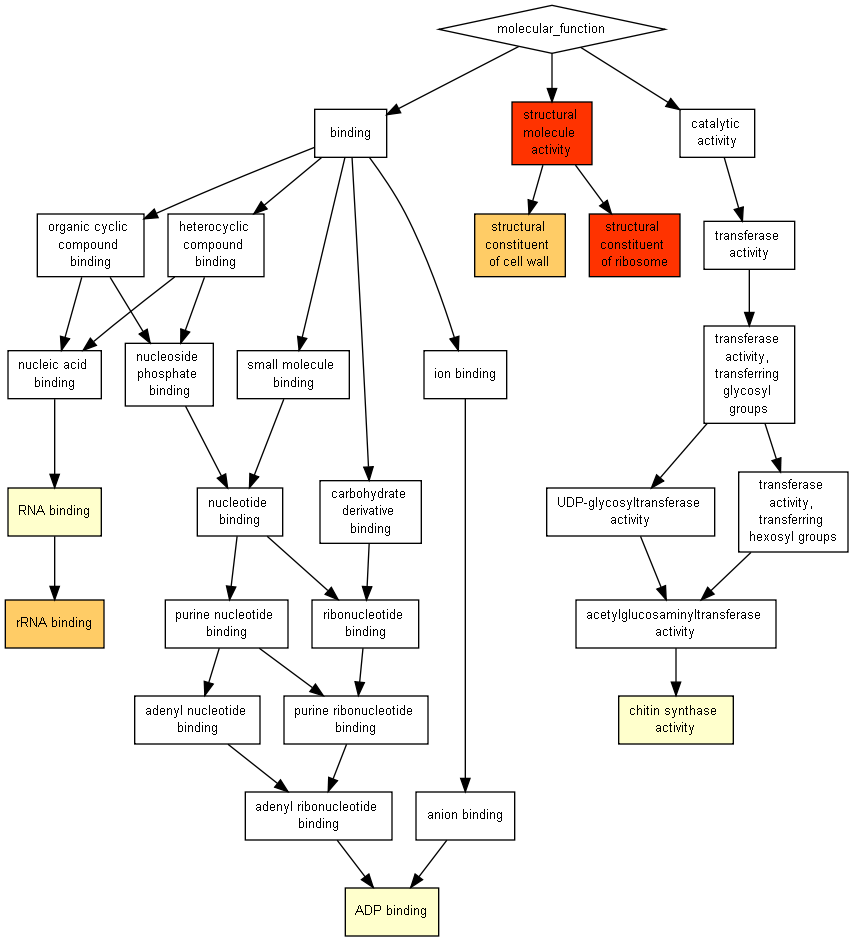

### GOPROCESS.png

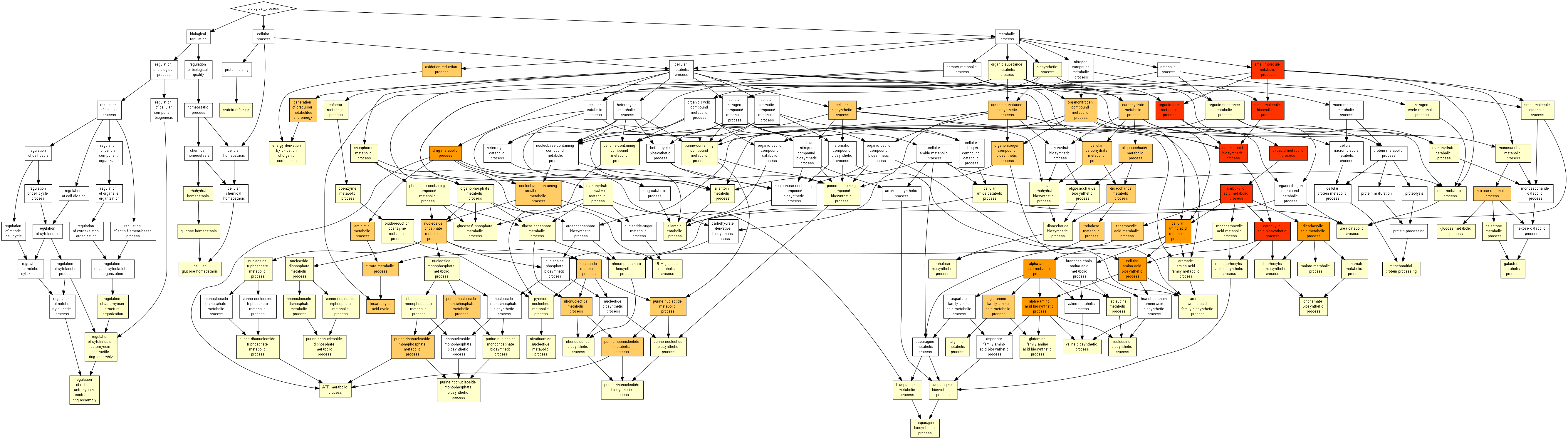

### GOPROCESS.png

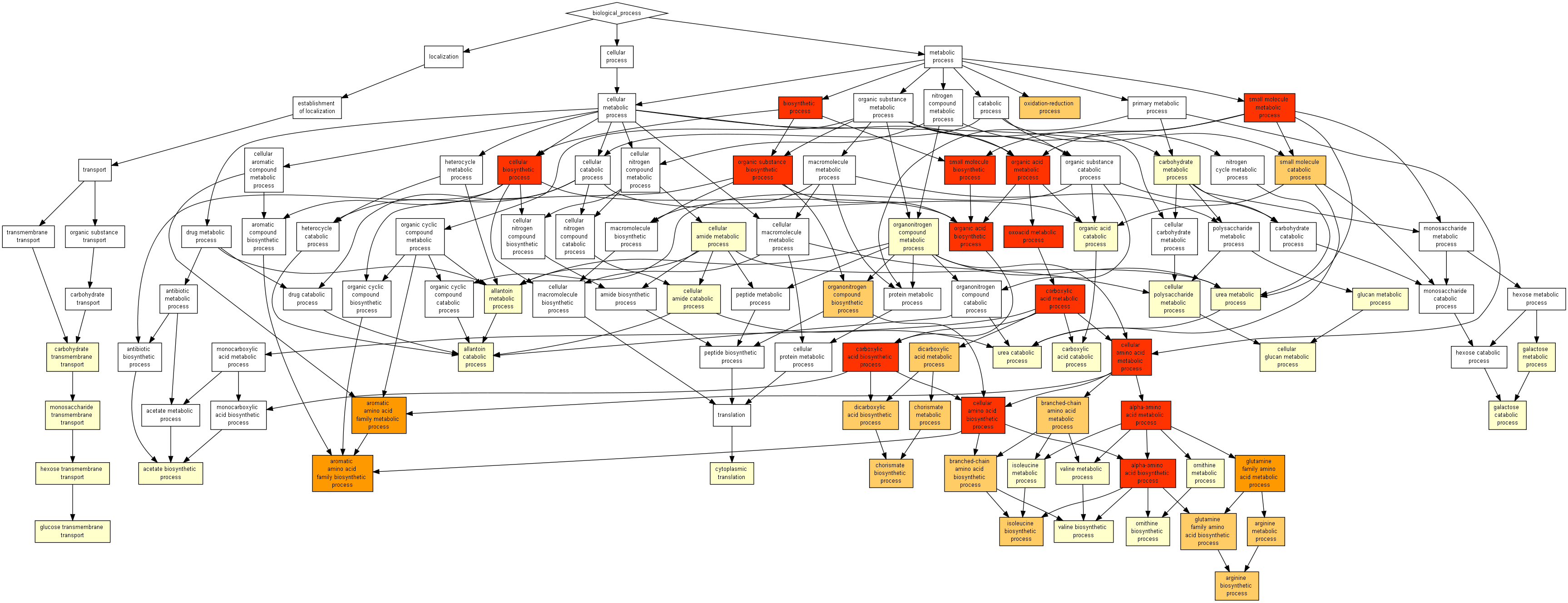

### GOPROCESS.png

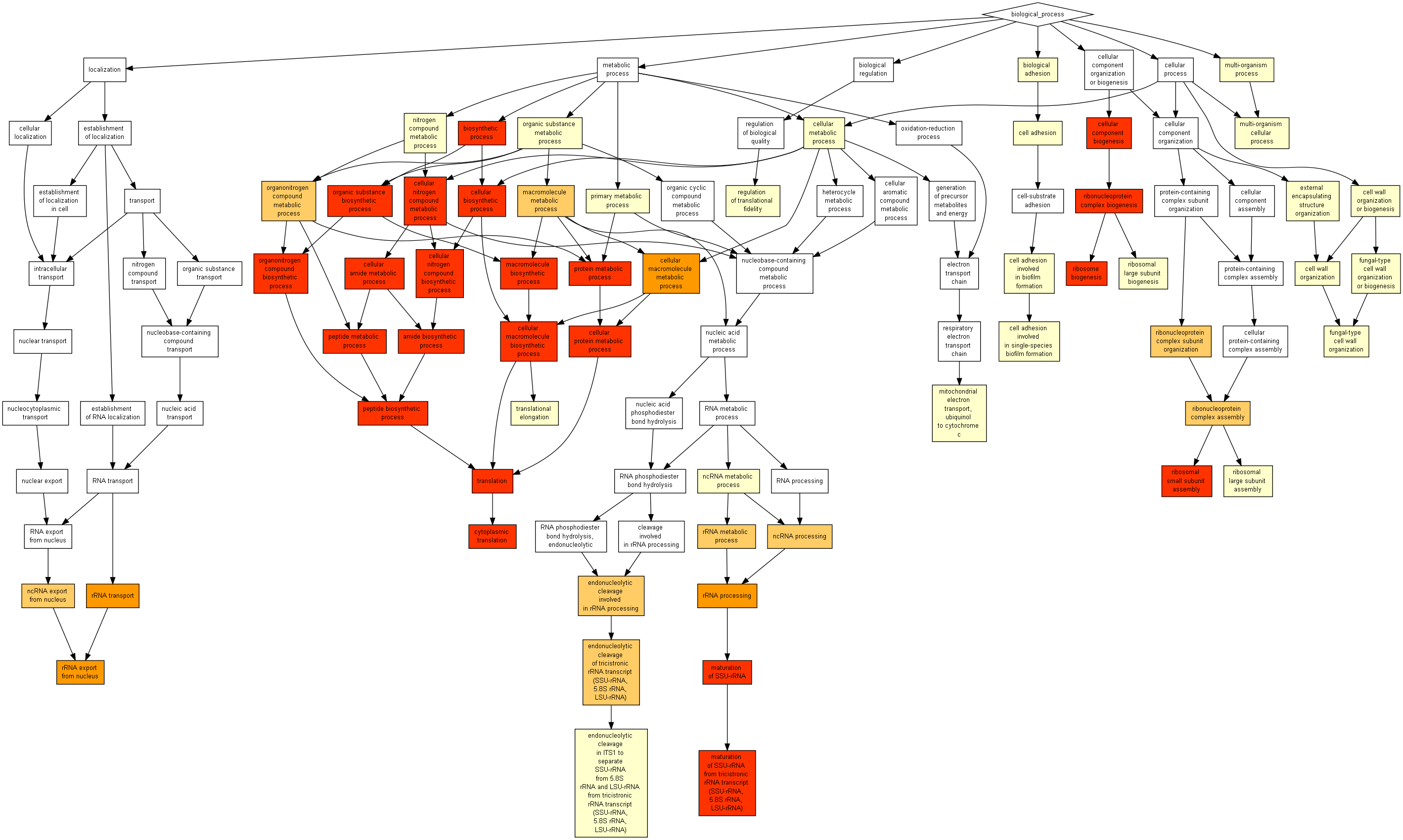
